## Supplementary File 1 for "BCIJelly: An integrated ecosystem for brain–computer interface research"

| Task type | Dataset | Metric | Algorithm 1 | Algorithm 2 | Performance 1 (%) | Performance 2 (%) | n_ pairs | t value | P value | significant |
|---|---|---|---|---|---|---|---|---|---|---|
| Regression | Indy | r2 | ndt1 | ndt2 | 45.3458 | 39.8417 | 12 | 2.7552 | 0.0187167 | * |
| Regression | Indy | r2 | ndt1 | poyo | 45.3458 | 30.5375 | 12 | 4.2705 | 0.00131963 | ** |
| Regression | Indy | r2 | ndt1 | UniBCI | 45.3458 | 34.53 | 12 | 5.9302 | 9.87111e-05 | *** |
| Regression | Indy | r2 | ndt1 | deepseek | 45.3458 | 38.009 | 12 | 2.5047 | 0.029262 | * |
| Regression | Indy | r2 | ndt1 | gpt | 45.3458 | 38.5654 | 12 | 2.0494 | 0.0650394 | n.s. |
| Regression | Indy | r2 | ndt1 | Claude | 45.3458 | 42.9534 | 12 | 0.7749 | 0.454721 | n.s. |
| Regression | Indy | r2 | ndt2 | poyo | 39.8417 | 30.5375 | 12 | 2.4002 | 0.0352191 | * |
| Regression | Indy | r2 | ndt2 | UniBCI | 39.8417 | 34.53 | 12 | 2.5746 | 0.0258378 | * |
| Regression | Indy | r2 | ndt2 | deepseek | 39.8417 | 38.009 | 12 | 0.7657 | 0.459944 | n.s. |
| Regression | Indy | r2 | ndt2 | gpt | 39.8417 | 38.5654 | 12 | 0.4783 | 0.641822 | n.s. |
| Regression | Indy | r2 | ndt2 | Claude | 39.8417 | 42.9534 | 12 | -1.2222 | 0.247174 | n.s. |
| Regression | Indy | r2 | poyo | UniBCI | 30.5375 | 34.53 | 12 | -1.1069 | 0.291973 | n.s. |
| Regression | Indy | r2 | poyo | deepseek | 30.5375 | 38.009 | 12 | -1.903 | 0.0835204 | n.s. |
| Regression | Indy | r2 | poyo | gpt | 30.5375 | 38.5654 | 12 | -1.9538 | 0.0766223 | n.s. |
| Regression | Indy | r2 | poyo | Claude | 30.5375 | 42.9534 | 12 | -2.9726 | 0.0126864 | * |
| Regression | Indy | r2 | UniBCI | deepseek | 34.53 | 38.009 | 12 | -0.9996 | 0.338972 | n.s. |
| Regression | Indy | r2 | UniBCI | gpt | 34.53 | 38.5654 | 12 | -1.0565 | 0.31338 | n.s. |
| Regression | Indy | r2 | UniBCI | Claude | 34.53 | 42.9534 | 12 | -2.2595 | 0.0451333 | * |
| Regression | Indy | r2 | deepseek | gpt | 38.009 | 38.5654 | 12 | -1.1234 | 0.285205 | n.s. |
| Regression | Indy | r2 | deepseek | Claude | 38.009 | 42.9534 | 12 | -11.0847 | 2.61784e-07 | *** |
| Regression | Indy | r2 | gpt | Claude | 38.5654 | 42.9534 | 12 | -8.665 | 3.0334e-06 | *** |
| Regression | Jango | r2 | ndt1 | ndt2 | 74.27 | 52.5079 | 14 | 5.2552 | 0.000155469 | *** |
| Regression | Jango | r2 | ndt1 | poyo | 74.27 | 73.0436 | 14 | 0.23 | 0.821682 | n.s. |
| Regression | Jango | r2 | ndt1 | UniBCI | 74.27 | 68.5279 | 14 | 3.3255 | 0.00547256 | ** |
| Regression | Jango | r2 | ndt1 | deepseek | 74.27 | 85.7647 | 14 | -1.8729 | 0.0837378 | n.s. |
| Regression | Jango | r2 | ndt1 | gpt | 74.27 | 85.7586 | 14 | -1.9061 | 0.0789815 | n.s. |
| Regression | Jango | r2 | ndt1 | Claude | 74.27 | 86.6858 | 14 | -2.0133 | 0.0652668 | n.s. |
| Regression | Jango | r2 | ndt2 | poyo | 52.5079 | 73.0436 | 14 | -4.1026 | 0.00124706 | ** |
| Regression | Jango | r2 | ndt2 | UniBCI | 52.5079 | 68.5279 | 14 | -3.4927 | 0.0039689 | ** |
| Regression | Jango | r2 | ndt2 | deepseek | 52.5079 | 85.7647 | 14 | -6.5185 | 1.94662e-05 | *** |
| Regression | Jango | r2 | ndt2 | gpt | 52.5079 | 85.7586 | 14 | -6.574 | 1.78573e-05 | *** |
| Regression | Jango | r2 | ndt2 | Claude | 52.5079 | 86.6858 | 14 | -6.5039 | 1.99139e-05 | *** |
| Regression | Jango | r2 | poyo | UniBCI | 73.0436 | 68.5279 | 14 | 0.8563 | 0.407316 | n.s. |
| Regression | Jango | r2 | poyo | deepseek | 73.0436 | 85.7647 | 14 | -3.5353 | 0.00365766 | ** |
| Regression | Jango | r2 | poyo | gpt | 73.0436 | 85.7586 | 14 | -3.7872 | 0.00226152 | ** |
| Regression | Jango | r2 | poyo | Claude | 73.0436 | 86.6858 | 14 | -3.8619 | 0.00196294 | ** |
| Regression | Jango | r2 | UniBCI | deepseek | 68.5279 | 85.7647 | 14 | -2.7521 | 0.0164729 | * |
| Regression | Jango | r2 | UniBCI | gpt | 68.5279 | 85.7586 | 14 | -2.8303 | 0.0141828 | * |
| Regression | Jango | r2 | UniBCI | Claude | 68.5279 | 86.6858 | 14 | -2.8953 | 0.0125201 | * |
| Regression | Jango | r2 | deepseek | gpt | 85.7647 | 85.7586 | 14 | 0.0105 | 0.991745 | n.s. |
| Regression | Jango | r2 | deepseek | Claude | 85.7647 | 86.6858 | 14 | -1.8096 | 0.0935331 | n.s. |
| Regression | Jango | r2 | gpt | Claude | 85.7586 | 86.6858 | 14 | -2.5433 | 0.0245023 | * |
| Classification | Finger PPC | accuracy | ndt1 | ndt2 | 36.18 | 33.0633 | 3 | 0.4431 | 0.701036 | n.s. |
| Classification | Finger PPC | accuracy | ndt1 | poyo | 36.18 | 36.9933 | 3 | -0.1136 | 0.9199 | n.s. |
| Classification | Finger PPC | accuracy | ndt1 | UniBCI | 36.18 | 38.2433 | 3 | -0.2504 | 0.825647 | n.s. |
| Classification | Finger PPC | accuracy | ndt1 | deepseek | 36.18 | 34.973 | 3 | 0.1577 | 0.889186 | n.s. |
| Classification | Finger PPC | accuracy | ndt1 | gpt | 36.18 | 32.3303 | 3 | 0.5076 | 0.662184 | n.s. |
| Classification | Finger PPC | accuracy | ndt1 | Claude | 36.18 | 32.3688 | 3 | 0.7028 | 0.55495 | n.s. |
| Classification | Finger PPC | accuracy | ndt2 | poyo | 33.0633 | 36.9933 | 3 | -29.5119 | 0.00114619 | ** |
| Classification | Finger PPC | accuracy | ndt2 | UniBCI | 33.0633 | 38.2433 | 3 | -0.7088 | 0.551907 | n.s. |
| Classification | Finger PPC | accuracy | ndt2 | deepseek | 33.0633 | 34.973 | 3 | -0.8718 | 0.475226 | n.s. |
| Classification | Finger PPC | accuracy | ndt2 | gpt | 33.0633 | 32.3303 | 3 | 0.547 | 0.639252 | n.s. |
| Classification | Finger PPC | accuracy | ndt2 | Claude | 33.0633 | 32.3688 | 3 | 0.2362 | 0.835248 | n.s. |
| Classification | Finger PPC | accuracy | poyo | UniBCI | 36.9933 | 38.2433 | 3 | -0.1689 | 0.881391 | n.s. |
| Classification | Finger PPC | accuracy | poyo | deepseek | 36.9933 | 34.973 | 3 | 0.9511 | 0.441925 | n.s. |
| Classification | Finger PPC | accuracy | poyo | gpt | 36.9933 | 32.3303 | 3 | 3.4391 | 0.0751427 | n.s. |
| Classification | Finger PPC | accuracy | poyo | Claude | 36.9933 | 32.3688 | 3 | 1.5468 | 0.261971 | n.s. |
| Classification | Finger PPC | accuracy | UniBCI | deepseek | 38.2433 | 34.973 | 3 | 0.3456 | 0.762592 | n.s. |
| Classification | Finger PPC | accuracy | UniBCI | gpt | 38.2433 | 32.3303 | 3 | 0.9346 | 0.448652 | n.s. |
| Classification | Finger PPC | accuracy | UniBCI | Claude | 38.2433 | 32.3688 | 3 | 0.6529 | 0.580868 | n.s. |

| | | | | | | | | | | |
|---|---|---|---|---|---|---|---|---|---|---|
| Classification | Finger PPC | accuracy | deepseek | gpt | 34.973 | 32.3303 | 3 | 0.7691 | 0.52224 | n.s. |
| Classification | Finger PPC | accuracy | deepseek | Claude | 34.973 | 32.3688 | 3 | 1.1088 | 0.382978 | n.s. |
| Classification | Finger PPC | accuracy | gpt | Claude | 32.3303 | 32.3688 | 3 | -0.0092 | 0.993506 | n.s. |
| Classification | Reaching B | accuracy | ndt1 | ndt2 | 75.498 | 65.1 | 10 | 2.7536 | 0.0223465 | * |
| Classification | Reaching B | accuracy | ndt1 | poyo | 75.498 | 80.117 | 10 | -1.5309 | 0.160164 | n.s. |
| Classification | Reaching B | accuracy | ndt1 | UniBCI | 75.498 | 66.831 | 10 | 2.7389 | 0.0228902 | * |
| Classification | Reaching B | accuracy | ndt1 | deepseek | 75.498 | 79.2549 | 10 | -1.1591 | 0.276262 | n.s. |
| Classification | Reaching B | accuracy | ndt1 | gpt | 75.498 | 78.1325 | 10 | -1.0073 | 0.340116 | n.s. |
| Classification | Reaching B | accuracy | ndt1 | Claude | 75.498 | 79.8623 | 10 | -1.4793 | 0.173189 | n.s. |
| Classification | Reaching B | accuracy | ndt2 | poyo | 65.1 | 80.117 | 10 | -3.2249 | 0.0104071 | * |
| Classification | Reaching B | accuracy | ndt2 | UniBCI | 65.1 | 66.831 | 10 | -0.3734 | 0.717471 | n.s. |
| Classification | Reaching B | accuracy | ndt2 | deepseek | 65.1 | 79.2549 | 10 | -2.6911 | 0.0247522 | * |
| Classification | Reaching B | accuracy | ndt2 | gpt | 65.1 | 78.1325 | 10 | -2.831 | 0.0196917 | * |
| Classification | Reaching B | accuracy | ndt2 | Claude | 65.1 | 79.8623 | 10 | -2.8228 | 0.019956 | * |
| Classification | Reaching B | accuracy | poyo | UniBCI | 80.117 | 66.831 | 10 | 4.7956 | 0.000979779 | *** |
| Classification | Reaching B | accuracy | poyo | deepseek | 80.117 | 79.2549 | 10 | 0.4291 | 0.677964 | n.s. |
| Classification | Reaching B | accuracy | poyo | gpt | 80.117 | 78.1325 | 10 | 1.2935 | 0.228036 | n.s. |
| Classification | Reaching B | accuracy | poyo | Claude | 80.117 | 79.8623 | 10 | 0.1499 | 0.884128 | n.s. |
| Classification | Reaching B | accuracy | UniBCI | deepseek | 66.831 | 79.2549 | 10 | -4.0828 | 0.00274648 | ** |
| Classification | Reaching B | accuracy | UniBCI | gpt | 66.831 | 78.1325 | 10 | -3.9249 | 0.00348469 | ** |
| Classification | Reaching B | accuracy | UniBCI | Claude | 66.831 | 79.8623 | 10 | -4.6675 | 0.00117275 | ** |
| Classification | Reaching B | accuracy | deepseek | gpt | 79.2549 | 78.1325 | 10 | 0.7899 | 0.449893 | n.s. |
| Classification | Reaching B | accuracy | deepseek | Claude | 79.2549 | 79.8623 | 10 | -0.6972 | 0.503306 | n.s. |
| Classification | Reaching B | accuracy | gpt | Claude | 78.1325 | 79.8623 | 10 | -1.7243 | 0.118738 | n.s. |
| Classification | Visual Coding | accuracy | ndt1 | ndt2 | 12.2233 | 15.47 | 6 | -1.9755 | 0.105182 | n.s. |
| Classification | Visual Coding | accuracy | ndt1 | poyo | 12.2233 | 13.5833 | 6 | -0.6884 | 0.521818 | n.s. |
| Classification | Visual Coding | accuracy | ndt1 | UniBCI | 12.2233 | 11.6383 | 6 | 0.2433 | 0.817411 | n.s. |
| Classification | Visual Coding | accuracy | ndt1 | deepseek | 12.2233 | 12.6111 | 6 | -0.1785 | 0.865366 | n.s. |
| Classification | Visual Coding | accuracy | ndt1 | gpt | 12.2233 | 13.0555 | 6 | -0.3428 | 0.745708 | n.s. |
| Classification | Visual Coding | accuracy | ndt1 | Claude | 12.2233 | 11.6111 | 6 | 0.2946 | 0.780113 | n.s. |
| Classification | Visual Coding | accuracy | ndt2 | poyo | 15.47 | 13.5833 | 6 | 0.8694 | 0.424403 | n.s. |
| Classification | Visual Coding | accuracy | ndt2 | UniBCI | 15.47 | 11.6383 | 6 | 1.1292 | 0.310075 | n.s. |
| Classification | Visual Coding | accuracy | ndt2 | deepseek | 15.47 | 12.6111 | 6 | 1.0977 | 0.322382 | n.s. |
| Classification | Visual Coding | accuracy | ndt2 | gpt | 15.47 | 13.0555 | 6 | 0.8634 | 0.427406 | n.s. |
| Classification | Visual Coding | accuracy | ndt2 | Claude | 15.47 | 11.6111 | 6 | 1.6112 | 0.168062 | n.s. |
| Classification | Visual Coding | accuracy | poyo | UniBCI | 13.5833 | 11.6383 | 6 | 0.6106 | 0.568159 | n.s. |
| Classification | Visual Coding | accuracy | poyo | deepseek | 13.5833 | 12.6111 | 6 | 0.394 | 0.709798 | n.s. |
| Classification | Visual Coding | accuracy | poyo | gpt | 13.5833 | 13.0555 | 6 | 0.2286 | 0.828244 | n.s. |
| Classification | Visual Coding | accuracy | poyo | Claude | 13.5833 | 11.6111 | 6 | 1.1293 | 0.310008 | n.s. |
| Classification | Visual Coding | accuracy | UniBCI | deepseek | 11.6383 | 12.6111 | 6 | -0.6655 | 0.535192 | n.s. |
| Classification | Visual Coding | accuracy | UniBCI | gpt | 11.6383 | 13.0555 | 6 | -0.9227 | 0.39852 | n.s. |
| Classification | Visual Coding | accuracy | UniBCI | Claude | 11.6383 | 11.6111 | 6 | 0.0149 | 0.988683 | n.s. |
| Classification | Visual Coding | accuracy | deepseek | gpt | 12.6111 | 13.0555 | 6 | -0.5946 | 0.577981 | n.s. |
| Classification | Visual Coding | accuracy | deepseek | Claude | 12.6111 | 11.6111 | 6 | 1.0617 | 0.336919 | n.s. |
| Classification | Visual Coding | accuracy | gpt | Claude | 13.0555 | 11.6111 | 6 | 2.3695 | 0.0639928 | n.s. |
| Human and macaque | Finger PPC | accuracy | ndt1 | ndt2 | 32.2067 | 29.85 | 3 | 0.9549 | 0.44041 | n.s. |
| Human and macaque | Finger PPC | accuracy | ndt1 | poyo | 32.2067 | 16.7067 | 3 | 7.2084 | 0.018707 | * |
| Human and macaque | Finger PPC | accuracy | ndt1 | UniBCI | 32.2067 | 32.9733 | 3 | -0.1078 | 0.923978 | n.s. |
| Human and macaque | Finger PPC | accuracy | ndt1 | deepseek | 32.2067 | 30.1601 | 3 | 0.35 | 0.759735 | n.s. |
| Human and macaque | Finger PPC | accuracy | ndt1 | gpt | 32.2067 | 33.8253 | 3 | -0.241 | 0.832032 | n.s. |
| Human and macaque | Finger PPC | accuracy | ndt1 | Claude | 32.2067 | 37.5579 | 3 | -1.1756 | 0.360746 | n.s. |
| Human and macaque | Finger PPC | accuracy | ndt2 | poyo | 29.85 | 16.7067 | 3 | 16.0626 | 0.00385347 | ** |
| Human and macaque | Finger PPC | accuracy | ndt2 | UniBCI | 29.85 | 32.9733 | 3 | -0.4107 | 0.721121 | n.s. |
| Human and macaque | Finger PPC | accuracy | ndt2 | deepseek | 29.85 | 30.1601 | 3 | -0.0634 | 0.955205 | n.s. |
| Human and macaque | Finger PPC | accuracy | ndt2 | gpt | 29.85 | 33.8253 | 3 | -0.7564 | 0.528377 | n.s. |
| Human and macaque | Finger PPC | accuracy | ndt2 | Claude | 29.85 | 37.5579 | 3 | -2.0323 | 0.179175 | n.s. |
| Human and macaque | Finger PPC | accuracy | poyo | UniBCI | 16.7067 | 32.9733 | 3 | -2.3908 | 0.139307 | n.s. |
| Human and macaque | Finger PPC | accuracy | poyo | deepseek | 16.7067 | 30.1601 | 3 | -2.3781 | 0.140497 | n.s. |
| Human and macaque | Finger PPC | accuracy | poyo | gpt | 16.7067 | 33.8253 | 3 | -2.8201 | 0.106101 | n.s. |
| Human and macaque | Finger PPC | accuracy | poyo | Claude | 16.7067 | 37.5579 | 3 | -4.6207 | 0.0437829 | * |
| Human and macaque | Finger PPC | accuracy | UniBCI | deepseek | 32.9733 | 30.1601 | 3 | 0.2261 | 0.842116 | n.s. |
| Human and macaque | Finger PPC | accuracy | UniBCI | gpt | 32.9733 | 33.8253 | 3 | -0.0663 | 0.953164 | n.s. |
| Human and macaque | Finger PPC | accuracy | UniBCI | Claude | 32.9733 | 37.5579 | 3 | -0.4078 | 0.722933 | n.s. |

| | | | | | | | | | | |
|---|---|---|---|---|---|---|---|---|---|---|
| Human and macaque | Finger PPC | accuracy | deepseek | gpt | 30.1601 | 33.8253 | 3 | -2.4979 | 0.12979 | n.s. |
| Human and macaque | Finger PPC | accuracy | deepseek | Claude | 30.1601 | 37.5579 | 3 | -5.7007 | 0.02942 | * |
| Human and macaque | Finger PPC | accuracy | gpt | Claude | 33.8253 | 37.5579 | 3 | -1.527 | 0.266322 | n.s. |
| Human and macaque | Reaching B | accuracy | ndt1 | ndt2 | 63.369 | 56.105 | 10 | 1.6441 | 0.134557 | n.s. |
| Human and macaque | Reaching B | accuracy | ndt1 | poyo | 63.369 | 56.585 | 10 | 1.2085 | 0.257656 | n.s. |
| Human and macaque | Reaching B | accuracy | ndt1 | UniBCI | 63.369 | 59.632 | 10 | 1.471 | 0.175372 | n.s. |
| Human and macaque | Reaching B | accuracy | ndt1 | deepseek | 63.369 | 82.93 | 10 | -5.3679 | 0.000451578 | *** |
| Human and macaque | Reaching B | accuracy | ndt1 | gpt | 63.369 | 82.5847 | 10 | -5.5376 | 0.000362169 | *** |
| Human and macaque | Reaching B | accuracy | ndt1 | Claude | 63.369 | 82.2448 | 10 | -5.1288 | 0.000620582 | *** |
| Human and macaque | Reaching B | accuracy | ndt2 | poyo | 56.105 | 56.585 | 10 | -0.0855 | 0.933763 | n.s. |
| Human and macaque | Reaching B | accuracy | ndt2 | UniBCI | 56.105 | 59.632 | 10 | -1.5139 | 0.16435 | n.s. |
| Human and macaque | Reaching B | accuracy | ndt2 | deepseek | 56.105 | 82.93 | 10 | -8.5922 | 1.24561e-05 | *** |
| Human and macaque | Reaching B | accuracy | ndt2 | gpt | 56.105 | 82.5847 | 10 | -8.2468 | 1.73493e-05 | *** |
| Human and macaque | Reaching B | accuracy | ndt2 | Claude | 56.105 | 82.2448 | 10 | -9.6324 | 4.88272e-06 | *** |
| Human and macaque | Reaching B | accuracy | poyo | UniBCI | 56.585 | 59.632 | 10 | -0.5496 | 0.595977 | n.s. |
| Human and macaque | Reaching B | accuracy | poyo | deepseek | 56.585 | 82.93 | 10 | -4.4352 | 0.00163491 | ** |
| Human and macaque | Reaching B | accuracy | poyo | gpt | 56.585 | 82.5847 | 10 | -4.422 | 0.00166625 | ** |
| Human and macaque | Reaching B | accuracy | poyo | Claude | 56.585 | 82.2448 | 10 | -4.1461 | 0.00249876 | ** |
| Human and macaque | Reaching B | accuracy | UniBCI | deepseek | 59.632 | 82.93 | 10 | -10.0093 | 3.55051e-06 | *** |
| Human and macaque | Reaching B | accuracy | UniBCI | gpt | 59.632 | 82.5847 | 10 | -10.2139 | 2.9995e-06 | *** |
| Human and macaque | Reaching B | accuracy | UniBCI | Claude | 59.632 | 82.2448 | 10 | -9.8279 | 4.13363e-06 | *** |
| Human and macaque | Reaching B | accuracy | deepseek | gpt | 82.93 | 82.5847 | 10 | 0.4766 | 0.64503 | n.s. |
| Human and macaque | Reaching B | accuracy | deepseek | Claude | 82.93 | 82.2448 | 10 | 0.7617 | 0.465741 | n.s. |
| Human and macaque | Reaching B | accuracy | gpt | Claude | 82.5847 | 82.2448 | 10 | 0.2806 | 0.785353 | n.s. |
| Human and macaque | Indy | r2 | ndt1 | ndt2 | 48.9442 | 42.4183 | 12 | 5.1002 | 0.000343913 | *** |
| Human and macaque | Indy | r2 | ndt1 | poyo | 48.9442 | 30.6908 | 12 | 6.1894 | 6.81688e-05 | *** |
| Human and macaque | Indy | r2 | ndt1 | UniBCI | 48.9442 | 33.9475 | 12 | 7.1853 | 1.78594e-05 | *** |
| Human and macaque | Indy | r2 | ndt1 | deepseek | 48.9442 | 35.7945 | 12 | 2.9978 | 0.0121264 | * |
| Human and macaque | Indy | r2 | ndt1 | gpt | 48.9442 | 34.1698 | 12 | 4.2845 | 0.00128905 | ** |
| Human and macaque | Indy | r2 | ndt1 | Claude | 48.9442 | 43.3146 | 12 | 1.9301 | 0.0797739 | n.s. |
| Human and macaque | Indy | r2 | ndt2 | poyo | 42.4183 | 30.6908 | 12 | 3.9362 | 0.00232719 | ** |
| Human and macaque | Indy | r2 | ndt2 | UniBCI | 42.4183 | 33.9475 | 12 | 5.6884 | 0.000140588 | *** |
| Human and macaque | Indy | r2 | ndt2 | deepseek | 42.4183 | 35.7945 | 12 | 1.3225 | 0.212854 | n.s. |
| Human and macaque | Indy | r2 | ndt2 | gpt | 42.4183 | 34.1698 | 12 | 2.2515 | 0.0457709 | * |
| Human and macaque | Indy | r2 | ndt2 | Claude | 42.4183 | 43.3146 | 12 | -0.2617 | 0.798406 | n.s. |
| Human and macaque | Indy | r2 | poyo | UniBCI | 30.6908 | 33.9475 | 12 | -0.892 | 0.391491 | n.s. |
| Human and macaque | Indy | r2 | poyo | deepseek | 30.6908 | 35.7945 | 12 | -1.0142 | 0.332262 | n.s. |
| Human and macaque | Indy | r2 | poyo | gpt | 30.6908 | 34.1698 | 12 | -0.8932 | 0.390892 | n.s. |
| Human and macaque | Indy | r2 | poyo | Claude | 30.6908 | 43.3146 | 12 | -3.1326 | 0.00953417 | ** |
| Human and macaque | Indy | r2 | UniBCI | deepseek | 33.9475 | 35.7945 | 12 | -0.3384 | 0.741436 | n.s. |
| Human and macaque | Indy | r2 | UniBCI | gpt | 33.9475 | 34.1698 | 12 | -0.0536 | 0.958189 | n.s. |
| Human and macaque | Indy | r2 | UniBCI | Claude | 33.9475 | 43.3146 | 12 | -2.2706 | 0.044259 | * |
| Human and macaque | Indy | r2 | deepseek | gpt | 35.7945 | 34.1698 | 12 | 0.7539 | 0.466731 | n.s. |
| Human and macaque | Indy | r2 | deepseek | Claude | 35.7945 | 43.3146 | 12 | -3.6185 | 0.00403698 | ** |
| Human and macaque | Indy | r2 | gpt | Claude | 34.1698 | 43.3146 | 12 | -6.7205 | 3.28474e-05 | *** |
| Human and macaque | Jango | r2 | ndt1 | ndt2 | 66.025 | 60.4271 | 14 | 2.2628 | 0.0414168 | * |
| Human and macaque | Jango | r2 | ndt1 | poyo | 66.025 | 60.6643 | 14 | 0.7796 | 0.449599 | n.s. |
| Human and macaque | Jango | r2 | ndt1 | UniBCI | 66.025 | 61.415 | 14 | 0.8385 | 0.416892 | n.s. |
| Human and macaque | Jango | r2 | ndt1 | deepseek | 66.025 | 84.9319 | 14 | -4.867 | 0.000307644 | *** |
| Human and macaque | Jango | r2 | ndt1 | gpt | 66.025 | 83.6957 | 14 | -4.6616 | 0.000445076 | *** |
| Human and macaque | Jango | r2 | ndt1 | Claude | 66.025 | 84.7661 | 14 | -5.4763 | 0.000106344 | *** |
| Human and macaque | Jango | r2 | ndt2 | poyo | 60.4271 | 60.6643 | 14 | -0.0318 | 0.975133 | n.s. |
| Human and macaque | Jango | r2 | ndt2 | UniBCI | 60.4271 | 61.415 | 14 | -0.1356 | 0.894208 | n.s. |
| Human and macaque | Jango | r2 | ndt2 | deepseek | 60.4271 | 84.9319 | 14 | -4.636 | 0.000466206 | *** |
| Human and macaque | Jango | r2 | ndt2 | gpt | 60.4271 | 83.6957 | 14 | -4.3906 | 0.000730143 | *** |
| Human and macaque | Jango | r2 | ndt2 | Claude | 60.4271 | 84.7661 | 14 | -4.9884 | 0.000247991 | *** |
| Human and macaque | Jango | r2 | poyo | UniBCI | 60.6643 | 61.415 | 14 | -0.0882 | 0.93106 | n.s. |
| Human and macaque | Jango | r2 | poyo | deepseek | 60.6643 | 84.9319 | 14 | -3.0905 | 0.00860201 | ** |
| Human and macaque | Jango | r2 | poyo | gpt | 60.6643 | 83.6957 | 14 | -2.9497 | 0.0112785 | * |
| Human and macaque | Jango | r2 | poyo | Claude | 60.6643 | 84.7661 | 14 | -3.1861 | 0.00715689 | ** |
| Human and macaque | Jango | r2 | UniBCI | deepseek | 61.415 | 84.9319 | 14 | -4.4136 | 0.000699891 | *** |
| Human and macaque | Jango | r2 | UniBCI | gpt | 61.415 | 83.6957 | 14 | -4.3534 | 0.00078202 | *** |
| Human and macaque | Jango | r2 | UniBCI | Claude | 61.415 | 84.7661 | 14 | -4.5386 | 0.000556633 | *** |

| | | | | | | | | | | |
|---|---|---|---|---|---|---|---|---|---|---|
| Human and macaque | Jango | r2 | deepseek | gpt | 84.9319 | 83.6957 | 14 | 1.423 | 0.178303 | n.s. |
| Human and macaque | Jango | r2 | deepseek | Claude | 84.9319 | 84.7661 | 14 | 0.2134 | 0.834362 | n.s. |
| Human and macaque | Jango | r2 | gpt | Claude | 83.6957 | 84.7661 | 14 | -1.8367 | 0.0892189 | n.s. |
| Human and mouse | Finger PPC | accuracy | ndt1 | ndt2 | 33.8533 | 36.3733 | 3 | -0.7324 | 0.540104 | n.s. |
| Human and mouse | Finger PPC | accuracy | ndt1 | poyo | 33.8533 | 39.9233 | 3 | -2.7736 | 0.109125 | n.s. |
| Human and mouse | Finger PPC | accuracy | ndt1 | UniBCI | 33.8533 | 45.92 | 3 | -1.159 | 0.366129 | n.s. |
| Human and mouse | Finger PPC | accuracy | ndt1 | deepseek | 33.8533 | 36.1111 | 3 | -0.6734 | 0.570088 | n.s. |
| Human and mouse | Finger PPC | accuracy | ndt1 | gpt | 33.8533 | 29.2824 | 3 | 1.794 | 0.214668 | n.s. |
| Human and mouse | Finger PPC | accuracy | ndt1 | Claude | 33.8533 | 35.0502 | 3 | -0.2524 | 0.824273 | n.s. |
| Human and mouse | Finger PPC | accuracy | ndt2 | poyo | 36.3733 | 39.9233 | 3 | -2.373 | 0.140985 | n.s. |
| Human and mouse | Finger PPC | accuracy | ndt2 | UniBCI | 36.3733 | 45.92 | 3 | -0.7375 | 0.537591 | n.s. |
| Human and mouse | Finger PPC | accuracy | ndt2 | deepseek | 36.3733 | 36.1111 | 3 | 0.0816 | 0.942416 | n.s. |
| Human and mouse | Finger PPC | accuracy | ndt2 | gpt | 36.3733 | 29.2824 | 3 | 2.7419 | 0.111255 | n.s. |
| Human and mouse | Finger PPC | accuracy | ndt2 | Claude | 36.3733 | 35.0502 | 3 | 0.3944 | 0.731355 | n.s. |
| Human and mouse | Finger PPC | accuracy | poyo | UniBCI | 39.9233 | 45.92 | 3 | -0.5222 | 0.65358 | n.s. |
| Human and mouse | Finger PPC | accuracy | poyo | deepseek | 39.9233 | 36.1111 | 3 | 1.763 | 0.219952 | n.s. |
| Human and mouse | Finger PPC | accuracy | poyo | gpt | 39.9233 | 29.2824 | 3 | 8.0538 | 0.0150692 | * |
| Human and mouse | Finger PPC | accuracy | poyo | Claude | 39.9233 | 35.0502 | 3 | 1.6317 | 0.244333 | n.s. |
| Human and mouse | Finger PPC | accuracy | UniBCI | deepseek | 45.92 | 36.1111 | 3 | 0.974 | 0.432773 | n.s. |
| Human and mouse | Finger PPC | accuracy | UniBCI | gpt | 45.92 | 29.2824 | 3 | 1.5983 | 0.251091 | n.s. |
| Human and mouse | Finger PPC | accuracy | UniBCI | Claude | 45.92 | 35.0502 | 3 | 0.974 | 0.432771 | n.s. |
| Human and mouse | Finger PPC | accuracy | deepseek | gpt | 36.1111 | 29.2824 | 3 | 7.5539 | 0.0170772 | * |
| Human and mouse | Finger PPC | accuracy | deepseek | Claude | 36.1111 | 35.0502 | 3 | 0.6631 | 0.575463 | n.s. |
| Human and mouse | Finger PPC | accuracy | gpt | Claude | 29.2824 | 35.0502 | 3 | -2.6185 | 0.120123 | n.s. |
| Human and mouse | Visual Coding | accuracy | ndt1 | ndt2 | 13.8033 | 13.9167 | 6 | -0.176 | 0.867223 | n.s. |
| Human and mouse | Visual Coding | accuracy | ndt1 | poyo | 13.8033 | 11.1117 | 6 | 1.7838 | 0.134539 | n.s. |
| Human and mouse | Visual Coding | accuracy | ndt1 | UniBCI | 13.8033 | 12.305 | 6 | 0.4482 | 0.672787 | n.s. |
| Human and mouse | Visual Coding | accuracy | ndt1 | deepseek | 13.8033 | 11.5278 | 6 | 1.2582 | 0.263865 | n.s. |
| Human and mouse | Visual Coding | accuracy | ndt1 | gpt | 13.8033 | 12.1111 | 6 | 0.8799 | 0.419202 | n.s. |
| Human and mouse | Visual Coding | accuracy | ndt1 | Claude | 13.8033 | 12.5 | 6 | 0.7154 | 0.506391 | n.s. |
| Human and mouse | Visual Coding | accuracy | ndt2 | poyo | 13.9167 | 11.1117 | 6 | 2.4979 | 0.054627 | n.s. |
| Human and mouse | Visual Coding | accuracy | ndt2 | UniBCI | 13.9167 | 12.305 | 6 | 0.5668 | 0.595361 | n.s. |
| Human and mouse | Visual Coding | accuracy | ndt2 | deepseek | 13.9167 | 11.5278 | 6 | 1.5586 | 0.179828 | n.s. |
| Human and mouse | Visual Coding | accuracy | ndt2 | gpt | 13.9167 | 12.1111 | 6 | 1.1504 | 0.302024 | n.s. |
| Human and mouse | Visual Coding | accuracy | ndt2 | Claude | 13.9167 | 12.5 | 6 | 1.0306 | 0.349964 | n.s. |
| Human and mouse | Visual Coding | accuracy | poyo | UniBCI | 11.1117 | 12.305 | 6 | -0.5806 | 0.586699 | n.s. |
| Human and mouse | Visual Coding | accuracy | poyo | deepseek | 11.1117 | 11.5278 | 6 | -0.5673 | 0.595037 | n.s. |
| Human and mouse | Visual Coding | accuracy | poyo | gpt | 11.1117 | 12.1111 | 6 | -0.9085 | 0.405265 | n.s. |
| Human and mouse | Visual Coding | accuracy | poyo | Claude | 11.1117 | 12.5 | 6 | -1.8307 | 0.126654 | n.s. |
| Human and mouse | Visual Coding | accuracy | UniBCI | deepseek | 12.305 | 11.5278 | 6 | 0.3586 | 0.734539 | n.s. |
| Human and mouse | Visual Coding | accuracy | UniBCI | gpt | 12.305 | 12.1111 | 6 | 0.1067 | 0.919163 | n.s. |
| Human and mouse | Visual Coding | accuracy | UniBCI | Claude | 12.305 | 12.5 | 6 | -0.118 | 0.910656 | n.s. |
| Human and mouse | Visual Coding | accuracy | deepseek | gpt | 11.5278 | 12.1111 | 6 | -0.6138 | 0.566202 | n.s. |
| Human and mouse | Visual Coding | accuracy | deepseek | Claude | 11.5278 | 12.5 | 6 | -0.9837 | 0.370418 | n.s. |
| Human and mouse | Visual Coding | accuracy | gpt | Claude | 12.1111 | 12.5 | 6 | -0.6818 | 0.525645 | n.s. |
| Macaque and mouse | Reaching B | accuracy | ndt1 | ndt2 | 52.182 | 63.182 | 10 | -2.3017 | 0.0468723 | * |
| Macaque and mouse | Reaching B | accuracy | ndt1 | poyo | 52.182 | 62.292 | 10 | -3.1976 | 0.0108732 | * |
| Macaque and mouse | Reaching B | accuracy | ndt1 | UniBCI | 52.182 | 63.939 | 10 | -3.0063 | 0.0148042 | * |
| Macaque and mouse | Reaching B | accuracy | ndt1 | deepseek | 52.182 | 81.5892 | 10 | -9.8285 | 4.13153e-06 | *** |
| Macaque and mouse | Reaching B | accuracy | ndt1 | gpt | 52.182 | 78.5497 | 10 | -9.1246 | 7.62892e-06 | *** |
| Macaque and mouse | Reaching B | accuracy | ndt1 | Claude | 52.182 | 79.5089 | 10 | -8.7819 | 1.04306e-05 | *** |
| Macaque and mouse | Reaching B | accuracy | ndt2 | poyo | 63.182 | 62.292 | 10 | 0.2502 | 0.808077 | n.s. |
| Macaque and mouse | Reaching B | accuracy | ndt2 | UniBCI | 63.182 | 63.939 | 10 | -0.1811 | 0.860308 | n.s. |
| Macaque and mouse | Reaching B | accuracy | ndt2 | deepseek | 63.182 | 81.5892 | 10 | -4.7859 | 0.000992992 | *** |
| Macaque and mouse | Reaching B | accuracy | ndt2 | gpt | 63.182 | 78.5497 | 10 | -3.8876 | 0.00368819 | ** |
| Macaque and mouse | Reaching B | accuracy | ndt2 | Claude | 63.182 | 79.5089 | 10 | -4.3513 | 0.00184666 | ** |
| Macaque and mouse | Reaching B | accuracy | poyo | UniBCI | 62.292 | 63.939 | 10 | -0.8489 | 0.417967 | n.s. |
| Macaque and mouse | Reaching B | accuracy | poyo | deepseek | 62.292 | 81.5892 | 10 | -8.1324 | 1.94099e-05 | *** |
| Macaque and mouse | Reaching B | accuracy | poyo | gpt | 62.292 | 78.5497 | 10 | -6.947 | 6.70716e-05 | *** |
| Macaque and mouse | Reaching B | accuracy | poyo | Claude | 62.292 | 79.5089 | 10 | -8.2562 | 1.71926e-05 | *** |
| Macaque and mouse | Reaching B | accuracy | UniBCI | deepseek | 63.939 | 81.5892 | 10 | -6.7871 | 8.02105e-05 | *** |
| Macaque and mouse | Reaching B | accuracy | UniBCI | gpt | 63.939 | 78.5497 | 10 | -6.5331 | 0.000107253 | *** |
| Macaque and mouse | Reaching B | accuracy | UniBCI | Claude | 63.939 | 79.5089 | 10 | -6.0534 | 0.000189696 | *** |

| | | | | | | | | | | |
|---|---|---|---|---|---|---|---|---|---|---|
| Macaque and mouse | Reaching B | accuracy | deepseek | gpt | 81.5892 | 78.5497 | 10 | 3.6141 | 0.00562348 | ** |
| Macaque and mouse | Reaching B | accuracy | deepseek | Claude | 81.5892 | 79.5089 | 10 | 3.0267 | 0.014323 | * |
| Macaque and mouse | Reaching B | accuracy | gpt | Claude | 78.5497 | 79.5089 | 10 | -0.7035 | 0.499553 | n.s. |
| Macaque and mouse | Indy | r2 | ndt1 | ndt2 | 50.3208 | 43.5125 | 12 | 4.3274 | 0.00119972 | ** |
| Macaque and mouse | Indy | r2 | ndt1 | poyo | 50.3208 | 30.8967 | 12 | 5.6085 | 0.00015829 | *** |
| Macaque and mouse | Indy | r2 | ndt1 | UniBCI | 50.3208 | 36.3258 | 12 | 4.8901 | 0.000479133 | *** |
| Macaque and mouse | Indy | r2 | ndt1 | deepseek | 50.3208 | 39.406 | 12 | 3.1113 | 0.00990304 | ** |
| Macaque and mouse | Indy | r2 | ndt1 | gpt | 50.3208 | 33.3555 | 12 | 4.2406 | 0.00138756 | ** |
| Macaque and mouse | Indy | r2 | ndt1 | Claude | 50.3208 | 43.152 | 12 | 2.7643 | 0.0184121 | * |
| Macaque and mouse | Indy | r2 | ndt2 | poyo | 43.5125 | 30.8967 | 12 | 4.2803 | 0.00129807 | ** |
| Macaque and mouse | Indy | r2 | ndt2 | UniBCI | 43.5125 | 36.3258 | 12 | 3.3579 | 0.00638761 | ** |
| Macaque and mouse | Indy | r2 | ndt2 | deepseek | 43.5125 | 39.406 | 12 | 1.2395 | 0.240941 | n.s. |
| Macaque and mouse | Indy | r2 | ndt2 | gpt | 43.5125 | 33.3555 | 12 | 2.5935 | 0.0249839 | * |
| Macaque and mouse | Indy | r2 | ndt2 | Claude | 43.5125 | 43.152 | 12 | 0.1467 | 0.886006 | n.s. |
| Macaque and mouse | Indy | r2 | poyo | UniBCI | 30.8967 | 36.3258 | 12 | -1.5716 | 0.144343 | n.s. |
| Macaque and mouse | Indy | r2 | poyo | deepseek | 30.8967 | 39.406 | 12 | -2.0205 | 0.0683632 | n.s. |
| Macaque and mouse | Indy | r2 | poyo | gpt | 30.8967 | 33.3555 | 12 | -0.5333 | 0.604465 | n.s. |
| Macaque and mouse | Indy | r2 | poyo | Claude | 30.8967 | 43.152 | 12 | -3.1456 | 0.00931527 | ** |
| Macaque and mouse | Indy | r2 | UniBCI | deepseek | 36.3258 | 39.406 | 12 | -0.7417 | 0.473793 | n.s. |
| Macaque and mouse | Indy | r2 | UniBCI | gpt | 36.3258 | 33.3555 | 12 | 0.6444 | 0.532508 | n.s. |
| Macaque and mouse | Indy | r2 | UniBCI | Claude | 36.3258 | 43.152 | 12 | -1.9389 | 0.0785938 | n.s. |
| Macaque and mouse | Indy | r2 | deepseek | gpt | 39.406 | 33.3555 | 12 | 4.7577 | 0.000592324 | *** |
| Macaque and mouse | Indy | r2 | deepseek | Claude | 39.406 | 43.152 | 12 | -3.2076 | 0.00834182 | ** |
| Macaque and mouse | Indy | r2 | gpt | Claude | 33.3555 | 43.152 | 12 | -4.8005 | 0.000552868 | *** |
| Macaque and mouse | Jango | r2 | ndt1 | ndt2 | 76.7829 | 67.9014 | 14 | 2.7426 | 0.0167739 | * |
| Macaque and mouse | Jango | r2 | ndt1 | poyo | 76.7829 | 69.5986 | 14 | 1.9339 | 0.0752017 | n.s. |
| Macaque and mouse | Jango | r2 | ndt1 | UniBCI | 76.7829 | 60.2329 | 14 | 4.5369 | 0.000558335 | *** |
| Macaque and mouse | Jango | r2 | ndt1 | deepseek | 76.7829 | 84.3872 | 14 | -2.1972 | 0.0467383 | * |
| Macaque and mouse | Jango | r2 | ndt1 | gpt | 76.7829 | 83.2622 | 14 | -2.0009 | 0.0667318 | n.s. |
| Macaque and mouse | Jango | r2 | ndt1 | Claude | 76.7829 | 86.9754 | 14 | -2.8998 | 0.0124118 | * |
| Macaque and mouse | Jango | r2 | ndt2 | poyo | 67.9014 | 69.5986 | 14 | -0.3178 | 0.755659 | n.s. |
| Macaque and mouse | Jango | r2 | ndt2 | UniBCI | 67.9014 | 60.2329 | 14 | 1.4065 | 0.183021 | n.s. |
| Macaque and mouse | Jango | r2 | ndt2 | deepseek | 67.9014 | 84.3872 | 14 | -3.0786 | 0.00880123 | ** |
| Macaque and mouse | Jango | r2 | ndt2 | gpt | 67.9014 | 83.2622 | 14 | -2.9106 | 0.0121571 | * |
| Macaque and mouse | Jango | r2 | ndt2 | Claude | 67.9014 | 86.9754 | 14 | -3.4782 | 0.00408093 | ** |
| Macaque and mouse | Jango | r2 | poyo | UniBCI | 69.5986 | 60.2329 | 14 | 2.512 | 0.0259915 | * |
| Macaque and mouse | Jango | r2 | poyo | deepseek | 69.5986 | 84.3872 | 14 | -4.8772 | 0.000302128 | *** |
| Macaque and mouse | Jango | r2 | poyo | gpt | 69.5986 | 83.2622 | 14 | -4.8268 | 0.00033056 | *** |
| Macaque and mouse | Jango | r2 | poyo | Claude | 69.5986 | 86.9754 | 14 | -5.4877 | 0.000104313 | *** |
| Macaque and mouse | Jango | r2 | UniBCI | deepseek | 60.2329 | 84.3872 | 14 | -6.3893 | 2.38302e-05 | *** |
| Macaque and mouse | Jango | r2 | UniBCI | gpt | 60.2329 | 83.2622 | 14 | -6.2242 | 3.0965e-05 | *** |
| Macaque and mouse | Jango | r2 | UniBCI | Claude | 60.2329 | 86.9754 | 14 | -6.8965 | 1.09096e-05 | *** |
| Macaque and mouse | Jango | r2 | deepseek | gpt | 84.3872 | 83.2622 | 14 | 1.8882 | 0.0815233 | n.s. |
| Macaque and mouse | Jango | r2 | deepseek | Claude | 84.3872 | 86.9754 | 14 | -5.6122 | 8.44934e-05 | *** |
| Macaque and mouse | Jango | r2 | gpt | Claude | 83.2622 | 86.9754 | 14 | -6.1202 | 3.65942e-05 | *** |
| Macaque and mouse | Visual Coding | accuracy | ndt1 | ndt2 | 15.2783 | 14.195 | 6 | 0.534 | 0.61623 | n.s. |
| Macaque and mouse | Visual Coding | accuracy | ndt1 | poyo | 15.2783 | 12.3583 | 6 | 0.9779 | 0.373047 | n.s. |
| Macaque and mouse | Visual Coding | accuracy | ndt1 | UniBCI | 15.2783 | 14.64 | 6 | 0.2673 | 0.799929 | n.s. |
| Macaque and mouse | Visual Coding | accuracy | ndt1 | deepseek | 15.2783 | 13.0833 | 6 | 1.1419 | 0.305202 | n.s. |
| Macaque and mouse | Visual Coding | accuracy | ndt1 | gpt | 15.2783 | 12.9167 | 6 | 1.6443 | 0.161028 | n.s. |
| Macaque and mouse | Visual Coding | accuracy | ndt1 | Claude | 15.2783 | 13.0556 | 6 | 1.0885 | 0.326041 | n.s. |
| Macaque and mouse | Visual Coding | accuracy | ndt2 | poyo | 14.195 | 12.3583 | 6 | 0.8705 | 0.423856 | n.s. |
| Macaque and mouse | Visual Coding | accuracy | ndt2 | UniBCI | 14.195 | 14.64 | 6 | -0.2256 | 0.830415 | n.s. |
| Macaque and mouse | Visual Coding | accuracy | ndt2 | deepseek | 14.195 | 13.0833 | 6 | 0.9234 | 0.398154 | n.s. |
| Macaque and mouse | Visual Coding | accuracy | ndt2 | gpt | 14.195 | 12.9167 | 6 | 0.8677 | 0.425248 | n.s. |
| Macaque and mouse | Visual Coding | accuracy | ndt2 | Claude | 14.195 | 13.0556 | 6 | 0.8192 | 0.449924 | n.s. |
| Macaque and mouse | Visual Coding | accuracy | poyo | UniBCI | 12.3583 | 14.64 | 6 | -1.227 | 0.274446 | n.s. |
| Macaque and mouse | Visual Coding | accuracy | poyo | deepseek | 12.3583 | 13.0833 | 6 | -0.3703 | 0.726311 | n.s. |
| Macaque and mouse | Visual Coding | accuracy | poyo | gpt | 12.3583 | 12.9167 | 6 | -0.2107 | 0.841457 | n.s. |
| Macaque and mouse | Visual Coding | accuracy | poyo | Claude | 12.3583 | 13.0556 | 6 | -0.2592 | 0.805833 | n.s. |
| Macaque and mouse | Visual Coding | accuracy | UniBCI | deepseek | 14.64 | 13.0833 | 6 | 1.6375 | 0.162456 | n.s. |
| Macaque and mouse | Visual Coding | accuracy | UniBCI | gpt | 14.64 | 12.9167 | 6 | 1.1037 | 0.319971 | n.s. |
| Macaque and mouse | Visual Coding | accuracy | UniBCI | Claude | 14.64 | 13.0556 | 6 | 0.8939 | 0.412323 | n.s. |

| | | | | | | | | | | |
|---|---|---|---|---|---|---|---|---|---|---|
| Macaque and mouse | Visual Coding | accuracy | deepseek | gpt | 13.0833 | 12.9167 | 6 | 0.1995 | 0.849767 | n.s. |
| Macaque and mouse | Visual Coding | accuracy | deepseek | Claude | 13.0833 | 13.0556 | 6 | 0.0271 | 0.979425 | n.s. |
| Macaque and mouse | Visual Coding | accuracy | gpt | Claude | 12.9167 | 13.0556 | 6 | -0.1472 | 0.888706 | n.s. |
| Single-task decoding setting | Finger PPC | accuracy | ndt1 | ndt2 | 33.6433 | 32.2067 | 3 | 4.1692 | 0.0529988 | n.s. |
| Single-task decoding setting | Finger PPC | accuracy | ndt1 | poyo | 33.6433 | 41.42 | 3 | -2.5769 | 0.123344 | n.s. |
| Single-task decoding setting | Finger PPC | accuracy | ndt1 | UniBCI | 33.6433 | 52.5567 | 3 | -1.6439 | 0.241924 | n.s. |
| Single-task decoding setting | Finger PPC | accuracy | ndt1 | deepseek | 33.6433 | 33.642 | 3 | 0.0003 | 0.999804 | n.s. |
| Single-task decoding setting | Finger PPC | accuracy | ndt1 | gpt | 33.6433 | 33.3816 | 3 | 0.0895 | 0.936861 | n.s. |
| Single-task decoding setting | Finger PPC | accuracy | ndt1 | Claude | 33.6433 | 32.2917 | 3 | 0.9233 | 0.453337 | n.s. |
| Single-task decoding setting | Finger PPC | accuracy | ndt2 | poyo | 32.2067 | 41.42 | 3 | -2.759 | 0.110097 | n.s. |
| Single-task decoding setting | Finger PPC | accuracy | ndt2 | UniBCI | 32.2067 | 52.5567 | 3 | -1.8072 | 0.212463 | n.s. |
| Single-task decoding setting | Finger PPC | accuracy | ndt2 | deepseek | 32.2067 | 33.642 | 3 | -0.2845 | 0.802779 | n.s. |
| Single-task decoding setting | Finger PPC | accuracy | ndt2 | gpt | 32.2067 | 33.3816 | 3 | -0.364 | 0.750722 | n.s. |
| Single-task decoding setting | Finger PPC | accuracy | ndt2 | Claude | 32.2067 | 32.2917 | 3 | -0.0754 | 0.946742 | n.s. |
| Single-task decoding setting | Finger PPC | accuracy | poyo | UniBCI | 41.42 | 52.5567 | 3 | -0.8556 | 0.48234 | n.s. |
| Single-task decoding setting | Finger PPC | accuracy | poyo | deepseek | 41.42 | 33.642 | 3 | 3.1161 | 0.0893898 | n.s. |
| Single-task decoding setting | Finger PPC | accuracy | poyo | gpt | 41.42 | 33.3816 | 3 | 18.4565 | 0.00292278 | ** |
| Single-task decoding setting | Finger PPC | accuracy | poyo | Claude | 41.42 | 32.2917 | 3 | 2.1119 | 0.169095 | n.s. |
| Single-task decoding setting | Finger PPC | accuracy | UniBCI | deepseek | 52.5567 | 33.642 | 3 | 1.5422 | 0.262961 | n.s. |
| Single-task decoding setting | Finger PPC | accuracy | UniBCI | gpt | 52.5567 | 33.3816 | 3 | 1.5211 | 0.267634 | n.s. |
| Single-task decoding setting | Finger PPC | accuracy | UniBCI | Claude | 52.5567 | 32.2917 | 3 | 1.9642 | 0.188457 | n.s. |
| Single-task decoding setting | Finger PPC | accuracy | deepseek | gpt | 33.642 | 33.3816 | 3 | 0.117 | 0.917515 | n.s. |
| Single-task decoding setting | Finger PPC | accuracy | deepseek | Claude | 33.642 | 32.2917 | 3 | 0.2368 | 0.83486 | n.s. |
| Single-task decoding setting | Finger PPC | accuracy | gpt | Claude | 33.3816 | 32.2917 | 3 | 0.263 | 0.81716 | n.s. |
| Single-task decoding setting | Reaching B | accuracy | ndt1 | ndt2 | 73.746 | 70.388 | 10 | 0.7174 | 0.491323 | n.s. |
| Single-task decoding setting | Reaching B | accuracy | ndt1 | poyo | 73.746 | 86.124 | 10 | -3.0761 | 0.0132228 | * |
| Single-task decoding setting | Reaching B | accuracy | ndt1 | UniBCI | 73.746 | 64.729 | 10 | 1.9464 | 0.0834416 | n.s. |
| Single-task decoding setting | Reaching B | accuracy | ndt1 | deepseek | 73.746 | 81.3972 | 10 | -2.0803 | 0.0672271 | n.s. |
| Single-task decoding setting | Reaching B | accuracy | ndt1 | gpt | 73.746 | 78.7837 | 10 | -1.4064 | 0.193172 | n.s. |
| Single-task decoding setting | Reaching B | accuracy | ndt1 | Claude | 73.746 | 80.5523 | 10 | -2.0134 | 0.0749201 | n.s. |
| Single-task decoding setting | Reaching B | accuracy | ndt2 | poyo | 70.388 | 86.124 | 10 | -3.1032 | 0.0126572 | * |
| Single-task decoding setting | Reaching B | accuracy | ndt2 | UniBCI | 70.388 | 64.729 | 10 | 1.393 | 0.197067 | n.s. |
| Single-task decoding setting | Reaching B | accuracy | ndt2 | deepseek | 70.388 | 81.3972 | 10 | -2.541 | 0.031661 | * |
| Single-task decoding setting | Reaching B | accuracy | ndt2 | gpt | 70.388 | 78.7837 | 10 | -1.8446 | 0.0981985 | n.s. |
| Single-task decoding setting | Reaching B | accuracy | ndt2 | Claude | 70.388 | 80.5523 | 10 | -2.4607 | 0.036118 | * |
| Single-task decoding setting | Reaching B | accuracy | poyo | UniBCI | 86.124 | 64.729 | 10 | 4.6076 | 0.00127661 | ** |
| Single-task decoding setting | Reaching B | accuracy | poyo | deepseek | 86.124 | 81.3972 | 10 | 1.9285 | 0.0858725 | n.s. |
| Single-task decoding setting | Reaching B | accuracy | poyo | gpt | 86.124 | 78.7837 | 10 | 2.9433 | 0.0163981 | * |
| Single-task decoding setting | Reaching B | accuracy | poyo | Claude | 86.124 | 80.5523 | 10 | 2.496 | 0.0340844 | * |
| Single-task decoding setting | Reaching B | accuracy | UniBCI | deepseek | 64.729 | 81.3972 | 10 | -4.5074 | 0.00147323 | ** |
| Single-task decoding setting | Reaching B | accuracy | UniBCI | gpt | 64.729 | 78.7837 | 10 | -4.3955 | 0.00173168 | ** |
| Single-task decoding setting | Reaching B | accuracy | UniBCI | Claude | 64.729 | 80.5523 | 10 | -4.8999 | 0.000847714 | *** |
| Single-task decoding setting | Reaching B | accuracy | deepseek | gpt | 81.3972 | 78.7837 | 10 | 2.4839 | 0.0347674 | * |
| Single-task decoding setting | Reaching B | accuracy | deepseek | Claude | 81.3972 | 80.5523 | 10 | 0.8155 | 0.435826 | n.s. |
| Single-task decoding setting | Reaching B | accuracy | gpt | Claude | 78.7837 | 80.5523 | 10 | -2.3915 | 0.0404586 | * |
| Single-task decoding setting | Indy | r2 | ndt1 | ndt2 | 50.405 | 43.3167 | 12 | 4.5224 | 0.00086864 | *** |
| Single-task decoding setting | Indy | r2 | ndt1 | poyo | 50.405 | 34.2567 | 12 | 4.5944 | 0.000771971 | *** |
| Single-task decoding setting | Indy | r2 | ndt1 | UniBCI | 50.405 | 37.2725 | 12 | 5.2498 | 0.00027269 | *** |
| Single-task decoding setting | Indy | r2 | ndt1 | deepseek | 50.405 | 38.5229 | 12 | 3.3774 | 0.00617108 | ** |
| Single-task decoding setting | Indy | r2 | ndt1 | gpt | 50.405 | 35.6175 | 12 | 4.1128 | 0.00172146 | ** |
| Single-task decoding setting | Indy | r2 | ndt1 | Claude | 50.405 | 41.5487 | 12 | 2.6531 | 0.0224622 | * |
| Single-task decoding setting | Indy | r2 | ndt2 | poyo | 43.3167 | 34.2567 | 12 | 2.5259 | 0.0281792 | * |
| Single-task decoding setting | Indy | r2 | ndt2 | UniBCI | 43.3167 | 37.2725 | 12 | 2.4342 | 0.0331626 | * |
| Single-task decoding setting | Indy | r2 | ndt2 | deepseek | 43.3167 | 38.5229 | 12 | 1.3681 | 0.198578 | n.s. |
| Single-task decoding setting | Indy | r2 | ndt2 | gpt | 43.3167 | 35.6175 | 12 | 2.3021 | 0.0418766 | * |
| Single-task decoding setting | Indy | r2 | ndt2 | Claude | 43.3167 | 41.5487 | 12 | 0.5505 | 0.592976 | n.s. |
| Single-task decoding setting | Indy | r2 | poyo | UniBCI | 34.2567 | 37.2725 | 12 | -0.6568 | 0.524817 | n.s. |
| Single-task decoding setting | Indy | r2 | poyo | deepseek | 34.2567 | 38.5229 | 12 | -0.9185 | 0.378054 | n.s. |
| Single-task decoding setting | Indy | r2 | poyo | gpt | 34.2567 | 35.6175 | 12 | -0.2992 | 0.770374 | n.s. |
| Single-task decoding setting | Indy | r2 | poyo | Claude | 34.2567 | 41.5487 | 12 | -1.5778 | 0.142906 | n.s. |
| Single-task decoding setting | Indy | r2 | UniBCI | deepseek | 37.2725 | 38.5229 | 12 | -0.2802 | 0.784538 | n.s. |
| Single-task decoding setting | Indy | r2 | UniBCI | gpt | 37.2725 | 35.6175 | 12 | 0.399 | 0.697556 | n.s. |
| Single-task decoding setting | Indy | r2 | UniBCI | Claude | 37.2725 | 41.5487 | 12 | -1.0067 | 0.335708 | n.s. |

| | | | | | | | | | | |
|---|---|---|---|---|---|---|---|---|---|---|
| Single-task decoding setting | Indy | r2 | deepseek | gpt | 38.5229 | 35.6175 | 12 | 3.3272 | 0.00674464 | ** |
| Single-task decoding setting | Indy | r2 | deepseek | Claude | 38.5229 | 41.5487 | 12 | -3.3349 | 0.00665365 | ** |
| Single-task decoding setting | Indy | r2 | gpt | Claude | 35.6175 | 41.5487 | 12 | -5.4709 | 0.000194585 | *** |
| Single-task decoding setting | Jango | r2 | ndt1 | ndt2 | 70.2293 | 63.1093 | 14 | 2.1819 | 0.0480653 | * |
| Single-task decoding setting | Jango | r2 | ndt1 | poyo | 70.2293 | 75.17 | 14 | -1.0663 | 0.305687 | n.s. |
| Single-task decoding setting | Jango | r2 | ndt1 | UniBCI | 70.2293 | 64.9436 | 14 | 1.2876 | 0.220324 | n.s. |
| Single-task decoding setting | Jango | r2 | ndt1 | deepseek | 70.2293 | 84.8031 | 14 | -3.7643 | 0.00236225 | ** |
| Single-task decoding setting | Jango | r2 | ndt1 | gpt | 70.2293 | 84.7268 | 14 | -3.8199 | 0.00212529 | ** |
| Single-task decoding setting | Jango | r2 | ndt1 | Claude | 70.2293 | 85.1124 | 14 | -3.9382 | 0.00169886 | ** |
| Single-task decoding setting | Jango | r2 | ndt2 | poyo | 63.1093 | 75.17 | 14 | -1.8975 | 0.0801985 | n.s. |
| Single-task decoding setting | Jango | r2 | ndt2 | UniBCI | 63.1093 | 64.9436 | 14 | -0.4057 | 0.691596 | n.s. |
| Single-task decoding setting | Jango | r2 | ndt2 | deepseek | 63.1093 | 84.8031 | 14 | -3.3583 | 0.00513801 | ** |
| Single-task decoding setting | Jango | r2 | ndt2 | gpt | 63.1093 | 84.7268 | 14 | -3.4131 | 0.00462402 | ** |
| Single-task decoding setting | Jango | r2 | ndt2 | Claude | 63.1093 | 85.1124 | 14 | -3.4911 | 0.00398134 | ** |
| Single-task decoding setting | Jango | r2 | poyo | UniBCI | 75.17 | 64.9436 | 14 | 2.1668 | 0.0494174 | * |
| Single-task decoding setting | Jango | r2 | poyo | deepseek | 75.17 | 84.8031 | 14 | -2.2553 | 0.0419948 | * |
| Single-task decoding setting | Jango | r2 | poyo | gpt | 75.17 | 84.7268 | 14 | -2.1705 | 0.0490785 | * |
| Single-task decoding setting | Jango | r2 | poyo | Claude | 75.17 | 85.1124 | 14 | -2.3007 | 0.0386061 | * |
| Single-task decoding setting | Jango | r2 | UniBCI | deepseek | 64.9436 | 84.8031 | 14 | -4.0786 | 0.00130446 | ** |
| Single-task decoding setting | Jango | r2 | UniBCI | gpt | 64.9436 | 84.7268 | 14 | -4.1103 | 0.0012293 | ** |
| Single-task decoding setting | Jango | r2 | UniBCI | Claude | 64.9436 | 85.1124 | 14 | -4.1184 | 0.0012106 | ** |
| Single-task decoding setting | Jango | r2 | deepseek | gpt | 84.8031 | 84.7268 | 14 | 0.1546 | 0.879542 | n.s. |
| Single-task decoding setting | Jango | r2 | deepseek | Claude | 84.8031 | 85.1124 | 14 | -0.5422 | 0.596828 | n.s. |
| Single-task decoding setting | Jango | r2 | gpt | Claude | 84.7268 | 85.1124 | 14 | -0.9643 | 0.352507 | n.s. |
| Single-task decoding setting | Visual Coding | accuracy | ndt1 | ndt2 | 13.4733 | 16.0017 | 6 | -1.3433 | 0.236914 | n.s. |
| Single-task decoding setting | Visual Coding | accuracy | ndt1 | poyo | 13.4733 | 15.7783 | 6 | -1.0686 | 0.334097 | n.s. |
| Single-task decoding setting | Visual Coding | accuracy | ndt1 | UniBCI | 13.4733 | 16.4717 | 6 | -1.0985 | 0.322058 | n.s. |
| Single-task decoding setting | Visual Coding | accuracy | ndt1 | deepseek | 13.4733 | 14.3333 | 6 | -0.6047 | 0.571759 | n.s. |
| Single-task decoding setting | Visual Coding | accuracy | ndt1 | gpt | 13.4733 | 10.9722 | 6 | 3.176 | 0.0246488 | * |
| Single-task decoding setting | Visual Coding | accuracy | ndt1 | Claude | 13.4733 | 15.1389 | 6 | -0.9353 | 0.392558 | n.s. |
| Single-task decoding setting | Visual Coding | accuracy | ndt2 | poyo | 16.0017 | 15.7783 | 6 | 0.0771 | 0.941563 | n.s. |
| Single-task decoding setting | Visual Coding | accuracy | ndt2 | UniBCI | 16.0017 | 16.4717 | 6 | -0.1139 | 0.913733 | n.s. |
| Single-task decoding setting | Visual Coding | accuracy | ndt2 | deepseek | 16.0017 | 14.3333 | 6 | 0.5929 | 0.579057 | n.s. |
| Single-task decoding setting | Visual Coding | accuracy | ndt2 | gpt | 16.0017 | 10.9722 | 6 | 2.0483 | 0.0958464 | n.s. |
| Single-task decoding setting | Visual Coding | accuracy | ndt2 | Claude | 16.0017 | 15.1389 | 6 | 0.2816 | 0.789498 | n.s. |
| Single-task decoding setting | Visual Coding | accuracy | poyo | UniBCI | 15.7783 | 16.4717 | 6 | -0.2333 | 0.824745 | n.s. |
| Single-task decoding setting | Visual Coding | accuracy | poyo | deepseek | 15.7783 | 14.3333 | 6 | 0.5116 | 0.630722 | n.s. |
| Single-task decoding setting | Visual Coding | accuracy | poyo | gpt | 15.7783 | 10.9722 | 6 | 2.236 | 0.0755962 | n.s. |
| Single-task decoding setting | Visual Coding | accuracy | poyo | Claude | 15.7783 | 15.1389 | 6 | 0.1918 | 0.855456 | n.s. |
| Single-task decoding setting | Visual Coding | accuracy | UniBCI | deepseek | 16.4717 | 14.3333 | 6 | 1.2739 | 0.258686 | n.s. |
| Single-task decoding setting | Visual Coding | accuracy | UniBCI | gpt | 16.4717 | 10.9722 | 6 | 2.4257 | 0.0596955 | n.s. |
| Single-task decoding setting | Visual Coding | accuracy | UniBCI | Claude | 16.4717 | 15.1389 | 6 | 0.6884 | 0.521821 | n.s. |
| Single-task decoding setting | Visual Coding | accuracy | deepseek | gpt | 14.3333 | 10.9722 | 6 | 2.9491 | 0.0319172 | * |
| Single-task decoding setting | Visual Coding | accuracy | deepseek | Claude | 14.3333 | 15.1389 | 6 | -1.4555 | 0.205301 | n.s. |
| Single-task decoding setting | Visual Coding | accuracy | gpt | Claude | 10.9722 | 15.1389 | 6 | -2.838 | 0.036332 | * |
| Species across human, macaque and m... | Finger PPC | accuracy | ndt1 | ndt2 | 28.59 | 31.0567 | 3 | -0.5606 | 0.63148 | n.s. |
| Species across human, macaque and m... | Finger PPC | accuracy | ndt1 | poyo | 28.59 | 29.99 | 3 | -0.5521 | 0.636358 | n.s. |
| Species across human, macaque and m... | Finger PPC | accuracy | ndt1 | UniBCI | 28.59 | 34.4133 | 3 | -0.5781 | 0.621594 | n.s. |
| Species across human, macaque and m... | Finger PPC | accuracy | ndt1 | deepseek | 28.59 | 30.5459 | 3 | -0.2736 | 0.810048 | n.s. |
| Species across human, macaque and m... | Finger PPC | accuracy | ndt1 | gpt | 28.59 | 35.137 | 3 | -1.2232 | 0.345825 | n.s. |
| Species across human, macaque and m... | Finger PPC | accuracy | ndt1 | Claude | 28.59 | 31.1536 | 3 | -0.3984 | 0.72882 | n.s. |
| Species across human, macaque and m... | Finger PPC | accuracy | ndt2 | poyo | 31.0567 | 29.99 | 3 | 0.535 | 0.646175 | n.s. |
| Species across human, macaque and m... | Finger PPC | accuracy | ndt2 | UniBCI | 31.0567 | 34.4133 | 3 | -0.2905 | 0.798796 | n.s. |
| Species across human, macaque and m... | Finger PPC | accuracy | ndt2 | deepseek | 31.0567 | 30.5459 | 3 | 0.1817 | 0.872555 | n.s. |
| Species across human, macaque and m... | Finger PPC | accuracy | ndt2 | gpt | 31.0567 | 35.137 | 3 | -2.5832 | 0.122843 | n.s. |
| Species across human, macaque and m... | Finger PPC | accuracy | ndt2 | Claude | 31.0567 | 31.1536 | 3 | -0.0431 | 0.96953 | n.s. |
| Species across human, macaque and m... | Finger PPC | accuracy | poyo | UniBCI | 29.99 | 34.4133 | 3 | -0.4329 | 0.707287 | n.s. |
| Species across human, macaque and m... | Finger PPC | accuracy | poyo | deepseek | 29.99 | 30.5459 | 3 | -0.1199 | 0.915495 | n.s. |
| Species across human, macaque and m... | Finger PPC | accuracy | poyo | gpt | 29.99 | 35.137 | 3 | -1.8239 | 0.209737 | n.s. |
| Species across human, macaque and m... | Finger PPC | accuracy | poyo | Claude | 29.99 | 31.1536 | 3 | -0.2768 | 0.807934 | n.s. |
| Species across human, macaque and m... | Finger PPC | accuracy | UniBCI | deepseek | 34.4133 | 30.5459 | 3 | 0.3075 | 0.787542 | n.s. |
| Species across human, macaque and m... | Finger PPC | accuracy | UniBCI | gpt | 34.4133 | 35.137 | 3 | -0.0667 | 0.952863 | n.s. |
| Species across human, macaque and m... | Finger PPC | accuracy | UniBCI | Claude | 34.4133 | 31.1536 | 3 | 0.2402 | 0.832518 | n.s. |

| | | | | | | | | | | |
|---|---|---|---|---|---|---|---|---|---|---|
| Species across human, macaque and m... | Finger PPC | accuracy | deepseek | gpt | 30.5459 | 35.137 | 3 | -2.2533 | 0.152996 | n.s. |
| Species across human, macaque and m... | Finger PPC | accuracy | deepseek | Claude | 30.5459 | 31.1536 | 3 | -0.2961 | 0.795093 | n.s. |
| Species across human, macaque and m... | Finger PPC | accuracy | gpt | Claude | 35.137 | 31.1536 | 3 | 1.4318 | 0.288528 | n.s. |
| Species across human, macaque and m... | Reaching B | accuracy | ndt1 | ndt2 | 52.48 | 66.963 | 10 | -2.7818 | 0.0213377 | * |
| Species across human, macaque and m... | Reaching B | accuracy | ndt1 | poyo | 52.48 | 58.161 | 10 | -1.3752 | 0.202327 | n.s. |
| Species across human, macaque and m... | Reaching B | accuracy | ndt1 | UniBCI | 52.48 | 60.495 | 10 | -1.7425 | 0.115395 | n.s. |
| Species across human, macaque and m... | Reaching B | accuracy | ndt1 | deepseek | 52.48 | 81.1879 | 10 | -7.783 | 2.75554e-05 | *** |
| Species across human, macaque and m... | Reaching B | accuracy | ndt1 | gpt | 52.48 | 78.7188 | 10 | -6.4943 | 0.00011219 | *** |
| Species across human, macaque and m... | Reaching B | accuracy | ndt1 | Claude | 52.48 | 78.48 | 10 | -6.4137 | 0.000123274 | *** |
| Species across human, macaque and m... | Reaching B | accuracy | ndt2 | poyo | 66.963 | 58.161 | 10 | 1.7546 | 0.113227 | n.s. |
| Species across human, macaque and m... | Reaching B | accuracy | ndt2 | UniBCI | 66.963 | 60.495 | 10 | 1.6234 | 0.138944 | n.s. |
| Species across human, macaque and m... | Reaching B | accuracy | ndt2 | deepseek | 66.963 | 81.1879 | 10 | -4.5159 | 0.00145524 | ** |
| Species across human, macaque and m... | Reaching B | accuracy | ndt2 | gpt | 66.963 | 78.7188 | 10 | -3.8831 | 0.00371358 | ** |
| Species across human, macaque and m... | Reaching B | accuracy | ndt2 | Claude | 66.963 | 78.48 | 10 | -3.7111 | 0.00483648 | ** |
| Species across human, macaque and m... | Reaching B | accuracy | poyo | UniBCI | 58.161 | 60.495 | 10 | -0.6903 | 0.507403 | n.s. |
| Species across human, macaque and m... | Reaching B | accuracy | poyo | deepseek | 58.161 | 81.1879 | 10 | -6.3963 | 0.000125817 | *** |
| Species across human, macaque and m... | Reaching B | accuracy | poyo | gpt | 58.161 | 78.7188 | 10 | -4.1837 | 0.00236299 | ** |
| Species across human, macaque and m... | Reaching B | accuracy | poyo | Claude | 58.161 | 78.48 | 10 | -4.4642 | 0.0015677 | ** |
| Species across human, macaque and m... | Reaching B | accuracy | UniBCI | deepseek | 60.495 | 81.1879 | 10 | -6.1601 | 0.000166674 | *** |
| Species across human, macaque and m... | Reaching B | accuracy | UniBCI | gpt | 60.495 | 78.7188 | 10 | -4.5829 | 0.00132226 | ** |
| Species across human, macaque and m... | Reaching B | accuracy | UniBCI | Claude | 60.495 | 78.48 | 10 | -5.0314 | 0.000707973 | *** |
| Species across human, macaque and m... | Reaching B | accuracy | deepseek | gpt | 81.1879 | 78.7188 | 10 | 1.5158 | 0.163874 | n.s. |
| Species across human, macaque and m... | Reaching B | accuracy | deepseek | Claude | 81.1879 | 78.48 | 10 | 1.6264 | 0.138305 | n.s. |
| Species across human, macaque and m... | Reaching B | accuracy | gpt | Claude | 78.7188 | 78.48 | 10 | 0.2615 | 0.799556 | n.s. |
| Species across human, macaque and m... | Indy | r2 | ndt1 | ndt2 | 51.8 | 43.7467 | 12 | 3.8868 | 0.0025334 | ** |
| Species across human, macaque and m... | Indy | r2 | ndt1 | poyo | 51.8 | 33.4433 | 12 | 5.3086 | 0.000249106 | *** |
| Species across human, macaque and m... | Indy | r2 | ndt1 | UniBCI | 51.8 | 36.2875 | 12 | 6.4777 | 4.56513e-05 | *** |
| Species across human, macaque and m... | Indy | r2 | ndt1 | deepseek | 51.8 | 32.724 | 12 | 4.6832 | 0.000668099 | *** |
| Species across human, macaque and m... | Indy | r2 | ndt1 | gpt | 51.8 | 37.9186 | 12 | 5.2669 | 0.000265567 | *** |
| Species across human, macaque and m... | Indy | r2 | ndt1 | Claude | 51.8 | 44.3597 | 12 | 3.4333 | 0.00559087 | ** |
| Species across human, macaque and m... | Indy | r2 | ndt2 | poyo | 43.7467 | 33.4433 | 12 | 2.6441 | 0.0228237 | * |
| Species across human, macaque and m... | Indy | r2 | ndt2 | UniBCI | 43.7467 | 36.2875 | 12 | 4.5364 | 0.000848835 | *** |
| Species across human, macaque and m... | Indy | r2 | ndt2 | deepseek | 43.7467 | 32.724 | 12 | 2.4161 | 0.0342443 | * |
| Species across human, macaque and m... | Indy | r2 | ndt2 | gpt | 43.7467 | 37.9186 | 12 | 1.8434 | 0.0923488 | n.s. |
| Species across human, macaque and m... | Indy | r2 | ndt2 | Claude | 43.7467 | 44.3597 | 12 | -0.2126 | 0.835489 | n.s. |
| Species across human, macaque and m... | Indy | r2 | poyo | UniBCI | 33.4433 | 36.2875 | 12 | -0.7977 | 0.44194 | n.s. |
| Species across human, macaque and m... | Indy | r2 | poyo | deepseek | 33.4433 | 32.724 | 12 | 0.157 | 0.878108 | n.s. |
| Species across human, macaque and m... | Indy | r2 | poyo | gpt | 33.4433 | 37.9186 | 12 | -1.1055 | 0.292537 | n.s. |
| Species across human, macaque and m... | Indy | r2 | poyo | Claude | 33.4433 | 44.3597 | 12 | -2.5713 | 0.0259921 | * |
| Species across human, macaque and m... | Indy | r2 | UniBCI | deepseek | 36.2875 | 32.724 | 12 | 0.7297 | 0.480842 | n.s. |
| Species across human, macaque and m... | Indy | r2 | UniBCI | gpt | 36.2875 | 37.9186 | 12 | -0.4483 | 0.662671 | n.s. |
| Species across human, macaque and m... | Indy | r2 | UniBCI | Claude | 36.2875 | 44.3597 | 12 | -2.3413 | 0.0390846 | * |
| Species across human, macaque and m... | Indy | r2 | deepseek | gpt | 32.724 | 37.9186 | 12 | -2.724 | 0.0197892 | * |
| Species across human, macaque and m... | Indy | r2 | deepseek | Claude | 32.724 | 44.3597 | 12 | -4.7224 | 0.000626964 | *** |
| Species across human, macaque and m... | Indy | r2 | gpt | Claude | 37.9186 | 44.3597 | 12 | -5.5383 | 0.00017581 | *** |
| Species across human, macaque and m... | Jango | r2 | ndt1 | ndt2 | 75.9607 | 64.6871 | 14 | 5.8761 | 5.44599e-05 | *** |
| Species across human, macaque and m... | Jango | r2 | ndt1 | poyo | 75.9607 | 67.065 | 14 | 1.7369 | 0.106015 | n.s. |
| Species across human, macaque and m... | Jango | r2 | ndt1 | UniBCI | 75.9607 | 64.13 | 14 | 2.5202 | 0.0255933 | * |
| Species across human, macaque and m... | Jango | r2 | ndt1 | deepseek | 75.9607 | 84.6062 | 14 | -1.6381 | 0.12536 | n.s. |
| Species across human, macaque and m... | Jango | r2 | ndt1 | gpt | 75.9607 | 85.4105 | 14 | -1.7951 | 0.0959057 | n.s. |
| Species across human, macaque and m... | Jango | r2 | ndt1 | Claude | 75.9607 | 85.2243 | 14 | -1.775 | 0.099294 | n.s. |
| Species across human, macaque and m... | Jango | r2 | ndt2 | poyo | 64.6871 | 67.065 | 14 | -0.4023 | 0.694013 | n.s. |
| Species across human, macaque and m... | Jango | r2 | ndt2 | UniBCI | 64.6871 | 64.13 | 14 | 0.1064 | 0.916902 | n.s. |
| Species across human, macaque and m... | Jango | r2 | ndt2 | deepseek | 64.6871 | 84.6062 | 14 | -3.4982 | 0.00392734 | ** |
| Species across human, macaque and m... | Jango | r2 | ndt2 | gpt | 64.6871 | 85.4105 | 14 | -3.6103 | 0.00316885 | ** |
| Species across human, macaque and m... | Jango | r2 | ndt2 | Claude | 64.6871 | 85.2243 | 14 | -3.6498 | 0.00293873 | ** |
| Species across human, macaque and m... | Jango | r2 | poyo | UniBCI | 67.065 | 64.13 | 14 | 0.9064 | 0.381215 | n.s. |
| Species across human, macaque and m... | Jango | r2 | poyo | deepseek | 67.065 | 84.6062 | 14 | -5.0162 | 0.000236088 | *** |
| Species across human, macaque and m... | Jango | r2 | poyo | gpt | 67.065 | 85.4105 | 14 | -5.3628 | 0.000129114 | *** |
| Species across human, macaque and m... | Jango | r2 | poyo | Claude | 67.065 | 85.2243 | 14 | -5.2128 | 0.000167326 | *** |
| Species across human, macaque and m... | Jango | r2 | UniBCI | deepseek | 64.13 | 84.6062 | 14 | -5.8405 | 5.77562e-05 | *** |
| Species across human, macaque and m... | Jango | r2 | UniBCI | gpt | 64.13 | 85.4105 | 14 | -6.1139 | 3.69665e-05 | *** |
| Species across human, macaque and m... | Jango | r2 | UniBCI | Claude | 64.13 | 85.2243 | 14 | -5.9986 | 4.4562e-05 | *** |

| | | | | | | | | | | |
|---|---|---|---|---|---|---|---|---|---|---|
| Species across human, macaque and m... | Jango | r2 | deepseek | gpt | 84.6062 | 85.4105 | 14 | -1.7584 | 0.102188 | n.s. |
| Species across human, macaque and m... | Jango | r2 | deepseek | Claude | 84.6062 | 85.2243 | 14 | -2.672 | 0.0191898 | * |
| Species across human, macaque and m... | Jango | r2 | gpt | Claude | 85.4105 | 85.2243 | 14 | 0.4265 | 0.676743 | n.s. |
| Species across human, macaque and m... | Visual Coding | accuracy | ndt1 | ndt2 | 14.75 | 15.1117 | 6 | -0.2767 | 0.793065 | n.s. |
| Species across human, macaque and m... | Visual Coding | accuracy | ndt1 | poyo | 14.75 | 13.835 | 6 | 0.4109 | 0.698132 | n.s. |
| Species across human, macaque and m... | Visual Coding | accuracy | ndt1 | UniBCI | 14.75 | 12.5 | 6 | 1.2126 | 0.279434 | n.s. |
| Species across human, macaque and m... | Visual Coding | accuracy | ndt1 | deepseek | 14.75 | 13.1111 | 6 | 0.833 | 0.442778 | n.s. |
| Species across human, macaque and m... | Visual Coding | accuracy | ndt1 | gpt | 14.75 | 12.1666 | 6 | 1.7861 | 0.134138 | n.s. |
| Species across human, macaque and m... | Visual Coding | accuracy | ndt1 | Claude | 14.75 | 13.1945 | 6 | 1.0884 | 0.326086 | n.s. |
| Species across human, macaque and m... | Visual Coding | accuracy | ndt2 | poyo | 15.1117 | 13.835 | 6 | 0.6842 | 0.524272 | n.s. |
| Species across human, macaque and m... | Visual Coding | accuracy | ndt2 | UniBCI | 15.1117 | 12.5 | 6 | 1.2005 | 0.283704 | n.s. |
| Species across human, macaque and m... | Visual Coding | accuracy | ndt2 | deepseek | 15.1117 | 13.1111 | 6 | 0.9505 | 0.385508 | n.s. |
| Species across human, macaque and m... | Visual Coding | accuracy | ndt2 | gpt | 15.1117 | 12.1666 | 6 | 1.6371 | 0.16254 | n.s. |
| Species across human, macaque and m... | Visual Coding | accuracy | ndt2 | Claude | 15.1117 | 13.1945 | 6 | 1.2406 | 0.269784 | n.s. |
| Species across human, macaque and m... | Visual Coding | accuracy | poyo | UniBCI | 13.835 | 12.5 | 6 | 0.4264 | 0.687562 | n.s. |
| Species across human, macaque and m... | Visual Coding | accuracy | poyo | deepseek | 13.835 | 13.1111 | 6 | 0.3372 | 0.749633 | n.s. |
| Species across human, macaque and m... | Visual Coding | accuracy | poyo | gpt | 13.835 | 12.1666 | 6 | 0.7619 | 0.480471 | n.s. |
| Species across human, macaque and m... | Visual Coding | accuracy | poyo | Claude | 13.835 | 13.1945 | 6 | 0.2888 | 0.784312 | n.s. |
| Species across human, macaque and m... | Visual Coding | accuracy | UniBCI | deepseek | 12.5 | 13.1111 | 6 | -0.3186 | 0.762937 | n.s. |
| Species across human, macaque and m... | Visual Coding | accuracy | UniBCI | gpt | 12.5 | 12.1666 | 6 | 0.1859 | 0.859815 | n.s. |
| Species across human, macaque and m... | Visual Coding | accuracy | UniBCI | Claude | 12.5 | 13.1945 | 6 | -0.5938 | 0.578487 | n.s. |
| Species across human, macaque and m... | Visual Coding | accuracy | deepseek | gpt | 13.1111 | 12.1666 | 6 | 1.4148 | 0.216289 | n.s. |
| Species across human, macaque and m... | Visual Coding | accuracy | deepseek | Claude | 13.1111 | 13.1945 | 6 | -0.0706 | 0.946456 | n.s. |
| Species across human, macaque and m... | Visual Coding | accuracy | gpt | Claude | 12.1666 | 13.1945 | 6 | -1.041 | 0.345576 | n.s. |
