## Supplementary File 2 for "BCIJelly: An integrated ecosystem for brain–computer interface research"

| Task | Dataset | Metric | Day_ or_ session_ id | NDT1 | NDT2 | POYO | UniBCI | AAS with DeepSeek-V4 Pro | AAS with ChatGPT-5.5 | AAS with Claude Opus-4.7 |
|---|---|---|---|---|---|---|---|---|---|---|
| Regression | Jango | r2 | 20150808 | 0.9423 | 0.819 | 0.8882 | 0.9116 | 0.907077 | 0.90486 | 0.914832 |
| Regression | Jango | r2 | 20150809 | 0.8717 | 0.6612 | 0.8379 | 0.8091 | 0.915925 | 0.916348 | 0.91279 |
| Regression | Jango | r2 | 20150820 | 0.8835 | 0.7664 | 0.8822 | 0.807 | 0.913678 | 0.910817 | 0.916626 |
| Regression | Jango | r2 | 20150824 | 0.9118 | 0.7483 | 0.8976 | 0.8387 | 0.908803 | 0.911523 | 0.911028 |
| Regression | Jango | r2 | 20150825 | 0.7689 | 0.3903 | 0.8465 | 0.7142 | 0.792943 | 0.850305 | 0.855014 |
| Regression | Jango | r2 | 20150826 | 0.8233 | 0.656 | 0.8715 | 0.8751 | 0.907708 | 0.917819 | 0.909687 |
| Regression | Jango | r2 | 20150827 | 0.8629 | 0.4444 | 0.8813 | 0.8762 | 0.909907 | 0.90928 | 0.915367 |
| Regression | Jango | r2 | 20150828 | 0.9141 | 0.4516 | 0.8326 | 0.8626 | 0.879547 | 0.878497 | 0.89192 |
| Regression | Jango | r2 | 20150831 | 0.6553 | 0.6734 | 0.6083 | 0.5396 | 0.904888 | 0.901807 | 0.915672 |
| Regression | Jango | r2 | 20150905 | 0.7624 | 0.4489 | 0.422 | 0.7796 | 0.801741 | 0.798409 | 0.805841 |
| Regression | Jango | r2 | 20150906 | 0.9193 | 0.5303 | 0.6061 | 0.779 | 0.830254 | 0.827437 | 0.855902 |
| Regression | Jango | r2 | 20150908 | 0 | 0 | 0.5296 | 0 | 0.826046 | 0.823548 | 0.850514 |
| Regression | Jango | r2 | 20151029 | 0.6954 | 0.6166 | 0.6793 | 0.5935 | 0.836662 | 0.835018 | 0.824554 |
| Regression | Jango | r2 | 20151102 | 0.3869 | 0.1447 | 0.443 | 0.2077 | 0.671884 | 0.620541 | 0.656269 |
| Regression | Indy | r2 | 20161011 | 0.6872 | 0.6027 | 0.3455 | 0.5544 | 0.541927 | 0.5467 | 0.591003 |
| Regression | Indy | r2 | 20161013 | 0.0075 | 0 | 0 | 0 | -0.139429 | -0.129595 | -0.109922 |
| Regression | Indy | r2 | 20161014 | 0.6002 | 0.6523 | 0.3738 | 0.5743 | 0.624304 | 0.62563 | 0.665755 |
| Regression | Indy | r2 | 20161017 | 0.471 | 0.5045 | 0.2347 | 0.299 | 0.46741 | 0.493461 | 0.529366 |
| Regression | Indy | r2 | 20161024 | 0.4107 | 0.3214 | 0.3686 | 0.2485 | 0.41649 | 0.449385 | 0.480049 |
| Regression | Indy | r2 | 20161025 | 0.5536 | 0.5456 | 0.4794 | 0.5211 | 0.555445 | 0.57566 | 0.607862 |
| Regression | Indy | r2 | 20161026 | 0.567 | 0.4997 | 0.4968 | 0.3841 | 0.565776 | 0.58247 | 0.634414 |
| Regression | Indy | r2 | 20161027 | 0.502 | 0.3589 | 0.2698 | 0.3936 | 0.171433 | 0.144781 | 0.221012 |
| Regression | Indy | r2 | 20161206 | 0.3094 | 0.2796 | 0.3582 | 0.1957 | 0.261235 | 0.255683 | 0.291421 |
| Regression | Indy | r2 | 20161207 | 0.4229 | 0.2442 | 0.3104 | 0.3113 | 0.317012 | 0.30908 | 0.339439 |
| Regression | Indy | r2 | 20161212 | 0.5793 | 0.4741 | 0.3258 | 0.3937 | 0.473994 | 0.460954 | 0.535199 |
| Regression | Indy | r2 | 20161220 | 0.3307 | 0.298 | 0.1015 | 0.2679 | 0.305486 | 0.313644 | 0.36881 |
| Classification | Finger PPC | accuracy | 20181015 | 0.2963 | 0.1898 | 0.2269 | 0.3843 | 0.168981 | 0.199074 | 0.168981 |
| Classification | Finger PPC | accuracy | 20181017 | 0.4714 | 0.375 | 0.4141 | 0.3281 | 0.429688 | 0.341146 | 0.424479 |
| Classification | Finger PPC | accuracy | 20181022 | 0.3177 | 0.4271 | 0.4688 | 0.4349 | 0.450521 | 0.429688 | 0.377604 |
| Classification | Visual Coding | accuracy | session_763673393 | 0.1267 | 0.185 | 0.11 | 0.1233 | 0.133333 | 0.126667 | 0.116667 |
| Classification | Visual Coding | accuracy | session_773418906 | 0.1217 | 0.1283 | 0.105 | 0.1633 | 0.116667 | 0.14 | 0.111667 |
| Classification | Visual Coding | accuracy | session_791319847 | 0.1183 | 0.18 | 0.2217 | 0.0717 | 0.113333 | 0.143333 | 0.146667 |
| Classification | Visual Coding | accuracy | session_797828357 | 0.2133 | 0.2633 | 0.1883 | 0.1117 | 0.141667 | 0.125 | 0.123333 |
| Classification | Visual Coding | accuracy | session_798911424 | 0.1267 | 0.0883 | 0.1333 | 0.1483 | 0.13 | 0.125 | 0.11 |
| Classification | Visual Coding | accuracy | session_799864342 | 0.0267 | 0.0833 | 0.0567 | 0.08 | 0.121667 | 0.123333 | 0.088333 |
| Classification | Reaching B | accuracy | 20221129 | 0.905 | 0.84 | 0.8267 | 0.7767 | 0.833333 | 0.821667 | 0.806667 |
| Classification | Reaching B | accuracy | 20221202 | 0.9125 | 0.7188 | 0.8625 | 0.82 | 0.8625 | 0.8625 | 0.9025 |
| Classification | Reaching B | accuracy | 20221205 | 0.87 | 0.82 | 0.93 | 0.745 | 0.815 | 0.89 | 0.86 |
| Classification | Reaching B | accuracy | 20221208 | 0.845 | 0.825 | 0.95 | 0.705 | 0.855 | 0.855 | 0.868333 |
| Classification | Reaching B | accuracy | 20221209 | 0.7425 | 0.89 | 0.8275 | 0.7775 | 0.8625 | 0.845 | 0.8475 |
| Classification | Reaching B | accuracy | 20221213 | 0.7725 | 0.5425 | 0.7375 | 0.56 | 0.845 | 0.7775 | 0.83 |
| Classification | Reaching B | accuracy | 20221214 | 0.77 | 0.62 | 0.765 | 0.565 | 0.77 | 0.815 | 0.795 |
| Classification | Reaching B | accuracy | 20221215 | 0.5577 | 0.4343 | 0.5626 | 0.541 | 0.520868 | 0.494157 | 0.539232 |
| Classification | Reaching B | accuracy | 20221219 | 0.6336 | 0.5464 | 0.7979 | 0.5529 | 0.794286 | 0.731429 | 0.76 |
| Classification | Reaching B | accuracy | 20221230 | 0.541 | 0.273 | 0.752 | 0.64 | 0.767 | 0.721 | 0.777 |
| Human and monkey | Finger PPC | accuracy | 20181015 | 0.2708 | 0.2106 | 0.0949 | 0.3773 | 0.136574 | 0.152778 | 0.236111 |
| Human and monkey | Finger PPC | accuracy | 20181017 | 0.3438 | 0.3099 | 0.1667 | 0.2135 | 0.403646 | 0.432292 | 0.460938 |
| Human and monkey | Finger PPC | accuracy | 20181022 | 0.3516 | 0.375 | 0.2396 | 0.3984 | 0.364583 | 0.429688 | 0.429688 |
| Human and monkey | Jango | r2 | 20150808 | 0.9317 | 0.8988 | 0.9182 | 0.7739 | 0.902536 | 0.888185 | 0.901013 |
| Human and monkey | Jango | r2 | 20150809 | 0.7533 | 0.7093 | 0.8888 | 0.8543 | 0.903565 | 0.895499 | 0.910525 |
| Human and monkey | Jango | r2 | 20150820 | 0.7457 | 0.7667 | 0.0458 | 0.8172 | 0.903322 | 0.889767 | 0.896579 |
| Human and monkey | Jango | r2 | 20150824 | 0.8349 | 0.7885 | 0.8893 | 0.8332 | 0.900417 | 0.884307 | 0.902318 |
| Human and monkey | Jango | r2 | 20150825 | 0.6454 | 0.4856 | 0.6933 | 0.6613 | 0.769357 | 0.803309 | 0.794789 |
| Human and monkey | Jango | r2 | 20150826 | 0.6803 | 0.635 | 0.867 | 0.6932 | 0.89654 | 0.897601 | 0.899296 |
| Human and monkey | Jango | r2 | 20150827 | 0.6523 | 0.5163 | 0.8348 | 0.7256 | 0.895816 | 0.882293 | 0.890958 |
| Human and monkey | Jango | r2 | 20150828 | 0.7326 | 0.5809 | 0.8133 | 0.7848 | 0.862157 | 0.860161 | 0.887532 |
| Human and monkey | Jango | r2 | 20150831 | 0.7635 | 0.8201 | 0.6097 | 0.1459 | 0.898023 | 0.885914 | 0.899461 |
| Human and monkey | Jango | r2 | 20150905 | 0.4853 | 0.5552 | 0.5616 | 0.3703 | 0.767729 | 0.79354 | 0.800308 |
| Human and monkey | Jango | r2 | 20150906 | 0.726 | 0.6394 | 0.2786 | 0.5225 | 0.828629 | 0.80808 | 0.839091 |
| Human and monkey | Jango | r2 | 20150908 | 0.2532 | 0 | 0 | 0.5349 | 0.81512 | 0.811041 | 0.759965 |

| | | | | | | | | | | |
|---|---|---|---|---|---|---|---|---|---|---|
| Human and monkey | Jango | r2 | 20151029 | 0.7045 | 0.7333 | 0.6381 | 0.6652 | 0.833628 | 0.81455 | 0.844392 |
| Human and monkey | Jango | r2 | 20151102 | 0.3348 | 0.3307 | 0.4545 | 0.2158 | 0.713623 | 0.603145 | 0.641031 |
| Human and monkey | Indy | r2 | 20161011 | 0.6863 | 0.5892 | 0.3755 | 0.5954 | 0.512949 | 0.462814 | 0.575615 |
| Human and monkey | Indy | r2 | 20161013 | 0.0075 | 0 | 0 | 0 | -0.193416 | -0.111858 | -0.141614 |
| Human and monkey | Indy | r2 | 20161014 | 0.6721 | 0.6806 | 0.4528 | 0.6249 | 0.601105 | 0.599932 | 0.665751 |
| Human and monkey | Indy | r2 | 20161017 | 0.4788 | 0.3807 | 0.2707 | 0.2788 | 0.485818 | 0.440808 | 0.537764 |
| Human and monkey | Indy | r2 | 20161024 | 0.4248 | 0.3912 | 0.3349 | 0.2341 | 0.413544 | 0.419388 | 0.487236 |
| Human and monkey | Indy | r2 | 20161025 | 0.6492 | 0.54 | 0.4628 | 0.4937 | 0.579586 | 0.535047 | 0.625797 |
| Human and monkey | Indy | r2 | 20161026 | 0.5977 | 0.5374 | 0.4846 | 0.4153 | 0.55864 | 0.557397 | 0.639637 |
| Human and monkey | Indy | r2 | 20161027 | 0.4802 | 0.4707 | 0.2471 | 0.3747 | -0.06694 | 0.086519 | 0.230996 |
| Human and monkey | Indy | r2 | 20161206 | 0.4012 | 0.2826 | 0.376 | 0.1449 | 0.289183 | 0.214634 | 0.327469 |
| Human and monkey | Indy | r2 | 20161207 | 0.5258 | 0.4403 | 0.3132 | 0.3236 | 0.2912 | 0.187565 | 0.334237 |
| Human and monkey | Indy | r2 | 20161212 | 0.5748 | 0.5053 | 0.2791 | 0.3961 | 0.463964 | 0.454854 | 0.532124 |
| Human and monkey | Indy | r2 | 20161220 | 0.3749 | 0.2722 | 0.0862 | 0.1922 | 0.359706 | 0.253274 | 0.382743 |
| Human and monkey | Reaching B | accuracy | 20221129 | 0.7167 | 0.4633 | 0.47 | 0.6367 | 0.878333 | 0.88 | 0.823333 |
| Human and monkey | Reaching B | accuracy | 20221202 | 0.8075 | 0.6163 | 0.5975 | 0.7112 | 0.8825 | 0.90875 | 0.8875 |
| Human and monkey | Reaching B | accuracy | 20221205 | 0.49 | 0.65 | 0.45 | 0.605 | 0.91 | 0.87 | 0.9 |
| Human and monkey | Reaching B | accuracy | 20221208 | 0.7017 | 0.5233 | 0.47 | 0.5967 | 0.916667 | 0.91 | 0.903333 |
| Human and monkey | Reaching B | accuracy | 20221209 | 0.645 | 0.695 | 0.335 | 0.6825 | 0.865 | 0.8725 | 0.91 |
| Human and monkey | Reaching B | accuracy | 20221213 | 0.6375 | 0.67 | 0.73 | 0.5975 | 0.835 | 0.845 | 0.845 |
| Human and monkey | Reaching B | accuracy | 20221214 | 0.61 | 0.545 | 0.65 | 0.55 | 0.86 | 0.825 | 0.87 |
| Human and monkey | Reaching B | accuracy | 20221215 | 0.5611 | 0.4267 | 0.5159 | 0.4727 | 0.555927 | 0.529215 | 0.557596 |
| Human and monkey | Reaching B | accuracy | 20221219 | 0.5214 | 0.5779 | 0.7671 | 0.5829 | 0.798571 | 0.815 | 0.755714 |
| Human and monkey | Reaching B | accuracy | 20221230 | 0.646 | 0.443 | 0.673 | 0.528 | 0.791 | 0.803 | 0.772 |
| Human and mouse | Finger PPC | accuracy | 20181015 | 0.2083 | 0.1667 | 0.2315 | 0.5 | 0.208333 | 0.138889 | 0.171296 |
| Human and mouse | Finger PPC | accuracy | 20181017 | 0.4323 | 0.4766 | 0.4922 | 0.3646 | 0.411458 | 0.359375 | 0.398438 |
| Human and mouse | Finger PPC | accuracy | 20181022 | 0.375 | 0.4479 | 0.474 | 0.513 | 0.463542 | 0.380208 | 0.481771 |
| Human and mouse | Visual Coding | accuracy | session_763673393 | 0.1283 | 0.14 | 0.12 | 0.1267 | 0.141667 | 0.121667 | 0.12 |
| Human and mouse | Visual Coding | accuracy | session_773418906 | 0.105 | 0.1067 | 0.0667 | 0.115 | 0.086667 | 0.13 | 0.111667 |
| Human and mouse | Visual Coding | accuracy | session_791319847 | 0.1533 | 0.1317 | 0.115 | 0.0683 | 0.133333 | 0.116667 | 0.115 |
| Human and mouse | Visual Coding | accuracy | session_797828357 | 0.2283 | 0.2183 | 0.14 | 0.09 | 0.12 | 0.128333 | 0.143333 |
| Human and mouse | Visual Coding | accuracy | session_798911424 | 0.1283 | 0.13 | 0.1217 | 0.1733 | 0.115 | 0.118333 | 0.128333 |
| Human and mouse | Visual Coding | accuracy | session_799864342 | 0.085 | 0.1083 | 0.1033 | 0.165 | 0.095 | 0.111667 | 0.131667 |
| Monkey and mouse | Jango | r2 | 20150808 | 0.9447 | 0.8585 | 0.9194 | 0.7557 | 0.901583 | 0.90436 | 0.918294 |
| Monkey and mouse | Jango | r2 | 20150809 | 0.8499 | 0.7845 | 0.8769 | 0.8335 | 0.899807 | 0.896303 | 0.918954 |
| Monkey and mouse | Jango | r2 | 20150820 | 0.8818 | 0.7969 | 0.8065 | 0.6369 | 0.910411 | 0.870283 | 0.919586 |
| Monkey and mouse | Jango | r2 | 20150824 | 0.9295 | 0.8823 | 0.783 | 0.8437 | 0.878701 | 0.893044 | 0.916519 |
| Monkey and mouse | Jango | r2 | 20150825 | 0.8067 | 0.729 | 0.709 | 0.5511 | 0.788209 | 0.792656 | 0.843591 |
| Monkey and mouse | Jango | r2 | 20150826 | 0.7407 | 0.7815 | 0.7636 | 0.6655 | 0.898304 | 0.900388 | 0.916355 |
| Monkey and mouse | Jango | r2 | 20150827 | 0.8209 | 0.7285 | 0.8418 | 0.6723 | 0.916326 | 0.890425 | 0.907296 |
| Monkey and mouse | Jango | r2 | 20150828 | 0.8474 | 0.7336 | 0.8151 | 0.71 | 0.845183 | 0.847535 | 0.874436 |
| Monkey and mouse | Jango | r2 | 20150831 | 0.7222 | 0.7283 | 0.6454 | 0.4294 | 0.893816 | 0.879558 | 0.915004 |
| Monkey and mouse | Jango | r2 | 20150905 | 0.6534 | 0.7236 | 0.4412 | 0.5452 | 0.790011 | 0.724782 | 0.818574 |
| Monkey and mouse | Jango | r2 | 20150906 | 0.9295 | 0.6959 | 0.4964 | 0.5772 | 0.814731 | 0.828491 | 0.852938 |
| Monkey and mouse | Jango | r2 | 20150908 | 0.4211 | 0 | 0.5569 | 0.5566 | 0.808109 | 0.783353 | 0.850182 |
| Monkey and mouse | Jango | r2 | 20151029 | 0.7663 | 0.7449 | 0.6096 | 0.5962 | 0.82866 | 0.807986 | 0.837769 |
| Monkey and mouse | Jango | r2 | 20151102 | 0.4355 | 0.3187 | 0.479 | 0.0593 | 0.640357 | 0.637551 | 0.687052 |
| Monkey and mouse | Indy | r2 | 20161011 | 0.6984 | 0.6149 | 0.3877 | 0.5974 | 0.529884 | 0.516586 | 0.625036 |
| Monkey and mouse | Indy | r2 | 20161013 | 0 | 0 | 0 | 0 | -0.185238 | -0.210296 | -0.154079 |
| Monkey and mouse | Indy | r2 | 20161014 | 0.6611 | 0.6726 | 0.4239 | 0.6229 | 0.633951 | 0.595685 | 0.65091 |
| Monkey and mouse | Indy | r2 | 20161017 | 0.5111 | 0.4737 | 0.3027 | 0.2375 | 0.497987 | 0.424276 | 0.53054 |
| Monkey and mouse | Indy | r2 | 20161024 | 0.4625 | 0.3867 | 0.3302 | 0.2489 | 0.453933 | 0.400584 | 0.462357 |
| Monkey and mouse | Indy | r2 | 20161025 | 0.6773 | 0.5507 | 0.4489 | 0.5717 | 0.586082 | 0.532851 | 0.6124 |
| Monkey and mouse | Indy | r2 | 20161026 | 0.5968 | 0.5168 | 0.4717 | 0.4386 | 0.592936 | 0.529456 | 0.601249 |
| Monkey and mouse | Indy | r2 | 20161027 | 0.6053 | 0.488 | 0.2448 | 0.3748 | 0.185281 | 0.038534 | 0.324206 |
| Monkey and mouse | Indy | r2 | 20161206 | 0.4794 | 0.3508 | 0.3741 | 0.2215 | 0.266514 | 0.253039 | 0.304054 |
| Monkey and mouse | Indy | r2 | 20161207 | 0.3785 | 0.4016 | 0.3674 | 0.4072 | 0.324494 | 0.179109 | 0.366047 |
| Monkey and mouse | Indy | r2 | 20161212 | 0.5836 | 0.4948 | 0.2536 | 0.4127 | 0.499829 | 0.451754 | 0.513968 |
| Monkey and mouse | Indy | r2 | 20161220 | 0.3845 | 0.2709 | 0.1026 | 0.2259 | 0.343065 | 0.291085 | 0.341552 |
| Monkey and mouse | Visual Coding | accuracy | session_763673393 | 0.2067 | 0.1467 | 0.0883 | 0.1067 | 0.125 | 0.155 | 0.138333 |
| Monkey and mouse | Visual Coding | accuracy | session_773418906 | 0.1233 | 0.1667 | 0.1133 | 0.1567 | 0.145 | 0.136667 | 0.13 |
| Monkey and mouse | Visual Coding | accuracy | session_791319847 | 0.1217 | 0.1783 | 0.1583 | 0.1367 | 0.138333 | 0.12 | 0.165 |
| Monkey and mouse | Visual Coding | accuracy | session_797828357 | 0.2067 | 0.155 | 0.1083 | 0.1483 | 0.126667 | 0.135 | 0.131667 |

| | | | | | | | | | | |
|---|---|---|---|---|---|---|---|---|---|---|
| Monkey and mouse | Visual Coding | accuracy | session_798911424 | 0.1183 | 0.08 | 0.0683 | 0.1583 | 0.121667 | 0.126667 | 0.133333 |
| Monkey and mouse | Visual Coding | accuracy | session_799864342 | 0.14 | 0.125 | 0.205 | 0.1717 | 0.128333 | 0.101667 | 0.085 |
| Monkey and mouse | Reaching B | accuracy | 20221129 | 0.655 | 0.635 | 0.605 | 0.7 | 0.866667 | 0.85 | 0.823333 |
| Monkey and mouse | Reaching B | accuracy | 20221202 | 0.6563 | 0.6675 | 0.7188 | 0.6925 | 0.885 | 0.88125 | 0.83625 |
| Monkey and mouse | Reaching B | accuracy | 20221205 | 0.69 | 0.73 | 0.685 | 0.685 | 0.93 | 0.905 | 0.91 |
| Monkey and mouse | Reaching B | accuracy | 20221208 | 0.5867 | 0.605 | 0.6933 | 0.6767 | 0.883333 | 0.836667 | 0.876667 |
| Monkey and mouse | Reaching B | accuracy | 20221209 | 0.46 | 0.82 | 0.69 | 0.8025 | 0.85 | 0.8425 | 0.8225 |
| Monkey and mouse | Reaching B | accuracy | 20221213 | 0.5025 | 0.7 | 0.535 | 0.525 | 0.845 | 0.8 | 0.7975 |
| Monkey and mouse | Reaching B | accuracy | 20221214 | 0.455 | 0.66 | 0.605 | 0.555 | 0.8 | 0.705 | 0.815 |
| Monkey and mouse | Reaching B | accuracy | 20221215 | 0.4151 | 0.449 | 0.4841 | 0.4389 | 0.524207 | 0.505843 | 0.524207 |
| Monkey and mouse | Reaching B | accuracy | 20221219 | 0.3786 | 0.7057 | 0.645 | 0.6543 | 0.780714 | 0.755714 | 0.776429 |
| Monkey and mouse | Reaching B | accuracy | 20221230 | 0.419 | 0.346 | 0.568 | 0.664 | 0.794 | 0.773 | 0.769 |
| Single-task decoding setting | Finger PPC | accuracy | 20181015 | 0.1759 | 0.1667 | 0.2269 | 0.5949 | 0.175926 | 0.155093 | 0.1875 |
| Single-task decoding setting | Finger PPC | accuracy | 20181017 | 0.4089 | 0.388 | 0.5469 | 0.4922 | 0.492188 | 0.463542 | 0.369792 |
| Single-task decoding setting | Finger PPC | accuracy | 20181022 | 0.4245 | 0.4115 | 0.4688 | 0.4896 | 0.341146 | 0.382812 | 0.411458 |
| Single-task decoding setting | Reaching B | accuracy | 20221129 | 0.9 | 0.9233 | 0.8 | 0.7617 | 0.856667 | 0.831667 | 0.855 |
| Single-task decoding setting | Reaching B | accuracy | 20221202 | 0.8875 | 0.8188 | 0.8988 | 0.7875 | 0.88 | 0.8925 | 0.8925 |
| Single-task decoding setting | Reaching B | accuracy | 20221205 | 0.89 | 0.82 | 0.97 | 0.865 | 0.91 | 0.88 | 0.875 |
| Single-task decoding setting | Reaching B | accuracy | 20221208 | 0.7617 | 0.8367 | 0.88 | 0.775 | 0.906667 | 0.86 | 0.9 |
| Single-task decoding setting | Reaching B | accuracy | 20221209 | 0.6175 | 0.9075 | 0.935 | 0.7525 | 0.9075 | 0.845 | 0.8725 |
| Single-task decoding setting | Reaching B | accuracy | 20221213 | 0.79 | 0.685 | 0.8525 | 0.51 | 0.84 | 0.785 | 0.8 |
| Single-task decoding setting | Reaching B | accuracy | 20221214 | 0.795 | 0.58 | 0.93 | 0.435 | 0.82 | 0.745 | 0.78 |
| Single-task decoding setting | Reaching B | accuracy | 20221215 | 0.5628 | 0.4494 | 0.6578 | 0.4762 | 0.504174 | 0.519199 | 0.565943 |
| Single-task decoding setting | Reaching B | accuracy | 20221219 | 0.6221 | 0.6621 | 0.9143 | 0.525 | 0.735714 | 0.73 | 0.754286 |
| Single-task decoding setting | Reaching B | accuracy | 20221230 | 0.548 | 0.356 | 0.774 | 0.585 | 0.779 | 0.79 | 0.76 |
| Single-task decoding setting | Indy | r2 | 20161011 | 0.6823 | 0.5761 | 0.4794 | 0.6291 | 0.560088 | 0.545555 | 0.58302 |
| Single-task decoding setting | Indy | r2 | 20161013 | 0.0118 | 0 | 0 | 0 | -0.152527 | -0.121954 | -0.117025 |
| Single-task decoding setting | Indy | r2 | 20161014 | 0.6404 | 0.6637 | 0.3614 | 0.6378 | 0.605696 | 0.631284 | 0.653402 |
| Single-task decoding setting | Indy | r2 | 20161017 | 0.5129 | 0.4827 | 0.2425 | 0.285 | 0.500093 | 0.4282 | 0.512463 |
| Single-task decoding setting | Indy | r2 | 20161024 | 0.5318 | 0.3947 | 0.4627 | 0.2456 | 0.444013 | 0.39744 | 0.471636 |
| Single-task decoding setting | Indy | r2 | 20161025 | 0.6286 | 0.5036 | 0.4623 | 0.5707 | 0.565642 | 0.517391 | 0.601142 |
| Single-task decoding setting | Indy | r2 | 20161026 | 0.589 | 0.5122 | 0.5021 | 0.3989 | 0.589811 | 0.551075 | 0.619455 |
| Single-task decoding setting | Indy | r2 | 20161027 | 0.5295 | 0.462 | 0.2547 | 0.3809 | 0.154347 | 0.121249 | 0.270226 |
| Single-task decoding setting | Indy | r2 | 20161206 | 0.3614 | 0.3573 | 0.4492 | 0.1787 | 0.263285 | 0.240256 | 0.287123 |
| Single-task decoding setting | Indy | r2 | 20161207 | 0.6087 | 0.482 | 0.4333 | 0.4756 | 0.275791 | 0.230516 | 0.258961 |
| Single-task decoding setting | Indy | r2 | 20161212 | 0.6232 | 0.5051 | 0.3073 | 0.4689 | 0.494424 | 0.446692 | 0.504511 |
| Single-task decoding setting | Indy | r2 | 20161220 | 0.329 | 0.2586 | 0.1559 | 0.2015 | 0.322079 | 0.286397 | 0.340927 |
| Single-task decoding setting | Jango | r2 | 20150808 | 0.9334 | 0.8958 | 0.921 | 0.875 | 0.91069 | 0.89777 | 0.898606 |
| Single-task decoding setting | Jango | r2 | 20150809 | 0.8261 | 0.791 | 0.8473 | 0.7978 | 0.897172 | 0.904873 | 0.913471 |
| Single-task decoding setting | Jango | r2 | 20150820 | 0.8348 | 0.9163 | 0.8821 | 0.7238 | 0.916126 | 0.907637 | 0.910559 |
| Single-task decoding setting | Jango | r2 | 20150824 | 0.8908 | 0.9216 | 0.8946 | 0.8562 | 0.89169 | 0.880877 | 0.90245 |
| Single-task decoding setting | Jango | r2 | 20150825 | 0.5765 | 0.6264 | 0.6943 | 0.7147 | 0.779296 | 0.82275 | 0.830212 |
| Single-task decoding setting | Jango | r2 | 20150826 | 0.7855 | 0.8123 | 0.911 | 0.7371 | 0.900093 | 0.904252 | 0.892119 |
| Single-task decoding setting | Jango | r2 | 20150827 | 0.6403 | 0.5513 | 0.8605 | 0.7993 | 0.894004 | 0.895775 | 0.892461 |
| Single-task decoding setting | Jango | r2 | 20150828 | 0.8482 | 0.8163 | 0.8858 | 0.8269 | 0.871076 | 0.868597 | 0.865869 |
| Single-task decoding setting | Jango | r2 | 20150831 | 0.7292 | 0.6482 | 0.4143 | 0.4744 | 0.888422 | 0.89388 | 0.902563 |
| Single-task decoding setting | Jango | r2 | 20150905 | 0.4389 | 0.2943 | 0.7248 | 0.6715 | 0.79258 | 0.754702 | 0.756489 |
| Single-task decoding setting | Jango | r2 | 20150906 | 0.8808 | 0.7363 | 0.837 | 0.6232 | 0.830265 | 0.841515 | 0.841079 |
| Single-task decoding setting | Jango | r2 | 20150908 | 0.4321 | 0.1165 | 0.8065 | 0.1511 | 0.809306 | 0.787433 | 0.820901 |
| Single-task decoding setting | Jango | r2 | 20151029 | 0.7082 | 0.709 | 0.5731 | 0.5951 | 0.823512 | 0.828318 | 0.845069 |
| Single-task decoding setting | Jango | r2 | 20151102 | 0.3073 | 0 | 0.2715 | 0.246 | 0.668204 | 0.673368 | 0.643884 |
| Single-task decoding setting | Visual Coding | accuracy | session_763673393 | 0.13 | 0.22 | 0.1233 | 0.16 | 0.158333 | 0.088333 | 0.171667 |
| Single-task decoding setting | Visual Coding | accuracy | session_773418906 | 0.1317 | 0.1283 | 0.1233 | 0.14 | 0.126667 | 0.086667 | 0.13 |
| Single-task decoding setting | Visual Coding | accuracy | session_791319847 | 0.1517 | 0.1867 | 0.1117 | 0.0983 | 0.14 | 0.128333 | 0.165 |
| Single-task decoding setting | Visual Coding | accuracy | session_797828357 | 0.1733 | 0.2267 | 0.2817 | 0.1633 | 0.135 | 0.138333 | 0.12 |
| Single-task decoding setting | Visual Coding | accuracy | session_798911424 | 0.1317 | 0.1517 | 0.18 | 0.1967 | 0.148333 | 0.123333 | 0.155 |
| Single-task decoding setting | Visual Coding | accuracy | session_799864342 | 0.09 | 0.0467 | 0.1267 | 0.23 | 0.151667 | 0.093333 | 0.166667 |
| Species across human, monkey and mo... | Finger PPC | accuracy | 20181015 | 0.2222 | 0.1713 | 0.1991 | 0.4074 | 0.127315 | 0.212963 | 0.127315 |
| Species across human, monkey and mo... | Finger PPC | accuracy | 20181017 | 0.3542 | 0.3776 | 0.3568 | 0.2135 | 0.356771 | 0.390625 | 0.401042 |
| Species across human, monkey and mo... | Finger PPC | accuracy | 20181022 | 0.2813 | 0.3828 | 0.3438 | 0.4115 | 0.432292 | 0.450521 | 0.40625 |
| Species across human, monkey and mo... | Jango | r2 | 20150808 | 0.9335 | 0.9055 | 0.8961 | 0.8353 | 0.906392 | 0.89164 | 0.915729 |
| Species across human, monkey and mo... | Jango | r2 | 20150809 | 0.8969 | 0.7769 | 0.8814 | 0.7977 | 0.91029 | 0.902336 | 0.914753 |
| Species across human, monkey and mo... | Jango | r2 | 20150820 | 0.9035 | 0.7102 | 0.8888 | 0.6415 | 0.905057 | 0.912309 | 0.911302 |

| | | | | | | | | | | |
|---|---|---|---|---|---|---|---|---|---|---|
| Species across human, monkey and mo... | Jango | r2 | 20150824 | 0.9088 | 0.9025 | 0.6773 | 0.7986 | 0.902353 | 0.905555 | 0.907806 |
| Species across human, monkey and mo... | Jango | r2 | 20150825 | 0.801 | 0.6587 | 0.5755 | 0.6799 | 0.787076 | 0.792428 | 0.807786 |
| Species across human, monkey and mo... | Jango | r2 | 20150826 | 0.8081 | 0.69 | 0.6156 | 0.7176 | 0.891449 | 0.908266 | 0.892396 |
| Species across human, monkey and mo... | Jango | r2 | 20150827 | 0.8602 | 0.6476 | 0.7623 | 0.7534 | 0.894364 | 0.910912 | 0.890596 |
| Species across human, monkey and mo... | Jango | r2 | 20150828 | 0.8764 | 0.7062 | 0.7512 | 0.7712 | 0.866577 | 0.874824 | 0.857773 |
| Species across human, monkey and mo... | Jango | r2 | 20150831 | 0.7447 | 0.71 | 0.6848 | 0.4423 | 0.894083 | 0.911071 | 0.906735 |
| Species across human, monkey and mo... | Jango | r2 | 20150905 | 0.7831 | 0.6739 | 0.5407 | 0.573 | 0.763145 | 0.799089 | 0.784071 |
| Species across human, monkey and mo... | Jango | r2 | 20150906 | 0.8992 | 0.7266 | 0.7541 | 0.6278 | 0.800402 | 0.835217 | 0.8148 |
| Species across human, monkey and mo... | Jango | r2 | 20150908 | 0.1528 | 0 | 0.6659 | 0.5893 | 0.826988 | 0.847289 | 0.828843 |
| Species across human, monkey and mo... | Jango | r2 | 20151029 | 0.7252 | 0.7393 | 0.4945 | 0.5974 | 0.834103 | 0.824387 | 0.834767 |
| Species across human, monkey and mo... | Jango | r2 | 20151102 | 0.3411 | 0.2088 | 0.2009 | 0.1532 | 0.662588 | 0.64215 | 0.66404 |
| Species across human, monkey and mo... | Indy | r2 | 20161011 | 0.7262 | 0.6282 | 0.3948 | 0.6058 | 0.43477 | 0.536731 | 0.646841 |
| Species across human, monkey and mo... | Indy | r2 | 20161013 | 0 | 0 | 0 | 0 | -0.271914 | -0.156761 | -0.164211 |
| Species across human, monkey and mo... | Indy | r2 | 20161014 | 0.6926 | 0.6864 | 0.4436 | 0.594 | 0.554757 | 0.617038 | 0.651077 |
| Species across human, monkey and mo... | Indy | r2 | 20161017 | 0.5163 | 0.5074 | 0.2796 | 0.3179 | 0.474198 | 0.48263 | 0.55189 |
| Species across human, monkey and mo... | Indy | r2 | 20161024 | 0.5389 | 0.413 | 0.4216 | 0.2584 | 0.394947 | 0.440619 | 0.468229 |
| Species across human, monkey and mo... | Indy | r2 | 20161025 | 0.6628 | 0.5602 | 0.4883 | 0.4885 | 0.543289 | 0.550621 | 0.632893 |
| Species across human, monkey and mo... | Indy | r2 | 20161026 | 0.6064 | 0.5301 | 0.5148 | 0.4768 | 0.53377 | 0.578065 | 0.633621 |
| Species across human, monkey and mo... | Indy | r2 | 20161027 | 0.5397 | 0.502 | 0.237 | 0.3852 | -0.023519 | 0.176486 | 0.323801 |
| Species across human, monkey and mo... | Indy | r2 | 20161206 | 0.4983 | 0.2759 | 0.3971 | 0.2314 | 0.241091 | 0.277378 | 0.338942 |
| Species across human, monkey and mo... | Indy | r2 | 20161207 | 0.4093 | 0.3725 | 0.4106 | 0.3535 | 0.314807 | 0.245438 | 0.328246 |
| Species across human, monkey and mo... | Indy | r2 | 20161212 | 0.5866 | 0.5267 | 0.2817 | 0.4381 | 0.439953 | 0.474883 | 0.513926 |
| Species across human, monkey and mo... | Indy | r2 | 20161220 | 0.4389 | 0.2472 | 0.1441 | 0.2049 | 0.290734 | 0.327102 | 0.39791 |
| Species across human, monkey and mo... | Visual Coding | accuracy | session_763673393 | 0.12 | 0.1317 | 0.12 | 0.135 | 0.125 | 0.113333 | 0.151667 |
| Species across human, monkey and mo... | Visual Coding | accuracy | session_773418906 | 0.105 | 0.1433 | 0.1533 | 0.0733 | 0.155 | 0.13 | 0.111667 |
| Species across human, monkey and mo... | Visual Coding | accuracy | session_791319847 | 0.1667 | 0.2083 | 0.1717 | 0.1617 | 0.118333 | 0.105 | 0.146667 |
| Species across human, monkey and mo... | Visual Coding | accuracy | session_797828357 | 0.2033 | 0.1917 | 0.1967 | 0.0933 | 0.121667 | 0.135 | 0.131667 |
| Species across human, monkey and mo... | Visual Coding | accuracy | session_798911424 | 0.135 | 0.1017 | 0.1467 | 0.1317 | 0.15 | 0.123333 | 0.125 |
| Species across human, monkey and mo... | Visual Coding | accuracy | session_799864342 | 0.155 | 0.13 | 0.0417 | 0.155 | 0.116667 | 0.123333 | 0.125 |
| Species across human, monkey and mo... | Reaching B | accuracy | 20221129 | 0.5883 | 0.675 | 0.5083 | 0.6067 | 0.841667 | 0.85 | 0.785 |
| Species across human, monkey and mo... | Reaching B | accuracy | 20221202 | 0.6838 | 0.61 | 0.7013 | 0.69 | 0.88125 | 0.84625 | 0.8825 |
| Species across human, monkey and mo... | Reaching B | accuracy | 20221205 | 0.64 | 0.75 | 0.605 | 0.51 | 0.86 | 0.875 | 0.86 |
| Species across human, monkey and mo... | Reaching B | accuracy | 20221208 | 0.5533 | 0.7383 | 0.53 | 0.61 | 0.813333 | 0.866667 | 0.868333 |
| Species across human, monkey and mo... | Reaching B | accuracy | 20221209 | 0.3375 | 0.79 | 0.4425 | 0.705 | 0.8225 | 0.86 | 0.8725 |
| Species across human, monkey and mo... | Reaching B | accuracy | 20221213 | 0.4675 | 0.7425 | 0.57 | 0.52 | 0.845 | 0.8025 | 0.7975 |
| Species across human, monkey and mo... | Reaching B | accuracy | 20221214 | 0.56 | 0.715 | 0.55 | 0.555 | 0.84 | 0.78 | 0.75 |
| Species across human, monkey and mo... | Reaching B | accuracy | 20221215 | 0.4793 | 0.5224 | 0.5309 | 0.5778 | 0.567613 | 0.509182 | 0.530885 |
| Species across human, monkey and mo... | Reaching B | accuracy | 20221219 | 0.4143 | 0.7021 | 0.8021 | 0.705 | 0.846429 | 0.734286 | 0.749286 |
| Species across human, monkey and mo... | Reaching B | accuracy | 20221230 | 0.524 | 0.451 | 0.576 | 0.57 | 0.801 | 0.748 | 0.752 |
