## Supplementary File 3 for "BCIJelly: An integrated ecosystem for brain–computer interface research"

| Task type | Algorithm | Dataset | Protocol | Training day | Test day | Seed | Metric | Score |
|---|---|---|---|---|---|---|---|---|
| classification | lstm | Finger PPC | single_day | 20180910 | 20180910 | 31 | accuracy | 0.844156 |
| classification | lstm | Finger PPC | single_day | 20180910 | 20180910 | 44 | accuracy | 0.909091 |
| classification | lstm | Finger PPC | single_day | 20180910 | 20180910 | 44 | accuracy | 0.961039 |
| classification | lstm | Finger PPC | single_day | 20180910 | 20180910 | 46 | accuracy | 0.909091 |
| classification | lstm | Finger PPC | single_day | 20180910 | 20180910 | 46 | accuracy | 0.948052 |
| classification | lstm | Finger PPC | single_day | 20180910 | 20180910 | 54 | accuracy | 0.961039 |
| classification | lstm | Finger PPC | single_day | 20180910 | 20180910 | 54 | accuracy | 0.974026 |
| classification | lstm | Finger PPC | single_day | 20180910 | 20180910 | 69 | accuracy | 0.896104 |
| classification | lstm | Finger PPC | single_day | 20180917a | 20180917a | 31 | accuracy | 0.922078 |
| classification | lstm | Finger PPC | single_day | 20180917a | 20180917a | 44 | accuracy | 0.922078 |
| classification | lstm | Finger PPC | single_day | 20180917a | 20180917a | 44 | accuracy | 0.961039 |
| classification | lstm | Finger PPC | single_day | 20180917a | 20180917a | 46 | accuracy | 0.857143 |
| classification | lstm | Finger PPC | single_day | 20180917a | 20180917a | 46 | accuracy | 0.922078 |
| classification | lstm | Finger PPC | single_day | 20180917a | 20180917a | 54 | accuracy | 0.87013 |
| classification | lstm | Finger PPC | single_day | 20180917a | 20180917a | 54 | accuracy | 0.909091 |
| classification | lstm | Finger PPC | single_day | 20180917a | 20180917a | 69 | accuracy | 0.792208 |
| classification | lstm | Finger PPC | single_day | 20180917b | 20180917b | 31 | accuracy | 0.867647 |
| classification | lstm | Finger PPC | single_day | 20180917b | 20180917b | 44 | accuracy | 0.941176 |
| classification | lstm | Finger PPC | single_day | 20180917b | 20180917b | 44 | accuracy | 0.955882 |
| classification | lstm | Finger PPC | single_day | 20180917b | 20180917b | 46 | accuracy | 0.941176 |
| classification | lstm | Finger PPC | single_day | 20180917b | 20180917b | 46 | accuracy | 0.955882 |
| classification | lstm | Finger PPC | single_day | 20180917b | 20180917b | 54 | accuracy | 0.794118 |
| classification | lstm | Finger PPC | single_day | 20180917b | 20180917b | 54 | accuracy | 0.911765 |
| classification | lstm | Finger PPC | single_day | 20180917b | 20180917b | 69 | accuracy | 0.911765 |
| classification | lstm | Finger PPC | single_day | 20180924 | 20180924 | 31 | accuracy | 0.895833 |
| classification | lstm | Finger PPC | single_day | 20180924 | 20180924 | 44 | accuracy | 0.958333 |
| classification | lstm | Finger PPC | single_day | 20180924 | 20180924 | 44 | accuracy | 0.947917 |
| classification | lstm | Finger PPC | single_day | 20180924 | 20180924 | 46 | accuracy | 0.979167 |
| classification | lstm | Finger PPC | single_day | 20180924 | 20180924 | 46 | accuracy | 0.947917 |
| classification | lstm | Finger PPC | single_day | 20180924 | 20180924 | 54 | accuracy | 0.927083 |
| classification | lstm | Finger PPC | single_day | 20180924 | 20180924 | 54 | accuracy | 0.9375 |
| classification | lstm | Finger PPC | single_day | 20180924 | 20180924 | 69 | accuracy | 0.979167 |
| classification | lstm | Finger PPC | single_day | 20180926 | 20180926 | 31 | accuracy | 0.873563 |
| classification | lstm | Finger PPC | single_day | 20180926 | 20180926 | 44 | accuracy | 0.931034 |
| classification | lstm | Finger PPC | single_day | 20180926 | 20180926 | 44 | accuracy | 0.896552 |
| classification | lstm | Finger PPC | single_day | 20180926 | 20180926 | 46 | accuracy | 0.931034 |
| classification | lstm | Finger PPC | single_day | 20180926 | 20180926 | 54 | accuracy | 0.931034 |
| classification | lstm | Finger PPC | single_day | 20180926 | 20180926 | 54 | accuracy | 0.977011 |
| classification | lstm | Finger PPC | single_day | 20180926 | 20180926 | 69 | accuracy | 0.885057 |
| classification | lstm | Finger PPC | single_day | 20181001 | 20181001 | 31 | accuracy | 0.931034 |
| classification | lstm | Finger PPC | single_day | 20181001 | 20181001 | 44 | accuracy | 0.896552 |
| classification | lstm | Finger PPC | single_day | 20181001 | 20181001 | 44 | accuracy | 0.91954 |
| classification | lstm | Finger PPC | single_day | 20181001 | 20181001 | 46 | accuracy | 0.942529 |
| classification | lstm | Finger PPC | single_day | 20181001 | 20181001 | 46 | accuracy | 0.977011 |
| classification | lstm | Finger PPC | single_day | 20181001 | 20181001 | 54 | accuracy | 0.91954 |
| classification | lstm | Finger PPC | single_day | 20181001 | 20181001 | 54 | accuracy | 0.965517 |
| classification | lstm | Finger PPC | single_day | 20181001 | 20181001 | 69 | accuracy | 0.91954 |
| classification | lstm | Finger PPC | single_day | 20181012 | 20181012 | 31 | accuracy | 0.83908 |
| classification | lstm | Finger PPC | single_day | 20181012 | 20181012 | 44 | accuracy | 0.862069 |
| classification | lstm | Finger PPC | single_day | 20181012 | 20181012 | 44 | accuracy | 0.91954 |
| classification | lstm | Finger PPC | single_day | 20181012 | 20181012 | 46 | accuracy | 0.873563 |
| classification | lstm | Finger PPC | single_day | 20181012 | 20181012 | 46 | accuracy | 0.977011 |
| classification | lstm | Finger PPC | single_day | 20181012 | 20181012 | 54 | accuracy | 0.942529 |
| classification | lstm | Finger PPC | single_day | 20181012 | 20181012 | 54 | accuracy | 0.977011 |
| classification | lstm | Finger PPC | single_day | 20181012 | 20181012 | 69 | accuracy | 0.931034 |
| classification | lstm | Finger PPC | single_day | 20181015 | 20181015 | 31 | accuracy | 0.896552 |
| classification | lstm | Finger PPC | single_day | 20181015 | 20181015 | 44 | accuracy | 0.954023 |
| classification | lstm | Finger PPC | single_day | 20181015 | 20181015 | 44 | accuracy | 0.942529 |
| classification | lstm | Finger PPC | single_day | 20181015 | 20181015 | 46 | accuracy | 0.931034 |
| classification | lstm | Finger PPC | single_day | 20181015 | 20181015 | 46 | accuracy | 0.954023 |

| | | | | | | | | |
|---|---|---|---|---|---|---|---|---|
| classification | lstm | Finger PPC | single_day | 20181015 | 20181015 | 54 | accuracy | 0.942529 |
| classification | lstm | Finger PPC | single_day | 20181015 | 20181015 | 54 | accuracy | 0.954023 |
| classification | lstm | Finger PPC | single_day | 20181015 | 20181015 | 69 | accuracy | 0.942529 |
| classification | lstm | Finger PPC | single_day | 20181017 | 20181017 | 31 | accuracy | 0.896104 |
| classification | lstm | Finger PPC | single_day | 20181017 | 20181017 | 44 | accuracy | 0.87013 |
| classification | lstm | Finger PPC | single_day | 20181017 | 20181017 | 44 | accuracy | 0.948052 |
| classification | lstm | Finger PPC | single_day | 20181017 | 20181017 | 46 | accuracy | 0.779221 |
| classification | lstm | Finger PPC | single_day | 20181017 | 20181017 | 46 | accuracy | 0.844156 |
| classification | lstm | Finger PPC | single_day | 20181017 | 20181017 | 54 | accuracy | 0.896104 |
| classification | lstm | Finger PPC | single_day | 20181017 | 20181017 | 54 | accuracy | 0.935065 |
| classification | lstm | Finger PPC | single_day | 20181017 | 20181017 | 69 | accuracy | 0.922078 |
| classification | lstm | Finger PPC | single_day | 20181022 | 20181022 | 31 | accuracy | 0.831169 |
| classification | lstm | Finger PPC | single_day | 20181022 | 20181022 | 44 | accuracy | 0.766234 |
| classification | lstm | Finger PPC | single_day | 20181022 | 20181022 | 44 | accuracy | 0.909091 |
| classification | lstm | Finger PPC | single_day | 20181022 | 20181022 | 46 | accuracy | 0.87013 |
| classification | lstm | Finger PPC | single_day | 20181022 | 20181022 | 46 | accuracy | 0.935065 |
| classification | lstm | Finger PPC | single_day | 20181022 | 20181022 | 54 | accuracy | 0.87013 |
| classification | lstm | Finger PPC | single_day | 20181022 | 20181022 | 54 | accuracy | 0.922078 |
| classification | lstm | Finger PPC | single_day | 20181022 | 20181022 | 69 | accuracy | 0.766234 |
| classification | transformer | Finger PPC | single_day | 20180910 | 20180910 | 31 | accuracy | 0.883117 |
| classification | transformer | Finger PPC | single_day | 20180910 | 20180910 | 44 | accuracy | 0.909091 |
| classification | transformer | Finger PPC | single_day | 20180910 | 20180910 | 44 | accuracy | 0.935065 |
| classification | transformer | Finger PPC | single_day | 20180910 | 20180910 | 46 | accuracy | 0.87013 |
| classification | transformer | Finger PPC | single_day | 20180910 | 20180910 | 46 | accuracy | 0.883117 |
| classification | transformer | Finger PPC | single_day | 20180910 | 20180910 | 54 | accuracy | 0.935065 |
| classification | transformer | Finger PPC | single_day | 20180910 | 20180910 | 54 | accuracy | 0.948052 |
| classification | transformer | Finger PPC | single_day | 20180910 | 20180910 | 69 | accuracy | 0.87013 |
| classification | transformer | Finger PPC | single_day | 20180917a | 20180917a | 31 | accuracy | 0.961039 |
| classification | transformer | Finger PPC | single_day | 20180917a | 20180917a | 44 | accuracy | 0.909091 |
| classification | transformer | Finger PPC | single_day | 20180917a | 20180917a | 46 | accuracy | 0.883117 |
| classification | transformer | Finger PPC | single_day | 20180917a | 20180917a | 46 | accuracy | 0.87013 |
| classification | transformer | Finger PPC | single_day | 20180917a | 20180917a | 54 | accuracy | 0.948052 |
| classification | transformer | Finger PPC | single_day | 20180917a | 20180917a | 54 | accuracy | 0.909091 |
| classification | transformer | Finger PPC | single_day | 20180917a | 20180917a | 69 | accuracy | 0.87013 |
| classification | transformer | Finger PPC | single_day | 20180917b | 20180917b | 31 | accuracy | 0.867647 |
| classification | transformer | Finger PPC | single_day | 20180917b | 20180917b | 44 | accuracy | 0.926471 |
| classification | transformer | Finger PPC | single_day | 20180917b | 20180917b | 44 | accuracy | 0.867647 |
| classification | transformer | Finger PPC | single_day | 20180917b | 20180917b | 46 | accuracy | 0.911765 |
| classification | transformer | Finger PPC | single_day | 20180917b | 20180917b | 46 | accuracy | 0.897059 |
| classification | transformer | Finger PPC | single_day | 20180917b | 20180917b | 54 | accuracy | 0.926471 |
| classification | transformer | Finger PPC | single_day | 20180917b | 20180917b | 54 | accuracy | 0.882353 |
| classification | transformer | Finger PPC | single_day | 20180917b | 20180917b | 69 | accuracy | 0.838235 |
| classification | transformer | Finger PPC | single_day | 20180924 | 20180924 | 31 | accuracy | 0.90625 |
| classification | transformer | Finger PPC | single_day | 20180924 | 20180924 | 44 | accuracy | 0.895833 |
| classification | transformer | Finger PPC | single_day | 20180924 | 20180924 | 44 | accuracy | 0.875 |
| classification | transformer | Finger PPC | single_day | 20180924 | 20180924 | 46 | accuracy | 0.947917 |
| classification | transformer | Finger PPC | single_day | 20180924 | 20180924 | 46 | accuracy | 0.833333 |
| classification | transformer | Finger PPC | single_day | 20180924 | 20180924 | 54 | accuracy | 0.916667 |
| classification | transformer | Finger PPC | single_day | 20180924 | 20180924 | 54 | accuracy | 0.864583 |
| classification | transformer | Finger PPC | single_day | 20180924 | 20180924 | 69 | accuracy | 0.96875 |
| classification | transformer | Finger PPC | single_day | 20180926 | 20180926 | 31 | accuracy | 0.896552 |
| classification | transformer | Finger PPC | single_day | 20180926 | 20180926 | 44 | accuracy | 0.896552 |
| classification | transformer | Finger PPC | single_day | 20180926 | 20180926 | 44 | accuracy | 0.83908 |
| classification | transformer | Finger PPC | single_day | 20180926 | 20180926 | 46 | accuracy | 0.931034 |
| classification | transformer | Finger PPC | single_day | 20180926 | 20180926 | 46 | accuracy | 0.91954 |
| classification | transformer | Finger PPC | single_day | 20180926 | 20180926 | 54 | accuracy | 0.91954 |
| classification | transformer | Finger PPC | single_day | 20180926 | 20180926 | 54 | accuracy | 0.908046 |
| classification | transformer | Finger PPC | single_day | 20180926 | 20180926 | 69 | accuracy | 0.931034 |
| classification | transformer | Finger PPC | single_day | 20181001 | 20181001 | 31 | accuracy | 0.908046 |
| classification | transformer | Finger PPC | single_day | 20181001 | 20181001 | 44 | accuracy | 0.931034 |
| classification | transformer | Finger PPC | single_day | 20181001 | 20181001 | 44 | accuracy | 0.83908 |
| classification | transformer | Finger PPC | single_day | 20181001 | 20181001 | 46 | accuracy | 0.954023 |
| classification | transformer | Finger PPC | single_day | 20181001 | 20181001 | 46 | accuracy | 0.965517 |

| | | | | | | | | |
|---|---|---|---|---|---|---|---|---|
| classification | transformer | Finger PPC | single_day | 20181001 | 20181001 | 54 | accuracy | 0.931034 |
| classification | transformer | Finger PPC | single_day | 20181001 | 20181001 | 54 | accuracy | 0.91954 |
| classification | transformer | Finger PPC | single_day | 20181001 | 20181001 | 69 | accuracy | 0.885057 |
| classification | transformer | Finger PPC | single_day | 20181012 | 20181012 | 31 | accuracy | 0.965517 |
| classification | transformer | Finger PPC | single_day | 20181012 | 20181012 | 44 | accuracy | 0.931034 |
| classification | transformer | Finger PPC | single_day | 20181012 | 20181012 | 44 | accuracy | 0.83908 |
| classification | transformer | Finger PPC | single_day | 20181012 | 20181012 | 46 | accuracy | 0.908046 |
| classification | transformer | Finger PPC | single_day | 20181012 | 20181012 | 46 | accuracy | 0.873563 |
| classification | transformer | Finger PPC | single_day | 20181012 | 20181012 | 54 | accuracy | 0.965517 |
| classification | transformer | Finger PPC | single_day | 20181012 | 20181012 | 54 | accuracy | 0.942529 |
| classification | transformer | Finger PPC | single_day | 20181012 | 20181012 | 69 | accuracy | 0.954023 |
| classification | transformer | Finger PPC | single_day | 20181015 | 20181015 | 31 | accuracy | 0.91954 |
| classification | transformer | Finger PPC | single_day | 20181015 | 20181015 | 44 | accuracy | 0.977011 |
| classification | transformer | Finger PPC | single_day | 20181015 | 20181015 | 44 | accuracy | 0.931034 |
| classification | transformer | Finger PPC | single_day | 20181015 | 20181015 | 46 | accuracy | 0.885057 |
| classification | transformer | Finger PPC | single_day | 20181015 | 20181015 | 46 | accuracy | 0.873563 |
| classification | transformer | Finger PPC | single_day | 20181015 | 20181015 | 54 | accuracy | 0.965517 |
| classification | transformer | Finger PPC | single_day | 20181015 | 20181015 | 54 | accuracy | 0.885057 |
| classification | transformer | Finger PPC | single_day | 20181015 | 20181015 | 69 | accuracy | 0.931034 |
| classification | transformer | Finger PPC | single_day | 20181017 | 20181017 | 31 | accuracy | 0.857143 |
| classification | transformer | Finger PPC | single_day | 20181017 | 20181017 | 44 | accuracy | 0.909091 |
| classification | transformer | Finger PPC | single_day | 20181017 | 20181017 | 44 | accuracy | 0.922078 |
| classification | transformer | Finger PPC | single_day | 20181017 | 20181017 | 46 | accuracy | 0.87013 |
| classification | transformer | Finger PPC | single_day | 20181017 | 20181017 | 46 | accuracy | 0.805195 |
| classification | transformer | Finger PPC | single_day | 20181017 | 20181017 | 54 | accuracy | 0.909091 |
| classification | transformer | Finger PPC | single_day | 20181017 | 20181017 | 54 | accuracy | 0.896104 |
| classification | transformer | Finger PPC | single_day | 20181017 | 20181017 | 69 | accuracy | 0.831169 |
| classification | transformer | Finger PPC | single_day | 20181022 | 20181022 | 31 | accuracy | 0.896104 |
| classification | transformer | Finger PPC | single_day | 20181022 | 20181022 | 44 | accuracy | 0.857143 |
| classification | transformer | Finger PPC | single_day | 20181022 | 20181022 | 46 | accuracy | 0.87013 |
| classification | transformer | Finger PPC | single_day | 20181022 | 20181022 | 46 | accuracy | 0.857143 |
| classification | transformer | Finger PPC | single_day | 20181022 | 20181022 | 54 | accuracy | 0.857143 |
| classification | transformer | Finger PPC | single_day | 20181022 | 20181022 | 54 | accuracy | 0.87013 |
| classification | transformer | Finger PPC | single_day | 20181022 | 20181022 | 69 | accuracy | 0.844156 |
| classification | lfads | Finger PPC | single_day | 20180910 | 20180910 | 44 | accuracy | 0.948052 |
| classification | lfads | Finger PPC | single_day | 20180910 | 20180910 | 46 | accuracy | 0.922078 |
| classification | lfads | Finger PPC | single_day | 20180910 | 20180910 | 54 | accuracy | 0.948052 |
| classification | lfads | Finger PPC | single_day | 20180917a | 20180917a | 44 | accuracy | 0.961039 |
| classification | lfads | Finger PPC | single_day | 20180917a | 20180917a | 46 | accuracy | 0.883117 |
| classification | lfads | Finger PPC | single_day | 20180917a | 20180917a | 54 | accuracy | 0.87013 |
| classification | lfads | Finger PPC | single_day | 20180917b | 20180917b | 44 | accuracy | 0.838235 |
| classification | lfads | Finger PPC | single_day | 20180917b | 20180917b | 46 | accuracy | 0.882353 |
| classification | lfads | Finger PPC | single_day | 20180917b | 20180917b | 54 | accuracy | 0.852941 |
| classification | lfads | Finger PPC | single_day | 20180924 | 20180924 | 44 | accuracy | 0.96875 |
| classification | lfads | Finger PPC | single_day | 20180924 | 20180924 | 46 | accuracy | 0.916667 |
| classification | lfads | Finger PPC | single_day | 20180924 | 20180924 | 54 | accuracy | 0.885417 |
| classification | lfads | Finger PPC | single_day | 20180926 | 20180926 | 44 | accuracy | 0.83908 |
| classification | lfads | Finger PPC | single_day | 20180926 | 20180926 | 46 | accuracy | 0.908046 |
| classification | lfads | Finger PPC | single_day | 20180926 | 20180926 | 54 | accuracy | 0.91954 |
| classification | lfads | Finger PPC | single_day | 20181001 | 20181001 | 44 | accuracy | 0.873563 |
| classification | lfads | Finger PPC | single_day | 20181001 | 20181001 | 46 | accuracy | 0.954023 |
| classification | lfads | Finger PPC | single_day | 20181001 | 20181001 | 54 | accuracy | 0.850575 |
| classification | lfads | Finger PPC | single_day | 20181012 | 20181012 | 44 | accuracy | 0.91954 |
| classification | lfads | Finger PPC | single_day | 20181012 | 20181012 | 46 | accuracy | 0.896552 |
| classification | lfads | Finger PPC | single_day | 20181012 | 20181012 | 54 | accuracy | 0.908046 |
| classification | lfads | Finger PPC | single_day | 20181015 | 20181015 | 44 | accuracy | 0.885057 |
| classification | lfads | Finger PPC | single_day | 20181015 | 20181015 | 46 | accuracy | 0.931034 |
| classification | lfads | Finger PPC | single_day | 20181015 | 20181015 | 54 | accuracy | 0.896552 |
| classification | lfads | Finger PPC | single_day | 20181017 | 20181017 | 44 | accuracy | 0.792208 |
| classification | lfads | Finger PPC | single_day | 20181017 | 20181017 | 46 | accuracy | 0.753247 |
| classification | lfads | Finger PPC | single_day | 20181017 | 20181017 | 54 | accuracy | 0.753247 |
| classification | lfads | Finger PPC | single_day | 20181022 | 20181022 | 44 | accuracy | 0.857143 |
| classification | lfads | Finger PPC | single_day | 20181022 | 20181022 | 46 | accuracy | 0.753247 |

| | | | | | | | | |
|---|---|---|---|---|---|---|---|---|
| classification | lfads | Finger PPC | single_day | 20181022 | 20181022 | 54 | accuracy | 0.844156 |
| classification | cycle_gan | Finger PPC | single_day | 20180910 | 20180910 | 31 | accuracy | 0.87013 |
| classification | cycle_gan | Finger PPC | single_day | 20180910 | 20180910 | 44 | accuracy | 0.87013 |
| classification | cycle_gan | Finger PPC | single_day | 20180910 | 20180910 | 46 | accuracy | 0.896104 |
| classification | cycle_gan | Finger PPC | single_day | 20180910 | 20180910 | 54 | accuracy | 0.922078 |
| classification | cycle_gan | Finger PPC | single_day | 20180910 | 20180910 | 69 | accuracy | 0.948052 |
| classification | cycle_gan | Finger PPC | single_day | 20180917a | 20180917a | 31 | accuracy | 0.935065 |
| classification | cycle_gan | Finger PPC | single_day | 20180917a | 20180917a | 44 | accuracy | 0.909091 |
| classification | cycle_gan | Finger PPC | single_day | 20180917a | 20180917a | 46 | accuracy | 0.792208 |
| classification | cycle_gan | Finger PPC | single_day | 20180917a | 20180917a | 54 | accuracy | 0.857143 |
| classification | cycle_gan | Finger PPC | single_day | 20180917a | 20180917a | 69 | accuracy | 0.818182 |
| classification | cycle_gan | Finger PPC | single_day | 20180917b | 20180917b | 31 | accuracy | 0.838235 |
| classification | cycle_gan | Finger PPC | single_day | 20180917b | 20180917b | 44 | accuracy | 0.867647 |
| classification | cycle_gan | Finger PPC | single_day | 20180917b | 20180917b | 46 | accuracy | 0.808824 |
| classification | cycle_gan | Finger PPC | single_day | 20180917b | 20180917b | 54 | accuracy | 0.808824 |
| classification | cycle_gan | Finger PPC | single_day | 20180917b | 20180917b | 69 | accuracy | 0.823529 |
| classification | cycle_gan | Finger PPC | single_day | 20180924 | 20180924 | 31 | accuracy | 0.958333 |
| classification | cycle_gan | Finger PPC | single_day | 20180924 | 20180924 | 44 | accuracy | 0.875 |
| classification | cycle_gan | Finger PPC | single_day | 20180924 | 20180924 | 46 | accuracy | 0.864583 |
| classification | cycle_gan | Finger PPC | single_day | 20180924 | 20180924 | 54 | accuracy | 0.916667 |
| classification | cycle_gan | Finger PPC | single_day | 20180924 | 20180924 | 69 | accuracy | 0.864583 |
| classification | cycle_gan | Finger PPC | single_day | 20180926 | 20180926 | 31 | accuracy | 0.850575 |
| classification | cycle_gan | Finger PPC | single_day | 20180926 | 20180926 | 44 | accuracy | 0.83908 |
| classification | cycle_gan | Finger PPC | single_day | 20180926 | 20180926 | 46 | accuracy | 0.896552 |
| classification | cycle_gan | Finger PPC | single_day | 20180926 | 20180926 | 54 | accuracy | 0.885057 |
| classification | cycle_gan | Finger PPC | single_day | 20180926 | 20180926 | 69 | accuracy | 0.908046 |
| classification | cycle_gan | Finger PPC | single_day | 20181001 | 20181001 | 31 | accuracy | 0.850575 |
| classification | cycle_gan | Finger PPC | single_day | 20181001 | 20181001 | 44 | accuracy | 0.862069 |
| classification | cycle_gan | Finger PPC | single_day | 20181001 | 20181001 | 46 | accuracy | 0.862069 |
| classification | cycle_gan | Finger PPC | single_day | 20181001 | 20181001 | 54 | accuracy | 0.885057 |
| classification | cycle_gan | Finger PPC | single_day | 20181001 | 20181001 | 69 | accuracy | 0.908046 |
| classification | cycle_gan | Finger PPC | single_day | 20181012 | 20181012 | 31 | accuracy | 0.862069 |
| classification | cycle_gan | Finger PPC | single_day | 20181012 | 20181012 | 44 | accuracy | 0.91954 |
| classification | cycle_gan | Finger PPC | single_day | 20181012 | 20181012 | 46 | accuracy | 0.896552 |
| classification | cycle_gan | Finger PPC | single_day | 20181012 | 20181012 | 54 | accuracy | 0.931034 |
| classification | cycle_gan | Finger PPC | single_day | 20181012 | 20181012 | 69 | accuracy | 0.896552 |
| classification | cycle_gan | Finger PPC | single_day | 20181015 | 20181015 | 31 | accuracy | 0.91954 |
| classification | cycle_gan | Finger PPC | single_day | 20181015 | 20181015 | 44 | accuracy | 0.850575 |
| classification | cycle_gan | Finger PPC | single_day | 20181015 | 20181015 | 46 | accuracy | 0.862069 |
| classification | cycle_gan | Finger PPC | single_day | 20181015 | 20181015 | 54 | accuracy | 0.862069 |
| classification | cycle_gan | Finger PPC | single_day | 20181015 | 20181015 | 69 | accuracy | 0.896552 |
| classification | cycle_gan | Finger PPC | single_day | 20181017 | 20181017 | 31 | accuracy | 0.792208 |
| classification | cycle_gan | Finger PPC | single_day | 20181017 | 20181017 | 44 | accuracy | 0.844156 |
| classification | cycle_gan | Finger PPC | single_day | 20181017 | 20181017 | 46 | accuracy | 0.766234 |
| classification | cycle_gan | Finger PPC | single_day | 20181017 | 20181017 | 54 | accuracy | 0.857143 |
| classification | cycle_gan | Finger PPC | single_day | 20181017 | 20181017 | 69 | accuracy | 0.831169 |
| classification | cycle_gan | Finger PPC | single_day | 20181022 | 20181022 | 31 | accuracy | 0.831169 |
| classification | cycle_gan | Finger PPC | single_day | 20181022 | 20181022 | 44 | accuracy | 0.805195 |
| classification | cycle_gan | Finger PPC | single_day | 20181022 | 20181022 | 46 | accuracy | 0.792208 |
| classification | cycle_gan | Finger PPC | single_day | 20181022 | 20181022 | 54 | accuracy | 0.779221 |
| classification | cycle_gan | Finger PPC | single_day | 20181022 | 20181022 | 69 | accuracy | 0.701299 |
| classification | stabilization | Finger PPC | single_day | 20180910 | 20180910 | 44 | accuracy | 0.842857 |
| classification | stabilization | Finger PPC | single_day | 20180910 | 20180910 | 46 | accuracy | 0.8 |
| classification | stabilization | Finger PPC | single_day | 20180910 | 20180910 | 54 | accuracy | 0.842857 |
| classification | stabilization | Finger PPC | single_day | 20180917a | 20180917a | 44 | accuracy | 0.557143 |
| classification | stabilization | Finger PPC | single_day | 20180917a | 20180917a | 46 | accuracy | 0.628571 |
| classification | stabilization | Finger PPC | single_day | 20180917a | 20180917a | 54 | accuracy | 0.571429 |
| classification | stabilization | Finger PPC | single_day | 20180917b | 20180917b | 44 | accuracy | 0.532258 |
| classification | stabilization | Finger PPC | single_day | 20180917b | 20180917b | 46 | accuracy | 0.645161 |
| classification | stabilization | Finger PPC | single_day | 20180917b | 20180917b | 54 | accuracy | 0.645161 |
| classification | stabilization | Finger PPC | single_day | 20180924 | 20180924 | 44 | accuracy | 0.678161 |
| classification | stabilization | Finger PPC | single_day | 20180924 | 20180924 | 46 | accuracy | 0.781609 |
| classification | stabilization | Finger PPC | single_day | 20180924 | 20180924 | 54 | accuracy | 0.701149 |

| | | | | | | | | |
|---|---|---|---|---|---|---|---|---|
| classification | stabilization | Finger PPC | single_day | 20180926 | 20180926 | 44 | accuracy | 0.746835 |
| classification | stabilization | Finger PPC | single_day | 20180926 | 20180926 | 46 | accuracy | 0.658228 |
| classification | stabilization | Finger PPC | single_day | 20180926 | 20180926 | 54 | accuracy | 0.860759 |
| classification | stabilization | Finger PPC | single_day | 20181001 | 20181001 | 44 | accuracy | 0.708861 |
| classification | stabilization | Finger PPC | single_day | 20181001 | 20181001 | 46 | accuracy | 0.670886 |
| classification | stabilization | Finger PPC | single_day | 20181001 | 20181001 | 54 | accuracy | 0.696203 |
| classification | stabilization | Finger PPC | single_day | 20181012 | 20181012 | 44 | accuracy | 0.721519 |
| classification | stabilization | Finger PPC | single_day | 20181012 | 20181012 | 46 | accuracy | 0.759494 |
| classification | stabilization | Finger PPC | single_day | 20181012 | 20181012 | 54 | accuracy | 0.797468 |
| classification | stabilization | Finger PPC | single_day | 20181015 | 20181015 | 44 | accuracy | 0.670886 |
| classification | stabilization | Finger PPC | single_day | 20181015 | 20181015 | 46 | accuracy | 0.772152 |
| classification | stabilization | Finger PPC | single_day | 20181015 | 20181015 | 54 | accuracy | 0.670886 |
| classification | stabilization | Finger PPC | single_day | 20181017 | 20181017 | 44 | accuracy | 0.628571 |
| classification | stabilization | Finger PPC | single_day | 20181017 | 20181017 | 46 | accuracy | 0.657143 |
| classification | stabilization | Finger PPC | single_day | 20181017 | 20181017 | 54 | accuracy | 0.671429 |
| classification | stabilization | Finger PPC | single_day | 20181022 | 20181022 | 44 | accuracy | 0.542857 |
| classification | stabilization | Finger PPC | single_day | 20181022 | 20181022 | 46 | accuracy | 0.628571 |
| classification | stabilization | Finger PPC | single_day | 20181022 | 20181022 | 54 | accuracy | 0.728571 |
| classification | mfsnn | Finger PPC | single_day | 20180910 | 20180910 | 44 | accuracy | 0.727273 |
| classification | mfsnn | Finger PPC | single_day | 20180910 | 20180910 | 46 | accuracy | 0.753247 |
| classification | mfsnn | Finger PPC | single_day | 20180910 | 20180910 | 54 | accuracy | 0.584416 |
| classification | mfsnn | Finger PPC | single_day | 20180917a | 20180917a | 44 | accuracy | 0.766234 |
| classification | mfsnn | Finger PPC | single_day | 20180917a | 20180917a | 46 | accuracy | 0.649351 |
| classification | mfsnn | Finger PPC | single_day | 20180917a | 20180917a | 54 | accuracy | 0.636364 |
| classification | mfsnn | Finger PPC | single_day | 20180917b | 20180917b | 44 | accuracy | 0.676471 |
| classification | mfsnn | Finger PPC | single_day | 20180917b | 20180917b | 46 | accuracy | 0.676471 |
| classification | mfsnn | Finger PPC | single_day | 20180917b | 20180917b | 54 | accuracy | 0.720588 |
| classification | mfsnn | Finger PPC | single_day | 20180924 | 20180924 | 44 | accuracy | 0.8125 |
| classification | mfsnn | Finger PPC | single_day | 20180924 | 20180924 | 46 | accuracy | 0.697917 |
| classification | mfsnn | Finger PPC | single_day | 20180924 | 20180924 | 54 | accuracy | 0.822917 |
| classification | mfsnn | Finger PPC | single_day | 20180926 | 20180926 | 44 | accuracy | 0.804598 |
| classification | mfsnn | Finger PPC | single_day | 20180926 | 20180926 | 46 | accuracy | 0.83908 |
| classification | mfsnn | Finger PPC | single_day | 20180926 | 20180926 | 54 | accuracy | 0.735632 |
| classification | mfsnn | Finger PPC | single_day | 20181001 | 20181001 | 44 | accuracy | 0.712644 |
| classification | mfsnn | Finger PPC | single_day | 20181001 | 20181001 | 46 | accuracy | 0.54023 |
| classification | mfsnn | Finger PPC | single_day | 20181001 | 20181001 | 54 | accuracy | 0.655172 |
| classification | mfsnn | Finger PPC | single_day | 20181012 | 20181012 | 44 | accuracy | 0.689655 |
| classification | mfsnn | Finger PPC | single_day | 20181012 | 20181012 | 46 | accuracy | 0.701149 |
| classification | mfsnn | Finger PPC | single_day | 20181012 | 20181012 | 54 | accuracy | 0.758621 |
| classification | mfsnn | Finger PPC | single_day | 20181015 | 20181015 | 44 | accuracy | 0.91954 |
| classification | mfsnn | Finger PPC | single_day | 20181015 | 20181015 | 46 | accuracy | 0.873563 |
| classification | mfsnn | Finger PPC | single_day | 20181015 | 20181015 | 54 | accuracy | 0.862069 |
| classification | mfsnn | Finger PPC | single_day | 20181017 | 20181017 | 44 | accuracy | 0.727273 |
| classification | mfsnn | Finger PPC | single_day | 20181017 | 20181017 | 46 | accuracy | 0.454545 |
| classification | mfsnn | Finger PPC | single_day | 20181017 | 20181017 | 54 | accuracy | 0.675325 |
| classification | mfsnn | Finger PPC | single_day | 20181022 | 20181022 | 44 | accuracy | 0.818182 |
| classification | mfsnn | Finger PPC | single_day | 20181022 | 20181022 | 46 | accuracy | 0.74026 |
| classification | mfsnn | Finger PPC | single_day | 20181022 | 20181022 | 54 | accuracy | 0.688312 |
| classification | mscformer | Finger PPC | single_day | 20180910 | 20180910 | 44 | accuracy | 0.792208 |
| classification | mscformer | Finger PPC | single_day | 20180910 | 20180910 | 46 | accuracy | 0.805195 |
| classification | mscformer | Finger PPC | single_day | 20180910 | 20180910 | 54 | accuracy | 0.779221 |
| classification | mscformer | Finger PPC | single_day | 20180917a | 20180917a | 44 | accuracy | 0.662338 |
| classification | mscformer | Finger PPC | single_day | 20180917a | 20180917a | 46 | accuracy | 0.805195 |
| classification | mscformer | Finger PPC | single_day | 20180917a | 20180917a | 54 | accuracy | 0.753247 |
| classification | mscformer | Finger PPC | single_day | 20180917b | 20180917b | 44 | accuracy | 0.779412 |
| classification | mscformer | Finger PPC | single_day | 20180917b | 20180917b | 46 | accuracy | 0.764706 |
| classification | mscformer | Finger PPC | single_day | 20180917b | 20180917b | 54 | accuracy | 0.75 |
| classification | mscformer | Finger PPC | single_day | 20180924 | 20180924 | 44 | accuracy | 0.78125 |
| classification | mscformer | Finger PPC | single_day | 20180924 | 20180924 | 46 | accuracy | 0.75 |
| classification | mscformer | Finger PPC | single_day | 20180924 | 20180924 | 54 | accuracy | 0.78125 |
| classification | mscformer | Finger PPC | single_day | 20180926 | 20180926 | 44 | accuracy | 0.804598 |
| classification | mscformer | Finger PPC | single_day | 20180926 | 20180926 | 46 | accuracy | 0.827586 |
| classification | mscformer | Finger PPC | single_day | 20180926 | 20180926 | 54 | accuracy | 0.770115 |

| | | | | | | | | |
|---|---|---|---|---|---|---|---|---|
| classification | mscformer | Finger PPC | single_day | 20181001 | 20181001 | 44 | accuracy | 0.781609 |
| classification | mscformer | Finger PPC | single_day | 20181001 | 20181001 | 46 | accuracy | 0.862069 |
| classification | mscformer | Finger PPC | single_day | 20181001 | 20181001 | 54 | accuracy | 0.747126 |
| classification | mscformer | Finger PPC | single_day | 20181012 | 20181012 | 44 | accuracy | 0.816092 |
| classification | mscformer | Finger PPC | single_day | 20181012 | 20181012 | 46 | accuracy | 0.850575 |
| classification | mscformer | Finger PPC | single_day | 20181012 | 20181012 | 54 | accuracy | 0.689655 |
| classification | mscformer | Finger PPC | single_day | 20181015 | 20181015 | 44 | accuracy | 0.850575 |
| classification | mscformer | Finger PPC | single_day | 20181015 | 20181015 | 46 | accuracy | 0.83908 |
| classification | mscformer | Finger PPC | single_day | 20181015 | 20181015 | 54 | accuracy | 0.735632 |
| classification | mscformer | Finger PPC | single_day | 20181017 | 20181017 | 44 | accuracy | 0.584416 |
| classification | mscformer | Finger PPC | single_day | 20181017 | 20181017 | 46 | accuracy | 0.753247 |
| classification | mscformer | Finger PPC | single_day | 20181017 | 20181017 | 54 | accuracy | 0.766234 |
| classification | mscformer | Finger PPC | single_day | 20181022 | 20181022 | 44 | accuracy | 0.675325 |
| classification | mscformer | Finger PPC | single_day | 20181022 | 20181022 | 46 | accuracy | 0.688312 |
| classification | mscformer | Finger PPC | single_day | 20181022 | 20181022 | 54 | accuracy | 0.779221 |
| classification | nomad | Finger PPC | single_day | 20180910 | 20180910 | 44 | accuracy | 0.948052 |
| classification | nomad | Finger PPC | single_day | 20180910 | 20180910 | 46 | accuracy | 0.883117 |
| classification | nomad | Finger PPC | single_day | 20180910 | 20180910 | 54 | accuracy | 0.818182 |
| classification | nomad | Finger PPC | single_day | 20180917a | 20180917a | 44 | accuracy | 0.831169 |
| classification | nomad | Finger PPC | single_day | 20180917a | 20180917a | 46 | accuracy | 0.818182 |
| classification | nomad | Finger PPC | single_day | 20180917a | 20180917a | 54 | accuracy | 0.831169 |
| classification | nomad | Finger PPC | single_day | 20180917b | 20180917b | 44 | accuracy | 0.897059 |
| classification | nomad | Finger PPC | single_day | 20180917b | 20180917b | 46 | accuracy | 0.838235 |
| classification | nomad | Finger PPC | single_day | 20180917b | 20180917b | 54 | accuracy | 0.705882 |
| classification | nomad | Finger PPC | single_day | 20180924 | 20180924 | 44 | accuracy | 0.802083 |
| classification | nomad | Finger PPC | single_day | 20180924 | 20180924 | 46 | accuracy | 0.916667 |
| classification | nomad | Finger PPC | single_day | 20180924 | 20180924 | 54 | accuracy | 0.895833 |
| classification | nomad | Finger PPC | single_day | 20180926 | 20180926 | 44 | accuracy | 0.850575 |
| classification | nomad | Finger PPC | single_day | 20180926 | 20180926 | 46 | accuracy | 0.827586 |
| classification | nomad | Finger PPC | single_day | 20180926 | 20180926 | 54 | accuracy | 0.91954 |
| classification | nomad | Finger PPC | single_day | 20181001 | 20181001 | 44 | accuracy | 0.816092 |
| classification | nomad | Finger PPC | single_day | 20181001 | 20181001 | 46 | accuracy | 0.862069 |
| classification | nomad | Finger PPC | single_day | 20181001 | 20181001 | 54 | accuracy | 0.896552 |
| classification | nomad | Finger PPC | single_day | 20181012 | 20181012 | 44 | accuracy | 0.885057 |
| classification | nomad | Finger PPC | single_day | 20181012 | 20181012 | 46 | accuracy | 0.793103 |
| classification | nomad | Finger PPC | single_day | 20181012 | 20181012 | 54 | accuracy | 0.850575 |
| classification | nomad | Finger PPC | single_day | 20181015 | 20181015 | 44 | accuracy | 0.873563 |
| classification | nomad | Finger PPC | single_day | 20181015 | 20181015 | 46 | accuracy | 0.827586 |
| classification | nomad | Finger PPC | single_day | 20181015 | 20181015 | 54 | accuracy | 0.896552 |
| classification | nomad | Finger PPC | single_day | 20181017 | 20181017 | 44 | accuracy | 0.779221 |
| classification | nomad | Finger PPC | single_day | 20181017 | 20181017 | 46 | accuracy | 0.805195 |
| classification | nomad | Finger PPC | single_day | 20181017 | 20181017 | 54 | accuracy | 0.636364 |
| classification | nomad | Finger PPC | single_day | 20181022 | 20181022 | 44 | accuracy | 0.805195 |
| classification | nomad | Finger PPC | single_day | 20181022 | 20181022 | 46 | accuracy | 0.766234 |
| classification | nomad | Finger PPC | single_day | 20181022 | 20181022 | 54 | accuracy | 0.87013 |
| classification | mlp | Finger PPC | single_day | 20180910 | 20180910 | 44 | accuracy | 0.974026 |
| classification | mlp | Finger PPC | single_day | 20180910 | 20180910 | 46 | accuracy | 0.909091 |
| classification | mlp | Finger PPC | single_day | 20180910 | 20180910 | 54 | accuracy | 0.948052 |
| classification | mlp | Finger PPC | single_day | 20180917a | 20180917a | 44 | accuracy | 0.896104 |
| classification | mlp | Finger PPC | single_day | 20180917a | 20180917a | 46 | accuracy | 0.857143 |
| classification | mlp | Finger PPC | single_day | 20180917a | 20180917a | 54 | accuracy | 0.844156 |
| classification | mlp | Finger PPC | single_day | 20180917b | 20180917b | 44 | accuracy | 0.926471 |
| classification | mlp | Finger PPC | single_day | 20180917b | 20180917b | 46 | accuracy | 0.897059 |
| classification | mlp | Finger PPC | single_day | 20180917b | 20180917b | 54 | accuracy | 0.882353 |
| classification | mlp | Finger PPC | single_day | 20180924 | 20180924 | 44 | accuracy | 0.9375 |
| classification | mlp | Finger PPC | single_day | 20180924 | 20180924 | 46 | accuracy | 0.885417 |
| classification | mlp | Finger PPC | single_day | 20180924 | 20180924 | 54 | accuracy | 0.90625 |
| classification | mlp | Finger PPC | single_day | 20180926 | 20180926 | 44 | accuracy | 0.91954 |
| classification | mlp | Finger PPC | single_day | 20180926 | 20180926 | 46 | accuracy | 0.931034 |
| classification | mlp | Finger PPC | single_day | 20180926 | 20180926 | 54 | accuracy | 0.942529 |
| classification | mlp | Finger PPC | single_day | 20181001 | 20181001 | 44 | accuracy | 0.862069 |
| classification | mlp | Finger PPC | single_day | 20181001 | 20181001 | 46 | accuracy | 0.965517 |
| classification | mlp | Finger PPC | single_day | 20181001 | 20181001 | 54 | accuracy | 0.862069 |

| | | | | | | | | |
|---|---|---|---|---|---|---|---|---|
| classification | mlp | Finger PPC | single_day | 20181012 | 20181012 | 44 | accuracy | 0.931034 |
| classification | mlp | Finger PPC | single_day | 20181012 | 20181012 | 46 | accuracy | 0.954023 |
| classification | mlp | Finger PPC | single_day | 20181012 | 20181012 | 54 | accuracy | 0.931034 |
| classification | mlp | Finger PPC | single_day | 20181015 | 20181015 | 44 | accuracy | 0.908046 |
| classification | mlp | Finger PPC | single_day | 20181015 | 20181015 | 46 | accuracy | 0.91954 |
| classification | mlp | Finger PPC | single_day | 20181015 | 20181015 | 54 | accuracy | 0.862069 |
| classification | mlp | Finger PPC | single_day | 20181017 | 20181017 | 44 | accuracy | 0.831169 |
| classification | mlp | Finger PPC | single_day | 20181017 | 20181017 | 46 | accuracy | 0.844156 |
| classification | mlp | Finger PPC | single_day | 20181017 | 20181017 | 54 | accuracy | 0.909091 |
| classification | mlp | Finger PPC | single_day | 20181022 | 20181022 | 44 | accuracy | 0.883117 |
| classification | mlp | Finger PPC | single_day | 20181022 | 20181022 | 46 | accuracy | 0.831169 |
| classification | mlp | Finger PPC | single_day | 20181022 | 20181022 | 54 | accuracy | 0.896104 |
| classification | convpi_vae | Finger PPC | single_day | 20180910 | 20180910 | 31 | accuracy | 1 |
| classification | convpi_vae | Finger PPC | single_day | 20180910 | 20180910 | 44 | accuracy | 1 |
| classification | convpi_vae | Finger PPC | single_day | 20180910 | 20180910 | 46 | accuracy | 1 |
| classification | convpi_vae | Finger PPC | single_day | 20180910 | 20180910 | 54 | accuracy | 1 |
| classification | convpi_vae | Finger PPC | single_day | 20180910 | 20180910 | 69 | accuracy | 1 |
| classification | convpi_vae | Finger PPC | single_day | 20180917a | 20180917a | 31 | accuracy | 1 |
| classification | convpi_vae | Finger PPC | single_day | 20180917a | 20180917a | 44 | accuracy | 1 |
| classification | convpi_vae | Finger PPC | single_day | 20180917a | 20180917a | 46 | accuracy | 1 |
| classification | convpi_vae | Finger PPC | single_day | 20180917a | 20180917a | 54 | accuracy | 1 |
| classification | convpi_vae | Finger PPC | single_day | 20180917a | 20180917a | 69 | accuracy | 1 |
| classification | convpi_vae | Finger PPC | single_day | 20180917b | 20180917b | 31 | accuracy | 1 |
| classification | convpi_vae | Finger PPC | single_day | 20180917b | 20180917b | 44 | accuracy | 1 |
| classification | convpi_vae | Finger PPC | single_day | 20180917b | 20180917b | 46 | accuracy | 1 |
| classification | convpi_vae | Finger PPC | single_day | 20180917b | 20180917b | 54 | accuracy | 1 |
| classification | convpi_vae | Finger PPC | single_day | 20180917b | 20180917b | 69 | accuracy | 1 |
| classification | convpi_vae | Finger PPC | single_day | 20180924 | 20180924 | 31 | accuracy | 1 |
| classification | convpi_vae | Finger PPC | single_day | 20180924 | 20180924 | 44 | accuracy | 1 |
| classification | convpi_vae | Finger PPC | single_day | 20180924 | 20180924 | 46 | accuracy | 1 |
| classification | convpi_vae | Finger PPC | single_day | 20180924 | 20180924 | 54 | accuracy | 1 |
| classification | convpi_vae | Finger PPC | single_day | 20180924 | 20180924 | 69 | accuracy | 1 |
| classification | convpi_vae | Finger PPC | single_day | 20180926 | 20180926 | 31 | accuracy | 1 |
| classification | convpi_vae | Finger PPC | single_day | 20180926 | 20180926 | 44 | accuracy | 1 |
| classification | convpi_vae | Finger PPC | single_day | 20180926 | 20180926 | 46 | accuracy | 1 |
| classification | convpi_vae | Finger PPC | single_day | 20180926 | 20180926 | 54 | accuracy | 1 |
| classification | convpi_vae | Finger PPC | single_day | 20180926 | 20180926 | 69 | accuracy | 1 |
| classification | convpi_vae | Finger PPC | single_day | 20181001 | 20181001 | 31 | accuracy | 1 |
| classification | convpi_vae | Finger PPC | single_day | 20181001 | 20181001 | 44 | accuracy | 1 |
| classification | convpi_vae | Finger PPC | single_day | 20181001 | 20181001 | 46 | accuracy | 1 |
| classification | convpi_vae | Finger PPC | single_day | 20181001 | 20181001 | 54 | accuracy | 1 |
| classification | convpi_vae | Finger PPC | single_day | 20181001 | 20181001 | 69 | accuracy | 1 |
| classification | convpi_vae | Finger PPC | single_day | 20181012 | 20181012 | 31 | accuracy | 1 |
| classification | convpi_vae | Finger PPC | single_day | 20181012 | 20181012 | 44 | accuracy | 1 |
| classification | convpi_vae | Finger PPC | single_day | 20181012 | 20181012 | 46 | accuracy | 1 |
| classification | convpi_vae | Finger PPC | single_day | 20181012 | 20181012 | 54 | accuracy | 1 |
| classification | convpi_vae | Finger PPC | single_day | 20181012 | 20181012 | 69 | accuracy | 1 |
| classification | convpi_vae | Finger PPC | single_day | 20181015 | 20181015 | 31 | accuracy | 1 |
| classification | convpi_vae | Finger PPC | single_day | 20181015 | 20181015 | 44 | accuracy | 1 |
| classification | convpi_vae | Finger PPC | single_day | 20181015 | 20181015 | 46 | accuracy | 1 |
| classification | convpi_vae | Finger PPC | single_day | 20181015 | 20181015 | 54 | accuracy | 1 |
| classification | convpi_vae | Finger PPC | single_day | 20181015 | 20181015 | 69 | accuracy | 1 |
| classification | convpi_vae | Finger PPC | single_day | 20181017 | 20181017 | 31 | accuracy | 1 |
| classification | convpi_vae | Finger PPC | single_day | 20181017 | 20181017 | 44 | accuracy | 1 |
| classification | convpi_vae | Finger PPC | single_day | 20181017 | 20181017 | 46 | accuracy | 1 |
| classification | convpi_vae | Finger PPC | single_day | 20181017 | 20181017 | 54 | accuracy | 1 |
| classification | convpi_vae | Finger PPC | single_day | 20181017 | 20181017 | 69 | accuracy | 1 |
| classification | convpi_vae | Finger PPC | single_day | 20181022 | 20181022 | 31 | accuracy | 1 |
| classification | convpi_vae | Finger PPC | single_day | 20181022 | 20181022 | 44 | accuracy | 1 |
| classification | convpi_vae | Finger PPC | single_day | 20181022 | 20181022 | 46 | accuracy | 1 |
| classification | convpi_vae | Finger PPC | single_day | 20181022 | 20181022 | 54 | accuracy | 1 |
| classification | convpi_vae | Finger PPC | single_day | 20181022 | 20181022 | 69 | accuracy | 1 |
| classification | lstm | Finger PPC | cross_day | 20180910__to__20181012 | 20181017__to__20181022 | 31 | accuracy | 0.346354 |

| | | | | | | | | |
|---|---|---|---|---|---|---|---|---|
| classification | lstm | Finger PPC | cross_day | 20180910__to__20181012 | 20181017__to__20181022 | 44 | accuracy | 0.289062 |
| classification | lstm | Finger PPC | cross_day | 20180910__to__20181012 | 20181017__to__20181022 | 46 | accuracy | 0.389323 |
| classification | lstm | Finger PPC | cross_day | 20180910__to__20181012 | 20181017__to__20181022 | 46 | accuracy | 0.389323 |
| classification | lstm | Finger PPC | cross_day | 20180910__to__20181012 | 20181017__to__20181022 | 54 | accuracy | 0.334635 |
| classification | lstm | Finger PPC | cross_day | 20180910__to__20181012 | 20181017__to__20181022 | 54 | accuracy | 0.334635 |
| classification | lstm | Finger PPC | cross_day | 20180910__to__20181012 | 20181017__to__20181022 | 69 | accuracy | 0.279948 |
| classification | lstm | Finger PPC | cross_day | 20180910__to__20181015 | 20181017__to__20181022 | 31 | accuracy | 0.34375 |
| classification | lstm | Finger PPC | cross_day | 20180910__to__20181015 | 20181017__to__20181022 | 44 | accuracy | 0.309896 |
| classification | lstm | Finger PPC | cross_day | 20180910__to__20181015 | 20181017__to__20181022 | 46 | accuracy | 0.377604 |
| classification | lstm | Finger PPC | cross_day | 20180910__to__20181015 | 20181017__to__20181022 | 54 | accuracy | 0.292969 |
| classification | lstm | Finger PPC | cross_day | 20180910__to__20181015 | 20181017__to__20181022 | 69 | accuracy | 0.30599 |
| classification | transformer | Finger PPC | cross_day | 20180910__to__20181012 | 20181017__to__20181022 | 31 | accuracy | 0.236979 |
| classification | transformer | Finger PPC | cross_day | 20180910__to__20181012 | 20181017__to__20181022 | 44 | accuracy | 0.325521 |
| classification | transformer | Finger PPC | cross_day | 20180910__to__20181012 | 20181017__to__20181022 | 44 | accuracy | 0.325521 |
| classification | transformer | Finger PPC | cross_day | 20180910__to__20181012 | 20181017__to__20181022 | 46 | accuracy | 0.299479 |
| classification | transformer | Finger PPC | cross_day | 20180910__to__20181012 | 20181017__to__20181022 | 46 | accuracy | 0.299479 |
| classification | transformer | Finger PPC | cross_day | 20180910__to__20181012 | 20181017__to__20181022 | 54 | accuracy | 0.196615 |
| classification | transformer | Finger PPC | cross_day | 20180910__to__20181012 | 20181017__to__20181022 | 54 | accuracy | 0.196615 |
| classification | transformer | Finger PPC | cross_day | 20180910__to__20181012 | 20181017__to__20181022 | 69 | accuracy | 0.322917 |
| classification | transformer | Finger PPC | cross_day | 20180910__to__20181015 | 20181017__to__20181022 | 31 | accuracy | 0.333333 |
| classification | transformer | Finger PPC | cross_day | 20180910__to__20181015 | 20181017__to__20181022 | 44 | accuracy | 0.371094 |
| classification | transformer | Finger PPC | cross_day | 20180910__to__20181015 | 20181017__to__20181022 | 46 | accuracy | 0.333333 |
| classification | transformer | Finger PPC | cross_day | 20180910__to__20181015 | 20181017__to__20181022 | 54 | accuracy | 0.348958 |
| classification | transformer | Finger PPC | cross_day | 20180910__to__20181015 | 20181017__to__20181022 | 69 | accuracy | 0.360677 |
| classification | lfads | Finger PPC | cross_day | 20180910__to__20181012 | 20181017__to__20181022 | 44 | accuracy | 0.303385 |
| classification | lfads | Finger PPC | cross_day | 20180910__to__20181012 | 20181017__to__20181022 | 44 | accuracy | 0.410156 |
| classification | lfads | Finger PPC | cross_day | 20180910__to__20181012 | 20181017__to__20181022 | 46 | accuracy | 0.328125 |
| classification | lfads | Finger PPC | cross_day | 20180910__to__20181012 | 20181017__to__20181022 | 46 | accuracy | 0.244792 |
| classification | lfads | Finger PPC | cross_day | 20180910__to__20181012 | 20181017__to__20181022 | 54 | accuracy | 0.309896 |
| classification | lfads | Finger PPC | cross_day | 20180910__to__20181012 | 20181017__to__20181022 | 54 | accuracy | 0.345052 |
| classification | cycle_gan | Finger PPC | cross_day | 20180910__to__20181012 | 20181017__to__20181022 | 31 | accuracy | 0.158854 |
| classification | cycle_gan | Finger PPC | cross_day | 20180910__to__20181012 | 20181017__to__20181022 | 44 | accuracy | 0.265583 |
| classification | cycle_gan | Finger PPC | cross_day | 20180910__to__20181012 | 20181017__to__20181022 | 44 | accuracy | 0.252825 |
| classification | cycle_gan | Finger PPC | cross_day | 20180910__to__20181012 | 20181017__to__20181022 | 44 | accuracy | 0.201977 |
| classification | cycle_gan | Finger PPC | cross_day | 20180910__to__20181012 | 20181017__to__20181022 | 44 | accuracy | 0.166667 |
| classification | cycle_gan | Finger PPC | cross_day | 20180910__to__20181012 | 20181017__to__20181022 | 46 | accuracy | 0.222222 |
| classification | cycle_gan | Finger PPC | cross_day | 20180910__to__20181012 | 20181017__to__20181022 | 46 | accuracy | 0.20339 |
| classification | cycle_gan | Finger PPC | cross_day | 20180910__to__20181012 | 20181017__to__20181022 | 46 | accuracy | 0.35452 |
| classification | cycle_gan | Finger PPC | cross_day | 20180910__to__20181012 | 20181017__to__20181022 | 46 | accuracy | 0.166667 |
| classification | cycle_gan | Finger PPC | cross_day | 20180910__to__20181012 | 20181017__to__20181022 | 54 | accuracy | 0.249322 |
| classification | cycle_gan | Finger PPC | cross_day | 20180910__to__20181012 | 20181017__to__20181022 | 54 | accuracy | 0.262712 |
| classification | cycle_gan | Finger PPC | cross_day | 20180910__to__20181012 | 20181017__to__20181022 | 54 | accuracy | 0.238701 |
| classification | cycle_gan | Finger PPC | cross_day | 20180910__to__20181012 | 20181017__to__20181022 | 54 | accuracy | 0.166667 |
| classification | cycle_gan | Finger PPC | cross_day | 20180910__to__20181012 | 20181017__to__20181022 | 69 | accuracy | 0.166667 |
| classification | cycle_gan | Finger PPC | cross_day | 20180910__to__20181015 | 20181017__to__20181022 | 31 | accuracy | 0.425781 |
| classification | cycle_gan | Finger PPC | cross_day | 20180910__to__20181015 | 20181017__to__20181022 | 44 | accuracy | 0.463542 |
| classification | cycle_gan | Finger PPC | cross_day | 20180910__to__20181015 | 20181017__to__20181022 | 46 | accuracy | 0.459635 |
| classification | cycle_gan | Finger PPC | cross_day | 20180910__to__20181015 | 20181017__to__20181022 | 54 | accuracy | 0.483073 |
| classification | cycle_gan | Finger PPC | cross_day | 20180910__to__20181015 | 20181017__to__20181022 | 69 | accuracy | 0.454427 |
| classification | stabilization | Finger PPC | cross_day | 20180910__to__20181012 | 20181017__to__20181022 | 44 | accuracy | 0.371274 |
| classification | stabilization | Finger PPC | cross_day | 20180910__to__20181012 | 20181017__to__20181022 | 44 | accuracy | 0.409605 |
| classification | stabilization | Finger PPC | cross_day | 20180910__to__20181012 | 20181017__to__20181022 | 44 | accuracy | 0.281073 |
| classification | stabilization | Finger PPC | cross_day | 20180910__to__20181012 | 20181017__to__20181022 | 46 | accuracy | 0.372629 |
| classification | stabilization | Finger PPC | cross_day | 20180910__to__20181012 | 20181017__to__20181022 | 46 | accuracy | 0.378531 |
| classification | stabilization | Finger PPC | cross_day | 20180910__to__20181012 | 20181017__to__20181022 | 46 | accuracy | 0.228814 |
| classification | stabilization | Finger PPC | cross_day | 20180910__to__20181012 | 20181017__to__20181022 | 54 | accuracy | 0.371274 |
| classification | stabilization | Finger PPC | cross_day | 20180910__to__20181012 | 20181017__to__20181022 | 54 | accuracy | 0.423729 |
| classification | stabilization | Finger PPC | cross_day | 20180910__to__20181012 | 20181017__to__20181022 | 54 | accuracy | 0.245763 |
| classification | mfsnn | Finger PPC | cross_day | 20180910__to__20181012 | 20181017 | 44 | accuracy | 0.166667 |
| classification | mfsnn | Finger PPC | cross_day | 20180910__to__20181012 | 20181017 | 46 | accuracy | 0.276042 |
| classification | mfsnn | Finger PPC | cross_day | 20180910__to__20181012 | 20181017 | 54 | accuracy | 0.231771 |
| classification | mfsnn | Finger PPC | cross_day | 20180910__to__20181012 | 20181022 | 44 | accuracy | 0.166667 |
| classification | mfsnn | Finger PPC | cross_day | 20180910__to__20181012 | 20181022 | 46 | accuracy | 0.213542 |

| | | | | | | | | |
|---|---|---|---|---|---|---|---|---|
| classification | mfsnn | Finger PPC | cross_day | 20180910__to__20181012 | 20181022 | 54 | accuracy | 0.403646 |
| classification | mscformer | Finger PPC | cross_day | 20180910__to__20181012 | 20181017 | 44 | accuracy | 0.382812 |
| classification | mscformer | Finger PPC | cross_day | 20180910__to__20181012 | 20181017 | 46 | accuracy | 0.393229 |
| classification | mscformer | Finger PPC | cross_day | 20180910__to__20181012 | 20181017 | 54 | accuracy | 0.328125 |
| classification | mscformer | Finger PPC | cross_day | 20180910__to__20181012 | 20181022 | 44 | accuracy | 0.427083 |
| classification | mscformer | Finger PPC | cross_day | 20180910__to__20181012 | 20181022 | 46 | accuracy | 0.46875 |
| classification | mscformer | Finger PPC | cross_day | 20180910__to__20181012 | 20181022 | 54 | accuracy | 0.447917 |
| classification | nomad | Finger PPC | cross_day | 20180910__to__20181012 | 20181017 | 44 | accuracy | 0.440104 |
| classification | nomad | Finger PPC | cross_day | 20180910__to__20181012 | 20181017 | 46 | accuracy | 0.53125 |
| classification | nomad | Finger PPC | cross_day | 20180910__to__20181012 | 20181017 | 54 | accuracy | 0.536458 |
| classification | nomad | Finger PPC | cross_day | 20180910__to__20181012 | 20181022 | 44 | accuracy | 0.479167 |
| classification | nomad | Finger PPC | cross_day | 20180910__to__20181012 | 20181022 | 46 | accuracy | 0.520833 |
| classification | nomad | Finger PPC | cross_day | 20180910__to__20181012 | 20181022 | 54 | accuracy | 0.536458 |
| classification | mlp | Finger PPC | cross_day | 20180910__to__20181012 | 20181017 | 44 | accuracy | 0.356771 |
| classification | mlp | Finger PPC | cross_day | 20180910__to__20181012 | 20181017 | 46 | accuracy | 0.520833 |
| classification | mlp | Finger PPC | cross_day | 20180910__to__20181012 | 20181017 | 54 | accuracy | 0.367188 |
| classification | mlp | Finger PPC | cross_day | 20180910__to__20181012 | 20181022 | 44 | accuracy | 0.385417 |
| classification | mlp | Finger PPC | cross_day | 20180910__to__20181012 | 20181022 | 46 | accuracy | 0.304688 |
| classification | mlp | Finger PPC | cross_day | 20180910__to__20181012 | 20181022 | 54 | accuracy | 0.395833 |
| classification | convpi_vae | Finger PPC | cross_day | 20180910__to__20181012 | 20181017__to__20181022 | 31 | accuracy | 0.161458 |
| classification | convpi_vae | Finger PPC | cross_day | 20180910__to__20181012 | 20181017__to__20181022 | 44 | accuracy | 0.184896 |
| classification | convpi_vae | Finger PPC | cross_day | 20180910__to__20181012 | 20181017__to__20181022 | 46 | accuracy | 0.186198 |
| classification | convpi_vae | Finger PPC | cross_day | 20180910__to__20181012 | 20181017__to__20181022 | 54 | accuracy | 0.203125 |
| classification | convpi_vae | Finger PPC | cross_day | 20180910__to__20181012 | 20181017__to__20181022 | 69 | accuracy | 0.170573 |
| classification | convpi_vae | Finger PPC | cross_day | 20180910__to__20181015 | 20181017__to__20181022 | 31 | accuracy | 0.927083 |
| classification | convpi_vae | Finger PPC | cross_day | 20180910__to__20181015 | 20181017__to__20181022 | 44 | accuracy | 0.979167 |
| classification | convpi_vae | Finger PPC | cross_day | 20180910__to__20181015 | 20181017__to__20181022 | 46 | accuracy | 1 |
| classification | convpi_vae | Finger PPC | cross_day | 20180910__to__20181015 | 20181017__to__20181022 | 54 | accuracy | 1 |
| classification | convpi_vae | Finger PPC | cross_day | 20180910__to__20181015 | 20181017__to__20181022 | 69 | accuracy | 1 |
| classification | lstm | Reaching C-J-M-T_1 | single_day | 20131003 | 20131003 | 31 | accuracy | 0.40625 |
| classification | lstm | Reaching C-J-M-T_1 | single_day | 20131003 | 20131003 | 44 | accuracy | 0.34375 |
| classification | lstm | Reaching C-J-M-T_1 | single_day | 20131003 | 20131003 | 46 | accuracy | 0.4375 |
| classification | lstm | Reaching C-J-M-T_1 | single_day | 20131003 | 20131003 | 54 | accuracy | 0.46875 |
| classification | lstm | Reaching C-J-M-T_1 | single_day | 20131003 | 20131003 | 69 | accuracy | 0.4375 |
| classification | lstm | Reaching C-J-M-T_1 | single_day | 20131022 | 20131022 | 31 | accuracy | 0.580645 |
| classification | lstm | Reaching C-J-M-T_1 | single_day | 20131022 | 20131022 | 44 | accuracy | 0.483871 |
| classification | lstm | Reaching C-J-M-T_1 | single_day | 20131022 | 20131022 | 46 | accuracy | 0.548387 |
| classification | lstm | Reaching C-J-M-T_1 | single_day | 20131022 | 20131022 | 54 | accuracy | 0.516129 |
| classification | lstm | Reaching C-J-M-T_1 | single_day | 20131022 | 20131022 | 69 | accuracy | 0.419355 |
| classification | lstm | Reaching C-J-M-T_1 | single_day | 20131023 | 20131023 | 31 | accuracy | 0.487179 |
| classification | lstm | Reaching C-J-M-T_1 | single_day | 20131023 | 20131023 | 44 | accuracy | 0.589744 |
| classification | lstm | Reaching C-J-M-T_1 | single_day | 20131023 | 20131023 | 46 | accuracy | 0.589744 |
| classification | lstm | Reaching C-J-M-T_1 | single_day | 20131023 | 20131023 | 54 | accuracy | 0.461538 |
| classification | lstm | Reaching C-J-M-T_1 | single_day | 20131023 | 20131023 | 69 | accuracy | 0.512821 |
| classification | lstm | Reaching C-J-M-T_1 | single_day | 20131031 | 20131031 | 31 | accuracy | 0.477273 |
| classification | lstm | Reaching C-J-M-T_1 | single_day | 20131031 | 20131031 | 44 | accuracy | 0.5 |
| classification | lstm | Reaching C-J-M-T_1 | single_day | 20131031 | 20131031 | 69 | accuracy | 0.454545 |
| classification | lstm | Reaching C-J-M-T_1 | single_day | 20131101 | 20131101 | 31 | accuracy | 0.54 |
| classification | lstm | Reaching C-J-M-T_1 | single_day | 20131101 | 20131101 | 44 | accuracy | 0.62 |
| classification | lstm | Reaching C-J-M-T_1 | single_day | 20131101 | 20131101 | 46 | accuracy | 0.68 |
| classification | lstm | Reaching C-J-M-T_1 | single_day | 20131101 | 20131101 | 54 | accuracy | 0.58 |
| classification | lstm | Reaching C-J-M-T_1 | single_day | 20131101 | 20131101 | 69 | accuracy | 0.6 |
| classification | lstm | Reaching C-J-M-T_1 | single_day | 20131203 | 20131203 | 31 | accuracy | 0.588235 |
| classification | lstm | Reaching C-J-M-T_1 | single_day | 20131203 | 20131203 | 44 | accuracy | 0.588235 |
| classification | lstm | Reaching C-J-M-T_1 | single_day | 20131203 | 20131203 | 46 | accuracy | 0.441176 |
| classification | lstm | Reaching C-J-M-T_1 | single_day | 20131203 | 20131203 | 54 | accuracy | 0.5 |
| classification | lstm | Reaching C-J-M-T_1 | single_day | 20131203 | 20131203 | 69 | accuracy | 0.470588 |
| classification | lstm | Reaching C-J-M-T_1 | single_day | 20131204 | 20131204 | 31 | accuracy | 0.363636 |
| classification | lstm | Reaching C-J-M-T_1 | single_day | 20131204 | 20131204 | 44 | accuracy | 0.454545 |
| classification | lstm | Reaching C-J-M-T_1 | single_day | 20131204 | 20131204 | 46 | accuracy | 0.393939 |
| classification | lstm | Reaching C-J-M-T_1 | single_day | 20131204 | 20131204 | 54 | accuracy | 0.333333 |
| classification | lstm | Reaching C-J-M-T_1 | single_day | 20131204 | 20131204 | 69 | accuracy | 0.272727 |
| classification | lstm | Reaching C-J-M-T_1 | single_day | 20131219 | 20131219 | 31 | accuracy | 0.416667 |

| | | | | | | | | |
|---|---|---|---|---|---|---|---|---|
| classification | lstm | Reaching C-J-M-T_1 | single_day | 20131219 | 20131219 | 44 | accuracy | 0.555556 |
| classification | lstm | Reaching C-J-M-T_1 | single_day | 20131219 | 20131219 | 46 | accuracy | 0.527778 |
| classification | lstm | Reaching C-J-M-T_1 | single_day | 20131219 | 20131219 | 54 | accuracy | 0.527778 |
| classification | lstm | Reaching C-J-M-T_1 | single_day | 20131219 | 20131219 | 69 | accuracy | 0.583333 |
| classification | lstm | Reaching C-J-M-T_1 | single_day | 20131220 | 20131220 | 31 | accuracy | 0.45 |
| classification | lstm | Reaching C-J-M-T_1 | single_day | 20131220 | 20131220 | 44 | accuracy | 0.35 |
| classification | lstm | Reaching C-J-M-T_1 | single_day | 20131220 | 20131220 | 46 | accuracy | 0.425 |
| classification | lstm | Reaching C-J-M-T_1 | single_day | 20131220 | 20131220 | 54 | accuracy | 0.45 |
| classification | lstm | Reaching C-J-M-T_1 | single_day | 20131220 | 20131220 | 69 | accuracy | 0.35 |
| classification | lstm | Reaching C-J-M-T_1 | single_day | 20150309 | 20150309 | 31 | accuracy | 0.754098 |
| classification | lstm | Reaching C-J-M-T_1 | single_day | 20150309 | 20150309 | 44 | accuracy | 0.713115 |
| classification | lstm | Reaching C-J-M-T_1 | single_day | 20150309 | 20150309 | 46 | accuracy | 0.721311 |
| classification | lstm | Reaching C-J-M-T_1 | single_day | 20150309 | 20150309 | 54 | accuracy | 0.737705 |
| classification | lstm | Reaching C-J-M-T_1 | single_day | 20150311 | 20150311 | 31 | accuracy | 0.961749 |
| classification | lstm | Reaching C-J-M-T_1 | single_day | 20150311 | 20150311 | 44 | accuracy | 0.928962 |
| classification | lstm | Reaching C-J-M-T_1 | single_day | 20150311 | 20150311 | 46 | accuracy | 0.928962 |
| classification | lstm | Reaching C-J-M-T_1 | single_day | 20150311 | 20150311 | 54 | accuracy | 0.956284 |
| classification | lstm | Reaching C-J-M-T_1 | single_day | 20150311 | 20150311 | 69 | accuracy | 0.912568 |
| classification | lstm | Reaching C-J-M-T_1 | single_day | 20150312 | 20150312 | 31 | accuracy | 0.917582 |
| classification | lstm | Reaching C-J-M-T_1 | single_day | 20150312 | 20150312 | 44 | accuracy | 0.934066 |
| classification | lstm | Reaching C-J-M-T_1 | single_day | 20150312 | 20150312 | 46 | accuracy | 0.972527 |
| classification | lstm | Reaching C-J-M-T_1 | single_day | 20150312 | 20150312 | 54 | accuracy | 0.93956 |
| classification | lstm | Reaching C-J-M-T_1 | single_day | 20150312 | 20150312 | 69 | accuracy | 0.961538 |
| classification | lstm | Reaching C-J-M-T_1 | single_day | 20150313 | 20150313 | 31 | accuracy | 0.923077 |
| classification | lstm | Reaching C-J-M-T_1 | single_day | 20150313 | 20150313 | 44 | accuracy | 0.971154 |
| classification | lstm | Reaching C-J-M-T_1 | single_day | 20150313 | 20150313 | 46 | accuracy | 0.971154 |
| classification | lstm | Reaching C-J-M-T_1 | single_day | 20150313 | 20150313 | 54 | accuracy | 0.961538 |
| classification | lstm | Reaching C-J-M-T_1 | single_day | 20150313 | 20150313 | 69 | accuracy | 0.966346 |
| classification | lstm | Reaching C-J-M-T_1 | single_day | 20150319 | 20150319 | 31 | accuracy | 0.898058 |
| classification | lstm | Reaching C-J-M-T_1 | single_day | 20150319 | 20150319 | 44 | accuracy | 0.932039 |
| classification | lstm | Reaching C-J-M-T_1 | single_day | 20150319 | 20150319 | 46 | accuracy | 0.956311 |
| classification | lstm | Reaching C-J-M-T_1 | single_day | 20150319 | 20150319 | 54 | accuracy | 0.946602 |
| classification | lstm | Reaching C-J-M-T_1 | single_day | 20150319 | 20150319 | 69 | accuracy | 0.980583 |
| classification | lstm | Reaching C-J-M-T_1 | single_day | 20150629 | 20150629 | 31 | accuracy | 0.555556 |
| classification | lstm | Reaching C-J-M-T_1 | single_day | 20150629 | 20150629 | 44 | accuracy | 0.666667 |
| classification | lstm | Reaching C-J-M-T_1 | single_day | 20150629 | 20150629 | 44 | accuracy | 0.694444 |
| classification | lstm | Reaching C-J-M-T_1 | single_day | 20150629 | 20150629 | 46 | accuracy | 0.472222 |
| classification | lstm | Reaching C-J-M-T_1 | single_day | 20150629 | 20150629 | 46 | accuracy | 0.583333 |
| classification | lstm | Reaching C-J-M-T_1 | single_day | 20150629 | 20150629 | 54 | accuracy | 0.472222 |
| classification | lstm | Reaching C-J-M-T_1 | single_day | 20150629 | 20150629 | 54 | accuracy | 0.555556 |
| classification | lstm | Reaching C-J-M-T_1 | single_day | 20150629 | 20150629 | 69 | accuracy | 0.638889 |
| classification | lstm | Reaching C-J-M-T_1 | single_day | 20150630 | 20150630 | 31 | accuracy | 0.722222 |
| classification | lstm | Reaching C-J-M-T_1 | single_day | 20150630 | 20150630 | 44 | accuracy | 0.583333 |
| classification | lstm | Reaching C-J-M-T_1 | single_day | 20150630 | 20150630 | 44 | accuracy | 0.722222 |
| classification | lstm | Reaching C-J-M-T_1 | single_day | 20150630 | 20150630 | 46 | accuracy | 0.666667 |
| classification | lstm | Reaching C-J-M-T_1 | single_day | 20150630 | 20150630 | 46 | accuracy | 0.861111 |
| classification | lstm | Reaching C-J-M-T_1 | single_day | 20150630 | 20150630 | 54 | accuracy | 0.75 |
| classification | lstm | Reaching C-J-M-T_1 | single_day | 20150630 | 20150630 | 54 | accuracy | 0.833333 |
| classification | lstm | Reaching C-J-M-T_1 | single_day | 20150630 | 20150630 | 69 | accuracy | 0.638889 |
| classification | lstm | Reaching C-J-M-T_1 | single_day | 20150701 | 20150701 | 31 | accuracy | 0.6 |
| classification | lstm | Reaching C-J-M-T_1 | single_day | 20150701 | 20150701 | 44 | accuracy | 0.7 |
| classification | lstm | Reaching C-J-M-T_1 | single_day | 20150701 | 20150701 | 44 | accuracy | 0.8 |
| classification | lstm | Reaching C-J-M-T_1 | single_day | 20150701 | 20150701 | 46 | accuracy | 0.65 |
| classification | lstm | Reaching C-J-M-T_1 | single_day | 20150701 | 20150701 | 46 | accuracy | 0.7 |
| classification | lstm | Reaching C-J-M-T_1 | single_day | 20150701 | 20150701 | 54 | accuracy | 0.65 |
| classification | lstm | Reaching C-J-M-T_1 | single_day | 20150701 | 20150701 | 54 | accuracy | 0.7 |
| classification | lstm | Reaching C-J-M-T_1 | single_day | 20150701 | 20150701 | 69 | accuracy | 0.7 |
| classification | lstm | Reaching C-J-M-T_1 | single_day | 20150703 | 20150703 | 31 | accuracy | 0.459459 |
| classification | lstm | Reaching C-J-M-T_1 | single_day | 20150703 | 20150703 | 44 | accuracy | 0.648649 |
| classification | lstm | Reaching C-J-M-T_1 | single_day | 20150703 | 20150703 | 46 | accuracy | 0.486486 |
| classification | lstm | Reaching C-J-M-T_1 | single_day | 20150703 | 20150703 | 54 | accuracy | 0.351351 |
| classification | lstm | Reaching C-J-M-T_1 | single_day | 20150706 | 20150706 | 31 | accuracy | 0.513514 |
| classification | lstm | Reaching C-J-M-T_1 | single_day | 20150706 | 20150706 | 44 | accuracy | 0.810811 |

| | | | | | | | | |
|---|---|---|---|---|---|---|---|---|
| classification | lstm | Reaching C-J-M-T_1 | single_day | 20150706 | 20150706 | 46 | accuracy | 0.648649 |
| classification | lstm | Reaching C-J-M-T_1 | single_day | 20150706 | 20150706 | 46 | accuracy | 0.756757 |
| classification | lstm | Reaching C-J-M-T_1 | single_day | 20150706 | 20150706 | 54 | accuracy | 0.567568 |
| classification | lstm | Reaching C-J-M-T_1 | single_day | 20150706 | 20150706 | 54 | accuracy | 0.702703 |
| classification | lstm | Reaching C-J-M-T_1 | single_day | 20150706 | 20150706 | 69 | accuracy | 0.648649 |
| classification | lstm | Reaching C-J-M-T_1 | single_day | 20150707 | 20150707 | 31 | accuracy | 0.473684 |
| classification | lstm | Reaching C-J-M-T_1 | single_day | 20150707 | 20150707 | 44 | accuracy | 0.631579 |
| classification | lstm | Reaching C-J-M-T_1 | single_day | 20150707 | 20150707 | 46 | accuracy | 0.5 |
| classification | lstm | Reaching C-J-M-T_1 | single_day | 20150707 | 20150707 | 54 | accuracy | 0.605263 |
| classification | lstm | Reaching C-J-M-T_1 | single_day | 20150707 | 20150707 | 69 | accuracy | 0.368421 |
| classification | lstm | Reaching C-J-M-T_1 | single_day | 20150708 | 20150708 | 31 | accuracy | 0.736842 |
| classification | lstm | Reaching C-J-M-T_1 | single_day | 20150708 | 20150708 | 44 | accuracy | 0.631579 |
| classification | lstm | Reaching C-J-M-T_1 | single_day | 20150708 | 20150708 | 44 | accuracy | 0.868421 |
| classification | lstm | Reaching C-J-M-T_1 | single_day | 20150708 | 20150708 | 46 | accuracy | 0.710526 |
| classification | lstm | Reaching C-J-M-T_1 | single_day | 20150708 | 20150708 | 46 | accuracy | 0.789474 |
| classification | lstm | Reaching C-J-M-T_1 | single_day | 20150708 | 20150708 | 54 | accuracy | 0.657895 |
| classification | lstm | Reaching C-J-M-T_1 | single_day | 20150708 | 20150708 | 54 | accuracy | 0.894737 |
| classification | lstm | Reaching C-J-M-T_1 | single_day | 20150708 | 20150708 | 69 | accuracy | 0.605263 |
| classification | lstm | Reaching C-J-M-T_1 | single_day | 20150709 | 20150709 | 31 | accuracy | 0.7 |
| classification | lstm | Reaching C-J-M-T_1 | single_day | 20150709 | 20150709 | 44 | accuracy | 0.75 |
| classification | lstm | Reaching C-J-M-T_1 | single_day | 20150709 | 20150709 | 44 | accuracy | 0.85 |
| classification | lstm | Reaching C-J-M-T_1 | single_day | 20150709 | 20150709 | 46 | accuracy | 0.675 |
| classification | lstm | Reaching C-J-M-T_1 | single_day | 20150709 | 20150709 | 46 | accuracy | 0.775 |
| classification | lstm | Reaching C-J-M-T_1 | single_day | 20150709 | 20150709 | 54 | accuracy | 0.65 |
| classification | lstm | Reaching C-J-M-T_1 | single_day | 20150709 | 20150709 | 54 | accuracy | 0.725 |
| classification | lstm | Reaching C-J-M-T_1 | single_day | 20150709 | 20150709 | 69 | accuracy | 0.6 |
| classification | lstm | Reaching C-J-M-T_1 | single_day | 20150710 | 20150710 | 31 | accuracy | 0.804878 |
| classification | lstm | Reaching C-J-M-T_1 | single_day | 20150710 | 20150710 | 44 | accuracy | 0.560976 |
| classification | lstm | Reaching C-J-M-T_1 | single_day | 20150710 | 20150710 | 44 | accuracy | 0.731707 |
| classification | lstm | Reaching C-J-M-T_1 | single_day | 20150710 | 20150710 | 46 | accuracy | 0.609756 |
| classification | lstm | Reaching C-J-M-T_1 | single_day | 20150710 | 20150710 | 46 | accuracy | 0.731707 |
| classification | lstm | Reaching C-J-M-T_1 | single_day | 20150710 | 20150710 | 54 | accuracy | 0.609756 |
| classification | lstm | Reaching C-J-M-T_1 | single_day | 20150710 | 20150710 | 54 | accuracy | 0.731707 |
| classification | lstm | Reaching C-J-M-T_1 | single_day | 20150710 | 20150710 | 69 | accuracy | 0.609756 |
| classification | lstm | Reaching C-J-M-T_1 | single_day | 20150713 | 20150713 | 31 | accuracy | 0.675676 |
| classification | lstm | Reaching C-J-M-T_1 | single_day | 20150713 | 20150713 | 44 | accuracy | 0.648649 |
| classification | lstm | Reaching C-J-M-T_1 | single_day | 20150713 | 20150713 | 46 | accuracy | 0.72973 |
| classification | lstm | Reaching C-J-M-T_1 | single_day | 20150713 | 20150713 | 54 | accuracy | 0.756757 |
| classification | lstm | Reaching C-J-M-T_1 | single_day | 20150713 | 20150713 | 69 | accuracy | 0.648649 |
| classification | lstm | Reaching C-J-M-T_1 | single_day | 20150714 | 20150714 | 31 | accuracy | 0.625 |
| classification | lstm | Reaching C-J-M-T_1 | single_day | 20150714 | 20150714 | 44 | accuracy | 0.6 |
| classification | lstm | Reaching C-J-M-T_1 | single_day | 20150714 | 20150714 | 46 | accuracy | 0.65 |
| classification | lstm | Reaching C-J-M-T_1 | single_day | 20150714 | 20150714 | 54 | accuracy | 0.75 |
| classification | lstm | Reaching C-J-M-T_1 | single_day | 20150714 | 20150714 | 69 | accuracy | 0.675 |
| classification | lstm | Reaching C-J-M-T_1 | single_day | 20150715 | 20150715 | 31 | accuracy | 0.666667 |
| classification | lstm | Reaching C-J-M-T_1 | single_day | 20150715 | 20150715 | 44 | accuracy | 0.820513 |
| classification | lstm | Reaching C-J-M-T_1 | single_day | 20150715 | 20150715 | 46 | accuracy | 0.769231 |
| classification | lstm | Reaching C-J-M-T_1 | single_day | 20150715 | 20150715 | 54 | accuracy | 0.74359 |
| classification | lstm | Reaching C-J-M-T_1 | single_day | 20150715 | 20150715 | 69 | accuracy | 0.74359 |
| classification | lstm | Reaching C-J-M-T_1 | single_day | 20150716 | 20150716 | 31 | accuracy | 0.820513 |
| classification | lstm | Reaching C-J-M-T_1 | single_day | 20150716 | 20150716 | 44 | accuracy | 0.666667 |
| classification | lstm | Reaching C-J-M-T_1 | single_day | 20150716 | 20150716 | 46 | accuracy | 0.717949 |
| classification | lstm | Reaching C-J-M-T_1 | single_day | 20150716 | 20150716 | 54 | accuracy | 0.769231 |
| classification | lstm | Reaching C-J-M-T_1 | single_day | 20150716 | 20150716 | 69 | accuracy | 0.769231 |
| classification | lstm | Reaching C-J-M-T_1 | single_day | 20151103 | 20151103 | 31 | accuracy | 0.56 |
| classification | lstm | Reaching C-J-M-T_1 | single_day | 20151103 | 20151103 | 44 | accuracy | 0.4 |
| classification | lstm | Reaching C-J-M-T_1 | single_day | 20151103 | 20151103 | 46 | accuracy | 0.6 |
| classification | lstm | Reaching C-J-M-T_1 | single_day | 20151103 | 20151103 | 54 | accuracy | 0.64 |
| classification | lstm | Reaching C-J-M-T_1 | single_day | 20151103 | 20151103 | 69 | accuracy | 0.54 |
| classification | lstm | Reaching C-J-M-T_1 | single_day | 20151104 | 20151104 | 31 | accuracy | 0.8125 |
| classification | lstm | Reaching C-J-M-T_1 | single_day | 20151104 | 20151104 | 54 | accuracy | 0.78125 |
| classification | lstm | Reaching C-J-M-T_1 | single_day | 20151104 | 20151104 | 69 | accuracy | 0.765625 |
| classification | lstm | Reaching C-J-M-T_1 | single_day | 20151106 | 20151106 | 31 | accuracy | 0.711538 |

| | | | | | | | | |
|---|---|---|---|---|---|---|---|---|
| classification | lstm | Reaching C-J-M-T_1 | single_day | 20151106 | 20151106 | 44 | accuracy | 0.538462 |
| classification | lstm | Reaching C-J-M-T_1 | single_day | 20151106 | 20151106 | 46 | accuracy | 0.730769 |
| classification | lstm | Reaching C-J-M-T_1 | single_day | 20151106 | 20151106 | 54 | accuracy | 0.75 |
| classification | lstm | Reaching C-J-M-T_1 | single_day | 20151106 | 20151106 | 69 | accuracy | 0.692308 |
| classification | lstm | Reaching C-J-M-T_1 | single_day | 20151109 | 20151109 | 31 | accuracy | 0.628571 |
| classification | lstm | Reaching C-J-M-T_1 | single_day | 20151109 | 20151109 | 44 | accuracy | 0.457143 |
| classification | lstm | Reaching C-J-M-T_1 | single_day | 20151109 | 20151109 | 46 | accuracy | 0.342857 |
| classification | lstm | Reaching C-J-M-T_1 | single_day | 20151109 | 20151109 | 54 | accuracy | 0.371429 |
| classification | lstm | Reaching C-J-M-T_1 | single_day | 20151109 | 20151109 | 69 | accuracy | 0.371429 |
| classification | lstm | Reaching C-J-M-T_1 | single_day | 20151110 | 20151110 | 31 | accuracy | 0.625 |
| classification | lstm | Reaching C-J-M-T_1 | single_day | 20151110 | 20151110 | 46 | accuracy | 0.6 |
| classification | lstm | Reaching C-J-M-T_1 | single_day | 20151110 | 20151110 | 54 | accuracy | 0.5 |
| classification | lstm | Reaching C-J-M-T_1 | single_day | 20151110 | 20151110 | 69 | accuracy | 0.475 |
| classification | lstm | Reaching C-J-M-T_1 | single_day | 20151112 | 20151112 | 31 | accuracy | 0.710526 |
| classification | lstm | Reaching C-J-M-T_1 | single_day | 20151112 | 20151112 | 44 | accuracy | 0.578947 |
| classification | lstm | Reaching C-J-M-T_1 | single_day | 20151112 | 20151112 | 46 | accuracy | 0.710526 |
| classification | lstm | Reaching C-J-M-T_1 | single_day | 20151112 | 20151112 | 54 | accuracy | 0.657895 |
| classification | lstm | Reaching C-J-M-T_1 | single_day | 20151112 | 20151112 | 69 | accuracy | 0.605263 |
| classification | lstm | Reaching C-J-M-T_1 | single_day | 20151113 | 20151113 | 31 | accuracy | 0.736842 |
| classification | lstm | Reaching C-J-M-T_1 | single_day | 20151113 | 20151113 | 44 | accuracy | 0.578947 |
| classification | lstm | Reaching C-J-M-T_1 | single_day | 20151113 | 20151113 | 46 | accuracy | 0.684211 |
| classification | lstm | Reaching C-J-M-T_1 | single_day | 20151113 | 20151113 | 54 | accuracy | 0.526316 |
| classification | lstm | Reaching C-J-M-T_1 | single_day | 20151113 | 20151113 | 69 | accuracy | 0.710526 |
| classification | lstm | Reaching C-J-M-T_1 | single_day | 20151116 | 20151116 | 31 | accuracy | 0.470588 |
| classification | lstm | Reaching C-J-M-T_1 | single_day | 20151116 | 20151116 | 44 | accuracy | 0.470588 |
| classification | lstm | Reaching C-J-M-T_1 | single_day | 20151116 | 20151116 | 46 | accuracy | 0.558824 |
| classification | lstm | Reaching C-J-M-T_1 | single_day | 20151116 | 20151116 | 54 | accuracy | 0.5 |
| classification | lstm | Reaching C-J-M-T_1 | single_day | 20151116 | 20151116 | 69 | accuracy | 0.617647 |
| classification | lstm | Reaching C-J-M-T_1 | single_day | 20151117 | 20151117 | 31 | accuracy | 0.605263 |
| classification | lstm | Reaching C-J-M-T_1 | single_day | 20151117 | 20151117 | 44 | accuracy | 0.684211 |
| classification | lstm | Reaching C-J-M-T_1 | single_day | 20151117 | 20151117 | 46 | accuracy | 0.552632 |
| classification | lstm | Reaching C-J-M-T_1 | single_day | 20151117 | 20151117 | 54 | accuracy | 0.631579 |
| classification | lstm | Reaching C-J-M-T_1 | single_day | 20151117 | 20151117 | 69 | accuracy | 0.605263 |
| classification | lstm | Reaching C-J-M-T_1 | single_day | 20151119 | 20151119 | 31 | accuracy | 0.714286 |
| classification | lstm | Reaching C-J-M-T_1 | single_day | 20151119 | 20151119 | 44 | accuracy | 0.666667 |
| classification | lstm | Reaching C-J-M-T_1 | single_day | 20151119 | 20151119 | 46 | accuracy | 0.738095 |
| classification | lstm | Reaching C-J-M-T_1 | single_day | 20151119 | 20151119 | 54 | accuracy | 0.666667 |
| classification | lstm | Reaching C-J-M-T_1 | single_day | 20151119 | 20151119 | 69 | accuracy | 0.833333 |
| classification | lstm | Reaching C-J-M-T_1 | single_day | 20151120 | 20151120 | 31 | accuracy | 0.352941 |
| classification | lstm | Reaching C-J-M-T_1 | single_day | 20151120 | 20151120 | 44 | accuracy | 0.617647 |
| classification | lstm | Reaching C-J-M-T_1 | single_day | 20151120 | 20151120 | 46 | accuracy | 0.676471 |
| classification | lstm | Reaching C-J-M-T_1 | single_day | 20151120 | 20151120 | 54 | accuracy | 0.558824 |
| classification | lstm | Reaching C-J-M-T_1 | single_day | 20151120 | 20151120 | 69 | accuracy | 0.558824 |
| classification | lstm | Reaching C-J-M-T_1 | single_day | 20151201 | 20151201 | 31 | accuracy | 0.657895 |
| classification | lstm | Reaching C-J-M-T_1 | single_day | 20151201 | 20151201 | 44 | accuracy | 0.710526 |
| classification | lstm | Reaching C-J-M-T_1 | single_day | 20151201 | 20151201 | 46 | accuracy | 0.736842 |
| classification | lstm | Reaching C-J-M-T_1 | single_day | 20151201 | 20151201 | 54 | accuracy | 0.657895 |
| classification | lstm | Reaching C-J-M-T_1 | single_day | 20151201 | 20151201 | 69 | accuracy | 0.763158 |
| classification | lstm | Reaching C-J-M-T_1 | single_day | 20160909 | 20160909 | 31 | accuracy | 0.645161 |
| classification | lstm | Reaching C-J-M-T_1 | single_day | 20160909 | 20160909 | 44 | accuracy | 0.709677 |
| classification | lstm | Reaching C-J-M-T_1 | single_day | 20160909 | 20160909 | 46 | accuracy | 0.612903 |
| classification | lstm | Reaching C-J-M-T_1 | single_day | 20160909 | 20160909 | 54 | accuracy | 0.645161 |
| classification | lstm | Reaching C-J-M-T_1 | single_day | 20160909 | 20160909 | 69 | accuracy | 0.612903 |
| classification | lstm | Reaching C-J-M-T_1 | single_day | 20160912 | 20160912 | 31 | accuracy | 0.682927 |
| classification | lstm | Reaching C-J-M-T_1 | single_day | 20160912 | 20160912 | 44 | accuracy | 0.756098 |
| classification | lstm | Reaching C-J-M-T_1 | single_day | 20160912 | 20160912 | 54 | accuracy | 0.731707 |
| classification | lstm | Reaching C-J-M-T_1 | single_day | 20160912 | 20160912 | 69 | accuracy | 0.829268 |
| classification | lstm | Reaching C-J-M-T_1 | single_day | 20160914 | 20160914 | 31 | accuracy | 0.820513 |
| classification | lstm | Reaching C-J-M-T_1 | single_day | 20160914 | 20160914 | 44 | accuracy | 0.769231 |
| classification | lstm | Reaching C-J-M-T_1 | single_day | 20160914 | 20160914 | 46 | accuracy | 0.769231 |
| classification | lstm | Reaching C-J-M-T_1 | single_day | 20160914 | 20160914 | 54 | accuracy | 0.692308 |
| classification | lstm | Reaching C-J-M-T_1 | single_day | 20160914 | 20160914 | 69 | accuracy | 0.820513 |
| classification | lstm | Reaching C-J-M-T_1 | single_day | 20160915 | 20160915 | 31 | accuracy | 0.864865 |

| | | | | | | | | |
|---|---|---|---|---|---|---|---|---|
| classification | lstm | Reaching C-J-M-T_1 | single_day | 20160915 | 20160915 | 44 | accuracy | 0.837838 |
| classification | lstm | Reaching C-J-M-T_1 | single_day | 20160915 | 20160915 | 46 | accuracy | 0.72973 |
| classification | lstm | Reaching C-J-M-T_1 | single_day | 20160915 | 20160915 | 54 | accuracy | 0.837838 |
| classification | lstm | Reaching C-J-M-T_1 | single_day | 20160915 | 20160915 | 69 | accuracy | 0.756757 |
| classification | lstm | Reaching C-J-M-T_1 | single_day | 20160919 | 20160919 | 31 | accuracy | 0.789474 |
| classification | lstm | Reaching C-J-M-T_1 | single_day | 20160919 | 20160919 | 44 | accuracy | 0.868421 |
| classification | lstm | Reaching C-J-M-T_1 | single_day | 20160919 | 20160919 | 46 | accuracy | 0.815789 |
| classification | lstm | Reaching C-J-M-T_1 | single_day | 20160919 | 20160919 | 54 | accuracy | 0.684211 |
| classification | lstm | Reaching C-J-M-T_1 | single_day | 20160919 | 20160919 | 69 | accuracy | 0.789474 |
| classification | lstm | Reaching C-J-M-T_1 | single_day | 20160921 | 20160921 | 31 | accuracy | 0.666667 |
| classification | lstm | Reaching C-J-M-T_1 | single_day | 20160921 | 20160921 | 44 | accuracy | 0.861111 |
| classification | lstm | Reaching C-J-M-T_1 | single_day | 20160921 | 20160921 | 46 | accuracy | 0.916667 |
| classification | lstm | Reaching C-J-M-T_1 | single_day | 20160921 | 20160921 | 54 | accuracy | 0.75 |
| classification | lstm | Reaching C-J-M-T_1 | single_day | 20160921 | 20160921 | 69 | accuracy | 0.75 |
| classification | lstm | Reaching C-J-M-T_1 | single_day | 20160923 | 20160923 | 31 | accuracy | 0.853659 |
| classification | lstm | Reaching C-J-M-T_1 | single_day | 20160923 | 20160923 | 44 | accuracy | 0.853659 |
| classification | lstm | Reaching C-J-M-T_1 | single_day | 20160923 | 20160923 | 46 | accuracy | 0.780488 |
| classification | lstm | Reaching C-J-M-T_1 | single_day | 20160923 | 20160923 | 54 | accuracy | 0.829268 |
| classification | lstm | Reaching C-J-M-T_1 | single_day | 20160923 | 20160923 | 69 | accuracy | 0.780488 |
| classification | lstm | Reaching C-J-M-T_1 | single_day | 20160929 | 20160929 | 31 | accuracy | 0.809524 |
| classification | lstm | Reaching C-J-M-T_1 | single_day | 20160929 | 20160929 | 44 | accuracy | 0.666667 |
| classification | lstm | Reaching C-J-M-T_1 | single_day | 20160929 | 20160929 | 46 | accuracy | 0.809524 |
| classification | lstm | Reaching C-J-M-T_1 | single_day | 20160929 | 20160929 | 54 | accuracy | 0.738095 |
| classification | lstm | Reaching C-J-M-T_1 | single_day | 20160929 | 20160929 | 69 | accuracy | 0.595238 |
| classification | lstm | Reaching C-J-M-T_1 | single_day | 20161005 | 20161005 | 31 | accuracy | 0.756098 |
| classification | lstm | Reaching C-J-M-T_1 | single_day | 20161005 | 20161005 | 44 | accuracy | 0.707317 |
| classification | lstm | Reaching C-J-M-T_1 | single_day | 20161005 | 20161005 | 46 | accuracy | 0.756098 |
| classification | lstm | Reaching C-J-M-T_1 | single_day | 20161005 | 20161005 | 54 | accuracy | 0.853659 |
| classification | lstm | Reaching C-J-M-T_1 | single_day | 20161005 | 20161005 | 69 | accuracy | 0.731707 |
| classification | lstm | Reaching C-J-M-T_1 | single_day | 20161006 | 20161006 | 31 | accuracy | 0.642857 |
| classification | lstm | Reaching C-J-M-T_1 | single_day | 20161006 | 20161006 | 44 | accuracy | 0.642857 |
| classification | lstm | Reaching C-J-M-T_1 | single_day | 20161006 | 20161006 | 46 | accuracy | 0.761905 |
| classification | lstm | Reaching C-J-M-T_1 | single_day | 20161006 | 20161006 | 54 | accuracy | 0.690476 |
| classification | lstm | Reaching C-J-M-T_1 | single_day | 20161006 | 20161006 | 69 | accuracy | 0.761905 |
| classification | lstm | Reaching C-J-M-T_1 | single_day | 20161007 | 20161007 | 31 | accuracy | 0.628571 |
| classification | lstm | Reaching C-J-M-T_1 | single_day | 20161007 | 20161007 | 54 | accuracy | 0.657143 |
| classification | lstm | Reaching C-J-M-T_1 | single_day | 20161007 | 20161007 | 69 | accuracy | 0.6 |
| classification | lstm | Reaching C-J-M-T_1 | single_day | 20161011 | 20161011 | 31 | accuracy | 0.825 |
| classification | lstm | Reaching C-J-M-T_1 | single_day | 20161011 | 20161011 | 44 | accuracy | 0.85 |
| classification | lstm | Reaching C-J-M-T_1 | single_day | 20161011 | 20161011 | 46 | accuracy | 0.85 |
| classification | lstm | Reaching C-J-M-T_1 | single_day | 20161011 | 20161011 | 54 | accuracy | 0.8 |
| classification | lstm | Reaching C-J-M-T_1 | single_day | 20161011 | 20161011 | 69 | accuracy | 0.8 |
| classification | lstm | Reaching C-J-M-T_1 | single_day | 20161013 | 20161013 | 31 | accuracy | 0.833333 |
| classification | lstm | Reaching C-J-M-T_1 | single_day | 20161013 | 20161013 | 44 | accuracy | 0.895833 |
| classification | lstm | Reaching C-J-M-T_1 | single_day | 20161013 | 20161013 | 46 | accuracy | 0.916667 |
| classification | lstm | Reaching C-J-M-T_1 | single_day | 20161013 | 20161013 | 54 | accuracy | 0.791667 |
| classification | lstm | Reaching C-J-M-T_1 | single_day | 20161013 | 20161013 | 69 | accuracy | 0.8125 |
| classification | lstm | Reaching C-J-M-T_1 | single_day | 20161021 | 20161021 | 31 | accuracy | 0.862069 |
| classification | lstm | Reaching C-J-M-T_1 | single_day | 20161021 | 20161021 | 44 | accuracy | 0.87931 |
| classification | lstm | Reaching C-J-M-T_1 | single_day | 20161021 | 20161021 | 46 | accuracy | 0.793103 |
| classification | lstm | Reaching C-J-M-T_1 | single_day | 20161021 | 20161021 | 54 | accuracy | 0.913793 |
| classification | lstm | Reaching C-J-M-T_1 | single_day | 20161021 | 20161021 | 69 | accuracy | 0.844828 |
| classification | transformer | Reaching C-J-M-T_1 | single_day | 20131003 | 20131003 | 31 | accuracy | 0.625 |
| classification | transformer | Reaching C-J-M-T_1 | single_day | 20131003 | 20131003 | 44 | accuracy | 0.875 |
| classification | transformer | Reaching C-J-M-T_1 | single_day | 20131003 | 20131003 | 46 | accuracy | 0.78125 |
| classification | transformer | Reaching C-J-M-T_1 | single_day | 20131003 | 20131003 | 54 | accuracy | 0.625 |
| classification | transformer | Reaching C-J-M-T_1 | single_day | 20131003 | 20131003 | 69 | accuracy | 0.78125 |
| classification | transformer | Reaching C-J-M-T_1 | single_day | 20131022 | 20131022 | 31 | accuracy | 0.774194 |
| classification | transformer | Reaching C-J-M-T_1 | single_day | 20131022 | 20131022 | 44 | accuracy | 0.870968 |
| classification | transformer | Reaching C-J-M-T_1 | single_day | 20131022 | 20131022 | 46 | accuracy | 0.806452 |
| classification | transformer | Reaching C-J-M-T_1 | single_day | 20131022 | 20131022 | 54 | accuracy | 0.870968 |
| classification | transformer | Reaching C-J-M-T_1 | single_day | 20131022 | 20131022 | 69 | accuracy | 0.741935 |
| classification | transformer | Reaching C-J-M-T_1 | single_day | 20131023 | 20131023 | 31 | accuracy | 0.846154 |

| | | | | | | | | |
|---|---|---|---|---|---|---|---|---|
| classification | transformer | Reaching C-J-M-T_1 | single_day | 20131023 | 20131023 | 44 | accuracy | 0.948718 |
| classification | transformer | Reaching C-J-M-T_1 | single_day | 20131023 | 20131023 | 46 | accuracy | 0.923077 |
| classification | transformer | Reaching C-J-M-T_1 | single_day | 20131023 | 20131023 | 54 | accuracy | 0.794872 |
| classification | transformer | Reaching C-J-M-T_1 | single_day | 20131023 | 20131023 | 69 | accuracy | 0.897436 |
| classification | transformer | Reaching C-J-M-T_1 | single_day | 20131031 | 20131031 | 31 | accuracy | 0.772727 |
| classification | transformer | Reaching C-J-M-T_1 | single_day | 20131031 | 20131031 | 44 | accuracy | 0.772727 |
| classification | transformer | Reaching C-J-M-T_1 | single_day | 20131031 | 20131031 | 69 | accuracy | 0.772727 |
| classification | transformer | Reaching C-J-M-T_1 | single_day | 20131101 | 20131101 | 31 | accuracy | 0.76 |
| classification | transformer | Reaching C-J-M-T_1 | single_day | 20131101 | 20131101 | 44 | accuracy | 0.82 |
| classification | transformer | Reaching C-J-M-T_1 | single_day | 20131101 | 20131101 | 46 | accuracy | 0.82 |
| classification | transformer | Reaching C-J-M-T_1 | single_day | 20131101 | 20131101 | 54 | accuracy | 0.92 |
| classification | transformer | Reaching C-J-M-T_1 | single_day | 20131101 | 20131101 | 69 | accuracy | 0.68 |
| classification | transformer | Reaching C-J-M-T_1 | single_day | 20131203 | 20131203 | 31 | accuracy | 0.823529 |
| classification | transformer | Reaching C-J-M-T_1 | single_day | 20131203 | 20131203 | 44 | accuracy | 0.823529 |
| classification | transformer | Reaching C-J-M-T_1 | single_day | 20131203 | 20131203 | 46 | accuracy | 0.794118 |
| classification | transformer | Reaching C-J-M-T_1 | single_day | 20131203 | 20131203 | 54 | accuracy | 0.823529 |
| classification | transformer | Reaching C-J-M-T_1 | single_day | 20131203 | 20131203 | 69 | accuracy | 0.852941 |
| classification | transformer | Reaching C-J-M-T_1 | single_day | 20131204 | 20131204 | 31 | accuracy | 0.666667 |
| classification | transformer | Reaching C-J-M-T_1 | single_day | 20131204 | 20131204 | 44 | accuracy | 0.636364 |
| classification | transformer | Reaching C-J-M-T_1 | single_day | 20131204 | 20131204 | 46 | accuracy | 0.484848 |
| classification | transformer | Reaching C-J-M-T_1 | single_day | 20131204 | 20131204 | 54 | accuracy | 0.757576 |
| classification | transformer | Reaching C-J-M-T_1 | single_day | 20131204 | 20131204 | 69 | accuracy | 0.818182 |
| classification | transformer | Reaching C-J-M-T_1 | single_day | 20131219 | 20131219 | 31 | accuracy | 0.777778 |
| classification | transformer | Reaching C-J-M-T_1 | single_day | 20131219 | 20131219 | 44 | accuracy | 0.638889 |
| classification | transformer | Reaching C-J-M-T_1 | single_day | 20131219 | 20131219 | 46 | accuracy | 0.777778 |
| classification | transformer | Reaching C-J-M-T_1 | single_day | 20131219 | 20131219 | 54 | accuracy | 0.722222 |
| classification | transformer | Reaching C-J-M-T_1 | single_day | 20131219 | 20131219 | 69 | accuracy | 0.694444 |
| classification | transformer | Reaching C-J-M-T_1 | single_day | 20131220 | 20131220 | 31 | accuracy | 0.7 |
| classification | transformer | Reaching C-J-M-T_1 | single_day | 20131220 | 20131220 | 44 | accuracy | 0.575 |
| classification | transformer | Reaching C-J-M-T_1 | single_day | 20131220 | 20131220 | 46 | accuracy | 0.7 |
| classification | transformer | Reaching C-J-M-T_1 | single_day | 20131220 | 20131220 | 54 | accuracy | 0.725 |
| classification | transformer | Reaching C-J-M-T_1 | single_day | 20131220 | 20131220 | 69 | accuracy | 0.65 |
| classification | transformer | Reaching C-J-M-T_1 | single_day | 20150309 | 20150309 | 31 | accuracy | 0.844262 |
| classification | transformer | Reaching C-J-M-T_1 | single_day | 20150309 | 20150309 | 44 | accuracy | 0.860656 |
| classification | transformer | Reaching C-J-M-T_1 | single_day | 20150309 | 20150309 | 46 | accuracy | 0.836066 |
| classification | transformer | Reaching C-J-M-T_1 | single_day | 20150309 | 20150309 | 54 | accuracy | 0.827869 |
| classification | transformer | Reaching C-J-M-T_1 | single_day | 20150311 | 20150311 | 31 | accuracy | 0.961749 |
| classification | transformer | Reaching C-J-M-T_1 | single_day | 20150311 | 20150311 | 44 | accuracy | 0.945355 |
| classification | transformer | Reaching C-J-M-T_1 | single_day | 20150311 | 20150311 | 46 | accuracy | 0.918033 |
| classification | transformer | Reaching C-J-M-T_1 | single_day | 20150311 | 20150311 | 54 | accuracy | 0.934426 |
| classification | transformer | Reaching C-J-M-T_1 | single_day | 20150311 | 20150311 | 69 | accuracy | 0.923497 |
| classification | transformer | Reaching C-J-M-T_1 | single_day | 20150312 | 20150312 | 31 | accuracy | 0.961538 |
| classification | transformer | Reaching C-J-M-T_1 | single_day | 20150312 | 20150312 | 44 | accuracy | 0.945055 |
| classification | transformer | Reaching C-J-M-T_1 | single_day | 20150312 | 20150312 | 46 | accuracy | 0.950549 |
| classification | transformer | Reaching C-J-M-T_1 | single_day | 20150312 | 20150312 | 54 | accuracy | 0.956044 |
| classification | transformer | Reaching C-J-M-T_1 | single_day | 20150312 | 20150312 | 69 | accuracy | 0.934066 |
| classification | transformer | Reaching C-J-M-T_1 | single_day | 20150313 | 20150313 | 31 | accuracy | 0.961538 |
| classification | transformer | Reaching C-J-M-T_1 | single_day | 20150313 | 20150313 | 44 | accuracy | 0.932692 |
| classification | transformer | Reaching C-J-M-T_1 | single_day | 20150313 | 20150313 | 46 | accuracy | 0.971154 |
| classification | transformer | Reaching C-J-M-T_1 | single_day | 20150313 | 20150313 | 54 | accuracy | 0.971154 |
| classification | transformer | Reaching C-J-M-T_1 | single_day | 20150313 | 20150313 | 69 | accuracy | 0.966346 |
| classification | transformer | Reaching C-J-M-T_1 | single_day | 20150319 | 20150319 | 31 | accuracy | 0.970874 |
| classification | transformer | Reaching C-J-M-T_1 | single_day | 20150319 | 20150319 | 44 | accuracy | 0.970874 |
| classification | transformer | Reaching C-J-M-T_1 | single_day | 20150319 | 20150319 | 46 | accuracy | 0.946602 |
| classification | transformer | Reaching C-J-M-T_1 | single_day | 20150319 | 20150319 | 54 | accuracy | 0.956311 |
| classification | transformer | Reaching C-J-M-T_1 | single_day | 20150319 | 20150319 | 69 | accuracy | 0.941748 |
| classification | transformer | Reaching C-J-M-T_1 | single_day | 20150629 | 20150629 | 31 | accuracy | 0.861111 |
| classification | transformer | Reaching C-J-M-T_1 | single_day | 20150629 | 20150629 | 44 | accuracy | 0.916667 |
| classification | transformer | Reaching C-J-M-T_1 | single_day | 20150629 | 20150629 | 46 | accuracy | 0.861111 |
| classification | transformer | Reaching C-J-M-T_1 | single_day | 20150629 | 20150629 | 46 | accuracy | 0.888889 |
| classification | transformer | Reaching C-J-M-T_1 | single_day | 20150629 | 20150629 | 54 | accuracy | 0.916667 |
| classification | transformer | Reaching C-J-M-T_1 | single_day | 20150629 | 20150629 | 54 | accuracy | 0.777778 |
| classification | transformer | Reaching C-J-M-T_1 | single_day | 20150629 | 20150629 | 69 | accuracy | 0.972222 |

| | | | | | | | | |
|---|---|---|---|---|---|---|---|---|
| classification | transformer | Reaching C-J-M-T_1 | single_day | 20150630 | 20150630 | 31 | accuracy | 0.972222 |
| classification | transformer | Reaching C-J-M-T_1 | single_day | 20150630 | 20150630 | 44 | accuracy | 0.944444 |
| classification | transformer | Reaching C-J-M-T_1 | single_day | 20150630 | 20150630 | 46 | accuracy | 0.888889 |
| classification | transformer | Reaching C-J-M-T_1 | single_day | 20150630 | 20150630 | 46 | accuracy | 0.916667 |
| classification | transformer | Reaching C-J-M-T_1 | single_day | 20150630 | 20150630 | 54 | accuracy | 0.972222 |
| classification | transformer | Reaching C-J-M-T_1 | single_day | 20150630 | 20150630 | 69 | accuracy | 0.944444 |
| classification | transformer | Reaching C-J-M-T_1 | single_day | 20150701 | 20150701 | 31 | accuracy | 0.875 |
| classification | transformer | Reaching C-J-M-T_1 | single_day | 20150701 | 20150701 | 44 | accuracy | 0.875 |
| classification | transformer | Reaching C-J-M-T_1 | single_day | 20150701 | 20150701 | 44 | accuracy | 0.825 |
| classification | transformer | Reaching C-J-M-T_1 | single_day | 20150701 | 20150701 | 46 | accuracy | 0.875 |
| classification | transformer | Reaching C-J-M-T_1 | single_day | 20150701 | 20150701 | 46 | accuracy | 0.8 |
| classification | transformer | Reaching C-J-M-T_1 | single_day | 20150701 | 20150701 | 54 | accuracy | 0.875 |
| classification | transformer | Reaching C-J-M-T_1 | single_day | 20150701 | 20150701 | 69 | accuracy | 0.875 |
| classification | transformer | Reaching C-J-M-T_1 | single_day | 20150703 | 20150703 | 31 | accuracy | 0.783784 |
| classification | transformer | Reaching C-J-M-T_1 | single_day | 20150703 | 20150703 | 44 | accuracy | 0.837838 |
| classification | transformer | Reaching C-J-M-T_1 | single_day | 20150703 | 20150703 | 46 | accuracy | 0.945946 |
| classification | transformer | Reaching C-J-M-T_1 | single_day | 20150703 | 20150703 | 54 | accuracy | 0.837838 |
| classification | transformer | Reaching C-J-M-T_1 | single_day | 20150706 | 20150706 | 31 | accuracy | 0.918919 |
| classification | transformer | Reaching C-J-M-T_1 | single_day | 20150706 | 20150706 | 44 | accuracy | 0.864865 |
| classification | transformer | Reaching C-J-M-T_1 | single_day | 20150706 | 20150706 | 46 | accuracy | 0.972973 |
| classification | transformer | Reaching C-J-M-T_1 | single_day | 20150706 | 20150706 | 46 | accuracy | 0.945946 |
| classification | transformer | Reaching C-J-M-T_1 | single_day | 20150706 | 20150706 | 54 | accuracy | 0.945946 |
| classification | transformer | Reaching C-J-M-T_1 | single_day | 20150706 | 20150706 | 69 | accuracy | 0.783784 |
| classification | transformer | Reaching C-J-M-T_1 | single_day | 20150707 | 20150707 | 31 | accuracy | 0.921053 |
| classification | transformer | Reaching C-J-M-T_1 | single_day | 20150707 | 20150707 | 44 | accuracy | 0.894737 |
| classification | transformer | Reaching C-J-M-T_1 | single_day | 20150707 | 20150707 | 46 | accuracy | 0.868421 |
| classification | transformer | Reaching C-J-M-T_1 | single_day | 20150707 | 20150707 | 54 | accuracy | 0.815789 |
| classification | transformer | Reaching C-J-M-T_1 | single_day | 20150707 | 20150707 | 69 | accuracy | 0.815789 |
| classification | transformer | Reaching C-J-M-T_1 | single_day | 20150708 | 20150708 | 31 | accuracy | 0.894737 |
| classification | transformer | Reaching C-J-M-T_1 | single_day | 20150708 | 20150708 | 44 | accuracy | 0.868421 |
| classification | transformer | Reaching C-J-M-T_1 | single_day | 20150708 | 20150708 | 46 | accuracy | 0.921053 |
| classification | transformer | Reaching C-J-M-T_1 | single_day | 20150708 | 20150708 | 46 | accuracy | 0.868421 |
| classification | transformer | Reaching C-J-M-T_1 | single_day | 20150708 | 20150708 | 54 | accuracy | 0.868421 |
| classification | transformer | Reaching C-J-M-T_1 | single_day | 20150708 | 20150708 | 54 | accuracy | 0.842105 |
| classification | transformer | Reaching C-J-M-T_1 | single_day | 20150708 | 20150708 | 69 | accuracy | 0.921053 |
| classification | transformer | Reaching C-J-M-T_1 | single_day | 20150709 | 20150709 | 31 | accuracy | 0.95 |
| classification | transformer | Reaching C-J-M-T_1 | single_day | 20150709 | 20150709 | 44 | accuracy | 0.95 |
| classification | transformer | Reaching C-J-M-T_1 | single_day | 20150709 | 20150709 | 44 | accuracy | 0.9 |
| classification | transformer | Reaching C-J-M-T_1 | single_day | 20150709 | 20150709 | 46 | accuracy | 0.9 |
| classification | transformer | Reaching C-J-M-T_1 | single_day | 20150709 | 20150709 | 46 | accuracy | 0.85 |
| classification | transformer | Reaching C-J-M-T_1 | single_day | 20150709 | 20150709 | 54 | accuracy | 0.975 |
| classification | transformer | Reaching C-J-M-T_1 | single_day | 20150709 | 20150709 | 54 | accuracy | 0.925 |
| classification | transformer | Reaching C-J-M-T_1 | single_day | 20150709 | 20150709 | 69 | accuracy | 0.975 |
| classification | transformer | Reaching C-J-M-T_1 | single_day | 20150710 | 20150710 | 31 | accuracy | 0.926829 |
| classification | transformer | Reaching C-J-M-T_1 | single_day | 20150710 | 20150710 | 44 | accuracy | 0.926829 |
| classification | transformer | Reaching C-J-M-T_1 | single_day | 20150710 | 20150710 | 44 | accuracy | 0.804878 |
| classification | transformer | Reaching C-J-M-T_1 | single_day | 20150710 | 20150710 | 46 | accuracy | 0.780488 |
| classification | transformer | Reaching C-J-M-T_1 | single_day | 20150710 | 20150710 | 46 | accuracy | 0.853659 |
| classification | transformer | Reaching C-J-M-T_1 | single_day | 20150710 | 20150710 | 54 | accuracy | 0.878049 |
| classification | transformer | Reaching C-J-M-T_1 | single_day | 20150710 | 20150710 | 54 | accuracy | 0.707317 |
| classification | transformer | Reaching C-J-M-T_1 | single_day | 20150710 | 20150710 | 69 | accuracy | 0.829268 |
| classification | transformer | Reaching C-J-M-T_1 | single_day | 20150713 | 20150713 | 31 | accuracy | 0.945946 |
| classification | transformer | Reaching C-J-M-T_1 | single_day | 20150713 | 20150713 | 44 | accuracy | 1 |
| classification | transformer | Reaching C-J-M-T_1 | single_day | 20150713 | 20150713 | 46 | accuracy | 1 |
| classification | transformer | Reaching C-J-M-T_1 | single_day | 20150713 | 20150713 | 54 | accuracy | 0.918919 |
| classification | transformer | Reaching C-J-M-T_1 | single_day | 20150713 | 20150713 | 69 | accuracy | 0.945946 |
| classification | transformer | Reaching C-J-M-T_1 | single_day | 20150714 | 20150714 | 31 | accuracy | 0.95 |
| classification | transformer | Reaching C-J-M-T_1 | single_day | 20150714 | 20150714 | 44 | accuracy | 0.975 |
| classification | transformer | Reaching C-J-M-T_1 | single_day | 20150714 | 20150714 | 46 | accuracy | 0.9 |
| classification | transformer | Reaching C-J-M-T_1 | single_day | 20150714 | 20150714 | 54 | accuracy | 0.875 |
| classification | transformer | Reaching C-J-M-T_1 | single_day | 20150714 | 20150714 | 69 | accuracy | 0.95 |
| classification | transformer | Reaching C-J-M-T_1 | single_day | 20150715 | 20150715 | 31 | accuracy | 0.974359 |
| classification | transformer | Reaching C-J-M-T_1 | single_day | 20150715 | 20150715 | 44 | accuracy | 0.974359 |

| | | | | | | | | |
|---|---|---|---|---|---|---|---|---|
| classification | transformer | Reaching C-J-M-T_1 | single_day | 20150715 | 20150715 | 46 | accuracy | 0.948718 |
| classification | transformer | Reaching C-J-M-T_1 | single_day | 20150715 | 20150715 | 54 | accuracy | 0.948718 |
| classification | transformer | Reaching C-J-M-T_1 | single_day | 20150715 | 20150715 | 69 | accuracy | 0.974359 |
| classification | transformer | Reaching C-J-M-T_1 | single_day | 20150716 | 20150716 | 31 | accuracy | 0.923077 |
| classification | transformer | Reaching C-J-M-T_1 | single_day | 20150716 | 20150716 | 44 | accuracy | 0.948718 |
| classification | transformer | Reaching C-J-M-T_1 | single_day | 20150716 | 20150716 | 46 | accuracy | 0.948718 |
| classification | transformer | Reaching C-J-M-T_1 | single_day | 20150716 | 20150716 | 54 | accuracy | 0.974359 |
| classification | transformer | Reaching C-J-M-T_1 | single_day | 20150716 | 20150716 | 69 | accuracy | 0.923077 |
| classification | transformer | Reaching C-J-M-T_1 | single_day | 20151103 | 20151103 | 31 | accuracy | 0.74 |
| classification | transformer | Reaching C-J-M-T_1 | single_day | 20151103 | 20151103 | 44 | accuracy | 0.82 |
| classification | transformer | Reaching C-J-M-T_1 | single_day | 20151103 | 20151103 | 46 | accuracy | 0.86 |
| classification | transformer | Reaching C-J-M-T_1 | single_day | 20151103 | 20151103 | 54 | accuracy | 0.78 |
| classification | transformer | Reaching C-J-M-T_1 | single_day | 20151103 | 20151103 | 69 | accuracy | 0.82 |
| classification | transformer | Reaching C-J-M-T_1 | single_day | 20151104 | 20151104 | 31 | accuracy | 0.90625 |
| classification | transformer | Reaching C-J-M-T_1 | single_day | 20151104 | 20151104 | 54 | accuracy | 0.921875 |
| classification | transformer | Reaching C-J-M-T_1 | single_day | 20151104 | 20151104 | 69 | accuracy | 0.890625 |
| classification | transformer | Reaching C-J-M-T_1 | single_day | 20151106 | 20151106 | 31 | accuracy | 0.826923 |
| classification | transformer | Reaching C-J-M-T_1 | single_day | 20151106 | 20151106 | 44 | accuracy | 0.865385 |
| classification | transformer | Reaching C-J-M-T_1 | single_day | 20151106 | 20151106 | 46 | accuracy | 0.884615 |
| classification | transformer | Reaching C-J-M-T_1 | single_day | 20151106 | 20151106 | 54 | accuracy | 1 |
| classification | transformer | Reaching C-J-M-T_1 | single_day | 20151106 | 20151106 | 69 | accuracy | 0.884615 |
| classification | transformer | Reaching C-J-M-T_1 | single_day | 20151109 | 20151109 | 31 | accuracy | 0.857143 |
| classification | transformer | Reaching C-J-M-T_1 | single_day | 20151109 | 20151109 | 44 | accuracy | 0.857143 |
| classification | transformer | Reaching C-J-M-T_1 | single_day | 20151109 | 20151109 | 46 | accuracy | 0.771429 |
| classification | transformer | Reaching C-J-M-T_1 | single_day | 20151109 | 20151109 | 54 | accuracy | 0.685714 |
| classification | transformer | Reaching C-J-M-T_1 | single_day | 20151109 | 20151109 | 69 | accuracy | 0.857143 |
| classification | transformer | Reaching C-J-M-T_1 | single_day | 20151110 | 20151110 | 31 | accuracy | 0.85 |
| classification | transformer | Reaching C-J-M-T_1 | single_day | 20151110 | 20151110 | 46 | accuracy | 0.975 |
| classification | transformer | Reaching C-J-M-T_1 | single_day | 20151110 | 20151110 | 54 | accuracy | 0.9 |
| classification | transformer | Reaching C-J-M-T_1 | single_day | 20151110 | 20151110 | 69 | accuracy | 0.95 |
| classification | transformer | Reaching C-J-M-T_1 | single_day | 20151112 | 20151112 | 31 | accuracy | 0.947368 |
| classification | transformer | Reaching C-J-M-T_1 | single_day | 20151112 | 20151112 | 44 | accuracy | 0.763158 |
| classification | transformer | Reaching C-J-M-T_1 | single_day | 20151112 | 20151112 | 46 | accuracy | 0.894737 |
| classification | transformer | Reaching C-J-M-T_1 | single_day | 20151112 | 20151112 | 54 | accuracy | 0.842105 |
| classification | transformer | Reaching C-J-M-T_1 | single_day | 20151112 | 20151112 | 69 | accuracy | 0.842105 |
| classification | transformer | Reaching C-J-M-T_1 | single_day | 20151113 | 20151113 | 31 | accuracy | 0.947368 |
| classification | transformer | Reaching C-J-M-T_1 | single_day | 20151113 | 20151113 | 44 | accuracy | 0.973684 |
| classification | transformer | Reaching C-J-M-T_1 | single_day | 20151113 | 20151113 | 46 | accuracy | 1 |
| classification | transformer | Reaching C-J-M-T_1 | single_day | 20151113 | 20151113 | 54 | accuracy | 0.947368 |
| classification | transformer | Reaching C-J-M-T_1 | single_day | 20151113 | 20151113 | 69 | accuracy | 0.868421 |
| classification | transformer | Reaching C-J-M-T_1 | single_day | 20151116 | 20151116 | 31 | accuracy | 0.911765 |
| classification | transformer | Reaching C-J-M-T_1 | single_day | 20151116 | 20151116 | 44 | accuracy | 0.941176 |
| classification | transformer | Reaching C-J-M-T_1 | single_day | 20151116 | 20151116 | 46 | accuracy | 0.882353 |
| classification | transformer | Reaching C-J-M-T_1 | single_day | 20151116 | 20151116 | 54 | accuracy | 0.852941 |
| classification | transformer | Reaching C-J-M-T_1 | single_day | 20151116 | 20151116 | 69 | accuracy | 0.882353 |
| classification | transformer | Reaching C-J-M-T_1 | single_day | 20151117 | 20151117 | 31 | accuracy | 0.868421 |
| classification | transformer | Reaching C-J-M-T_1 | single_day | 20151117 | 20151117 | 44 | accuracy | 0.947368 |
| classification | transformer | Reaching C-J-M-T_1 | single_day | 20151117 | 20151117 | 46 | accuracy | 0.921053 |
| classification | transformer | Reaching C-J-M-T_1 | single_day | 20151117 | 20151117 | 54 | accuracy | 0.789474 |
| classification | transformer | Reaching C-J-M-T_1 | single_day | 20151117 | 20151117 | 69 | accuracy | 0.763158 |
| classification | transformer | Reaching C-J-M-T_1 | single_day | 20151119 | 20151119 | 31 | accuracy | 0.928571 |
| classification | transformer | Reaching C-J-M-T_1 | single_day | 20151119 | 20151119 | 44 | accuracy | 1 |
| classification | transformer | Reaching C-J-M-T_1 | single_day | 20151119 | 20151119 | 46 | accuracy | 0.952381 |
| classification | transformer | Reaching C-J-M-T_1 | single_day | 20151119 | 20151119 | 54 | accuracy | 0.928571 |
| classification | transformer | Reaching C-J-M-T_1 | single_day | 20151119 | 20151119 | 69 | accuracy | 0.857143 |
| classification | transformer | Reaching C-J-M-T_1 | single_day | 20151120 | 20151120 | 31 | accuracy | 0.970588 |
| classification | transformer | Reaching C-J-M-T_1 | single_day | 20151120 | 20151120 | 44 | accuracy | 0.941176 |
| classification | transformer | Reaching C-J-M-T_1 | single_day | 20151120 | 20151120 | 46 | accuracy | 0.970588 |
| classification | transformer | Reaching C-J-M-T_1 | single_day | 20151120 | 20151120 | 54 | accuracy | 0.911765 |
| classification | transformer | Reaching C-J-M-T_1 | single_day | 20151120 | 20151120 | 69 | accuracy | 0.911765 |
| classification | transformer | Reaching C-J-M-T_1 | single_day | 20151201 | 20151201 | 31 | accuracy | 0.868421 |
| classification | transformer | Reaching C-J-M-T_1 | single_day | 20151201 | 20151201 | 44 | accuracy | 0.947368 |
| classification | transformer | Reaching C-J-M-T_1 | single_day | 20151201 | 20151201 | 46 | accuracy | 0.973684 |

| | | | | | | | | |
|---|---|---|---|---|---|---|---|---|
| classification | transformer | Reaching C-J-M-T_1 | single_day | 20151201 | 20151201 | 54 | accuracy | 0.973684 |
| classification | transformer | Reaching C-J-M-T_1 | single_day | 20151201 | 20151201 | 69 | accuracy | 0.868421 |
| classification | transformer | Reaching C-J-M-T_1 | single_day | 20160909 | 20160909 | 31 | accuracy | 0.870968 |
| classification | transformer | Reaching C-J-M-T_1 | single_day | 20160909 | 20160909 | 44 | accuracy | 0.806452 |
| classification | transformer | Reaching C-J-M-T_1 | single_day | 20160909 | 20160909 | 46 | accuracy | 0.903226 |
| classification | transformer | Reaching C-J-M-T_1 | single_day | 20160909 | 20160909 | 54 | accuracy | 0.903226 |
| classification | transformer | Reaching C-J-M-T_1 | single_day | 20160909 | 20160909 | 69 | accuracy | 0.935484 |
| classification | transformer | Reaching C-J-M-T_1 | single_day | 20160912 | 20160912 | 31 | accuracy | 1 |
| classification | transformer | Reaching C-J-M-T_1 | single_day | 20160912 | 20160912 | 44 | accuracy | 1 |
| classification | transformer | Reaching C-J-M-T_1 | single_day | 20160912 | 20160912 | 54 | accuracy | 0.97561 |
| classification | transformer | Reaching C-J-M-T_1 | single_day | 20160912 | 20160912 | 69 | accuracy | 0.97561 |
| classification | transformer | Reaching C-J-M-T_1 | single_day | 20160914 | 20160914 | 31 | accuracy | 0.948718 |
| classification | transformer | Reaching C-J-M-T_1 | single_day | 20160914 | 20160914 | 44 | accuracy | 0.948718 |
| classification | transformer | Reaching C-J-M-T_1 | single_day | 20160914 | 20160914 | 46 | accuracy | 1 |
| classification | transformer | Reaching C-J-M-T_1 | single_day | 20160914 | 20160914 | 54 | accuracy | 0.948718 |
| classification | transformer | Reaching C-J-M-T_1 | single_day | 20160914 | 20160914 | 69 | accuracy | 0.974359 |
| classification | transformer | Reaching C-J-M-T_1 | single_day | 20160915 | 20160915 | 31 | accuracy | 0.972973 |
| classification | transformer | Reaching C-J-M-T_1 | single_day | 20160915 | 20160915 | 44 | accuracy | 0.945946 |
| classification | transformer | Reaching C-J-M-T_1 | single_day | 20160915 | 20160915 | 46 | accuracy | 0.972973 |
| classification | transformer | Reaching C-J-M-T_1 | single_day | 20160915 | 20160915 | 54 | accuracy | 1 |
| classification | transformer | Reaching C-J-M-T_1 | single_day | 20160915 | 20160915 | 69 | accuracy | 0.972973 |
| classification | transformer | Reaching C-J-M-T_1 | single_day | 20160919 | 20160919 | 31 | accuracy | 0.973684 |
| classification | transformer | Reaching C-J-M-T_1 | single_day | 20160919 | 20160919 | 44 | accuracy | 0.947368 |
| classification | transformer | Reaching C-J-M-T_1 | single_day | 20160919 | 20160919 | 46 | accuracy | 1 |
| classification | transformer | Reaching C-J-M-T_1 | single_day | 20160919 | 20160919 | 54 | accuracy | 0.973684 |
| classification | transformer | Reaching C-J-M-T_1 | single_day | 20160919 | 20160919 | 69 | accuracy | 0.947368 |
| classification | transformer | Reaching C-J-M-T_1 | single_day | 20160921 | 20160921 | 31 | accuracy | 0.972222 |
| classification | transformer | Reaching C-J-M-T_1 | single_day | 20160921 | 20160921 | 44 | accuracy | 0.944444 |
| classification | transformer | Reaching C-J-M-T_1 | single_day | 20160921 | 20160921 | 46 | accuracy | 0.888889 |
| classification | transformer | Reaching C-J-M-T_1 | single_day | 20160921 | 20160921 | 54 | accuracy | 0.972222 |
| classification | transformer | Reaching C-J-M-T_1 | single_day | 20160921 | 20160921 | 69 | accuracy | 0.916667 |
| classification | transformer | Reaching C-J-M-T_1 | single_day | 20160923 | 20160923 | 31 | accuracy | 1 |
| classification | transformer | Reaching C-J-M-T_1 | single_day | 20160923 | 20160923 | 44 | accuracy | 0.97561 |
| classification | transformer | Reaching C-J-M-T_1 | single_day | 20160923 | 20160923 | 46 | accuracy | 1 |
| classification | transformer | Reaching C-J-M-T_1 | single_day | 20160923 | 20160923 | 54 | accuracy | 0.97561 |
| classification | transformer | Reaching C-J-M-T_1 | single_day | 20160923 | 20160923 | 69 | accuracy | 0.926829 |
| classification | transformer | Reaching C-J-M-T_1 | single_day | 20160929 | 20160929 | 31 | accuracy | 0.952381 |
| classification | transformer | Reaching C-J-M-T_1 | single_day | 20160929 | 20160929 | 44 | accuracy | 0.952381 |
| classification | transformer | Reaching C-J-M-T_1 | single_day | 20160929 | 20160929 | 46 | accuracy | 1 |
| classification | transformer | Reaching C-J-M-T_1 | single_day | 20160929 | 20160929 | 54 | accuracy | 0.97619 |
| classification | transformer | Reaching C-J-M-T_1 | single_day | 20160929 | 20160929 | 69 | accuracy | 0.928571 |
| classification | transformer | Reaching C-J-M-T_1 | single_day | 20161005 | 20161005 | 31 | accuracy | 1 |
| classification | transformer | Reaching C-J-M-T_1 | single_day | 20161005 | 20161005 | 44 | accuracy | 0.95122 |
| classification | transformer | Reaching C-J-M-T_1 | single_day | 20161005 | 20161005 | 46 | accuracy | 1 |
| classification | transformer | Reaching C-J-M-T_1 | single_day | 20161005 | 20161005 | 54 | accuracy | 1 |
| classification | transformer | Reaching C-J-M-T_1 | single_day | 20161005 | 20161005 | 69 | accuracy | 0.97561 |
| classification | transformer | Reaching C-J-M-T_1 | single_day | 20161006 | 20161006 | 31 | accuracy | 0.97619 |
| classification | transformer | Reaching C-J-M-T_1 | single_day | 20161006 | 20161006 | 44 | accuracy | 0.928571 |
| classification | transformer | Reaching C-J-M-T_1 | single_day | 20161006 | 20161006 | 46 | accuracy | 0.97619 |
| classification | transformer | Reaching C-J-M-T_1 | single_day | 20161006 | 20161006 | 54 | accuracy | 0.952381 |
| classification | transformer | Reaching C-J-M-T_1 | single_day | 20161006 | 20161006 | 69 | accuracy | 0.97619 |
| classification | transformer | Reaching C-J-M-T_1 | single_day | 20161007 | 20161007 | 31 | accuracy | 0.828571 |
| classification | transformer | Reaching C-J-M-T_1 | single_day | 20161007 | 20161007 | 54 | accuracy | 0.971429 |
| classification | transformer | Reaching C-J-M-T_1 | single_day | 20161007 | 20161007 | 69 | accuracy | 1 |
| classification | transformer | Reaching C-J-M-T_1 | single_day | 20161011 | 20161011 | 31 | accuracy | 0.975 |
| classification | transformer | Reaching C-J-M-T_1 | single_day | 20161011 | 20161011 | 44 | accuracy | 0.925 |
| classification | transformer | Reaching C-J-M-T_1 | single_day | 20161011 | 20161011 | 46 | accuracy | 0.975 |
| classification | transformer | Reaching C-J-M-T_1 | single_day | 20161011 | 20161011 | 54 | accuracy | 0.95 |
| classification | transformer | Reaching C-J-M-T_1 | single_day | 20161011 | 20161011 | 69 | accuracy | 0.9 |
| classification | transformer | Reaching C-J-M-T_1 | single_day | 20161013 | 20161013 | 31 | accuracy | 0.958333 |
| classification | transformer | Reaching C-J-M-T_1 | single_day | 20161013 | 20161013 | 44 | accuracy | 0.9375 |
| classification | transformer | Reaching C-J-M-T_1 | single_day | 20161013 | 20161013 | 46 | accuracy | 0.9375 |
| classification | transformer | Reaching C-J-M-T_1 | single_day | 20161013 | 20161013 | 54 | accuracy | 0.958333 |

| | | | | | | | | |
|---|---|---|---|---|---|---|---|---|
| classification | transformer | Reaching C-J-M-T_1 | single_day | 20161013 | 20161013 | 69 | accuracy | 1 |
| classification | transformer | Reaching C-J-M-T_1 | single_day | 20161021 | 20161021 | 31 | accuracy | 0.982759 |
| classification | transformer | Reaching C-J-M-T_1 | single_day | 20161021 | 20161021 | 44 | accuracy | 1 |
| classification | transformer | Reaching C-J-M-T_1 | single_day | 20161021 | 20161021 | 46 | accuracy | 0.982759 |
| classification | transformer | Reaching C-J-M-T_1 | single_day | 20161021 | 20161021 | 54 | accuracy | 1 |
| classification | transformer | Reaching C-J-M-T_1 | single_day | 20161021 | 20161021 | 69 | accuracy | 1 |
| classification | lfads | Reaching C-J-M-T_1 | single_day | 20150629 | 20150629 | 44 | accuracy | 0.777778 |
| classification | lfads | Reaching C-J-M-T_1 | single_day | 20150629 | 20150629 | 46 | accuracy | 0.722222 |
| classification | lfads | Reaching C-J-M-T_1 | single_day | 20150629 | 20150629 | 54 | accuracy | 0.583333 |
| classification | lfads | Reaching C-J-M-T_1 | single_day | 20150630 | 20150630 | 44 | accuracy | 0.611111 |
| classification | lfads | Reaching C-J-M-T_1 | single_day | 20150630 | 20150630 | 46 | accuracy | 0.694444 |
| classification | lfads | Reaching C-J-M-T_1 | single_day | 20150630 | 20150630 | 54 | accuracy | 0.833333 |
| classification | lfads | Reaching C-J-M-T_1 | single_day | 20150701 | 20150701 | 44 | accuracy | 0.65 |
| classification | lfads | Reaching C-J-M-T_1 | single_day | 20150701 | 20150701 | 46 | accuracy | 0.75 |
| classification | lfads | Reaching C-J-M-T_1 | single_day | 20150701 | 20150701 | 54 | accuracy | 0.8 |
| classification | lfads | Reaching C-J-M-T_1 | single_day | 20150706 | 20150706 | 44 | accuracy | 0.648649 |
| classification | lfads | Reaching C-J-M-T_1 | single_day | 20150706 | 20150706 | 46 | accuracy | 0.621622 |
| classification | lfads | Reaching C-J-M-T_1 | single_day | 20150706 | 20150706 | 54 | accuracy | 0.675676 |
| classification | lfads | Reaching C-J-M-T_1 | single_day | 20150708 | 20150708 | 44 | accuracy | 0.894737 |
| classification | lfads | Reaching C-J-M-T_1 | single_day | 20150708 | 20150708 | 46 | accuracy | 0.736842 |
| classification | lfads | Reaching C-J-M-T_1 | single_day | 20150708 | 20150708 | 54 | accuracy | 0.736842 |
| classification | lfads | Reaching C-J-M-T_1 | single_day | 20150709 | 20150709 | 44 | accuracy | 0.7 |
| classification | lfads | Reaching C-J-M-T_1 | single_day | 20150709 | 20150709 | 46 | accuracy | 0.775 |
| classification | lfads | Reaching C-J-M-T_1 | single_day | 20150709 | 20150709 | 54 | accuracy | 0.75 |
| classification | lfads | Reaching C-J-M-T_1 | single_day | 20150710 | 20150710 | 44 | accuracy | 0.853659 |
| classification | lfads | Reaching C-J-M-T_1 | single_day | 20150710 | 20150710 | 46 | accuracy | 0.731707 |
| classification | lfads | Reaching C-J-M-T_1 | single_day | 20150710 | 20150710 | 54 | accuracy | 0.609756 |
| classification | cycle_gan | Reaching C-J-M-T_1 | single_day | 20131003 | 20131003 | 31 | accuracy | 0.625 |
| classification | cycle_gan | Reaching C-J-M-T_1 | single_day | 20131003 | 20131003 | 44 | accuracy | 0.625 |
| classification | cycle_gan | Reaching C-J-M-T_1 | single_day | 20131003 | 20131003 | 46 | accuracy | 0.25 |
| classification | cycle_gan | Reaching C-J-M-T_1 | single_day | 20131003 | 20131003 | 54 | accuracy | 0.5625 |
| classification | cycle_gan | Reaching C-J-M-T_1 | single_day | 20131003 | 20131003 | 69 | accuracy | 0.53125 |
| classification | cycle_gan | Reaching C-J-M-T_1 | single_day | 20131022 | 20131022 | 31 | accuracy | 0.677419 |
| classification | cycle_gan | Reaching C-J-M-T_1 | single_day | 20131022 | 20131022 | 44 | accuracy | 0.612903 |
| classification | cycle_gan | Reaching C-J-M-T_1 | single_day | 20131022 | 20131022 | 46 | accuracy | 0.774194 |
| classification | cycle_gan | Reaching C-J-M-T_1 | single_day | 20131022 | 20131022 | 54 | accuracy | 0.677419 |
| classification | cycle_gan | Reaching C-J-M-T_1 | single_day | 20131022 | 20131022 | 69 | accuracy | 0.548387 |
| classification | cycle_gan | Reaching C-J-M-T_1 | single_day | 20131023 | 20131023 | 31 | accuracy | 0.717949 |
| classification | cycle_gan | Reaching C-J-M-T_1 | single_day | 20131023 | 20131023 | 44 | accuracy | 0.615385 |
| classification | cycle_gan | Reaching C-J-M-T_1 | single_day | 20131023 | 20131023 | 46 | accuracy | 0.769231 |
| classification | cycle_gan | Reaching C-J-M-T_1 | single_day | 20131023 | 20131023 | 54 | accuracy | 0.615385 |
| classification | cycle_gan | Reaching C-J-M-T_1 | single_day | 20131023 | 20131023 | 69 | accuracy | 0.74359 |
| classification | cycle_gan | Reaching C-J-M-T_1 | single_day | 20131101 | 20131101 | 31 | accuracy | 0.7 |
| classification | cycle_gan | Reaching C-J-M-T_1 | single_day | 20131101 | 20131101 | 44 | accuracy | 0.54 |
| classification | cycle_gan | Reaching C-J-M-T_1 | single_day | 20131101 | 20131101 | 46 | accuracy | 0.68 |
| classification | cycle_gan | Reaching C-J-M-T_1 | single_day | 20131101 | 20131101 | 54 | accuracy | 0.7 |
| classification | cycle_gan | Reaching C-J-M-T_1 | single_day | 20131101 | 20131101 | 69 | accuracy | 0.56 |
| classification | cycle_gan | Reaching C-J-M-T_1 | single_day | 20131203 | 20131203 | 31 | accuracy | 0.705882 |
| classification | cycle_gan | Reaching C-J-M-T_1 | single_day | 20131203 | 20131203 | 44 | accuracy | 0.705882 |
| classification | cycle_gan | Reaching C-J-M-T_1 | single_day | 20131203 | 20131203 | 46 | accuracy | 0.735294 |
| classification | cycle_gan | Reaching C-J-M-T_1 | single_day | 20131203 | 20131203 | 54 | accuracy | 0.588235 |
| classification | cycle_gan | Reaching C-J-M-T_1 | single_day | 20131203 | 20131203 | 69 | accuracy | 0.647059 |
| classification | cycle_gan | Reaching C-J-M-T_1 | single_day | 20131204 | 20131204 | 31 | accuracy | 0.515152 |
| classification | cycle_gan | Reaching C-J-M-T_1 | single_day | 20131204 | 20131204 | 44 | accuracy | 0.69697 |
| classification | cycle_gan | Reaching C-J-M-T_1 | single_day | 20131204 | 20131204 | 46 | accuracy | 0.575758 |
| classification | cycle_gan | Reaching C-J-M-T_1 | single_day | 20131204 | 20131204 | 54 | accuracy | 0.545455 |
| classification | cycle_gan | Reaching C-J-M-T_1 | single_day | 20131204 | 20131204 | 69 | accuracy | 0.545455 |
| classification | cycle_gan | Reaching C-J-M-T_1 | single_day | 20131219 | 20131219 | 31 | accuracy | 0.388889 |
| classification | cycle_gan | Reaching C-J-M-T_1 | single_day | 20131219 | 20131219 | 44 | accuracy | 0.416667 |
| classification | cycle_gan | Reaching C-J-M-T_1 | single_day | 20131219 | 20131219 | 46 | accuracy | 0.583333 |
| classification | cycle_gan | Reaching C-J-M-T_1 | single_day | 20131219 | 20131219 | 54 | accuracy | 0.5 |
| classification | cycle_gan | Reaching C-J-M-T_1 | single_day | 20131219 | 20131219 | 69 | accuracy | 0.472222 |
| classification | cycle_gan | Reaching C-J-M-T_1 | single_day | 20131220 | 20131220 | 31 | accuracy | 0.575 |

| | | | | | | | | |
|---|---|---|---|---|---|---|---|---|
| classification | cycle_gan | Reaching C-J-M-T_1 | single_day | 20131220 | 20131220 | 44 | accuracy | 0.6 |
| classification | cycle_gan | Reaching C-J-M-T_1 | single_day | 20131220 | 20131220 | 46 | accuracy | 0.65 |
| classification | cycle_gan | Reaching C-J-M-T_1 | single_day | 20131220 | 20131220 | 54 | accuracy | 0.475 |
| classification | cycle_gan | Reaching C-J-M-T_1 | single_day | 20131220 | 20131220 | 69 | accuracy | 0.625 |
| classification | cycle_gan | Reaching C-J-M-T_1 | single_day | 20150311 | 20150311 | 31 | accuracy | 0.912568 |
| classification | cycle_gan | Reaching C-J-M-T_1 | single_day | 20150311 | 20150311 | 44 | accuracy | 0.874317 |
| classification | cycle_gan | Reaching C-J-M-T_1 | single_day | 20150311 | 20150311 | 46 | accuracy | 0.863388 |
| classification | cycle_gan | Reaching C-J-M-T_1 | single_day | 20150311 | 20150311 | 54 | accuracy | 0.896175 |
| classification | cycle_gan | Reaching C-J-M-T_1 | single_day | 20150311 | 20150311 | 69 | accuracy | 0.928962 |
| classification | cycle_gan | Reaching C-J-M-T_1 | single_day | 20150319 | 20150319 | 31 | accuracy | 0.912621 |
| classification | cycle_gan | Reaching C-J-M-T_1 | single_day | 20150319 | 20150319 | 44 | accuracy | 0.941748 |
| classification | cycle_gan | Reaching C-J-M-T_1 | single_day | 20150319 | 20150319 | 46 | accuracy | 0.854369 |
| classification | cycle_gan | Reaching C-J-M-T_1 | single_day | 20150319 | 20150319 | 54 | accuracy | 0.92233 |
| classification | cycle_gan | Reaching C-J-M-T_1 | single_day | 20150319 | 20150319 | 69 | accuracy | 0.898058 |
| classification | cycle_gan | Reaching C-J-M-T_1 | single_day | 20150629 | 20150629 | 31 | accuracy | 0.611111 |
| classification | cycle_gan | Reaching C-J-M-T_1 | single_day | 20150629 | 20150629 | 44 | accuracy | 0.75 |
| classification | cycle_gan | Reaching C-J-M-T_1 | single_day | 20150629 | 20150629 | 46 | accuracy | 0.638889 |
| classification | cycle_gan | Reaching C-J-M-T_1 | single_day | 20150629 | 20150629 | 54 | accuracy | 0.638889 |
| classification | cycle_gan | Reaching C-J-M-T_1 | single_day | 20150629 | 20150629 | 69 | accuracy | 0.583333 |
| classification | cycle_gan | Reaching C-J-M-T_1 | single_day | 20150630 | 20150630 | 31 | accuracy | 0.861111 |
| classification | cycle_gan | Reaching C-J-M-T_1 | single_day | 20150630 | 20150630 | 44 | accuracy | 0.694444 |
| classification | cycle_gan | Reaching C-J-M-T_1 | single_day | 20150630 | 20150630 | 46 | accuracy | 0.777778 |
| classification | cycle_gan | Reaching C-J-M-T_1 | single_day | 20150630 | 20150630 | 54 | accuracy | 0.888889 |
| classification | cycle_gan | Reaching C-J-M-T_1 | single_day | 20150630 | 20150630 | 69 | accuracy | 0.75 |
| classification | cycle_gan | Reaching C-J-M-T_1 | single_day | 20150701 | 20150701 | 31 | accuracy | 0.65 |
| classification | cycle_gan | Reaching C-J-M-T_1 | single_day | 20150701 | 20150701 | 44 | accuracy | 0.725 |
| classification | cycle_gan | Reaching C-J-M-T_1 | single_day | 20150701 | 20150701 | 46 | accuracy | 0.75 |
| classification | cycle_gan | Reaching C-J-M-T_1 | single_day | 20150701 | 20150701 | 54 | accuracy | 0.7 |
| classification | cycle_gan | Reaching C-J-M-T_1 | single_day | 20150701 | 20150701 | 69 | accuracy | 0.7 |
| classification | cycle_gan | Reaching C-J-M-T_1 | single_day | 20150706 | 20150706 | 31 | accuracy | 0.756757 |
| classification | cycle_gan | Reaching C-J-M-T_1 | single_day | 20150706 | 20150706 | 44 | accuracy | 0.72973 |
| classification | cycle_gan | Reaching C-J-M-T_1 | single_day | 20150706 | 20150706 | 46 | accuracy | 0.783784 |
| classification | cycle_gan | Reaching C-J-M-T_1 | single_day | 20150706 | 20150706 | 54 | accuracy | 0.540541 |
| classification | cycle_gan | Reaching C-J-M-T_1 | single_day | 20150706 | 20150706 | 69 | accuracy | 0.513514 |
| classification | cycle_gan | Reaching C-J-M-T_1 | single_day | 20150708 | 20150708 | 31 | accuracy | 0.815789 |
| classification | cycle_gan | Reaching C-J-M-T_1 | single_day | 20150708 | 20150708 | 44 | accuracy | 0.736842 |
| classification | cycle_gan | Reaching C-J-M-T_1 | single_day | 20150708 | 20150708 | 46 | accuracy | 0.605263 |
| classification | cycle_gan | Reaching C-J-M-T_1 | single_day | 20150708 | 20150708 | 54 | accuracy | 0.657895 |
| classification | cycle_gan | Reaching C-J-M-T_1 | single_day | 20150708 | 20150708 | 69 | accuracy | 0.842105 |
| classification | cycle_gan | Reaching C-J-M-T_1 | single_day | 20150709 | 20150709 | 31 | accuracy | 0.85 |
| classification | cycle_gan | Reaching C-J-M-T_1 | single_day | 20150709 | 20150709 | 44 | accuracy | 0.875 |
| classification | cycle_gan | Reaching C-J-M-T_1 | single_day | 20150709 | 20150709 | 46 | accuracy | 0.625 |
| classification | cycle_gan | Reaching C-J-M-T_1 | single_day | 20150709 | 20150709 | 54 | accuracy | 0.85 |
| classification | cycle_gan | Reaching C-J-M-T_1 | single_day | 20150709 | 20150709 | 69 | accuracy | 0.7 |
| classification | cycle_gan | Reaching C-J-M-T_1 | single_day | 20150710 | 20150710 | 31 | accuracy | 0.682927 |
| classification | cycle_gan | Reaching C-J-M-T_1 | single_day | 20150710 | 20150710 | 44 | accuracy | 0.707317 |
| classification | cycle_gan | Reaching C-J-M-T_1 | single_day | 20150710 | 20150710 | 46 | accuracy | 0.682927 |
| classification | cycle_gan | Reaching C-J-M-T_1 | single_day | 20150710 | 20150710 | 54 | accuracy | 0.658537 |
| classification | cycle_gan | Reaching C-J-M-T_1 | single_day | 20150710 | 20150710 | 69 | accuracy | 0.731707 |
| classification | cycle_gan | Reaching C-J-M-T_1 | single_day | 20150713 | 20150713 | 31 | accuracy | 0.810811 |
| classification | cycle_gan | Reaching C-J-M-T_1 | single_day | 20150713 | 20150713 | 44 | accuracy | 0.918919 |
| classification | cycle_gan | Reaching C-J-M-T_1 | single_day | 20150713 | 20150713 | 46 | accuracy | 0.72973 |
| classification | cycle_gan | Reaching C-J-M-T_1 | single_day | 20150713 | 20150713 | 54 | accuracy | 0.864865 |
| classification | cycle_gan | Reaching C-J-M-T_1 | single_day | 20150713 | 20150713 | 69 | accuracy | 0.702703 |
| classification | cycle_gan | Reaching C-J-M-T_1 | single_day | 20150714 | 20150714 | 31 | accuracy | 0.8 |
| classification | cycle_gan | Reaching C-J-M-T_1 | single_day | 20150714 | 20150714 | 44 | accuracy | 0.9 |
| classification | cycle_gan | Reaching C-J-M-T_1 | single_day | 20150714 | 20150714 | 46 | accuracy | 0.825 |
| classification | cycle_gan | Reaching C-J-M-T_1 | single_day | 20150714 | 20150714 | 54 | accuracy | 0.675 |
| classification | cycle_gan | Reaching C-J-M-T_1 | single_day | 20150714 | 20150714 | 69 | accuracy | 0.825 |
| classification | cycle_gan | Reaching C-J-M-T_1 | single_day | 20150715 | 20150715 | 31 | accuracy | 0.666667 |
| classification | cycle_gan | Reaching C-J-M-T_1 | single_day | 20150715 | 20150715 | 44 | accuracy | 0.717949 |
| classification | cycle_gan | Reaching C-J-M-T_1 | single_day | 20150715 | 20150715 | 46 | accuracy | 0.846154 |
| classification | cycle_gan | Reaching C-J-M-T_1 | single_day | 20150715 | 20150715 | 54 | accuracy | 0.820513 |

| | | | | | | | | |
|---|---|---|---|---|---|---|---|---|
| classification | cycle_gan | Reaching C-J-M-T_1 | single_day | 20150715 | 20150715 | 69 | accuracy | 0.794872 |
| classification | cycle_gan | Reaching C-J-M-T_1 | single_day | 20150716 | 20150716 | 31 | accuracy | 0.769231 |
| classification | cycle_gan | Reaching C-J-M-T_1 | single_day | 20150716 | 20150716 | 44 | accuracy | 0.794872 |
| classification | cycle_gan | Reaching C-J-M-T_1 | single_day | 20150716 | 20150716 | 46 | accuracy | 0.589744 |
| classification | cycle_gan | Reaching C-J-M-T_1 | single_day | 20150716 | 20150716 | 54 | accuracy | 0.717949 |
| classification | cycle_gan | Reaching C-J-M-T_1 | single_day | 20150716 | 20150716 | 69 | accuracy | 0.692308 |
| classification | cycle_gan | Reaching C-J-M-T_1 | single_day | 20151103 | 20151103 | 31 | accuracy | 0.6 |
| classification | cycle_gan | Reaching C-J-M-T_1 | single_day | 20151103 | 20151103 | 44 | accuracy | 0.64 |
| classification | cycle_gan | Reaching C-J-M-T_1 | single_day | 20151103 | 20151103 | 46 | accuracy | 0.76 |
| classification | cycle_gan | Reaching C-J-M-T_1 | single_day | 20151103 | 20151103 | 54 | accuracy | 0.64 |
| classification | cycle_gan | Reaching C-J-M-T_1 | single_day | 20151103 | 20151103 | 69 | accuracy | 0.72 |
| classification | cycle_gan | Reaching C-J-M-T_1 | single_day | 20151106 | 20151106 | 31 | accuracy | 0.75 |
| classification | cycle_gan | Reaching C-J-M-T_1 | single_day | 20151106 | 20151106 | 44 | accuracy | 0.442308 |
| classification | cycle_gan | Reaching C-J-M-T_1 | single_day | 20151106 | 20151106 | 46 | accuracy | 0.75 |
| classification | cycle_gan | Reaching C-J-M-T_1 | single_day | 20151106 | 20151106 | 54 | accuracy | 0.615385 |
| classification | cycle_gan | Reaching C-J-M-T_1 | single_day | 20151106 | 20151106 | 69 | accuracy | 0.769231 |
| classification | cycle_gan | Reaching C-J-M-T_1 | single_day | 20151112 | 20151112 | 31 | accuracy | 0.631579 |
| classification | cycle_gan | Reaching C-J-M-T_1 | single_day | 20151112 | 20151112 | 44 | accuracy | 0.815789 |
| classification | cycle_gan | Reaching C-J-M-T_1 | single_day | 20151112 | 20151112 | 46 | accuracy | 0.789474 |
| classification | cycle_gan | Reaching C-J-M-T_1 | single_day | 20151112 | 20151112 | 54 | accuracy | 0.710526 |
| classification | cycle_gan | Reaching C-J-M-T_1 | single_day | 20151112 | 20151112 | 69 | accuracy | 0.605263 |
| classification | cycle_gan | Reaching C-J-M-T_1 | single_day | 20151113 | 20151113 | 31 | accuracy | 0.789474 |
| classification | cycle_gan | Reaching C-J-M-T_1 | single_day | 20151113 | 20151113 | 44 | accuracy | 0.684211 |
| classification | cycle_gan | Reaching C-J-M-T_1 | single_day | 20151113 | 20151113 | 46 | accuracy | 0.815789 |
| classification | cycle_gan | Reaching C-J-M-T_1 | single_day | 20151113 | 20151113 | 54 | accuracy | 0.763158 |
| classification | cycle_gan | Reaching C-J-M-T_1 | single_day | 20151113 | 20151113 | 69 | accuracy | 0.657895 |
| classification | cycle_gan | Reaching C-J-M-T_1 | single_day | 20151116 | 20151116 | 31 | accuracy | 0.735294 |
| classification | cycle_gan | Reaching C-J-M-T_1 | single_day | 20151116 | 20151116 | 44 | accuracy | 0.705882 |
| classification | cycle_gan | Reaching C-J-M-T_1 | single_day | 20151116 | 20151116 | 46 | accuracy | 0.676471 |
| classification | cycle_gan | Reaching C-J-M-T_1 | single_day | 20151116 | 20151116 | 54 | accuracy | 0.588235 |
| classification | cycle_gan | Reaching C-J-M-T_1 | single_day | 20151116 | 20151116 | 69 | accuracy | 0.647059 |
| classification | cycle_gan | Reaching C-J-M-T_1 | single_day | 20151119 | 20151119 | 31 | accuracy | 0.5 |
| classification | cycle_gan | Reaching C-J-M-T_1 | single_day | 20151119 | 20151119 | 44 | accuracy | 0.571429 |
| classification | cycle_gan | Reaching C-J-M-T_1 | single_day | 20151119 | 20151119 | 46 | accuracy | 0.857143 |
| classification | cycle_gan | Reaching C-J-M-T_1 | single_day | 20151119 | 20151119 | 54 | accuracy | 0.52381 |
| classification | cycle_gan | Reaching C-J-M-T_1 | single_day | 20151119 | 20151119 | 69 | accuracy | 0.619048 |
| classification | cycle_gan | Reaching C-J-M-T_1 | single_day | 20151120 | 20151120 | 31 | accuracy | 0.764706 |
| classification | cycle_gan | Reaching C-J-M-T_1 | single_day | 20151120 | 20151120 | 44 | accuracy | 0.735294 |
| classification | cycle_gan | Reaching C-J-M-T_1 | single_day | 20151120 | 20151120 | 46 | accuracy | 0.647059 |
| classification | cycle_gan | Reaching C-J-M-T_1 | single_day | 20151120 | 20151120 | 54 | accuracy | 0.764706 |
| classification | cycle_gan | Reaching C-J-M-T_1 | single_day | 20151120 | 20151120 | 69 | accuracy | 0.705882 |
| classification | cycle_gan | Reaching C-J-M-T_1 | single_day | 20151201 | 20151201 | 31 | accuracy | 0.684211 |
| classification | cycle_gan | Reaching C-J-M-T_1 | single_day | 20151201 | 20151201 | 44 | accuracy | 0.684211 |
| classification | cycle_gan | Reaching C-J-M-T_1 | single_day | 20151201 | 20151201 | 46 | accuracy | 0.789474 |
| classification | cycle_gan | Reaching C-J-M-T_1 | single_day | 20151201 | 20151201 | 54 | accuracy | 0.763158 |
| classification | cycle_gan | Reaching C-J-M-T_1 | single_day | 20151201 | 20151201 | 69 | accuracy | 0.657895 |
| classification | cycle_gan | Reaching C-J-M-T_1 | single_day | 20160914 | 20160914 | 31 | accuracy | 0.615385 |
| classification | cycle_gan | Reaching C-J-M-T_1 | single_day | 20160914 | 20160914 | 44 | accuracy | 0.692308 |
| classification | cycle_gan | Reaching C-J-M-T_1 | single_day | 20160914 | 20160914 | 46 | accuracy | 0.769231 |
| classification | cycle_gan | Reaching C-J-M-T_1 | single_day | 20160914 | 20160914 | 54 | accuracy | 0.615385 |
| classification | cycle_gan | Reaching C-J-M-T_1 | single_day | 20160914 | 20160914 | 69 | accuracy | 0.666667 |
| classification | cycle_gan | Reaching C-J-M-T_1 | single_day | 20160915 | 20160915 | 31 | accuracy | 0.675676 |
| classification | cycle_gan | Reaching C-J-M-T_1 | single_day | 20160915 | 20160915 | 44 | accuracy | 0.783784 |
| classification | cycle_gan | Reaching C-J-M-T_1 | single_day | 20160915 | 20160915 | 46 | accuracy | 0.567568 |
| classification | cycle_gan | Reaching C-J-M-T_1 | single_day | 20160915 | 20160915 | 54 | accuracy | 0.675676 |
| classification | cycle_gan | Reaching C-J-M-T_1 | single_day | 20160915 | 20160915 | 69 | accuracy | 0.810811 |
| classification | cycle_gan | Reaching C-J-M-T_1 | single_day | 20160919 | 20160919 | 31 | accuracy | 0.815789 |
| classification | cycle_gan | Reaching C-J-M-T_1 | single_day | 20160919 | 20160919 | 44 | accuracy | 0.631579 |
| classification | cycle_gan | Reaching C-J-M-T_1 | single_day | 20160919 | 20160919 | 46 | accuracy | 0.763158 |
| classification | cycle_gan | Reaching C-J-M-T_1 | single_day | 20160919 | 20160919 | 54 | accuracy | 0.684211 |
| classification | cycle_gan | Reaching C-J-M-T_1 | single_day | 20160919 | 20160919 | 69 | accuracy | 0.763158 |
| classification | cycle_gan | Reaching C-J-M-T_1 | single_day | 20160921 | 20160921 | 31 | accuracy | 0.694444 |
| classification | cycle_gan | Reaching C-J-M-T_1 | single_day | 20160921 | 20160921 | 44 | accuracy | 0.694444 |

| | | | | | | | | |
|---|---|---|---|---|---|---|---|---|
| classification | cycle_gan | Reaching C-J-M-T_1 | single_day | 20160921 | 20160921 | 46 | accuracy | 0.694444 |
| classification | cycle_gan | Reaching C-J-M-T_1 | single_day | 20160921 | 20160921 | 54 | accuracy | 0.694444 |
| classification | cycle_gan | Reaching C-J-M-T_1 | single_day | 20160921 | 20160921 | 69 | accuracy | 0.805556 |
| classification | cycle_gan | Reaching C-J-M-T_1 | single_day | 20160923 | 20160923 | 31 | accuracy | 0.853659 |
| classification | cycle_gan | Reaching C-J-M-T_1 | single_day | 20160923 | 20160923 | 44 | accuracy | 0.829268 |
| classification | cycle_gan | Reaching C-J-M-T_1 | single_day | 20160923 | 20160923 | 46 | accuracy | 0.756098 |
| classification | cycle_gan | Reaching C-J-M-T_1 | single_day | 20160923 | 20160923 | 54 | accuracy | 0.682927 |
| classification | cycle_gan | Reaching C-J-M-T_1 | single_day | 20160923 | 20160923 | 69 | accuracy | 0.634146 |
| classification | cycle_gan | Reaching C-J-M-T_1 | single_day | 20160929 | 20160929 | 31 | accuracy | 0.833333 |
| classification | cycle_gan | Reaching C-J-M-T_1 | single_day | 20160929 | 20160929 | 44 | accuracy | 0.642857 |
| classification | cycle_gan | Reaching C-J-M-T_1 | single_day | 20160929 | 20160929 | 46 | accuracy | 0.690476 |
| classification | cycle_gan | Reaching C-J-M-T_1 | single_day | 20160929 | 20160929 | 54 | accuracy | 0.761905 |
| classification | cycle_gan | Reaching C-J-M-T_1 | single_day | 20160929 | 20160929 | 69 | accuracy | 0.666667 |
| classification | cycle_gan | Reaching C-J-M-T_1 | single_day | 20161005 | 20161005 | 31 | accuracy | 0.560976 |
| classification | cycle_gan | Reaching C-J-M-T_1 | single_day | 20161005 | 20161005 | 44 | accuracy | 0.804878 |
| classification | cycle_gan | Reaching C-J-M-T_1 | single_day | 20161005 | 20161005 | 46 | accuracy | 0.658537 |
| classification | cycle_gan | Reaching C-J-M-T_1 | single_day | 20161005 | 20161005 | 54 | accuracy | 0.707317 |
| classification | cycle_gan | Reaching C-J-M-T_1 | single_day | 20161005 | 20161005 | 69 | accuracy | 0.780488 |
| classification | cycle_gan | Reaching C-J-M-T_1 | single_day | 20161011 | 20161011 | 31 | accuracy | 0.725 |
| classification | cycle_gan | Reaching C-J-M-T_1 | single_day | 20161011 | 20161011 | 44 | accuracy | 0.725 |
| classification | cycle_gan | Reaching C-J-M-T_1 | single_day | 20161011 | 20161011 | 46 | accuracy | 0.75 |
| classification | cycle_gan | Reaching C-J-M-T_1 | single_day | 20161011 | 20161011 | 54 | accuracy | 0.7 |
| classification | cycle_gan | Reaching C-J-M-T_1 | single_day | 20161011 | 20161011 | 69 | accuracy | 0.6 |
| classification | cycle_gan | Reaching C-J-M-T_1 | single_day | 20161013 | 20161013 | 31 | accuracy | 0.75 |
| classification | cycle_gan | Reaching C-J-M-T_1 | single_day | 20161013 | 20161013 | 44 | accuracy | 0.75 |
| classification | cycle_gan | Reaching C-J-M-T_1 | single_day | 20161013 | 20161013 | 46 | accuracy | 0.791667 |
| classification | cycle_gan | Reaching C-J-M-T_1 | single_day | 20161013 | 20161013 | 54 | accuracy | 0.833333 |
| classification | cycle_gan | Reaching C-J-M-T_1 | single_day | 20161013 | 20161013 | 69 | accuracy | 0.895833 |
| classification | cycle_gan | Reaching C-J-M-T_1 | single_day | 20161021 | 20161021 | 31 | accuracy | 0.87931 |
| classification | cycle_gan | Reaching C-J-M-T_1 | single_day | 20161021 | 20161021 | 44 | accuracy | 0.775862 |
| classification | cycle_gan | Reaching C-J-M-T_1 | single_day | 20161021 | 20161021 | 46 | accuracy | 0.758621 |
| classification | cycle_gan | Reaching C-J-M-T_1 | single_day | 20161021 | 20161021 | 54 | accuracy | 0.775862 |
| classification | cycle_gan | Reaching C-J-M-T_1 | single_day | 20161021 | 20161021 | 69 | accuracy | 0.844828 |
| classification | stabilization | Reaching C-J-M-T_1 | single_day | 20150629 | 20150629 | 44 | accuracy | 0.580645 |
| classification | stabilization | Reaching C-J-M-T_1 | single_day | 20150629 | 20150629 | 46 | accuracy | 0.580645 |
| classification | stabilization | Reaching C-J-M-T_1 | single_day | 20150629 | 20150629 | 54 | accuracy | 0.419355 |
| classification | stabilization | Reaching C-J-M-T_1 | single_day | 20150630 | 20150630 | 44 | accuracy | 0.580645 |
| classification | stabilization | Reaching C-J-M-T_1 | single_day | 20150630 | 20150630 | 46 | accuracy | 0.548387 |
| classification | stabilization | Reaching C-J-M-T_1 | single_day | 20150630 | 20150630 | 54 | accuracy | 0.483871 |
| classification | stabilization | Reaching C-J-M-T_1 | single_day | 20150701 | 20150701 | 44 | accuracy | 0.628571 |
| classification | stabilization | Reaching C-J-M-T_1 | single_day | 20150701 | 20150701 | 46 | accuracy | 0.657143 |
| classification | stabilization | Reaching C-J-M-T_1 | single_day | 20150701 | 20150701 | 54 | accuracy | 0.657143 |
| classification | stabilization | Reaching C-J-M-T_1 | single_day | 20150706 | 20150706 | 44 | accuracy | 0.5625 |
| classification | stabilization | Reaching C-J-M-T_1 | single_day | 20150706 | 20150706 | 46 | accuracy | 0.5625 |
| classification | stabilization | Reaching C-J-M-T_1 | single_day | 20150706 | 20150706 | 54 | accuracy | 0.46875 |
| classification | stabilization | Reaching C-J-M-T_1 | single_day | 20150708 | 20150708 | 44 | accuracy | 0.545455 |
| classification | stabilization | Reaching C-J-M-T_1 | single_day | 20150708 | 20150708 | 46 | accuracy | 0.757576 |
| classification | stabilization | Reaching C-J-M-T_1 | single_day | 20150708 | 20150708 | 54 | accuracy | 0.606061 |
| classification | stabilization | Reaching C-J-M-T_1 | single_day | 20150709 | 20150709 | 44 | accuracy | 0.6 |
| classification | stabilization | Reaching C-J-M-T_1 | single_day | 20150709 | 20150709 | 46 | accuracy | 0.742857 |
| classification | stabilization | Reaching C-J-M-T_1 | single_day | 20150709 | 20150709 | 54 | accuracy | 0.485714 |
| classification | stabilization | Reaching C-J-M-T_1 | single_day | 20150710 | 20150710 | 44 | accuracy | 0.583333 |
| classification | stabilization | Reaching C-J-M-T_1 | single_day | 20150710 | 20150710 | 46 | accuracy | 0.583333 |
| classification | stabilization | Reaching C-J-M-T_1 | single_day | 20150710 | 20150710 | 54 | accuracy | 0.527778 |
| classification | mfsnn | Reaching C-J-M-T_1 | single_day | 20131003 | 20131003 | 44 | accuracy | 0.375 |
| classification | mfsnn | Reaching C-J-M-T_1 | single_day | 20131003 | 20131003 | 46 | accuracy | 0.21875 |
| classification | mfsnn | Reaching C-J-M-T_1 | single_day | 20131003 | 20131003 | 54 | accuracy | 0.125 |
| classification | mfsnn | Reaching C-J-M-T_1 | single_day | 20131022 | 20131022 | 44 | accuracy | 0.225806 |
| classification | mfsnn | Reaching C-J-M-T_1 | single_day | 20131022 | 20131022 | 46 | accuracy | 0.193548 |
| classification | mfsnn | Reaching C-J-M-T_1 | single_day | 20131022 | 20131022 | 54 | accuracy | 0.129032 |
| classification | mfsnn | Reaching C-J-M-T_1 | single_day | 20131023 | 20131023 | 44 | accuracy | 0.333333 |
| classification | mfsnn | Reaching C-J-M-T_1 | single_day | 20131023 | 20131023 | 46 | accuracy | 0.435897 |
| classification | mfsnn | Reaching C-J-M-T_1 | single_day | 20131023 | 20131023 | 54 | accuracy | 0.487179 |

| | | | | | | | | |
|---|---|---|---|---|---|---|---|---|
| classification | mfsnn | Reaching C-J-M-T_1 | single_day | 20131101 | 20131101 | 44 | accuracy | 0.14 |
| classification | mfsnn | Reaching C-J-M-T_1 | single_day | 20131101 | 20131101 | 46 | accuracy | 0.14 |
| classification | mfsnn | Reaching C-J-M-T_1 | single_day | 20131101 | 20131101 | 54 | accuracy | 0.52 |
| classification | mfsnn | Reaching C-J-M-T_1 | single_day | 20131203 | 20131203 | 44 | accuracy | 0.117647 |
| classification | mfsnn | Reaching C-J-M-T_1 | single_day | 20131203 | 20131203 | 46 | accuracy | 0.117647 |
| classification | mfsnn | Reaching C-J-M-T_1 | single_day | 20131203 | 20131203 | 54 | accuracy | 0.147059 |
| classification | mfsnn | Reaching C-J-M-T_1 | single_day | 20131219 | 20131219 | 44 | accuracy | 0.138889 |
| classification | mfsnn | Reaching C-J-M-T_1 | single_day | 20131219 | 20131219 | 46 | accuracy | 0.138889 |
| classification | mfsnn | Reaching C-J-M-T_1 | single_day | 20131219 | 20131219 | 54 | accuracy | 0.222222 |
| classification | mfsnn | Reaching C-J-M-T_1 | single_day | 20131220 | 20131220 | 44 | accuracy | 0.075 |
| classification | mfsnn | Reaching C-J-M-T_1 | single_day | 20131220 | 20131220 | 46 | accuracy | 0.15 |
| classification | mfsnn | Reaching C-J-M-T_1 | single_day | 20131220 | 20131220 | 54 | accuracy | 0.2 |
| classification | mfsnn | Reaching C-J-M-T_1 | single_day | 20150311 | 20150311 | 44 | accuracy | 0.142077 |
| classification | mfsnn | Reaching C-J-M-T_1 | single_day | 20150311 | 20150311 | 46 | accuracy | 0.300546 |
| classification | mfsnn | Reaching C-J-M-T_1 | single_day | 20150311 | 20150311 | 54 | accuracy | 0.131148 |
| classification | mfsnn | Reaching C-J-M-T_1 | single_day | 20150312 | 20150312 | 44 | accuracy | 0.137363 |
| classification | mfsnn | Reaching C-J-M-T_1 | single_day | 20150312 | 20150312 | 46 | accuracy | 0.131868 |
| classification | mfsnn | Reaching C-J-M-T_1 | single_day | 20150312 | 20150312 | 54 | accuracy | 0.208791 |
| classification | mfsnn | Reaching C-J-M-T_1 | single_day | 20150313 | 20150313 | 44 | accuracy | 0.235577 |
| classification | mfsnn | Reaching C-J-M-T_1 | single_day | 20150313 | 20150313 | 46 | accuracy | 0.293269 |
| classification | mfsnn | Reaching C-J-M-T_1 | single_day | 20150313 | 20150313 | 54 | accuracy | 0.384615 |
| classification | mfsnn | Reaching C-J-M-T_1 | single_day | 20150319 | 20150319 | 44 | accuracy | 0.203883 |
| classification | mfsnn | Reaching C-J-M-T_1 | single_day | 20150319 | 20150319 | 46 | accuracy | 0.315534 |
| classification | mfsnn | Reaching C-J-M-T_1 | single_day | 20150319 | 20150319 | 54 | accuracy | 0.15534 |
| classification | mfsnn | Reaching C-J-M-T_1 | single_day | 20150629 | 20150629 | 44 | accuracy | 0.194444 |
| classification | mfsnn | Reaching C-J-M-T_1 | single_day | 20150629 | 20150629 | 46 | accuracy | 0.361111 |
| classification | mfsnn | Reaching C-J-M-T_1 | single_day | 20150629 | 20150629 | 54 | accuracy | 0.166667 |
| classification | mfsnn | Reaching C-J-M-T_1 | single_day | 20150630 | 20150630 | 44 | accuracy | 0.5 |
| classification | mfsnn | Reaching C-J-M-T_1 | single_day | 20150630 | 20150630 | 46 | accuracy | 0.472222 |
| classification | mfsnn | Reaching C-J-M-T_1 | single_day | 20150630 | 20150630 | 54 | accuracy | 0.361111 |
| classification | mfsnn | Reaching C-J-M-T_1 | single_day | 20150701 | 20150701 | 44 | accuracy | 0.125 |
| classification | mfsnn | Reaching C-J-M-T_1 | single_day | 20150701 | 20150701 | 46 | accuracy | 0.225 |
| classification | mfsnn | Reaching C-J-M-T_1 | single_day | 20150701 | 20150701 | 54 | accuracy | 0.15 |
| classification | mfsnn | Reaching C-J-M-T_1 | single_day | 20150706 | 20150706 | 44 | accuracy | 0.135135 |
| classification | mfsnn | Reaching C-J-M-T_1 | single_day | 20150706 | 20150706 | 46 | accuracy | 0.351351 |
| classification | mfsnn | Reaching C-J-M-T_1 | single_day | 20150706 | 20150706 | 54 | accuracy | 0.27027 |
| classification | mfsnn | Reaching C-J-M-T_1 | single_day | 20150708 | 20150708 | 44 | accuracy | 0.315789 |
| classification | mfsnn | Reaching C-J-M-T_1 | single_day | 20150708 | 20150708 | 46 | accuracy | 0.315789 |
| classification | mfsnn | Reaching C-J-M-T_1 | single_day | 20150708 | 20150708 | 54 | accuracy | 0.394737 |
| classification | mfsnn | Reaching C-J-M-T_1 | single_day | 20150709 | 20150709 | 44 | accuracy | 0.6 |
| classification | mfsnn | Reaching C-J-M-T_1 | single_day | 20150709 | 20150709 | 46 | accuracy | 0.525 |
| classification | mfsnn | Reaching C-J-M-T_1 | single_day | 20150709 | 20150709 | 54 | accuracy | 0.675 |
| classification | mfsnn | Reaching C-J-M-T_1 | single_day | 20150713 | 20150713 | 44 | accuracy | 0.162162 |
| classification | mfsnn | Reaching C-J-M-T_1 | single_day | 20150713 | 20150713 | 46 | accuracy | 0.27027 |
| classification | mfsnn | Reaching C-J-M-T_1 | single_day | 20150713 | 20150713 | 54 | accuracy | 0.243243 |
| classification | mfsnn | Reaching C-J-M-T_1 | single_day | 20150714 | 20150714 | 44 | accuracy | 0.15 |
| classification | mfsnn | Reaching C-J-M-T_1 | single_day | 20150714 | 20150714 | 46 | accuracy | 0.275 |
| classification | mfsnn | Reaching C-J-M-T_1 | single_day | 20150714 | 20150714 | 54 | accuracy | 0.375 |
| classification | mfsnn | Reaching C-J-M-T_1 | single_day | 20150715 | 20150715 | 44 | accuracy | 0.230769 |
| classification | mfsnn | Reaching C-J-M-T_1 | single_day | 20150715 | 20150715 | 46 | accuracy | 0.333333 |
| classification | mfsnn | Reaching C-J-M-T_1 | single_day | 20150715 | 20150715 | 54 | accuracy | 0.282051 |
| classification | mfsnn | Reaching C-J-M-T_1 | single_day | 20150716 | 20150716 | 44 | accuracy | 0.384615 |
| classification | mfsnn | Reaching C-J-M-T_1 | single_day | 20150716 | 20150716 | 46 | accuracy | 0.179487 |
| classification | mfsnn | Reaching C-J-M-T_1 | single_day | 20150716 | 20150716 | 54 | accuracy | 0.128205 |
| classification | mfsnn | Reaching C-J-M-T_1 | single_day | 20151103 | 20151103 | 44 | accuracy | 0.2 |
| classification | mfsnn | Reaching C-J-M-T_1 | single_day | 20151103 | 20151103 | 46 | accuracy | 0.1 |
| classification | mfsnn | Reaching C-J-M-T_1 | single_day | 20151103 | 20151103 | 54 | accuracy | 0.2 |
| classification | mfsnn | Reaching C-J-M-T_1 | single_day | 20151106 | 20151106 | 44 | accuracy | 0.115385 |
| classification | mfsnn | Reaching C-J-M-T_1 | single_day | 20151106 | 20151106 | 46 | accuracy | 0.0961538 |
| classification | mfsnn | Reaching C-J-M-T_1 | single_day | 20151106 | 20151106 | 54 | accuracy | 0.115385 |
| classification | mfsnn | Reaching C-J-M-T_1 | single_day | 20151109 | 20151109 | 44 | accuracy | 0.2 |
| classification | mfsnn | Reaching C-J-M-T_1 | single_day | 20151109 | 20151109 | 46 | accuracy | 0.2 |
| classification | mfsnn | Reaching C-J-M-T_1 | single_day | 20151109 | 20151109 | 54 | accuracy | 0.142857 |

| | | | | | | | | |
|---|---|---|---|---|---|---|---|---|
| classification | mfsnn | Reaching C-J-M-T_1 | single_day | 20151112 | 20151112 | 44 | accuracy | 0.289474 |
| classification | mfsnn | Reaching C-J-M-T_1 | single_day | 20151112 | 20151112 | 46 | accuracy | 0.631579 |
| classification | mfsnn | Reaching C-J-M-T_1 | single_day | 20151112 | 20151112 | 54 | accuracy | 0.263158 |
| classification | mfsnn | Reaching C-J-M-T_1 | single_day | 20151113 | 20151113 | 44 | accuracy | 0.368421 |
| classification | mfsnn | Reaching C-J-M-T_1 | single_day | 20151113 | 20151113 | 46 | accuracy | 0.394737 |
| classification | mfsnn | Reaching C-J-M-T_1 | single_day | 20151113 | 20151113 | 54 | accuracy | 0.131579 |
| classification | mfsnn | Reaching C-J-M-T_1 | single_day | 20151116 | 20151116 | 44 | accuracy | 0.294118 |
| classification | mfsnn | Reaching C-J-M-T_1 | single_day | 20151116 | 20151116 | 46 | accuracy | 0.117647 |
| classification | mfsnn | Reaching C-J-M-T_1 | single_day | 20151116 | 20151116 | 54 | accuracy | 0.205882 |
| classification | mfsnn | Reaching C-J-M-T_1 | single_day | 20151119 | 20151119 | 44 | accuracy | 0.238095 |
| classification | mfsnn | Reaching C-J-M-T_1 | single_day | 20151119 | 20151119 | 46 | accuracy | 0.5 |
| classification | mfsnn | Reaching C-J-M-T_1 | single_day | 20151119 | 20151119 | 54 | accuracy | 0.761905 |
| classification | mfsnn | Reaching C-J-M-T_1 | single_day | 20151120 | 20151120 | 44 | accuracy | 0.176471 |
| classification | mfsnn | Reaching C-J-M-T_1 | single_day | 20151120 | 20151120 | 46 | accuracy | 0.176471 |
| classification | mfsnn | Reaching C-J-M-T_1 | single_day | 20151120 | 20151120 | 54 | accuracy | 0.147059 |
| classification | mfsnn | Reaching C-J-M-T_1 | single_day | 20151201 | 20151201 | 44 | accuracy | 0.263158 |
| classification | mfsnn | Reaching C-J-M-T_1 | single_day | 20151201 | 20151201 | 46 | accuracy | 0.447368 |
| classification | mfsnn | Reaching C-J-M-T_1 | single_day | 20151201 | 20151201 | 54 | accuracy | 0.315789 |
| classification | mfsnn | Reaching C-J-M-T_1 | single_day | 20160914 | 20160914 | 44 | accuracy | 0.820513 |
| classification | mfsnn | Reaching C-J-M-T_1 | single_day | 20160914 | 20160914 | 46 | accuracy | 0.641026 |
| classification | mfsnn | Reaching C-J-M-T_1 | single_day | 20160914 | 20160914 | 54 | accuracy | 0.666667 |
| classification | mfsnn | Reaching C-J-M-T_1 | single_day | 20160915 | 20160915 | 44 | accuracy | 0.72973 |
| classification | mfsnn | Reaching C-J-M-T_1 | single_day | 20160915 | 20160915 | 46 | accuracy | 0.378378 |
| classification | mfsnn | Reaching C-J-M-T_1 | single_day | 20160915 | 20160915 | 54 | accuracy | 0.486486 |
| classification | mfsnn | Reaching C-J-M-T_1 | single_day | 20160919 | 20160919 | 44 | accuracy | 0.394737 |
| classification | mfsnn | Reaching C-J-M-T_1 | single_day | 20160919 | 20160919 | 46 | accuracy | 0.447368 |
| classification | mfsnn | Reaching C-J-M-T_1 | single_day | 20160919 | 20160919 | 54 | accuracy | 0.473684 |
| classification | mfsnn | Reaching C-J-M-T_1 | single_day | 20160921 | 20160921 | 44 | accuracy | 0.722222 |
| classification | mfsnn | Reaching C-J-M-T_1 | single_day | 20160921 | 20160921 | 46 | accuracy | 0.527778 |
| classification | mfsnn | Reaching C-J-M-T_1 | single_day | 20160921 | 20160921 | 54 | accuracy | 0.638889 |
| classification | mfsnn | Reaching C-J-M-T_1 | single_day | 20160923 | 20160923 | 44 | accuracy | 0.682927 |
| classification | mfsnn | Reaching C-J-M-T_1 | single_day | 20160923 | 20160923 | 46 | accuracy | 0.634146 |
| classification | mfsnn | Reaching C-J-M-T_1 | single_day | 20160923 | 20160923 | 54 | accuracy | 0.634146 |
| classification | mfsnn | Reaching C-J-M-T_1 | single_day | 20160929 | 20160929 | 44 | accuracy | 0.333333 |
| classification | mfsnn | Reaching C-J-M-T_1 | single_day | 20160929 | 20160929 | 46 | accuracy | 0.52381 |
| classification | mfsnn | Reaching C-J-M-T_1 | single_day | 20160929 | 20160929 | 54 | accuracy | 0.333333 |
| classification | mfsnn | Reaching C-J-M-T_1 | single_day | 20161005 | 20161005 | 44 | accuracy | 0.243902 |
| classification | mfsnn | Reaching C-J-M-T_1 | single_day | 20161005 | 20161005 | 46 | accuracy | 0.268293 |
| classification | mfsnn | Reaching C-J-M-T_1 | single_day | 20161005 | 20161005 | 54 | accuracy | 0.536585 |
| classification | mfsnn | Reaching C-J-M-T_1 | single_day | 20161011 | 20161011 | 44 | accuracy | 0.625 |
| classification | mfsnn | Reaching C-J-M-T_1 | single_day | 20161011 | 20161011 | 46 | accuracy | 0.475 |
| classification | mfsnn | Reaching C-J-M-T_1 | single_day | 20161011 | 20161011 | 54 | accuracy | 0.55 |
| classification | mfsnn | Reaching C-J-M-T_1 | single_day | 20161013 | 20161013 | 44 | accuracy | 0.708333 |
| classification | mfsnn | Reaching C-J-M-T_1 | single_day | 20161013 | 20161013 | 46 | accuracy | 0.708333 |
| classification | mfsnn | Reaching C-J-M-T_1 | single_day | 20161013 | 20161013 | 54 | accuracy | 0.625 |
| classification | mfsnn | Reaching C-J-M-T_1 | single_day | 20161021 | 20161021 | 44 | accuracy | 0.534483 |
| classification | mfsnn | Reaching C-J-M-T_1 | single_day | 20161021 | 20161021 | 46 | accuracy | 0.517241 |
| classification | mfsnn | Reaching C-J-M-T_1 | single_day | 20161021 | 20161021 | 54 | accuracy | 0.551724 |
| classification | mscformer | Reaching C-J-M-T_1 | single_day | 20131003 | 20131003 | 44 | accuracy | 0.75 |
| classification | mscformer | Reaching C-J-M-T_1 | single_day | 20131003 | 20131003 | 46 | accuracy | 0.71875 |
| classification | mscformer | Reaching C-J-M-T_1 | single_day | 20131003 | 20131003 | 54 | accuracy | 0.65625 |
| classification | mscformer | Reaching C-J-M-T_1 | single_day | 20131022 | 20131022 | 44 | accuracy | 0.935484 |
| classification | mscformer | Reaching C-J-M-T_1 | single_day | 20131022 | 20131022 | 46 | accuracy | 0.774194 |
| classification | mscformer | Reaching C-J-M-T_1 | single_day | 20131022 | 20131022 | 54 | accuracy | 0.709677 |
| classification | mscformer | Reaching C-J-M-T_1 | single_day | 20131023 | 20131023 | 44 | accuracy | 0.846154 |
| classification | mscformer | Reaching C-J-M-T_1 | single_day | 20131023 | 20131023 | 46 | accuracy | 0.846154 |
| classification | mscformer | Reaching C-J-M-T_1 | single_day | 20131023 | 20131023 | 54 | accuracy | 0.871795 |
| classification | mscformer | Reaching C-J-M-T_1 | single_day | 20131101 | 20131101 | 44 | accuracy | 0.78 |
| classification | mscformer | Reaching C-J-M-T_1 | single_day | 20131101 | 20131101 | 46 | accuracy | 0.68 |
| classification | mscformer | Reaching C-J-M-T_1 | single_day | 20131101 | 20131101 | 54 | accuracy | 0.64 |
| classification | mscformer | Reaching C-J-M-T_1 | single_day | 20131203 | 20131203 | 44 | accuracy | 0.735294 |
| classification | mscformer | Reaching C-J-M-T_1 | single_day | 20131203 | 20131203 | 46 | accuracy | 0.764706 |
| classification | mscformer | Reaching C-J-M-T_1 | single_day | 20131203 | 20131203 | 54 | accuracy | 0.588235 |

| | | | | | | | | |
|---|---|---|---|---|---|---|---|---|
| classification | mscformer | Reaching C-J-M-T_1 | single_day | 20131204 | 20131204 | 44 | accuracy | 0.727273 |
| classification | mscformer | Reaching C-J-M-T_1 | single_day | 20131204 | 20131204 | 46 | accuracy | 0.636364 |
| classification | mscformer | Reaching C-J-M-T_1 | single_day | 20131204 | 20131204 | 54 | accuracy | 0.787879 |
| classification | mscformer | Reaching C-J-M-T_1 | single_day | 20131219 | 20131219 | 44 | accuracy | 0.583333 |
| classification | mscformer | Reaching C-J-M-T_1 | single_day | 20131219 | 20131219 | 46 | accuracy | 0.722222 |
| classification | mscformer | Reaching C-J-M-T_1 | single_day | 20131219 | 20131219 | 54 | accuracy | 0.805556 |
| classification | mscformer | Reaching C-J-M-T_1 | single_day | 20131220 | 20131220 | 44 | accuracy | 0.675 |
| classification | mscformer | Reaching C-J-M-T_1 | single_day | 20131220 | 20131220 | 46 | accuracy | 0.5 |
| classification | mscformer | Reaching C-J-M-T_1 | single_day | 20131220 | 20131220 | 54 | accuracy | 0.65 |
| classification | mscformer | Reaching C-J-M-T_1 | single_day | 20150311 | 20150311 | 44 | accuracy | 0.874317 |
| classification | mscformer | Reaching C-J-M-T_1 | single_day | 20150311 | 20150311 | 46 | accuracy | 0.918033 |
| classification | mscformer | Reaching C-J-M-T_1 | single_day | 20150311 | 20150311 | 54 | accuracy | 0.89071 |
| classification | mscformer | Reaching C-J-M-T_1 | single_day | 20150312 | 20150312 | 44 | accuracy | 0.93956 |
| classification | mscformer | Reaching C-J-M-T_1 | single_day | 20150312 | 20150312 | 46 | accuracy | 0.928571 |
| classification | mscformer | Reaching C-J-M-T_1 | single_day | 20150312 | 20150312 | 54 | accuracy | 0.945055 |
| classification | mscformer | Reaching C-J-M-T_1 | single_day | 20150313 | 20150313 | 44 | accuracy | 0.966346 |
| classification | mscformer | Reaching C-J-M-T_1 | single_day | 20150313 | 20150313 | 46 | accuracy | 0.918269 |
| classification | mscformer | Reaching C-J-M-T_1 | single_day | 20150313 | 20150313 | 54 | accuracy | 0.918269 |
| classification | mscformer | Reaching C-J-M-T_1 | single_day | 20150319 | 20150319 | 44 | accuracy | 0.941748 |
| classification | mscformer | Reaching C-J-M-T_1 | single_day | 20150319 | 20150319 | 46 | accuracy | 0.946602 |
| classification | mscformer | Reaching C-J-M-T_1 | single_day | 20150319 | 20150319 | 54 | accuracy | 0.946602 |
| classification | mscformer | Reaching C-J-M-T_1 | single_day | 20150629 | 20150629 | 44 | accuracy | 0.805556 |
| classification | mscformer | Reaching C-J-M-T_1 | single_day | 20150629 | 20150629 | 46 | accuracy | 0.722222 |
| classification | mscformer | Reaching C-J-M-T_1 | single_day | 20150629 | 20150629 | 54 | accuracy | 0.777778 |
| classification | mscformer | Reaching C-J-M-T_1 | single_day | 20150630 | 20150630 | 44 | accuracy | 0.777778 |
| classification | mscformer | Reaching C-J-M-T_1 | single_day | 20150630 | 20150630 | 46 | accuracy | 0.916667 |
| classification | mscformer | Reaching C-J-M-T_1 | single_day | 20150630 | 20150630 | 54 | accuracy | 0.805556 |
| classification | mscformer | Reaching C-J-M-T_1 | single_day | 20150701 | 20150701 | 44 | accuracy | 0.9 |
| classification | mscformer | Reaching C-J-M-T_1 | single_day | 20150701 | 20150701 | 46 | accuracy | 0.775 |
| classification | mscformer | Reaching C-J-M-T_1 | single_day | 20150701 | 20150701 | 54 | accuracy | 0.7 |
| classification | mscformer | Reaching C-J-M-T_1 | single_day | 20150706 | 20150706 | 44 | accuracy | 0.945946 |
| classification | mscformer | Reaching C-J-M-T_1 | single_day | 20150706 | 20150706 | 46 | accuracy | 0.864865 |
| classification | mscformer | Reaching C-J-M-T_1 | single_day | 20150706 | 20150706 | 54 | accuracy | 0.837838 |
| classification | mscformer | Reaching C-J-M-T_1 | single_day | 20150708 | 20150708 | 44 | accuracy | 0.894737 |
| classification | mscformer | Reaching C-J-M-T_1 | single_day | 20150708 | 20150708 | 46 | accuracy | 0.842105 |
| classification | mscformer | Reaching C-J-M-T_1 | single_day | 20150708 | 20150708 | 54 | accuracy | 0.815789 |
| classification | mscformer | Reaching C-J-M-T_1 | single_day | 20150709 | 20150709 | 44 | accuracy | 0.85 |
| classification | mscformer | Reaching C-J-M-T_1 | single_day | 20150709 | 20150709 | 46 | accuracy | 0.925 |
| classification | mscformer | Reaching C-J-M-T_1 | single_day | 20150709 | 20150709 | 54 | accuracy | 0.9 |
| classification | mscformer | Reaching C-J-M-T_1 | single_day | 20150710 | 20150710 | 44 | accuracy | 0.804878 |
| classification | mscformer | Reaching C-J-M-T_1 | single_day | 20150710 | 20150710 | 46 | accuracy | 0.902439 |
| classification | mscformer | Reaching C-J-M-T_1 | single_day | 20150710 | 20150710 | 54 | accuracy | 0.804878 |
| classification | mscformer | Reaching C-J-M-T_1 | single_day | 20150713 | 20150713 | 44 | accuracy | 0.891892 |
| classification | mscformer | Reaching C-J-M-T_1 | single_day | 20150713 | 20150713 | 46 | accuracy | 0.756757 |
| classification | mscformer | Reaching C-J-M-T_1 | single_day | 20150713 | 20150713 | 54 | accuracy | 0.891892 |
| classification | mscformer | Reaching C-J-M-T_1 | single_day | 20150714 | 20150714 | 44 | accuracy | 0.975 |
| classification | mscformer | Reaching C-J-M-T_1 | single_day | 20150714 | 20150714 | 46 | accuracy | 0.95 |
| classification | mscformer | Reaching C-J-M-T_1 | single_day | 20150714 | 20150714 | 54 | accuracy | 0.85 |
| classification | mscformer | Reaching C-J-M-T_1 | single_day | 20150715 | 20150715 | 44 | accuracy | 0.846154 |
| classification | mscformer | Reaching C-J-M-T_1 | single_day | 20150715 | 20150715 | 46 | accuracy | 0.897436 |
| classification | mscformer | Reaching C-J-M-T_1 | single_day | 20150715 | 20150715 | 54 | accuracy | 0.769231 |
| classification | mscformer | Reaching C-J-M-T_1 | single_day | 20150716 | 20150716 | 44 | accuracy | 0.794872 |
| classification | mscformer | Reaching C-J-M-T_1 | single_day | 20150716 | 20150716 | 46 | accuracy | 0.846154 |
| classification | mscformer | Reaching C-J-M-T_1 | single_day | 20150716 | 20150716 | 54 | accuracy | 0.820513 |
| classification | mscformer | Reaching C-J-M-T_1 | single_day | 20151103 | 20151103 | 44 | accuracy | 0.78 |
| classification | mscformer | Reaching C-J-M-T_1 | single_day | 20151103 | 20151103 | 46 | accuracy | 0.78 |
| classification | mscformer | Reaching C-J-M-T_1 | single_day | 20151103 | 20151103 | 54 | accuracy | 0.8 |
| classification | mscformer | Reaching C-J-M-T_1 | single_day | 20151106 | 20151106 | 44 | accuracy | 0.846154 |
| classification | mscformer | Reaching C-J-M-T_1 | single_day | 20151106 | 20151106 | 46 | accuracy | 0.826923 |
| classification | mscformer | Reaching C-J-M-T_1 | single_day | 20151106 | 20151106 | 54 | accuracy | 0.75 |
| classification | mscformer | Reaching C-J-M-T_1 | single_day | 20151109 | 20151109 | 44 | accuracy | 0.742857 |
| classification | mscformer | Reaching C-J-M-T_1 | single_day | 20151109 | 20151109 | 46 | accuracy | 0.742857 |
| classification | mscformer | Reaching C-J-M-T_1 | single_day | 20151109 | 20151109 | 54 | accuracy | 0.742857 |

| | | | | | | | | |
|---|---|---|---|---|---|---|---|---|
| classification | mscformer | Reaching C-J-M-T_1 | single_day | 20151112 | 20151112 | 44 | accuracy | 0.789474 |
| classification | mscformer | Reaching C-J-M-T_1 | single_day | 20151112 | 20151112 | 46 | accuracy | 0.815789 |
| classification | mscformer | Reaching C-J-M-T_1 | single_day | 20151112 | 20151112 | 54 | accuracy | 0.763158 |
| classification | mscformer | Reaching C-J-M-T_1 | single_day | 20151113 | 20151113 | 44 | accuracy | 0.868421 |
| classification | mscformer | Reaching C-J-M-T_1 | single_day | 20151113 | 20151113 | 46 | accuracy | 0.789474 |
| classification | mscformer | Reaching C-J-M-T_1 | single_day | 20151113 | 20151113 | 54 | accuracy | 0.815789 |
| classification | mscformer | Reaching C-J-M-T_1 | single_day | 20151116 | 20151116 | 44 | accuracy | 0.735294 |
| classification | mscformer | Reaching C-J-M-T_1 | single_day | 20151116 | 20151116 | 46 | accuracy | 0.882353 |
| classification | mscformer | Reaching C-J-M-T_1 | single_day | 20151116 | 20151116 | 54 | accuracy | 0.823529 |
| classification | mscformer | Reaching C-J-M-T_1 | single_day | 20151119 | 20151119 | 44 | accuracy | 0.928571 |
| classification | mscformer | Reaching C-J-M-T_1 | single_day | 20151119 | 20151119 | 46 | accuracy | 0.904762 |
| classification | mscformer | Reaching C-J-M-T_1 | single_day | 20151119 | 20151119 | 54 | accuracy | 0.880952 |
| classification | mscformer | Reaching C-J-M-T_1 | single_day | 20151120 | 20151120 | 44 | accuracy | 0.794118 |
| classification | mscformer | Reaching C-J-M-T_1 | single_day | 20151120 | 20151120 | 46 | accuracy | 0.970588 |
| classification | mscformer | Reaching C-J-M-T_1 | single_day | 20151120 | 20151120 | 54 | accuracy | 0.852941 |
| classification | mscformer | Reaching C-J-M-T_1 | single_day | 20151201 | 20151201 | 44 | accuracy | 0.789474 |
| classification | mscformer | Reaching C-J-M-T_1 | single_day | 20151201 | 20151201 | 46 | accuracy | 0.842105 |
| classification | mscformer | Reaching C-J-M-T_1 | single_day | 20151201 | 20151201 | 54 | accuracy | 0.921053 |
| classification | mscformer | Reaching C-J-M-T_1 | single_day | 20160914 | 20160914 | 44 | accuracy | 0.846154 |
| classification | mscformer | Reaching C-J-M-T_1 | single_day | 20160914 | 20160914 | 46 | accuracy | 0.897436 |
| classification | mscformer | Reaching C-J-M-T_1 | single_day | 20160914 | 20160914 | 54 | accuracy | 0.897436 |
| classification | mscformer | Reaching C-J-M-T_1 | single_day | 20160915 | 20160915 | 44 | accuracy | 0.972973 |
| classification | mscformer | Reaching C-J-M-T_1 | single_day | 20160915 | 20160915 | 46 | accuracy | 0.864865 |
| classification | mscformer | Reaching C-J-M-T_1 | single_day | 20160915 | 20160915 | 54 | accuracy | 0.891892 |
| classification | mscformer | Reaching C-J-M-T_1 | single_day | 20160919 | 20160919 | 44 | accuracy | 0.868421 |
| classification | mscformer | Reaching C-J-M-T_1 | single_day | 20160919 | 20160919 | 46 | accuracy | 0.868421 |
| classification | mscformer | Reaching C-J-M-T_1 | single_day | 20160919 | 20160919 | 54 | accuracy | 0.789474 |
| classification | mscformer | Reaching C-J-M-T_1 | single_day | 20160921 | 20160921 | 44 | accuracy | 0.888889 |
| classification | mscformer | Reaching C-J-M-T_1 | single_day | 20160921 | 20160921 | 46 | accuracy | 0.861111 |
| classification | mscformer | Reaching C-J-M-T_1 | single_day | 20160921 | 20160921 | 54 | accuracy | 0.777778 |
| classification | mscformer | Reaching C-J-M-T_1 | single_day | 20160923 | 20160923 | 44 | accuracy | 0.926829 |
| classification | mscformer | Reaching C-J-M-T_1 | single_day | 20160923 | 20160923 | 46 | accuracy | 0.926829 |
| classification | mscformer | Reaching C-J-M-T_1 | single_day | 20160923 | 20160923 | 54 | accuracy | 0.878049 |
| classification | mscformer | Reaching C-J-M-T_1 | single_day | 20160929 | 20160929 | 44 | accuracy | 0.928571 |
| classification | mscformer | Reaching C-J-M-T_1 | single_day | 20160929 | 20160929 | 46 | accuracy | 0.880952 |
| classification | mscformer | Reaching C-J-M-T_1 | single_day | 20160929 | 20160929 | 54 | accuracy | 0.952381 |
| classification | mscformer | Reaching C-J-M-T_1 | single_day | 20161005 | 20161005 | 44 | accuracy | 0.926829 |
| classification | mscformer | Reaching C-J-M-T_1 | single_day | 20161005 | 20161005 | 46 | accuracy | 0.95122 |
| classification | mscformer | Reaching C-J-M-T_1 | single_day | 20161005 | 20161005 | 54 | accuracy | 0.95122 |
| classification | mscformer | Reaching C-J-M-T_1 | single_day | 20161011 | 20161011 | 44 | accuracy | 0.95 |
| classification | mscformer | Reaching C-J-M-T_1 | single_day | 20161011 | 20161011 | 46 | accuracy | 0.85 |
| classification | mscformer | Reaching C-J-M-T_1 | single_day | 20161011 | 20161011 | 54 | accuracy | 0.825 |
| classification | mscformer | Reaching C-J-M-T_1 | single_day | 20161013 | 20161013 | 44 | accuracy | 0.9375 |
| classification | mscformer | Reaching C-J-M-T_1 | single_day | 20161013 | 20161013 | 46 | accuracy | 0.875 |
| classification | mscformer | Reaching C-J-M-T_1 | single_day | 20161013 | 20161013 | 54 | accuracy | 0.9375 |
| classification | mscformer | Reaching C-J-M-T_1 | single_day | 20161021 | 20161021 | 44 | accuracy | 0.965517 |
| classification | mscformer | Reaching C-J-M-T_1 | single_day | 20161021 | 20161021 | 46 | accuracy | 0.948276 |
| classification | mscformer | Reaching C-J-M-T_1 | single_day | 20161021 | 20161021 | 54 | accuracy | 0.965517 |
| classification | nomad | Reaching C-J-M-T_1 | single_day | 20131003 | 20131003 | 44 | accuracy | 0.21875 |
| classification | nomad | Reaching C-J-M-T_1 | single_day | 20131003 | 20131003 | 46 | accuracy | 0.53125 |
| classification | nomad | Reaching C-J-M-T_1 | single_day | 20131003 | 20131003 | 54 | accuracy | 0.15625 |
| classification | nomad | Reaching C-J-M-T_1 | single_day | 20131022 | 20131022 | 44 | accuracy | 0.677419 |
| classification | nomad | Reaching C-J-M-T_1 | single_day | 20131022 | 20131022 | 46 | accuracy | 0.645161 |
| classification | nomad | Reaching C-J-M-T_1 | single_day | 20131022 | 20131022 | 54 | accuracy | 0.516129 |
| classification | nomad | Reaching C-J-M-T_1 | single_day | 20131023 | 20131023 | 44 | accuracy | 0.769231 |
| classification | nomad | Reaching C-J-M-T_1 | single_day | 20131023 | 20131023 | 46 | accuracy | 0.589744 |
| classification | nomad | Reaching C-J-M-T_1 | single_day | 20131023 | 20131023 | 54 | accuracy | 0.435897 |
| classification | nomad | Reaching C-J-M-T_1 | single_day | 20131101 | 20131101 | 44 | accuracy | 0.76 |
| classification | nomad | Reaching C-J-M-T_1 | single_day | 20131101 | 20131101 | 46 | accuracy | 0.76 |
| classification | nomad | Reaching C-J-M-T_1 | single_day | 20131101 | 20131101 | 54 | accuracy | 0.64 |
| classification | nomad | Reaching C-J-M-T_1 | single_day | 20131203 | 20131203 | 44 | accuracy | 0.647059 |
| classification | nomad | Reaching C-J-M-T_1 | single_day | 20131203 | 20131203 | 46 | accuracy | 0.176471 |
| classification | nomad | Reaching C-J-M-T_1 | single_day | 20131203 | 20131203 | 54 | accuracy | 0.5 |

| | | | | | | | | |
|---|---|---|---|---|---|---|---|---|
| classification | nomad | Reaching C-J-M-T_1 | single_day | 20131204 | 20131204 | 44 | accuracy | 0.454545 |
| classification | nomad | Reaching C-J-M-T_1 | single_day | 20131204 | 20131204 | 46 | accuracy | 0.606061 |
| classification | nomad | Reaching C-J-M-T_1 | single_day | 20131204 | 20131204 | 54 | accuracy | 0.69697 |
| classification | nomad | Reaching C-J-M-T_1 | single_day | 20131219 | 20131219 | 44 | accuracy | 0.305556 |
| classification | nomad | Reaching C-J-M-T_1 | single_day | 20131219 | 20131219 | 46 | accuracy | 0.555556 |
| classification | nomad | Reaching C-J-M-T_1 | single_day | 20131219 | 20131219 | 54 | accuracy | 0.472222 |
| classification | nomad | Reaching C-J-M-T_1 | single_day | 20131220 | 20131220 | 44 | accuracy | 0.125 |
| classification | nomad | Reaching C-J-M-T_1 | single_day | 20131220 | 20131220 | 46 | accuracy | 0.425 |
| classification | nomad | Reaching C-J-M-T_1 | single_day | 20131220 | 20131220 | 54 | accuracy | 0.1 |
| classification | nomad | Reaching C-J-M-T_1 | single_day | 20150311 | 20150311 | 44 | accuracy | 0.945355 |
| classification | nomad | Reaching C-J-M-T_1 | single_day | 20150311 | 20150311 | 46 | accuracy | 0.928962 |
| classification | nomad | Reaching C-J-M-T_1 | single_day | 20150311 | 20150311 | 54 | accuracy | 0.939891 |
| classification | nomad | Reaching C-J-M-T_1 | single_day | 20150312 | 20150312 | 44 | accuracy | 0.945055 |
| classification | nomad | Reaching C-J-M-T_1 | single_day | 20150312 | 20150312 | 46 | accuracy | 0.950549 |
| classification | nomad | Reaching C-J-M-T_1 | single_day | 20150312 | 20150312 | 54 | accuracy | 0.950549 |
| classification | nomad | Reaching C-J-M-T_1 | single_day | 20150313 | 20150313 | 44 | accuracy | 0.966346 |
| classification | nomad | Reaching C-J-M-T_1 | single_day | 20150313 | 20150313 | 46 | accuracy | 0.966346 |
| classification | nomad | Reaching C-J-M-T_1 | single_day | 20150313 | 20150313 | 54 | accuracy | 0.951923 |
| classification | nomad | Reaching C-J-M-T_1 | single_day | 20150319 | 20150319 | 44 | accuracy | 0.966019 |
| classification | nomad | Reaching C-J-M-T_1 | single_day | 20150319 | 20150319 | 46 | accuracy | 0.936893 |
| classification | nomad | Reaching C-J-M-T_1 | single_day | 20150319 | 20150319 | 54 | accuracy | 0.932039 |
| classification | nomad | Reaching C-J-M-T_1 | single_day | 20150629 | 20150629 | 44 | accuracy | 0.805556 |
| classification | nomad | Reaching C-J-M-T_1 | single_day | 20150629 | 20150629 | 46 | accuracy | 0.805556 |
| classification | nomad | Reaching C-J-M-T_1 | single_day | 20150629 | 20150629 | 54 | accuracy | 0.861111 |
| classification | nomad | Reaching C-J-M-T_1 | single_day | 20150630 | 20150630 | 44 | accuracy | 0.888889 |
| classification | nomad | Reaching C-J-M-T_1 | single_day | 20150630 | 20150630 | 46 | accuracy | 0.861111 |
| classification | nomad | Reaching C-J-M-T_1 | single_day | 20150630 | 20150630 | 54 | accuracy | 0.722222 |
| classification | nomad | Reaching C-J-M-T_1 | single_day | 20150701 | 20150701 | 44 | accuracy | 0.775 |
| classification | nomad | Reaching C-J-M-T_1 | single_day | 20150701 | 20150701 | 46 | accuracy | 0.225 |
| classification | nomad | Reaching C-J-M-T_1 | single_day | 20150701 | 20150701 | 54 | accuracy | 0.9 |
| classification | nomad | Reaching C-J-M-T_1 | single_day | 20150706 | 20150706 | 44 | accuracy | 0.837838 |
| classification | nomad | Reaching C-J-M-T_1 | single_day | 20150706 | 20150706 | 46 | accuracy | 0.918919 |
| classification | nomad | Reaching C-J-M-T_1 | single_day | 20150706 | 20150706 | 54 | accuracy | 0.837838 |
| classification | nomad | Reaching C-J-M-T_1 | single_day | 20150708 | 20150708 | 44 | accuracy | 0.894737 |
| classification | nomad | Reaching C-J-M-T_1 | single_day | 20150708 | 20150708 | 46 | accuracy | 0.921053 |
| classification | nomad | Reaching C-J-M-T_1 | single_day | 20150708 | 20150708 | 54 | accuracy | 0.921053 |
| classification | nomad | Reaching C-J-M-T_1 | single_day | 20150709 | 20150709 | 44 | accuracy | 0.85 |
| classification | nomad | Reaching C-J-M-T_1 | single_day | 20150709 | 20150709 | 46 | accuracy | 0.8 |
| classification | nomad | Reaching C-J-M-T_1 | single_day | 20150709 | 20150709 | 54 | accuracy | 0.775 |
| classification | nomad | Reaching C-J-M-T_1 | single_day | 20150710 | 20150710 | 44 | accuracy | 0.804878 |
| classification | nomad | Reaching C-J-M-T_1 | single_day | 20150710 | 20150710 | 46 | accuracy | 0.902439 |
| classification | nomad | Reaching C-J-M-T_1 | single_day | 20150710 | 20150710 | 54 | accuracy | 0.682927 |
| classification | nomad | Reaching C-J-M-T_1 | single_day | 20150713 | 20150713 | 44 | accuracy | 0.189189 |
| classification | nomad | Reaching C-J-M-T_1 | single_day | 20150713 | 20150713 | 46 | accuracy | 0.864865 |
| classification | nomad | Reaching C-J-M-T_1 | single_day | 20150713 | 20150713 | 54 | accuracy | 0.621622 |
| classification | nomad | Reaching C-J-M-T_1 | single_day | 20150714 | 20150714 | 44 | accuracy | 0.625 |
| classification | nomad | Reaching C-J-M-T_1 | single_day | 20150714 | 20150714 | 46 | accuracy | 0.425 |
| classification | nomad | Reaching C-J-M-T_1 | single_day | 20150714 | 20150714 | 54 | accuracy | 0.7 |
| classification | nomad | Reaching C-J-M-T_1 | single_day | 20150715 | 20150715 | 44 | accuracy | 0.871795 |
| classification | nomad | Reaching C-J-M-T_1 | single_day | 20150715 | 20150715 | 46 | accuracy | 0.897436 |
| classification | nomad | Reaching C-J-M-T_1 | single_day | 20150715 | 20150715 | 54 | accuracy | 0.974359 |
| classification | nomad | Reaching C-J-M-T_1 | single_day | 20150716 | 20150716 | 44 | accuracy | 0.871795 |
| classification | nomad | Reaching C-J-M-T_1 | single_day | 20150716 | 20150716 | 46 | accuracy | 0.74359 |
| classification | nomad | Reaching C-J-M-T_1 | single_day | 20150716 | 20150716 | 54 | accuracy | 0.717949 |
| classification | nomad | Reaching C-J-M-T_1 | single_day | 20151103 | 20151103 | 44 | accuracy | 0.74 |
| classification | nomad | Reaching C-J-M-T_1 | single_day | 20151103 | 20151103 | 46 | accuracy | 0.68 |
| classification | nomad | Reaching C-J-M-T_1 | single_day | 20151103 | 20151103 | 54 | accuracy | 0.68 |
| classification | nomad | Reaching C-J-M-T_1 | single_day | 20151106 | 20151106 | 44 | accuracy | 0.788462 |
| classification | nomad | Reaching C-J-M-T_1 | single_day | 20151106 | 20151106 | 46 | accuracy | 0.769231 |
| classification | nomad | Reaching C-J-M-T_1 | single_day | 20151106 | 20151106 | 54 | accuracy | 0.865385 |
| classification | nomad | Reaching C-J-M-T_1 | single_day | 20151109 | 20151109 | 44 | accuracy | 0.2 |
| classification | nomad | Reaching C-J-M-T_1 | single_day | 20151109 | 20151109 | 46 | accuracy | 0.742857 |
| classification | nomad | Reaching C-J-M-T_1 | single_day | 20151109 | 20151109 | 54 | accuracy | 0.514286 |

| | | | | | | | | |
|---|---|---|---|---|---|---|---|---|
| classification | nomad | Reaching C-J-M-T_1 | single_day | 20151112 | 20151112 | 44 | accuracy | 0.763158 |
| classification | nomad | Reaching C-J-M-T_1 | single_day | 20151112 | 20151112 | 46 | accuracy | 0.842105 |
| classification | nomad | Reaching C-J-M-T_1 | single_day | 20151112 | 20151112 | 54 | accuracy | 0.894737 |
| classification | nomad | Reaching C-J-M-T_1 | single_day | 20151113 | 20151113 | 44 | accuracy | 0.789474 |
| classification | nomad | Reaching C-J-M-T_1 | single_day | 20151113 | 20151113 | 46 | accuracy | 0.315789 |
| classification | nomad | Reaching C-J-M-T_1 | single_day | 20151113 | 20151113 | 54 | accuracy | 0.763158 |
| classification | nomad | Reaching C-J-M-T_1 | single_day | 20151116 | 20151116 | 44 | accuracy | 0.205882 |
| classification | nomad | Reaching C-J-M-T_1 | single_day | 20151116 | 20151116 | 46 | accuracy | 0.735294 |
| classification | nomad | Reaching C-J-M-T_1 | single_day | 20151116 | 20151116 | 54 | accuracy | 0.647059 |
| classification | nomad | Reaching C-J-M-T_1 | single_day | 20151119 | 20151119 | 44 | accuracy | 0.666667 |
| classification | nomad | Reaching C-J-M-T_1 | single_day | 20151119 | 20151119 | 46 | accuracy | 0.809524 |
| classification | nomad | Reaching C-J-M-T_1 | single_day | 20151119 | 20151119 | 54 | accuracy | 0.952381 |
| classification | nomad | Reaching C-J-M-T_1 | single_day | 20151120 | 20151120 | 44 | accuracy | 0.823529 |
| classification | nomad | Reaching C-J-M-T_1 | single_day | 20151120 | 20151120 | 46 | accuracy | 0.588235 |
| classification | nomad | Reaching C-J-M-T_1 | single_day | 20151120 | 20151120 | 54 | accuracy | 0.794118 |
| classification | nomad | Reaching C-J-M-T_1 | single_day | 20151201 | 20151201 | 44 | accuracy | 0.763158 |
| classification | nomad | Reaching C-J-M-T_1 | single_day | 20151201 | 20151201 | 46 | accuracy | 0.710526 |
| classification | nomad | Reaching C-J-M-T_1 | single_day | 20151201 | 20151201 | 54 | accuracy | 0.157895 |
| classification | nomad | Reaching C-J-M-T_1 | single_day | 20160914 | 20160914 | 44 | accuracy | 0.692308 |
| classification | nomad | Reaching C-J-M-T_1 | single_day | 20160914 | 20160914 | 46 | accuracy | 0.717949 |
| classification | nomad | Reaching C-J-M-T_1 | single_day | 20160914 | 20160914 | 54 | accuracy | 0.74359 |
| classification | nomad | Reaching C-J-M-T_1 | single_day | 20160915 | 20160915 | 44 | accuracy | 0.108108 |
| classification | nomad | Reaching C-J-M-T_1 | single_day | 20160915 | 20160915 | 46 | accuracy | 0.945946 |
| classification | nomad | Reaching C-J-M-T_1 | single_day | 20160915 | 20160915 | 54 | accuracy | 0.918919 |
| classification | nomad | Reaching C-J-M-T_1 | single_day | 20160919 | 20160919 | 44 | accuracy | 0.894737 |
| classification | nomad | Reaching C-J-M-T_1 | single_day | 20160919 | 20160919 | 46 | accuracy | 0.578947 |
| classification | nomad | Reaching C-J-M-T_1 | single_day | 20160919 | 20160919 | 54 | accuracy | 0.342105 |
| classification | nomad | Reaching C-J-M-T_1 | single_day | 20160921 | 20160921 | 44 | accuracy | 0.805556 |
| classification | nomad | Reaching C-J-M-T_1 | single_day | 20160921 | 20160921 | 46 | accuracy | 0.75 |
| classification | nomad | Reaching C-J-M-T_1 | single_day | 20160921 | 20160921 | 54 | accuracy | 0.666667 |
| classification | nomad | Reaching C-J-M-T_1 | single_day | 20160923 | 20160923 | 44 | accuracy | 0.780488 |
| classification | nomad | Reaching C-J-M-T_1 | single_day | 20160923 | 20160923 | 46 | accuracy | 0.97561 |
| classification | nomad | Reaching C-J-M-T_1 | single_day | 20160923 | 20160923 | 54 | accuracy | 0.853659 |
| classification | nomad | Reaching C-J-M-T_1 | single_day | 20160929 | 20160929 | 44 | accuracy | 0.857143 |
| classification | nomad | Reaching C-J-M-T_1 | single_day | 20160929 | 20160929 | 46 | accuracy | 0.833333 |
| classification | nomad | Reaching C-J-M-T_1 | single_day | 20160929 | 20160929 | 54 | accuracy | 0.47619 |
| classification | nomad | Reaching C-J-M-T_1 | single_day | 20161005 | 20161005 | 44 | accuracy | 0.756098 |
| classification | nomad | Reaching C-J-M-T_1 | single_day | 20161005 | 20161005 | 46 | accuracy | 0.804878 |
| classification | nomad | Reaching C-J-M-T_1 | single_day | 20161005 | 20161005 | 54 | accuracy | 0.902439 |
| classification | nomad | Reaching C-J-M-T_1 | single_day | 20161011 | 20161011 | 44 | accuracy | 0.8 |
| classification | nomad | Reaching C-J-M-T_1 | single_day | 20161011 | 20161011 | 46 | accuracy | 0.7 |
| classification | nomad | Reaching C-J-M-T_1 | single_day | 20161011 | 20161011 | 54 | accuracy | 0.9 |
| classification | nomad | Reaching C-J-M-T_1 | single_day | 20161013 | 20161013 | 44 | accuracy | 0.958333 |
| classification | nomad | Reaching C-J-M-T_1 | single_day | 20161013 | 20161013 | 46 | accuracy | 0.895833 |
| classification | nomad | Reaching C-J-M-T_1 | single_day | 20161013 | 20161013 | 54 | accuracy | 0.833333 |
| classification | nomad | Reaching C-J-M-T_1 | single_day | 20161021 | 20161021 | 44 | accuracy | 0.982759 |
| classification | nomad | Reaching C-J-M-T_1 | single_day | 20161021 | 20161021 | 46 | accuracy | 0.948276 |
| classification | nomad | Reaching C-J-M-T_1 | single_day | 20161021 | 20161021 | 54 | accuracy | 0.862069 |
| classification | mlp | Reaching C-J-M-T_1 | single_day | 20131003 | 20131003 | 44 | accuracy | 0.5 |
| classification | mlp | Reaching C-J-M-T_1 | single_day | 20131003 | 20131003 | 46 | accuracy | 0.6875 |
| classification | mlp | Reaching C-J-M-T_1 | single_day | 20131003 | 20131003 | 54 | accuracy | 0.65625 |
| classification | mlp | Reaching C-J-M-T_1 | single_day | 20131022 | 20131022 | 44 | accuracy | 0.741935 |
| classification | mlp | Reaching C-J-M-T_1 | single_day | 20131022 | 20131022 | 46 | accuracy | 0.645161 |
| classification | mlp | Reaching C-J-M-T_1 | single_day | 20131022 | 20131022 | 54 | accuracy | 0.709677 |
| classification | mlp | Reaching C-J-M-T_1 | single_day | 20131023 | 20131023 | 44 | accuracy | 0.74359 |
| classification | mlp | Reaching C-J-M-T_1 | single_day | 20131023 | 20131023 | 46 | accuracy | 0.769231 |
| classification | mlp | Reaching C-J-M-T_1 | single_day | 20131023 | 20131023 | 54 | accuracy | 0.769231 |
| classification | mlp | Reaching C-J-M-T_1 | single_day | 20131101 | 20131101 | 44 | accuracy | 0.7 |
| classification | mlp | Reaching C-J-M-T_1 | single_day | 20131101 | 20131101 | 46 | accuracy | 0.78 |
| classification | mlp | Reaching C-J-M-T_1 | single_day | 20131101 | 20131101 | 54 | accuracy | 0.82 |
| classification | mlp | Reaching C-J-M-T_1 | single_day | 20131203 | 20131203 | 44 | accuracy | 0.676471 |
| classification | mlp | Reaching C-J-M-T_1 | single_day | 20131203 | 20131203 | 46 | accuracy | 0.558824 |
| classification | mlp | Reaching C-J-M-T_1 | single_day | 20131203 | 20131203 | 54 | accuracy | 0.647059 |

| | | | | | | | | |
|---|---|---|---|---|---|---|---|---|
| classification | mlp | Reaching C-J-M-T_1 | single_day | 20131204 | 20131204 | 44 | accuracy | 0.69697 |
| classification | mlp | Reaching C-J-M-T_1 | single_day | 20131204 | 20131204 | 46 | accuracy | 0.575758 |
| classification | mlp | Reaching C-J-M-T_1 | single_day | 20131204 | 20131204 | 54 | accuracy | 0.666667 |
| classification | mlp | Reaching C-J-M-T_1 | single_day | 20131219 | 20131219 | 44 | accuracy | 0.611111 |
| classification | mlp | Reaching C-J-M-T_1 | single_day | 20131219 | 20131219 | 46 | accuracy | 0.611111 |
| classification | mlp | Reaching C-J-M-T_1 | single_day | 20131219 | 20131219 | 54 | accuracy | 0.555556 |
| classification | mlp | Reaching C-J-M-T_1 | single_day | 20131220 | 20131220 | 44 | accuracy | 0.575 |
| classification | mlp | Reaching C-J-M-T_1 | single_day | 20131220 | 20131220 | 46 | accuracy | 0.65 |
| classification | mlp | Reaching C-J-M-T_1 | single_day | 20131220 | 20131220 | 54 | accuracy | 0.55 |
| classification | mlp | Reaching C-J-M-T_1 | single_day | 20150311 | 20150311 | 44 | accuracy | 0.939891 |
| classification | mlp | Reaching C-J-M-T_1 | single_day | 20150311 | 20150311 | 46 | accuracy | 0.912568 |
| classification | mlp | Reaching C-J-M-T_1 | single_day | 20150311 | 20150311 | 54 | accuracy | 0.907104 |
| classification | mlp | Reaching C-J-M-T_1 | single_day | 20150312 | 20150312 | 44 | accuracy | 0.945055 |
| classification | mlp | Reaching C-J-M-T_1 | single_day | 20150312 | 20150312 | 46 | accuracy | 0.950549 |
| classification | mlp | Reaching C-J-M-T_1 | single_day | 20150312 | 20150312 | 54 | accuracy | 0.956044 |
| classification | mlp | Reaching C-J-M-T_1 | single_day | 20150313 | 20150313 | 44 | accuracy | 0.966346 |
| classification | mlp | Reaching C-J-M-T_1 | single_day | 20150313 | 20150313 | 46 | accuracy | 0.971154 |
| classification | mlp | Reaching C-J-M-T_1 | single_day | 20150313 | 20150313 | 54 | accuracy | 0.971154 |
| classification | mlp | Reaching C-J-M-T_1 | single_day | 20150319 | 20150319 | 44 | accuracy | 0.970874 |
| classification | mlp | Reaching C-J-M-T_1 | single_day | 20150319 | 20150319 | 46 | accuracy | 0.946602 |
| classification | mlp | Reaching C-J-M-T_1 | single_day | 20150319 | 20150319 | 54 | accuracy | 0.932039 |
| classification | mlp | Reaching C-J-M-T_1 | single_day | 20150629 | 20150629 | 44 | accuracy | 0.611111 |
| classification | mlp | Reaching C-J-M-T_1 | single_day | 20150629 | 20150629 | 46 | accuracy | 0.638889 |
| classification | mlp | Reaching C-J-M-T_1 | single_day | 20150629 | 20150629 | 54 | accuracy | 0.666667 |
| classification | mlp | Reaching C-J-M-T_1 | single_day | 20150630 | 20150630 | 44 | accuracy | 0.805556 |
| classification | mlp | Reaching C-J-M-T_1 | single_day | 20150630 | 20150630 | 46 | accuracy | 0.888889 |
| classification | mlp | Reaching C-J-M-T_1 | single_day | 20150630 | 20150630 | 54 | accuracy | 0.75 |
| classification | mlp | Reaching C-J-M-T_1 | single_day | 20150701 | 20150701 | 44 | accuracy | 0.8 |
| classification | mlp | Reaching C-J-M-T_1 | single_day | 20150701 | 20150701 | 46 | accuracy | 0.825 |
| classification | mlp | Reaching C-J-M-T_1 | single_day | 20150701 | 20150701 | 54 | accuracy | 0.75 |
| classification | mlp | Reaching C-J-M-T_1 | single_day | 20150706 | 20150706 | 44 | accuracy | 0.810811 |
| classification | mlp | Reaching C-J-M-T_1 | single_day | 20150706 | 20150706 | 46 | accuracy | 0.810811 |
| classification | mlp | Reaching C-J-M-T_1 | single_day | 20150706 | 20150706 | 54 | accuracy | 0.540541 |
| classification | mlp | Reaching C-J-M-T_1 | single_day | 20150708 | 20150708 | 44 | accuracy | 0.763158 |
| classification | mlp | Reaching C-J-M-T_1 | single_day | 20150708 | 20150708 | 46 | accuracy | 0.789474 |
| classification | mlp | Reaching C-J-M-T_1 | single_day | 20150708 | 20150708 | 54 | accuracy | 0.842105 |
| classification | mlp | Reaching C-J-M-T_1 | single_day | 20150709 | 20150709 | 44 | accuracy | 0.925 |
| classification | mlp | Reaching C-J-M-T_1 | single_day | 20150709 | 20150709 | 46 | accuracy | 0.75 |
| classification | mlp | Reaching C-J-M-T_1 | single_day | 20150709 | 20150709 | 54 | accuracy | 0.7 |
| classification | mlp | Reaching C-J-M-T_1 | single_day | 20150710 | 20150710 | 44 | accuracy | 0.804878 |
| classification | mlp | Reaching C-J-M-T_1 | single_day | 20150710 | 20150710 | 46 | accuracy | 0.780488 |
| classification | mlp | Reaching C-J-M-T_1 | single_day | 20150710 | 20150710 | 54 | accuracy | 0.878049 |
| classification | mlp | Reaching C-J-M-T_1 | single_day | 20150713 | 20150713 | 44 | accuracy | 0.864865 |
| classification | mlp | Reaching C-J-M-T_1 | single_day | 20150713 | 20150713 | 46 | accuracy | 0.837838 |
| classification | mlp | Reaching C-J-M-T_1 | single_day | 20150713 | 20150713 | 54 | accuracy | 0.72973 |
| classification | mlp | Reaching C-J-M-T_1 | single_day | 20150714 | 20150714 | 44 | accuracy | 0.7 |
| classification | mlp | Reaching C-J-M-T_1 | single_day | 20150714 | 20150714 | 46 | accuracy | 0.75 |
| classification | mlp | Reaching C-J-M-T_1 | single_day | 20150714 | 20150714 | 54 | accuracy | 0.8 |
| classification | mlp | Reaching C-J-M-T_1 | single_day | 20150715 | 20150715 | 44 | accuracy | 0.794872 |
| classification | mlp | Reaching C-J-M-T_1 | single_day | 20150715 | 20150715 | 46 | accuracy | 0.923077 |
| classification | mlp | Reaching C-J-M-T_1 | single_day | 20150715 | 20150715 | 54 | accuracy | 0.820513 |
| classification | mlp | Reaching C-J-M-T_1 | single_day | 20150716 | 20150716 | 44 | accuracy | 0.794872 |
| classification | mlp | Reaching C-J-M-T_1 | single_day | 20150716 | 20150716 | 46 | accuracy | 0.820513 |
| classification | mlp | Reaching C-J-M-T_1 | single_day | 20150716 | 20150716 | 54 | accuracy | 0.641026 |
| classification | mlp | Reaching C-J-M-T_1 | single_day | 20151103 | 20151103 | 44 | accuracy | 0.72 |
| classification | mlp | Reaching C-J-M-T_1 | single_day | 20151103 | 20151103 | 46 | accuracy | 0.64 |
| classification | mlp | Reaching C-J-M-T_1 | single_day | 20151103 | 20151103 | 54 | accuracy | 0.76 |
| classification | mlp | Reaching C-J-M-T_1 | single_day | 20151106 | 20151106 | 44 | accuracy | 0.653846 |
| classification | mlp | Reaching C-J-M-T_1 | single_day | 20151106 | 20151106 | 46 | accuracy | 0.75 |
| classification | mlp | Reaching C-J-M-T_1 | single_day | 20151106 | 20151106 | 54 | accuracy | 0.730769 |
| classification | mlp | Reaching C-J-M-T_1 | single_day | 20151109 | 20151109 | 44 | accuracy | 0.742857 |
| classification | mlp | Reaching C-J-M-T_1 | single_day | 20151109 | 20151109 | 46 | accuracy | 0.628571 |
| classification | mlp | Reaching C-J-M-T_1 | single_day | 20151109 | 20151109 | 54 | accuracy | 0.571429 |

| | | | | | | | | |
|---|---|---|---|---|---|---|---|---|
| classification | mlp | Reaching C-J-M-T_1 | single_day | 20151112 | 20151112 | 44 | accuracy | 0.868421 |
| classification | mlp | Reaching C-J-M-T_1 | single_day | 20151112 | 20151112 | 46 | accuracy | 0.763158 |
| classification | mlp | Reaching C-J-M-T_1 | single_day | 20151112 | 20151112 | 54 | accuracy | 0.815789 |
| classification | mlp | Reaching C-J-M-T_1 | single_day | 20151113 | 20151113 | 44 | accuracy | 0.894737 |
| classification | mlp | Reaching C-J-M-T_1 | single_day | 20151113 | 20151113 | 46 | accuracy | 0.815789 |
| classification | mlp | Reaching C-J-M-T_1 | single_day | 20151113 | 20151113 | 54 | accuracy | 0.815789 |
| classification | mlp | Reaching C-J-M-T_1 | single_day | 20151116 | 20151116 | 44 | accuracy | 0.794118 |
| classification | mlp | Reaching C-J-M-T_1 | single_day | 20151116 | 20151116 | 46 | accuracy | 0.647059 |
| classification | mlp | Reaching C-J-M-T_1 | single_day | 20151116 | 20151116 | 54 | accuracy | 0.705882 |
| classification | mlp | Reaching C-J-M-T_1 | single_day | 20151119 | 20151119 | 44 | accuracy | 0.738095 |
| classification | mlp | Reaching C-J-M-T_1 | single_day | 20151119 | 20151119 | 46 | accuracy | 0.833333 |
| classification | mlp | Reaching C-J-M-T_1 | single_day | 20151119 | 20151119 | 54 | accuracy | 0.785714 |
| classification | mlp | Reaching C-J-M-T_1 | single_day | 20151120 | 20151120 | 44 | accuracy | 0.794118 |
| classification | mlp | Reaching C-J-M-T_1 | single_day | 20151120 | 20151120 | 46 | accuracy | 0.911765 |
| classification | mlp | Reaching C-J-M-T_1 | single_day | 20151120 | 20151120 | 54 | accuracy | 0.852941 |
| classification | mlp | Reaching C-J-M-T_1 | single_day | 20151201 | 20151201 | 44 | accuracy | 0.947368 |
| classification | mlp | Reaching C-J-M-T_1 | single_day | 20151201 | 20151201 | 46 | accuracy | 0.789474 |
| classification | mlp | Reaching C-J-M-T_1 | single_day | 20151201 | 20151201 | 54 | accuracy | 0.894737 |
| classification | mlp | Reaching C-J-M-T_1 | single_day | 20160914 | 20160914 | 44 | accuracy | 0.717949 |
| classification | mlp | Reaching C-J-M-T_1 | single_day | 20160914 | 20160914 | 46 | accuracy | 0.769231 |
| classification | mlp | Reaching C-J-M-T_1 | single_day | 20160914 | 20160914 | 54 | accuracy | 0.74359 |
| classification | mlp | Reaching C-J-M-T_1 | single_day | 20160915 | 20160915 | 44 | accuracy | 0.756757 |
| classification | mlp | Reaching C-J-M-T_1 | single_day | 20160915 | 20160915 | 46 | accuracy | 0.783784 |
| classification | mlp | Reaching C-J-M-T_1 | single_day | 20160915 | 20160915 | 54 | accuracy | 0.756757 |
| classification | mlp | Reaching C-J-M-T_1 | single_day | 20160919 | 20160919 | 44 | accuracy | 0.815789 |
| classification | mlp | Reaching C-J-M-T_1 | single_day | 20160919 | 20160919 | 46 | accuracy | 0.789474 |
| classification | mlp | Reaching C-J-M-T_1 | single_day | 20160919 | 20160919 | 54 | accuracy | 0.605263 |
| classification | mlp | Reaching C-J-M-T_1 | single_day | 20160921 | 20160921 | 44 | accuracy | 0.861111 |
| classification | mlp | Reaching C-J-M-T_1 | single_day | 20160921 | 20160921 | 46 | accuracy | 0.805556 |
| classification | mlp | Reaching C-J-M-T_1 | single_day | 20160921 | 20160921 | 54 | accuracy | 0.805556 |
| classification | mlp | Reaching C-J-M-T_1 | single_day | 20160923 | 20160923 | 44 | accuracy | 0.829268 |
| classification | mlp | Reaching C-J-M-T_1 | single_day | 20160923 | 20160923 | 46 | accuracy | 0.829268 |
| classification | mlp | Reaching C-J-M-T_1 | single_day | 20160923 | 20160923 | 54 | accuracy | 0.829268 |
| classification | mlp | Reaching C-J-M-T_1 | single_day | 20160929 | 20160929 | 44 | accuracy | 0.880952 |
| classification | mlp | Reaching C-J-M-T_1 | single_day | 20160929 | 20160929 | 46 | accuracy | 0.928571 |
| classification | mlp | Reaching C-J-M-T_1 | single_day | 20160929 | 20160929 | 54 | accuracy | 0.785714 |
| classification | mlp | Reaching C-J-M-T_1 | single_day | 20161005 | 20161005 | 44 | accuracy | 0.853659 |
| classification | mlp | Reaching C-J-M-T_1 | single_day | 20161005 | 20161005 | 46 | accuracy | 0.902439 |
| classification | mlp | Reaching C-J-M-T_1 | single_day | 20161005 | 20161005 | 54 | accuracy | 0.853659 |
| classification | mlp | Reaching C-J-M-T_1 | single_day | 20161011 | 20161011 | 44 | accuracy | 0.775 |
| classification | mlp | Reaching C-J-M-T_1 | single_day | 20161011 | 20161011 | 46 | accuracy | 0.875 |
| classification | mlp | Reaching C-J-M-T_1 | single_day | 20161011 | 20161011 | 54 | accuracy | 0.725 |
| classification | mlp | Reaching C-J-M-T_1 | single_day | 20161013 | 20161013 | 44 | accuracy | 0.875 |
| classification | mlp | Reaching C-J-M-T_1 | single_day | 20161013 | 20161013 | 46 | accuracy | 0.9375 |
| classification | mlp | Reaching C-J-M-T_1 | single_day | 20161013 | 20161013 | 54 | accuracy | 0.645833 |
| classification | mlp | Reaching C-J-M-T_1 | single_day | 20161021 | 20161021 | 44 | accuracy | 0.810345 |
| classification | mlp | Reaching C-J-M-T_1 | single_day | 20161021 | 20161021 | 46 | accuracy | 0.913793 |
| classification | mlp | Reaching C-J-M-T_1 | single_day | 20161021 | 20161021 | 54 | accuracy | 0.931034 |
| classification | convpi_vae | Reaching C-J-M-T_1 | single_day | 20131003 | 20131003 | 31 | accuracy | 0.96875 |
| classification | convpi_vae | Reaching C-J-M-T_1 | single_day | 20131003 | 20131003 | 44 | accuracy | 1 |
| classification | convpi_vae | Reaching C-J-M-T_1 | single_day | 20131003 | 20131003 | 46 | accuracy | 0.96875 |
| classification | convpi_vae | Reaching C-J-M-T_1 | single_day | 20131003 | 20131003 | 54 | accuracy | 1 |
| classification | convpi_vae | Reaching C-J-M-T_1 | single_day | 20131003 | 20131003 | 69 | accuracy | 0.96875 |
| classification | convpi_vae | Reaching C-J-M-T_1 | single_day | 20131022 | 20131022 | 31 | accuracy | 1 |
| classification | convpi_vae | Reaching C-J-M-T_1 | single_day | 20131022 | 20131022 | 44 | accuracy | 1 |
| classification | convpi_vae | Reaching C-J-M-T_1 | single_day | 20131022 | 20131022 | 46 | accuracy | 1 |
| classification | convpi_vae | Reaching C-J-M-T_1 | single_day | 20131022 | 20131022 | 54 | accuracy | 1 |
| classification | convpi_vae | Reaching C-J-M-T_1 | single_day | 20131022 | 20131022 | 69 | accuracy | 1 |
| classification | convpi_vae | Reaching C-J-M-T_1 | single_day | 20131023 | 20131023 | 31 | accuracy | 1 |
| classification | convpi_vae | Reaching C-J-M-T_1 | single_day | 20131023 | 20131023 | 44 | accuracy | 1 |
| classification | convpi_vae | Reaching C-J-M-T_1 | single_day | 20131023 | 20131023 | 46 | accuracy | 1 |
| classification | convpi_vae | Reaching C-J-M-T_1 | single_day | 20131023 | 20131023 | 54 | accuracy | 1 |
| classification | convpi_vae | Reaching C-J-M-T_1 | single_day | 20131023 | 20131023 | 69 | accuracy | 1 |

| | | | | | | | | |
|---|---|---|---|---|---|---|---|---|
| classification | convpi_vae | Reaching C-J-M-T_1 | single_day | 20131101 | 20131101 | 31 | accuracy | 1 |
| classification | convpi_vae | Reaching C-J-M-T_1 | single_day | 20131101 | 20131101 | 44 | accuracy | 0.98 |
| classification | convpi_vae | Reaching C-J-M-T_1 | single_day | 20131101 | 20131101 | 46 | accuracy | 0.96 |
| classification | convpi_vae | Reaching C-J-M-T_1 | single_day | 20131101 | 20131101 | 54 | accuracy | 0.98 |
| classification | convpi_vae | Reaching C-J-M-T_1 | single_day | 20131101 | 20131101 | 69 | accuracy | 1 |
| classification | convpi_vae | Reaching C-J-M-T_1 | single_day | 20131203 | 20131203 | 31 | accuracy | 0.970588 |
| classification | convpi_vae | Reaching C-J-M-T_1 | single_day | 20131203 | 20131203 | 44 | accuracy | 0.970588 |
| classification | convpi_vae | Reaching C-J-M-T_1 | single_day | 20131203 | 20131203 | 46 | accuracy | 1 |
| classification | convpi_vae | Reaching C-J-M-T_1 | single_day | 20131203 | 20131203 | 54 | accuracy | 0.970588 |
| classification | convpi_vae | Reaching C-J-M-T_1 | single_day | 20131203 | 20131203 | 69 | accuracy | 1 |
| classification | convpi_vae | Reaching C-J-M-T_1 | single_day | 20131204 | 20131204 | 31 | accuracy | 0.939394 |
| classification | convpi_vae | Reaching C-J-M-T_1 | single_day | 20131204 | 20131204 | 44 | accuracy | 0.939394 |
| classification | convpi_vae | Reaching C-J-M-T_1 | single_day | 20131204 | 20131204 | 46 | accuracy | 0.969697 |
| classification | convpi_vae | Reaching C-J-M-T_1 | single_day | 20131204 | 20131204 | 54 | accuracy | 0.939394 |
| classification | convpi_vae | Reaching C-J-M-T_1 | single_day | 20131204 | 20131204 | 69 | accuracy | 0.909091 |
| classification | convpi_vae | Reaching C-J-M-T_1 | single_day | 20131219 | 20131219 | 31 | accuracy | 1 |
| classification | convpi_vae | Reaching C-J-M-T_1 | single_day | 20131219 | 20131219 | 44 | accuracy | 0.972222 |
| classification | convpi_vae | Reaching C-J-M-T_1 | single_day | 20131219 | 20131219 | 46 | accuracy | 0.972222 |
| classification | convpi_vae | Reaching C-J-M-T_1 | single_day | 20131219 | 20131219 | 54 | accuracy | 1 |
| classification | convpi_vae | Reaching C-J-M-T_1 | single_day | 20131219 | 20131219 | 69 | accuracy | 1 |
| classification | convpi_vae | Reaching C-J-M-T_1 | single_day | 20131220 | 20131220 | 31 | accuracy | 0.975 |
| classification | convpi_vae | Reaching C-J-M-T_1 | single_day | 20131220 | 20131220 | 44 | accuracy | 1 |
| classification | convpi_vae | Reaching C-J-M-T_1 | single_day | 20131220 | 20131220 | 46 | accuracy | 0.975 |
| classification | convpi_vae | Reaching C-J-M-T_1 | single_day | 20131220 | 20131220 | 54 | accuracy | 1 |
| classification | convpi_vae | Reaching C-J-M-T_1 | single_day | 20131220 | 20131220 | 69 | accuracy | 1 |
| classification | convpi_vae | Reaching C-J-M-T_1 | single_day | 20150311 | 20150311 | 31 | accuracy | 0.989071 |
| classification | convpi_vae | Reaching C-J-M-T_1 | single_day | 20150311 | 20150311 | 44 | accuracy | 0.978142 |
| classification | convpi_vae | Reaching C-J-M-T_1 | single_day | 20150311 | 20150311 | 46 | accuracy | 0.978142 |
| classification | convpi_vae | Reaching C-J-M-T_1 | single_day | 20150311 | 20150311 | 54 | accuracy | 1 |
| classification | convpi_vae | Reaching C-J-M-T_1 | single_day | 20150311 | 20150311 | 69 | accuracy | 1 |
| classification | convpi_vae | Reaching C-J-M-T_1 | single_day | 20150319 | 20150319 | 31 | accuracy | 0.985437 |
| classification | convpi_vae | Reaching C-J-M-T_1 | single_day | 20150319 | 20150319 | 44 | accuracy | 0.995146 |
| classification | convpi_vae | Reaching C-J-M-T_1 | single_day | 20150319 | 20150319 | 46 | accuracy | 0.980583 |
| classification | convpi_vae | Reaching C-J-M-T_1 | single_day | 20150319 | 20150319 | 54 | accuracy | 0.990291 |
| classification | convpi_vae | Reaching C-J-M-T_1 | single_day | 20150319 | 20150319 | 69 | accuracy | 0.990291 |
| classification | convpi_vae | Reaching C-J-M-T_1 | single_day | 20150629 | 20150629 | 31 | accuracy | 0.916667 |
| classification | convpi_vae | Reaching C-J-M-T_1 | single_day | 20150629 | 20150629 | 44 | accuracy | 1 |
| classification | convpi_vae | Reaching C-J-M-T_1 | single_day | 20150629 | 20150629 | 44 | accuracy | 0.222222 |
| classification | convpi_vae | Reaching C-J-M-T_1 | single_day | 20150629 | 20150629 | 46 | accuracy | 1 |
| classification | convpi_vae | Reaching C-J-M-T_1 | single_day | 20150629 | 20150629 | 46 | accuracy | 0.194444 |
| classification | convpi_vae | Reaching C-J-M-T_1 | single_day | 20150629 | 20150629 | 54 | accuracy | 1 |
| classification | convpi_vae | Reaching C-J-M-T_1 | single_day | 20150629 | 20150629 | 54 | accuracy | 0.25 |
| classification | convpi_vae | Reaching C-J-M-T_1 | single_day | 20150629 | 20150629 | 69 | accuracy | 1 |
| classification | convpi_vae | Reaching C-J-M-T_1 | single_day | 20150630 | 20150630 | 31 | accuracy | 1 |
| classification | convpi_vae | Reaching C-J-M-T_1 | single_day | 20150630 | 20150630 | 44 | accuracy | 1 |
| classification | convpi_vae | Reaching C-J-M-T_1 | single_day | 20150630 | 20150630 | 44 | accuracy | 0.0833333 |
| classification | convpi_vae | Reaching C-J-M-T_1 | single_day | 20150630 | 20150630 | 46 | accuracy | 1 |
| classification | convpi_vae | Reaching C-J-M-T_1 | single_day | 20150630 | 20150630 | 46 | accuracy | 0.25 |
| classification | convpi_vae | Reaching C-J-M-T_1 | single_day | 20150630 | 20150630 | 54 | accuracy | 1 |
| classification | convpi_vae | Reaching C-J-M-T_1 | single_day | 20150630 | 20150630 | 54 | accuracy | 0.194444 |
| classification | convpi_vae | Reaching C-J-M-T_1 | single_day | 20150630 | 20150630 | 69 | accuracy | 1 |
| classification | convpi_vae | Reaching C-J-M-T_1 | single_day | 20150701 | 20150701 | 31 | accuracy | 0.975 |
| classification | convpi_vae | Reaching C-J-M-T_1 | single_day | 20150701 | 20150701 | 44 | accuracy | 1 |
| classification | convpi_vae | Reaching C-J-M-T_1 | single_day | 20150701 | 20150701 | 44 | accuracy | 0.1 |
| classification | convpi_vae | Reaching C-J-M-T_1 | single_day | 20150701 | 20150701 | 46 | accuracy | 1 |
| classification | convpi_vae | Reaching C-J-M-T_1 | single_day | 20150701 | 20150701 | 46 | accuracy | 0.275 |
| classification | convpi_vae | Reaching C-J-M-T_1 | single_day | 20150701 | 20150701 | 54 | accuracy | 1 |
| classification | convpi_vae | Reaching C-J-M-T_1 | single_day | 20150701 | 20150701 | 54 | accuracy | 0.1 |
| classification | convpi_vae | Reaching C-J-M-T_1 | single_day | 20150701 | 20150701 | 69 | accuracy | 1 |
| classification | convpi_vae | Reaching C-J-M-T_1 | single_day | 20150703 | 20150703 | 44 | accuracy | 0.189189 |
| classification | convpi_vae | Reaching C-J-M-T_1 | single_day | 20150703 | 20150703 | 46 | accuracy | 0.162162 |
| classification | convpi_vae | Reaching C-J-M-T_1 | single_day | 20150703 | 20150703 | 54 | accuracy | 0.135135 |
| classification | convpi_vae | Reaching C-J-M-T_1 | single_day | 20150706 | 20150706 | 31 | accuracy | 0.972973 |

| | | | | | | | | |
|---|---|---|---|---|---|---|---|---|
| classification | convpi_vae | Reaching C-J-M-T_1 | single_day | 20150706 | 20150706 | 44 | accuracy | 1 |
| classification | convpi_vae | Reaching C-J-M-T_1 | single_day | 20150706 | 20150706 | 44 | accuracy | 0.0540541 |
| classification | convpi_vae | Reaching C-J-M-T_1 | single_day | 20150706 | 20150706 | 46 | accuracy | 1 |
| classification | convpi_vae | Reaching C-J-M-T_1 | single_day | 20150706 | 20150706 | 46 | accuracy | 0.135135 |
| classification | convpi_vae | Reaching C-J-M-T_1 | single_day | 20150706 | 20150706 | 54 | accuracy | 1 |
| classification | convpi_vae | Reaching C-J-M-T_1 | single_day | 20150706 | 20150706 | 54 | accuracy | 0.162162 |
| classification | convpi_vae | Reaching C-J-M-T_1 | single_day | 20150706 | 20150706 | 69 | accuracy | 1 |
| classification | convpi_vae | Reaching C-J-M-T_1 | single_day | 20150707 | 20150707 | 44 | accuracy | 0.210526 |
| classification | convpi_vae | Reaching C-J-M-T_1 | single_day | 20150707 | 20150707 | 46 | accuracy | 0.105263 |
| classification | convpi_vae | Reaching C-J-M-T_1 | single_day | 20150707 | 20150707 | 54 | accuracy | 0.184211 |
| classification | convpi_vae | Reaching C-J-M-T_1 | single_day | 20150708 | 20150708 | 31 | accuracy | 1 |
| classification | convpi_vae | Reaching C-J-M-T_1 | single_day | 20150708 | 20150708 | 44 | accuracy | 1 |
| classification | convpi_vae | Reaching C-J-M-T_1 | single_day | 20150708 | 20150708 | 44 | accuracy | 0.105263 |
| classification | convpi_vae | Reaching C-J-M-T_1 | single_day | 20150708 | 20150708 | 46 | accuracy | 1 |
| classification | convpi_vae | Reaching C-J-M-T_1 | single_day | 20150708 | 20150708 | 46 | accuracy | 0.157895 |
| classification | convpi_vae | Reaching C-J-M-T_1 | single_day | 20150708 | 20150708 | 54 | accuracy | 1 |
| classification | convpi_vae | Reaching C-J-M-T_1 | single_day | 20150708 | 20150708 | 54 | accuracy | 0.236842 |
| classification | convpi_vae | Reaching C-J-M-T_1 | single_day | 20150708 | 20150708 | 69 | accuracy | 1 |
| classification | convpi_vae | Reaching C-J-M-T_1 | single_day | 20150709 | 20150709 | 31 | accuracy | 1 |
| classification | convpi_vae | Reaching C-J-M-T_1 | single_day | 20150709 | 20150709 | 44 | accuracy | 1 |
| classification | convpi_vae | Reaching C-J-M-T_1 | single_day | 20150709 | 20150709 | 44 | accuracy | 0.275 |
| classification | convpi_vae | Reaching C-J-M-T_1 | single_day | 20150709 | 20150709 | 46 | accuracy | 1 |
| classification | convpi_vae | Reaching C-J-M-T_1 | single_day | 20150709 | 20150709 | 46 | accuracy | 0.075 |
| classification | convpi_vae | Reaching C-J-M-T_1 | single_day | 20150709 | 20150709 | 54 | accuracy | 1 |
| classification | convpi_vae | Reaching C-J-M-T_1 | single_day | 20150709 | 20150709 | 54 | accuracy | 0.225 |
| classification | convpi_vae | Reaching C-J-M-T_1 | single_day | 20150709 | 20150709 | 69 | accuracy | 1 |
| classification | convpi_vae | Reaching C-J-M-T_1 | single_day | 20150710 | 20150710 | 31 | accuracy | 1 |
| classification | convpi_vae | Reaching C-J-M-T_1 | single_day | 20150710 | 20150710 | 44 | accuracy | 1 |
| classification | convpi_vae | Reaching C-J-M-T_1 | single_day | 20150710 | 20150710 | 44 | accuracy | 0.097561 |
| classification | convpi_vae | Reaching C-J-M-T_1 | single_day | 20150710 | 20150710 | 46 | accuracy | 1 |
| classification | convpi_vae | Reaching C-J-M-T_1 | single_day | 20150710 | 20150710 | 46 | accuracy | 0.219512 |
| classification | convpi_vae | Reaching C-J-M-T_1 | single_day | 20150710 | 20150710 | 54 | accuracy | 1 |
| classification | convpi_vae | Reaching C-J-M-T_1 | single_day | 20150710 | 20150710 | 54 | accuracy | 0.243902 |
| classification | convpi_vae | Reaching C-J-M-T_1 | single_day | 20150710 | 20150710 | 69 | accuracy | 1 |
| classification | convpi_vae | Reaching C-J-M-T_1 | single_day | 20150713 | 20150713 | 31 | accuracy | 0.972973 |
| classification | convpi_vae | Reaching C-J-M-T_1 | single_day | 20150713 | 20150713 | 44 | accuracy | 1 |
| classification | convpi_vae | Reaching C-J-M-T_1 | single_day | 20150713 | 20150713 | 46 | accuracy | 1 |
| classification | convpi_vae | Reaching C-J-M-T_1 | single_day | 20150713 | 20150713 | 54 | accuracy | 1 |
| classification | convpi_vae | Reaching C-J-M-T_1 | single_day | 20150713 | 20150713 | 69 | accuracy | 1 |
| classification | convpi_vae | Reaching C-J-M-T_1 | single_day | 20150714 | 20150714 | 31 | accuracy | 1 |
| classification | convpi_vae | Reaching C-J-M-T_1 | single_day | 20150714 | 20150714 | 44 | accuracy | 1 |
| classification | convpi_vae | Reaching C-J-M-T_1 | single_day | 20150714 | 20150714 | 46 | accuracy | 0.975 |
| classification | convpi_vae | Reaching C-J-M-T_1 | single_day | 20150714 | 20150714 | 54 | accuracy | 1 |
| classification | convpi_vae | Reaching C-J-M-T_1 | single_day | 20150714 | 20150714 | 69 | accuracy | 1 |
| classification | convpi_vae | Reaching C-J-M-T_1 | single_day | 20150715 | 20150715 | 31 | accuracy | 0.974359 |
| classification | convpi_vae | Reaching C-J-M-T_1 | single_day | 20150715 | 20150715 | 44 | accuracy | 1 |
| classification | convpi_vae | Reaching C-J-M-T_1 | single_day | 20150715 | 20150715 | 46 | accuracy | 1 |
| classification | convpi_vae | Reaching C-J-M-T_1 | single_day | 20150715 | 20150715 | 54 | accuracy | 1 |
| classification | convpi_vae | Reaching C-J-M-T_1 | single_day | 20150715 | 20150715 | 69 | accuracy | 1 |
| classification | convpi_vae | Reaching C-J-M-T_1 | single_day | 20150716 | 20150716 | 31 | accuracy | 0.974359 |
| classification | convpi_vae | Reaching C-J-M-T_1 | single_day | 20150716 | 20150716 | 44 | accuracy | 1 |
| classification | convpi_vae | Reaching C-J-M-T_1 | single_day | 20150716 | 20150716 | 46 | accuracy | 0.974359 |
| classification | convpi_vae | Reaching C-J-M-T_1 | single_day | 20150716 | 20150716 | 54 | accuracy | 0.974359 |
| classification | convpi_vae | Reaching C-J-M-T_1 | single_day | 20150716 | 20150716 | 69 | accuracy | 1 |
| classification | convpi_vae | Reaching C-J-M-T_1 | single_day | 20151103 | 20151103 | 31 | accuracy | 1 |
| classification | convpi_vae | Reaching C-J-M-T_1 | single_day | 20151103 | 20151103 | 44 | accuracy | 1 |
| classification | convpi_vae | Reaching C-J-M-T_1 | single_day | 20151103 | 20151103 | 46 | accuracy | 1 |
| classification | convpi_vae | Reaching C-J-M-T_1 | single_day | 20151103 | 20151103 | 54 | accuracy | 1 |
| classification | convpi_vae | Reaching C-J-M-T_1 | single_day | 20151103 | 20151103 | 69 | accuracy | 0.98 |
| classification | convpi_vae | Reaching C-J-M-T_1 | single_day | 20151106 | 20151106 | 31 | accuracy | 0.923077 |
| classification | convpi_vae | Reaching C-J-M-T_1 | single_day | 20151106 | 20151106 | 44 | accuracy | 0.980769 |
| classification | convpi_vae | Reaching C-J-M-T_1 | single_day | 20151106 | 20151106 | 46 | accuracy | 0.942308 |
| classification | convpi_vae | Reaching C-J-M-T_1 | single_day | 20151106 | 20151106 | 54 | accuracy | 0.942308 |

| | | | | | | | | |
|---|---|---|---|---|---|---|---|---|
| classification | convpi_vae | Reaching C-J-M-T_1 | single_day | 20151106 | 20151106 | 69 | accuracy | 0.980769 |
| classification | convpi_vae | Reaching C-J-M-T_1 | single_day | 20151112 | 20151112 | 31 | accuracy | 1 |
| classification | convpi_vae | Reaching C-J-M-T_1 | single_day | 20151112 | 20151112 | 44 | accuracy | 1 |
| classification | convpi_vae | Reaching C-J-M-T_1 | single_day | 20151112 | 20151112 | 46 | accuracy | 0.947368 |
| classification | convpi_vae | Reaching C-J-M-T_1 | single_day | 20151112 | 20151112 | 54 | accuracy | 1 |
| classification | convpi_vae | Reaching C-J-M-T_1 | single_day | 20151112 | 20151112 | 69 | accuracy | 1 |
| classification | convpi_vae | Reaching C-J-M-T_1 | single_day | 20151113 | 20151113 | 31 | accuracy | 1 |
| classification | convpi_vae | Reaching C-J-M-T_1 | single_day | 20151113 | 20151113 | 44 | accuracy | 1 |
| classification | convpi_vae | Reaching C-J-M-T_1 | single_day | 20151113 | 20151113 | 46 | accuracy | 1 |
| classification | convpi_vae | Reaching C-J-M-T_1 | single_day | 20151113 | 20151113 | 54 | accuracy | 1 |
| classification | convpi_vae | Reaching C-J-M-T_1 | single_day | 20151113 | 20151113 | 69 | accuracy | 1 |
| classification | convpi_vae | Reaching C-J-M-T_1 | single_day | 20151116 | 20151116 | 31 | accuracy | 0.941176 |
| classification | convpi_vae | Reaching C-J-M-T_1 | single_day | 20151116 | 20151116 | 44 | accuracy | 1 |
| classification | convpi_vae | Reaching C-J-M-T_1 | single_day | 20151116 | 20151116 | 46 | accuracy | 1 |
| classification | convpi_vae | Reaching C-J-M-T_1 | single_day | 20151116 | 20151116 | 54 | accuracy | 1 |
| classification | convpi_vae | Reaching C-J-M-T_1 | single_day | 20151116 | 20151116 | 69 | accuracy | 1 |
| classification | convpi_vae | Reaching C-J-M-T_1 | single_day | 20151119 | 20151119 | 31 | accuracy | 1 |
| classification | convpi_vae | Reaching C-J-M-T_1 | single_day | 20151119 | 20151119 | 44 | accuracy | 1 |
| classification | convpi_vae | Reaching C-J-M-T_1 | single_day | 20151119 | 20151119 | 46 | accuracy | 1 |
| classification | convpi_vae | Reaching C-J-M-T_1 | single_day | 20151119 | 20151119 | 54 | accuracy | 0.97619 |
| classification | convpi_vae | Reaching C-J-M-T_1 | single_day | 20151119 | 20151119 | 69 | accuracy | 1 |
| classification | convpi_vae | Reaching C-J-M-T_1 | single_day | 20151120 | 20151120 | 31 | accuracy | 1 |
| classification | convpi_vae | Reaching C-J-M-T_1 | single_day | 20151120 | 20151120 | 44 | accuracy | 1 |
| classification | convpi_vae | Reaching C-J-M-T_1 | single_day | 20151120 | 20151120 | 46 | accuracy | 1 |
| classification | convpi_vae | Reaching C-J-M-T_1 | single_day | 20151120 | 20151120 | 54 | accuracy | 1 |
| classification | convpi_vae | Reaching C-J-M-T_1 | single_day | 20151120 | 20151120 | 69 | accuracy | 1 |
| classification | convpi_vae | Reaching C-J-M-T_1 | single_day | 20151201 | 20151201 | 31 | accuracy | 0.947368 |
| classification | convpi_vae | Reaching C-J-M-T_1 | single_day | 20151201 | 20151201 | 44 | accuracy | 0.921053 |
| classification | convpi_vae | Reaching C-J-M-T_1 | single_day | 20151201 | 20151201 | 46 | accuracy | 1 |
| classification | convpi_vae | Reaching C-J-M-T_1 | single_day | 20151201 | 20151201 | 54 | accuracy | 0.973684 |
| classification | convpi_vae | Reaching C-J-M-T_1 | single_day | 20151201 | 20151201 | 69 | accuracy | 1 |
| classification | convpi_vae | Reaching C-J-M-T_1 | single_day | 20160914 | 20160914 | 31 | accuracy | 1 |
| classification | convpi_vae | Reaching C-J-M-T_1 | single_day | 20160914 | 20160914 | 44 | accuracy | 1 |
| classification | convpi_vae | Reaching C-J-M-T_1 | single_day | 20160914 | 20160914 | 46 | accuracy | 1 |
| classification | convpi_vae | Reaching C-J-M-T_1 | single_day | 20160914 | 20160914 | 54 | accuracy | 1 |
| classification | convpi_vae | Reaching C-J-M-T_1 | single_day | 20160914 | 20160914 | 69 | accuracy | 0.974359 |
| classification | convpi_vae | Reaching C-J-M-T_1 | single_day | 20160915 | 20160915 | 31 | accuracy | 0.972973 |
| classification | convpi_vae | Reaching C-J-M-T_1 | single_day | 20160915 | 20160915 | 44 | accuracy | 1 |
| classification | convpi_vae | Reaching C-J-M-T_1 | single_day | 20160915 | 20160915 | 46 | accuracy | 0.918919 |
| classification | convpi_vae | Reaching C-J-M-T_1 | single_day | 20160915 | 20160915 | 54 | accuracy | 1 |
| classification | convpi_vae | Reaching C-J-M-T_1 | single_day | 20160915 | 20160915 | 69 | accuracy | 0.945946 |
| classification | convpi_vae | Reaching C-J-M-T_1 | single_day | 20160919 | 20160919 | 31 | accuracy | 0.973684 |
| classification | convpi_vae | Reaching C-J-M-T_1 | single_day | 20160919 | 20160919 | 44 | accuracy | 1 |
| classification | convpi_vae | Reaching C-J-M-T_1 | single_day | 20160919 | 20160919 | 46 | accuracy | 1 |
| classification | convpi_vae | Reaching C-J-M-T_1 | single_day | 20160919 | 20160919 | 54 | accuracy | 1 |
| classification | convpi_vae | Reaching C-J-M-T_1 | single_day | 20160919 | 20160919 | 69 | accuracy | 1 |
| classification | convpi_vae | Reaching C-J-M-T_1 | single_day | 20160921 | 20160921 | 31 | accuracy | 0.972222 |
| classification | convpi_vae | Reaching C-J-M-T_1 | single_day | 20160921 | 20160921 | 44 | accuracy | 0.944444 |
| classification | convpi_vae | Reaching C-J-M-T_1 | single_day | 20160921 | 20160921 | 46 | accuracy | 0.972222 |
| classification | convpi_vae | Reaching C-J-M-T_1 | single_day | 20160921 | 20160921 | 54 | accuracy | 1 |
| classification | convpi_vae | Reaching C-J-M-T_1 | single_day | 20160921 | 20160921 | 69 | accuracy | 1 |
| classification | convpi_vae | Reaching C-J-M-T_1 | single_day | 20160923 | 20160923 | 31 | accuracy | 0.926829 |
| classification | convpi_vae | Reaching C-J-M-T_1 | single_day | 20160923 | 20160923 | 44 | accuracy | 1 |
| classification | convpi_vae | Reaching C-J-M-T_1 | single_day | 20160923 | 20160923 | 46 | accuracy | 1 |
| classification | convpi_vae | Reaching C-J-M-T_1 | single_day | 20160923 | 20160923 | 54 | accuracy | 0.95122 |
| classification | convpi_vae | Reaching C-J-M-T_1 | single_day | 20160923 | 20160923 | 69 | accuracy | 0.95122 |
| classification | convpi_vae | Reaching C-J-M-T_1 | single_day | 20160929 | 20160929 | 31 | accuracy | 0.97619 |
| classification | convpi_vae | Reaching C-J-M-T_1 | single_day | 20160929 | 20160929 | 44 | accuracy | 1 |
| classification | convpi_vae | Reaching C-J-M-T_1 | single_day | 20160929 | 20160929 | 46 | accuracy | 0.952381 |
| classification | convpi_vae | Reaching C-J-M-T_1 | single_day | 20160929 | 20160929 | 54 | accuracy | 1 |
| classification | convpi_vae | Reaching C-J-M-T_1 | single_day | 20160929 | 20160929 | 69 | accuracy | 1 |
| classification | convpi_vae | Reaching C-J-M-T_1 | single_day | 20161005 | 20161005 | 31 | accuracy | 1 |
| classification | convpi_vae | Reaching C-J-M-T_1 | single_day | 20161005 | 20161005 | 44 | accuracy | 1 |

| | | | | | | | | |
|---|---|---|---|---|---|---|---|---|
| classification | convpi_vae | Reaching C-J-M-T_1 | single_day | 20161005 | 20161005 | 46 | accuracy | 0.95122 |
| classification | convpi_vae | Reaching C-J-M-T_1 | single_day | 20161005 | 20161005 | 54 | accuracy | 1 |
| classification | convpi_vae | Reaching C-J-M-T_1 | single_day | 20161005 | 20161005 | 69 | accuracy | 1 |
| classification | convpi_vae | Reaching C-J-M-T_1 | single_day | 20161011 | 20161011 | 31 | accuracy | 1 |
| classification | convpi_vae | Reaching C-J-M-T_1 | single_day | 20161011 | 20161011 | 44 | accuracy | 1 |
| classification | convpi_vae | Reaching C-J-M-T_1 | single_day | 20161011 | 20161011 | 46 | accuracy | 1 |
| classification | convpi_vae | Reaching C-J-M-T_1 | single_day | 20161011 | 20161011 | 54 | accuracy | 1 |
| classification | convpi_vae | Reaching C-J-M-T_1 | single_day | 20161011 | 20161011 | 69 | accuracy | 0.975 |
| classification | convpi_vae | Reaching C-J-M-T_1 | single_day | 20161013 | 20161013 | 31 | accuracy | 1 |
| classification | convpi_vae | Reaching C-J-M-T_1 | single_day | 20161013 | 20161013 | 44 | accuracy | 0.979167 |
| classification | convpi_vae | Reaching C-J-M-T_1 | single_day | 20161013 | 20161013 | 46 | accuracy | 0.958333 |
| classification | convpi_vae | Reaching C-J-M-T_1 | single_day | 20161013 | 20161013 | 54 | accuracy | 1 |
| classification | convpi_vae | Reaching C-J-M-T_1 | single_day | 20161013 | 20161013 | 69 | accuracy | 1 |
| classification | convpi_vae | Reaching C-J-M-T_1 | single_day | 20161021 | 20161021 | 31 | accuracy | 1 |
| classification | convpi_vae | Reaching C-J-M-T_1 | single_day | 20161021 | 20161021 | 44 | accuracy | 0.948276 |
| classification | convpi_vae | Reaching C-J-M-T_1 | single_day | 20161021 | 20161021 | 46 | accuracy | 1 |
| classification | convpi_vae | Reaching C-J-M-T_1 | single_day | 20161021 | 20161021 | 54 | accuracy | 1 |
| classification | convpi_vae | Reaching C-J-M-T_1 | single_day | 20161021 | 20161021 | 69 | accuracy | 0.982759 |
| classification | lstm | Reaching C-J-M-T_1 | cross_day | 20131003__to__20160914 | 20160915__to__20161021 | 31 | accuracy | 0.150685 |
| classification | lstm | Reaching C-J-M-T_1 | cross_day | 20131003__to__20160914 | 20160915__to__20161021 | 44 | accuracy | 0.135661 |
| classification | lstm | Reaching C-J-M-T_1 | cross_day | 20131003__to__20160914 | 20160915__to__20161021 | 46 | accuracy | 0.126823 |
| classification | lstm | Reaching C-J-M-T_1 | cross_day | 20131003__to__20160914 | 20160915__to__20161021 | 54 | accuracy | 0.124613 |
| classification | lstm | Reaching C-J-M-T_1 | cross_day | 20131003__to__20160914 | 20160915__to__20161021 | 69 | accuracy | 0.138754 |
| classification | lstm | Reaching C-J-M-T_1 | cross_day | 20150629__to__20150629 | 20150701__to__20150701 | 31 | accuracy | 0.142857 |
| classification | lstm | Reaching C-J-M-T_1 | cross_day | 20150629__to__20150629 | 20150701__to__20150701 | 44 | accuracy | 0.142857 |
| classification | lstm | Reaching C-J-M-T_1 | cross_day | 20150629__to__20150629 | 20150701__to__20150701 | 46 | accuracy | 0.147959 |
| classification | lstm | Reaching C-J-M-T_1 | cross_day | 20150629__to__20150707 | 20150709__to__20150710 | 44 | accuracy | 0.110276 |
| classification | lstm | Reaching C-J-M-T_1 | cross_day | 20150629__to__20150707 | 20150709__to__20150710 | 44 | accuracy | 0.110398 |
| classification | lstm | Reaching C-J-M-T_1 | cross_day | 20150629__to__20150707 | 20150709__to__20150710 | 46 | accuracy | 0.110276 |
| classification | lstm | Reaching C-J-M-T_1 | cross_day | 20150629__to__20150707 | 20150709__to__20150710 | 46 | accuracy | 0.110398 |
| classification | lstm | Reaching C-J-M-T_1 | cross_day | 20150629__to__20150707 | 20150709__to__20150710 | 54 | accuracy | 0.102757 |
| classification | lstm | Reaching C-J-M-T_1 | cross_day | 20150629__to__20150707 | 20150709__to__20150710 | 54 | accuracy | 0.102745 |
| classification | transformer | Reaching C-J-M-T_1 | cross_day | 20131003__to__20160914 | 20160915__to__20161021 | 31 | accuracy | 0.132567 |
| classification | transformer | Reaching C-J-M-T_1 | cross_day | 20131003__to__20160914 | 20160915__to__20161021 | 44 | accuracy | 0.126823 |
| classification | transformer | Reaching C-J-M-T_1 | cross_day | 20131003__to__20160914 | 20160915__to__20161021 | 46 | accuracy | 0.120636 |
| classification | transformer | Reaching C-J-M-T_1 | cross_day | 20131003__to__20160914 | 20160915__to__20161021 | 54 | accuracy | 0.116217 |
| classification | transformer | Reaching C-J-M-T_1 | cross_day | 20131003__to__20160914 | 20160915__to__20161021 | 69 | accuracy | 0.124171 |
| classification | transformer | Reaching C-J-M-T_1 | cross_day | 20150629__to__20150629 | 20150701__to__20150701 | 31 | accuracy | 0.142857 |
| classification | transformer | Reaching C-J-M-T_1 | cross_day | 20150629__to__20150629 | 20150701__to__20150701 | 44 | accuracy | 0.168367 |
| classification | transformer | Reaching C-J-M-T_1 | cross_day | 20150629__to__20150629 | 20150701__to__20150701 | 46 | accuracy | 0.142857 |
| classification | transformer | Reaching C-J-M-T_1 | cross_day | 20150629__to__20150707 | 20150709__to__20150710 | 44 | accuracy | 0.100251 |
| classification | transformer | Reaching C-J-M-T_1 | cross_day | 20150629__to__20150707 | 20150709__to__20150710 | 44 | accuracy | 0.100194 |
| classification | transformer | Reaching C-J-M-T_1 | cross_day | 20150629__to__20150707 | 20150709__to__20150710 | 46 | accuracy | 0.117794 |
| classification | transformer | Reaching C-J-M-T_1 | cross_day | 20150629__to__20150707 | 20150709__to__20150710 | 46 | accuracy | 0.117875 |
| classification | transformer | Reaching C-J-M-T_1 | cross_day | 20150629__to__20150707 | 20150709__to__20150710 | 54 | accuracy | 0.122807 |
| classification | transformer | Reaching C-J-M-T_1 | cross_day | 20150629__to__20150707 | 20150709__to__20150710 | 54 | accuracy | 0.122537 |
| classification | lfads | Reaching C-J-M-T_1 | cross_day | 20150629__to__20150707 | 20150709__to__20150710 | 44 | accuracy | 0.175439 |
| classification | lfads | Reaching C-J-M-T_1 | cross_day | 20150629__to__20150707 | 20150709__to__20150710 | 44 | accuracy | 0.159395 |
| classification | lfads | Reaching C-J-M-T_1 | cross_day | 20150629__to__20150707 | 20150709__to__20150710 | 46 | accuracy | 0.170426 |
| classification | lfads | Reaching C-J-M-T_1 | cross_day | 20150629__to__20150707 | 20150709__to__20150710 | 46 | accuracy | 0.246833 |
| classification | lfads | Reaching C-J-M-T_1 | cross_day | 20150629__to__20150707 | 20150709__to__20150710 | 54 | accuracy | 0.270677 |
| classification | lfads | Reaching C-J-M-T_1 | cross_day | 20150629__to__20150707 | 20150709__to__20150710 | 54 | accuracy | 0.214462 |
| classification | cycle_gan | Reaching C-J-M-T_1 | cross_day | 20131003__to__20160914 | 20160915__to__20161021 | 31 | accuracy | 0.115776 |
| classification | cycle_gan | Reaching C-J-M-T_1 | cross_day | 20131003__to__20160914 | 20160915__to__20161021 | 44 | accuracy | 0.118427 |
| classification | cycle_gan | Reaching C-J-M-T_1 | cross_day | 20131003__to__20160914 | 20160915__to__20161021 | 46 | accuracy | 0.124613 |
| classification | cycle_gan | Reaching C-J-M-T_1 | cross_day | 20131003__to__20160914 | 20160915__to__20161021 | 54 | accuracy | 0.112682 |
| classification | cycle_gan | Reaching C-J-M-T_1 | cross_day | 20131003__to__20160914 | 20160915__to__20161021 | 69 | accuracy | 0.122404 |
| classification | cycle_gan | Reaching C-J-M-T_1 | cross_day | 20150629__to__20150629 | 20150701__to__20150701 | 31 | accuracy | 0.142857 |
| classification | cycle_gan | Reaching C-J-M-T_1 | cross_day | 20150629__to__20150629 | 20150701__to__20150701 | 44 | accuracy | 0.142857 |
| classification | cycle_gan | Reaching C-J-M-T_1 | cross_day | 20150629__to__20150707 | 20150709__to__20150710 | 44 | accuracy | 0.135501 |
| classification | cycle_gan | Reaching C-J-M-T_1 | cross_day | 20150629__to__20150707 | 20150709__to__20150710 | 44 | accuracy | 0.108132 |
| classification | cycle_gan | Reaching C-J-M-T_1 | cross_day | 20150629__to__20150707 | 20150709__to__20150710 | 44 | accuracy | 0.127535 |
| classification | cycle_gan | Reaching C-J-M-T_1 | cross_day | 20150629__to__20150707 | 20150709__to__20150710 | 46 | accuracy | 0.168022 |

| | | | | | | | | |
|---|---|---|---|---|---|---|---|---|
| classification | cycle_gan | Reaching C-J-M-T_1 | cross_day | 20150629__to__20150707 | 20150709__to__20150710 | 46 | accuracy | 0.119524 |
| classification | cycle_gan | Reaching C-J-M-T_1 | cross_day | 20150629__to__20150707 | 20150709__to__20150710 | 46 | accuracy | 0.111235 |
| classification | cycle_gan | Reaching C-J-M-T_1 | cross_day | 20150629__to__20150707 | 20150709__to__20150710 | 54 | accuracy | 0.132791 |
| classification | cycle_gan | Reaching C-J-M-T_1 | cross_day | 20150629__to__20150707 | 20150709__to__20150710 | 54 | accuracy | 0.158701 |
| classification | cycle_gan | Reaching C-J-M-T_1 | cross_day | 20150629__to__20150707 | 20150709__to__20150710 | 54 | accuracy | 0.161526 |
| classification | stabilization | Reaching C-J-M-T_1 | cross_day | 20150629__to__20150707 | 20150709__to__20150710 | 44 | accuracy | 0.371274 |
| classification | stabilization | Reaching C-J-M-T_1 | cross_day | 20150629__to__20150707 | 20150709__to__20150710 | 44 | accuracy | 0.258451 |
| classification | stabilization | Reaching C-J-M-T_1 | cross_day | 20150629__to__20150707 | 20150709__to__20150710 | 44 | accuracy | 0.33338 |
| classification | stabilization | Reaching C-J-M-T_1 | cross_day | 20150629__to__20150707 | 20150709__to__20150710 | 46 | accuracy | 0.287263 |
| classification | stabilization | Reaching C-J-M-T_1 | cross_day | 20150629__to__20150707 | 20150709__to__20150710 | 46 | accuracy | 0.185931 |
| classification | stabilization | Reaching C-J-M-T_1 | cross_day | 20150629__to__20150707 | 20150709__to__20150710 | 46 | accuracy | 0.308789 |
| classification | stabilization | Reaching C-J-M-T_1 | cross_day | 20150629__to__20150707 | 20150709__to__20150710 | 54 | accuracy | 0.311653 |
| classification | stabilization | Reaching C-J-M-T_1 | cross_day | 20150629__to__20150707 | 20150709__to__20150710 | 54 | accuracy | 0.271464 |
| classification | stabilization | Reaching C-J-M-T_1 | cross_day | 20150629__to__20150707 | 20150709__to__20150710 | 54 | accuracy | 0.320182 |
| classification | mfsnn | Reaching C-J-M-T_1 | cross_day | 20150629__to__20150707 | 20150709 | 44 | accuracy | 0.244898 |
| classification | mfsnn | Reaching C-J-M-T_1 | cross_day | 20150629__to__20150707 | 20150709 | 46 | accuracy | 0.122449 |
| classification | mfsnn | Reaching C-J-M-T_1 | cross_day | 20150629__to__20150707 | 20150709 | 54 | accuracy | 0.147959 |
| classification | mfsnn | Reaching C-J-M-T_1 | cross_day | 20150629__to__20150707 | 20150710 | 44 | accuracy | 0.133005 |
| classification | mfsnn | Reaching C-J-M-T_1 | cross_day | 20150629__to__20150707 | 20150710 | 46 | accuracy | 0.133005 |
| classification | mfsnn | Reaching C-J-M-T_1 | cross_day | 20150629__to__20150707 | 20150710 | 54 | accuracy | 0.182266 |
| classification | mscformer | Reaching C-J-M-T_1 | cross_day | 20150629__to__20150707 | 20150709 | 44 | accuracy | 0.0867347 |
| classification | mscformer | Reaching C-J-M-T_1 | cross_day | 20150629__to__20150707 | 20150709 | 46 | accuracy | 0.132653 |
| classification | mscformer | Reaching C-J-M-T_1 | cross_day | 20150629__to__20150707 | 20150709 | 54 | accuracy | 0.163265 |
| classification | mscformer | Reaching C-J-M-T_1 | cross_day | 20150629__to__20150707 | 20150710 | 44 | accuracy | 0.17734 |
| classification | mscformer | Reaching C-J-M-T_1 | cross_day | 20150629__to__20150707 | 20150710 | 46 | accuracy | 0.236453 |
| classification | mscformer | Reaching C-J-M-T_1 | cross_day | 20150629__to__20150707 | 20150710 | 54 | accuracy | 0.197044 |
| classification | nomad | Reaching C-J-M-T_1 | cross_day | 20150629__to__20150707 | 20150709 | 44 | accuracy | 0.153061 |
| classification | nomad | Reaching C-J-M-T_1 | cross_day | 20150629__to__20150707 | 20150709 | 46 | accuracy | 0.0969388 |
| classification | nomad | Reaching C-J-M-T_1 | cross_day | 20150629__to__20150707 | 20150709 | 54 | accuracy | 0.132653 |
| classification | nomad | Reaching C-J-M-T_1 | cross_day | 20150629__to__20150707 | 20150710 | 44 | accuracy | 0.251232 |
| classification | nomad | Reaching C-J-M-T_1 | cross_day | 20150629__to__20150707 | 20150710 | 46 | accuracy | 0.192118 |
| classification | nomad | Reaching C-J-M-T_1 | cross_day | 20150629__to__20150707 | 20150710 | 54 | accuracy | 0.182266 |
| classification | mlp | Reaching C-J-M-T_1 | cross_day | 20131003__to__20151119 | 20160915 | 44 | accuracy | 0.10989 |
| classification | mlp | Reaching C-J-M-T_1 | cross_day | 20131003__to__20151119 | 20160915 | 46 | accuracy | 0.148352 |
| classification | mlp | Reaching C-J-M-T_1 | cross_day | 20131003__to__20151119 | 20160915 | 54 | accuracy | 0.120879 |
| classification | mlp | Reaching C-J-M-T_1 | cross_day | 20131003__to__20151119 | 20160919 | 44 | accuracy | 0.274194 |
| classification | mlp | Reaching C-J-M-T_1 | cross_day | 20131003__to__20151119 | 20160919 | 46 | accuracy | 0.252688 |
| classification | mlp | Reaching C-J-M-T_1 | cross_day | 20131003__to__20151119 | 20160919 | 54 | accuracy | 0.263441 |
| classification | mlp | Reaching C-J-M-T_1 | cross_day | 20131003__to__20151119 | 20160921 | 44 | accuracy | 0.147727 |
| classification | mlp | Reaching C-J-M-T_1 | cross_day | 20131003__to__20151119 | 20160921 | 46 | accuracy | 0.147727 |
| classification | mlp | Reaching C-J-M-T_1 | cross_day | 20131003__to__20151119 | 20160921 | 54 | accuracy | 0.181818 |
| classification | mlp | Reaching C-J-M-T_1 | cross_day | 20131003__to__20151119 | 20160923 | 44 | accuracy | 0.133005 |
| classification | mlp | Reaching C-J-M-T_1 | cross_day | 20131003__to__20151119 | 20160923 | 46 | accuracy | 0.182266 |
| classification | mlp | Reaching C-J-M-T_1 | cross_day | 20131003__to__20151119 | 20160923 | 54 | accuracy | 0.152709 |
| classification | mlp | Reaching C-J-M-T_1 | cross_day | 20131003__to__20151119 | 20160929 | 44 | accuracy | 0.0956938 |
| classification | mlp | Reaching C-J-M-T_1 | cross_day | 20131003__to__20151119 | 20160929 | 46 | accuracy | 0.0813397 |
| classification | mlp | Reaching C-J-M-T_1 | cross_day | 20131003__to__20151119 | 20160929 | 54 | accuracy | 0.076555 |
| classification | mlp | Reaching C-J-M-T_1 | cross_day | 20131003__to__20151119 | 20161005 | 44 | accuracy | 0.0891089 |
| classification | mlp | Reaching C-J-M-T_1 | cross_day | 20131003__to__20151119 | 20161005 | 46 | accuracy | 0.108911 |
| classification | mlp | Reaching C-J-M-T_1 | cross_day | 20131003__to__20151119 | 20161005 | 54 | accuracy | 0.108911 |
| classification | mlp | Reaching C-J-M-T_1 | cross_day | 20131003__to__20151119 | 20161006 | 44 | accuracy | 0.0904762 |
| classification | mlp | Reaching C-J-M-T_1 | cross_day | 20131003__to__20151119 | 20161006 | 46 | accuracy | 0.0857143 |
| classification | mlp | Reaching C-J-M-T_1 | cross_day | 20131003__to__20151119 | 20161006 | 54 | accuracy | 0.0904762 |
| classification | mlp | Reaching C-J-M-T_1 | cross_day | 20131003__to__20151119 | 20161007 | 44 | accuracy | 0.0935673 |
| classification | mlp | Reaching C-J-M-T_1 | cross_day | 20131003__to__20151119 | 20161007 | 46 | accuracy | 0.111111 |
| classification | mlp | Reaching C-J-M-T_1 | cross_day | 20131003__to__20151119 | 20161007 | 54 | accuracy | 0.0877193 |
| classification | mlp | Reaching C-J-M-T_1 | cross_day | 20131003__to__20151119 | 20161011 | 44 | accuracy | 0.175879 |
| classification | mlp | Reaching C-J-M-T_1 | cross_day | 20131003__to__20151119 | 20161011 | 46 | accuracy | 0.19598 |
| classification | mlp | Reaching C-J-M-T_1 | cross_day | 20131003__to__20151119 | 20161011 | 54 | accuracy | 0.135678 |
| classification | mlp | Reaching C-J-M-T_1 | cross_day | 20131003__to__20151119 | 20161013 | 44 | accuracy | 0.0962343 |
| classification | mlp | Reaching C-J-M-T_1 | cross_day | 20131003__to__20151119 | 20161013 | 46 | accuracy | 0.108787 |
| classification | mlp | Reaching C-J-M-T_1 | cross_day | 20131003__to__20151119 | 20161013 | 54 | accuracy | 0.117155 |
| classification | mlp | Reaching C-J-M-T_1 | cross_day | 20131003__to__20151119 | 20161021 | 44 | accuracy | 0.0909091 |

| | | | | | | | | |
|---|---|---|---|---|---|---|---|---|
| classification | mlp | Reaching C-J-M-T_1 | cross_day | 20131003__to__20151119 | 20161021 | 46 | accuracy | 0.122378 |
| classification | mlp | Reaching C-J-M-T_1 | cross_day | 20131003__to__20151119 | 20161021 | 54 | accuracy | 0.122378 |
| classification | convpi_vae | Reaching C-J-M-T_1 | cross_day | 20131003__to__20160914 | 20160915__to__20161021 | 31 | accuracy | 1 |
| classification | convpi_vae | Reaching C-J-M-T_1 | cross_day | 20131003__to__20160914 | 20160915__to__20161021 | 44 | accuracy | 1 |
| classification | convpi_vae | Reaching C-J-M-T_1 | cross_day | 20131003__to__20160914 | 20160915__to__20161021 | 46 | accuracy | 1 |
| classification | convpi_vae | Reaching C-J-M-T_1 | cross_day | 20131003__to__20160914 | 20160915__to__20161021 | 54 | accuracy | 1 |
| classification | convpi_vae | Reaching C-J-M-T_1 | cross_day | 20131003__to__20160914 | 20160915__to__20161021 | 69 | accuracy | 1 |
| classification | convpi_vae | Reaching C-J-M-T_1 | cross_day | 20150629__to__20150629 | 20150701__to__20150701 | 31 | accuracy | 0.153061 |
| classification | convpi_vae | Reaching C-J-M-T_1 | cross_day | 20150629__to__20150629 | 20150701__to__20150701 | 44 | accuracy | 0.137755 |
| classification | lstm | Reaching B | single_day | GY_data_day-1 | GY_data_day-1 | 31 | accuracy | 0.870588 |
| classification | lstm | Reaching B | single_day | GY_data_day-1 | GY_data_day-1 | 44 | accuracy | 0.870588 |
| classification | lstm | Reaching B | single_day | GY_data_day-1 | GY_data_day-1 | 44 | accuracy | 0.935294 |
| classification | lstm | Reaching B | single_day | GY_data_day-1 | GY_data_day-1 | 46 | accuracy | 0.905882 |
| classification | lstm | Reaching B | single_day | GY_data_day-1 | GY_data_day-1 | 46 | accuracy | 0.923529 |
| classification | lstm | Reaching B | single_day | GY_data_day-1 | GY_data_day-1 | 54 | accuracy | 0.911765 |
| classification | lstm | Reaching B | single_day | GY_data_day-1 | GY_data_day-1 | 54 | accuracy | 0.917647 |
| classification | lstm | Reaching B | single_day | GY_data_day-1 | GY_data_day-1 | 69 | accuracy | 0.947059 |
| classification | lstm | Reaching B | single_day | GY_data_day-10 | GY_data_day-10 | 31 | accuracy | 0.65625 |
| classification | lstm | Reaching B | single_day | GY_data_day-10 | GY_data_day-10 | 44 | accuracy | 0.71875 |
| classification | lstm | Reaching B | single_day | GY_data_day-10 | GY_data_day-10 | 44 | accuracy | 0.70625 |
| classification | lstm | Reaching B | single_day | GY_data_day-10 | GY_data_day-10 | 46 | accuracy | 0.625 |
| classification | lstm | Reaching B | single_day | GY_data_day-10 | GY_data_day-10 | 46 | accuracy | 0.725 |
| classification | lstm | Reaching B | single_day | GY_data_day-10 | GY_data_day-10 | 54 | accuracy | 0.69375 |
| classification | lstm | Reaching B | single_day | GY_data_day-10 | GY_data_day-10 | 54 | accuracy | 0.7 |
| classification | lstm | Reaching B | single_day | GY_data_day-10 | GY_data_day-10 | 69 | accuracy | 0.70625 |
| classification | lstm | Reaching B | single_day | GY_data_day-11 | GY_data_day-11 | 31 | accuracy | 0.9625 |
| classification | lstm | Reaching B | single_day | GY_data_day-11 | GY_data_day-11 | 44 | accuracy | 0.9125 |
| classification | lstm | Reaching B | single_day | GY_data_day-11 | GY_data_day-11 | 44 | accuracy | 0.9375 |
| classification | lstm | Reaching B | single_day | GY_data_day-11 | GY_data_day-11 | 46 | accuracy | 0.8625 |
| classification | lstm | Reaching B | single_day | GY_data_day-11 | GY_data_day-11 | 46 | accuracy | 0.9375 |
| classification | lstm | Reaching B | single_day | GY_data_day-11 | GY_data_day-11 | 54 | accuracy | 0.85 |
| classification | lstm | Reaching B | single_day | GY_data_day-11 | GY_data_day-11 | 54 | accuracy | 0.9375 |
| classification | lstm | Reaching B | single_day | GY_data_day-11 | GY_data_day-11 | 69 | accuracy | 0.925 |
| classification | lstm | Reaching B | single_day | GY_data_day-12 | GY_data_day-12 | 31 | accuracy | 0.924419 |
| classification | lstm | Reaching B | single_day | GY_data_day-12 | GY_data_day-12 | 44 | accuracy | 0.936047 |
| classification | lstm | Reaching B | single_day | GY_data_day-12 | GY_data_day-12 | 44 | accuracy | 0.982558 |
| classification | lstm | Reaching B | single_day | GY_data_day-12 | GY_data_day-12 | 46 | accuracy | 0.97093 |
| classification | lstm | Reaching B | single_day | GY_data_day-12 | GY_data_day-12 | 46 | accuracy | 0.965116 |
| classification | lstm | Reaching B | single_day | GY_data_day-12 | GY_data_day-12 | 54 | accuracy | 0.854651 |
| classification | lstm | Reaching B | single_day | GY_data_day-12 | GY_data_day-12 | 54 | accuracy | 0.959302 |
| classification | lstm | Reaching B | single_day | GY_data_day-12 | GY_data_day-12 | 69 | accuracy | 0.877907 |
| classification | lstm | Reaching B | single_day | GY_data_day-13 | GY_data_day-13 | 31 | accuracy | 0.825 |
| classification | lstm | Reaching B | single_day | GY_data_day-13 | GY_data_day-13 | 44 | accuracy | 0.875 |
| classification | lstm | Reaching B | single_day | GY_data_day-13 | GY_data_day-13 | 46 | accuracy | 0.8375 |
| classification | lstm | Reaching B | single_day | GY_data_day-13 | GY_data_day-13 | 46 | accuracy | 0.7875 |
| classification | lstm | Reaching B | single_day | GY_data_day-13 | GY_data_day-13 | 54 | accuracy | 0.8875 |
| classification | lstm | Reaching B | single_day | GY_data_day-13 | GY_data_day-13 | 54 | accuracy | 0.7875 |
| classification | lstm | Reaching B | single_day | GY_data_day-13 | GY_data_day-13 | 69 | accuracy | 0.8625 |
| classification | lstm | Reaching B | single_day | GY_data_day-14 | GY_data_day-14 | 31 | accuracy | 0.866667 |
| classification | lstm | Reaching B | single_day | GY_data_day-14 | GY_data_day-14 | 44 | accuracy | 0.866667 |
| classification | lstm | Reaching B | single_day | GY_data_day-14 | GY_data_day-14 | 44 | accuracy | 0.925 |
| classification | lstm | Reaching B | single_day | GY_data_day-14 | GY_data_day-14 | 46 | accuracy | 0.941667 |
| classification | lstm | Reaching B | single_day | GY_data_day-14 | GY_data_day-14 | 46 | accuracy | 0.966667 |
| classification | lstm | Reaching B | single_day | GY_data_day-14 | GY_data_day-14 | 54 | accuracy | 0.858333 |
| classification | lstm | Reaching B | single_day | GY_data_day-14 | GY_data_day-14 | 54 | accuracy | 0.933333 |
| classification | lstm | Reaching B | single_day | GY_data_day-14 | GY_data_day-14 | 69 | accuracy | 0.875 |
| classification | lstm | Reaching B | single_day | GY_data_day-15 | GY_data_day-15 | 31 | accuracy | 0.98125 |
| classification | lstm | Reaching B | single_day | GY_data_day-15 | GY_data_day-15 | 44 | accuracy | 0.94375 |
| classification | lstm | Reaching B | single_day | GY_data_day-15 | GY_data_day-15 | 44 | accuracy | 0.9625 |
| classification | lstm | Reaching B | single_day | GY_data_day-15 | GY_data_day-15 | 46 | accuracy | 0.90625 |
| classification | lstm | Reaching B | single_day | GY_data_day-15 | GY_data_day-15 | 46 | accuracy | 0.96875 |
| classification | lstm | Reaching B | single_day | GY_data_day-15 | GY_data_day-15 | 54 | accuracy | 0.975 |
| classification | lstm | Reaching B | single_day | GY_data_day-15 | GY_data_day-15 | 54 | accuracy | 0.94375 |

| | | | | | | | | |
|---|---|---|---|---|---|---|---|---|
| classification | lstm | Reaching B | single_day | GY_data_day-15 | GY_data_day-15 | 69 | accuracy | 0.93125 |
| classification | lstm | Reaching B | single_day | GY_data_day-16 | GY_data_day-16 | 31 | accuracy | 0.675 |
| classification | lstm | Reaching B | single_day | GY_data_day-16 | GY_data_day-16 | 44 | accuracy | 0.725 |
| classification | lstm | Reaching B | single_day | GY_data_day-16 | GY_data_day-16 | 44 | accuracy | 0.775 |
| classification | lstm | Reaching B | single_day | GY_data_day-16 | GY_data_day-16 | 46 | accuracy | 0.575 |
| classification | lstm | Reaching B | single_day | GY_data_day-16 | GY_data_day-16 | 46 | accuracy | 0.725 |
| classification | lstm | Reaching B | single_day | GY_data_day-16 | GY_data_day-16 | 54 | accuracy | 0.625 |
| classification | lstm | Reaching B | single_day | GY_data_day-16 | GY_data_day-16 | 54 | accuracy | 0.8 |
| classification | lstm | Reaching B | single_day | GY_data_day-16 | GY_data_day-16 | 69 | accuracy | 0.625 |
| classification | lstm | Reaching B | single_day | GY_data_day-17 | GY_data_day-17 | 31 | accuracy | 0.983333 |
| classification | lstm | Reaching B | single_day | GY_data_day-17 | GY_data_day-17 | 44 | accuracy | 0.866667 |
| classification | lstm | Reaching B | single_day | GY_data_day-17 | GY_data_day-17 | 44 | accuracy | 0.925 |
| classification | lstm | Reaching B | single_day | GY_data_day-17 | GY_data_day-17 | 46 | accuracy | 0.916667 |
| classification | lstm | Reaching B | single_day | GY_data_day-17 | GY_data_day-17 | 46 | accuracy | 0.975 |
| classification | lstm | Reaching B | single_day | GY_data_day-17 | GY_data_day-17 | 54 | accuracy | 0.933333 |
| classification | lstm | Reaching B | single_day | GY_data_day-17 | GY_data_day-17 | 54 | accuracy | 0.908333 |
| classification | lstm | Reaching B | single_day | GY_data_day-17 | GY_data_day-17 | 69 | accuracy | 0.883333 |
| classification | lstm | Reaching B | single_day | GY_data_day-18 | GY_data_day-18 | 31 | accuracy | 0.9 |
| classification | lstm | Reaching B | single_day | GY_data_day-18 | GY_data_day-18 | 44 | accuracy | 0.9125 |
| classification | lstm | Reaching B | single_day | GY_data_day-18 | GY_data_day-18 | 44 | accuracy | 0.925 |
| classification | lstm | Reaching B | single_day | GY_data_day-18 | GY_data_day-18 | 46 | accuracy | 0.8875 |
| classification | lstm | Reaching B | single_day | GY_data_day-18 | GY_data_day-18 | 46 | accuracy | 0.95 |
| classification | lstm | Reaching B | single_day | GY_data_day-18 | GY_data_day-18 | 54 | accuracy | 0.875 |
| classification | lstm | Reaching B | single_day | GY_data_day-18 | GY_data_day-18 | 54 | accuracy | 0.95 |
| classification | lstm | Reaching B | single_day | GY_data_day-18 | GY_data_day-18 | 69 | accuracy | 0.925 |
| classification | lstm | Reaching B | single_day | GY_data_day-19 | GY_data_day-19 | 31 | accuracy | 0.825 |
| classification | lstm | Reaching B | single_day | GY_data_day-19 | GY_data_day-19 | 44 | accuracy | 0.9 |
| classification | lstm | Reaching B | single_day | GY_data_day-19 | GY_data_day-19 | 44 | accuracy | 0.9125 |
| classification | lstm | Reaching B | single_day | GY_data_day-19 | GY_data_day-19 | 46 | accuracy | 0.8375 |
| classification | lstm | Reaching B | single_day | GY_data_day-19 | GY_data_day-19 | 46 | accuracy | 0.9125 |
| classification | lstm | Reaching B | single_day | GY_data_day-19 | GY_data_day-19 | 54 | accuracy | 0.7125 |
| classification | lstm | Reaching B | single_day | GY_data_day-19 | GY_data_day-19 | 54 | accuracy | 0.825 |
| classification | lstm | Reaching B | single_day | GY_data_day-19 | GY_data_day-19 | 69 | accuracy | 0.8375 |
| classification | lstm | Reaching B | single_day | GY_data_day-2 | GY_data_day-2 | 31 | accuracy | 0.935 |
| classification | lstm | Reaching B | single_day | GY_data_day-2 | GY_data_day-2 | 44 | accuracy | 0.935 |
| classification | lstm | Reaching B | single_day | GY_data_day-2 | GY_data_day-2 | 44 | accuracy | 0.965 |
| classification | lstm | Reaching B | single_day | GY_data_day-2 | GY_data_day-2 | 46 | accuracy | 0.95 |
| classification | lstm | Reaching B | single_day | GY_data_day-2 | GY_data_day-2 | 46 | accuracy | 0.97 |
| classification | lstm | Reaching B | single_day | GY_data_day-2 | GY_data_day-2 | 54 | accuracy | 0.925 |
| classification | lstm | Reaching B | single_day | GY_data_day-2 | GY_data_day-2 | 54 | accuracy | 0.995 |
| classification | lstm | Reaching B | single_day | GY_data_day-2 | GY_data_day-2 | 69 | accuracy | 0.96 |
| classification | lstm | Reaching B | single_day | GY_data_day-20 | GY_data_day-20 | 31 | accuracy | 0.675 |
| classification | lstm | Reaching B | single_day | GY_data_day-20 | GY_data_day-20 | 44 | accuracy | 0.7 |
| classification | lstm | Reaching B | single_day | GY_data_day-20 | GY_data_day-20 | 44 | accuracy | 0.85 |
| classification | lstm | Reaching B | single_day | GY_data_day-20 | GY_data_day-20 | 46 | accuracy | 0.75 |
| classification | lstm | Reaching B | single_day | GY_data_day-20 | GY_data_day-20 | 46 | accuracy | 0.9 |
| classification | lstm | Reaching B | single_day | GY_data_day-20 | GY_data_day-20 | 54 | accuracy | 0.475 |
| classification | lstm | Reaching B | single_day | GY_data_day-20 | GY_data_day-20 | 54 | accuracy | 0.85 |
| classification | lstm | Reaching B | single_day | GY_data_day-20 | GY_data_day-20 | 69 | accuracy | 0.75 |
| classification | lstm | Reaching B | single_day | GY_data_day-21 | GY_data_day-21 | 31 | accuracy | 0.458333 |
| classification | lstm | Reaching B | single_day | GY_data_day-21 | GY_data_day-21 | 44 | accuracy | 0.483333 |
| classification | lstm | Reaching B | single_day | GY_data_day-21 | GY_data_day-21 | 44 | accuracy | 0.583333 |
| classification | lstm | Reaching B | single_day | GY_data_day-21 | GY_data_day-21 | 46 | accuracy | 0.516667 |
| classification | lstm | Reaching B | single_day | GY_data_day-21 | GY_data_day-21 | 46 | accuracy | 0.575 |
| classification | lstm | Reaching B | single_day | GY_data_day-21 | GY_data_day-21 | 54 | accuracy | 0.533333 |
| classification | lstm | Reaching B | single_day | GY_data_day-21 | GY_data_day-21 | 69 | accuracy | 0.475 |
| classification | lstm | Reaching B | single_day | GY_data_day-22 | GY_data_day-22 | 31 | accuracy | 0.875 |
| classification | lstm | Reaching B | single_day | GY_data_day-22 | GY_data_day-22 | 44 | accuracy | 0.960714 |
| classification | lstm | Reaching B | single_day | GY_data_day-22 | GY_data_day-22 | 44 | accuracy | 0.975 |
| classification | lstm | Reaching B | single_day | GY_data_day-22 | GY_data_day-22 | 46 | accuracy | 0.964286 |
| classification | lstm | Reaching B | single_day | GY_data_day-22 | GY_data_day-22 | 46 | accuracy | 0.971429 |
| classification | lstm | Reaching B | single_day | GY_data_day-22 | GY_data_day-22 | 54 | accuracy | 0.907143 |
| classification | lstm | Reaching B | single_day | GY_data_day-22 | GY_data_day-22 | 54 | accuracy | 0.953571 |

| | | | | | | | | |
|---|---|---|---|---|---|---|---|---|
| classification | lstm | Reaching B | single_day | GY_data_day-22 | GY_data_day-22 | 69 | accuracy | 0.882143 |
| classification | lstm | Reaching B | single_day | GY_data_day-23 | GY_data_day-23 | 31 | accuracy | 0.93 |
| classification | lstm | Reaching B | single_day | GY_data_day-23 | GY_data_day-23 | 44 | accuracy | 0.93 |
| classification | lstm | Reaching B | single_day | GY_data_day-23 | GY_data_day-23 | 44 | accuracy | 0.95 |
| classification | lstm | Reaching B | single_day | GY_data_day-23 | GY_data_day-23 | 46 | accuracy | 0.875 |
| classification | lstm | Reaching B | single_day | GY_data_day-23 | GY_data_day-23 | 46 | accuracy | 0.935 |
| classification | lstm | Reaching B | single_day | GY_data_day-23 | GY_data_day-23 | 54 | accuracy | 0.925 |
| classification | lstm | Reaching B | single_day | GY_data_day-23 | GY_data_day-23 | 54 | accuracy | 0.93 |
| classification | lstm | Reaching B | single_day | GY_data_day-23 | GY_data_day-23 | 69 | accuracy | 0.905 |
| classification | lstm | Reaching B | single_day | GY_data_day-24 | GY_data_day-24 | 31 | accuracy | 0.7375 |
| classification | lstm | Reaching B | single_day | GY_data_day-24 | GY_data_day-24 | 44 | accuracy | 0.675 |
| classification | lstm | Reaching B | single_day | GY_data_day-24 | GY_data_day-24 | 44 | accuracy | 0.85 |
| classification | lstm | Reaching B | single_day | GY_data_day-24 | GY_data_day-24 | 46 | accuracy | 0.875 |
| classification | lstm | Reaching B | single_day | GY_data_day-24 | GY_data_day-24 | 46 | accuracy | 0.8625 |
| classification | lstm | Reaching B | single_day | GY_data_day-24 | GY_data_day-24 | 54 | accuracy | 0.7625 |
| classification | lstm | Reaching B | single_day | GY_data_day-24 | GY_data_day-24 | 54 | accuracy | 0.875 |
| classification | lstm | Reaching B | single_day | GY_data_day-24 | GY_data_day-24 | 69 | accuracy | 0.8125 |
| classification | lstm | Reaching B | single_day | GY_data_day-25 | GY_data_day-25 | 31 | accuracy | 0.8625 |
| classification | lstm | Reaching B | single_day | GY_data_day-25 | GY_data_day-25 | 44 | accuracy | 0.7875 |
| classification | lstm | Reaching B | single_day | GY_data_day-25 | GY_data_day-25 | 44 | accuracy | 0.8875 |
| classification | lstm | Reaching B | single_day | GY_data_day-25 | GY_data_day-25 | 46 | accuracy | 0.8875 |
| classification | lstm | Reaching B | single_day | GY_data_day-25 | GY_data_day-25 | 46 | accuracy | 0.825 |
| classification | lstm | Reaching B | single_day | GY_data_day-25 | GY_data_day-25 | 54 | accuracy | 0.8375 |
| classification | lstm | Reaching B | single_day | GY_data_day-25 | GY_data_day-25 | 54 | accuracy | 0.7875 |
| classification | lstm | Reaching B | single_day | GY_data_day-25 | GY_data_day-25 | 69 | accuracy | 0.85 |
| classification | lstm | Reaching B | single_day | GY_data_day-26 | GY_data_day-26 | 31 | accuracy | 0.7375 |
| classification | lstm | Reaching B | single_day | GY_data_day-26 | GY_data_day-26 | 44 | accuracy | 0.725 |
| classification | lstm | Reaching B | single_day | GY_data_day-26 | GY_data_day-26 | 44 | accuracy | 0.825 |
| classification | lstm | Reaching B | single_day | GY_data_day-26 | GY_data_day-26 | 46 | accuracy | 0.85 |
| classification | lstm | Reaching B | single_day | GY_data_day-26 | GY_data_day-26 | 46 | accuracy | 0.7875 |
| classification | lstm | Reaching B | single_day | GY_data_day-26 | GY_data_day-26 | 54 | accuracy | 0.5875 |
| classification | lstm | Reaching B | single_day | GY_data_day-26 | GY_data_day-26 | 54 | accuracy | 0.825 |
| classification | lstm | Reaching B | single_day | GY_data_day-26 | GY_data_day-26 | 69 | accuracy | 0.85 |
| classification | lstm | Reaching B | single_day | GY_data_day-27 | GY_data_day-27 | 31 | accuracy | 0.8875 |
| classification | lstm | Reaching B | single_day | GY_data_day-27 | GY_data_day-27 | 44 | accuracy | 0.8375 |
| classification | lstm | Reaching B | single_day | GY_data_day-27 | GY_data_day-27 | 44 | accuracy | 0.95 |
| classification | lstm | Reaching B | single_day | GY_data_day-27 | GY_data_day-27 | 46 | accuracy | 0.75 |
| classification | lstm | Reaching B | single_day | GY_data_day-27 | GY_data_day-27 | 46 | accuracy | 0.9375 |
| classification | lstm | Reaching B | single_day | GY_data_day-27 | GY_data_day-27 | 54 | accuracy | 0.75 |
| classification | lstm | Reaching B | single_day | GY_data_day-27 | GY_data_day-27 | 54 | accuracy | 0.9 |
| classification | lstm | Reaching B | single_day | GY_data_day-27 | GY_data_day-27 | 69 | accuracy | 0.85 |
| classification | lstm | Reaching B | single_day | GY_data_day-28 | GY_data_day-28 | 31 | accuracy | 0.85 |
| classification | lstm | Reaching B | single_day | GY_data_day-28 | GY_data_day-28 | 44 | accuracy | 0.6875 |
| classification | lstm | Reaching B | single_day | GY_data_day-28 | GY_data_day-28 | 44 | accuracy | 0.9625 |
| classification | lstm | Reaching B | single_day | GY_data_day-28 | GY_data_day-28 | 46 | accuracy | 0.725 |
| classification | lstm | Reaching B | single_day | GY_data_day-28 | GY_data_day-28 | 46 | accuracy | 0.8375 |
| classification | lstm | Reaching B | single_day | GY_data_day-28 | GY_data_day-28 | 54 | accuracy | 0.775 |
| classification | lstm | Reaching B | single_day | GY_data_day-28 | GY_data_day-28 | 54 | accuracy | 0.9125 |
| classification | lstm | Reaching B | single_day | GY_data_day-28 | GY_data_day-28 | 69 | accuracy | 0.75 |
| classification | lstm | Reaching B | single_day | GY_data_day-29 | GY_data_day-29 | 31 | accuracy | 0.775 |
| classification | lstm | Reaching B | single_day | GY_data_day-29 | GY_data_day-29 | 44 | accuracy | 0.85 |
| classification | lstm | Reaching B | single_day | GY_data_day-29 | GY_data_day-29 | 44 | accuracy | 0.8625 |
| classification | lstm | Reaching B | single_day | GY_data_day-29 | GY_data_day-29 | 46 | accuracy | 0.8 |
| classification | lstm | Reaching B | single_day | GY_data_day-29 | GY_data_day-29 | 46 | accuracy | 0.875 |
| classification | lstm | Reaching B | single_day | GY_data_day-29 | GY_data_day-29 | 54 | accuracy | 0.725 |
| classification | lstm | Reaching B | single_day | GY_data_day-29 | GY_data_day-29 | 54 | accuracy | 0.85 |
| classification | lstm | Reaching B | single_day | GY_data_day-29 | GY_data_day-29 | 69 | accuracy | 0.875 |
| classification | lstm | Reaching B | single_day | GY_data_day-3 | GY_data_day-3 | 31 | accuracy | 0.9 |
| classification | lstm | Reaching B | single_day | GY_data_day-3 | GY_data_day-3 | 44 | accuracy | 0.925 |
| classification | lstm | Reaching B | single_day | GY_data_day-3 | GY_data_day-3 | 44 | accuracy | 0.958333 |
| classification | lstm | Reaching B | single_day | GY_data_day-3 | GY_data_day-3 | 46 | accuracy | 0.683333 |
| classification | lstm | Reaching B | single_day | GY_data_day-3 | GY_data_day-3 | 46 | accuracy | 0.883333 |
| classification | lstm | Reaching B | single_day | GY_data_day-3 | GY_data_day-3 | 54 | accuracy | 0.883333 |

| | | | | | | | | |
|---|---|---|---|---|---|---|---|---|
| classification | lstm | Reaching B | single_day | GY_data_day-3 | GY_data_day-3 | 54 | accuracy | 0.866667 |
| classification | lstm | Reaching B | single_day | GY_data_day-3 | GY_data_day-3 | 69 | accuracy | 0.941667 |
| classification | lstm | Reaching B | single_day | GY_data_day-30 | GY_data_day-30 | 31 | accuracy | 0.7125 |
| classification | lstm | Reaching B | single_day | GY_data_day-30 | GY_data_day-30 | 44 | accuracy | 0.775 |
| classification | lstm | Reaching B | single_day | GY_data_day-30 | GY_data_day-30 | 44 | accuracy | 0.875 |
| classification | lstm | Reaching B | single_day | GY_data_day-30 | GY_data_day-30 | 46 | accuracy | 0.7375 |
| classification | lstm | Reaching B | single_day | GY_data_day-30 | GY_data_day-30 | 46 | accuracy | 0.8625 |
| classification | lstm | Reaching B | single_day | GY_data_day-30 | GY_data_day-30 | 54 | accuracy | 0.85 |
| classification | lstm | Reaching B | single_day | GY_data_day-30 | GY_data_day-30 | 54 | accuracy | 0.875 |
| classification | lstm | Reaching B | single_day | GY_data_day-30 | GY_data_day-30 | 69 | accuracy | 0.85 |
| classification | lstm | Reaching B | single_day | GY_data_day-31 | GY_data_day-31 | 31 | accuracy | 0.6 |
| classification | lstm | Reaching B | single_day | GY_data_day-31 | GY_data_day-31 | 44 | accuracy | 0.625 |
| classification | lstm | Reaching B | single_day | GY_data_day-31 | GY_data_day-31 | 44 | accuracy | 0.575 |
| classification | lstm | Reaching B | single_day | GY_data_day-31 | GY_data_day-31 | 46 | accuracy | 0.5 |
| classification | lstm | Reaching B | single_day | GY_data_day-31 | GY_data_day-31 | 46 | accuracy | 0.65 |
| classification | lstm | Reaching B | single_day | GY_data_day-31 | GY_data_day-31 | 54 | accuracy | 0.55 |
| classification | lstm | Reaching B | single_day | GY_data_day-31 | GY_data_day-31 | 54 | accuracy | 0.625 |
| classification | lstm | Reaching B | single_day | GY_data_day-31 | GY_data_day-31 | 69 | accuracy | 0.55 |
| classification | lstm | Reaching B | single_day | GY_data_day-32 | GY_data_day-32 | 31 | accuracy | 0.7625 |
| classification | lstm | Reaching B | single_day | GY_data_day-32 | GY_data_day-32 | 44 | accuracy | 0.7625 |
| classification | lstm | Reaching B | single_day | GY_data_day-32 | GY_data_day-32 | 44 | accuracy | 0.875 |
| classification | lstm | Reaching B | single_day | GY_data_day-32 | GY_data_day-32 | 46 | accuracy | 0.8625 |
| classification | lstm | Reaching B | single_day | GY_data_day-32 | GY_data_day-32 | 46 | accuracy | 0.9 |
| classification | lstm | Reaching B | single_day | GY_data_day-32 | GY_data_day-32 | 54 | accuracy | 0.675 |
| classification | lstm | Reaching B | single_day | GY_data_day-32 | GY_data_day-32 | 54 | accuracy | 0.825 |
| classification | lstm | Reaching B | single_day | GY_data_day-32 | GY_data_day-32 | 69 | accuracy | 0.7375 |
| classification | lstm | Reaching B | single_day | GY_data_day-33 | GY_data_day-33 | 31 | accuracy | 0.725 |
| classification | lstm | Reaching B | single_day | GY_data_day-33 | GY_data_day-33 | 44 | accuracy | 0.733333 |
| classification | lstm | Reaching B | single_day | GY_data_day-33 | GY_data_day-33 | 44 | accuracy | 0.816667 |
| classification | lstm | Reaching B | single_day | GY_data_day-33 | GY_data_day-33 | 46 | accuracy | 0.833333 |
| classification | lstm | Reaching B | single_day | GY_data_day-33 | GY_data_day-33 | 54 | accuracy | 0.75 |
| classification | lstm | Reaching B | single_day | GY_data_day-33 | GY_data_day-33 | 54 | accuracy | 0.733333 |
| classification | lstm | Reaching B | single_day | GY_data_day-33 | GY_data_day-33 | 69 | accuracy | 0.616667 |
| classification | lstm | Reaching B | single_day | GY_data_day-34 | GY_data_day-34 | 31 | accuracy | 0.83125 |
| classification | lstm | Reaching B | single_day | GY_data_day-34 | GY_data_day-34 | 44 | accuracy | 0.91875 |
| classification | lstm | Reaching B | single_day | GY_data_day-34 | GY_data_day-34 | 44 | accuracy | 0.925 |
| classification | lstm | Reaching B | single_day | GY_data_day-34 | GY_data_day-34 | 46 | accuracy | 0.93125 |
| classification | lstm | Reaching B | single_day | GY_data_day-34 | GY_data_day-34 | 46 | accuracy | 0.91875 |
| classification | lstm | Reaching B | single_day | GY_data_day-34 | GY_data_day-34 | 54 | accuracy | 0.83125 |
| classification | lstm | Reaching B | single_day | GY_data_day-34 | GY_data_day-34 | 54 | accuracy | 0.85 |
| classification | lstm | Reaching B | single_day | GY_data_day-34 | GY_data_day-34 | 69 | accuracy | 0.81875 |
| classification | lstm | Reaching B | single_day | GY_data_day-35 | GY_data_day-35 | 31 | accuracy | 0.79375 |
| classification | lstm | Reaching B | single_day | GY_data_day-35 | GY_data_day-35 | 44 | accuracy | 0.7 |
| classification | lstm | Reaching B | single_day | GY_data_day-35 | GY_data_day-35 | 44 | accuracy | 0.90625 |
| classification | lstm | Reaching B | single_day | GY_data_day-35 | GY_data_day-35 | 46 | accuracy | 0.85625 |
| classification | lstm | Reaching B | single_day | GY_data_day-35 | GY_data_day-35 | 46 | accuracy | 0.89375 |
| classification | lstm | Reaching B | single_day | GY_data_day-35 | GY_data_day-35 | 54 | accuracy | 0.85625 |
| classification | lstm | Reaching B | single_day | GY_data_day-35 | GY_data_day-35 | 54 | accuracy | 0.83125 |
| classification | lstm | Reaching B | single_day | GY_data_day-35 | GY_data_day-35 | 69 | accuracy | 0.8625 |
| classification | lstm | Reaching B | single_day | GY_data_day-36 | GY_data_day-36 | 31 | accuracy | 0.825 |
| classification | lstm | Reaching B | single_day | GY_data_day-36 | GY_data_day-36 | 44 | accuracy | 0.88125 |
| classification | lstm | Reaching B | single_day | GY_data_day-36 | GY_data_day-36 | 44 | accuracy | 0.93125 |
| classification | lstm | Reaching B | single_day | GY_data_day-36 | GY_data_day-36 | 46 | accuracy | 0.8625 |
| classification | lstm | Reaching B | single_day | GY_data_day-36 | GY_data_day-36 | 46 | accuracy | 0.91875 |
| classification | lstm | Reaching B | single_day | GY_data_day-36 | GY_data_day-36 | 54 | accuracy | 0.875 |
| classification | lstm | Reaching B | single_day | GY_data_day-36 | GY_data_day-36 | 54 | accuracy | 0.8875 |
| classification | lstm | Reaching B | single_day | GY_data_day-36 | GY_data_day-36 | 69 | accuracy | 0.86875 |
| classification | lstm | Reaching B | single_day | GY_data_day-37 | GY_data_day-37 | 31 | accuracy | 0.9 |
| classification | lstm | Reaching B | single_day | GY_data_day-37 | GY_data_day-37 | 44 | accuracy | 0.9125 |
| classification | lstm | Reaching B | single_day | GY_data_day-37 | GY_data_day-37 | 44 | accuracy | 0.925 |
| classification | lstm | Reaching B | single_day | GY_data_day-37 | GY_data_day-37 | 46 | accuracy | 0.95 |
| classification | lstm | Reaching B | single_day | GY_data_day-37 | GY_data_day-37 | 46 | accuracy | 0.9625 |
| classification | lstm | Reaching B | single_day | GY_data_day-37 | GY_data_day-37 | 54 | accuracy | 0.975 |

| | | | | | | | | |
|---|---|---|---|---|---|---|---|---|
| classification | lstm | Reaching B | single_day | GY_data_day-37 | GY_data_day-37 | 54 | accuracy | 0.9625 |
| classification | lstm | Reaching B | single_day | GY_data_day-37 | GY_data_day-37 | 69 | accuracy | 0.975 |
| classification | lstm | Reaching B | single_day | GY_data_day-4 | GY_data_day-4 | 31 | accuracy | 0.965 |
| classification | lstm | Reaching B | single_day | GY_data_day-4 | GY_data_day-4 | 44 | accuracy | 0.97 |
| classification | lstm | Reaching B | single_day | GY_data_day-4 | GY_data_day-4 | 44 | accuracy | 0.975 |
| classification | lstm | Reaching B | single_day | GY_data_day-4 | GY_data_day-4 | 46 | accuracy | 0.88 |
| classification | lstm | Reaching B | single_day | GY_data_day-4 | GY_data_day-4 | 46 | accuracy | 0.985 |
| classification | lstm | Reaching B | single_day | GY_data_day-4 | GY_data_day-4 | 54 | accuracy | 0.93 |
| classification | lstm | Reaching B | single_day | GY_data_day-4 | GY_data_day-4 | 54 | accuracy | 0.975 |
| classification | lstm | Reaching B | single_day | GY_data_day-4 | GY_data_day-4 | 69 | accuracy | 0.925 |
| classification | lstm | Reaching B | single_day | GY_data_day-5 | GY_data_day-5 | 31 | accuracy | 0.9125 |
| classification | lstm | Reaching B | single_day | GY_data_day-5 | GY_data_day-5 | 44 | accuracy | 0.958333 |
| classification | lstm | Reaching B | single_day | GY_data_day-5 | GY_data_day-5 | 44 | accuracy | 0.975 |
| classification | lstm | Reaching B | single_day | GY_data_day-5 | GY_data_day-5 | 46 | accuracy | 0.870833 |
| classification | lstm | Reaching B | single_day | GY_data_day-5 | GY_data_day-5 | 46 | accuracy | 0.933333 |
| classification | lstm | Reaching B | single_day | GY_data_day-5 | GY_data_day-5 | 54 | accuracy | 0.9625 |
| classification | lstm | Reaching B | single_day | GY_data_day-5 | GY_data_day-5 | 54 | accuracy | 0.954167 |
| classification | lstm | Reaching B | single_day | GY_data_day-5 | GY_data_day-5 | 69 | accuracy | 0.941667 |
| classification | lstm | Reaching B | single_day | GY_data_day-6 | GY_data_day-6 | 31 | accuracy | 0.770833 |
| classification | lstm | Reaching B | single_day | GY_data_day-6 | GY_data_day-6 | 44 | accuracy | 0.729167 |
| classification | lstm | Reaching B | single_day | GY_data_day-6 | GY_data_day-6 | 44 | accuracy | 0.795833 |
| classification | lstm | Reaching B | single_day | GY_data_day-6 | GY_data_day-6 | 46 | accuracy | 0.720833 |
| classification | lstm | Reaching B | single_day | GY_data_day-6 | GY_data_day-6 | 46 | accuracy | 0.7875 |
| classification | lstm | Reaching B | single_day | GY_data_day-6 | GY_data_day-6 | 54 | accuracy | 0.791667 |
| classification | lstm | Reaching B | single_day | GY_data_day-6 | GY_data_day-6 | 54 | accuracy | 0.795833 |
| classification | lstm | Reaching B | single_day | GY_data_day-6 | GY_data_day-6 | 69 | accuracy | 0.754167 |
| classification | lstm | Reaching B | single_day | GY_data_day-7 | GY_data_day-7 | 31 | accuracy | 0.483333 |
| classification | lstm | Reaching B | single_day | GY_data_day-7 | GY_data_day-7 | 44 | accuracy | 0.483333 |
| classification | lstm | Reaching B | single_day | GY_data_day-7 | GY_data_day-7 | 44 | accuracy | 0.479167 |
| classification | lstm | Reaching B | single_day | GY_data_day-7 | GY_data_day-7 | 46 | accuracy | 0.525 |
| classification | lstm | Reaching B | single_day | GY_data_day-7 | GY_data_day-7 | 46 | accuracy | 0.466667 |
| classification | lstm | Reaching B | single_day | GY_data_day-7 | GY_data_day-7 | 54 | accuracy | 0.45 |
| classification | lstm | Reaching B | single_day | GY_data_day-7 | GY_data_day-7 | 54 | accuracy | 0.458333 |
| classification | lstm | Reaching B | single_day | GY_data_day-7 | GY_data_day-7 | 69 | accuracy | 0.495833 |
| classification | lstm | Reaching B | single_day | GY_data_day-8 | GY_data_day-8 | 31 | accuracy | 0.95 |
| classification | lstm | Reaching B | single_day | GY_data_day-8 | GY_data_day-8 | 44 | accuracy | 0.9 |
| classification | lstm | Reaching B | single_day | GY_data_day-8 | GY_data_day-8 | 44 | accuracy | 0.95625 |
| classification | lstm | Reaching B | single_day | GY_data_day-8 | GY_data_day-8 | 46 | accuracy | 0.90625 |
| classification | lstm | Reaching B | single_day | GY_data_day-8 | GY_data_day-8 | 46 | accuracy | 0.9375 |
| classification | lstm | Reaching B | single_day | GY_data_day-8 | GY_data_day-8 | 54 | accuracy | 0.89375 |
| classification | lstm | Reaching B | single_day | GY_data_day-8 | GY_data_day-8 | 54 | accuracy | 0.91875 |
| classification | lstm | Reaching B | single_day | GY_data_day-8 | GY_data_day-8 | 69 | accuracy | 0.88125 |
| classification | lstm | Reaching B | single_day | GY_data_day-9 | GY_data_day-9 | 31 | accuracy | 0.925 |
| classification | lstm | Reaching B | single_day | GY_data_day-9 | GY_data_day-9 | 44 | accuracy | 0.9 |
| classification | lstm | Reaching B | single_day | GY_data_day-9 | GY_data_day-9 | 44 | accuracy | 0.7875 |
| classification | lstm | Reaching B | single_day | GY_data_day-9 | GY_data_day-9 | 46 | accuracy | 0.8875 |
| classification | lstm | Reaching B | single_day | GY_data_day-9 | GY_data_day-9 | 46 | accuracy | 0.95 |
| classification | lstm | Reaching B | single_day | GY_data_day-9 | GY_data_day-9 | 54 | accuracy | 0.825 |
| classification | lstm | Reaching B | single_day | GY_data_day-9 | GY_data_day-9 | 54 | accuracy | 0.9 |
| classification | lstm | Reaching B | single_day | GY_data_day-9 | GY_data_day-9 | 69 | accuracy | 0.9 |
| classification | transformer | Reaching B | single_day | GY_data_day-1 | GY_data_day-1 | 31 | accuracy | 0.952941 |
| classification | transformer | Reaching B | single_day | GY_data_day-1 | GY_data_day-1 | 44 | accuracy | 0.947059 |
| classification | transformer | Reaching B | single_day | GY_data_day-1 | GY_data_day-1 | 44 | accuracy | 0.917647 |
| classification | transformer | Reaching B | single_day | GY_data_day-1 | GY_data_day-1 | 46 | accuracy | 0.958824 |
| classification | transformer | Reaching B | single_day | GY_data_day-1 | GY_data_day-1 | 46 | accuracy | 0.929412 |
| classification | transformer | Reaching B | single_day | GY_data_day-1 | GY_data_day-1 | 54 | accuracy | 0.952941 |
| classification | transformer | Reaching B | single_day | GY_data_day-1 | GY_data_day-1 | 54 | accuracy | 0.935294 |
| classification | transformer | Reaching B | single_day | GY_data_day-1 | GY_data_day-1 | 69 | accuracy | 0.958824 |
| classification | transformer | Reaching B | single_day | GY_data_day-10 | GY_data_day-10 | 31 | accuracy | 0.70625 |
| classification | transformer | Reaching B | single_day | GY_data_day-10 | GY_data_day-10 | 44 | accuracy | 0.7375 |
| classification | transformer | Reaching B | single_day | GY_data_day-10 | GY_data_day-10 | 44 | accuracy | 0.75625 |
| classification | transformer | Reaching B | single_day | GY_data_day-10 | GY_data_day-10 | 46 | accuracy | 0.76875 |
| classification | transformer | Reaching B | single_day | GY_data_day-10 | GY_data_day-10 | 46 | accuracy | 0.7375 |

| | | | | | | | | |
|---|---|---|---|---|---|---|---|---|
| classification | transformer | Reaching B | single_day | GY_data_day-10 | GY_data_day-10 | 54 | accuracy | 0.75625 |
| classification | transformer | Reaching B | single_day | GY_data_day-10 | GY_data_day-10 | 54 | accuracy | 0.75 |
| classification | transformer | Reaching B | single_day | GY_data_day-10 | GY_data_day-10 | 69 | accuracy | 0.75 |
| classification | transformer | Reaching B | single_day | GY_data_day-11 | GY_data_day-11 | 31 | accuracy | 0.9375 |
| classification | transformer | Reaching B | single_day | GY_data_day-11 | GY_data_day-11 | 44 | accuracy | 0.9875 |
| classification | transformer | Reaching B | single_day | GY_data_day-11 | GY_data_day-11 | 44 | accuracy | 0.9375 |
| classification | transformer | Reaching B | single_day | GY_data_day-11 | GY_data_day-11 | 46 | accuracy | 0.95 |
| classification | transformer | Reaching B | single_day | GY_data_day-11 | GY_data_day-11 | 46 | accuracy | 0.9625 |
| classification | transformer | Reaching B | single_day | GY_data_day-11 | GY_data_day-11 | 54 | accuracy | 0.875 |
| classification | transformer | Reaching B | single_day | GY_data_day-11 | GY_data_day-11 | 54 | accuracy | 0.925 |
| classification | transformer | Reaching B | single_day | GY_data_day-11 | GY_data_day-11 | 69 | accuracy | 0.975 |
| classification | transformer | Reaching B | single_day | GY_data_day-12 | GY_data_day-12 | 31 | accuracy | 0.982558 |
| classification | transformer | Reaching B | single_day | GY_data_day-12 | GY_data_day-12 | 44 | accuracy | 0.976744 |
| classification | transformer | Reaching B | single_day | GY_data_day-12 | GY_data_day-12 | 44 | accuracy | 0.97093 |
| classification | transformer | Reaching B | single_day | GY_data_day-12 | GY_data_day-12 | 46 | accuracy | 0.988372 |
| classification | transformer | Reaching B | single_day | GY_data_day-12 | GY_data_day-12 | 46 | accuracy | 0.982558 |
| classification | transformer | Reaching B | single_day | GY_data_day-12 | GY_data_day-12 | 54 | accuracy | 0.959302 |
| classification | transformer | Reaching B | single_day | GY_data_day-12 | GY_data_day-12 | 54 | accuracy | 0.953488 |
| classification | transformer | Reaching B | single_day | GY_data_day-12 | GY_data_day-12 | 69 | accuracy | 0.976744 |
| classification | transformer | Reaching B | single_day | GY_data_day-13 | GY_data_day-13 | 31 | accuracy | 0.975 |
| classification | transformer | Reaching B | single_day | GY_data_day-13 | GY_data_day-13 | 44 | accuracy | 0.9125 |
| classification | transformer | Reaching B | single_day | GY_data_day-13 | GY_data_day-13 | 44 | accuracy | 0.95 |
| classification | transformer | Reaching B | single_day | GY_data_day-13 | GY_data_day-13 | 46 | accuracy | 0.9375 |
| classification | transformer | Reaching B | single_day | GY_data_day-13 | GY_data_day-13 | 46 | accuracy | 0.9625 |
| classification | transformer | Reaching B | single_day | GY_data_day-13 | GY_data_day-13 | 54 | accuracy | 0.9625 |
| classification | transformer | Reaching B | single_day | GY_data_day-13 | GY_data_day-13 | 54 | accuracy | 0.95 |
| classification | transformer | Reaching B | single_day | GY_data_day-13 | GY_data_day-13 | 69 | accuracy | 0.8875 |
| classification | transformer | Reaching B | single_day | GY_data_day-14 | GY_data_day-14 | 31 | accuracy | 0.916667 |
| classification | transformer | Reaching B | single_day | GY_data_day-14 | GY_data_day-14 | 44 | accuracy | 0.933333 |
| classification | transformer | Reaching B | single_day | GY_data_day-14 | GY_data_day-14 | 44 | accuracy | 0.916667 |
| classification | transformer | Reaching B | single_day | GY_data_day-14 | GY_data_day-14 | 46 | accuracy | 0.925 |
| classification | transformer | Reaching B | single_day | GY_data_day-14 | GY_data_day-14 | 54 | accuracy | 0.883333 |
| classification | transformer | Reaching B | single_day | GY_data_day-14 | GY_data_day-14 | 54 | accuracy | 0.916667 |
| classification | transformer | Reaching B | single_day | GY_data_day-14 | GY_data_day-14 | 69 | accuracy | 0.925 |
| classification | transformer | Reaching B | single_day | GY_data_day-15 | GY_data_day-15 | 31 | accuracy | 0.94375 |
| classification | transformer | Reaching B | single_day | GY_data_day-15 | GY_data_day-15 | 44 | accuracy | 0.95625 |
| classification | transformer | Reaching B | single_day | GY_data_day-15 | GY_data_day-15 | 44 | accuracy | 0.96875 |
| classification | transformer | Reaching B | single_day | GY_data_day-15 | GY_data_day-15 | 46 | accuracy | 0.95 |
| classification | transformer | Reaching B | single_day | GY_data_day-15 | GY_data_day-15 | 46 | accuracy | 0.9375 |
| classification | transformer | Reaching B | single_day | GY_data_day-15 | GY_data_day-15 | 54 | accuracy | 0.98125 |
| classification | transformer | Reaching B | single_day | GY_data_day-15 | GY_data_day-15 | 54 | accuracy | 0.96875 |
| classification | transformer | Reaching B | single_day | GY_data_day-15 | GY_data_day-15 | 69 | accuracy | 0.9625 |
| classification | transformer | Reaching B | single_day | GY_data_day-16 | GY_data_day-16 | 31 | accuracy | 0.9 |
| classification | transformer | Reaching B | single_day | GY_data_day-16 | GY_data_day-16 | 44 | accuracy | 0.95 |
| classification | transformer | Reaching B | single_day | GY_data_day-16 | GY_data_day-16 | 44 | accuracy | 0.8 |
| classification | transformer | Reaching B | single_day | GY_data_day-16 | GY_data_day-16 | 46 | accuracy | 0.925 |
| classification | transformer | Reaching B | single_day | GY_data_day-16 | GY_data_day-16 | 46 | accuracy | 0.95 |
| classification | transformer | Reaching B | single_day | GY_data_day-16 | GY_data_day-16 | 54 | accuracy | 1 |
| classification | transformer | Reaching B | single_day | GY_data_day-16 | GY_data_day-16 | 69 | accuracy | 0.975 |
| classification | transformer | Reaching B | single_day | GY_data_day-17 | GY_data_day-17 | 31 | accuracy | 0.975 |
| classification | transformer | Reaching B | single_day | GY_data_day-17 | GY_data_day-17 | 44 | accuracy | 0.975 |
| classification | transformer | Reaching B | single_day | GY_data_day-17 | GY_data_day-17 | 44 | accuracy | 0.991667 |
| classification | transformer | Reaching B | single_day | GY_data_day-17 | GY_data_day-17 | 46 | accuracy | 0.991667 |
| classification | transformer | Reaching B | single_day | GY_data_day-17 | GY_data_day-17 | 46 | accuracy | 0.966667 |
| classification | transformer | Reaching B | single_day | GY_data_day-17 | GY_data_day-17 | 54 | accuracy | 1 |
| classification | transformer | Reaching B | single_day | GY_data_day-17 | GY_data_day-17 | 69 | accuracy | 0.975 |
| classification | transformer | Reaching B | single_day | GY_data_day-18 | GY_data_day-18 | 31 | accuracy | 0.925 |
| classification | transformer | Reaching B | single_day | GY_data_day-18 | GY_data_day-18 | 44 | accuracy | 0.9875 |
| classification | transformer | Reaching B | single_day | GY_data_day-18 | GY_data_day-18 | 44 | accuracy | 0.9375 |
| classification | transformer | Reaching B | single_day | GY_data_day-18 | GY_data_day-18 | 46 | accuracy | 1 |
| classification | transformer | Reaching B | single_day | GY_data_day-18 | GY_data_day-18 | 46 | accuracy | 0.975 |
| classification | transformer | Reaching B | single_day | GY_data_day-18 | GY_data_day-18 | 54 | accuracy | 0.975 |
| classification | transformer | Reaching B | single_day | GY_data_day-18 | GY_data_day-18 | 54 | accuracy | 0.9125 |

| | | | | | | | | |
|---|---|---|---|---|---|---|---|---|
| classification | transformer | Reaching B | single_day | GY_data_day-18 | GY_data_day-18 | 69 | accuracy | 0.975 |
| classification | transformer | Reaching B | single_day | GY_data_day-19 | GY_data_day-19 | 31 | accuracy | 0.975 |
| classification | transformer | Reaching B | single_day | GY_data_day-19 | GY_data_day-19 | 44 | accuracy | 0.95 |
| classification | transformer | Reaching B | single_day | GY_data_day-19 | GY_data_day-19 | 44 | accuracy | 0.975 |
| classification | transformer | Reaching B | single_day | GY_data_day-19 | GY_data_day-19 | 46 | accuracy | 0.95 |
| classification | transformer | Reaching B | single_day | GY_data_day-19 | GY_data_day-19 | 46 | accuracy | 0.9625 |
| classification | transformer | Reaching B | single_day | GY_data_day-19 | GY_data_day-19 | 54 | accuracy | 0.9375 |
| classification | transformer | Reaching B | single_day | GY_data_day-19 | GY_data_day-19 | 69 | accuracy | 0.975 |
| classification | transformer | Reaching B | single_day | GY_data_day-2 | GY_data_day-2 | 31 | accuracy | 0.99 |
| classification | transformer | Reaching B | single_day | GY_data_day-2 | GY_data_day-2 | 44 | accuracy | 0.995 |
| classification | transformer | Reaching B | single_day | GY_data_day-2 | GY_data_day-2 | 44 | accuracy | 0.98 |
| classification | transformer | Reaching B | single_day | GY_data_day-2 | GY_data_day-2 | 46 | accuracy | 0.985 |
| classification | transformer | Reaching B | single_day | GY_data_day-2 | GY_data_day-2 | 54 | accuracy | 0.995 |
| classification | transformer | Reaching B | single_day | GY_data_day-2 | GY_data_day-2 | 54 | accuracy | 0.98 |
| classification | transformer | Reaching B | single_day | GY_data_day-2 | GY_data_day-2 | 69 | accuracy | 0.99 |
| classification | transformer | Reaching B | single_day | GY_data_day-20 | GY_data_day-20 | 31 | accuracy | 1 |
| classification | transformer | Reaching B | single_day | GY_data_day-20 | GY_data_day-20 | 44 | accuracy | 0.975 |
| classification | transformer | Reaching B | single_day | GY_data_day-20 | GY_data_day-20 | 44 | accuracy | 0.95 |
| classification | transformer | Reaching B | single_day | GY_data_day-20 | GY_data_day-20 | 46 | accuracy | 0.925 |
| classification | transformer | Reaching B | single_day | GY_data_day-20 | GY_data_day-20 | 54 | accuracy | 0.95 |
| classification | transformer | Reaching B | single_day | GY_data_day-20 | GY_data_day-20 | 54 | accuracy | 0.85 |
| classification | transformer | Reaching B | single_day | GY_data_day-20 | GY_data_day-20 | 69 | accuracy | 0.95 |
| classification | transformer | Reaching B | single_day | GY_data_day-21 | GY_data_day-21 | 31 | accuracy | 0.475 |
| classification | transformer | Reaching B | single_day | GY_data_day-21 | GY_data_day-21 | 44 | accuracy | 0.45 |
| classification | transformer | Reaching B | single_day | GY_data_day-21 | GY_data_day-21 | 44 | accuracy | 0.55 |
| classification | transformer | Reaching B | single_day | GY_data_day-21 | GY_data_day-21 | 46 | accuracy | 0.475 |
| classification | transformer | Reaching B | single_day | GY_data_day-21 | GY_data_day-21 | 46 | accuracy | 0.533333 |
| classification | transformer | Reaching B | single_day | GY_data_day-21 | GY_data_day-21 | 54 | accuracy | 0.408333 |
| classification | transformer | Reaching B | single_day | GY_data_day-21 | GY_data_day-21 | 54 | accuracy | 0.55 |
| classification | transformer | Reaching B | single_day | GY_data_day-21 | GY_data_day-21 | 69 | accuracy | 0.525 |
| classification | transformer | Reaching B | single_day | GY_data_day-22 | GY_data_day-22 | 31 | accuracy | 0.971429 |
| classification | transformer | Reaching B | single_day | GY_data_day-22 | GY_data_day-22 | 44 | accuracy | 0.960714 |
| classification | transformer | Reaching B | single_day | GY_data_day-22 | GY_data_day-22 | 44 | accuracy | 0.957143 |
| classification | transformer | Reaching B | single_day | GY_data_day-22 | GY_data_day-22 | 46 | accuracy | 0.975 |
| classification | transformer | Reaching B | single_day | GY_data_day-22 | GY_data_day-22 | 46 | accuracy | 0.942857 |
| classification | transformer | Reaching B | single_day | GY_data_day-22 | GY_data_day-22 | 54 | accuracy | 0.975 |
| classification | transformer | Reaching B | single_day | GY_data_day-22 | GY_data_day-22 | 54 | accuracy | 0.967857 |
| classification | transformer | Reaching B | single_day | GY_data_day-22 | GY_data_day-22 | 69 | accuracy | 0.960714 |
| classification | transformer | Reaching B | single_day | GY_data_day-23 | GY_data_day-23 | 31 | accuracy | 0.95 |
| classification | transformer | Reaching B | single_day | GY_data_day-23 | GY_data_day-23 | 44 | accuracy | 0.945 |
| classification | transformer | Reaching B | single_day | GY_data_day-23 | GY_data_day-23 | 44 | accuracy | 0.935 |
| classification | transformer | Reaching B | single_day | GY_data_day-23 | GY_data_day-23 | 46 | accuracy | 0.955 |
| classification | transformer | Reaching B | single_day | GY_data_day-23 | GY_data_day-23 | 46 | accuracy | 0.93 |
| classification | transformer | Reaching B | single_day | GY_data_day-23 | GY_data_day-23 | 54 | accuracy | 0.965 |
| classification | transformer | Reaching B | single_day | GY_data_day-23 | GY_data_day-23 | 54 | accuracy | 0.945 |
| classification | transformer | Reaching B | single_day | GY_data_day-23 | GY_data_day-23 | 69 | accuracy | 0.94 |
| classification | transformer | Reaching B | single_day | GY_data_day-24 | GY_data_day-24 | 31 | accuracy | 0.9875 |
| classification | transformer | Reaching B | single_day | GY_data_day-24 | GY_data_day-24 | 44 | accuracy | 0.925 |
| classification | transformer | Reaching B | single_day | GY_data_day-24 | GY_data_day-24 | 44 | accuracy | 0.8875 |
| classification | transformer | Reaching B | single_day | GY_data_day-24 | GY_data_day-24 | 46 | accuracy | 0.9375 |
| classification | transformer | Reaching B | single_day | GY_data_day-24 | GY_data_day-24 | 46 | accuracy | 0.8875 |
| classification | transformer | Reaching B | single_day | GY_data_day-24 | GY_data_day-24 | 54 | accuracy | 0.9375 |
| classification | transformer | Reaching B | single_day | GY_data_day-24 | GY_data_day-24 | 54 | accuracy | 0.9 |
| classification | transformer | Reaching B | single_day | GY_data_day-24 | GY_data_day-24 | 69 | accuracy | 0.9375 |
| classification | transformer | Reaching B | single_day | GY_data_day-25 | GY_data_day-25 | 31 | accuracy | 0.9875 |
| classification | transformer | Reaching B | single_day | GY_data_day-25 | GY_data_day-25 | 44 | accuracy | 0.9 |
| classification | transformer | Reaching B | single_day | GY_data_day-25 | GY_data_day-25 | 44 | accuracy | 0.9375 |
| classification | transformer | Reaching B | single_day | GY_data_day-25 | GY_data_day-25 | 46 | accuracy | 0.95 |
| classification | transformer | Reaching B | single_day | GY_data_day-25 | GY_data_day-25 | 46 | accuracy | 0.9125 |
| classification | transformer | Reaching B | single_day | GY_data_day-25 | GY_data_day-25 | 54 | accuracy | 0.9625 |
| classification | transformer | Reaching B | single_day | GY_data_day-25 | GY_data_day-25 | 69 | accuracy | 0.9375 |
| classification | transformer | Reaching B | single_day | GY_data_day-26 | GY_data_day-26 | 31 | accuracy | 0.925 |
| classification | transformer | Reaching B | single_day | GY_data_day-26 | GY_data_day-26 | 44 | accuracy | 0.9125 |

| | | | | | | | | |
|---|---|---|---|---|---|---|---|---|
| classification | transformer | Reaching B | single_day | GY_data_day-26 | GY_data_day-26 | 44 | accuracy | 0.8875 |
| classification | transformer | Reaching B | single_day | GY_data_day-26 | GY_data_day-26 | 46 | accuracy | 0.85 |
| classification | transformer | Reaching B | single_day | GY_data_day-26 | GY_data_day-26 | 46 | accuracy | 0.8625 |
| classification | transformer | Reaching B | single_day | GY_data_day-26 | GY_data_day-26 | 54 | accuracy | 0.925 |
| classification | transformer | Reaching B | single_day | GY_data_day-26 | GY_data_day-26 | 54 | accuracy | 0.9 |
| classification | transformer | Reaching B | single_day | GY_data_day-26 | GY_data_day-26 | 69 | accuracy | 0.925 |
| classification | transformer | Reaching B | single_day | GY_data_day-27 | GY_data_day-27 | 31 | accuracy | 0.925 |
| classification | transformer | Reaching B | single_day | GY_data_day-27 | GY_data_day-27 | 44 | accuracy | 0.925 |
| classification | transformer | Reaching B | single_day | GY_data_day-27 | GY_data_day-27 | 44 | accuracy | 0.8875 |
| classification | transformer | Reaching B | single_day | GY_data_day-27 | GY_data_day-27 | 46 | accuracy | 0.9125 |
| classification | transformer | Reaching B | single_day | GY_data_day-27 | GY_data_day-27 | 54 | accuracy | 0.95 |
| classification | transformer | Reaching B | single_day | GY_data_day-27 | GY_data_day-27 | 54 | accuracy | 0.9375 |
| classification | transformer | Reaching B | single_day | GY_data_day-27 | GY_data_day-27 | 69 | accuracy | 0.95 |
| classification | transformer | Reaching B | single_day | GY_data_day-28 | GY_data_day-28 | 31 | accuracy | 0.9625 |
| classification | transformer | Reaching B | single_day | GY_data_day-28 | GY_data_day-28 | 44 | accuracy | 0.9375 |
| classification | transformer | Reaching B | single_day | GY_data_day-28 | GY_data_day-28 | 44 | accuracy | 0.85 |
| classification | transformer | Reaching B | single_day | GY_data_day-28 | GY_data_day-28 | 46 | accuracy | 0.975 |
| classification | transformer | Reaching B | single_day | GY_data_day-28 | GY_data_day-28 | 46 | accuracy | 0.95 |
| classification | transformer | Reaching B | single_day | GY_data_day-28 | GY_data_day-28 | 54 | accuracy | 0.925 |
| classification | transformer | Reaching B | single_day | GY_data_day-28 | GY_data_day-28 | 54 | accuracy | 0.9 |
| classification | transformer | Reaching B | single_day | GY_data_day-28 | GY_data_day-28 | 69 | accuracy | 0.925 |
| classification | transformer | Reaching B | single_day | GY_data_day-29 | GY_data_day-29 | 31 | accuracy | 0.9125 |
| classification | transformer | Reaching B | single_day | GY_data_day-29 | GY_data_day-29 | 44 | accuracy | 0.9375 |
| classification | transformer | Reaching B | single_day | GY_data_day-29 | GY_data_day-29 | 44 | accuracy | 0.925 |
| classification | transformer | Reaching B | single_day | GY_data_day-29 | GY_data_day-29 | 46 | accuracy | 0.9125 |
| classification | transformer | Reaching B | single_day | GY_data_day-29 | GY_data_day-29 | 46 | accuracy | 0.9 |
| classification | transformer | Reaching B | single_day | GY_data_day-29 | GY_data_day-29 | 54 | accuracy | 0.875 |
| classification | transformer | Reaching B | single_day | GY_data_day-29 | GY_data_day-29 | 54 | accuracy | 0.9125 |
| classification | transformer | Reaching B | single_day | GY_data_day-29 | GY_data_day-29 | 69 | accuracy | 0.9125 |
| classification | transformer | Reaching B | single_day | GY_data_day-3 | GY_data_day-3 | 31 | accuracy | 0.941667 |
| classification | transformer | Reaching B | single_day | GY_data_day-3 | GY_data_day-3 | 44 | accuracy | 0.966667 |
| classification | transformer | Reaching B | single_day | GY_data_day-3 | GY_data_day-3 | 44 | accuracy | 0.925 |
| classification | transformer | Reaching B | single_day | GY_data_day-3 | GY_data_day-3 | 46 | accuracy | 0.941667 |
| classification | transformer | Reaching B | single_day | GY_data_day-3 | GY_data_day-3 | 46 | accuracy | 0.916667 |
| classification | transformer | Reaching B | single_day | GY_data_day-3 | GY_data_day-3 | 54 | accuracy | 0.966667 |
| classification | transformer | Reaching B | single_day | GY_data_day-3 | GY_data_day-3 | 54 | accuracy | 0.9 |
| classification | transformer | Reaching B | single_day | GY_data_day-3 | GY_data_day-3 | 69 | accuracy | 0.941667 |
| classification | transformer | Reaching B | single_day | GY_data_day-30 | GY_data_day-30 | 31 | accuracy | 0.9625 |
| classification | transformer | Reaching B | single_day | GY_data_day-30 | GY_data_day-30 | 44 | accuracy | 0.9625 |
| classification | transformer | Reaching B | single_day | GY_data_day-30 | GY_data_day-30 | 46 | accuracy | 0.95 |
| classification | transformer | Reaching B | single_day | GY_data_day-30 | GY_data_day-30 | 46 | accuracy | 0.925 |
| classification | transformer | Reaching B | single_day | GY_data_day-30 | GY_data_day-30 | 54 | accuracy | 0.9625 |
| classification | transformer | Reaching B | single_day | GY_data_day-30 | GY_data_day-30 | 69 | accuracy | 0.9375 |
| classification | transformer | Reaching B | single_day | GY_data_day-31 | GY_data_day-31 | 31 | accuracy | 0.85 |
| classification | transformer | Reaching B | single_day | GY_data_day-31 | GY_data_day-31 | 44 | accuracy | 0.875 |
| classification | transformer | Reaching B | single_day | GY_data_day-31 | GY_data_day-31 | 44 | accuracy | 0.825 |
| classification | transformer | Reaching B | single_day | GY_data_day-31 | GY_data_day-31 | 46 | accuracy | 0.9 |
| classification | transformer | Reaching B | single_day | GY_data_day-31 | GY_data_day-31 | 46 | accuracy | 0.775 |
| classification | transformer | Reaching B | single_day | GY_data_day-31 | GY_data_day-31 | 54 | accuracy | 0.725 |
| classification | transformer | Reaching B | single_day | GY_data_day-31 | GY_data_day-31 | 54 | accuracy | 0.8 |
| classification | transformer | Reaching B | single_day | GY_data_day-31 | GY_data_day-31 | 69 | accuracy | 0.825 |
| classification | transformer | Reaching B | single_day | GY_data_day-32 | GY_data_day-32 | 31 | accuracy | 0.9375 |
| classification | transformer | Reaching B | single_day | GY_data_day-32 | GY_data_day-32 | 44 | accuracy | 0.9 |
| classification | transformer | Reaching B | single_day | GY_data_day-32 | GY_data_day-32 | 44 | accuracy | 0.8875 |
| classification | transformer | Reaching B | single_day | GY_data_day-32 | GY_data_day-32 | 46 | accuracy | 0.9375 |
| classification | transformer | Reaching B | single_day | GY_data_day-32 | GY_data_day-32 | 46 | accuracy | 0.95 |
| classification | transformer | Reaching B | single_day | GY_data_day-32 | GY_data_day-32 | 54 | accuracy | 0.9375 |
| classification | transformer | Reaching B | single_day | GY_data_day-32 | GY_data_day-32 | 54 | accuracy | 0.8875 |
| classification | transformer | Reaching B | single_day | GY_data_day-32 | GY_data_day-32 | 69 | accuracy | 0.9125 |
| classification | transformer | Reaching B | single_day | GY_data_day-33 | GY_data_day-33 | 31 | accuracy | 0.866667 |
| classification | transformer | Reaching B | single_day | GY_data_day-33 | GY_data_day-33 | 44 | accuracy | 0.833333 |
| classification | transformer | Reaching B | single_day | GY_data_day-33 | GY_data_day-33 | 44 | accuracy | 0.808333 |
| classification | transformer | Reaching B | single_day | GY_data_day-33 | GY_data_day-33 | 46 | accuracy | 0.85 |

| | | | | | | | | |
|---|---|---|---|---|---|---|---|---|
| classification | transformer | Reaching B | single_day | GY_data_day-33 | GY_data_day-33 | 46 | accuracy | 0.775 |
| classification | transformer | Reaching B | single_day | GY_data_day-33 | GY_data_day-33 | 54 | accuracy | 0.833333 |
| classification | transformer | Reaching B | single_day | GY_data_day-33 | GY_data_day-33 | 69 | accuracy | 0.833333 |
| classification | transformer | Reaching B | single_day | GY_data_day-34 | GY_data_day-34 | 31 | accuracy | 0.95625 |
| classification | transformer | Reaching B | single_day | GY_data_day-34 | GY_data_day-34 | 44 | accuracy | 0.9375 |
| classification | transformer | Reaching B | single_day | GY_data_day-34 | GY_data_day-34 | 44 | accuracy | 0.93125 |
| classification | transformer | Reaching B | single_day | GY_data_day-34 | GY_data_day-34 | 46 | accuracy | 0.93125 |
| classification | transformer | Reaching B | single_day | GY_data_day-34 | GY_data_day-34 | 46 | accuracy | 0.925 |
| classification | transformer | Reaching B | single_day | GY_data_day-34 | GY_data_day-34 | 54 | accuracy | 0.9375 |
| classification | transformer | Reaching B | single_day | GY_data_day-34 | GY_data_day-34 | 54 | accuracy | 0.89375 |
| classification | transformer | Reaching B | single_day | GY_data_day-34 | GY_data_day-34 | 69 | accuracy | 0.925 |
| classification | transformer | Reaching B | single_day | GY_data_day-35 | GY_data_day-35 | 31 | accuracy | 0.925 |
| classification | transformer | Reaching B | single_day | GY_data_day-35 | GY_data_day-35 | 44 | accuracy | 0.85625 |
| classification | transformer | Reaching B | single_day | GY_data_day-35 | GY_data_day-35 | 44 | accuracy | 0.88125 |
| classification | transformer | Reaching B | single_day | GY_data_day-35 | GY_data_day-35 | 46 | accuracy | 0.89375 |
| classification | transformer | Reaching B | single_day | GY_data_day-35 | GY_data_day-35 | 46 | accuracy | 0.8875 |
| classification | transformer | Reaching B | single_day | GY_data_day-35 | GY_data_day-35 | 54 | accuracy | 0.89375 |
| classification | transformer | Reaching B | single_day | GY_data_day-35 | GY_data_day-35 | 54 | accuracy | 0.85625 |
| classification | transformer | Reaching B | single_day | GY_data_day-35 | GY_data_day-35 | 69 | accuracy | 0.9 |
| classification | transformer | Reaching B | single_day | GY_data_day-36 | GY_data_day-36 | 31 | accuracy | 0.925 |
| classification | transformer | Reaching B | single_day | GY_data_day-36 | GY_data_day-36 | 44 | accuracy | 0.89375 |
| classification | transformer | Reaching B | single_day | GY_data_day-36 | GY_data_day-36 | 46 | accuracy | 0.86875 |
| classification | transformer | Reaching B | single_day | GY_data_day-36 | GY_data_day-36 | 46 | accuracy | 0.8875 |
| classification | transformer | Reaching B | single_day | GY_data_day-36 | GY_data_day-36 | 54 | accuracy | 0.9125 |
| classification | transformer | Reaching B | single_day | GY_data_day-36 | GY_data_day-36 | 69 | accuracy | 0.8875 |
| classification | transformer | Reaching B | single_day | GY_data_day-37 | GY_data_day-37 | 31 | accuracy | 0.9875 |
| classification | transformer | Reaching B | single_day | GY_data_day-37 | GY_data_day-37 | 44 | accuracy | 0.95 |
| classification | transformer | Reaching B | single_day | GY_data_day-37 | GY_data_day-37 | 44 | accuracy | 0.9 |
| classification | transformer | Reaching B | single_day | GY_data_day-37 | GY_data_day-37 | 46 | accuracy | 0.975 |
| classification | transformer | Reaching B | single_day | GY_data_day-37 | GY_data_day-37 | 54 | accuracy | 0.9375 |
| classification | transformer | Reaching B | single_day | GY_data_day-37 | GY_data_day-37 | 69 | accuracy | 0.9125 |
| classification | transformer | Reaching B | single_day | GY_data_day-4 | GY_data_day-4 | 31 | accuracy | 0.98 |
| classification | transformer | Reaching B | single_day | GY_data_day-4 | GY_data_day-4 | 44 | accuracy | 0.995 |
| classification | transformer | Reaching B | single_day | GY_data_day-4 | GY_data_day-4 | 44 | accuracy | 0.99 |
| classification | transformer | Reaching B | single_day | GY_data_day-4 | GY_data_day-4 | 46 | accuracy | 0.965 |
| classification | transformer | Reaching B | single_day | GY_data_day-4 | GY_data_day-4 | 46 | accuracy | 0.97 |
| classification | transformer | Reaching B | single_day | GY_data_day-4 | GY_data_day-4 | 54 | accuracy | 0.985 |
| classification | transformer | Reaching B | single_day | GY_data_day-4 | GY_data_day-4 | 54 | accuracy | 0.965 |
| classification | transformer | Reaching B | single_day | GY_data_day-4 | GY_data_day-4 | 69 | accuracy | 0.975 |
| classification | transformer | Reaching B | single_day | GY_data_day-5 | GY_data_day-5 | 31 | accuracy | 0.979167 |
| classification | transformer | Reaching B | single_day | GY_data_day-5 | GY_data_day-5 | 44 | accuracy | 0.958333 |
| classification | transformer | Reaching B | single_day | GY_data_day-5 | GY_data_day-5 | 44 | accuracy | 0.95 |
| classification | transformer | Reaching B | single_day | GY_data_day-5 | GY_data_day-5 | 46 | accuracy | 0.970833 |
| classification | transformer | Reaching B | single_day | GY_data_day-5 | GY_data_day-5 | 46 | accuracy | 0.966667 |
| classification | transformer | Reaching B | single_day | GY_data_day-5 | GY_data_day-5 | 54 | accuracy | 0.966667 |
| classification | transformer | Reaching B | single_day | GY_data_day-5 | GY_data_day-5 | 54 | accuracy | 0.941667 |
| classification | transformer | Reaching B | single_day | GY_data_day-5 | GY_data_day-5 | 69 | accuracy | 0.9625 |
| classification | transformer | Reaching B | single_day | GY_data_day-6 | GY_data_day-6 | 31 | accuracy | 0.808333 |
| classification | transformer | Reaching B | single_day | GY_data_day-6 | GY_data_day-6 | 44 | accuracy | 0.820833 |
| classification | transformer | Reaching B | single_day | GY_data_day-6 | GY_data_day-6 | 44 | accuracy | 0.808333 |
| classification | transformer | Reaching B | single_day | GY_data_day-6 | GY_data_day-6 | 46 | accuracy | 0.820833 |
| classification | transformer | Reaching B | single_day | GY_data_day-6 | GY_data_day-6 | 46 | accuracy | 0.825 |
| classification | transformer | Reaching B | single_day | GY_data_day-6 | GY_data_day-6 | 54 | accuracy | 0.833333 |
| classification | transformer | Reaching B | single_day | GY_data_day-6 | GY_data_day-6 | 54 | accuracy | 0.8375 |
| classification | transformer | Reaching B | single_day | GY_data_day-6 | GY_data_day-6 | 69 | accuracy | 0.833333 |
| classification | transformer | Reaching B | single_day | GY_data_day-7 | GY_data_day-7 | 31 | accuracy | 0.466667 |
| classification | transformer | Reaching B | single_day | GY_data_day-7 | GY_data_day-7 | 44 | accuracy | 0.5 |
| classification | transformer | Reaching B | single_day | GY_data_day-7 | GY_data_day-7 | 44 | accuracy | 0.4875 |
| classification | transformer | Reaching B | single_day | GY_data_day-7 | GY_data_day-7 | 46 | accuracy | 0.4625 |
| classification | transformer | Reaching B | single_day | GY_data_day-7 | GY_data_day-7 | 46 | accuracy | 0.520833 |
| classification | transformer | Reaching B | single_day | GY_data_day-7 | GY_data_day-7 | 54 | accuracy | 0.454167 |
| classification | transformer | Reaching B | single_day | GY_data_day-7 | GY_data_day-7 | 54 | accuracy | 0.475 |
| classification | transformer | Reaching B | single_day | GY_data_day-7 | GY_data_day-7 | 69 | accuracy | 0.395833 |

| | | | | | | | | |
|---|---|---|---|---|---|---|---|---|
| classification | transformer | Reaching B | single_day | GY_data_day-8 | GY_data_day-8 | 31 | accuracy | 0.98125 |
| classification | transformer | Reaching B | single_day | GY_data_day-8 | GY_data_day-8 | 44 | accuracy | 0.95625 |
| classification | transformer | Reaching B | single_day | GY_data_day-8 | GY_data_day-8 | 44 | accuracy | 0.9125 |
| classification | transformer | Reaching B | single_day | GY_data_day-8 | GY_data_day-8 | 46 | accuracy | 0.9625 |
| classification | transformer | Reaching B | single_day | GY_data_day-8 | GY_data_day-8 | 46 | accuracy | 0.9375 |
| classification | transformer | Reaching B | single_day | GY_data_day-8 | GY_data_day-8 | 54 | accuracy | 0.9625 |
| classification | transformer | Reaching B | single_day | GY_data_day-8 | GY_data_day-8 | 54 | accuracy | 0.94375 |
| classification | transformer | Reaching B | single_day | GY_data_day-8 | GY_data_day-8 | 69 | accuracy | 0.94375 |
| classification | transformer | Reaching B | single_day | GY_data_day-9 | GY_data_day-9 | 31 | accuracy | 0.9875 |
| classification | transformer | Reaching B | single_day | GY_data_day-9 | GY_data_day-9 | 44 | accuracy | 0.975 |
| classification | transformer | Reaching B | single_day | GY_data_day-9 | GY_data_day-9 | 46 | accuracy | 0.9875 |
| classification | transformer | Reaching B | single_day | GY_data_day-9 | GY_data_day-9 | 46 | accuracy | 0.95 |
| classification | transformer | Reaching B | single_day | GY_data_day-9 | GY_data_day-9 | 54 | accuracy | 1 |
| classification | transformer | Reaching B | single_day | GY_data_day-9 | GY_data_day-9 | 54 | accuracy | 0.95 |
| classification | transformer | Reaching B | single_day | GY_data_day-9 | GY_data_day-9 | 69 | accuracy | 1 |
| classification | lfads | Reaching B | single_day | GY_data_day-1 | GY_data_day-1 | 44 | accuracy | 0.888235 |
| classification | lfads | Reaching B | single_day | GY_data_day-1 | GY_data_day-1 | 46 | accuracy | 0.864706 |
| classification | lfads | Reaching B | single_day | GY_data_day-1 | GY_data_day-1 | 54 | accuracy | 0.876471 |
| classification | lfads | Reaching B | single_day | GY_data_day-10 | GY_data_day-10 | 44 | accuracy | 0.66875 |
| classification | lfads | Reaching B | single_day | GY_data_day-10 | GY_data_day-10 | 46 | accuracy | 0.675 |
| classification | lfads | Reaching B | single_day | GY_data_day-10 | GY_data_day-10 | 54 | accuracy | 0.65 |
| classification | lfads | Reaching B | single_day | GY_data_day-11 | GY_data_day-11 | 44 | accuracy | 0.8625 |
| classification | lfads | Reaching B | single_day | GY_data_day-11 | GY_data_day-11 | 46 | accuracy | 0.9 |
| classification | lfads | Reaching B | single_day | GY_data_day-11 | GY_data_day-11 | 54 | accuracy | 0.85 |
| classification | lfads | Reaching B | single_day | GY_data_day-12 | GY_data_day-12 | 44 | accuracy | 0.901163 |
| classification | lfads | Reaching B | single_day | GY_data_day-12 | GY_data_day-12 | 46 | accuracy | 0.94186 |
| classification | lfads | Reaching B | single_day | GY_data_day-12 | GY_data_day-12 | 54 | accuracy | 0.866279 |
| classification | lfads | Reaching B | single_day | GY_data_day-13 | GY_data_day-13 | 44 | accuracy | 0.7875 |
| classification | lfads | Reaching B | single_day | GY_data_day-13 | GY_data_day-13 | 46 | accuracy | 0.7375 |
| classification | lfads | Reaching B | single_day | GY_data_day-13 | GY_data_day-13 | 54 | accuracy | 0.8 |
| classification | lfads | Reaching B | single_day | GY_data_day-14 | GY_data_day-14 | 44 | accuracy | 0.816667 |
| classification | lfads | Reaching B | single_day | GY_data_day-14 | GY_data_day-14 | 46 | accuracy | 0.758333 |
| classification | lfads | Reaching B | single_day | GY_data_day-14 | GY_data_day-14 | 54 | accuracy | 0.825 |
| classification | lfads | Reaching B | single_day | GY_data_day-15 | GY_data_day-15 | 44 | accuracy | 0.875 |
| classification | lfads | Reaching B | single_day | GY_data_day-15 | GY_data_day-15 | 46 | accuracy | 0.8625 |
| classification | lfads | Reaching B | single_day | GY_data_day-15 | GY_data_day-15 | 54 | accuracy | 0.89375 |
| classification | lfads | Reaching B | single_day | GY_data_day-16 | GY_data_day-16 | 44 | accuracy | 0.75 |
| classification | lfads | Reaching B | single_day | GY_data_day-16 | GY_data_day-16 | 46 | accuracy | 0.725 |
| classification | lfads | Reaching B | single_day | GY_data_day-16 | GY_data_day-16 | 54 | accuracy | 0.7 |
| classification | lfads | Reaching B | single_day | GY_data_day-17 | GY_data_day-17 | 44 | accuracy | 0.875 |
| classification | lfads | Reaching B | single_day | GY_data_day-17 | GY_data_day-17 | 46 | accuracy | 0.875 |
| classification | lfads | Reaching B | single_day | GY_data_day-17 | GY_data_day-17 | 54 | accuracy | 0.875 |
| classification | lfads | Reaching B | single_day | GY_data_day-18 | GY_data_day-18 | 44 | accuracy | 0.9375 |
| classification | lfads | Reaching B | single_day | GY_data_day-18 | GY_data_day-18 | 46 | accuracy | 0.95 |
| classification | lfads | Reaching B | single_day | GY_data_day-18 | GY_data_day-18 | 54 | accuracy | 0.95 |
| classification | lfads | Reaching B | single_day | GY_data_day-19 | GY_data_day-19 | 44 | accuracy | 0.7125 |
| classification | lfads | Reaching B | single_day | GY_data_day-19 | GY_data_day-19 | 46 | accuracy | 0.6375 |
| classification | lfads | Reaching B | single_day | GY_data_day-19 | GY_data_day-19 | 54 | accuracy | 0.85 |
| classification | lfads | Reaching B | single_day | GY_data_day-2 | GY_data_day-2 | 44 | accuracy | 0.935 |
| classification | lfads | Reaching B | single_day | GY_data_day-2 | GY_data_day-2 | 46 | accuracy | 0.96 |
| classification | lfads | Reaching B | single_day | GY_data_day-2 | GY_data_day-2 | 54 | accuracy | 0.955 |
| classification | lfads | Reaching B | single_day | GY_data_day-20 | GY_data_day-20 | 44 | accuracy | 0.7 |
| classification | lfads | Reaching B | single_day | GY_data_day-20 | GY_data_day-20 | 46 | accuracy | 0.675 |
| classification | lfads | Reaching B | single_day | GY_data_day-20 | GY_data_day-20 | 54 | accuracy | 0.85 |
| classification | lfads | Reaching B | single_day | GY_data_day-21 | GY_data_day-21 | 44 | accuracy | 0.416667 |
| classification | lfads | Reaching B | single_day | GY_data_day-21 | GY_data_day-21 | 46 | accuracy | 0.491667 |
| classification | lfads | Reaching B | single_day | GY_data_day-21 | GY_data_day-21 | 54 | accuracy | 0.45 |
| classification | lfads | Reaching B | single_day | GY_data_day-22 | GY_data_day-22 | 44 | accuracy | 0.95 |
| classification | lfads | Reaching B | single_day | GY_data_day-22 | GY_data_day-22 | 46 | accuracy | 0.95 |
| classification | lfads | Reaching B | single_day | GY_data_day-22 | GY_data_day-22 | 54 | accuracy | 0.942857 |
| classification | lfads | Reaching B | single_day | GY_data_day-23 | GY_data_day-23 | 44 | accuracy | 0.945 |
| classification | lfads | Reaching B | single_day | GY_data_day-23 | GY_data_day-23 | 46 | accuracy | 0.935 |
| classification | lfads | Reaching B | single_day | GY_data_day-23 | GY_data_day-23 | 54 | accuracy | 0.91 |

| | | | | | | | | |
|---|---|---|---|---|---|---|---|---|
| classification | lfads | Reaching B | single_day | GY_data_day-24 | GY_data_day-24 | 44 | accuracy | 0.6625 |
| classification | lfads | Reaching B | single_day | GY_data_day-24 | GY_data_day-24 | 46 | accuracy | 0.6625 |
| classification | lfads | Reaching B | single_day | GY_data_day-24 | GY_data_day-24 | 54 | accuracy | 0.6625 |
| classification | lfads | Reaching B | single_day | GY_data_day-25 | GY_data_day-25 | 44 | accuracy | 0.7 |
| classification | lfads | Reaching B | single_day | GY_data_day-25 | GY_data_day-25 | 46 | accuracy | 0.7625 |
| classification | lfads | Reaching B | single_day | GY_data_day-25 | GY_data_day-25 | 54 | accuracy | 0.7375 |
| classification | lfads | Reaching B | single_day | GY_data_day-26 | GY_data_day-26 | 44 | accuracy | 0.825 |
| classification | lfads | Reaching B | single_day | GY_data_day-26 | GY_data_day-26 | 46 | accuracy | 0.8625 |
| classification | lfads | Reaching B | single_day | GY_data_day-26 | GY_data_day-26 | 54 | accuracy | 0.8125 |
| classification | lfads | Reaching B | single_day | GY_data_day-27 | GY_data_day-27 | 44 | accuracy | 0.8 |
| classification | lfads | Reaching B | single_day | GY_data_day-27 | GY_data_day-27 | 46 | accuracy | 0.75 |
| classification | lfads | Reaching B | single_day | GY_data_day-27 | GY_data_day-27 | 54 | accuracy | 0.9 |
| classification | lfads | Reaching B | single_day | GY_data_day-28 | GY_data_day-28 | 44 | accuracy | 0.8625 |
| classification | lfads | Reaching B | single_day | GY_data_day-28 | GY_data_day-28 | 46 | accuracy | 0.9 |
| classification | lfads | Reaching B | single_day | GY_data_day-28 | GY_data_day-28 | 54 | accuracy | 0.8 |
| classification | lfads | Reaching B | single_day | GY_data_day-29 | GY_data_day-29 | 44 | accuracy | 0.7875 |
| classification | lfads | Reaching B | single_day | GY_data_day-29 | GY_data_day-29 | 46 | accuracy | 0.8375 |
| classification | lfads | Reaching B | single_day | GY_data_day-29 | GY_data_day-29 | 54 | accuracy | 0.825 |
| classification | lfads | Reaching B | single_day | GY_data_day-3 | GY_data_day-3 | 44 | accuracy | 0.816667 |
| classification | lfads | Reaching B | single_day | GY_data_day-3 | GY_data_day-3 | 46 | accuracy | 0.758333 |
| classification | lfads | Reaching B | single_day | GY_data_day-3 | GY_data_day-3 | 54 | accuracy | 0.833333 |
| classification | lfads | Reaching B | single_day | GY_data_day-30 | GY_data_day-30 | 44 | accuracy | 0.8625 |
| classification | lfads | Reaching B | single_day | GY_data_day-30 | GY_data_day-30 | 46 | accuracy | 0.775 |
| classification | lfads | Reaching B | single_day | GY_data_day-30 | GY_data_day-30 | 54 | accuracy | 0.8625 |
| classification | lfads | Reaching B | single_day | GY_data_day-31 | GY_data_day-31 | 44 | accuracy | 0.625 |
| classification | lfads | Reaching B | single_day | GY_data_day-31 | GY_data_day-31 | 46 | accuracy | 0.7 |
| classification | lfads | Reaching B | single_day | GY_data_day-31 | GY_data_day-31 | 54 | accuracy | 0.525 |
| classification | lfads | Reaching B | single_day | GY_data_day-32 | GY_data_day-32 | 44 | accuracy | 0.75 |
| classification | lfads | Reaching B | single_day | GY_data_day-32 | GY_data_day-32 | 46 | accuracy | 0.7375 |
| classification | lfads | Reaching B | single_day | GY_data_day-32 | GY_data_day-32 | 54 | accuracy | 0.725 |
| classification | lfads | Reaching B | single_day | GY_data_day-33 | GY_data_day-33 | 44 | accuracy | 0.7 |
| classification | lfads | Reaching B | single_day | GY_data_day-33 | GY_data_day-33 | 46 | accuracy | 0.716667 |
| classification | lfads | Reaching B | single_day | GY_data_day-33 | GY_data_day-33 | 54 | accuracy | 0.641667 |
| classification | lfads | Reaching B | single_day | GY_data_day-34 | GY_data_day-34 | 44 | accuracy | 0.90625 |
| classification | lfads | Reaching B | single_day | GY_data_day-34 | GY_data_day-34 | 46 | accuracy | 0.9 |
| classification | lfads | Reaching B | single_day | GY_data_day-34 | GY_data_day-34 | 54 | accuracy | 0.9125 |
| classification | lfads | Reaching B | single_day | GY_data_day-35 | GY_data_day-35 | 44 | accuracy | 0.81875 |
| classification | lfads | Reaching B | single_day | GY_data_day-35 | GY_data_day-35 | 46 | accuracy | 0.8375 |
| classification | lfads | Reaching B | single_day | GY_data_day-35 | GY_data_day-35 | 54 | accuracy | 0.83125 |
| classification | lfads | Reaching B | single_day | GY_data_day-36 | GY_data_day-36 | 44 | accuracy | 0.8875 |
| classification | lfads | Reaching B | single_day | GY_data_day-36 | GY_data_day-36 | 46 | accuracy | 0.85625 |
| classification | lfads | Reaching B | single_day | GY_data_day-36 | GY_data_day-36 | 54 | accuracy | 0.875 |
| classification | lfads | Reaching B | single_day | GY_data_day-37 | GY_data_day-37 | 44 | accuracy | 0.825 |
| classification | lfads | Reaching B | single_day | GY_data_day-37 | GY_data_day-37 | 46 | accuracy | 0.8375 |
| classification | lfads | Reaching B | single_day | GY_data_day-37 | GY_data_day-37 | 54 | accuracy | 0.7875 |
| classification | lfads | Reaching B | single_day | GY_data_day-4 | GY_data_day-4 | 44 | accuracy | 0.96 |
| classification | lfads | Reaching B | single_day | GY_data_day-4 | GY_data_day-4 | 46 | accuracy | 0.95 |
| classification | lfads | Reaching B | single_day | GY_data_day-4 | GY_data_day-4 | 54 | accuracy | 0.935 |
| classification | lfads | Reaching B | single_day | GY_data_day-5 | GY_data_day-5 | 44 | accuracy | 0.875 |
| classification | lfads | Reaching B | single_day | GY_data_day-5 | GY_data_day-5 | 46 | accuracy | 0.879167 |
| classification | lfads | Reaching B | single_day | GY_data_day-5 | GY_data_day-5 | 54 | accuracy | 0.845833 |
| classification | lfads | Reaching B | single_day | GY_data_day-6 | GY_data_day-6 | 44 | accuracy | 0.766667 |
| classification | lfads | Reaching B | single_day | GY_data_day-6 | GY_data_day-6 | 46 | accuracy | 0.7375 |
| classification | lfads | Reaching B | single_day | GY_data_day-6 | GY_data_day-6 | 54 | accuracy | 0.745833 |
| classification | lfads | Reaching B | single_day | GY_data_day-7 | GY_data_day-7 | 44 | accuracy | 0.416667 |
| classification | lfads | Reaching B | single_day | GY_data_day-7 | GY_data_day-7 | 46 | accuracy | 0.420833 |
| classification | lfads | Reaching B | single_day | GY_data_day-7 | GY_data_day-7 | 54 | accuracy | 0.45 |
| classification | lfads | Reaching B | single_day | GY_data_day-8 | GY_data_day-8 | 44 | accuracy | 0.78125 |
| classification | lfads | Reaching B | single_day | GY_data_day-8 | GY_data_day-8 | 46 | accuracy | 0.74375 |
| classification | lfads | Reaching B | single_day | GY_data_day-8 | GY_data_day-8 | 54 | accuracy | 0.76875 |
| classification | lfads | Reaching B | single_day | GY_data_day-9 | GY_data_day-9 | 44 | accuracy | 0.9375 |
| classification | lfads | Reaching B | single_day | GY_data_day-9 | GY_data_day-9 | 46 | accuracy | 0.9 |
| classification | lfads | Reaching B | single_day | GY_data_day-9 | GY_data_day-9 | 54 | accuracy | 0.8125 |

| | | | | | | | | |
|---|---|---|---|---|---|---|---|---|
| classification | cycle_gan | Reaching B | single_day | GY_data_day-1 | GY_data_day-1 | 31 | accuracy | 0.823529 |
| classification | cycle_gan | Reaching B | single_day | GY_data_day-1 | GY_data_day-1 | 44 | accuracy | 0.905882 |
| classification | cycle_gan | Reaching B | single_day | GY_data_day-1 | GY_data_day-1 | 46 | accuracy | 0.823529 |
| classification | cycle_gan | Reaching B | single_day | GY_data_day-1 | GY_data_day-1 | 54 | accuracy | 0.864706 |
| classification | cycle_gan | Reaching B | single_day | GY_data_day-1 | GY_data_day-1 | 69 | accuracy | 0.805882 |
| classification | cycle_gan | Reaching B | single_day | GY_data_day-10 | GY_data_day-10 | 31 | accuracy | 0.6625 |
| classification | cycle_gan | Reaching B | single_day | GY_data_day-10 | GY_data_day-10 | 44 | accuracy | 0.6625 |
| classification | cycle_gan | Reaching B | single_day | GY_data_day-10 | GY_data_day-10 | 46 | accuracy | 0.71875 |
| classification | cycle_gan | Reaching B | single_day | GY_data_day-10 | GY_data_day-10 | 54 | accuracy | 0.7125 |
| classification | cycle_gan | Reaching B | single_day | GY_data_day-10 | GY_data_day-10 | 69 | accuracy | 0.65625 |
| classification | cycle_gan | Reaching B | single_day | GY_data_day-11 | GY_data_day-11 | 31 | accuracy | 0.8625 |
| classification | cycle_gan | Reaching B | single_day | GY_data_day-11 | GY_data_day-11 | 44 | accuracy | 0.825 |
| classification | cycle_gan | Reaching B | single_day | GY_data_day-11 | GY_data_day-11 | 46 | accuracy | 0.8375 |
| classification | cycle_gan | Reaching B | single_day | GY_data_day-11 | GY_data_day-11 | 54 | accuracy | 0.7875 |
| classification | cycle_gan | Reaching B | single_day | GY_data_day-11 | GY_data_day-11 | 69 | accuracy | 0.8375 |
| classification | cycle_gan | Reaching B | single_day | GY_data_day-12 | GY_data_day-12 | 31 | accuracy | 0.924419 |
| classification | cycle_gan | Reaching B | single_day | GY_data_day-12 | GY_data_day-12 | 44 | accuracy | 0.906977 |
| classification | cycle_gan | Reaching B | single_day | GY_data_day-12 | GY_data_day-12 | 46 | accuracy | 0.936047 |
| classification | cycle_gan | Reaching B | single_day | GY_data_day-12 | GY_data_day-12 | 54 | accuracy | 0.901163 |
| classification | cycle_gan | Reaching B | single_day | GY_data_day-12 | GY_data_day-12 | 69 | accuracy | 0.895349 |
| classification | cycle_gan | Reaching B | single_day | GY_data_day-13 | GY_data_day-13 | 31 | accuracy | 0.8375 |
| classification | cycle_gan | Reaching B | single_day | GY_data_day-13 | GY_data_day-13 | 44 | accuracy | 0.75 |
| classification | cycle_gan | Reaching B | single_day | GY_data_day-13 | GY_data_day-13 | 46 | accuracy | 0.8375 |
| classification | cycle_gan | Reaching B | single_day | GY_data_day-13 | GY_data_day-13 | 54 | accuracy | 0.825 |
| classification | cycle_gan | Reaching B | single_day | GY_data_day-13 | GY_data_day-13 | 69 | accuracy | 0.8 |
| classification | cycle_gan | Reaching B | single_day | GY_data_day-14 | GY_data_day-14 | 31 | accuracy | 0.866667 |
| classification | cycle_gan | Reaching B | single_day | GY_data_day-14 | GY_data_day-14 | 44 | accuracy | 0.816667 |
| classification | cycle_gan | Reaching B | single_day | GY_data_day-14 | GY_data_day-14 | 46 | accuracy | 0.8 |
| classification | cycle_gan | Reaching B | single_day | GY_data_day-14 | GY_data_day-14 | 54 | accuracy | 0.75 |
| classification | cycle_gan | Reaching B | single_day | GY_data_day-14 | GY_data_day-14 | 69 | accuracy | 0.908333 |
| classification | cycle_gan | Reaching B | single_day | GY_data_day-15 | GY_data_day-15 | 31 | accuracy | 0.91875 |
| classification | cycle_gan | Reaching B | single_day | GY_data_day-15 | GY_data_day-15 | 44 | accuracy | 0.9 |
| classification | cycle_gan | Reaching B | single_day | GY_data_day-15 | GY_data_day-15 | 46 | accuracy | 0.9375 |
| classification | cycle_gan | Reaching B | single_day | GY_data_day-15 | GY_data_day-15 | 54 | accuracy | 0.90625 |
| classification | cycle_gan | Reaching B | single_day | GY_data_day-15 | GY_data_day-15 | 69 | accuracy | 0.925 |
| classification | cycle_gan | Reaching B | single_day | GY_data_day-16 | GY_data_day-16 | 31 | accuracy | 0.975 |
| classification | cycle_gan | Reaching B | single_day | GY_data_day-16 | GY_data_day-16 | 44 | accuracy | 0.65 |
| classification | cycle_gan | Reaching B | single_day | GY_data_day-16 | GY_data_day-16 | 46 | accuracy | 0.9 |
| classification | cycle_gan | Reaching B | single_day | GY_data_day-16 | GY_data_day-16 | 54 | accuracy | 0.75 |
| classification | cycle_gan | Reaching B | single_day | GY_data_day-16 | GY_data_day-16 | 69 | accuracy | 0.775 |
| classification | cycle_gan | Reaching B | single_day | GY_data_day-17 | GY_data_day-17 | 31 | accuracy | 0.95 |
| classification | cycle_gan | Reaching B | single_day | GY_data_day-17 | GY_data_day-17 | 44 | accuracy | 0.925 |
| classification | cycle_gan | Reaching B | single_day | GY_data_day-17 | GY_data_day-17 | 46 | accuracy | 0.958333 |
| classification | cycle_gan | Reaching B | single_day | GY_data_day-17 | GY_data_day-17 | 54 | accuracy | 0.933333 |
| classification | cycle_gan | Reaching B | single_day | GY_data_day-17 | GY_data_day-17 | 69 | accuracy | 0.941667 |
| classification | cycle_gan | Reaching B | single_day | GY_data_day-18 | GY_data_day-18 | 31 | accuracy | 0.9625 |
| classification | cycle_gan | Reaching B | single_day | GY_data_day-18 | GY_data_day-18 | 44 | accuracy | 0.95 |
| classification | cycle_gan | Reaching B | single_day | GY_data_day-18 | GY_data_day-18 | 46 | accuracy | 0.9375 |
| classification | cycle_gan | Reaching B | single_day | GY_data_day-18 | GY_data_day-18 | 54 | accuracy | 0.9 |
| classification | cycle_gan | Reaching B | single_day | GY_data_day-18 | GY_data_day-18 | 69 | accuracy | 0.9 |
| classification | cycle_gan | Reaching B | single_day | GY_data_day-19 | GY_data_day-19 | 31 | accuracy | 0.85 |
| classification | cycle_gan | Reaching B | single_day | GY_data_day-19 | GY_data_day-19 | 44 | accuracy | 0.7125 |
| classification | cycle_gan | Reaching B | single_day | GY_data_day-19 | GY_data_day-19 | 46 | accuracy | 0.9 |
| classification | cycle_gan | Reaching B | single_day | GY_data_day-19 | GY_data_day-19 | 54 | accuracy | 0.825 |
| classification | cycle_gan | Reaching B | single_day | GY_data_day-19 | GY_data_day-19 | 69 | accuracy | 0.8875 |
| classification | cycle_gan | Reaching B | single_day | GY_data_day-2 | GY_data_day-2 | 31 | accuracy | 0.905 |
| classification | cycle_gan | Reaching B | single_day | GY_data_day-2 | GY_data_day-2 | 44 | accuracy | 0.86 |
| classification | cycle_gan | Reaching B | single_day | GY_data_day-2 | GY_data_day-2 | 46 | accuracy | 0.905 |
| classification | cycle_gan | Reaching B | single_day | GY_data_day-2 | GY_data_day-2 | 54 | accuracy | 0.91 |
| classification | cycle_gan | Reaching B | single_day | GY_data_day-2 | GY_data_day-2 | 69 | accuracy | 0.86 |
| classification | cycle_gan | Reaching B | single_day | GY_data_day-20 | GY_data_day-20 | 31 | accuracy | 0.875 |
| classification | cycle_gan | Reaching B | single_day | GY_data_day-20 | GY_data_day-20 | 44 | accuracy | 0.75 |
| classification | cycle_gan | Reaching B | single_day | GY_data_day-20 | GY_data_day-20 | 46 | accuracy | 0.925 |

| | | | | | | | | |
|---|---|---|---|---|---|---|---|---|
| classification | cycle_gan | Reaching B | single_day | GY_data_day-20 | GY_data_day-20 | 54 | accuracy | 0.725 |
| classification | cycle_gan | Reaching B | single_day | GY_data_day-20 | GY_data_day-20 | 69 | accuracy | 0.85 |
| classification | cycle_gan | Reaching B | single_day | GY_data_day-21 | GY_data_day-21 | 31 | accuracy | 0.566667 |
| classification | cycle_gan | Reaching B | single_day | GY_data_day-21 | GY_data_day-21 | 44 | accuracy | 0.558333 |
| classification | cycle_gan | Reaching B | single_day | GY_data_day-21 | GY_data_day-21 | 46 | accuracy | 0.516667 |
| classification | cycle_gan | Reaching B | single_day | GY_data_day-21 | GY_data_day-21 | 54 | accuracy | 0.591667 |
| classification | cycle_gan | Reaching B | single_day | GY_data_day-21 | GY_data_day-21 | 69 | accuracy | 0.6 |
| classification | cycle_gan | Reaching B | single_day | GY_data_day-22 | GY_data_day-22 | 31 | accuracy | 0.942857 |
| classification | cycle_gan | Reaching B | single_day | GY_data_day-22 | GY_data_day-22 | 44 | accuracy | 0.942857 |
| classification | cycle_gan | Reaching B | single_day | GY_data_day-22 | GY_data_day-22 | 46 | accuracy | 0.925 |
| classification | cycle_gan | Reaching B | single_day | GY_data_day-22 | GY_data_day-22 | 54 | accuracy | 0.932143 |
| classification | cycle_gan | Reaching B | single_day | GY_data_day-22 | GY_data_day-22 | 69 | accuracy | 0.9 |
| classification | cycle_gan | Reaching B | single_day | GY_data_day-23 | GY_data_day-23 | 31 | accuracy | 0.95 |
| classification | cycle_gan | Reaching B | single_day | GY_data_day-23 | GY_data_day-23 | 44 | accuracy | 0.955 |
| classification | cycle_gan | Reaching B | single_day | GY_data_day-23 | GY_data_day-23 | 46 | accuracy | 0.945 |
| classification | cycle_gan | Reaching B | single_day | GY_data_day-23 | GY_data_day-23 | 54 | accuracy | 0.925 |
| classification | cycle_gan | Reaching B | single_day | GY_data_day-23 | GY_data_day-23 | 69 | accuracy | 0.955 |
| classification | cycle_gan | Reaching B | single_day | GY_data_day-24 | GY_data_day-24 | 31 | accuracy | 0.7875 |
| classification | cycle_gan | Reaching B | single_day | GY_data_day-24 | GY_data_day-24 | 44 | accuracy | 0.85 |
| classification | cycle_gan | Reaching B | single_day | GY_data_day-24 | GY_data_day-24 | 46 | accuracy | 0.9 |
| classification | cycle_gan | Reaching B | single_day | GY_data_day-24 | GY_data_day-24 | 54 | accuracy | 0.7375 |
| classification | cycle_gan | Reaching B | single_day | GY_data_day-24 | GY_data_day-24 | 69 | accuracy | 0.825 |
| classification | cycle_gan | Reaching B | single_day | GY_data_day-25 | GY_data_day-25 | 31 | accuracy | 0.875 |
| classification | cycle_gan | Reaching B | single_day | GY_data_day-25 | GY_data_day-25 | 44 | accuracy | 0.8125 |
| classification | cycle_gan | Reaching B | single_day | GY_data_day-25 | GY_data_day-25 | 46 | accuracy | 0.8 |
| classification | cycle_gan | Reaching B | single_day | GY_data_day-25 | GY_data_day-25 | 54 | accuracy | 0.8375 |
| classification | cycle_gan | Reaching B | single_day | GY_data_day-25 | GY_data_day-25 | 69 | accuracy | 0.825 |
| classification | cycle_gan | Reaching B | single_day | GY_data_day-26 | GY_data_day-26 | 31 | accuracy | 0.825 |
| classification | cycle_gan | Reaching B | single_day | GY_data_day-26 | GY_data_day-26 | 44 | accuracy | 0.875 |
| classification | cycle_gan | Reaching B | single_day | GY_data_day-26 | GY_data_day-26 | 46 | accuracy | 0.8125 |
| classification | cycle_gan | Reaching B | single_day | GY_data_day-26 | GY_data_day-26 | 54 | accuracy | 0.85 |
| classification | cycle_gan | Reaching B | single_day | GY_data_day-26 | GY_data_day-26 | 69 | accuracy | 0.7625 |
| classification | cycle_gan | Reaching B | single_day | GY_data_day-27 | GY_data_day-27 | 31 | accuracy | 0.8625 |
| classification | cycle_gan | Reaching B | single_day | GY_data_day-27 | GY_data_day-27 | 44 | accuracy | 0.85 |
| classification | cycle_gan | Reaching B | single_day | GY_data_day-27 | GY_data_day-27 | 46 | accuracy | 0.8875 |
| classification | cycle_gan | Reaching B | single_day | GY_data_day-27 | GY_data_day-27 | 54 | accuracy | 0.8375 |
| classification | cycle_gan | Reaching B | single_day | GY_data_day-27 | GY_data_day-27 | 69 | accuracy | 0.875 |
| classification | cycle_gan | Reaching B | single_day | GY_data_day-28 | GY_data_day-28 | 31 | accuracy | 0.9125 |
| classification | cycle_gan | Reaching B | single_day | GY_data_day-28 | GY_data_day-28 | 44 | accuracy | 0.875 |
| classification | cycle_gan | Reaching B | single_day | GY_data_day-28 | GY_data_day-28 | 46 | accuracy | 0.9 |
| classification | cycle_gan | Reaching B | single_day | GY_data_day-28 | GY_data_day-28 | 54 | accuracy | 0.8375 |
| classification | cycle_gan | Reaching B | single_day | GY_data_day-28 | GY_data_day-28 | 69 | accuracy | 0.9125 |
| classification | cycle_gan | Reaching B | single_day | GY_data_day-29 | GY_data_day-29 | 31 | accuracy | 0.825 |
| classification | cycle_gan | Reaching B | single_day | GY_data_day-29 | GY_data_day-29 | 44 | accuracy | 0.8 |
| classification | cycle_gan | Reaching B | single_day | GY_data_day-29 | GY_data_day-29 | 46 | accuracy | 0.8125 |
| classification | cycle_gan | Reaching B | single_day | GY_data_day-29 | GY_data_day-29 | 54 | accuracy | 0.8125 |
| classification | cycle_gan | Reaching B | single_day | GY_data_day-29 | GY_data_day-29 | 69 | accuracy | 0.825 |
| classification | cycle_gan | Reaching B | single_day | GY_data_day-3 | GY_data_day-3 | 31 | accuracy | 0.683333 |
| classification | cycle_gan | Reaching B | single_day | GY_data_day-3 | GY_data_day-3 | 44 | accuracy | 0.741667 |
| classification | cycle_gan | Reaching B | single_day | GY_data_day-3 | GY_data_day-3 | 46 | accuracy | 0.716667 |
| classification | cycle_gan | Reaching B | single_day | GY_data_day-3 | GY_data_day-3 | 54 | accuracy | 0.641667 |
| classification | cycle_gan | Reaching B | single_day | GY_data_day-3 | GY_data_day-3 | 69 | accuracy | 0.708333 |
| classification | cycle_gan | Reaching B | single_day | GY_data_day-30 | GY_data_day-30 | 31 | accuracy | 0.8875 |
| classification | cycle_gan | Reaching B | single_day | GY_data_day-30 | GY_data_day-30 | 44 | accuracy | 0.875 |
| classification | cycle_gan | Reaching B | single_day | GY_data_day-30 | GY_data_day-30 | 46 | accuracy | 0.8125 |
| classification | cycle_gan | Reaching B | single_day | GY_data_day-30 | GY_data_day-30 | 54 | accuracy | 0.85 |
| classification | cycle_gan | Reaching B | single_day | GY_data_day-30 | GY_data_day-30 | 69 | accuracy | 0.8875 |
| classification | cycle_gan | Reaching B | single_day | GY_data_day-31 | GY_data_day-31 | 31 | accuracy | 0.425 |
| classification | cycle_gan | Reaching B | single_day | GY_data_day-31 | GY_data_day-31 | 44 | accuracy | 0.6 |
| classification | cycle_gan | Reaching B | single_day | GY_data_day-31 | GY_data_day-31 | 46 | accuracy | 0.6 |
| classification | cycle_gan | Reaching B | single_day | GY_data_day-31 | GY_data_day-31 | 54 | accuracy | 0.75 |
| classification | cycle_gan | Reaching B | single_day | GY_data_day-31 | GY_data_day-31 | 69 | accuracy | 0.55 |
| classification | cycle_gan | Reaching B | single_day | GY_data_day-32 | GY_data_day-32 | 31 | accuracy | 0.825 |

| | | | | | | | | |
|---|---|---|---|---|---|---|---|---|
| classification | cycle_gan | Reaching B | single_day | GY_data_day-32 | GY_data_day-32 | 44 | accuracy | 0.7125 |
| classification | cycle_gan | Reaching B | single_day | GY_data_day-32 | GY_data_day-32 | 46 | accuracy | 0.8 |
| classification | cycle_gan | Reaching B | single_day | GY_data_day-32 | GY_data_day-32 | 54 | accuracy | 0.8125 |
| classification | cycle_gan | Reaching B | single_day | GY_data_day-32 | GY_data_day-32 | 69 | accuracy | 0.75 |
| classification | cycle_gan | Reaching B | single_day | GY_data_day-33 | GY_data_day-33 | 31 | accuracy | 0.875 |
| classification | cycle_gan | Reaching B | single_day | GY_data_day-33 | GY_data_day-33 | 44 | accuracy | 0.791667 |
| classification | cycle_gan | Reaching B | single_day | GY_data_day-33 | GY_data_day-33 | 46 | accuracy | 0.783333 |
| classification | cycle_gan | Reaching B | single_day | GY_data_day-33 | GY_data_day-33 | 54 | accuracy | 0.833333 |
| classification | cycle_gan | Reaching B | single_day | GY_data_day-33 | GY_data_day-33 | 69 | accuracy | 0.816667 |
| classification | cycle_gan | Reaching B | single_day | GY_data_day-34 | GY_data_day-34 | 31 | accuracy | 0.875 |
| classification | cycle_gan | Reaching B | single_day | GY_data_day-34 | GY_data_day-34 | 44 | accuracy | 0.93125 |
| classification | cycle_gan | Reaching B | single_day | GY_data_day-34 | GY_data_day-34 | 46 | accuracy | 0.9375 |
| classification | cycle_gan | Reaching B | single_day | GY_data_day-34 | GY_data_day-34 | 54 | accuracy | 0.925 |
| classification | cycle_gan | Reaching B | single_day | GY_data_day-34 | GY_data_day-34 | 69 | accuracy | 0.9375 |
| classification | cycle_gan | Reaching B | single_day | GY_data_day-35 | GY_data_day-35 | 31 | accuracy | 0.88125 |
| classification | cycle_gan | Reaching B | single_day | GY_data_day-35 | GY_data_day-35 | 44 | accuracy | 0.85625 |
| classification | cycle_gan | Reaching B | single_day | GY_data_day-35 | GY_data_day-35 | 46 | accuracy | 0.9375 |
| classification | cycle_gan | Reaching B | single_day | GY_data_day-35 | GY_data_day-35 | 54 | accuracy | 0.8125 |
| classification | cycle_gan | Reaching B | single_day | GY_data_day-35 | GY_data_day-35 | 69 | accuracy | 0.86875 |
| classification | cycle_gan | Reaching B | single_day | GY_data_day-36 | GY_data_day-36 | 31 | accuracy | 0.90625 |
| classification | cycle_gan | Reaching B | single_day | GY_data_day-36 | GY_data_day-36 | 44 | accuracy | 0.85 |
| classification | cycle_gan | Reaching B | single_day | GY_data_day-36 | GY_data_day-36 | 46 | accuracy | 0.925 |
| classification | cycle_gan | Reaching B | single_day | GY_data_day-36 | GY_data_day-36 | 54 | accuracy | 0.8625 |
| classification | cycle_gan | Reaching B | single_day | GY_data_day-36 | GY_data_day-36 | 69 | accuracy | 0.8375 |
| classification | cycle_gan | Reaching B | single_day | GY_data_day-37 | GY_data_day-37 | 31 | accuracy | 0.875 |
| classification | cycle_gan | Reaching B | single_day | GY_data_day-37 | GY_data_day-37 | 44 | accuracy | 0.8 |
| classification | cycle_gan | Reaching B | single_day | GY_data_day-37 | GY_data_day-37 | 46 | accuracy | 0.8125 |
| classification | cycle_gan | Reaching B | single_day | GY_data_day-37 | GY_data_day-37 | 54 | accuracy | 0.7875 |
| classification | cycle_gan | Reaching B | single_day | GY_data_day-37 | GY_data_day-37 | 69 | accuracy | 0.8125 |
| classification | cycle_gan | Reaching B | single_day | GY_data_day-4 | GY_data_day-4 | 31 | accuracy | 0.925 |
| classification | cycle_gan | Reaching B | single_day | GY_data_day-4 | GY_data_day-4 | 44 | accuracy | 0.915 |
| classification | cycle_gan | Reaching B | single_day | GY_data_day-4 | GY_data_day-4 | 46 | accuracy | 0.9 |
| classification | cycle_gan | Reaching B | single_day | GY_data_day-4 | GY_data_day-4 | 54 | accuracy | 0.915 |
| classification | cycle_gan | Reaching B | single_day | GY_data_day-4 | GY_data_day-4 | 69 | accuracy | 0.925 |
| classification | cycle_gan | Reaching B | single_day | GY_data_day-5 | GY_data_day-5 | 31 | accuracy | 0.804167 |
| classification | cycle_gan | Reaching B | single_day | GY_data_day-5 | GY_data_day-5 | 44 | accuracy | 0.770833 |
| classification | cycle_gan | Reaching B | single_day | GY_data_day-5 | GY_data_day-5 | 46 | accuracy | 0.783333 |
| classification | cycle_gan | Reaching B | single_day | GY_data_day-5 | GY_data_day-5 | 54 | accuracy | 0.804167 |
| classification | cycle_gan | Reaching B | single_day | GY_data_day-5 | GY_data_day-5 | 69 | accuracy | 0.833333 |
| classification | cycle_gan | Reaching B | single_day | GY_data_day-6 | GY_data_day-6 | 31 | accuracy | 0.725 |
| classification | cycle_gan | Reaching B | single_day | GY_data_day-6 | GY_data_day-6 | 44 | accuracy | 0.7375 |
| classification | cycle_gan | Reaching B | single_day | GY_data_day-6 | GY_data_day-6 | 46 | accuracy | 0.7625 |
| classification | cycle_gan | Reaching B | single_day | GY_data_day-6 | GY_data_day-6 | 54 | accuracy | 0.745833 |
| classification | cycle_gan | Reaching B | single_day | GY_data_day-6 | GY_data_day-6 | 69 | accuracy | 0.745833 |
| classification | cycle_gan | Reaching B | single_day | GY_data_day-7 | GY_data_day-7 | 31 | accuracy | 0.4375 |
| classification | cycle_gan | Reaching B | single_day | GY_data_day-7 | GY_data_day-7 | 44 | accuracy | 0.408333 |
| classification | cycle_gan | Reaching B | single_day | GY_data_day-7 | GY_data_day-7 | 46 | accuracy | 0.379167 |
| classification | cycle_gan | Reaching B | single_day | GY_data_day-7 | GY_data_day-7 | 54 | accuracy | 0.458333 |
| classification | cycle_gan | Reaching B | single_day | GY_data_day-7 | GY_data_day-7 | 69 | accuracy | 0.395833 |
| classification | cycle_gan | Reaching B | single_day | GY_data_day-8 | GY_data_day-8 | 31 | accuracy | 0.78125 |
| classification | cycle_gan | Reaching B | single_day | GY_data_day-8 | GY_data_day-8 | 44 | accuracy | 0.71875 |
| classification | cycle_gan | Reaching B | single_day | GY_data_day-8 | GY_data_day-8 | 46 | accuracy | 0.725 |
| classification | cycle_gan | Reaching B | single_day | GY_data_day-8 | GY_data_day-8 | 54 | accuracy | 0.69375 |
| classification | cycle_gan | Reaching B | single_day | GY_data_day-8 | GY_data_day-8 | 69 | accuracy | 0.7125 |
| classification | cycle_gan | Reaching B | single_day | GY_data_day-9 | GY_data_day-9 | 31 | accuracy | 0.8625 |
| classification | cycle_gan | Reaching B | single_day | GY_data_day-9 | GY_data_day-9 | 44 | accuracy | 0.875 |
| classification | cycle_gan | Reaching B | single_day | GY_data_day-9 | GY_data_day-9 | 46 | accuracy | 0.9 |
| classification | cycle_gan | Reaching B | single_day | GY_data_day-9 | GY_data_day-9 | 54 | accuracy | 0.875 |
| classification | cycle_gan | Reaching B | single_day | GY_data_day-9 | GY_data_day-9 | 69 | accuracy | 0.9125 |
| classification | stabilization | Reaching B | single_day | GY_data_day-1 | GY_data_day-1 | 44 | accuracy | 0.862745 |
| classification | stabilization | Reaching B | single_day | GY_data_day-1 | GY_data_day-1 | 46 | accuracy | 0.869281 |
| classification | stabilization | Reaching B | single_day | GY_data_day-1 | GY_data_day-1 | 54 | accuracy | 0.882353 |
| classification | stabilization | Reaching B | single_day | GY_data_day-10 | GY_data_day-10 | 44 | accuracy | 0.638889 |

| | | | | | | | | |
|---|---|---|---|---|---|---|---|---|
| classification | stabilization | Reaching B | single_day | GY_data_day-10 | GY_data_day-10 | 46 | accuracy | 0.618056 |
| classification | stabilization | Reaching B | single_day | GY_data_day-10 | GY_data_day-10 | 54 | accuracy | 0.659722 |
| classification | stabilization | Reaching B | single_day | GY_data_day-11 | GY_data_day-11 | 44 | accuracy | 0.791667 |
| classification | stabilization | Reaching B | single_day | GY_data_day-11 | GY_data_day-11 | 46 | accuracy | 0.791667 |
| classification | stabilization | Reaching B | single_day | GY_data_day-11 | GY_data_day-11 | 54 | accuracy | 0.805556 |
| classification | stabilization | Reaching B | single_day | GY_data_day-12 | GY_data_day-12 | 44 | accuracy | 0.851613 |
| classification | stabilization | Reaching B | single_day | GY_data_day-12 | GY_data_day-12 | 46 | accuracy | 0.864516 |
| classification | stabilization | Reaching B | single_day | GY_data_day-12 | GY_data_day-12 | 54 | accuracy | 0.812903 |
| classification | stabilization | Reaching B | single_day | GY_data_day-13 | GY_data_day-13 | 44 | accuracy | 0.777778 |
| classification | stabilization | Reaching B | single_day | GY_data_day-13 | GY_data_day-13 | 46 | accuracy | 0.791667 |
| classification | stabilization | Reaching B | single_day | GY_data_day-13 | GY_data_day-13 | 54 | accuracy | 0.791667 |
| classification | stabilization | Reaching B | single_day | GY_data_day-14 | GY_data_day-14 | 44 | accuracy | 0.907407 |
| classification | stabilization | Reaching B | single_day | GY_data_day-14 | GY_data_day-14 | 46 | accuracy | 0.907407 |
| classification | stabilization | Reaching B | single_day | GY_data_day-14 | GY_data_day-14 | 54 | accuracy | 0.898148 |
| classification | stabilization | Reaching B | single_day | GY_data_day-15 | GY_data_day-15 | 44 | accuracy | 0.923611 |
| classification | stabilization | Reaching B | single_day | GY_data_day-15 | GY_data_day-15 | 46 | accuracy | 0.916667 |
| classification | stabilization | Reaching B | single_day | GY_data_day-15 | GY_data_day-15 | 54 | accuracy | 0.930556 |
| classification | stabilization | Reaching B | single_day | GY_data_day-16 | GY_data_day-16 | 44 | accuracy | 0.685714 |
| classification | stabilization | Reaching B | single_day | GY_data_day-16 | GY_data_day-16 | 46 | accuracy | 0.742857 |
| classification | stabilization | Reaching B | single_day | GY_data_day-16 | GY_data_day-16 | 54 | accuracy | 0.8 |
| classification | stabilization | Reaching B | single_day | GY_data_day-17 | GY_data_day-17 | 44 | accuracy | 0.861111 |
| classification | stabilization | Reaching B | single_day | GY_data_day-17 | GY_data_day-17 | 46 | accuracy | 0.898148 |
| classification | stabilization | Reaching B | single_day | GY_data_day-17 | GY_data_day-17 | 54 | accuracy | 0.925926 |
| classification | stabilization | Reaching B | single_day | GY_data_day-18 | GY_data_day-18 | 44 | accuracy | 0.861111 |
| classification | stabilization | Reaching B | single_day | GY_data_day-18 | GY_data_day-18 | 46 | accuracy | 0.902778 |
| classification | stabilization | Reaching B | single_day | GY_data_day-18 | GY_data_day-18 | 54 | accuracy | 0.819444 |
| classification | stabilization | Reaching B | single_day | GY_data_day-19 | GY_data_day-19 | 44 | accuracy | 0.833333 |
| classification | stabilization | Reaching B | single_day | GY_data_day-19 | GY_data_day-19 | 46 | accuracy | 0.736111 |
| classification | stabilization | Reaching B | single_day | GY_data_day-19 | GY_data_day-19 | 54 | accuracy | 0.791667 |
| classification | stabilization | Reaching B | single_day | GY_data_day-2 | GY_data_day-2 | 44 | accuracy | 0.9 |
| classification | stabilization | Reaching B | single_day | GY_data_day-2 | GY_data_day-2 | 46 | accuracy | 0.883333 |
| classification | stabilization | Reaching B | single_day | GY_data_day-2 | GY_data_day-2 | 54 | accuracy | 0.883333 |
| classification | stabilization | Reaching B | single_day | GY_data_day-20 | GY_data_day-20 | 44 | accuracy | 0.685714 |
| classification | stabilization | Reaching B | single_day | GY_data_day-20 | GY_data_day-20 | 46 | accuracy | 0.857143 |
| classification | stabilization | Reaching B | single_day | GY_data_day-20 | GY_data_day-20 | 54 | accuracy | 0.828571 |
| classification | stabilization | Reaching B | single_day | GY_data_day-21 | GY_data_day-21 | 44 | accuracy | 0.564815 |
| classification | stabilization | Reaching B | single_day | GY_data_day-21 | GY_data_day-21 | 46 | accuracy | 0.583333 |
| classification | stabilization | Reaching B | single_day | GY_data_day-21 | GY_data_day-21 | 54 | accuracy | 0.537037 |
| classification | stabilization | Reaching B | single_day | GY_data_day-22 | GY_data_day-22 | 44 | accuracy | 0.924603 |
| classification | stabilization | Reaching B | single_day | GY_data_day-22 | GY_data_day-22 | 46 | accuracy | 0.892857 |
| classification | stabilization | Reaching B | single_day | GY_data_day-22 | GY_data_day-22 | 54 | accuracy | 0.924603 |
| classification | stabilization | Reaching B | single_day | GY_data_day-23 | GY_data_day-23 | 44 | accuracy | 0.927778 |
| classification | stabilization | Reaching B | single_day | GY_data_day-23 | GY_data_day-23 | 46 | accuracy | 0.955556 |
| classification | stabilization | Reaching B | single_day | GY_data_day-23 | GY_data_day-23 | 54 | accuracy | 0.922222 |
| classification | stabilization | Reaching B | single_day | GY_data_day-24 | GY_data_day-24 | 44 | accuracy | 0.805556 |
| classification | stabilization | Reaching B | single_day | GY_data_day-24 | GY_data_day-24 | 46 | accuracy | 0.847222 |
| classification | stabilization | Reaching B | single_day | GY_data_day-24 | GY_data_day-24 | 54 | accuracy | 0.875 |
| classification | stabilization | Reaching B | single_day | GY_data_day-25 | GY_data_day-25 | 44 | accuracy | 0.805556 |
| classification | stabilization | Reaching B | single_day | GY_data_day-25 | GY_data_day-25 | 46 | accuracy | 0.791667 |
| classification | stabilization | Reaching B | single_day | GY_data_day-25 | GY_data_day-25 | 54 | accuracy | 0.791667 |
| classification | stabilization | Reaching B | single_day | GY_data_day-26 | GY_data_day-26 | 44 | accuracy | 0.902778 |
| classification | stabilization | Reaching B | single_day | GY_data_day-26 | GY_data_day-26 | 46 | accuracy | 0.833333 |
| classification | stabilization | Reaching B | single_day | GY_data_day-26 | GY_data_day-26 | 54 | accuracy | 0.875 |
| classification | stabilization | Reaching B | single_day | GY_data_day-27 | GY_data_day-27 | 44 | accuracy | 0.861111 |
| classification | stabilization | Reaching B | single_day | GY_data_day-27 | GY_data_day-27 | 46 | accuracy | 0.888889 |
| classification | stabilization | Reaching B | single_day | GY_data_day-27 | GY_data_day-27 | 54 | accuracy | 0.791667 |
| classification | stabilization | Reaching B | single_day | GY_data_day-28 | GY_data_day-28 | 44 | accuracy | 0.819444 |
| classification | stabilization | Reaching B | single_day | GY_data_day-28 | GY_data_day-28 | 46 | accuracy | 0.722222 |
| classification | stabilization | Reaching B | single_day | GY_data_day-28 | GY_data_day-28 | 54 | accuracy | 0.902778 |
| classification | stabilization | Reaching B | single_day | GY_data_day-29 | GY_data_day-29 | 44 | accuracy | 0.777778 |
| classification | stabilization | Reaching B | single_day | GY_data_day-29 | GY_data_day-29 | 46 | accuracy | 0.875 |
| classification | stabilization | Reaching B | single_day | GY_data_day-29 | GY_data_day-29 | 54 | accuracy | 0.875 |
| classification | stabilization | Reaching B | single_day | GY_data_day-3 | GY_data_day-3 | 44 | accuracy | 0.787037 |

| | | | | | | | | |
|---|---|---|---|---|---|---|---|---|
| classification | stabilization | Reaching B | single_day | GY_data_day-3 | GY_data_day-3 | 46 | accuracy | 0.777778 |
| classification | stabilization | Reaching B | single_day | GY_data_day-3 | GY_data_day-3 | 54 | accuracy | 0.768519 |
| classification | stabilization | Reaching B | single_day | GY_data_day-30 | GY_data_day-30 | 44 | accuracy | 0.736111 |
| classification | stabilization | Reaching B | single_day | GY_data_day-30 | GY_data_day-30 | 46 | accuracy | 0.819444 |
| classification | stabilization | Reaching B | single_day | GY_data_day-30 | GY_data_day-30 | 54 | accuracy | 0.875 |
| classification | stabilization | Reaching B | single_day | GY_data_day-31 | GY_data_day-31 | 44 | accuracy | 0.4 |
| classification | stabilization | Reaching B | single_day | GY_data_day-31 | GY_data_day-31 | 46 | accuracy | 0.457143 |
| classification | stabilization | Reaching B | single_day | GY_data_day-31 | GY_data_day-31 | 54 | accuracy | 0.628571 |
| classification | stabilization | Reaching B | single_day | GY_data_day-32 | GY_data_day-32 | 44 | accuracy | 0.736111 |
| classification | stabilization | Reaching B | single_day | GY_data_day-32 | GY_data_day-32 | 46 | accuracy | 0.805556 |
| classification | stabilization | Reaching B | single_day | GY_data_day-32 | GY_data_day-32 | 54 | accuracy | 0.763889 |
| classification | stabilization | Reaching B | single_day | GY_data_day-33 | GY_data_day-33 | 44 | accuracy | 0.777778 |
| classification | stabilization | Reaching B | single_day | GY_data_day-33 | GY_data_day-33 | 46 | accuracy | 0.75 |
| classification | stabilization | Reaching B | single_day | GY_data_day-33 | GY_data_day-33 | 54 | accuracy | 0.796296 |
| classification | stabilization | Reaching B | single_day | GY_data_day-34 | GY_data_day-34 | 44 | accuracy | 0.9375 |
| classification | stabilization | Reaching B | single_day | GY_data_day-34 | GY_data_day-34 | 46 | accuracy | 0.951389 |
| classification | stabilization | Reaching B | single_day | GY_data_day-34 | GY_data_day-34 | 54 | accuracy | 0.923611 |
| classification | stabilization | Reaching B | single_day | GY_data_day-35 | GY_data_day-35 | 44 | accuracy | 0.805556 |
| classification | stabilization | Reaching B | single_day | GY_data_day-35 | GY_data_day-35 | 46 | accuracy | 0.8125 |
| classification | stabilization | Reaching B | single_day | GY_data_day-35 | GY_data_day-35 | 54 | accuracy | 0.868056 |
| classification | stabilization | Reaching B | single_day | GY_data_day-36 | GY_data_day-36 | 44 | accuracy | 0.895833 |
| classification | stabilization | Reaching B | single_day | GY_data_day-36 | GY_data_day-36 | 46 | accuracy | 0.888889 |
| classification | stabilization | Reaching B | single_day | GY_data_day-36 | GY_data_day-36 | 54 | accuracy | 0.868056 |
| classification | stabilization | Reaching B | single_day | GY_data_day-37 | GY_data_day-37 | 44 | accuracy | 0.888889 |
| classification | stabilization | Reaching B | single_day | GY_data_day-37 | GY_data_day-37 | 46 | accuracy | 0.847222 |
| classification | stabilization | Reaching B | single_day | GY_data_day-37 | GY_data_day-37 | 54 | accuracy | 0.819444 |
| classification | stabilization | Reaching B | single_day | GY_data_day-4 | GY_data_day-4 | 44 | accuracy | 0.883333 |
| classification | stabilization | Reaching B | single_day | GY_data_day-4 | GY_data_day-4 | 46 | accuracy | 0.916667 |
| classification | stabilization | Reaching B | single_day | GY_data_day-4 | GY_data_day-4 | 54 | accuracy | 0.905556 |
| classification | stabilization | Reaching B | single_day | GY_data_day-5 | GY_data_day-5 | 44 | accuracy | 0.74537 |
| classification | stabilization | Reaching B | single_day | GY_data_day-5 | GY_data_day-5 | 46 | accuracy | 0.791667 |
| classification | stabilization | Reaching B | single_day | GY_data_day-5 | GY_data_day-5 | 54 | accuracy | 0.824074 |
| classification | stabilization | Reaching B | single_day | GY_data_day-6 | GY_data_day-6 | 44 | accuracy | 0.699074 |
| classification | stabilization | Reaching B | single_day | GY_data_day-6 | GY_data_day-6 | 46 | accuracy | 0.699074 |
| classification | stabilization | Reaching B | single_day | GY_data_day-6 | GY_data_day-6 | 54 | accuracy | 0.657407 |
| classification | stabilization | Reaching B | single_day | GY_data_day-7 | GY_data_day-7 | 44 | accuracy | 0.444444 |
| classification | stabilization | Reaching B | single_day | GY_data_day-7 | GY_data_day-7 | 46 | accuracy | 0.462963 |
| classification | stabilization | Reaching B | single_day | GY_data_day-7 | GY_data_day-7 | 54 | accuracy | 0.453704 |
| classification | stabilization | Reaching B | single_day | GY_data_day-8 | GY_data_day-8 | 44 | accuracy | 0.847222 |
| classification | stabilization | Reaching B | single_day | GY_data_day-8 | GY_data_day-8 | 46 | accuracy | 0.8125 |
| classification | stabilization | Reaching B | single_day | GY_data_day-8 | GY_data_day-8 | 54 | accuracy | 0.854167 |
| classification | stabilization | Reaching B | single_day | GY_data_day-9 | GY_data_day-9 | 44 | accuracy | 0.861111 |
| classification | stabilization | Reaching B | single_day | GY_data_day-9 | GY_data_day-9 | 46 | accuracy | 0.930556 |
| classification | stabilization | Reaching B | single_day | GY_data_day-9 | GY_data_day-9 | 54 | accuracy | 0.888889 |
| classification | mfsnn | Reaching B | single_day | GY_data_day-1 | GY_data_day-1 | 44 | accuracy | 0.782353 |
| classification | mfsnn | Reaching B | single_day | GY_data_day-1 | GY_data_day-1 | 46 | accuracy | 0.623529 |
| classification | mfsnn | Reaching B | single_day | GY_data_day-1 | GY_data_day-1 | 54 | accuracy | 0.529412 |
| classification | mfsnn | Reaching B | single_day | GY_data_day-10 | GY_data_day-10 | 44 | accuracy | 0.45 |
| classification | mfsnn | Reaching B | single_day | GY_data_day-10 | GY_data_day-10 | 46 | accuracy | 0.4875 |
| classification | mfsnn | Reaching B | single_day | GY_data_day-10 | GY_data_day-10 | 54 | accuracy | 0.6375 |
| classification | mfsnn | Reaching B | single_day | GY_data_day-11 | GY_data_day-11 | 44 | accuracy | 0.7875 |
| classification | mfsnn | Reaching B | single_day | GY_data_day-11 | GY_data_day-11 | 46 | accuracy | 0.75 |
| classification | mfsnn | Reaching B | single_day | GY_data_day-11 | GY_data_day-11 | 54 | accuracy | 0.8125 |
| classification | mfsnn | Reaching B | single_day | GY_data_day-12 | GY_data_day-12 | 44 | accuracy | 0.639535 |
| classification | mfsnn | Reaching B | single_day | GY_data_day-12 | GY_data_day-12 | 46 | accuracy | 0.563953 |
| classification | mfsnn | Reaching B | single_day | GY_data_day-12 | GY_data_day-12 | 54 | accuracy | 0.447674 |
| classification | mfsnn | Reaching B | single_day | GY_data_day-13 | GY_data_day-13 | 44 | accuracy | 0.7875 |
| classification | mfsnn | Reaching B | single_day | GY_data_day-13 | GY_data_day-13 | 46 | accuracy | 0.825 |
| classification | mfsnn | Reaching B | single_day | GY_data_day-13 | GY_data_day-13 | 54 | accuracy | 0.8375 |
| classification | mfsnn | Reaching B | single_day | GY_data_day-14 | GY_data_day-14 | 44 | accuracy | 0.75 |
| classification | mfsnn | Reaching B | single_day | GY_data_day-14 | GY_data_day-14 | 46 | accuracy | 0.766667 |
| classification | mfsnn | Reaching B | single_day | GY_data_day-14 | GY_data_day-14 | 54 | accuracy | 0.675 |
| classification | mfsnn | Reaching B | single_day | GY_data_day-15 | GY_data_day-15 | 44 | accuracy | 0.66875 |

| | | | | | | | | |
|---|---|---|---|---|---|---|---|---|
| classification | mfsnn | Reaching B | single_day | GY_data_day-15 | GY_data_day-15 | 46 | accuracy | 0.775 |
| classification | mfsnn | Reaching B | single_day | GY_data_day-15 | GY_data_day-15 | 54 | accuracy | 0.625 |
| classification | mfsnn | Reaching B | single_day | GY_data_day-16 | GY_data_day-16 | 44 | accuracy | 0.7 |
| classification | mfsnn | Reaching B | single_day | GY_data_day-16 | GY_data_day-16 | 46 | accuracy | 0.9 |
| classification | mfsnn | Reaching B | single_day | GY_data_day-16 | GY_data_day-16 | 54 | accuracy | 0.75 |
| classification | mfsnn | Reaching B | single_day | GY_data_day-17 | GY_data_day-17 | 44 | accuracy | 0.85 |
| classification | mfsnn | Reaching B | single_day | GY_data_day-17 | GY_data_day-17 | 46 | accuracy | 0.8 |
| classification | mfsnn | Reaching B | single_day | GY_data_day-17 | GY_data_day-17 | 54 | accuracy | 0.841667 |
| classification | mfsnn | Reaching B | single_day | GY_data_day-18 | GY_data_day-18 | 44 | accuracy | 0.775 |
| classification | mfsnn | Reaching B | single_day | GY_data_day-18 | GY_data_day-18 | 46 | accuracy | 0.7875 |
| classification | mfsnn | Reaching B | single_day | GY_data_day-18 | GY_data_day-18 | 54 | accuracy | 0.7 |
| classification | mfsnn | Reaching B | single_day | GY_data_day-19 | GY_data_day-19 | 44 | accuracy | 0.4125 |
| classification | mfsnn | Reaching B | single_day | GY_data_day-19 | GY_data_day-19 | 46 | accuracy | 0.475 |
| classification | mfsnn | Reaching B | single_day | GY_data_day-19 | GY_data_day-19 | 54 | accuracy | 0.45 |
| classification | mfsnn | Reaching B | single_day | GY_data_day-2 | GY_data_day-2 | 44 | accuracy | 0.785 |
| classification | mfsnn | Reaching B | single_day | GY_data_day-2 | GY_data_day-2 | 46 | accuracy | 0.845 |
| classification | mfsnn | Reaching B | single_day | GY_data_day-2 | GY_data_day-2 | 54 | accuracy | 0.745 |
| classification | mfsnn | Reaching B | single_day | GY_data_day-20 | GY_data_day-20 | 44 | accuracy | 0.675 |
| classification | mfsnn | Reaching B | single_day | GY_data_day-20 | GY_data_day-20 | 46 | accuracy | 0.625 |
| classification | mfsnn | Reaching B | single_day | GY_data_day-20 | GY_data_day-20 | 54 | accuracy | 0.725 |
| classification | mfsnn | Reaching B | single_day | GY_data_day-21 | GY_data_day-21 | 44 | accuracy | 0.466667 |
| classification | mfsnn | Reaching B | single_day | GY_data_day-21 | GY_data_day-21 | 46 | accuracy | 0.491667 |
| classification | mfsnn | Reaching B | single_day | GY_data_day-21 | GY_data_day-21 | 54 | accuracy | 0.4 |
| classification | mfsnn | Reaching B | single_day | GY_data_day-22 | GY_data_day-22 | 44 | accuracy | 0.685714 |
| classification | mfsnn | Reaching B | single_day | GY_data_day-22 | GY_data_day-22 | 46 | accuracy | 0.735714 |
| classification | mfsnn | Reaching B | single_day | GY_data_day-22 | GY_data_day-22 | 54 | accuracy | 0.821429 |
| classification | mfsnn | Reaching B | single_day | GY_data_day-23 | GY_data_day-23 | 44 | accuracy | 0.715 |
| classification | mfsnn | Reaching B | single_day | GY_data_day-23 | GY_data_day-23 | 46 | accuracy | 0.645 |
| classification | mfsnn | Reaching B | single_day | GY_data_day-23 | GY_data_day-23 | 54 | accuracy | 0.68 |
| classification | mfsnn | Reaching B | single_day | GY_data_day-24 | GY_data_day-24 | 44 | accuracy | 0.775 |
| classification | mfsnn | Reaching B | single_day | GY_data_day-24 | GY_data_day-24 | 46 | accuracy | 0.6875 |
| classification | mfsnn | Reaching B | single_day | GY_data_day-24 | GY_data_day-24 | 54 | accuracy | 0.7375 |
| classification | mfsnn | Reaching B | single_day | GY_data_day-25 | GY_data_day-25 | 44 | accuracy | 0.725 |
| classification | mfsnn | Reaching B | single_day | GY_data_day-25 | GY_data_day-25 | 46 | accuracy | 0.75 |
| classification | mfsnn | Reaching B | single_day | GY_data_day-25 | GY_data_day-25 | 54 | accuracy | 0.5875 |
| classification | mfsnn | Reaching B | single_day | GY_data_day-26 | GY_data_day-26 | 44 | accuracy | 0.6875 |
| classification | mfsnn | Reaching B | single_day | GY_data_day-26 | GY_data_day-26 | 46 | accuracy | 0.6375 |
| classification | mfsnn | Reaching B | single_day | GY_data_day-26 | GY_data_day-26 | 54 | accuracy | 0.725 |
| classification | mfsnn | Reaching B | single_day | GY_data_day-27 | GY_data_day-27 | 44 | accuracy | 0.5625 |
| classification | mfsnn | Reaching B | single_day | GY_data_day-27 | GY_data_day-27 | 46 | accuracy | 0.775 |
| classification | mfsnn | Reaching B | single_day | GY_data_day-27 | GY_data_day-27 | 54 | accuracy | 0.725 |
| classification | mfsnn | Reaching B | single_day | GY_data_day-28 | GY_data_day-28 | 44 | accuracy | 0.6625 |
| classification | mfsnn | Reaching B | single_day | GY_data_day-28 | GY_data_day-28 | 46 | accuracy | 0.6875 |
| classification | mfsnn | Reaching B | single_day | GY_data_day-28 | GY_data_day-28 | 54 | accuracy | 0.75 |
| classification | mfsnn | Reaching B | single_day | GY_data_day-29 | GY_data_day-29 | 44 | accuracy | 0.6125 |
| classification | mfsnn | Reaching B | single_day | GY_data_day-29 | GY_data_day-29 | 46 | accuracy | 0.8375 |
| classification | mfsnn | Reaching B | single_day | GY_data_day-29 | GY_data_day-29 | 54 | accuracy | 0.7375 |
| classification | mfsnn | Reaching B | single_day | GY_data_day-3 | GY_data_day-3 | 44 | accuracy | 0.733333 |
| classification | mfsnn | Reaching B | single_day | GY_data_day-3 | GY_data_day-3 | 46 | accuracy | 0.7 |
| classification | mfsnn | Reaching B | single_day | GY_data_day-3 | GY_data_day-3 | 54 | accuracy | 0.675 |
| classification | mfsnn | Reaching B | single_day | GY_data_day-30 | GY_data_day-30 | 44 | accuracy | 0.675 |
| classification | mfsnn | Reaching B | single_day | GY_data_day-30 | GY_data_day-30 | 46 | accuracy | 0.7125 |
| classification | mfsnn | Reaching B | single_day | GY_data_day-30 | GY_data_day-30 | 54 | accuracy | 0.4625 |
| classification | mfsnn | Reaching B | single_day | GY_data_day-31 | GY_data_day-31 | 44 | accuracy | 0.725 |
| classification | mfsnn | Reaching B | single_day | GY_data_day-31 | GY_data_day-31 | 46 | accuracy | 0.45 |
| classification | mfsnn | Reaching B | single_day | GY_data_day-31 | GY_data_day-31 | 54 | accuracy | 0.6 |
| classification | mfsnn | Reaching B | single_day | GY_data_day-32 | GY_data_day-32 | 44 | accuracy | 0.6375 |
| classification | mfsnn | Reaching B | single_day | GY_data_day-32 | GY_data_day-32 | 46 | accuracy | 0.7125 |
| classification | mfsnn | Reaching B | single_day | GY_data_day-32 | GY_data_day-32 | 54 | accuracy | 0.7375 |
| classification | mfsnn | Reaching B | single_day | GY_data_day-33 | GY_data_day-33 | 44 | accuracy | 0.75 |
| classification | mfsnn | Reaching B | single_day | GY_data_day-33 | GY_data_day-33 | 46 | accuracy | 0.625 |
| classification | mfsnn | Reaching B | single_day | GY_data_day-33 | GY_data_day-33 | 54 | accuracy | 0.766667 |
| classification | mfsnn | Reaching B | single_day | GY_data_day-34 | GY_data_day-34 | 44 | accuracy | 0.73125 |

| | | | | | | | | |
|---|---|---|---|---|---|---|---|---|
| classification | mfsnn | Reaching B | single_day | GY_data_day-34 | GY_data_day-34 | 46 | accuracy | 0.6125 |
| classification | mfsnn | Reaching B | single_day | GY_data_day-34 | GY_data_day-34 | 54 | accuracy | 0.8 |
| classification | mfsnn | Reaching B | single_day | GY_data_day-35 | GY_data_day-35 | 44 | accuracy | 0.7875 |
| classification | mfsnn | Reaching B | single_day | GY_data_day-35 | GY_data_day-35 | 46 | accuracy | 0.71875 |
| classification | mfsnn | Reaching B | single_day | GY_data_day-35 | GY_data_day-35 | 54 | accuracy | 0.75 |
| classification | mfsnn | Reaching B | single_day | GY_data_day-36 | GY_data_day-36 | 44 | accuracy | 0.6375 |
| classification | mfsnn | Reaching B | single_day | GY_data_day-36 | GY_data_day-36 | 46 | accuracy | 0.6125 |
| classification | mfsnn | Reaching B | single_day | GY_data_day-36 | GY_data_day-36 | 54 | accuracy | 0.75625 |
| classification | mfsnn | Reaching B | single_day | GY_data_day-37 | GY_data_day-37 | 44 | accuracy | 0.8 |
| classification | mfsnn | Reaching B | single_day | GY_data_day-37 | GY_data_day-37 | 46 | accuracy | 0.725 |
| classification | mfsnn | Reaching B | single_day | GY_data_day-37 | GY_data_day-37 | 54 | accuracy | 0.7875 |
| classification | mfsnn | Reaching B | single_day | GY_data_day-4 | GY_data_day-4 | 44 | accuracy | 0.645 |
| classification | mfsnn | Reaching B | single_day | GY_data_day-4 | GY_data_day-4 | 46 | accuracy | 0.64 |
| classification | mfsnn | Reaching B | single_day | GY_data_day-4 | GY_data_day-4 | 54 | accuracy | 0.825 |
| classification | mfsnn | Reaching B | single_day | GY_data_day-5 | GY_data_day-5 | 44 | accuracy | 0.729167 |
| classification | mfsnn | Reaching B | single_day | GY_data_day-5 | GY_data_day-5 | 46 | accuracy | 0.775 |
| classification | mfsnn | Reaching B | single_day | GY_data_day-5 | GY_data_day-5 | 54 | accuracy | 0.733333 |
| classification | mfsnn | Reaching B | single_day | GY_data_day-6 | GY_data_day-6 | 44 | accuracy | 0.6375 |
| classification | mfsnn | Reaching B | single_day | GY_data_day-6 | GY_data_day-6 | 46 | accuracy | 0.716667 |
| classification | mfsnn | Reaching B | single_day | GY_data_day-6 | GY_data_day-6 | 54 | accuracy | 0.629167 |
| classification | mfsnn | Reaching B | single_day | GY_data_day-7 | GY_data_day-7 | 44 | accuracy | 0.3875 |
| classification | mfsnn | Reaching B | single_day | GY_data_day-7 | GY_data_day-7 | 46 | accuracy | 0.4375 |
| classification | mfsnn | Reaching B | single_day | GY_data_day-7 | GY_data_day-7 | 54 | accuracy | 0.345833 |
| classification | mfsnn | Reaching B | single_day | GY_data_day-8 | GY_data_day-8 | 44 | accuracy | 0.61875 |
| classification | mfsnn | Reaching B | single_day | GY_data_day-8 | GY_data_day-8 | 46 | accuracy | 0.7375 |
| classification | mfsnn | Reaching B | single_day | GY_data_day-8 | GY_data_day-8 | 54 | accuracy | 0.4125 |
| classification | mfsnn | Reaching B | single_day | GY_data_day-9 | GY_data_day-9 | 44 | accuracy | 0.725 |
| classification | mfsnn | Reaching B | single_day | GY_data_day-9 | GY_data_day-9 | 46 | accuracy | 0.7375 |
| classification | mfsnn | Reaching B | single_day | GY_data_day-9 | GY_data_day-9 | 54 | accuracy | 0.6625 |
| classification | mscformer | Reaching B | single_day | GY_data_day-1 | GY_data_day-1 | 44 | accuracy | 0.941176 |
| classification | mscformer | Reaching B | single_day | GY_data_day-1 | GY_data_day-1 | 46 | accuracy | 0.923529 |
| classification | mscformer | Reaching B | single_day | GY_data_day-1 | GY_data_day-1 | 54 | accuracy | 0.952941 |
| classification | mscformer | Reaching B | single_day | GY_data_day-10 | GY_data_day-10 | 44 | accuracy | 0.625 |
| classification | mscformer | Reaching B | single_day | GY_data_day-10 | GY_data_day-10 | 46 | accuracy | 0.7125 |
| classification | mscformer | Reaching B | single_day | GY_data_day-10 | GY_data_day-10 | 54 | accuracy | 0.6875 |
| classification | mscformer | Reaching B | single_day | GY_data_day-11 | GY_data_day-11 | 44 | accuracy | 0.875 |
| classification | mscformer | Reaching B | single_day | GY_data_day-11 | GY_data_day-11 | 46 | accuracy | 0.9375 |
| classification | mscformer | Reaching B | single_day | GY_data_day-11 | GY_data_day-11 | 54 | accuracy | 0.9 |
| classification | mscformer | Reaching B | single_day | GY_data_day-12 | GY_data_day-12 | 44 | accuracy | 0.994186 |
| classification | mscformer | Reaching B | single_day | GY_data_day-12 | GY_data_day-12 | 46 | accuracy | 0.959302 |
| classification | mscformer | Reaching B | single_day | GY_data_day-12 | GY_data_day-12 | 54 | accuracy | 0.930233 |
| classification | mscformer | Reaching B | single_day | GY_data_day-13 | GY_data_day-13 | 44 | accuracy | 0.9125 |
| classification | mscformer | Reaching B | single_day | GY_data_day-13 | GY_data_day-13 | 46 | accuracy | 0.9125 |
| classification | mscformer | Reaching B | single_day | GY_data_day-13 | GY_data_day-13 | 54 | accuracy | 0.9 |
| classification | mscformer | Reaching B | single_day | GY_data_day-14 | GY_data_day-14 | 44 | accuracy | 0.958333 |
| classification | mscformer | Reaching B | single_day | GY_data_day-14 | GY_data_day-14 | 46 | accuracy | 0.883333 |
| classification | mscformer | Reaching B | single_day | GY_data_day-14 | GY_data_day-14 | 54 | accuracy | 0.933333 |
| classification | mscformer | Reaching B | single_day | GY_data_day-15 | GY_data_day-15 | 44 | accuracy | 0.93125 |
| classification | mscformer | Reaching B | single_day | GY_data_day-15 | GY_data_day-15 | 46 | accuracy | 0.95 |
| classification | mscformer | Reaching B | single_day | GY_data_day-15 | GY_data_day-15 | 54 | accuracy | 0.94375 |
| classification | mscformer | Reaching B | single_day | GY_data_day-16 | GY_data_day-16 | 44 | accuracy | 0.975 |
| classification | mscformer | Reaching B | single_day | GY_data_day-16 | GY_data_day-16 | 46 | accuracy | 0.95 |
| classification | mscformer | Reaching B | single_day | GY_data_day-16 | GY_data_day-16 | 54 | accuracy | 0.9 |
| classification | mscformer | Reaching B | single_day | GY_data_day-17 | GY_data_day-17 | 44 | accuracy | 0.966667 |
| classification | mscformer | Reaching B | single_day | GY_data_day-17 | GY_data_day-17 | 46 | accuracy | 0.966667 |
| classification | mscformer | Reaching B | single_day | GY_data_day-17 | GY_data_day-17 | 54 | accuracy | 0.983333 |
| classification | mscformer | Reaching B | single_day | GY_data_day-18 | GY_data_day-18 | 44 | accuracy | 0.95 |
| classification | mscformer | Reaching B | single_day | GY_data_day-18 | GY_data_day-18 | 46 | accuracy | 0.95 |
| classification | mscformer | Reaching B | single_day | GY_data_day-18 | GY_data_day-18 | 54 | accuracy | 0.9625 |
| classification | mscformer | Reaching B | single_day | GY_data_day-19 | GY_data_day-19 | 44 | accuracy | 0.9375 |
| classification | mscformer | Reaching B | single_day | GY_data_day-19 | GY_data_day-19 | 46 | accuracy | 0.9375 |
| classification | mscformer | Reaching B | single_day | GY_data_day-19 | GY_data_day-19 | 54 | accuracy | 0.95 |
| classification | mscformer | Reaching B | single_day | GY_data_day-2 | GY_data_day-2 | 44 | accuracy | 0.96 |

| | | | | | | | | |
|---|---|---|---|---|---|---|---|---|
| classification | mscformer | Reaching B | single_day | GY_data_day-2 | GY_data_day-2 | 46 | accuracy | 0.915 |
| classification | mscformer | Reaching B | single_day | GY_data_day-2 | GY_data_day-2 | 54 | accuracy | 0.935 |
| classification | mscformer | Reaching B | single_day | GY_data_day-20 | GY_data_day-20 | 44 | accuracy | 0.9 |
| classification | mscformer | Reaching B | single_day | GY_data_day-20 | GY_data_day-20 | 46 | accuracy | 0.9 |
| classification | mscformer | Reaching B | single_day | GY_data_day-20 | GY_data_day-20 | 54 | accuracy | 0.8 |
| classification | mscformer | Reaching B | single_day | GY_data_day-21 | GY_data_day-21 | 44 | accuracy | 0.591667 |
| classification | mscformer | Reaching B | single_day | GY_data_day-21 | GY_data_day-21 | 46 | accuracy | 0.483333 |
| classification | mscformer | Reaching B | single_day | GY_data_day-21 | GY_data_day-21 | 54 | accuracy | 0.55 |
| classification | mscformer | Reaching B | single_day | GY_data_day-22 | GY_data_day-22 | 44 | accuracy | 0.964286 |
| classification | mscformer | Reaching B | single_day | GY_data_day-22 | GY_data_day-22 | 46 | accuracy | 0.978571 |
| classification | mscformer | Reaching B | single_day | GY_data_day-22 | GY_data_day-22 | 54 | accuracy | 0.95 |
| classification | mscformer | Reaching B | single_day | GY_data_day-23 | GY_data_day-23 | 44 | accuracy | 0.96 |
| classification | mscformer | Reaching B | single_day | GY_data_day-23 | GY_data_day-23 | 46 | accuracy | 0.97 |
| classification | mscformer | Reaching B | single_day | GY_data_day-23 | GY_data_day-23 | 54 | accuracy | 0.975 |
| classification | mscformer | Reaching B | single_day | GY_data_day-24 | GY_data_day-24 | 44 | accuracy | 0.925 |
| classification | mscformer | Reaching B | single_day | GY_data_day-24 | GY_data_day-24 | 46 | accuracy | 0.9 |
| classification | mscformer | Reaching B | single_day | GY_data_day-24 | GY_data_day-24 | 54 | accuracy | 0.8875 |
| classification | mscformer | Reaching B | single_day | GY_data_day-25 | GY_data_day-25 | 44 | accuracy | 0.875 |
| classification | mscformer | Reaching B | single_day | GY_data_day-25 | GY_data_day-25 | 46 | accuracy | 0.9125 |
| classification | mscformer | Reaching B | single_day | GY_data_day-25 | GY_data_day-25 | 54 | accuracy | 0.8625 |
| classification | mscformer | Reaching B | single_day | GY_data_day-26 | GY_data_day-26 | 44 | accuracy | 0.9 |
| classification | mscformer | Reaching B | single_day | GY_data_day-26 | GY_data_day-26 | 46 | accuracy | 0.9125 |
| classification | mscformer | Reaching B | single_day | GY_data_day-26 | GY_data_day-26 | 54 | accuracy | 0.8875 |
| classification | mscformer | Reaching B | single_day | GY_data_day-27 | GY_data_day-27 | 44 | accuracy | 0.9625 |
| classification | mscformer | Reaching B | single_day | GY_data_day-27 | GY_data_day-27 | 46 | accuracy | 0.9375 |
| classification | mscformer | Reaching B | single_day | GY_data_day-27 | GY_data_day-27 | 54 | accuracy | 0.95 |
| classification | mscformer | Reaching B | single_day | GY_data_day-28 | GY_data_day-28 | 44 | accuracy | 0.925 |
| classification | mscformer | Reaching B | single_day | GY_data_day-28 | GY_data_day-28 | 46 | accuracy | 0.925 |
| classification | mscformer | Reaching B | single_day | GY_data_day-28 | GY_data_day-28 | 54 | accuracy | 0.9 |
| classification | mscformer | Reaching B | single_day | GY_data_day-29 | GY_data_day-29 | 44 | accuracy | 0.9 |
| classification | mscformer | Reaching B | single_day | GY_data_day-29 | GY_data_day-29 | 46 | accuracy | 0.9 |
| classification | mscformer | Reaching B | single_day | GY_data_day-29 | GY_data_day-29 | 54 | accuracy | 0.9375 |
| classification | mscformer | Reaching B | single_day | GY_data_day-3 | GY_data_day-3 | 44 | accuracy | 0.875 |
| classification | mscformer | Reaching B | single_day | GY_data_day-3 | GY_data_day-3 | 46 | accuracy | 0.866667 |
| classification | mscformer | Reaching B | single_day | GY_data_day-3 | GY_data_day-3 | 54 | accuracy | 0.858333 |
| classification | mscformer | Reaching B | single_day | GY_data_day-30 | GY_data_day-30 | 44 | accuracy | 0.8875 |
| classification | mscformer | Reaching B | single_day | GY_data_day-30 | GY_data_day-30 | 46 | accuracy | 0.95 |
| classification | mscformer | Reaching B | single_day | GY_data_day-30 | GY_data_day-30 | 54 | accuracy | 0.9125 |
| classification | mscformer | Reaching B | single_day | GY_data_day-31 | GY_data_day-31 | 44 | accuracy | 0.85 |
| classification | mscformer | Reaching B | single_day | GY_data_day-31 | GY_data_day-31 | 46 | accuracy | 0.825 |
| classification | mscformer | Reaching B | single_day | GY_data_day-31 | GY_data_day-31 | 54 | accuracy | 0.725 |
| classification | mscformer | Reaching B | single_day | GY_data_day-32 | GY_data_day-32 | 44 | accuracy | 0.9125 |
| classification | mscformer | Reaching B | single_day | GY_data_day-32 | GY_data_day-32 | 46 | accuracy | 0.9125 |
| classification | mscformer | Reaching B | single_day | GY_data_day-32 | GY_data_day-32 | 54 | accuracy | 0.8875 |
| classification | mscformer | Reaching B | single_day | GY_data_day-33 | GY_data_day-33 | 44 | accuracy | 0.916667 |
| classification | mscformer | Reaching B | single_day | GY_data_day-33 | GY_data_day-33 | 46 | accuracy | 0.908333 |
| classification | mscformer | Reaching B | single_day | GY_data_day-33 | GY_data_day-33 | 54 | accuracy | 0.908333 |
| classification | mscformer | Reaching B | single_day | GY_data_day-34 | GY_data_day-34 | 44 | accuracy | 0.88125 |
| classification | mscformer | Reaching B | single_day | GY_data_day-34 | GY_data_day-34 | 46 | accuracy | 0.88125 |
| classification | mscformer | Reaching B | single_day | GY_data_day-34 | GY_data_day-34 | 54 | accuracy | 0.85625 |
| classification | mscformer | Reaching B | single_day | GY_data_day-35 | GY_data_day-35 | 44 | accuracy | 0.93125 |
| classification | mscformer | Reaching B | single_day | GY_data_day-35 | GY_data_day-35 | 46 | accuracy | 0.8875 |
| classification | mscformer | Reaching B | single_day | GY_data_day-35 | GY_data_day-35 | 54 | accuracy | 0.8625 |
| classification | mscformer | Reaching B | single_day | GY_data_day-36 | GY_data_day-36 | 44 | accuracy | 0.8875 |
| classification | mscformer | Reaching B | single_day | GY_data_day-36 | GY_data_day-36 | 46 | accuracy | 0.9 |
| classification | mscformer | Reaching B | single_day | GY_data_day-36 | GY_data_day-36 | 54 | accuracy | 0.9 |
| classification | mscformer | Reaching B | single_day | GY_data_day-37 | GY_data_day-37 | 44 | accuracy | 0.9125 |
| classification | mscformer | Reaching B | single_day | GY_data_day-37 | GY_data_day-37 | 46 | accuracy | 0.95 |
| classification | mscformer | Reaching B | single_day | GY_data_day-37 | GY_data_day-37 | 54 | accuracy | 0.9125 |
| classification | mscformer | Reaching B | single_day | GY_data_day-4 | GY_data_day-4 | 44 | accuracy | 0.915 |
| classification | mscformer | Reaching B | single_day | GY_data_day-4 | GY_data_day-4 | 46 | accuracy | 0.97 |
| classification | mscformer | Reaching B | single_day | GY_data_day-4 | GY_data_day-4 | 54 | accuracy | 0.96 |
| classification | mscformer | Reaching B | single_day | GY_data_day-5 | GY_data_day-5 | 44 | accuracy | 0.925 |

| | | | | | | | | |
|---|---|---|---|---|---|---|---|---|
| classification | mscformer | Reaching B | single_day | GY_data_day-5 | GY_data_day-5 | 46 | accuracy | 0.908333 |
| classification | mscformer | Reaching B | single_day | GY_data_day-5 | GY_data_day-5 | 54 | accuracy | 0.9 |
| classification | mscformer | Reaching B | single_day | GY_data_day-6 | GY_data_day-6 | 44 | accuracy | 0.754167 |
| classification | mscformer | Reaching B | single_day | GY_data_day-6 | GY_data_day-6 | 46 | accuracy | 0.75 |
| classification | mscformer | Reaching B | single_day | GY_data_day-6 | GY_data_day-6 | 54 | accuracy | 0.7625 |
| classification | mscformer | Reaching B | single_day | GY_data_day-7 | GY_data_day-7 | 44 | accuracy | 0.470833 |
| classification | mscformer | Reaching B | single_day | GY_data_day-7 | GY_data_day-7 | 46 | accuracy | 0.433333 |
| classification | mscformer | Reaching B | single_day | GY_data_day-7 | GY_data_day-7 | 54 | accuracy | 0.45 |
| classification | mscformer | Reaching B | single_day | GY_data_day-8 | GY_data_day-8 | 44 | accuracy | 0.95625 |
| classification | mscformer | Reaching B | single_day | GY_data_day-8 | GY_data_day-8 | 46 | accuracy | 0.93125 |
| classification | mscformer | Reaching B | single_day | GY_data_day-8 | GY_data_day-8 | 54 | accuracy | 0.89375 |
| classification | mscformer | Reaching B | single_day | GY_data_day-9 | GY_data_day-9 | 44 | accuracy | 0.9625 |
| classification | mscformer | Reaching B | single_day | GY_data_day-9 | GY_data_day-9 | 46 | accuracy | 0.9375 |
| classification | mscformer | Reaching B | single_day | GY_data_day-9 | GY_data_day-9 | 54 | accuracy | 0.9375 |
| classification | nomad | Reaching B | single_day | GY_data_day-1 | GY_data_day-1 | 44 | accuracy | 0.958824 |
| classification | nomad | Reaching B | single_day | GY_data_day-1 | GY_data_day-1 | 46 | accuracy | 0.929412 |
| classification | nomad | Reaching B | single_day | GY_data_day-1 | GY_data_day-1 | 54 | accuracy | 0.958824 |
| classification | nomad | Reaching B | single_day | GY_data_day-10 | GY_data_day-10 | 44 | accuracy | 0.68125 |
| classification | nomad | Reaching B | single_day | GY_data_day-10 | GY_data_day-10 | 46 | accuracy | 0.6875 |
| classification | nomad | Reaching B | single_day | GY_data_day-10 | GY_data_day-10 | 54 | accuracy | 0.64375 |
| classification | nomad | Reaching B | single_day | GY_data_day-11 | GY_data_day-11 | 44 | accuracy | 0.8625 |
| classification | nomad | Reaching B | single_day | GY_data_day-11 | GY_data_day-11 | 46 | accuracy | 0.825 |
| classification | nomad | Reaching B | single_day | GY_data_day-11 | GY_data_day-11 | 54 | accuracy | 0.8 |
| classification | nomad | Reaching B | single_day | GY_data_day-12 | GY_data_day-12 | 44 | accuracy | 0.982558 |
| classification | nomad | Reaching B | single_day | GY_data_day-12 | GY_data_day-12 | 46 | accuracy | 0.97093 |
| classification | nomad | Reaching B | single_day | GY_data_day-12 | GY_data_day-12 | 54 | accuracy | 0.965116 |
| classification | nomad | Reaching B | single_day | GY_data_day-13 | GY_data_day-13 | 44 | accuracy | 0.85 |
| classification | nomad | Reaching B | single_day | GY_data_day-13 | GY_data_day-13 | 46 | accuracy | 0.8375 |
| classification | nomad | Reaching B | single_day | GY_data_day-13 | GY_data_day-13 | 54 | accuracy | 0.9 |
| classification | nomad | Reaching B | single_day | GY_data_day-14 | GY_data_day-14 | 44 | accuracy | 0.95 |
| classification | nomad | Reaching B | single_day | GY_data_day-14 | GY_data_day-14 | 46 | accuracy | 0.966667 |
| classification | nomad | Reaching B | single_day | GY_data_day-14 | GY_data_day-14 | 54 | accuracy | 0.916667 |
| classification | nomad | Reaching B | single_day | GY_data_day-15 | GY_data_day-15 | 44 | accuracy | 0.95 |
| classification | nomad | Reaching B | single_day | GY_data_day-15 | GY_data_day-15 | 46 | accuracy | 0.9875 |
| classification | nomad | Reaching B | single_day | GY_data_day-15 | GY_data_day-15 | 54 | accuracy | 0.90625 |
| classification | nomad | Reaching B | single_day | GY_data_day-16 | GY_data_day-16 | 44 | accuracy | 0.975 |
| classification | nomad | Reaching B | single_day | GY_data_day-16 | GY_data_day-16 | 46 | accuracy | 0.825 |
| classification | nomad | Reaching B | single_day | GY_data_day-16 | GY_data_day-16 | 54 | accuracy | 0.775 |
| classification | nomad | Reaching B | single_day | GY_data_day-17 | GY_data_day-17 | 44 | accuracy | 0.95 |
| classification | nomad | Reaching B | single_day | GY_data_day-17 | GY_data_day-17 | 46 | accuracy | 0.975 |
| classification | nomad | Reaching B | single_day | GY_data_day-17 | GY_data_day-17 | 54 | accuracy | 0.958333 |
| classification | nomad | Reaching B | single_day | GY_data_day-18 | GY_data_day-18 | 44 | accuracy | 0.9 |
| classification | nomad | Reaching B | single_day | GY_data_day-18 | GY_data_day-18 | 46 | accuracy | 0.9375 |
| classification | nomad | Reaching B | single_day | GY_data_day-18 | GY_data_day-18 | 54 | accuracy | 0.9625 |
| classification | nomad | Reaching B | single_day | GY_data_day-19 | GY_data_day-19 | 44 | accuracy | 0.95 |
| classification | nomad | Reaching B | single_day | GY_data_day-19 | GY_data_day-19 | 46 | accuracy | 0.7375 |
| classification | nomad | Reaching B | single_day | GY_data_day-19 | GY_data_day-19 | 54 | accuracy | 0.925 |
| classification | nomad | Reaching B | single_day | GY_data_day-2 | GY_data_day-2 | 44 | accuracy | 0.975 |
| classification | nomad | Reaching B | single_day | GY_data_day-2 | GY_data_day-2 | 46 | accuracy | 0.97 |
| classification | nomad | Reaching B | single_day | GY_data_day-2 | GY_data_day-2 | 54 | accuracy | 0.97 |
| classification | nomad | Reaching B | single_day | GY_data_day-20 | GY_data_day-20 | 44 | accuracy | 0.85 |
| classification | nomad | Reaching B | single_day | GY_data_day-20 | GY_data_day-20 | 46 | accuracy | 0.75 |
| classification | nomad | Reaching B | single_day | GY_data_day-20 | GY_data_day-20 | 54 | accuracy | 0.85 |
| classification | nomad | Reaching B | single_day | GY_data_day-21 | GY_data_day-21 | 44 | accuracy | 0.633333 |
| classification | nomad | Reaching B | single_day | GY_data_day-21 | GY_data_day-21 | 46 | accuracy | 0.541667 |
| classification | nomad | Reaching B | single_day | GY_data_day-21 | GY_data_day-21 | 54 | accuracy | 0.575 |
| classification | nomad | Reaching B | single_day | GY_data_day-22 | GY_data_day-22 | 44 | accuracy | 0.957143 |
| classification | nomad | Reaching B | single_day | GY_data_day-22 | GY_data_day-22 | 46 | accuracy | 0.975 |
| classification | nomad | Reaching B | single_day | GY_data_day-22 | GY_data_day-22 | 54 | accuracy | 0.985714 |
| classification | nomad | Reaching B | single_day | GY_data_day-23 | GY_data_day-23 | 44 | accuracy | 0.965 |
| classification | nomad | Reaching B | single_day | GY_data_day-23 | GY_data_day-23 | 46 | accuracy | 0.935 |
| classification | nomad | Reaching B | single_day | GY_data_day-23 | GY_data_day-23 | 54 | accuracy | 0.965 |
| classification | nomad | Reaching B | single_day | GY_data_day-24 | GY_data_day-24 | 44 | accuracy | 0.9 |

| | | | | | | | | |
|---|---|---|---|---|---|---|---|---|
| classification | nomad | Reaching B | single_day | GY_data_day-24 | GY_data_day-24 | 46 | accuracy | 0.9125 |
| classification | nomad | Reaching B | single_day | GY_data_day-24 | GY_data_day-24 | 54 | accuracy | 0.9 |
| classification | nomad | Reaching B | single_day | GY_data_day-25 | GY_data_day-25 | 44 | accuracy | 0.9 |
| classification | nomad | Reaching B | single_day | GY_data_day-25 | GY_data_day-25 | 46 | accuracy | 0.925 |
| classification | nomad | Reaching B | single_day | GY_data_day-25 | GY_data_day-25 | 54 | accuracy | 0.8625 |
| classification | nomad | Reaching B | single_day | GY_data_day-26 | GY_data_day-26 | 44 | accuracy | 0.8625 |
| classification | nomad | Reaching B | single_day | GY_data_day-26 | GY_data_day-26 | 46 | accuracy | 0.875 |
| classification | nomad | Reaching B | single_day | GY_data_day-26 | GY_data_day-26 | 54 | accuracy | 0.8625 |
| classification | nomad | Reaching B | single_day | GY_data_day-27 | GY_data_day-27 | 44 | accuracy | 0.95 |
| classification | nomad | Reaching B | single_day | GY_data_day-27 | GY_data_day-27 | 46 | accuracy | 0.95 |
| classification | nomad | Reaching B | single_day | GY_data_day-27 | GY_data_day-27 | 54 | accuracy | 0.8125 |
| classification | nomad | Reaching B | single_day | GY_data_day-28 | GY_data_day-28 | 44 | accuracy | 0.975 |
| classification | nomad | Reaching B | single_day | GY_data_day-28 | GY_data_day-28 | 46 | accuracy | 0.9125 |
| classification | nomad | Reaching B | single_day | GY_data_day-28 | GY_data_day-28 | 54 | accuracy | 0.8625 |
| classification | nomad | Reaching B | single_day | GY_data_day-29 | GY_data_day-29 | 44 | accuracy | 0.925 |
| classification | nomad | Reaching B | single_day | GY_data_day-29 | GY_data_day-29 | 46 | accuracy | 0.8625 |
| classification | nomad | Reaching B | single_day | GY_data_day-29 | GY_data_day-29 | 54 | accuracy | 0.7875 |
| classification | nomad | Reaching B | single_day | GY_data_day-3 | GY_data_day-3 | 44 | accuracy | 0.933333 |
| classification | nomad | Reaching B | single_day | GY_data_day-3 | GY_data_day-3 | 46 | accuracy | 0.958333 |
| classification | nomad | Reaching B | single_day | GY_data_day-3 | GY_data_day-3 | 54 | accuracy | 0.933333 |
| classification | nomad | Reaching B | single_day | GY_data_day-30 | GY_data_day-30 | 44 | accuracy | 0.8375 |
| classification | nomad | Reaching B | single_day | GY_data_day-30 | GY_data_day-30 | 46 | accuracy | 0.875 |
| classification | nomad | Reaching B | single_day | GY_data_day-30 | GY_data_day-30 | 54 | accuracy | 0.9125 |
| classification | nomad | Reaching B | single_day | GY_data_day-31 | GY_data_day-31 | 44 | accuracy | 0.675 |
| classification | nomad | Reaching B | single_day | GY_data_day-31 | GY_data_day-31 | 46 | accuracy | 0.575 |
| classification | nomad | Reaching B | single_day | GY_data_day-31 | GY_data_day-31 | 54 | accuracy | 0.675 |
| classification | nomad | Reaching B | single_day | GY_data_day-32 | GY_data_day-32 | 44 | accuracy | 0.9 |
| classification | nomad | Reaching B | single_day | GY_data_day-32 | GY_data_day-32 | 46 | accuracy | 0.8625 |
| classification | nomad | Reaching B | single_day | GY_data_day-32 | GY_data_day-32 | 54 | accuracy | 0.7875 |
| classification | nomad | Reaching B | single_day | GY_data_day-33 | GY_data_day-33 | 44 | accuracy | 0.933333 |
| classification | nomad | Reaching B | single_day | GY_data_day-33 | GY_data_day-33 | 46 | accuracy | 0.891667 |
| classification | nomad | Reaching B | single_day | GY_data_day-33 | GY_data_day-33 | 54 | accuracy | 0.791667 |
| classification | nomad | Reaching B | single_day | GY_data_day-34 | GY_data_day-34 | 44 | accuracy | 0.925 |
| classification | nomad | Reaching B | single_day | GY_data_day-34 | GY_data_day-34 | 46 | accuracy | 0.93125 |
| classification | nomad | Reaching B | single_day | GY_data_day-34 | GY_data_day-34 | 54 | accuracy | 0.93125 |
| classification | nomad | Reaching B | single_day | GY_data_day-35 | GY_data_day-35 | 44 | accuracy | 0.88125 |
| classification | nomad | Reaching B | single_day | GY_data_day-35 | GY_data_day-35 | 46 | accuracy | 0.9 |
| classification | nomad | Reaching B | single_day | GY_data_day-35 | GY_data_day-35 | 54 | accuracy | 0.9 |
| classification | nomad | Reaching B | single_day | GY_data_day-36 | GY_data_day-36 | 44 | accuracy | 0.93125 |
| classification | nomad | Reaching B | single_day | GY_data_day-36 | GY_data_day-36 | 46 | accuracy | 0.89375 |
| classification | nomad | Reaching B | single_day | GY_data_day-36 | GY_data_day-36 | 54 | accuracy | 0.8625 |
| classification | nomad | Reaching B | single_day | GY_data_day-37 | GY_data_day-37 | 44 | accuracy | 0.925 |
| classification | nomad | Reaching B | single_day | GY_data_day-37 | GY_data_day-37 | 46 | accuracy | 0.9375 |
| classification | nomad | Reaching B | single_day | GY_data_day-37 | GY_data_day-37 | 54 | accuracy | 0.95 |
| classification | nomad | Reaching B | single_day | GY_data_day-4 | GY_data_day-4 | 44 | accuracy | 0.975 |
| classification | nomad | Reaching B | single_day | GY_data_day-4 | GY_data_day-4 | 46 | accuracy | 0.97 |
| classification | nomad | Reaching B | single_day | GY_data_day-4 | GY_data_day-4 | 54 | accuracy | 0.975 |
| classification | nomad | Reaching B | single_day | GY_data_day-5 | GY_data_day-5 | 44 | accuracy | 0.9625 |
| classification | nomad | Reaching B | single_day | GY_data_day-5 | GY_data_day-5 | 46 | accuracy | 0.958333 |
| classification | nomad | Reaching B | single_day | GY_data_day-5 | GY_data_day-5 | 54 | accuracy | 0.920833 |
| classification | nomad | Reaching B | single_day | GY_data_day-6 | GY_data_day-6 | 44 | accuracy | 0.804167 |
| classification | nomad | Reaching B | single_day | GY_data_day-6 | GY_data_day-6 | 46 | accuracy | 0.8125 |
| classification | nomad | Reaching B | single_day | GY_data_day-6 | GY_data_day-6 | 54 | accuracy | 0.804167 |
| classification | nomad | Reaching B | single_day | GY_data_day-7 | GY_data_day-7 | 44 | accuracy | 0.391667 |
| classification | nomad | Reaching B | single_day | GY_data_day-7 | GY_data_day-7 | 46 | accuracy | 0.4875 |
| classification | nomad | Reaching B | single_day | GY_data_day-7 | GY_data_day-7 | 54 | accuracy | 0.491667 |
| classification | nomad | Reaching B | single_day | GY_data_day-8 | GY_data_day-8 | 44 | accuracy | 0.95 |
| classification | nomad | Reaching B | single_day | GY_data_day-8 | GY_data_day-8 | 46 | accuracy | 0.95625 |
| classification | nomad | Reaching B | single_day | GY_data_day-8 | GY_data_day-8 | 54 | accuracy | 0.95625 |
| classification | nomad | Reaching B | single_day | GY_data_day-9 | GY_data_day-9 | 44 | accuracy | 0.9 |
| classification | nomad | Reaching B | single_day | GY_data_day-9 | GY_data_day-9 | 46 | accuracy | 0.9 |
| classification | nomad | Reaching B | single_day | GY_data_day-9 | GY_data_day-9 | 54 | accuracy | 0.825 |
| classification | mlp | Reaching B | single_day | GY_data_day-1 | GY_data_day-1 | 44 | accuracy | 0.894118 |

| | | | | | | | | |
|---|---|---|---|---|---|---|---|---|
| classification | mlp | Reaching B | single_day | GY_data_day-1 | GY_data_day-1 | 46 | accuracy | 0.823529 |
| classification | mlp | Reaching B | single_day | GY_data_day-1 | GY_data_day-1 | 54 | accuracy | 0.847059 |
| classification | mlp | Reaching B | single_day | GY_data_day-10 | GY_data_day-10 | 44 | accuracy | 0.64375 |
| classification | mlp | Reaching B | single_day | GY_data_day-10 | GY_data_day-10 | 46 | accuracy | 0.71875 |
| classification | mlp | Reaching B | single_day | GY_data_day-10 | GY_data_day-10 | 54 | accuracy | 0.70625 |
| classification | mlp | Reaching B | single_day | GY_data_day-11 | GY_data_day-11 | 44 | accuracy | 0.7875 |
| classification | mlp | Reaching B | single_day | GY_data_day-11 | GY_data_day-11 | 46 | accuracy | 0.7875 |
| classification | mlp | Reaching B | single_day | GY_data_day-11 | GY_data_day-11 | 54 | accuracy | 0.8875 |
| classification | mlp | Reaching B | single_day | GY_data_day-12 | GY_data_day-12 | 44 | accuracy | 0.953488 |
| classification | mlp | Reaching B | single_day | GY_data_day-12 | GY_data_day-12 | 46 | accuracy | 0.947674 |
| classification | mlp | Reaching B | single_day | GY_data_day-12 | GY_data_day-12 | 54 | accuracy | 0.924419 |
| classification | mlp | Reaching B | single_day | GY_data_day-13 | GY_data_day-13 | 44 | accuracy | 0.9 |
| classification | mlp | Reaching B | single_day | GY_data_day-13 | GY_data_day-13 | 46 | accuracy | 0.8875 |
| classification | mlp | Reaching B | single_day | GY_data_day-13 | GY_data_day-13 | 54 | accuracy | 0.825 |
| classification | mlp | Reaching B | single_day | GY_data_day-14 | GY_data_day-14 | 44 | accuracy | 0.833333 |
| classification | mlp | Reaching B | single_day | GY_data_day-14 | GY_data_day-14 | 46 | accuracy | 0.883333 |
| classification | mlp | Reaching B | single_day | GY_data_day-14 | GY_data_day-14 | 54 | accuracy | 0.816667 |
| classification | mlp | Reaching B | single_day | GY_data_day-15 | GY_data_day-15 | 44 | accuracy | 0.9125 |
| classification | mlp | Reaching B | single_day | GY_data_day-15 | GY_data_day-15 | 46 | accuracy | 0.96875 |
| classification | mlp | Reaching B | single_day | GY_data_day-15 | GY_data_day-15 | 54 | accuracy | 0.89375 |
| classification | mlp | Reaching B | single_day | GY_data_day-16 | GY_data_day-16 | 44 | accuracy | 0.875 |
| classification | mlp | Reaching B | single_day | GY_data_day-16 | GY_data_day-16 | 46 | accuracy | 0.95 |
| classification | mlp | Reaching B | single_day | GY_data_day-16 | GY_data_day-16 | 54 | accuracy | 0.9 |
| classification | mlp | Reaching B | single_day | GY_data_day-17 | GY_data_day-17 | 44 | accuracy | 0.908333 |
| classification | mlp | Reaching B | single_day | GY_data_day-17 | GY_data_day-17 | 46 | accuracy | 0.941667 |
| classification | mlp | Reaching B | single_day | GY_data_day-17 | GY_data_day-17 | 54 | accuracy | 0.925 |
| classification | mlp | Reaching B | single_day | GY_data_day-18 | GY_data_day-18 | 44 | accuracy | 0.8625 |
| classification | mlp | Reaching B | single_day | GY_data_day-18 | GY_data_day-18 | 46 | accuracy | 0.9625 |
| classification | mlp | Reaching B | single_day | GY_data_day-18 | GY_data_day-18 | 54 | accuracy | 0.9125 |
| classification | mlp | Reaching B | single_day | GY_data_day-19 | GY_data_day-19 | 44 | accuracy | 0.7375 |
| classification | mlp | Reaching B | single_day | GY_data_day-19 | GY_data_day-19 | 46 | accuracy | 0.8875 |
| classification | mlp | Reaching B | single_day | GY_data_day-19 | GY_data_day-19 | 54 | accuracy | 0.825 |
| classification | mlp | Reaching B | single_day | GY_data_day-2 | GY_data_day-2 | 44 | accuracy | 0.885 |
| classification | mlp | Reaching B | single_day | GY_data_day-2 | GY_data_day-2 | 46 | accuracy | 0.95 |
| classification | mlp | Reaching B | single_day | GY_data_day-2 | GY_data_day-2 | 54 | accuracy | 0.88 |
| classification | mlp | Reaching B | single_day | GY_data_day-20 | GY_data_day-20 | 44 | accuracy | 0.875 |
| classification | mlp | Reaching B | single_day | GY_data_day-20 | GY_data_day-20 | 46 | accuracy | 0.925 |
| classification | mlp | Reaching B | single_day | GY_data_day-20 | GY_data_day-20 | 54 | accuracy | 0.825 |
| classification | mlp | Reaching B | single_day | GY_data_day-21 | GY_data_day-21 | 44 | accuracy | 0.675 |
| classification | mlp | Reaching B | single_day | GY_data_day-21 | GY_data_day-21 | 46 | accuracy | 0.666667 |
| classification | mlp | Reaching B | single_day | GY_data_day-21 | GY_data_day-21 | 54 | accuracy | 0.6 |
| classification | mlp | Reaching B | single_day | GY_data_day-22 | GY_data_day-22 | 44 | accuracy | 0.953571 |
| classification | mlp | Reaching B | single_day | GY_data_day-22 | GY_data_day-22 | 46 | accuracy | 0.935714 |
| classification | mlp | Reaching B | single_day | GY_data_day-22 | GY_data_day-22 | 54 | accuracy | 0.975 |
| classification | mlp | Reaching B | single_day | GY_data_day-23 | GY_data_day-23 | 44 | accuracy | 0.965 |
| classification | mlp | Reaching B | single_day | GY_data_day-23 | GY_data_day-23 | 46 | accuracy | 0.955 |
| classification | mlp | Reaching B | single_day | GY_data_day-23 | GY_data_day-23 | 54 | accuracy | 0.97 |
| classification | mlp | Reaching B | single_day | GY_data_day-24 | GY_data_day-24 | 44 | accuracy | 0.825 |
| classification | mlp | Reaching B | single_day | GY_data_day-24 | GY_data_day-24 | 46 | accuracy | 0.7875 |
| classification | mlp | Reaching B | single_day | GY_data_day-24 | GY_data_day-24 | 54 | accuracy | 0.7875 |
| classification | mlp | Reaching B | single_day | GY_data_day-25 | GY_data_day-25 | 44 | accuracy | 0.875 |
| classification | mlp | Reaching B | single_day | GY_data_day-25 | GY_data_day-25 | 46 | accuracy | 0.875 |
| classification | mlp | Reaching B | single_day | GY_data_day-25 | GY_data_day-25 | 54 | accuracy | 0.8125 |
| classification | mlp | Reaching B | single_day | GY_data_day-26 | GY_data_day-26 | 44 | accuracy | 0.85 |
| classification | mlp | Reaching B | single_day | GY_data_day-26 | GY_data_day-26 | 46 | accuracy | 0.9 |
| classification | mlp | Reaching B | single_day | GY_data_day-26 | GY_data_day-26 | 54 | accuracy | 0.7625 |
| classification | mlp | Reaching B | single_day | GY_data_day-27 | GY_data_day-27 | 44 | accuracy | 0.8875 |
| classification | mlp | Reaching B | single_day | GY_data_day-27 | GY_data_day-27 | 46 | accuracy | 0.9375 |
| classification | mlp | Reaching B | single_day | GY_data_day-27 | GY_data_day-27 | 54 | accuracy | 0.9375 |
| classification | mlp | Reaching B | single_day | GY_data_day-28 | GY_data_day-28 | 44 | accuracy | 0.925 |
| classification | mlp | Reaching B | single_day | GY_data_day-28 | GY_data_day-28 | 46 | accuracy | 0.9125 |
| classification | mlp | Reaching B | single_day | GY_data_day-28 | GY_data_day-28 | 54 | accuracy | 0.9 |
| classification | mlp | Reaching B | single_day | GY_data_day-29 | GY_data_day-29 | 44 | accuracy | 0.8375 |

| | | | | | | | | |
|---|---|---|---|---|---|---|---|---|
| classification | mlp | Reaching B | single_day | GY_data_day-29 | GY_data_day-29 | 46 | accuracy | 0.8875 |
| classification | mlp | Reaching B | single_day | GY_data_day-29 | GY_data_day-29 | 54 | accuracy | 0.8875 |
| classification | mlp | Reaching B | single_day | GY_data_day-3 | GY_data_day-3 | 44 | accuracy | 0.791667 |
| classification | mlp | Reaching B | single_day | GY_data_day-3 | GY_data_day-3 | 46 | accuracy | 0.775 |
| classification | mlp | Reaching B | single_day | GY_data_day-3 | GY_data_day-3 | 54 | accuracy | 0.766667 |
| classification | mlp | Reaching B | single_day | GY_data_day-30 | GY_data_day-30 | 44 | accuracy | 0.9 |
| classification | mlp | Reaching B | single_day | GY_data_day-30 | GY_data_day-30 | 46 | accuracy | 0.8875 |
| classification | mlp | Reaching B | single_day | GY_data_day-30 | GY_data_day-30 | 54 | accuracy | 0.8875 |
| classification | mlp | Reaching B | single_day | GY_data_day-31 | GY_data_day-31 | 44 | accuracy | 0.825 |
| classification | mlp | Reaching B | single_day | GY_data_day-31 | GY_data_day-31 | 46 | accuracy | 0.775 |
| classification | mlp | Reaching B | single_day | GY_data_day-31 | GY_data_day-31 | 54 | accuracy | 0.725 |
| classification | mlp | Reaching B | single_day | GY_data_day-32 | GY_data_day-32 | 44 | accuracy | 0.825 |
| classification | mlp | Reaching B | single_day | GY_data_day-32 | GY_data_day-32 | 46 | accuracy | 0.925 |
| classification | mlp | Reaching B | single_day | GY_data_day-32 | GY_data_day-32 | 54 | accuracy | 0.7875 |
| classification | mlp | Reaching B | single_day | GY_data_day-33 | GY_data_day-33 | 44 | accuracy | 0.8 |
| classification | mlp | Reaching B | single_day | GY_data_day-33 | GY_data_day-33 | 46 | accuracy | 0.841667 |
| classification | mlp | Reaching B | single_day | GY_data_day-33 | GY_data_day-33 | 54 | accuracy | 0.825 |
| classification | mlp | Reaching B | single_day | GY_data_day-34 | GY_data_day-34 | 44 | accuracy | 0.86875 |
| classification | mlp | Reaching B | single_day | GY_data_day-34 | GY_data_day-34 | 46 | accuracy | 0.91875 |
| classification | mlp | Reaching B | single_day | GY_data_day-34 | GY_data_day-34 | 54 | accuracy | 0.9 |
| classification | mlp | Reaching B | single_day | GY_data_day-35 | GY_data_day-35 | 44 | accuracy | 0.84375 |
| classification | mlp | Reaching B | single_day | GY_data_day-35 | GY_data_day-35 | 46 | accuracy | 0.9125 |
| classification | mlp | Reaching B | single_day | GY_data_day-35 | GY_data_day-35 | 54 | accuracy | 0.7875 |
| classification | mlp | Reaching B | single_day | GY_data_day-36 | GY_data_day-36 | 44 | accuracy | 0.91875 |
| classification | mlp | Reaching B | single_day | GY_data_day-36 | GY_data_day-36 | 46 | accuracy | 0.8875 |
| classification | mlp | Reaching B | single_day | GY_data_day-36 | GY_data_day-36 | 54 | accuracy | 0.85 |
| classification | mlp | Reaching B | single_day | GY_data_day-37 | GY_data_day-37 | 44 | accuracy | 0.925 |
| classification | mlp | Reaching B | single_day | GY_data_day-37 | GY_data_day-37 | 46 | accuracy | 0.875 |
| classification | mlp | Reaching B | single_day | GY_data_day-37 | GY_data_day-37 | 54 | accuracy | 0.875 |
| classification | mlp | Reaching B | single_day | GY_data_day-4 | GY_data_day-4 | 44 | accuracy | 0.945 |
| classification | mlp | Reaching B | single_day | GY_data_day-4 | GY_data_day-4 | 46 | accuracy | 0.97 |
| classification | mlp | Reaching B | single_day | GY_data_day-4 | GY_data_day-4 | 54 | accuracy | 0.925 |
| classification | mlp | Reaching B | single_day | GY_data_day-5 | GY_data_day-5 | 44 | accuracy | 0.891667 |
| classification | mlp | Reaching B | single_day | GY_data_day-5 | GY_data_day-5 | 46 | accuracy | 0.8375 |
| classification | mlp | Reaching B | single_day | GY_data_day-5 | GY_data_day-5 | 54 | accuracy | 0.879167 |
| classification | mlp | Reaching B | single_day | GY_data_day-6 | GY_data_day-6 | 44 | accuracy | 0.75 |
| classification | mlp | Reaching B | single_day | GY_data_day-6 | GY_data_day-6 | 46 | accuracy | 0.779167 |
| classification | mlp | Reaching B | single_day | GY_data_day-6 | GY_data_day-6 | 54 | accuracy | 0.758333 |
| classification | mlp | Reaching B | single_day | GY_data_day-7 | GY_data_day-7 | 44 | accuracy | 0.441667 |
| classification | mlp | Reaching B | single_day | GY_data_day-7 | GY_data_day-7 | 46 | accuracy | 0.520833 |
| classification | mlp | Reaching B | single_day | GY_data_day-7 | GY_data_day-7 | 54 | accuracy | 0.4875 |
| classification | mlp | Reaching B | single_day | GY_data_day-8 | GY_data_day-8 | 44 | accuracy | 0.7625 |
| classification | mlp | Reaching B | single_day | GY_data_day-8 | GY_data_day-8 | 46 | accuracy | 0.78125 |
| classification | mlp | Reaching B | single_day | GY_data_day-8 | GY_data_day-8 | 54 | accuracy | 0.84375 |
| classification | mlp | Reaching B | single_day | GY_data_day-9 | GY_data_day-9 | 44 | accuracy | 0.8875 |
| classification | mlp | Reaching B | single_day | GY_data_day-9 | GY_data_day-9 | 46 | accuracy | 0.8 |
| classification | mlp | Reaching B | single_day | GY_data_day-9 | GY_data_day-9 | 54 | accuracy | 0.825 |
| classification | convpi_vae | Reaching B | single_day | GY_data_day-1 | GY_data_day-1 | 31 | accuracy | 0.729412 |
| classification | convpi_vae | Reaching B | single_day | GY_data_day-1 | GY_data_day-1 | 44 | accuracy | 0.994118 |
| classification | convpi_vae | Reaching B | single_day | GY_data_day-1 | GY_data_day-1 | 46 | accuracy | 0.994118 |
| classification | convpi_vae | Reaching B | single_day | GY_data_day-1 | GY_data_day-1 | 54 | accuracy | 0.8 |
| classification | convpi_vae | Reaching B | single_day | GY_data_day-1 | GY_data_day-1 | 69 | accuracy | 1 |
| classification | convpi_vae | Reaching B | single_day | GY_data_day-10 | GY_data_day-10 | 31 | accuracy | 0.6625 |
| classification | convpi_vae | Reaching B | single_day | GY_data_day-10 | GY_data_day-10 | 44 | accuracy | 0.96875 |
| classification | convpi_vae | Reaching B | single_day | GY_data_day-10 | GY_data_day-10 | 46 | accuracy | 0.94375 |
| classification | convpi_vae | Reaching B | single_day | GY_data_day-10 | GY_data_day-10 | 54 | accuracy | 0.91875 |
| classification | convpi_vae | Reaching B | single_day | GY_data_day-10 | GY_data_day-10 | 69 | accuracy | 1 |
| classification | convpi_vae | Reaching B | single_day | GY_data_day-11 | GY_data_day-11 | 31 | accuracy | 0.8 |
| classification | convpi_vae | Reaching B | single_day | GY_data_day-11 | GY_data_day-11 | 44 | accuracy | 0.975 |
| classification | convpi_vae | Reaching B | single_day | GY_data_day-11 | GY_data_day-11 | 46 | accuracy | 0.95 |
| classification | convpi_vae | Reaching B | single_day | GY_data_day-11 | GY_data_day-11 | 54 | accuracy | 0.7375 |
| classification | convpi_vae | Reaching B | single_day | GY_data_day-11 | GY_data_day-11 | 69 | accuracy | 0.95 |
| classification | convpi_vae | Reaching B | single_day | GY_data_day-12 | GY_data_day-12 | 31 | accuracy | 0.994186 |

| | | | | | | | | |
|---|---|---|---|---|---|---|---|---|
| classification | convpi_vae | Reaching B | single_day | GY_data_day-12 | GY_data_day-12 | 44 | accuracy | 1 |
| classification | convpi_vae | Reaching B | single_day | GY_data_day-12 | GY_data_day-12 | 46 | accuracy | 0.936047 |
| classification | convpi_vae | Reaching B | single_day | GY_data_day-12 | GY_data_day-12 | 54 | accuracy | 0.674419 |
| classification | convpi_vae | Reaching B | single_day | GY_data_day-12 | GY_data_day-12 | 69 | accuracy | 0.918605 |
| classification | convpi_vae | Reaching B | single_day | GY_data_day-13 | GY_data_day-13 | 31 | accuracy | 0.5625 |
| classification | convpi_vae | Reaching B | single_day | GY_data_day-13 | GY_data_day-13 | 44 | accuracy | 0.9125 |
| classification | convpi_vae | Reaching B | single_day | GY_data_day-13 | GY_data_day-13 | 46 | accuracy | 0.7125 |
| classification | convpi_vae | Reaching B | single_day | GY_data_day-13 | GY_data_day-13 | 54 | accuracy | 0.65 |
| classification | convpi_vae | Reaching B | single_day | GY_data_day-13 | GY_data_day-13 | 69 | accuracy | 0.45 |
| classification | convpi_vae | Reaching B | single_day | GY_data_day-14 | GY_data_day-14 | 31 | accuracy | 0.491667 |
| classification | convpi_vae | Reaching B | single_day | GY_data_day-14 | GY_data_day-14 | 44 | accuracy | 0.95 |
| classification | convpi_vae | Reaching B | single_day | GY_data_day-14 | GY_data_day-14 | 46 | accuracy | 0.65 |
| classification | convpi_vae | Reaching B | single_day | GY_data_day-14 | GY_data_day-14 | 54 | accuracy | 0.766667 |
| classification | convpi_vae | Reaching B | single_day | GY_data_day-14 | GY_data_day-14 | 69 | accuracy | 0.725 |
| classification | convpi_vae | Reaching B | single_day | GY_data_day-15 | GY_data_day-15 | 31 | accuracy | 0.9625 |
| classification | convpi_vae | Reaching B | single_day | GY_data_day-15 | GY_data_day-15 | 44 | accuracy | 0.95625 |
| classification | convpi_vae | Reaching B | single_day | GY_data_day-15 | GY_data_day-15 | 46 | accuracy | 0.9625 |
| classification | convpi_vae | Reaching B | single_day | GY_data_day-15 | GY_data_day-15 | 54 | accuracy | 0.975 |
| classification | convpi_vae | Reaching B | single_day | GY_data_day-15 | GY_data_day-15 | 69 | accuracy | 0.96875 |
| classification | convpi_vae | Reaching B | single_day | GY_data_day-16 | GY_data_day-16 | 31 | accuracy | 0.65 |
| classification | convpi_vae | Reaching B | single_day | GY_data_day-16 | GY_data_day-16 | 44 | accuracy | 1 |
| classification | convpi_vae | Reaching B | single_day | GY_data_day-16 | GY_data_day-16 | 46 | accuracy | 0.9 |
| classification | convpi_vae | Reaching B | single_day | GY_data_day-16 | GY_data_day-16 | 54 | accuracy | 0.975 |
| classification | convpi_vae | Reaching B | single_day | GY_data_day-16 | GY_data_day-16 | 69 | accuracy | 1 |
| classification | convpi_vae | Reaching B | single_day | GY_data_day-17 | GY_data_day-17 | 31 | accuracy | 0.808333 |
| classification | convpi_vae | Reaching B | single_day | GY_data_day-17 | GY_data_day-17 | 44 | accuracy | 0.991667 |
| classification | convpi_vae | Reaching B | single_day | GY_data_day-17 | GY_data_day-17 | 46 | accuracy | 1 |
| classification | convpi_vae | Reaching B | single_day | GY_data_day-17 | GY_data_day-17 | 54 | accuracy | 0.941667 |
| classification | convpi_vae | Reaching B | single_day | GY_data_day-17 | GY_data_day-17 | 69 | accuracy | 1 |
| classification | convpi_vae | Reaching B | single_day | GY_data_day-18 | GY_data_day-18 | 31 | accuracy | 0.7625 |
| classification | convpi_vae | Reaching B | single_day | GY_data_day-18 | GY_data_day-18 | 44 | accuracy | 0.975 |
| classification | convpi_vae | Reaching B | single_day | GY_data_day-18 | GY_data_day-18 | 46 | accuracy | 0.85 |
| classification | convpi_vae | Reaching B | single_day | GY_data_day-18 | GY_data_day-18 | 54 | accuracy | 0.925 |
| classification | convpi_vae | Reaching B | single_day | GY_data_day-18 | GY_data_day-18 | 69 | accuracy | 0.9 |
| classification | convpi_vae | Reaching B | single_day | GY_data_day-19 | GY_data_day-19 | 31 | accuracy | 0.8375 |
| classification | convpi_vae | Reaching B | single_day | GY_data_day-19 | GY_data_day-19 | 44 | accuracy | 0.95 |
| classification | convpi_vae | Reaching B | single_day | GY_data_day-19 | GY_data_day-19 | 46 | accuracy | 0.9125 |
| classification | convpi_vae | Reaching B | single_day | GY_data_day-19 | GY_data_day-19 | 54 | accuracy | 0.775 |
| classification | convpi_vae | Reaching B | single_day | GY_data_day-19 | GY_data_day-19 | 69 | accuracy | 0.9125 |
| classification | convpi_vae | Reaching B | single_day | GY_data_day-2 | GY_data_day-2 | 31 | accuracy | 0.82 |
| classification | convpi_vae | Reaching B | single_day | GY_data_day-2 | GY_data_day-2 | 44 | accuracy | 0.985 |
| classification | convpi_vae | Reaching B | single_day | GY_data_day-2 | GY_data_day-2 | 46 | accuracy | 0.98 |
| classification | convpi_vae | Reaching B | single_day | GY_data_day-2 | GY_data_day-2 | 54 | accuracy | 0.79 |
| classification | convpi_vae | Reaching B | single_day | GY_data_day-2 | GY_data_day-2 | 69 | accuracy | 1 |
| classification | convpi_vae | Reaching B | single_day | GY_data_day-20 | GY_data_day-20 | 31 | accuracy | 0.975 |
| classification | convpi_vae | Reaching B | single_day | GY_data_day-20 | GY_data_day-20 | 44 | accuracy | 1 |
| classification | convpi_vae | Reaching B | single_day | GY_data_day-20 | GY_data_day-20 | 46 | accuracy | 0.975 |
| classification | convpi_vae | Reaching B | single_day | GY_data_day-20 | GY_data_day-20 | 54 | accuracy | 1 |
| classification | convpi_vae | Reaching B | single_day | GY_data_day-20 | GY_data_day-20 | 69 | accuracy | 0.975 |
| classification | convpi_vae | Reaching B | single_day | GY_data_day-21 | GY_data_day-21 | 31 | accuracy | 0.5 |
| classification | convpi_vae | Reaching B | single_day | GY_data_day-21 | GY_data_day-21 | 44 | accuracy | 0.941667 |
| classification | convpi_vae | Reaching B | single_day | GY_data_day-21 | GY_data_day-21 | 46 | accuracy | 0.966667 |
| classification | convpi_vae | Reaching B | single_day | GY_data_day-21 | GY_data_day-21 | 54 | accuracy | 0.983333 |
| classification | convpi_vae | Reaching B | single_day | GY_data_day-21 | GY_data_day-21 | 69 | accuracy | 0.55 |
| classification | convpi_vae | Reaching B | single_day | GY_data_day-22 | GY_data_day-22 | 31 | accuracy | 1 |
| classification | convpi_vae | Reaching B | single_day | GY_data_day-22 | GY_data_day-22 | 44 | accuracy | 1 |
| classification | convpi_vae | Reaching B | single_day | GY_data_day-22 | GY_data_day-22 | 46 | accuracy | 1 |
| classification | convpi_vae | Reaching B | single_day | GY_data_day-22 | GY_data_day-22 | 54 | accuracy | 1 |
| classification | convpi_vae | Reaching B | single_day | GY_data_day-22 | GY_data_day-22 | 69 | accuracy | 1 |
| classification | convpi_vae | Reaching B | single_day | GY_data_day-23 | GY_data_day-23 | 31 | accuracy | 0.98 |
| classification | convpi_vae | Reaching B | single_day | GY_data_day-23 | GY_data_day-23 | 44 | accuracy | 0.995 |
| classification | convpi_vae | Reaching B | single_day | GY_data_day-23 | GY_data_day-23 | 46 | accuracy | 1 |
| classification | convpi_vae | Reaching B | single_day | GY_data_day-23 | GY_data_day-23 | 54 | accuracy | 1 |

| | | | | | | | | |
|---|---|---|---|---|---|---|---|---|
| classification | convpi_vae | Reaching B | single_day | GY_data_day-23 | GY_data_day-23 | 69 | accuracy | 0.995 |
| classification | convpi_vae | Reaching B | single_day | GY_data_day-24 | GY_data_day-24 | 31 | accuracy | 0.6375 |
| classification | convpi_vae | Reaching B | single_day | GY_data_day-24 | GY_data_day-24 | 44 | accuracy | 0.9875 |
| classification | convpi_vae | Reaching B | single_day | GY_data_day-24 | GY_data_day-24 | 46 | accuracy | 0.875 |
| classification | convpi_vae | Reaching B | single_day | GY_data_day-24 | GY_data_day-24 | 54 | accuracy | 0.7 |
| classification | convpi_vae | Reaching B | single_day | GY_data_day-24 | GY_data_day-24 | 69 | accuracy | 0.7875 |
| classification | convpi_vae | Reaching B | single_day | GY_data_day-25 | GY_data_day-25 | 31 | accuracy | 0.8125 |
| classification | convpi_vae | Reaching B | single_day | GY_data_day-25 | GY_data_day-25 | 44 | accuracy | 1 |
| classification | convpi_vae | Reaching B | single_day | GY_data_day-25 | GY_data_day-25 | 46 | accuracy | 0.9125 |
| classification | convpi_vae | Reaching B | single_day | GY_data_day-25 | GY_data_day-25 | 54 | accuracy | 0.95 |
| classification | convpi_vae | Reaching B | single_day | GY_data_day-25 | GY_data_day-25 | 69 | accuracy | 0.9375 |
| classification | convpi_vae | Reaching B | single_day | GY_data_day-26 | GY_data_day-26 | 31 | accuracy | 0.925 |
| classification | convpi_vae | Reaching B | single_day | GY_data_day-26 | GY_data_day-26 | 44 | accuracy | 0.975 |
| classification | convpi_vae | Reaching B | single_day | GY_data_day-26 | GY_data_day-26 | 46 | accuracy | 0.9375 |
| classification | convpi_vae | Reaching B | single_day | GY_data_day-26 | GY_data_day-26 | 54 | accuracy | 0.9625 |
| classification | convpi_vae | Reaching B | single_day | GY_data_day-26 | GY_data_day-26 | 69 | accuracy | 0.9625 |
| classification | convpi_vae | Reaching B | single_day | GY_data_day-27 | GY_data_day-27 | 31 | accuracy | 0.875 |
| classification | convpi_vae | Reaching B | single_day | GY_data_day-27 | GY_data_day-27 | 44 | accuracy | 0.8625 |
| classification | convpi_vae | Reaching B | single_day | GY_data_day-27 | GY_data_day-27 | 46 | accuracy | 0.8125 |
| classification | convpi_vae | Reaching B | single_day | GY_data_day-27 | GY_data_day-27 | 54 | accuracy | 0.8875 |
| classification | convpi_vae | Reaching B | single_day | GY_data_day-27 | GY_data_day-27 | 69 | accuracy | 0.975 |
| classification | convpi_vae | Reaching B | single_day | GY_data_day-28 | GY_data_day-28 | 31 | accuracy | 0.85 |
| classification | convpi_vae | Reaching B | single_day | GY_data_day-28 | GY_data_day-28 | 44 | accuracy | 0.975 |
| classification | convpi_vae | Reaching B | single_day | GY_data_day-28 | GY_data_day-28 | 46 | accuracy | 0.7875 |
| classification | convpi_vae | Reaching B | single_day | GY_data_day-28 | GY_data_day-28 | 54 | accuracy | 0.9125 |
| classification | convpi_vae | Reaching B | single_day | GY_data_day-28 | GY_data_day-28 | 69 | accuracy | 0.7125 |
| classification | convpi_vae | Reaching B | single_day | GY_data_day-29 | GY_data_day-29 | 31 | accuracy | 0.8625 |
| classification | convpi_vae | Reaching B | single_day | GY_data_day-29 | GY_data_day-29 | 44 | accuracy | 1 |
| classification | convpi_vae | Reaching B | single_day | GY_data_day-29 | GY_data_day-29 | 46 | accuracy | 1 |
| classification | convpi_vae | Reaching B | single_day | GY_data_day-29 | GY_data_day-29 | 54 | accuracy | 0.975 |
| classification | convpi_vae | Reaching B | single_day | GY_data_day-29 | GY_data_day-29 | 69 | accuracy | 0.825 |
| classification | convpi_vae | Reaching B | single_day | GY_data_day-3 | GY_data_day-3 | 31 | accuracy | 0.616667 |
| classification | convpi_vae | Reaching B | single_day | GY_data_day-3 | GY_data_day-3 | 44 | accuracy | 0.966667 |
| classification | convpi_vae | Reaching B | single_day | GY_data_day-3 | GY_data_day-3 | 46 | accuracy | 0.941667 |
| classification | convpi_vae | Reaching B | single_day | GY_data_day-3 | GY_data_day-3 | 54 | accuracy | 0.741667 |
| classification | convpi_vae | Reaching B | single_day | GY_data_day-3 | GY_data_day-3 | 69 | accuracy | 0.916667 |
| classification | convpi_vae | Reaching B | single_day | GY_data_day-30 | GY_data_day-30 | 31 | accuracy | 0.95 |
| classification | convpi_vae | Reaching B | single_day | GY_data_day-30 | GY_data_day-30 | 44 | accuracy | 1 |
| classification | convpi_vae | Reaching B | single_day | GY_data_day-30 | GY_data_day-30 | 46 | accuracy | 1 |
| classification | convpi_vae | Reaching B | single_day | GY_data_day-30 | GY_data_day-30 | 54 | accuracy | 0.9875 |
| classification | convpi_vae | Reaching B | single_day | GY_data_day-30 | GY_data_day-30 | 69 | accuracy | 0.9875 |
| classification | convpi_vae | Reaching B | single_day | GY_data_day-31 | GY_data_day-31 | 31 | accuracy | 0.975 |
| classification | convpi_vae | Reaching B | single_day | GY_data_day-31 | GY_data_day-31 | 44 | accuracy | 1 |
| classification | convpi_vae | Reaching B | single_day | GY_data_day-31 | GY_data_day-31 | 46 | accuracy | 0.85 |
| classification | convpi_vae | Reaching B | single_day | GY_data_day-31 | GY_data_day-31 | 54 | accuracy | 0.95 |
| classification | convpi_vae | Reaching B | single_day | GY_data_day-31 | GY_data_day-31 | 69 | accuracy | 0.95 |
| classification | convpi_vae | Reaching B | single_day | GY_data_day-32 | GY_data_day-32 | 31 | accuracy | 0.9375 |
| classification | convpi_vae | Reaching B | single_day | GY_data_day-32 | GY_data_day-32 | 44 | accuracy | 1 |
| classification | convpi_vae | Reaching B | single_day | GY_data_day-32 | GY_data_day-32 | 46 | accuracy | 0.9875 |
| classification | convpi_vae | Reaching B | single_day | GY_data_day-32 | GY_data_day-32 | 54 | accuracy | 0.975 |
| classification | convpi_vae | Reaching B | single_day | GY_data_day-32 | GY_data_day-32 | 69 | accuracy | 0.975 |
| classification | convpi_vae | Reaching B | single_day | GY_data_day-33 | GY_data_day-33 | 31 | accuracy | 0.975 |
| classification | convpi_vae | Reaching B | single_day | GY_data_day-33 | GY_data_day-33 | 44 | accuracy | 1 |
| classification | convpi_vae | Reaching B | single_day | GY_data_day-33 | GY_data_day-33 | 46 | accuracy | 0.983333 |
| classification | convpi_vae | Reaching B | single_day | GY_data_day-33 | GY_data_day-33 | 54 | accuracy | 0.991667 |
| classification | convpi_vae | Reaching B | single_day | GY_data_day-33 | GY_data_day-33 | 69 | accuracy | 0.933333 |
| classification | convpi_vae | Reaching B | single_day | GY_data_day-34 | GY_data_day-34 | 31 | accuracy | 0.975 |
| classification | convpi_vae | Reaching B | single_day | GY_data_day-34 | GY_data_day-34 | 44 | accuracy | 1 |
| classification | convpi_vae | Reaching B | single_day | GY_data_day-34 | GY_data_day-34 | 46 | accuracy | 1 |
| classification | convpi_vae | Reaching B | single_day | GY_data_day-34 | GY_data_day-34 | 54 | accuracy | 1 |
| classification | convpi_vae | Reaching B | single_day | GY_data_day-34 | GY_data_day-34 | 69 | accuracy | 0.99375 |
| classification | convpi_vae | Reaching B | single_day | GY_data_day-35 | GY_data_day-35 | 31 | accuracy | 1 |
| classification | convpi_vae | Reaching B | single_day | GY_data_day-35 | GY_data_day-35 | 44 | accuracy | 1 |

| | | | | | | | | |
|---|---|---|---|---|---|---|---|---|
| classification | convpi_vae | Reaching B | single_day | GY_data_day-35 | GY_data_day-35 | 46 | accuracy | 1 |
| classification | convpi_vae | Reaching B | single_day | GY_data_day-35 | GY_data_day-35 | 54 | accuracy | 1 |
| classification | convpi_vae | Reaching B | single_day | GY_data_day-35 | GY_data_day-35 | 69 | accuracy | 1 |
| classification | convpi_vae | Reaching B | single_day | GY_data_day-36 | GY_data_day-36 | 31 | accuracy | 1 |
| classification | convpi_vae | Reaching B | single_day | GY_data_day-36 | GY_data_day-36 | 44 | accuracy | 0.9875 |
| classification | convpi_vae | Reaching B | single_day | GY_data_day-36 | GY_data_day-36 | 46 | accuracy | 0.99375 |
| classification | convpi_vae | Reaching B | single_day | GY_data_day-36 | GY_data_day-36 | 54 | accuracy | 1 |
| classification | convpi_vae | Reaching B | single_day | GY_data_day-36 | GY_data_day-36 | 69 | accuracy | 1 |
| classification | convpi_vae | Reaching B | single_day | GY_data_day-37 | GY_data_day-37 | 31 | accuracy | 0.6375 |
| classification | convpi_vae | Reaching B | single_day | GY_data_day-37 | GY_data_day-37 | 44 | accuracy | 0.9125 |
| classification | convpi_vae | Reaching B | single_day | GY_data_day-37 | GY_data_day-37 | 46 | accuracy | 0.7375 |
| classification | convpi_vae | Reaching B | single_day | GY_data_day-37 | GY_data_day-37 | 54 | accuracy | 0.6875 |
| classification | convpi_vae | Reaching B | single_day | GY_data_day-37 | GY_data_day-37 | 69 | accuracy | 0.975 |
| classification | convpi_vae | Reaching B | single_day | GY_data_day-4 | GY_data_day-4 | 31 | accuracy | 0.98 |
| classification | convpi_vae | Reaching B | single_day | GY_data_day-4 | GY_data_day-4 | 44 | accuracy | 1 |
| classification | convpi_vae | Reaching B | single_day | GY_data_day-4 | GY_data_day-4 | 46 | accuracy | 0.965 |
| classification | convpi_vae | Reaching B | single_day | GY_data_day-4 | GY_data_day-4 | 54 | accuracy | 1 |
| classification | convpi_vae | Reaching B | single_day | GY_data_day-4 | GY_data_day-4 | 69 | accuracy | 0.995 |
| classification | convpi_vae | Reaching B | single_day | GY_data_day-5 | GY_data_day-5 | 31 | accuracy | 0.7375 |
| classification | convpi_vae | Reaching B | single_day | GY_data_day-5 | GY_data_day-5 | 44 | accuracy | 0.95 |
| classification | convpi_vae | Reaching B | single_day | GY_data_day-5 | GY_data_day-5 | 46 | accuracy | 0.991667 |
| classification | convpi_vae | Reaching B | single_day | GY_data_day-5 | GY_data_day-5 | 54 | accuracy | 1 |
| classification | convpi_vae | Reaching B | single_day | GY_data_day-5 | GY_data_day-5 | 69 | accuracy | 1 |
| classification | convpi_vae | Reaching B | single_day | GY_data_day-6 | GY_data_day-6 | 31 | accuracy | 0.7875 |
| classification | convpi_vae | Reaching B | single_day | GY_data_day-6 | GY_data_day-6 | 44 | accuracy | 1 |
| classification | convpi_vae | Reaching B | single_day | GY_data_day-6 | GY_data_day-6 | 46 | accuracy | 0.9875 |
| classification | convpi_vae | Reaching B | single_day | GY_data_day-6 | GY_data_day-6 | 54 | accuracy | 0.791667 |
| classification | convpi_vae | Reaching B | single_day | GY_data_day-6 | GY_data_day-6 | 69 | accuracy | 1 |
| classification | convpi_vae | Reaching B | single_day | GY_data_day-7 | GY_data_day-7 | 31 | accuracy | 1 |
| classification | convpi_vae | Reaching B | single_day | GY_data_day-7 | GY_data_day-7 | 44 | accuracy | 1 |
| classification | convpi_vae | Reaching B | single_day | GY_data_day-7 | GY_data_day-7 | 46 | accuracy | 0.95 |
| classification | convpi_vae | Reaching B | single_day | GY_data_day-7 | GY_data_day-7 | 54 | accuracy | 1 |
| classification | convpi_vae | Reaching B | single_day | GY_data_day-7 | GY_data_day-7 | 69 | accuracy | 0.995833 |
| classification | convpi_vae | Reaching B | single_day | GY_data_day-8 | GY_data_day-8 | 31 | accuracy | 0.875 |
| classification | convpi_vae | Reaching B | single_day | GY_data_day-8 | GY_data_day-8 | 44 | accuracy | 0.90625 |
| classification | convpi_vae | Reaching B | single_day | GY_data_day-8 | GY_data_day-8 | 46 | accuracy | 0.89375 |
| classification | convpi_vae | Reaching B | single_day | GY_data_day-8 | GY_data_day-8 | 54 | accuracy | 0.8875 |
| classification | convpi_vae | Reaching B | single_day | GY_data_day-8 | GY_data_day-8 | 69 | accuracy | 0.91875 |
| classification | convpi_vae | Reaching B | single_day | GY_data_day-9 | GY_data_day-9 | 31 | accuracy | 0.8 |
| classification | convpi_vae | Reaching B | single_day | GY_data_day-9 | GY_data_day-9 | 44 | accuracy | 1 |
| classification | convpi_vae | Reaching B | single_day | GY_data_day-9 | GY_data_day-9 | 46 | accuracy | 0.925 |
| classification | convpi_vae | Reaching B | single_day | GY_data_day-9 | GY_data_day-9 | 54 | accuracy | 0.8375 |
| classification | convpi_vae | Reaching B | single_day | GY_data_day-9 | GY_data_day-9 | 69 | accuracy | 0.8625 |
| classification | lstm | Reaching B | cross_day | GY_data_day-1__to__GY_data_day-26 | GY_data_day-30 | 44 | accuracy | 0.5325 |
| classification | lstm | Reaching B | cross_day | GY_data_day-1__to__GY_data_day-26 | GY_data_day-30 | 46 | accuracy | 0.4725 |
| classification | lstm | Reaching B | cross_day | GY_data_day-1__to__GY_data_day-26 | GY_data_day-30 | 54 | accuracy | 0.5525 |
| classification | lstm | Reaching B | cross_day | GY_data_day-1__to__GY_data_day-26 | GY_data_day-31 | 44 | accuracy | 0.67 |
| classification | lstm | Reaching B | cross_day | GY_data_day-1__to__GY_data_day-26 | GY_data_day-31 | 46 | accuracy | 0.56 |
| classification | lstm | Reaching B | cross_day | GY_data_day-1__to__GY_data_day-26 | GY_data_day-31 | 54 | accuracy | 0.66 |
| classification | lstm | Reaching B | cross_day | GY_data_day-1__to__GY_data_day-26 | GY_data_day-32 | 44 | accuracy | 0.6175 |
| classification | lstm | Reaching B | cross_day | GY_data_day-1__to__GY_data_day-26 | GY_data_day-32 | 46 | accuracy | 0.7325 |
| classification | lstm | Reaching B | cross_day | GY_data_day-1__to__GY_data_day-26 | GY_data_day-32 | 54 | accuracy | 0.645 |
| classification | lstm | Reaching B | cross_day | GY_data_day-1__to__GY_data_day-26 | GY_data_day-33 | 44 | accuracy | 0.648333 |
| classification | lstm | Reaching B | cross_day | GY_data_day-1__to__GY_data_day-26 | GY_data_day-33 | 46 | accuracy | 0.541667 |
| classification | lstm | Reaching B | cross_day | GY_data_day-1__to__GY_data_day-26 | GY_data_day-33 | 54 | accuracy | 0.67 |
| classification | lstm | Reaching B | cross_day | GY_data_day-1__to__GY_data_day-26 | GY_data_day-34 | 44 | accuracy | 0.5575 |
| classification | lstm | Reaching B | cross_day | GY_data_day-1__to__GY_data_day-26 | GY_data_day-34 | 46 | accuracy | 0.4325 |
| classification | lstm | Reaching B | cross_day | GY_data_day-1__to__GY_data_day-26 | GY_data_day-34 | 54 | accuracy | 0.5325 |
| classification | lstm | Reaching B | cross_day | GY_data_day-1__to__GY_data_day-26 | GY_data_day-35 | 44 | accuracy | 0.36125 |
| classification | lstm | Reaching B | cross_day | GY_data_day-1__to__GY_data_day-26 | GY_data_day-35 | 46 | accuracy | 0.2925 |
| classification | lstm | Reaching B | cross_day | GY_data_day-1__to__GY_data_day-26 | GY_data_day-35 | 54 | accuracy | 0.29 |
| classification | lstm | Reaching B | cross_day | GY_data_day-1__to__GY_data_day-26 | GY_data_day-36 | 44 | accuracy | 0.50625 |
| classification | lstm | Reaching B | cross_day | GY_data_day-1__to__GY_data_day-26 | GY_data_day-36 | 46 | accuracy | 0.36875 |

| | | | | | | | | |
|---|---|---|---|---|---|---|---|---|
| classification | lstm | Reaching B | cross_day | GY_data_day-1__to__GY_data_day-26 | GY_data_day-36 | 54 | accuracy | 0.44875 |
| classification | lstm | Reaching B | cross_day | GY_data_day-1__to__GY_data_day-26 | GY_data_day-37 | 44 | accuracy | 0.7525 |
| classification | lstm | Reaching B | cross_day | GY_data_day-1__to__GY_data_day-26 | GY_data_day-37 | 46 | accuracy | 0.7825 |
| classification | lstm | Reaching B | cross_day | GY_data_day-1__to__GY_data_day-26 | GY_data_day-37 | 54 | accuracy | 0.52 |
| classification | transformer | Reaching B | cross_day | GY_data_day-1__to__GY_data_day-26 | GY_data_day-30 | 44 | accuracy | 0.3225 |
| classification | transformer | Reaching B | cross_day | GY_data_day-1__to__GY_data_day-26 | GY_data_day-30 | 46 | accuracy | 0.39 |
| classification | transformer | Reaching B | cross_day | GY_data_day-1__to__GY_data_day-26 | GY_data_day-30 | 54 | accuracy | 0.3925 |
| classification | transformer | Reaching B | cross_day | GY_data_day-1__to__GY_data_day-26 | GY_data_day-31 | 44 | accuracy | 0.51 |
| classification | transformer | Reaching B | cross_day | GY_data_day-1__to__GY_data_day-26 | GY_data_day-31 | 46 | accuracy | 0.37 |
| classification | transformer | Reaching B | cross_day | GY_data_day-1__to__GY_data_day-26 | GY_data_day-31 | 54 | accuracy | 0.49 |
| classification | transformer | Reaching B | cross_day | GY_data_day-1__to__GY_data_day-26 | GY_data_day-32 | 44 | accuracy | 0.4 |
| classification | transformer | Reaching B | cross_day | GY_data_day-1__to__GY_data_day-26 | GY_data_day-32 | 46 | accuracy | 0.515 |
| classification | transformer | Reaching B | cross_day | GY_data_day-1__to__GY_data_day-26 | GY_data_day-32 | 54 | accuracy | 0.52 |
| classification | transformer | Reaching B | cross_day | GY_data_day-1__to__GY_data_day-26 | GY_data_day-33 | 44 | accuracy | 0.335 |
| classification | transformer | Reaching B | cross_day | GY_data_day-1__to__GY_data_day-26 | GY_data_day-33 | 46 | accuracy | 0.355 |
| classification | transformer | Reaching B | cross_day | GY_data_day-1__to__GY_data_day-26 | GY_data_day-33 | 54 | accuracy | 0.358333 |
| classification | transformer | Reaching B | cross_day | GY_data_day-1__to__GY_data_day-26 | GY_data_day-34 | 44 | accuracy | 0.34375 |
| classification | transformer | Reaching B | cross_day | GY_data_day-1__to__GY_data_day-26 | GY_data_day-34 | 46 | accuracy | 0.34125 |
| classification | transformer | Reaching B | cross_day | GY_data_day-1__to__GY_data_day-26 | GY_data_day-34 | 54 | accuracy | 0.375 |
| classification | transformer | Reaching B | cross_day | GY_data_day-1__to__GY_data_day-26 | GY_data_day-35 | 44 | accuracy | 0.2375 |
| classification | transformer | Reaching B | cross_day | GY_data_day-1__to__GY_data_day-26 | GY_data_day-35 | 46 | accuracy | 0.23375 |
| classification | transformer | Reaching B | cross_day | GY_data_day-1__to__GY_data_day-26 | GY_data_day-35 | 54 | accuracy | 0.32875 |
| classification | transformer | Reaching B | cross_day | GY_data_day-1__to__GY_data_day-26 | GY_data_day-36 | 44 | accuracy | 0.33625 |
| classification | transformer | Reaching B | cross_day | GY_data_day-1__to__GY_data_day-26 | GY_data_day-36 | 46 | accuracy | 0.3475 |
| classification | transformer | Reaching B | cross_day | GY_data_day-1__to__GY_data_day-26 | GY_data_day-36 | 54 | accuracy | 0.4375 |
| classification | transformer | Reaching B | cross_day | GY_data_day-1__to__GY_data_day-26 | GY_data_day-37 | 44 | accuracy | 0.7425 |
| classification | transformer | Reaching B | cross_day | GY_data_day-1__to__GY_data_day-26 | GY_data_day-37 | 46 | accuracy | 0.6625 |
| classification | transformer | Reaching B | cross_day | GY_data_day-1__to__GY_data_day-26 | GY_data_day-37 | 54 | accuracy | 0.7275 |
| classification | lfads | Reaching B | cross_day | GY_data_day-1__to__GY_data_day-26 | GY_data_day-30 | 44 | accuracy | 0.58 |
| classification | lfads | Reaching B | cross_day | GY_data_day-1__to__GY_data_day-26 | GY_data_day-30 | 46 | accuracy | 0.65 |
| classification | lfads | Reaching B | cross_day | GY_data_day-1__to__GY_data_day-26 | GY_data_day-30 | 54 | accuracy | 0.5825 |
| classification | lfads | Reaching B | cross_day | GY_data_day-1__to__GY_data_day-26 | GY_data_day-31 | 44 | accuracy | 0.585 |
| classification | lfads | Reaching B | cross_day | GY_data_day-1__to__GY_data_day-26 | GY_data_day-31 | 46 | accuracy | 0.685 |
| classification | lfads | Reaching B | cross_day | GY_data_day-1__to__GY_data_day-26 | GY_data_day-31 | 54 | accuracy | 0.635 |
| classification | lfads | Reaching B | cross_day | GY_data_day-1__to__GY_data_day-26 | GY_data_day-32 | 44 | accuracy | 0.76 |
| classification | lfads | Reaching B | cross_day | GY_data_day-1__to__GY_data_day-26 | GY_data_day-32 | 46 | accuracy | 0.605 |
| classification | lfads | Reaching B | cross_day | GY_data_day-1__to__GY_data_day-26 | GY_data_day-32 | 54 | accuracy | 0.7275 |
| classification | lfads | Reaching B | cross_day | GY_data_day-1__to__GY_data_day-26 | GY_data_day-33 | 44 | accuracy | 0.556667 |
| classification | lfads | Reaching B | cross_day | GY_data_day-1__to__GY_data_day-26 | GY_data_day-33 | 46 | accuracy | 0.508333 |
| classification | lfads | Reaching B | cross_day | GY_data_day-1__to__GY_data_day-26 | GY_data_day-33 | 54 | accuracy | 0.566667 |
| classification | lfads | Reaching B | cross_day | GY_data_day-1__to__GY_data_day-26 | GY_data_day-34 | 44 | accuracy | 0.5125 |
| classification | lfads | Reaching B | cross_day | GY_data_day-1__to__GY_data_day-26 | GY_data_day-34 | 46 | accuracy | 0.38625 |
| classification | lfads | Reaching B | cross_day | GY_data_day-1__to__GY_data_day-26 | GY_data_day-34 | 54 | accuracy | 0.56875 |
| classification | lfads | Reaching B | cross_day | GY_data_day-1__to__GY_data_day-26 | GY_data_day-35 | 44 | accuracy | 0.4025 |
| classification | lfads | Reaching B | cross_day | GY_data_day-1__to__GY_data_day-26 | GY_data_day-35 | 46 | accuracy | 0.28375 |
| classification | lfads | Reaching B | cross_day | GY_data_day-1__to__GY_data_day-26 | GY_data_day-35 | 54 | accuracy | 0.40125 |
| classification | lfads | Reaching B | cross_day | GY_data_day-1__to__GY_data_day-26 | GY_data_day-36 | 44 | accuracy | 0.51875 |
| classification | lfads | Reaching B | cross_day | GY_data_day-1__to__GY_data_day-26 | GY_data_day-36 | 46 | accuracy | 0.4225 |
| classification | lfads | Reaching B | cross_day | GY_data_day-1__to__GY_data_day-26 | GY_data_day-36 | 54 | accuracy | 0.405 |
| classification | lfads | Reaching B | cross_day | GY_data_day-1__to__GY_data_day-26 | GY_data_day-37 | 44 | accuracy | 0.43 |
| classification | lfads | Reaching B | cross_day | GY_data_day-1__to__GY_data_day-26 | GY_data_day-37 | 46 | accuracy | 0.4925 |
| classification | lfads | Reaching B | cross_day | GY_data_day-1__to__GY_data_day-26 | GY_data_day-37 | 54 | accuracy | 0.465 |
| classification | cycle_gan | Reaching B | cross_day | GY_data_day-1__to__GY_data_day-26 | GY_data_day-30 | 44 | accuracy | 0.67027 |
| classification | cycle_gan | Reaching B | cross_day | GY_data_day-1__to__GY_data_day-26 | GY_data_day-30 | 46 | accuracy | 0.602703 |
| classification | cycle_gan | Reaching B | cross_day | GY_data_day-1__to__GY_data_day-26 | GY_data_day-30 | 54 | accuracy | 0.694595 |
| classification | cycle_gan | Reaching B | cross_day | GY_data_day-1__to__GY_data_day-26 | GY_data_day-31 | 44 | accuracy | 0.838889 |
| classification | cycle_gan | Reaching B | cross_day | GY_data_day-1__to__GY_data_day-26 | GY_data_day-31 | 46 | accuracy | 0.766667 |
| classification | cycle_gan | Reaching B | cross_day | GY_data_day-1__to__GY_data_day-26 | GY_data_day-31 | 54 | accuracy | 0.855556 |
| classification | cycle_gan | Reaching B | cross_day | GY_data_day-1__to__GY_data_day-26 | GY_data_day-32 | 44 | accuracy | 0.856757 |
| classification | cycle_gan | Reaching B | cross_day | GY_data_day-1__to__GY_data_day-26 | GY_data_day-32 | 46 | accuracy | 0.813514 |
| classification | cycle_gan | Reaching B | cross_day | GY_data_day-1__to__GY_data_day-26 | GY_data_day-32 | 54 | accuracy | 0.82973 |
| classification | cycle_gan | Reaching B | cross_day | GY_data_day-1__to__GY_data_day-26 | GY_data_day-33 | 44 | accuracy | 0.682456 |
| classification | cycle_gan | Reaching B | cross_day | GY_data_day-1__to__GY_data_day-26 | GY_data_day-33 | 46 | accuracy | 0.67193 |

| | | | | | | | | |
|---|---|---|---|---|---|---|---|---|
| classification | cycle_gan | Reaching B | cross_day | GY_data_day-1__to__GY_data_day-26 | GY_data_day-33 | 54 | accuracy | 0.714035 |
| classification | cycle_gan | Reaching B | cross_day | GY_data_day-1__to__GY_data_day-26 | GY_data_day-34 | 44 | accuracy | 0.62987 |
| classification | cycle_gan | Reaching B | cross_day | GY_data_day-1__to__GY_data_day-26 | GY_data_day-34 | 46 | accuracy | 0.581818 |
| classification | cycle_gan | Reaching B | cross_day | GY_data_day-1__to__GY_data_day-26 | GY_data_day-34 | 54 | accuracy | 0.658442 |
| classification | cycle_gan | Reaching B | cross_day | GY_data_day-1__to__GY_data_day-26 | GY_data_day-35 | 44 | accuracy | 0.576623 |
| classification | cycle_gan | Reaching B | cross_day | GY_data_day-1__to__GY_data_day-26 | GY_data_day-35 | 46 | accuracy | 0.441558 |
| classification | cycle_gan | Reaching B | cross_day | GY_data_day-1__to__GY_data_day-26 | GY_data_day-35 | 54 | accuracy | 0.605195 |
| classification | cycle_gan | Reaching B | cross_day | GY_data_day-1__to__GY_data_day-26 | GY_data_day-36 | 44 | accuracy | 0.502597 |
| classification | cycle_gan | Reaching B | cross_day | GY_data_day-1__to__GY_data_day-26 | GY_data_day-36 | 46 | accuracy | 0.394805 |
| classification | cycle_gan | Reaching B | cross_day | GY_data_day-1__to__GY_data_day-26 | GY_data_day-36 | 54 | accuracy | 0.503896 |
| classification | cycle_gan | Reaching B | cross_day | GY_data_day-1__to__GY_data_day-26 | GY_data_day-37 | 44 | accuracy | 0.781081 |
| classification | cycle_gan | Reaching B | cross_day | GY_data_day-1__to__GY_data_day-26 | GY_data_day-37 | 46 | accuracy | 0.772973 |
| classification | cycle_gan | Reaching B | cross_day | GY_data_day-1__to__GY_data_day-26 | GY_data_day-37 | 54 | accuracy | 0.832432 |
| classification | stabilization | Reaching B | cross_day | GY_data_day-1__to__GY_data_day-26 | GY_data_day-30 | 44 | accuracy | 0.87027 |
| classification | stabilization | Reaching B | cross_day | GY_data_day-1__to__GY_data_day-26 | GY_data_day-30 | 46 | accuracy | 0.859459 |
| classification | stabilization | Reaching B | cross_day | GY_data_day-1__to__GY_data_day-26 | GY_data_day-30 | 54 | accuracy | 0.821622 |
| classification | stabilization | Reaching B | cross_day | GY_data_day-1__to__GY_data_day-26 | GY_data_day-31 | 44 | accuracy | 0.794444 |
| classification | stabilization | Reaching B | cross_day | GY_data_day-1__to__GY_data_day-26 | GY_data_day-31 | 46 | accuracy | 0.85 |
| classification | stabilization | Reaching B | cross_day | GY_data_day-1__to__GY_data_day-26 | GY_data_day-31 | 54 | accuracy | 0.827778 |
| classification | stabilization | Reaching B | cross_day | GY_data_day-1__to__GY_data_day-26 | GY_data_day-32 | 44 | accuracy | 0.87027 |
| classification | stabilization | Reaching B | cross_day | GY_data_day-1__to__GY_data_day-26 | GY_data_day-32 | 46 | accuracy | 0.875676 |
| classification | stabilization | Reaching B | cross_day | GY_data_day-1__to__GY_data_day-26 | GY_data_day-32 | 54 | accuracy | 0.82973 |
| classification | stabilization | Reaching B | cross_day | GY_data_day-1__to__GY_data_day-26 | GY_data_day-33 | 44 | accuracy | 0.733333 |
| classification | stabilization | Reaching B | cross_day | GY_data_day-1__to__GY_data_day-26 | GY_data_day-33 | 46 | accuracy | 0.791228 |
| classification | stabilization | Reaching B | cross_day | GY_data_day-1__to__GY_data_day-26 | GY_data_day-33 | 54 | accuracy | 0.719298 |
| classification | stabilization | Reaching B | cross_day | GY_data_day-1__to__GY_data_day-26 | GY_data_day-34 | 44 | accuracy | 0.775325 |
| classification | stabilization | Reaching B | cross_day | GY_data_day-1__to__GY_data_day-26 | GY_data_day-34 | 46 | accuracy | 0.675325 |
| classification | stabilization | Reaching B | cross_day | GY_data_day-1__to__GY_data_day-26 | GY_data_day-34 | 54 | accuracy | 0.816883 |
| classification | stabilization | Reaching B | cross_day | GY_data_day-1__to__GY_data_day-26 | GY_data_day-35 | 44 | accuracy | 0.748052 |
| classification | stabilization | Reaching B | cross_day | GY_data_day-1__to__GY_data_day-26 | GY_data_day-35 | 46 | accuracy | 0.720779 |
| classification | stabilization | Reaching B | cross_day | GY_data_day-1__to__GY_data_day-26 | GY_data_day-35 | 54 | accuracy | 0.732468 |
| classification | stabilization | Reaching B | cross_day | GY_data_day-1__to__GY_data_day-26 | GY_data_day-36 | 44 | accuracy | 0.741558 |
| classification | stabilization | Reaching B | cross_day | GY_data_day-1__to__GY_data_day-26 | GY_data_day-36 | 46 | accuracy | 0.711688 |
| classification | stabilization | Reaching B | cross_day | GY_data_day-1__to__GY_data_day-26 | GY_data_day-36 | 54 | accuracy | 0.757143 |
| classification | stabilization | Reaching B | cross_day | GY_data_day-1__to__GY_data_day-26 | GY_data_day-37 | 44 | accuracy | 0.708108 |
| classification | stabilization | Reaching B | cross_day | GY_data_day-1__to__GY_data_day-26 | GY_data_day-37 | 46 | accuracy | 0.643243 |
| classification | stabilization | Reaching B | cross_day | GY_data_day-1__to__GY_data_day-26 | GY_data_day-37 | 54 | accuracy | 0.610811 |
| classification | mfsnn | Reaching B | cross_day | GY_data_day-1__to__GY_data_day-26 | GY_data_day-30 | 44 | accuracy | 0.3125 |
| classification | mfsnn | Reaching B | cross_day | GY_data_day-1__to__GY_data_day-26 | GY_data_day-30 | 46 | accuracy | 0.5225 |
| classification | mfsnn | Reaching B | cross_day | GY_data_day-1__to__GY_data_day-26 | GY_data_day-30 | 54 | accuracy | 0.2475 |
| classification | mfsnn | Reaching B | cross_day | GY_data_day-1__to__GY_data_day-26 | GY_data_day-31 | 44 | accuracy | 0.195 |
| classification | mfsnn | Reaching B | cross_day | GY_data_day-1__to__GY_data_day-26 | GY_data_day-31 | 46 | accuracy | 0.25 |
| classification | mfsnn | Reaching B | cross_day | GY_data_day-1__to__GY_data_day-26 | GY_data_day-31 | 54 | accuracy | 0.135 |
| classification | mfsnn | Reaching B | cross_day | GY_data_day-1__to__GY_data_day-26 | GY_data_day-32 | 44 | accuracy | 0.17 |
| classification | mfsnn | Reaching B | cross_day | GY_data_day-1__to__GY_data_day-26 | GY_data_day-32 | 46 | accuracy | 0.255 |
| classification | mfsnn | Reaching B | cross_day | GY_data_day-1__to__GY_data_day-26 | GY_data_day-32 | 54 | accuracy | 0.2225 |
| classification | mfsnn | Reaching B | cross_day | GY_data_day-1__to__GY_data_day-26 | GY_data_day-33 | 44 | accuracy | 0.195 |
| classification | mfsnn | Reaching B | cross_day | GY_data_day-1__to__GY_data_day-26 | GY_data_day-33 | 46 | accuracy | 0.235 |
| classification | mfsnn | Reaching B | cross_day | GY_data_day-1__to__GY_data_day-26 | GY_data_day-33 | 54 | accuracy | 0.176667 |
| classification | mfsnn | Reaching B | cross_day | GY_data_day-1__to__GY_data_day-26 | GY_data_day-34 | 44 | accuracy | 0.25 |
| classification | mfsnn | Reaching B | cross_day | GY_data_day-1__to__GY_data_day-26 | GY_data_day-34 | 46 | accuracy | 0.3525 |
| classification | mfsnn | Reaching B | cross_day | GY_data_day-1__to__GY_data_day-26 | GY_data_day-34 | 54 | accuracy | 0.27125 |
| classification | mfsnn | Reaching B | cross_day | GY_data_day-1__to__GY_data_day-26 | GY_data_day-35 | 44 | accuracy | 0.21875 |
| classification | mfsnn | Reaching B | cross_day | GY_data_day-1__to__GY_data_day-26 | GY_data_day-35 | 46 | accuracy | 0.38 |
| classification | mfsnn | Reaching B | cross_day | GY_data_day-1__to__GY_data_day-26 | GY_data_day-35 | 54 | accuracy | 0.245 |
| classification | mfsnn | Reaching B | cross_day | GY_data_day-1__to__GY_data_day-26 | GY_data_day-36 | 44 | accuracy | 0.1425 |
| classification | mfsnn | Reaching B | cross_day | GY_data_day-1__to__GY_data_day-26 | GY_data_day-36 | 46 | accuracy | 0.355 |
| classification | mfsnn | Reaching B | cross_day | GY_data_day-1__to__GY_data_day-26 | GY_data_day-36 | 54 | accuracy | 0.14 |
| classification | mfsnn | Reaching B | cross_day | GY_data_day-1__to__GY_data_day-26 | GY_data_day-37 | 44 | accuracy | 0.2725 |
| classification | mfsnn | Reaching B | cross_day | GY_data_day-1__to__GY_data_day-26 | GY_data_day-37 | 46 | accuracy | 0.405 |
| classification | mfsnn | Reaching B | cross_day | GY_data_day-1__to__GY_data_day-26 | GY_data_day-37 | 54 | accuracy | 0.145 |
| classification | mscformer | Reaching B | cross_day | GY_data_day-1__to__GY_data_day-26 | GY_data_day-30 | 44 | accuracy | 0.8825 |
| classification | mscformer | Reaching B | cross_day | GY_data_day-1__to__GY_data_day-26 | GY_data_day-30 | 46 | accuracy | 0.8575 |

| | | | | | | | | |
|---|---|---|---|---|---|---|---|---|
| classification | mscformer | Reaching B | cross_day | GY_data_day-1__to__GY_data_day-26 | GY_data_day-30 | 54 | accuracy | 0.88 |
| classification | mscformer | Reaching B | cross_day | GY_data_day-1__to__GY_data_day-26 | GY_data_day-31 | 44 | accuracy | 0.885 |
| classification | mscformer | Reaching B | cross_day | GY_data_day-1__to__GY_data_day-26 | GY_data_day-31 | 46 | accuracy | 0.855 |
| classification | mscformer | Reaching B | cross_day | GY_data_day-1__to__GY_data_day-26 | GY_data_day-31 | 54 | accuracy | 0.83 |
| classification | mscformer | Reaching B | cross_day | GY_data_day-1__to__GY_data_day-26 | GY_data_day-32 | 44 | accuracy | 0.7125 |
| classification | mscformer | Reaching B | cross_day | GY_data_day-1__to__GY_data_day-26 | GY_data_day-32 | 46 | accuracy | 0.7275 |
| classification | mscformer | Reaching B | cross_day | GY_data_day-1__to__GY_data_day-26 | GY_data_day-32 | 54 | accuracy | 0.74 |
| classification | mscformer | Reaching B | cross_day | GY_data_day-1__to__GY_data_day-26 | GY_data_day-33 | 44 | accuracy | 0.74 |
| classification | mscformer | Reaching B | cross_day | GY_data_day-1__to__GY_data_day-26 | GY_data_day-33 | 46 | accuracy | 0.705 |
| classification | mscformer | Reaching B | cross_day | GY_data_day-1__to__GY_data_day-26 | GY_data_day-33 | 54 | accuracy | 0.673333 |
| classification | mscformer | Reaching B | cross_day | GY_data_day-1__to__GY_data_day-26 | GY_data_day-34 | 44 | accuracy | 0.805 |
| classification | mscformer | Reaching B | cross_day | GY_data_day-1__to__GY_data_day-26 | GY_data_day-34 | 46 | accuracy | 0.81375 |
| classification | mscformer | Reaching B | cross_day | GY_data_day-1__to__GY_data_day-26 | GY_data_day-34 | 54 | accuracy | 0.77875 |
| classification | mscformer | Reaching B | cross_day | GY_data_day-1__to__GY_data_day-26 | GY_data_day-35 | 44 | accuracy | 0.7275 |
| classification | mscformer | Reaching B | cross_day | GY_data_day-1__to__GY_data_day-26 | GY_data_day-35 | 46 | accuracy | 0.72375 |
| classification | mscformer | Reaching B | cross_day | GY_data_day-1__to__GY_data_day-26 | GY_data_day-35 | 54 | accuracy | 0.74 |
| classification | mscformer | Reaching B | cross_day | GY_data_day-1__to__GY_data_day-26 | GY_data_day-36 | 44 | accuracy | 0.64625 |
| classification | mscformer | Reaching B | cross_day | GY_data_day-1__to__GY_data_day-26 | GY_data_day-36 | 46 | accuracy | 0.715 |
| classification | mscformer | Reaching B | cross_day | GY_data_day-1__to__GY_data_day-26 | GY_data_day-36 | 54 | accuracy | 0.70875 |
| classification | mscformer | Reaching B | cross_day | GY_data_day-1__to__GY_data_day-26 | GY_data_day-37 | 44 | accuracy | 0.77 |
| classification | mscformer | Reaching B | cross_day | GY_data_day-1__to__GY_data_day-26 | GY_data_day-37 | 46 | accuracy | 0.8325 |
| classification | mscformer | Reaching B | cross_day | GY_data_day-1__to__GY_data_day-26 | GY_data_day-37 | 54 | accuracy | 0.845 |
| classification | nomad | Reaching B | cross_day | GY_data_day-1__to__GY_data_day-26 | GY_data_day-30 | 44 | accuracy | 0.9175 |
| classification | nomad | Reaching B | cross_day | GY_data_day-1__to__GY_data_day-26 | GY_data_day-30 | 46 | accuracy | 0.8875 |
| classification | nomad | Reaching B | cross_day | GY_data_day-1__to__GY_data_day-26 | GY_data_day-30 | 54 | accuracy | 0.83 |
| classification | nomad | Reaching B | cross_day | GY_data_day-1__to__GY_data_day-26 | GY_data_day-31 | 44 | accuracy | 0.9 |
| classification | nomad | Reaching B | cross_day | GY_data_day-1__to__GY_data_day-26 | GY_data_day-31 | 46 | accuracy | 0.855 |
| classification | nomad | Reaching B | cross_day | GY_data_day-1__to__GY_data_day-26 | GY_data_day-31 | 54 | accuracy | 0.865 |
| classification | nomad | Reaching B | cross_day | GY_data_day-1__to__GY_data_day-26 | GY_data_day-32 | 44 | accuracy | 0.7925 |
| classification | nomad | Reaching B | cross_day | GY_data_day-1__to__GY_data_day-26 | GY_data_day-32 | 46 | accuracy | 0.7725 |
| classification | nomad | Reaching B | cross_day | GY_data_day-1__to__GY_data_day-26 | GY_data_day-32 | 54 | accuracy | 0.7125 |
| classification | nomad | Reaching B | cross_day | GY_data_day-1__to__GY_data_day-26 | GY_data_day-33 | 44 | accuracy | 0.823333 |
| classification | nomad | Reaching B | cross_day | GY_data_day-1__to__GY_data_day-26 | GY_data_day-33 | 46 | accuracy | 0.833333 |
| classification | nomad | Reaching B | cross_day | GY_data_day-1__to__GY_data_day-26 | GY_data_day-33 | 54 | accuracy | 0.781667 |
| classification | nomad | Reaching B | cross_day | GY_data_day-1__to__GY_data_day-26 | GY_data_day-34 | 44 | accuracy | 0.8125 |
| classification | nomad | Reaching B | cross_day | GY_data_day-1__to__GY_data_day-26 | GY_data_day-34 | 46 | accuracy | 0.82125 |
| classification | nomad | Reaching B | cross_day | GY_data_day-1__to__GY_data_day-26 | GY_data_day-34 | 54 | accuracy | 0.8275 |
| classification | nomad | Reaching B | cross_day | GY_data_day-1__to__GY_data_day-26 | GY_data_day-35 | 44 | accuracy | 0.72125 |
| classification | nomad | Reaching B | cross_day | GY_data_day-1__to__GY_data_day-26 | GY_data_day-35 | 46 | accuracy | 0.72 |
| classification | nomad | Reaching B | cross_day | GY_data_day-1__to__GY_data_day-26 | GY_data_day-35 | 54 | accuracy | 0.73125 |
| classification | nomad | Reaching B | cross_day | GY_data_day-1__to__GY_data_day-26 | GY_data_day-36 | 44 | accuracy | 0.65625 |
| classification | nomad | Reaching B | cross_day | GY_data_day-1__to__GY_data_day-26 | GY_data_day-36 | 46 | accuracy | 0.59625 |
| classification | nomad | Reaching B | cross_day | GY_data_day-1__to__GY_data_day-26 | GY_data_day-36 | 54 | accuracy | 0.61375 |
| classification | nomad | Reaching B | cross_day | GY_data_day-1__to__GY_data_day-26 | GY_data_day-37 | 44 | accuracy | 0.91 |
| classification | nomad | Reaching B | cross_day | GY_data_day-1__to__GY_data_day-26 | GY_data_day-37 | 46 | accuracy | 0.9175 |
| classification | nomad | Reaching B | cross_day | GY_data_day-1__to__GY_data_day-26 | GY_data_day-37 | 54 | accuracy | 0.89 |
| classification | mlp | Reaching B | cross_day | GY_data_day-1__to__GY_data_day-26 | GY_data_day-30 | 44 | accuracy | 0.6825 |
| classification | mlp | Reaching B | cross_day | GY_data_day-1__to__GY_data_day-26 | GY_data_day-30 | 46 | accuracy | 0.6575 |
| classification | mlp | Reaching B | cross_day | GY_data_day-1__to__GY_data_day-26 | GY_data_day-30 | 54 | accuracy | 0.7425 |
| classification | mlp | Reaching B | cross_day | GY_data_day-1__to__GY_data_day-26 | GY_data_day-31 | 44 | accuracy | 0.83 |
| classification | mlp | Reaching B | cross_day | GY_data_day-1__to__GY_data_day-26 | GY_data_day-31 | 46 | accuracy | 0.815 |
| classification | mlp | Reaching B | cross_day | GY_data_day-1__to__GY_data_day-26 | GY_data_day-31 | 54 | accuracy | 0.785 |
| classification | mlp | Reaching B | cross_day | GY_data_day-1__to__GY_data_day-26 | GY_data_day-32 | 44 | accuracy | 0.8525 |
| classification | mlp | Reaching B | cross_day | GY_data_day-1__to__GY_data_day-26 | GY_data_day-32 | 46 | accuracy | 0.8275 |
| classification | mlp | Reaching B | cross_day | GY_data_day-1__to__GY_data_day-26 | GY_data_day-32 | 54 | accuracy | 0.83 |
| classification | mlp | Reaching B | cross_day | GY_data_day-1__to__GY_data_day-26 | GY_data_day-33 | 44 | accuracy | 0.738333 |
| classification | mlp | Reaching B | cross_day | GY_data_day-1__to__GY_data_day-26 | GY_data_day-33 | 46 | accuracy | 0.756667 |
| classification | mlp | Reaching B | cross_day | GY_data_day-1__to__GY_data_day-26 | GY_data_day-33 | 54 | accuracy | 0.796667 |
| classification | mlp | Reaching B | cross_day | GY_data_day-1__to__GY_data_day-26 | GY_data_day-34 | 44 | accuracy | 0.53875 |
| classification | mlp | Reaching B | cross_day | GY_data_day-1__to__GY_data_day-26 | GY_data_day-34 | 46 | accuracy | 0.55625 |
| classification | mlp | Reaching B | cross_day | GY_data_day-1__to__GY_data_day-26 | GY_data_day-34 | 54 | accuracy | 0.5925 |
| classification | mlp | Reaching B | cross_day | GY_data_day-1__to__GY_data_day-26 | GY_data_day-35 | 44 | accuracy | 0.41 |
| classification | mlp | Reaching B | cross_day | GY_data_day-1__to__GY_data_day-26 | GY_data_day-35 | 46 | accuracy | 0.46375 |

| | | | | | | | | |
|---|---|---|---|---|---|---|---|---|
| classification | mlp | Reaching B | cross_day | GY_data_day-1__to__GY_data_day-26 | GY_data_day-35 | 54 | accuracy | 0.49625 |
| classification | mlp | Reaching B | cross_day | GY_data_day-1__to__GY_data_day-26 | GY_data_day-36 | 44 | accuracy | 0.50375 |
| classification | mlp | Reaching B | cross_day | GY_data_day-1__to__GY_data_day-26 | GY_data_day-36 | 46 | accuracy | 0.47125 |
| classification | mlp | Reaching B | cross_day | GY_data_day-1__to__GY_data_day-26 | GY_data_day-36 | 54 | accuracy | 0.5125 |
| classification | mlp | Reaching B | cross_day | GY_data_day-1__to__GY_data_day-26 | GY_data_day-37 | 44 | accuracy | 0.7775 |
| classification | mlp | Reaching B | cross_day | GY_data_day-1__to__GY_data_day-26 | GY_data_day-37 | 46 | accuracy | 0.7325 |
| classification | mlp | Reaching B | cross_day | GY_data_day-1__to__GY_data_day-26 | GY_data_day-37 | 54 | accuracy | 0.755 |
| classification | lstm | Jango | single_day | Jango_20150730_001 | Jango_20150730_001 | 31 | accuracy | 0.939394 |
| classification | lstm | Jango | single_day | Jango_20150730_001 | Jango_20150730_001 | 44 | accuracy | 0.969697 |
| classification | lstm | Jango | single_day | Jango_20150730_001 | Jango_20150730_001 | 44 | accuracy | 0.939394 |
| classification | lstm | Jango | single_day | Jango_20150730_001 | Jango_20150730_001 | 46 | accuracy | 0.939394 |
| classification | lstm | Jango | single_day | Jango_20150730_001 | Jango_20150730_001 | 46 | accuracy | 0.878788 |
| classification | lstm | Jango | single_day | Jango_20150730_001 | Jango_20150730_001 | 54 | accuracy | 0.909091 |
| classification | lstm | Jango | single_day | Jango_20150730_001 | Jango_20150730_001 | 54 | accuracy | 0.939394 |
| classification | lstm | Jango | single_day | Jango_20150730_001 | Jango_20150730_001 | 69 | accuracy | 0.909091 |
| classification | lstm | Jango | single_day | Jango_20150731_001 | Jango_20150731_001 | 31 | accuracy | 0.95 |
| classification | lstm | Jango | single_day | Jango_20150731_001 | Jango_20150731_001 | 44 | accuracy | 0.975 |
| classification | lstm | Jango | single_day | Jango_20150731_001 | Jango_20150731_001 | 44 | accuracy | 0.925 |
| classification | lstm | Jango | single_day | Jango_20150731_001 | Jango_20150731_001 | 46 | accuracy | 0.95 |
| classification | lstm | Jango | single_day | Jango_20150731_001 | Jango_20150731_001 | 46 | accuracy | 0.975 |
| classification | lstm | Jango | single_day | Jango_20150731_001 | Jango_20150731_001 | 54 | accuracy | 0.975 |
| classification | lstm | Jango | single_day | Jango_20150731_001 | Jango_20150731_001 | 54 | accuracy | 0.9 |
| classification | lstm | Jango | single_day | Jango_20150731_001 | Jango_20150731_001 | 69 | accuracy | 0.95 |
| classification | lstm | Jango | single_day | Jango_20150801_001 | Jango_20150801_001 | 31 | accuracy | 0.954545 |
| classification | lstm | Jango | single_day | Jango_20150801_001 | Jango_20150801_001 | 44 | accuracy | 0.954545 |
| classification | lstm | Jango | single_day | Jango_20150801_001 | Jango_20150801_001 | 44 | accuracy | 0.977273 |
| classification | lstm | Jango | single_day | Jango_20150801_001 | Jango_20150801_001 | 46 | accuracy | 0.977273 |
| classification | lstm | Jango | single_day | Jango_20150801_001 | Jango_20150801_001 | 46 | accuracy | 0.954545 |
| classification | lstm | Jango | single_day | Jango_20150801_001 | Jango_20150801_001 | 54 | accuracy | 0.977273 |
| classification | lstm | Jango | single_day | Jango_20150801_001 | Jango_20150801_001 | 54 | accuracy | 0.863636 |
| classification | lstm | Jango | single_day | Jango_20150801_001 | Jango_20150801_001 | 69 | accuracy | 0.909091 |
| classification | lstm | Jango | single_day | Jango_20150805_001 | Jango_20150805_001 | 31 | accuracy | 1 |
| classification | lstm | Jango | single_day | Jango_20150805_001 | Jango_20150805_001 | 44 | accuracy | 0.977778 |
| classification | lstm | Jango | single_day | Jango_20150805_001 | Jango_20150805_001 | 44 | accuracy | 0.933333 |
| classification | lstm | Jango | single_day | Jango_20150805_001 | Jango_20150805_001 | 46 | accuracy | 0.977778 |
| classification | lstm | Jango | single_day | Jango_20150805_001 | Jango_20150805_001 | 46 | accuracy | 0.933333 |
| classification | lstm | Jango | single_day | Jango_20150805_001 | Jango_20150805_001 | 54 | accuracy | 0.933333 |
| classification | lstm | Jango | single_day | Jango_20150805_001 | Jango_20150805_001 | 54 | accuracy | 0.911111 |
| classification | lstm | Jango | single_day | Jango_20150805_001 | Jango_20150805_001 | 69 | accuracy | 1 |
| classification | lstm | Jango | single_day | Jango_20150806_001 | Jango_20150806_001 | 31 | accuracy | 0.977778 |
| classification | lstm | Jango | single_day | Jango_20150806_001 | Jango_20150806_001 | 44 | accuracy | 0.977778 |
| classification | lstm | Jango | single_day | Jango_20150806_001 | Jango_20150806_001 | 44 | accuracy | 0.844444 |
| classification | lstm | Jango | single_day | Jango_20150806_001 | Jango_20150806_001 | 46 | accuracy | 0.977778 |
| classification | lstm | Jango | single_day | Jango_20150806_001 | Jango_20150806_001 | 46 | accuracy | 1 |
| classification | lstm | Jango | single_day | Jango_20150806_001 | Jango_20150806_001 | 54 | accuracy | 0.955556 |
| classification | lstm | Jango | single_day | Jango_20150806_001 | Jango_20150806_001 | 54 | accuracy | 0.844444 |
| classification | lstm | Jango | single_day | Jango_20150806_001 | Jango_20150806_001 | 69 | accuracy | 0.955556 |
| classification | lstm | Jango | single_day | Jango_20150807_001 | Jango_20150807_001 | 31 | accuracy | 0.955556 |
| classification | lstm | Jango | single_day | Jango_20150807_001 | Jango_20150807_001 | 44 | accuracy | 0.977778 |
| classification | lstm | Jango | single_day | Jango_20150807_001 | Jango_20150807_001 | 46 | accuracy | 1 |
| classification | lstm | Jango | single_day | Jango_20150807_001 | Jango_20150807_001 | 46 | accuracy | 0.933333 |
| classification | lstm | Jango | single_day | Jango_20150807_001 | Jango_20150807_001 | 54 | accuracy | 0.977778 |
| classification | lstm | Jango | single_day | Jango_20150807_001 | Jango_20150807_001 | 54 | accuracy | 0.888889 |
| classification | lstm | Jango | single_day | Jango_20150807_001 | Jango_20150807_001 | 69 | accuracy | 1 |
| classification | lstm | Jango | single_day | Jango_20150808_001 | Jango_20150808_001 | 31 | accuracy | 0.933333 |
| classification | lstm | Jango | single_day | Jango_20150808_001 | Jango_20150808_001 | 44 | accuracy | 0.933333 |
| classification | lstm | Jango | single_day | Jango_20150808_001 | Jango_20150808_001 | 44 | accuracy | 0.955556 |
| classification | lstm | Jango | single_day | Jango_20150808_001 | Jango_20150808_001 | 46 | accuracy | 0.955556 |
| classification | lstm | Jango | single_day | Jango_20150808_001 | Jango_20150808_001 | 46 | accuracy | 0.911111 |
| classification | lstm | Jango | single_day | Jango_20150808_001 | Jango_20150808_001 | 54 | accuracy | 0.933333 |
| classification | lstm | Jango | single_day | Jango_20150808_001 | Jango_20150808_001 | 54 | accuracy | 0.888889 |
| classification | lstm | Jango | single_day | Jango_20150808_001 | Jango_20150808_001 | 69 | accuracy | 0.977778 |
| classification | lstm | Jango | single_day | Jango_20150809_001 | Jango_20150809_001 | 31 | accuracy | 0.955556 |

| | | | | | | | | |
|---|---|---|---|---|---|---|---|---|
| classification | lstm | Jango | single_day | Jango_20150809_001 | Jango_20150809_001 | 44 | accuracy | 0.955556 |
| classification | lstm | Jango | single_day | Jango_20150809_001 | Jango_20150809_001 | 46 | accuracy | 1 |
| classification | lstm | Jango | single_day | Jango_20150809_001 | Jango_20150809_001 | 46 | accuracy | 0.911111 |
| classification | lstm | Jango | single_day | Jango_20150809_001 | Jango_20150809_001 | 54 | accuracy | 0.977778 |
| classification | lstm | Jango | single_day | Jango_20150809_001 | Jango_20150809_001 | 54 | accuracy | 0.955556 |
| classification | lstm | Jango | single_day | Jango_20150809_001 | Jango_20150809_001 | 69 | accuracy | 0.955556 |
| classification | lstm | Jango | single_day | Jango_20150820_001 | Jango_20150820_001 | 31 | accuracy | 0.911111 |
| classification | lstm | Jango | single_day | Jango_20150820_001 | Jango_20150820_001 | 44 | accuracy | 0.977778 |
| classification | lstm | Jango | single_day | Jango_20150820_001 | Jango_20150820_001 | 46 | accuracy | 0.888889 |
| classification | lstm | Jango | single_day | Jango_20150820_001 | Jango_20150820_001 | 54 | accuracy | 0.955556 |
| classification | lstm | Jango | single_day | Jango_20150820_001 | Jango_20150820_001 | 69 | accuracy | 0.977778 |
| classification | lstm | Jango | single_day | Jango_20150824_001 | Jango_20150824_001 | 31 | accuracy | 0.909091 |
| classification | lstm | Jango | single_day | Jango_20150824_001 | Jango_20150824_001 | 44 | accuracy | 0.909091 |
| classification | lstm | Jango | single_day | Jango_20150824_001 | Jango_20150824_001 | 44 | accuracy | 0.931818 |
| classification | lstm | Jango | single_day | Jango_20150824_001 | Jango_20150824_001 | 46 | accuracy | 0.931818 |
| classification | lstm | Jango | single_day | Jango_20150824_001 | Jango_20150824_001 | 46 | accuracy | 0.886364 |
| classification | lstm | Jango | single_day | Jango_20150824_001 | Jango_20150824_001 | 54 | accuracy | 0.954545 |
| classification | lstm | Jango | single_day | Jango_20150824_001 | Jango_20150824_001 | 54 | accuracy | 0.886364 |
| classification | lstm | Jango | single_day | Jango_20150824_001 | Jango_20150824_001 | 69 | accuracy | 0.931818 |
| classification | lstm | Jango | single_day | Jango_20150825_001 | Jango_20150825_001 | 31 | accuracy | 0.956522 |
| classification | lstm | Jango | single_day | Jango_20150825_001 | Jango_20150825_001 | 44 | accuracy | 0.956522 |
| classification | lstm | Jango | single_day | Jango_20150825_001 | Jango_20150825_001 | 44 | accuracy | 0.869565 |
| classification | lstm | Jango | single_day | Jango_20150825_001 | Jango_20150825_001 | 46 | accuracy | 0.978261 |
| classification | lstm | Jango | single_day | Jango_20150825_001 | Jango_20150825_001 | 46 | accuracy | 0.826087 |
| classification | lstm | Jango | single_day | Jango_20150825_001 | Jango_20150825_001 | 54 | accuracy | 0.913043 |
| classification | lstm | Jango | single_day | Jango_20150825_001 | Jango_20150825_001 | 69 | accuracy | 0.978261 |
| classification | lstm | Jango | single_day | Jango_20150826_001 | Jango_20150826_001 | 31 | accuracy | 1 |
| classification | lstm | Jango | single_day | Jango_20150826_001 | Jango_20150826_001 | 44 | accuracy | 0.933333 |
| classification | lstm | Jango | single_day | Jango_20150826_001 | Jango_20150826_001 | 44 | accuracy | 0.911111 |
| classification | lstm | Jango | single_day | Jango_20150826_001 | Jango_20150826_001 | 46 | accuracy | 0.933333 |
| classification | lstm | Jango | single_day | Jango_20150826_001 | Jango_20150826_001 | 46 | accuracy | 0.777778 |
| classification | lstm | Jango | single_day | Jango_20150826_001 | Jango_20150826_001 | 54 | accuracy | 1 |
| classification | lstm | Jango | single_day | Jango_20150826_001 | Jango_20150826_001 | 54 | accuracy | 0.933333 |
| classification | lstm | Jango | single_day | Jango_20150826_001 | Jango_20150826_001 | 69 | accuracy | 0.977778 |
| classification | lstm | Jango | single_day | Jango_20150827_001 | Jango_20150827_001 | 31 | accuracy | 0.978261 |
| classification | lstm | Jango | single_day | Jango_20150827_001 | Jango_20150827_001 | 44 | accuracy | 0.956522 |
| classification | lstm | Jango | single_day | Jango_20150827_001 | Jango_20150827_001 | 44 | accuracy | 0.934783 |
| classification | lstm | Jango | single_day | Jango_20150827_001 | Jango_20150827_001 | 46 | accuracy | 0.978261 |
| classification | lstm | Jango | single_day | Jango_20150827_001 | Jango_20150827_001 | 46 | accuracy | 0.956522 |
| classification | lstm | Jango | single_day | Jango_20150827_001 | Jango_20150827_001 | 54 | accuracy | 0.978261 |
| classification | lstm | Jango | single_day | Jango_20150827_001 | Jango_20150827_001 | 54 | accuracy | 0.956522 |
| classification | lstm | Jango | single_day | Jango_20150827_001 | Jango_20150827_001 | 69 | accuracy | 0.934783 |
| classification | lstm | Jango | single_day | Jango_20150828_001 | Jango_20150828_001 | 31 | accuracy | 0.930233 |
| classification | lstm | Jango | single_day | Jango_20150828_001 | Jango_20150828_001 | 44 | accuracy | 0.976744 |
| classification | lstm | Jango | single_day | Jango_20150828_001 | Jango_20150828_001 | 44 | accuracy | 0.953488 |
| classification | lstm | Jango | single_day | Jango_20150828_001 | Jango_20150828_001 | 46 | accuracy | 0.976744 |
| classification | lstm | Jango | single_day | Jango_20150828_001 | Jango_20150828_001 | 46 | accuracy | 0.906977 |
| classification | lstm | Jango | single_day | Jango_20150828_001 | Jango_20150828_001 | 54 | accuracy | 1 |
| classification | lstm | Jango | single_day | Jango_20150828_001 | Jango_20150828_001 | 54 | accuracy | 0.953488 |
| classification | lstm | Jango | single_day | Jango_20150828_001 | Jango_20150828_001 | 69 | accuracy | 0.976744 |
| classification | lstm | Jango | single_day | Jango_20150831_001 | Jango_20150831_001 | 31 | accuracy | 0.945946 |
| classification | lstm | Jango | single_day | Jango_20150831_001 | Jango_20150831_001 | 44 | accuracy | 1 |
| classification | lstm | Jango | single_day | Jango_20150831_001 | Jango_20150831_001 | 44 | accuracy | 0.918919 |
| classification | lstm | Jango | single_day | Jango_20150831_001 | Jango_20150831_001 | 46 | accuracy | 0.972973 |
| classification | lstm | Jango | single_day | Jango_20150831_001 | Jango_20150831_001 | 46 | accuracy | 0.945946 |
| classification | lstm | Jango | single_day | Jango_20150831_001 | Jango_20150831_001 | 54 | accuracy | 0.864865 |
| classification | lstm | Jango | single_day | Jango_20150831_001 | Jango_20150831_001 | 54 | accuracy | 1 |
| classification | lstm | Jango | single_day | Jango_20150831_001 | Jango_20150831_001 | 69 | accuracy | 0.945946 |
| classification | lstm | Jango | single_day | Jango_20150905_001 | Jango_20150905_001 | 31 | accuracy | 0.891304 |
| classification | lstm | Jango | single_day | Jango_20150905_001 | Jango_20150905_001 | 44 | accuracy | 0.978261 |
| classification | lstm | Jango | single_day | Jango_20150905_001 | Jango_20150905_001 | 44 | accuracy | 0.934783 |
| classification | lstm | Jango | single_day | Jango_20150905_001 | Jango_20150905_001 | 46 | accuracy | 0.913043 |
| classification | lstm | Jango | single_day | Jango_20150905_001 | Jango_20150905_001 | 46 | accuracy | 0.847826 |

| | | | | | | | | |
|---|---|---|---|---|---|---|---|---|
| classification | lstm | Jango | single_day | Jango_20150905_001 | Jango_20150905_001 | 54 | accuracy | 1 |
| classification | lstm | Jango | single_day | Jango_20150905_001 | Jango_20150905_001 | 54 | accuracy | 0.934783 |
| classification | lstm | Jango | single_day | Jango_20150905_001 | Jango_20150905_001 | 69 | accuracy | 0.956522 |
| classification | lstm | Jango | single_day | Jango_20150906_001 | Jango_20150906_001 | 31 | accuracy | 0.902439 |
| classification | lstm | Jango | single_day | Jango_20150906_001 | Jango_20150906_001 | 44 | accuracy | 0.902439 |
| classification | lstm | Jango | single_day | Jango_20150906_001 | Jango_20150906_001 | 44 | accuracy | 0.926829 |
| classification | lstm | Jango | single_day | Jango_20150906_001 | Jango_20150906_001 | 46 | accuracy | 0.95122 |
| classification | lstm | Jango | single_day | Jango_20150906_001 | Jango_20150906_001 | 46 | accuracy | 0.902439 |
| classification | lstm | Jango | single_day | Jango_20150906_001 | Jango_20150906_001 | 54 | accuracy | 0.926829 |
| classification | lstm | Jango | single_day | Jango_20150906_001 | Jango_20150906_001 | 54 | accuracy | 0.97561 |
| classification | lstm | Jango | single_day | Jango_20150906_001 | Jango_20150906_001 | 69 | accuracy | 0.902439 |
| classification | lstm | Jango | single_day | Jango_20150908_001 | Jango_20150908_001 | 31 | accuracy | 0.860465 |
| classification | lstm | Jango | single_day | Jango_20150908_001 | Jango_20150908_001 | 44 | accuracy | 0.860465 |
| classification | lstm | Jango | single_day | Jango_20150908_001 | Jango_20150908_001 | 46 | accuracy | 0.813953 |
| classification | lstm | Jango | single_day | Jango_20150908_001 | Jango_20150908_001 | 46 | accuracy | 0.953488 |
| classification | lstm | Jango | single_day | Jango_20150908_001 | Jango_20150908_001 | 54 | accuracy | 0.860465 |
| classification | lstm | Jango | single_day | Jango_20150908_001 | Jango_20150908_001 | 54 | accuracy | 0.976744 |
| classification | lstm | Jango | single_day | Jango_20150908_001 | Jango_20150908_001 | 69 | accuracy | 0.813953 |
| classification | lstm | Jango | single_day | Jango_20151029_001 | Jango_20151029_001 | 31 | accuracy | 0.977273 |
| classification | lstm | Jango | single_day | Jango_20151029_001 | Jango_20151029_001 | 44 | accuracy | 0.909091 |
| classification | lstm | Jango | single_day | Jango_20151029_001 | Jango_20151029_001 | 44 | accuracy | 0.954545 |
| classification | lstm | Jango | single_day | Jango_20151029_001 | Jango_20151029_001 | 46 | accuracy | 0.954545 |
| classification | lstm | Jango | single_day | Jango_20151029_001 | Jango_20151029_001 | 46 | accuracy | 0.977273 |
| classification | lstm | Jango | single_day | Jango_20151029_001 | Jango_20151029_001 | 54 | accuracy | 1 |
| classification | lstm | Jango | single_day | Jango_20151029_001 | Jango_20151029_001 | 54 | accuracy | 0.977273 |
| classification | lstm | Jango | single_day | Jango_20151029_001 | Jango_20151029_001 | 69 | accuracy | 0.931818 |
| classification | lstm | Jango | single_day | Jango_20151102_001 | Jango_20151102_001 | 31 | accuracy | 0.977273 |
| classification | lstm | Jango | single_day | Jango_20151102_001 | Jango_20151102_001 | 44 | accuracy | 0.931818 |
| classification | lstm | Jango | single_day | Jango_20151102_001 | Jango_20151102_001 | 44 | accuracy | 0.727273 |
| classification | lstm | Jango | single_day | Jango_20151102_001 | Jango_20151102_001 | 46 | accuracy | 0.909091 |
| classification | lstm | Jango | single_day | Jango_20151102_001 | Jango_20151102_001 | 46 | accuracy | 0.636364 |
| classification | lstm | Jango | single_day | Jango_20151102_001 | Jango_20151102_001 | 54 | accuracy | 0.954545 |
| classification | lstm | Jango | single_day | Jango_20151102_001 | Jango_20151102_001 | 54 | accuracy | 0.772727 |
| classification | lstm | Jango | single_day | Jango_20151102_001 | Jango_20151102_001 | 69 | accuracy | 0.863636 |
| classification | transformer | Jango | single_day | Jango_20150730_001 | Jango_20150730_001 | 31 | accuracy | 1 |
| classification | transformer | Jango | single_day | Jango_20150730_001 | Jango_20150730_001 | 44 | accuracy | 1 |
| classification | transformer | Jango | single_day | Jango_20150730_001 | Jango_20150730_001 | 44 | accuracy | 0.969697 |
| classification | transformer | Jango | single_day | Jango_20150730_001 | Jango_20150730_001 | 46 | accuracy | 0.969697 |
| classification | transformer | Jango | single_day | Jango_20150730_001 | Jango_20150730_001 | 46 | accuracy | 0.939394 |
| classification | transformer | Jango | single_day | Jango_20150730_001 | Jango_20150730_001 | 54 | accuracy | 0.969697 |
| classification | transformer | Jango | single_day | Jango_20150730_001 | Jango_20150730_001 | 54 | accuracy | 0.878788 |
| classification | transformer | Jango | single_day | Jango_20150730_001 | Jango_20150730_001 | 69 | accuracy | 1 |
| classification | transformer | Jango | single_day | Jango_20150731_001 | Jango_20150731_001 | 31 | accuracy | 0.975 |
| classification | transformer | Jango | single_day | Jango_20150731_001 | Jango_20150731_001 | 44 | accuracy | 0.975 |
| classification | transformer | Jango | single_day | Jango_20150731_001 | Jango_20150731_001 | 46 | accuracy | 0.925 |
| classification | transformer | Jango | single_day | Jango_20150731_001 | Jango_20150731_001 | 46 | accuracy | 0.9 |
| classification | transformer | Jango | single_day | Jango_20150731_001 | Jango_20150731_001 | 54 | accuracy | 0.95 |
| classification | transformer | Jango | single_day | Jango_20150731_001 | Jango_20150731_001 | 69 | accuracy | 1 |
| classification | transformer | Jango | single_day | Jango_20150801_001 | Jango_20150801_001 | 31 | accuracy | 0.977273 |
| classification | transformer | Jango | single_day | Jango_20150801_001 | Jango_20150801_001 | 44 | accuracy | 1 |
| classification | transformer | Jango | single_day | Jango_20150801_001 | Jango_20150801_001 | 44 | accuracy | 0.977273 |
| classification | transformer | Jango | single_day | Jango_20150801_001 | Jango_20150801_001 | 46 | accuracy | 0.954545 |
| classification | transformer | Jango | single_day | Jango_20150801_001 | Jango_20150801_001 | 54 | accuracy | 1 |
| classification | transformer | Jango | single_day | Jango_20150801_001 | Jango_20150801_001 | 54 | accuracy | 0.840909 |
| classification | transformer | Jango | single_day | Jango_20150801_001 | Jango_20150801_001 | 69 | accuracy | 1 |
| classification | transformer | Jango | single_day | Jango_20150805_001 | Jango_20150805_001 | 31 | accuracy | 0.977778 |
| classification | transformer | Jango | single_day | Jango_20150805_001 | Jango_20150805_001 | 44 | accuracy | 1 |
| classification | transformer | Jango | single_day | Jango_20150805_001 | Jango_20150805_001 | 44 | accuracy | 0.977778 |
| classification | transformer | Jango | single_day | Jango_20150805_001 | Jango_20150805_001 | 46 | accuracy | 0.977778 |
| classification | transformer | Jango | single_day | Jango_20150805_001 | Jango_20150805_001 | 54 | accuracy | 0.977778 |
| classification | transformer | Jango | single_day | Jango_20150805_001 | Jango_20150805_001 | 54 | accuracy | 0.911111 |
| classification | transformer | Jango | single_day | Jango_20150805_001 | Jango_20150805_001 | 69 | accuracy | 1 |
| classification | transformer | Jango | single_day | Jango_20150806_001 | Jango_20150806_001 | 31 | accuracy | 1 |

| | | | | | | | | |
|---|---|---|---|---|---|---|---|---|
| classification | transformer | Jango | single_day | Jango_20150806_001 | Jango_20150806_001 | 44 | accuracy | 1 |
| classification | transformer | Jango | single_day | Jango_20150806_001 | Jango_20150806_001 | 44 | accuracy | 0.688889 |
| classification | transformer | Jango | single_day | Jango_20150806_001 | Jango_20150806_001 | 46 | accuracy | 0.977778 |
| classification | transformer | Jango | single_day | Jango_20150806_001 | Jango_20150806_001 | 46 | accuracy | 0.933333 |
| classification | transformer | Jango | single_day | Jango_20150806_001 | Jango_20150806_001 | 54 | accuracy | 0.977778 |
| classification | transformer | Jango | single_day | Jango_20150806_001 | Jango_20150806_001 | 54 | accuracy | 0.844444 |
| classification | transformer | Jango | single_day | Jango_20150806_001 | Jango_20150806_001 | 69 | accuracy | 1 |
| classification | transformer | Jango | single_day | Jango_20150807_001 | Jango_20150807_001 | 31 | accuracy | 0.977778 |
| classification | transformer | Jango | single_day | Jango_20150807_001 | Jango_20150807_001 | 44 | accuracy | 1 |
| classification | transformer | Jango | single_day | Jango_20150807_001 | Jango_20150807_001 | 44 | accuracy | 0.933333 |
| classification | transformer | Jango | single_day | Jango_20150807_001 | Jango_20150807_001 | 46 | accuracy | 1 |
| classification | transformer | Jango | single_day | Jango_20150807_001 | Jango_20150807_001 | 46 | accuracy | 0.977778 |
| classification | transformer | Jango | single_day | Jango_20150807_001 | Jango_20150807_001 | 54 | accuracy | 1 |
| classification | transformer | Jango | single_day | Jango_20150807_001 | Jango_20150807_001 | 69 | accuracy | 1 |
| classification | transformer | Jango | single_day | Jango_20150808_001 | Jango_20150808_001 | 31 | accuracy | 0.977778 |
| classification | transformer | Jango | single_day | Jango_20150808_001 | Jango_20150808_001 | 44 | accuracy | 0.977778 |
| classification | transformer | Jango | single_day | Jango_20150808_001 | Jango_20150808_001 | 44 | accuracy | 0.933333 |
| classification | transformer | Jango | single_day | Jango_20150808_001 | Jango_20150808_001 | 46 | accuracy | 0.977778 |
| classification | transformer | Jango | single_day | Jango_20150808_001 | Jango_20150808_001 | 46 | accuracy | 0.955556 |
| classification | transformer | Jango | single_day | Jango_20150808_001 | Jango_20150808_001 | 54 | accuracy | 0.933333 |
| classification | transformer | Jango | single_day | Jango_20150808_001 | Jango_20150808_001 | 54 | accuracy | 0.866667 |
| classification | transformer | Jango | single_day | Jango_20150808_001 | Jango_20150808_001 | 69 | accuracy | 1 |
| classification | transformer | Jango | single_day | Jango_20150809_001 | Jango_20150809_001 | 31 | accuracy | 1 |
| classification | transformer | Jango | single_day | Jango_20150809_001 | Jango_20150809_001 | 44 | accuracy | 1 |
| classification | transformer | Jango | single_day | Jango_20150809_001 | Jango_20150809_001 | 44 | accuracy | 0.955556 |
| classification | transformer | Jango | single_day | Jango_20150809_001 | Jango_20150809_001 | 46 | accuracy | 1 |
| classification | transformer | Jango | single_day | Jango_20150809_001 | Jango_20150809_001 | 46 | accuracy | 0.955556 |
| classification | transformer | Jango | single_day | Jango_20150809_001 | Jango_20150809_001 | 54 | accuracy | 0.977778 |
| classification | transformer | Jango | single_day | Jango_20150809_001 | Jango_20150809_001 | 69 | accuracy | 0.977778 |
| classification | transformer | Jango | single_day | Jango_20150820_001 | Jango_20150820_001 | 31 | accuracy | 1 |
| classification | transformer | Jango | single_day | Jango_20150820_001 | Jango_20150820_001 | 44 | accuracy | 1 |
| classification | transformer | Jango | single_day | Jango_20150820_001 | Jango_20150820_001 | 44 | accuracy | 0.933333 |
| classification | transformer | Jango | single_day | Jango_20150820_001 | Jango_20150820_001 | 46 | accuracy | 1 |
| classification | transformer | Jango | single_day | Jango_20150820_001 | Jango_20150820_001 | 46 | accuracy | 0.911111 |
| classification | transformer | Jango | single_day | Jango_20150820_001 | Jango_20150820_001 | 54 | accuracy | 0.977778 |
| classification | transformer | Jango | single_day | Jango_20150820_001 | Jango_20150820_001 | 54 | accuracy | 0.955556 |
| classification | transformer | Jango | single_day | Jango_20150820_001 | Jango_20150820_001 | 69 | accuracy | 1 |
| classification | transformer | Jango | single_day | Jango_20150824_001 | Jango_20150824_001 | 31 | accuracy | 1 |
| classification | transformer | Jango | single_day | Jango_20150824_001 | Jango_20150824_001 | 44 | accuracy | 0.954545 |
| classification | transformer | Jango | single_day | Jango_20150824_001 | Jango_20150824_001 | 44 | accuracy | 0.931818 |
| classification | transformer | Jango | single_day | Jango_20150824_001 | Jango_20150824_001 | 46 | accuracy | 0.977273 |
| classification | transformer | Jango | single_day | Jango_20150824_001 | Jango_20150824_001 | 46 | accuracy | 0.954545 |
| classification | transformer | Jango | single_day | Jango_20150824_001 | Jango_20150824_001 | 54 | accuracy | 0.977273 |
| classification | transformer | Jango | single_day | Jango_20150824_001 | Jango_20150824_001 | 54 | accuracy | 0.863636 |
| classification | transformer | Jango | single_day | Jango_20150824_001 | Jango_20150824_001 | 69 | accuracy | 0.977273 |
| classification | transformer | Jango | single_day | Jango_20150825_001 | Jango_20150825_001 | 31 | accuracy | 1 |
| classification | transformer | Jango | single_day | Jango_20150825_001 | Jango_20150825_001 | 44 | accuracy | 1 |
| classification | transformer | Jango | single_day | Jango_20150825_001 | Jango_20150825_001 | 44 | accuracy | 0.891304 |
| classification | transformer | Jango | single_day | Jango_20150825_001 | Jango_20150825_001 | 46 | accuracy | 1 |
| classification | transformer | Jango | single_day | Jango_20150825_001 | Jango_20150825_001 | 46 | accuracy | 0.934783 |
| classification | transformer | Jango | single_day | Jango_20150825_001 | Jango_20150825_001 | 54 | accuracy | 1 |
| classification | transformer | Jango | single_day | Jango_20150825_001 | Jango_20150825_001 | 69 | accuracy | 1 |
| classification | transformer | Jango | single_day | Jango_20150826_001 | Jango_20150826_001 | 31 | accuracy | 1 |
| classification | transformer | Jango | single_day | Jango_20150826_001 | Jango_20150826_001 | 44 | accuracy | 1 |
| classification | transformer | Jango | single_day | Jango_20150826_001 | Jango_20150826_001 | 44 | accuracy | 0.933333 |
| classification | transformer | Jango | single_day | Jango_20150826_001 | Jango_20150826_001 | 46 | accuracy | 0.977778 |
| classification | transformer | Jango | single_day | Jango_20150826_001 | Jango_20150826_001 | 46 | accuracy | 0.844444 |
| classification | transformer | Jango | single_day | Jango_20150826_001 | Jango_20150826_001 | 54 | accuracy | 1 |
| classification | transformer | Jango | single_day | Jango_20150826_001 | Jango_20150826_001 | 69 | accuracy | 1 |
| classification | transformer | Jango | single_day | Jango_20150827_001 | Jango_20150827_001 | 31 | accuracy | 1 |
| classification | transformer | Jango | single_day | Jango_20150827_001 | Jango_20150827_001 | 44 | accuracy | 0.978261 |
| classification | transformer | Jango | single_day | Jango_20150827_001 | Jango_20150827_001 | 44 | accuracy | 0.956522 |
| classification | transformer | Jango | single_day | Jango_20150827_001 | Jango_20150827_001 | 46 | accuracy | 1 |

| | | | | | | | | |
|---|---|---|---|---|---|---|---|---|
| classification | transformer | Jango | single_day | Jango_20150827_001 | Jango_20150827_001 | 54 | accuracy | 1 |
| classification | transformer | Jango | single_day | Jango_20150827_001 | Jango_20150827_001 | 54 | accuracy | 0.956522 |
| classification | transformer | Jango | single_day | Jango_20150827_001 | Jango_20150827_001 | 69 | accuracy | 1 |
| classification | transformer | Jango | single_day | Jango_20150828_001 | Jango_20150828_001 | 31 | accuracy | 1 |
| classification | transformer | Jango | single_day | Jango_20150828_001 | Jango_20150828_001 | 44 | accuracy | 1 |
| classification | transformer | Jango | single_day | Jango_20150828_001 | Jango_20150828_001 | 44 | accuracy | 0.953488 |
| classification | transformer | Jango | single_day | Jango_20150828_001 | Jango_20150828_001 | 46 | accuracy | 0.953488 |
| classification | transformer | Jango | single_day | Jango_20150828_001 | Jango_20150828_001 | 46 | accuracy | 0.930233 |
| classification | transformer | Jango | single_day | Jango_20150828_001 | Jango_20150828_001 | 54 | accuracy | 1 |
| classification | transformer | Jango | single_day | Jango_20150828_001 | Jango_20150828_001 | 54 | accuracy | 0.953488 |
| classification | transformer | Jango | single_day | Jango_20150828_001 | Jango_20150828_001 | 69 | accuracy | 1 |
| classification | transformer | Jango | single_day | Jango_20150831_001 | Jango_20150831_001 | 31 | accuracy | 0.972973 |
| classification | transformer | Jango | single_day | Jango_20150831_001 | Jango_20150831_001 | 44 | accuracy | 0.972973 |
| classification | transformer | Jango | single_day | Jango_20150831_001 | Jango_20150831_001 | 44 | accuracy | 0.918919 |
| classification | transformer | Jango | single_day | Jango_20150831_001 | Jango_20150831_001 | 46 | accuracy | 1 |
| classification | transformer | Jango | single_day | Jango_20150831_001 | Jango_20150831_001 | 46 | accuracy | 0.945946 |
| classification | transformer | Jango | single_day | Jango_20150831_001 | Jango_20150831_001 | 54 | accuracy | 1 |
| classification | transformer | Jango | single_day | Jango_20150831_001 | Jango_20150831_001 | 54 | accuracy | 0.864865 |
| classification | transformer | Jango | single_day | Jango_20150831_001 | Jango_20150831_001 | 69 | accuracy | 0.972973 |
| classification | transformer | Jango | single_day | Jango_20150905_001 | Jango_20150905_001 | 31 | accuracy | 0.978261 |
| classification | transformer | Jango | single_day | Jango_20150905_001 | Jango_20150905_001 | 44 | accuracy | 1 |
| classification | transformer | Jango | single_day | Jango_20150905_001 | Jango_20150905_001 | 44 | accuracy | 0.913043 |
| classification | transformer | Jango | single_day | Jango_20150905_001 | Jango_20150905_001 | 46 | accuracy | 0.978261 |
| classification | transformer | Jango | single_day | Jango_20150905_001 | Jango_20150905_001 | 46 | accuracy | 0.891304 |
| classification | transformer | Jango | single_day | Jango_20150905_001 | Jango_20150905_001 | 54 | accuracy | 0.934783 |
| classification | transformer | Jango | single_day | Jango_20150905_001 | Jango_20150905_001 | 69 | accuracy | 0.978261 |
| classification | transformer | Jango | single_day | Jango_20150906_001 | Jango_20150906_001 | 31 | accuracy | 0.926829 |
| classification | transformer | Jango | single_day | Jango_20150906_001 | Jango_20150906_001 | 44 | accuracy | 1 |
| classification | transformer | Jango | single_day | Jango_20150906_001 | Jango_20150906_001 | 44 | accuracy | 0.926829 |
| classification | transformer | Jango | single_day | Jango_20150906_001 | Jango_20150906_001 | 46 | accuracy | 0.95122 |
| classification | transformer | Jango | single_day | Jango_20150906_001 | Jango_20150906_001 | 54 | accuracy | 1 |
| classification | transformer | Jango | single_day | Jango_20150906_001 | Jango_20150906_001 | 54 | accuracy | 0.878049 |
| classification | transformer | Jango | single_day | Jango_20150906_001 | Jango_20150906_001 | 69 | accuracy | 0.95122 |
| classification | transformer | Jango | single_day | Jango_20150908_001 | Jango_20150908_001 | 31 | accuracy | 0.953488 |
| classification | transformer | Jango | single_day | Jango_20150908_001 | Jango_20150908_001 | 44 | accuracy | 0.953488 |
| classification | transformer | Jango | single_day | Jango_20150908_001 | Jango_20150908_001 | 44 | accuracy | 0.906977 |
| classification | transformer | Jango | single_day | Jango_20150908_001 | Jango_20150908_001 | 46 | accuracy | 0.953488 |
| classification | transformer | Jango | single_day | Jango_20150908_001 | Jango_20150908_001 | 54 | accuracy | 0.976744 |
| classification | transformer | Jango | single_day | Jango_20150908_001 | Jango_20150908_001 | 54 | accuracy | 0.837209 |
| classification | transformer | Jango | single_day | Jango_20150908_001 | Jango_20150908_001 | 69 | accuracy | 1 |
| classification | transformer | Jango | single_day | Jango_20151029_001 | Jango_20151029_001 | 31 | accuracy | 1 |
| classification | transformer | Jango | single_day | Jango_20151029_001 | Jango_20151029_001 | 44 | accuracy | 1 |
| classification | transformer | Jango | single_day | Jango_20151029_001 | Jango_20151029_001 | 44 | accuracy | 0.977273 |
| classification | transformer | Jango | single_day | Jango_20151029_001 | Jango_20151029_001 | 46 | accuracy | 0.977273 |
| classification | transformer | Jango | single_day | Jango_20151029_001 | Jango_20151029_001 | 46 | accuracy | 0.931818 |
| classification | transformer | Jango | single_day | Jango_20151029_001 | Jango_20151029_001 | 54 | accuracy | 1 |
| classification | transformer | Jango | single_day | Jango_20151029_001 | Jango_20151029_001 | 54 | accuracy | 0.931818 |
| classification | transformer | Jango | single_day | Jango_20151029_001 | Jango_20151029_001 | 69 | accuracy | 0.977273 |
| classification | transformer | Jango | single_day | Jango_20151102_001 | Jango_20151102_001 | 31 | accuracy | 1 |
| classification | transformer | Jango | single_day | Jango_20151102_001 | Jango_20151102_001 | 44 | accuracy | 0.977273 |
| classification | transformer | Jango | single_day | Jango_20151102_001 | Jango_20151102_001 | 46 | accuracy | 1 |
| classification | transformer | Jango | single_day | Jango_20151102_001 | Jango_20151102_001 | 54 | accuracy | 0.977273 |
| classification | transformer | Jango | single_day | Jango_20151102_001 | Jango_20151102_001 | 54 | accuracy | 0.954545 |
| classification | transformer | Jango | single_day | Jango_20151102_001 | Jango_20151102_001 | 69 | accuracy | 0.977273 |
| classification | lfads | Jango | single_day | Jango_20150730_001 | Jango_20150730_001 | 44 | accuracy | 1 |
| classification | lfads | Jango | single_day | Jango_20150730_001 | Jango_20150730_001 | 46 | accuracy | 0.969697 |
| classification | lfads | Jango | single_day | Jango_20150730_001 | Jango_20150730_001 | 54 | accuracy | 0.848485 |
| classification | lfads | Jango | single_day | Jango_20150731_001 | Jango_20150731_001 | 44 | accuracy | 0.95 |
| classification | lfads | Jango | single_day | Jango_20150731_001 | Jango_20150731_001 | 46 | accuracy | 0.975 |
| classification | lfads | Jango | single_day | Jango_20150731_001 | Jango_20150731_001 | 54 | accuracy | 0.975 |
| classification | lfads | Jango | single_day | Jango_20150801_001 | Jango_20150801_001 | 44 | accuracy | 0.977273 |
| classification | lfads | Jango | single_day | Jango_20150801_001 | Jango_20150801_001 | 46 | accuracy | 0.954545 |
| classification | lfads | Jango | single_day | Jango_20150801_001 | Jango_20150801_001 | 54 | accuracy | 0.931818 |

| | | | | | | | | |
|---|---|---|---|---|---|---|---|---|
| classification | lfads | Jango | single_day | Jango_20150805_001 | Jango_20150805_001 | 44 | accuracy | 0.977778 |
| classification | lfads | Jango | single_day | Jango_20150805_001 | Jango_20150805_001 | 46 | accuracy | 0.933333 |
| classification | lfads | Jango | single_day | Jango_20150805_001 | Jango_20150805_001 | 54 | accuracy | 0.955556 |
| classification | lfads | Jango | single_day | Jango_20150806_001 | Jango_20150806_001 | 44 | accuracy | 0.955556 |
| classification | lfads | Jango | single_day | Jango_20150806_001 | Jango_20150806_001 | 46 | accuracy | 0.977778 |
| classification | lfads | Jango | single_day | Jango_20150806_001 | Jango_20150806_001 | 54 | accuracy | 1 |
| classification | lfads | Jango | single_day | Jango_20150807_001 | Jango_20150807_001 | 44 | accuracy | 1 |
| classification | lfads | Jango | single_day | Jango_20150807_001 | Jango_20150807_001 | 46 | accuracy | 1 |
| classification | lfads | Jango | single_day | Jango_20150807_001 | Jango_20150807_001 | 54 | accuracy | 1 |
| classification | lfads | Jango | single_day | Jango_20150808_001 | Jango_20150808_001 | 44 | accuracy | 0.866667 |
| classification | lfads | Jango | single_day | Jango_20150808_001 | Jango_20150808_001 | 46 | accuracy | 0.977778 |
| classification | lfads | Jango | single_day | Jango_20150808_001 | Jango_20150808_001 | 54 | accuracy | 0.933333 |
| classification | lfads | Jango | single_day | Jango_20150809_001 | Jango_20150809_001 | 44 | accuracy | 1 |
| classification | lfads | Jango | single_day | Jango_20150809_001 | Jango_20150809_001 | 46 | accuracy | 0.955556 |
| classification | lfads | Jango | single_day | Jango_20150809_001 | Jango_20150809_001 | 54 | accuracy | 0.977778 |
| classification | lfads | Jango | single_day | Jango_20150820_001 | Jango_20150820_001 | 44 | accuracy | 0.977778 |
| classification | lfads | Jango | single_day | Jango_20150820_001 | Jango_20150820_001 | 46 | accuracy | 0.955556 |
| classification | lfads | Jango | single_day | Jango_20150820_001 | Jango_20150820_001 | 54 | accuracy | 0.955556 |
| classification | lfads | Jango | single_day | Jango_20150824_001 | Jango_20150824_001 | 44 | accuracy | 0.977273 |
| classification | lfads | Jango | single_day | Jango_20150824_001 | Jango_20150824_001 | 46 | accuracy | 0.931818 |
| classification | lfads | Jango | single_day | Jango_20150824_001 | Jango_20150824_001 | 54 | accuracy | 0.954545 |
| classification | lfads | Jango | single_day | Jango_20150825_001 | Jango_20150825_001 | 44 | accuracy | 0.934783 |
| classification | lfads | Jango | single_day | Jango_20150825_001 | Jango_20150825_001 | 46 | accuracy | 0.956522 |
| classification | lfads | Jango | single_day | Jango_20150825_001 | Jango_20150825_001 | 54 | accuracy | 1 |
| classification | lfads | Jango | single_day | Jango_20150826_001 | Jango_20150826_001 | 44 | accuracy | 0.977778 |
| classification | lfads | Jango | single_day | Jango_20150826_001 | Jango_20150826_001 | 46 | accuracy | 0.955556 |
| classification | lfads | Jango | single_day | Jango_20150826_001 | Jango_20150826_001 | 54 | accuracy | 1 |
| classification | lfads | Jango | single_day | Jango_20150827_001 | Jango_20150827_001 | 44 | accuracy | 0.978261 |
| classification | lfads | Jango | single_day | Jango_20150827_001 | Jango_20150827_001 | 46 | accuracy | 0.978261 |
| classification | lfads | Jango | single_day | Jango_20150827_001 | Jango_20150827_001 | 54 | accuracy | 1 |
| classification | lfads | Jango | single_day | Jango_20150828_001 | Jango_20150828_001 | 44 | accuracy | 0.976744 |
| classification | lfads | Jango | single_day | Jango_20150828_001 | Jango_20150828_001 | 46 | accuracy | 0.953488 |
| classification | lfads | Jango | single_day | Jango_20150828_001 | Jango_20150828_001 | 54 | accuracy | 1 |
| classification | lfads | Jango | single_day | Jango_20150831_001 | Jango_20150831_001 | 44 | accuracy | 1 |
| classification | lfads | Jango | single_day | Jango_20150831_001 | Jango_20150831_001 | 46 | accuracy | 0.945946 |
| classification | lfads | Jango | single_day | Jango_20150831_001 | Jango_20150831_001 | 54 | accuracy | 0.972973 |
| classification | lfads | Jango | single_day | Jango_20150905_001 | Jango_20150905_001 | 44 | accuracy | 0.934783 |
| classification | lfads | Jango | single_day | Jango_20150905_001 | Jango_20150905_001 | 46 | accuracy | 0.934783 |
| classification | lfads | Jango | single_day | Jango_20150905_001 | Jango_20150905_001 | 54 | accuracy | 0.978261 |
| classification | lfads | Jango | single_day | Jango_20150906_001 | Jango_20150906_001 | 44 | accuracy | 1 |
| classification | lfads | Jango | single_day | Jango_20150906_001 | Jango_20150906_001 | 46 | accuracy | 0.926829 |
| classification | lfads | Jango | single_day | Jango_20150906_001 | Jango_20150906_001 | 54 | accuracy | 0.926829 |
| classification | lfads | Jango | single_day | Jango_20150908_001 | Jango_20150908_001 | 44 | accuracy | 0.837209 |
| classification | lfads | Jango | single_day | Jango_20150908_001 | Jango_20150908_001 | 46 | accuracy | 0.976744 |
| classification | lfads | Jango | single_day | Jango_20150908_001 | Jango_20150908_001 | 54 | accuracy | 0.930233 |
| classification | lfads | Jango | single_day | Jango_20151029_001 | Jango_20151029_001 | 44 | accuracy | 0.977273 |
| classification | lfads | Jango | single_day | Jango_20151029_001 | Jango_20151029_001 | 46 | accuracy | 1 |
| classification | lfads | Jango | single_day | Jango_20151029_001 | Jango_20151029_001 | 54 | accuracy | 0.977273 |
| classification | lfads | Jango | single_day | Jango_20151102_001 | Jango_20151102_001 | 44 | accuracy | 0.931818 |
| classification | lfads | Jango | single_day | Jango_20151102_001 | Jango_20151102_001 | 46 | accuracy | 0.954545 |
| classification | lfads | Jango | single_day | Jango_20151102_001 | Jango_20151102_001 | 54 | accuracy | 0.909091 |
| classification | cycle_gan | Jango | single_day | Jango_20150730_001 | Jango_20150730_001 | 31 | accuracy | 0.757576 |
| classification | cycle_gan | Jango | single_day | Jango_20150730_001 | Jango_20150730_001 | 44 | accuracy | 0.969697 |
| classification | cycle_gan | Jango | single_day | Jango_20150730_001 | Jango_20150730_001 | 46 | accuracy | 0.909091 |
| classification | cycle_gan | Jango | single_day | Jango_20150730_001 | Jango_20150730_001 | 54 | accuracy | 0.939394 |
| classification | cycle_gan | Jango | single_day | Jango_20150730_001 | Jango_20150730_001 | 69 | accuracy | 1 |
| classification | cycle_gan | Jango | single_day | Jango_20150731_001 | Jango_20150731_001 | 31 | accuracy | 0.975 |
| classification | cycle_gan | Jango | single_day | Jango_20150731_001 | Jango_20150731_001 | 44 | accuracy | 0.95 |
| classification | cycle_gan | Jango | single_day | Jango_20150731_001 | Jango_20150731_001 | 46 | accuracy | 0.975 |
| classification | cycle_gan | Jango | single_day | Jango_20150731_001 | Jango_20150731_001 | 54 | accuracy | 0.95 |
| classification | cycle_gan | Jango | single_day | Jango_20150731_001 | Jango_20150731_001 | 69 | accuracy | 0.975 |
| classification | cycle_gan | Jango | single_day | Jango_20150801_001 | Jango_20150801_001 | 31 | accuracy | 0.954545 |
| classification | cycle_gan | Jango | single_day | Jango_20150801_001 | Jango_20150801_001 | 44 | accuracy | 0.954545 |

| | | | | | | | | |
|---|---|---|---|---|---|---|---|---|
| classification | cycle_gan | Jango | single_day | Jango_20150801_001 | Jango_20150801_001 | 46 | accuracy | 0.954545 |
| classification | cycle_gan | Jango | single_day | Jango_20150801_001 | Jango_20150801_001 | 54 | accuracy | 0.909091 |
| classification | cycle_gan | Jango | single_day | Jango_20150801_001 | Jango_20150801_001 | 69 | accuracy | 0.977273 |
| classification | cycle_gan | Jango | single_day | Jango_20150805_001 | Jango_20150805_001 | 31 | accuracy | 0.955556 |
| classification | cycle_gan | Jango | single_day | Jango_20150805_001 | Jango_20150805_001 | 44 | accuracy | 0.955556 |
| classification | cycle_gan | Jango | single_day | Jango_20150805_001 | Jango_20150805_001 | 46 | accuracy | 1 |
| classification | cycle_gan | Jango | single_day | Jango_20150805_001 | Jango_20150805_001 | 54 | accuracy | 0.911111 |
| classification | cycle_gan | Jango | single_day | Jango_20150805_001 | Jango_20150805_001 | 69 | accuracy | 0.866667 |
| classification | cycle_gan | Jango | single_day | Jango_20150806_001 | Jango_20150806_001 | 31 | accuracy | 0.977778 |
| classification | cycle_gan | Jango | single_day | Jango_20150806_001 | Jango_20150806_001 | 44 | accuracy | 1 |
| classification | cycle_gan | Jango | single_day | Jango_20150806_001 | Jango_20150806_001 | 46 | accuracy | 1 |
| classification | cycle_gan | Jango | single_day | Jango_20150806_001 | Jango_20150806_001 | 54 | accuracy | 0.933333 |
| classification | cycle_gan | Jango | single_day | Jango_20150806_001 | Jango_20150806_001 | 69 | accuracy | 0.977778 |
| classification | cycle_gan | Jango | single_day | Jango_20150807_001 | Jango_20150807_001 | 31 | accuracy | 1 |
| classification | cycle_gan | Jango | single_day | Jango_20150807_001 | Jango_20150807_001 | 44 | accuracy | 1 |
| classification | cycle_gan | Jango | single_day | Jango_20150807_001 | Jango_20150807_001 | 46 | accuracy | 1 |
| classification | cycle_gan | Jango | single_day | Jango_20150807_001 | Jango_20150807_001 | 54 | accuracy | 0.955556 |
| classification | cycle_gan | Jango | single_day | Jango_20150807_001 | Jango_20150807_001 | 69 | accuracy | 0.955556 |
| classification | cycle_gan | Jango | single_day | Jango_20150808_001 | Jango_20150808_001 | 31 | accuracy | 0.977778 |
| classification | cycle_gan | Jango | single_day | Jango_20150808_001 | Jango_20150808_001 | 44 | accuracy | 0.933333 |
| classification | cycle_gan | Jango | single_day | Jango_20150808_001 | Jango_20150808_001 | 46 | accuracy | 0.933333 |
| classification | cycle_gan | Jango | single_day | Jango_20150808_001 | Jango_20150808_001 | 54 | accuracy | 0.911111 |
| classification | cycle_gan | Jango | single_day | Jango_20150808_001 | Jango_20150808_001 | 69 | accuracy | 0.955556 |
| classification | cycle_gan | Jango | single_day | Jango_20150809_001 | Jango_20150809_001 | 31 | accuracy | 1 |
| classification | cycle_gan | Jango | single_day | Jango_20150809_001 | Jango_20150809_001 | 44 | accuracy | 1 |
| classification | cycle_gan | Jango | single_day | Jango_20150809_001 | Jango_20150809_001 | 46 | accuracy | 1 |
| classification | cycle_gan | Jango | single_day | Jango_20150809_001 | Jango_20150809_001 | 54 | accuracy | 1 |
| classification | cycle_gan | Jango | single_day | Jango_20150809_001 | Jango_20150809_001 | 69 | accuracy | 0.933333 |
| classification | cycle_gan | Jango | single_day | Jango_20150820_001 | Jango_20150820_001 | 31 | accuracy | 0.955556 |
| classification | cycle_gan | Jango | single_day | Jango_20150820_001 | Jango_20150820_001 | 44 | accuracy | 0.911111 |
| classification | cycle_gan | Jango | single_day | Jango_20150820_001 | Jango_20150820_001 | 46 | accuracy | 0.977778 |
| classification | cycle_gan | Jango | single_day | Jango_20150820_001 | Jango_20150820_001 | 54 | accuracy | 0.822222 |
| classification | cycle_gan | Jango | single_day | Jango_20150820_001 | Jango_20150820_001 | 69 | accuracy | 0.955556 |
| classification | cycle_gan | Jango | single_day | Jango_20150824_001 | Jango_20150824_001 | 31 | accuracy | 0.954545 |
| classification | cycle_gan | Jango | single_day | Jango_20150824_001 | Jango_20150824_001 | 44 | accuracy | 0.954545 |
| classification | cycle_gan | Jango | single_day | Jango_20150824_001 | Jango_20150824_001 | 46 | accuracy | 0.954545 |
| classification | cycle_gan | Jango | single_day | Jango_20150824_001 | Jango_20150824_001 | 54 | accuracy | 0.909091 |
| classification | cycle_gan | Jango | single_day | Jango_20150824_001 | Jango_20150824_001 | 69 | accuracy | 0.931818 |
| classification | cycle_gan | Jango | single_day | Jango_20150825_001 | Jango_20150825_001 | 31 | accuracy | 1 |
| classification | cycle_gan | Jango | single_day | Jango_20150825_001 | Jango_20150825_001 | 44 | accuracy | 0.847826 |
| classification | cycle_gan | Jango | single_day | Jango_20150825_001 | Jango_20150825_001 | 46 | accuracy | 0.913043 |
| classification | cycle_gan | Jango | single_day | Jango_20150825_001 | Jango_20150825_001 | 54 | accuracy | 0.847826 |
| classification | cycle_gan | Jango | single_day | Jango_20150825_001 | Jango_20150825_001 | 69 | accuracy | 0.934783 |
| classification | cycle_gan | Jango | single_day | Jango_20150826_001 | Jango_20150826_001 | 31 | accuracy | 0.888889 |
| classification | cycle_gan | Jango | single_day | Jango_20150826_001 | Jango_20150826_001 | 44 | accuracy | 0.933333 |
| classification | cycle_gan | Jango | single_day | Jango_20150826_001 | Jango_20150826_001 | 46 | accuracy | 0.955556 |
| classification | cycle_gan | Jango | single_day | Jango_20150826_001 | Jango_20150826_001 | 54 | accuracy | 1 |
| classification | cycle_gan | Jango | single_day | Jango_20150826_001 | Jango_20150826_001 | 69 | accuracy | 1 |
| classification | cycle_gan | Jango | single_day | Jango_20150827_001 | Jango_20150827_001 | 31 | accuracy | 0.978261 |
| classification | cycle_gan | Jango | single_day | Jango_20150827_001 | Jango_20150827_001 | 44 | accuracy | 0.891304 |
| classification | cycle_gan | Jango | single_day | Jango_20150827_001 | Jango_20150827_001 | 46 | accuracy | 0.956522 |
| classification | cycle_gan | Jango | single_day | Jango_20150827_001 | Jango_20150827_001 | 54 | accuracy | 0.956522 |
| classification | cycle_gan | Jango | single_day | Jango_20150827_001 | Jango_20150827_001 | 69 | accuracy | 0.956522 |
| classification | cycle_gan | Jango | single_day | Jango_20150828_001 | Jango_20150828_001 | 31 | accuracy | 0.953488 |
| classification | cycle_gan | Jango | single_day | Jango_20150828_001 | Jango_20150828_001 | 44 | accuracy | 0.976744 |
| classification | cycle_gan | Jango | single_day | Jango_20150828_001 | Jango_20150828_001 | 46 | accuracy | 0.953488 |
| classification | cycle_gan | Jango | single_day | Jango_20150828_001 | Jango_20150828_001 | 54 | accuracy | 0.976744 |
| classification | cycle_gan | Jango | single_day | Jango_20150828_001 | Jango_20150828_001 | 69 | accuracy | 1 |
| classification | cycle_gan | Jango | single_day | Jango_20150831_001 | Jango_20150831_001 | 31 | accuracy | 0.864865 |
| classification | cycle_gan | Jango | single_day | Jango_20150831_001 | Jango_20150831_001 | 44 | accuracy | 1 |
| classification | cycle_gan | Jango | single_day | Jango_20150831_001 | Jango_20150831_001 | 46 | accuracy | 1 |
| classification | cycle_gan | Jango | single_day | Jango_20150831_001 | Jango_20150831_001 | 54 | accuracy | 0.945946 |
| classification | cycle_gan | Jango | single_day | Jango_20150831_001 | Jango_20150831_001 | 69 | accuracy | 1 |

| | | | | | | | | |
|---|---|---|---|---|---|---|---|---|
| classification | cycle_gan | Jango | single_day | Jango_20150905_001 | Jango_20150905_001 | 31 | accuracy | 0.934783 |
| classification | cycle_gan | Jango | single_day | Jango_20150905_001 | Jango_20150905_001 | 44 | accuracy | 0.934783 |
| classification | cycle_gan | Jango | single_day | Jango_20150905_001 | Jango_20150905_001 | 46 | accuracy | 0.956522 |
| classification | cycle_gan | Jango | single_day | Jango_20150905_001 | Jango_20150905_001 | 54 | accuracy | 0.934783 |
| classification | cycle_gan | Jango | single_day | Jango_20150905_001 | Jango_20150905_001 | 69 | accuracy | 0.978261 |
| classification | cycle_gan | Jango | single_day | Jango_20150906_001 | Jango_20150906_001 | 31 | accuracy | 0.97561 |
| classification | cycle_gan | Jango | single_day | Jango_20150906_001 | Jango_20150906_001 | 44 | accuracy | 0.97561 |
| classification | cycle_gan | Jango | single_day | Jango_20150906_001 | Jango_20150906_001 | 46 | accuracy | 0.829268 |
| classification | cycle_gan | Jango | single_day | Jango_20150906_001 | Jango_20150906_001 | 54 | accuracy | 0.97561 |
| classification | cycle_gan | Jango | single_day | Jango_20150906_001 | Jango_20150906_001 | 69 | accuracy | 0.926829 |
| classification | cycle_gan | Jango | single_day | Jango_20150908_001 | Jango_20150908_001 | 31 | accuracy | 0.627907 |
| classification | cycle_gan | Jango | single_day | Jango_20150908_001 | Jango_20150908_001 | 44 | accuracy | 0.72093 |
| classification | cycle_gan | Jango | single_day | Jango_20150908_001 | Jango_20150908_001 | 46 | accuracy | 0.697674 |
| classification | cycle_gan | Jango | single_day | Jango_20150908_001 | Jango_20150908_001 | 54 | accuracy | 0.837209 |
| classification | cycle_gan | Jango | single_day | Jango_20150908_001 | Jango_20150908_001 | 69 | accuracy | 0.813953 |
| classification | cycle_gan | Jango | single_day | Jango_20151029_001 | Jango_20151029_001 | 31 | accuracy | 0.977273 |
| classification | cycle_gan | Jango | single_day | Jango_20151029_001 | Jango_20151029_001 | 44 | accuracy | 0.954545 |
| classification | cycle_gan | Jango | single_day | Jango_20151029_001 | Jango_20151029_001 | 46 | accuracy | 0.977273 |
| classification | cycle_gan | Jango | single_day | Jango_20151029_001 | Jango_20151029_001 | 54 | accuracy | 0.886364 |
| classification | cycle_gan | Jango | single_day | Jango_20151029_001 | Jango_20151029_001 | 69 | accuracy | 0.909091 |
| classification | cycle_gan | Jango | single_day | Jango_20151102_001 | Jango_20151102_001 | 31 | accuracy | 1 |
| classification | cycle_gan | Jango | single_day | Jango_20151102_001 | Jango_20151102_001 | 44 | accuracy | 0.909091 |
| classification | cycle_gan | Jango | single_day | Jango_20151102_001 | Jango_20151102_001 | 46 | accuracy | 0.909091 |
| classification | cycle_gan | Jango | single_day | Jango_20151102_001 | Jango_20151102_001 | 54 | accuracy | 0.909091 |
| classification | cycle_gan | Jango | single_day | Jango_20151102_001 | Jango_20151102_001 | 69 | accuracy | 0.977273 |
| classification | stabilization | Jango | single_day | Jango_20150730_001 | Jango_20150730_001 | 44 | accuracy | 0.785714 |
| classification | stabilization | Jango | single_day | Jango_20150730_001 | Jango_20150730_001 | 46 | accuracy | 0.678571 |
| classification | stabilization | Jango | single_day | Jango_20150730_001 | Jango_20150730_001 | 54 | accuracy | 0.928571 |
| classification | stabilization | Jango | single_day | Jango_20150731_001 | Jango_20150731_001 | 44 | accuracy | 0.885714 |
| classification | stabilization | Jango | single_day | Jango_20150731_001 | Jango_20150731_001 | 46 | accuracy | 0.914286 |
| classification | stabilization | Jango | single_day | Jango_20150731_001 | Jango_20150731_001 | 54 | accuracy | 0.771429 |
| classification | stabilization | Jango | single_day | Jango_20150801_001 | Jango_20150801_001 | 44 | accuracy | 0.74359 |
| classification | stabilization | Jango | single_day | Jango_20150801_001 | Jango_20150801_001 | 46 | accuracy | 0.717949 |
| classification | stabilization | Jango | single_day | Jango_20150801_001 | Jango_20150801_001 | 54 | accuracy | 0.769231 |
| classification | stabilization | Jango | single_day | Jango_20150805_001 | Jango_20150805_001 | 44 | accuracy | 0.575 |
| classification | stabilization | Jango | single_day | Jango_20150805_001 | Jango_20150805_001 | 46 | accuracy | 0.75 |
| classification | stabilization | Jango | single_day | Jango_20150805_001 | Jango_20150805_001 | 54 | accuracy | 0.8 |
| classification | stabilization | Jango | single_day | Jango_20150806_001 | Jango_20150806_001 | 44 | accuracy | 0.475 |
| classification | stabilization | Jango | single_day | Jango_20150806_001 | Jango_20150806_001 | 46 | accuracy | 1 |
| classification | stabilization | Jango | single_day | Jango_20150806_001 | Jango_20150806_001 | 54 | accuracy | 0.85 |
| classification | stabilization | Jango | single_day | Jango_20150807_001 | Jango_20150807_001 | 44 | accuracy | 0.625 |
| classification | stabilization | Jango | single_day | Jango_20150807_001 | Jango_20150807_001 | 46 | accuracy | 1 |
| classification | stabilization | Jango | single_day | Jango_20150807_001 | Jango_20150807_001 | 54 | accuracy | 0.975 |
| classification | stabilization | Jango | single_day | Jango_20150808_001 | Jango_20150808_001 | 44 | accuracy | 0.625 |
| classification | stabilization | Jango | single_day | Jango_20150808_001 | Jango_20150808_001 | 46 | accuracy | 0.75 |
| classification | stabilization | Jango | single_day | Jango_20150808_001 | Jango_20150808_001 | 54 | accuracy | 0.675 |
| classification | stabilization | Jango | single_day | Jango_20150809_001 | Jango_20150809_001 | 44 | accuracy | 0.7 |
| classification | stabilization | Jango | single_day | Jango_20150809_001 | Jango_20150809_001 | 46 | accuracy | 0.825 |
| classification | stabilization | Jango | single_day | Jango_20150809_001 | Jango_20150809_001 | 54 | accuracy | 0.9 |
| classification | stabilization | Jango | single_day | Jango_20150820_001 | Jango_20150820_001 | 44 | accuracy | 0.4 |
| classification | stabilization | Jango | single_day | Jango_20150820_001 | Jango_20150820_001 | 46 | accuracy | 0.825 |
| classification | stabilization | Jango | single_day | Jango_20150820_001 | Jango_20150820_001 | 54 | accuracy | 0.925 |
| classification | stabilization | Jango | single_day | Jango_20150824_001 | Jango_20150824_001 | 44 | accuracy | 0.717949 |
| classification | stabilization | Jango | single_day | Jango_20150824_001 | Jango_20150824_001 | 46 | accuracy | 0.769231 |
| classification | stabilization | Jango | single_day | Jango_20150824_001 | Jango_20150824_001 | 54 | accuracy | 0.846154 |
| classification | stabilization | Jango | single_day | Jango_20150825_001 | Jango_20150825_001 | 44 | accuracy | 0.634146 |
| classification | stabilization | Jango | single_day | Jango_20150825_001 | Jango_20150825_001 | 46 | accuracy | 0.658537 |
| classification | stabilization | Jango | single_day | Jango_20150825_001 | Jango_20150825_001 | 54 | accuracy | 0.609756 |
| classification | stabilization | Jango | single_day | Jango_20150826_001 | Jango_20150826_001 | 44 | accuracy | 0.4 |
| classification | stabilization | Jango | single_day | Jango_20150826_001 | Jango_20150826_001 | 46 | accuracy | 0.9 |
| classification | stabilization | Jango | single_day | Jango_20150826_001 | Jango_20150826_001 | 54 | accuracy | 0.85 |
| classification | stabilization | Jango | single_day | Jango_20150827_001 | Jango_20150827_001 | 44 | accuracy | 0.512195 |
| classification | stabilization | Jango | single_day | Jango_20150827_001 | Jango_20150827_001 | 46 | accuracy | 0.95122 |

| | | | | | | | | |
|---|---|---|---|---|---|---|---|---|
| classification | stabilization | Jango | single_day | Jango_20150827_001 | Jango_20150827_001 | 54 | accuracy | 0.878049 |
| classification | stabilization | Jango | single_day | Jango_20150828_001 | Jango_20150828_001 | 44 | accuracy | 0.421053 |
| classification | stabilization | Jango | single_day | Jango_20150828_001 | Jango_20150828_001 | 46 | accuracy | 0.710526 |
| classification | stabilization | Jango | single_day | Jango_20150828_001 | Jango_20150828_001 | 54 | accuracy | 0.921053 |
| classification | stabilization | Jango | single_day | Jango_20150831_001 | Jango_20150831_001 | 44 | accuracy | 0.78125 |
| classification | stabilization | Jango | single_day | Jango_20150831_001 | Jango_20150831_001 | 46 | accuracy | 0.84375 |
| classification | stabilization | Jango | single_day | Jango_20150831_001 | Jango_20150831_001 | 54 | accuracy | 0.53125 |
| classification | stabilization | Jango | single_day | Jango_20150905_001 | Jango_20150905_001 | 44 | accuracy | 0.804878 |
| classification | stabilization | Jango | single_day | Jango_20150905_001 | Jango_20150905_001 | 46 | accuracy | 0.707317 |
| classification | stabilization | Jango | single_day | Jango_20150905_001 | Jango_20150905_001 | 54 | accuracy | 0.609756 |
| classification | stabilization | Jango | single_day | Jango_20150906_001 | Jango_20150906_001 | 44 | accuracy | 0.416667 |
| classification | stabilization | Jango | single_day | Jango_20150906_001 | Jango_20150906_001 | 46 | accuracy | 0.805556 |
| classification | stabilization | Jango | single_day | Jango_20150906_001 | Jango_20150906_001 | 54 | accuracy | 0.638889 |
| classification | stabilization | Jango | single_day | Jango_20150908_001 | Jango_20150908_001 | 44 | accuracy | 0.763158 |
| classification | stabilization | Jango | single_day | Jango_20150908_001 | Jango_20150908_001 | 46 | accuracy | 0.710526 |
| classification | stabilization | Jango | single_day | Jango_20150908_001 | Jango_20150908_001 | 54 | accuracy | 0.894737 |
| classification | stabilization | Jango | single_day | Jango_20151029_001 | Jango_20151029_001 | 44 | accuracy | 0.769231 |
| classification | stabilization | Jango | single_day | Jango_20151029_001 | Jango_20151029_001 | 46 | accuracy | 0.666667 |
| classification | stabilization | Jango | single_day | Jango_20151029_001 | Jango_20151029_001 | 54 | accuracy | 0.820513 |
| classification | stabilization | Jango | single_day | Jango_20151102_001 | Jango_20151102_001 | 44 | accuracy | 0.666667 |
| classification | stabilization | Jango | single_day | Jango_20151102_001 | Jango_20151102_001 | 46 | accuracy | 0.538462 |
| classification | stabilization | Jango | single_day | Jango_20151102_001 | Jango_20151102_001 | 54 | accuracy | 0.692308 |
| classification | mfsnn | Jango | single_day | Jango_20150730_001 | Jango_20150730_001 | 44 | accuracy | 0.727273 |
| classification | mfsnn | Jango | single_day | Jango_20150731_001 | Jango_20150731_001 | 44 | accuracy | 0.775 |
| classification | mfsnn | Jango | single_day | Jango_20150801_001 | Jango_20150801_001 | 44 | accuracy | 0.75 |
| classification | mfsnn | Jango | single_day | Jango_20150801_001 | Jango_20150801_001 | 44 | accuracy | 0.886364 |
| classification | mfsnn | Jango | single_day | Jango_20150805_001 | Jango_20150805_001 | 44 | accuracy | 0.844444 |
| classification | mfsnn | Jango | single_day | Jango_20150805_001 | Jango_20150805_001 | 44 | accuracy | 0.822222 |
| classification | mfsnn | Jango | single_day | Jango_20150806_001 | Jango_20150806_001 | 44 | accuracy | 0.777778 |
| classification | mfsnn | Jango | single_day | Jango_20150806_001 | Jango_20150806_001 | 44 | accuracy | 0.8 |
| classification | mfsnn | Jango | single_day | Jango_20150807_001 | Jango_20150807_001 | 44 | accuracy | 0.6 |
| classification | mfsnn | Jango | single_day | Jango_20150807_001 | Jango_20150807_001 | 44 | accuracy | 0.755556 |
| classification | mfsnn | Jango | single_day | Jango_20150808_001 | Jango_20150808_001 | 44 | accuracy | 0.777778 |
| classification | mfsnn | Jango | single_day | Jango_20150808_001 | Jango_20150808_001 | 44 | accuracy | 0.733333 |
| classification | mfsnn | Jango | single_day | Jango_20150809_001 | Jango_20150809_001 | 44 | accuracy | 0.955556 |
| classification | mfsnn | Jango | single_day | Jango_20150809_001 | Jango_20150809_001 | 44 | accuracy | 0.8 |
| classification | mfsnn | Jango | single_day | Jango_20150820_001 | Jango_20150820_001 | 44 | accuracy | 0.822222 |
| classification | mfsnn | Jango | single_day | Jango_20150820_001 | Jango_20150820_001 | 44 | accuracy | 0.911111 |
| classification | mfsnn | Jango | single_day | Jango_20150824_001 | Jango_20150824_001 | 44 | accuracy | 0.795455 |
| classification | mfsnn | Jango | single_day | Jango_20150825_001 | Jango_20150825_001 | 44 | accuracy | 0.847826 |
| classification | mfsnn | Jango | single_day | Jango_20150825_001 | Jango_20150825_001 | 44 | accuracy | 0.804348 |
| classification | mfsnn | Jango | single_day | Jango_20150826_001 | Jango_20150826_001 | 44 | accuracy | 0.711111 |
| classification | mfsnn | Jango | single_day | Jango_20150826_001 | Jango_20150826_001 | 44 | accuracy | 0.911111 |
| classification | mfsnn | Jango | single_day | Jango_20150827_001 | Jango_20150827_001 | 44 | accuracy | 0.826087 |
| classification | mfsnn | Jango | single_day | Jango_20150827_001 | Jango_20150827_001 | 44 | accuracy | 0.717391 |
| classification | mfsnn | Jango | single_day | Jango_20150828_001 | Jango_20150828_001 | 44 | accuracy | 0.813953 |
| classification | mfsnn | Jango | single_day | Jango_20150828_001 | Jango_20150828_001 | 44 | accuracy | 0.72093 |
| classification | mfsnn | Jango | single_day | Jango_20150831_001 | Jango_20150831_001 | 44 | accuracy | 0.621622 |
| classification | mfsnn | Jango | single_day | Jango_20150831_001 | Jango_20150831_001 | 44 | accuracy | 0.783784 |
| classification | mfsnn | Jango | single_day | Jango_20150905_001 | Jango_20150905_001 | 44 | accuracy | 0.891304 |
| classification | mfsnn | Jango | single_day | Jango_20150905_001 | Jango_20150905_001 | 44 | accuracy | 0.717391 |
| classification | mfsnn | Jango | single_day | Jango_20150906_001 | Jango_20150906_001 | 44 | accuracy | 0.829268 |
| classification | mfsnn | Jango | single_day | Jango_20150906_001 | Jango_20150906_001 | 44 | accuracy | 0.536585 |
| classification | mfsnn | Jango | single_day | Jango_20150908_001 | Jango_20150908_001 | 44 | accuracy | 0.790698 |
| classification | mfsnn | Jango | single_day | Jango_20151029_001 | Jango_20151029_001 | 44 | accuracy | 0.681818 |
| classification | mfsnn | Jango | single_day | Jango_20151029_001 | Jango_20151029_001 | 44 | accuracy | 0.613636 |
| classification | mfsnn | Jango | single_day | Jango_20151102_001 | Jango_20151102_001 | 44 | accuracy | 0.704545 |
| classification | mfsnn | Jango | single_day | Jango_20151102_001 | Jango_20151102_001 | 44 | accuracy | 0.613636 |
| classification | mscformer | Jango | single_day | Jango_20150730_001 | Jango_20150730_001 | 44 | accuracy | 0.939394 |
| classification | mscformer | Jango | single_day | Jango_20150731_001 | Jango_20150731_001 | 44 | accuracy | 0.85 |
| classification | mscformer | Jango | single_day | Jango_20150801_001 | Jango_20150801_001 | 44 | accuracy | 0.818182 |
| classification | mscformer | Jango | single_day | Jango_20150805_001 | Jango_20150805_001 | 44 | accuracy | 0.844444 |
| classification | mscformer | Jango | single_day | Jango_20150806_001 | Jango_20150806_001 | 44 | accuracy | 0.977778 |

| | | | | | | | | |
|---|---|---|---|---|---|---|---|---|
| classification | mscformer | Jango | single_day | Jango_20150807_001 | Jango_20150807_001 | 44 | accuracy | 0.8 |
| classification | mscformer | Jango | single_day | Jango_20150808_001 | Jango_20150808_001 | 44 | accuracy | 0.955556 |
| classification | mscformer | Jango | single_day | Jango_20150809_001 | Jango_20150809_001 | 44 | accuracy | 0.866667 |
| classification | mscformer | Jango | single_day | Jango_20150820_001 | Jango_20150820_001 | 44 | accuracy | 0.888889 |
| classification | mscformer | Jango | single_day | Jango_20150824_001 | Jango_20150824_001 | 44 | accuracy | 0.795455 |
| classification | mscformer | Jango | single_day | Jango_20150825_001 | Jango_20150825_001 | 44 | accuracy | 0.891304 |
| classification | mscformer | Jango | single_day | Jango_20150826_001 | Jango_20150826_001 | 44 | accuracy | 0.777778 |
| classification | mscformer | Jango | single_day | Jango_20150827_001 | Jango_20150827_001 | 44 | accuracy | 0.869565 |
| classification | mscformer | Jango | single_day | Jango_20150828_001 | Jango_20150828_001 | 44 | accuracy | 0.906977 |
| classification | mscformer | Jango | single_day | Jango_20150831_001 | Jango_20150831_001 | 44 | accuracy | 0.837838 |
| classification | mscformer | Jango | single_day | Jango_20150905_001 | Jango_20150905_001 | 44 | accuracy | 0.891304 |
| classification | mscformer | Jango | single_day | Jango_20150906_001 | Jango_20150906_001 | 44 | accuracy | 0.780488 |
| classification | mscformer | Jango | single_day | Jango_20150908_001 | Jango_20150908_001 | 44 | accuracy | 0.906977 |
| classification | mscformer | Jango | single_day | Jango_20151029_001 | Jango_20151029_001 | 44 | accuracy | 0.818182 |
| classification | mscformer | Jango | single_day | Jango_20151102_001 | Jango_20151102_001 | 44 | accuracy | 0.840909 |
| classification | nomad | Jango | single_day | Jango_20150730_001 | Jango_20150730_001 | 44 | accuracy | 0.636364 |
| classification | nomad | Jango | single_day | Jango_20150730_001 | Jango_20150730_001 | 44 | accuracy | 0.969697 |
| classification | nomad | Jango | single_day | Jango_20150731_001 | Jango_20150731_001 | 44 | accuracy | 0.625 |
| classification | nomad | Jango | single_day | Jango_20150731_001 | Jango_20150731_001 | 44 | accuracy | 0.975 |
| classification | nomad | Jango | single_day | Jango_20150801_001 | Jango_20150801_001 | 44 | accuracy | 0.727273 |
| classification | nomad | Jango | single_day | Jango_20150801_001 | Jango_20150801_001 | 44 | accuracy | 0.977273 |
| classification | nomad | Jango | single_day | Jango_20150805_001 | Jango_20150805_001 | 44 | accuracy | 0.777778 |
| classification | nomad | Jango | single_day | Jango_20150805_001 | Jango_20150805_001 | 44 | accuracy | 0.911111 |
| classification | nomad | Jango | single_day | Jango_20150806_001 | Jango_20150806_001 | 44 | accuracy | 0.888889 |
| classification | nomad | Jango | single_day | Jango_20150807_001 | Jango_20150807_001 | 44 | accuracy | 0.911111 |
| classification | nomad | Jango | single_day | Jango_20150807_001 | Jango_20150807_001 | 44 | accuracy | 0.977778 |
| classification | nomad | Jango | single_day | Jango_20150808_001 | Jango_20150808_001 | 44 | accuracy | 0.844444 |
| classification | nomad | Jango | single_day | Jango_20150808_001 | Jango_20150808_001 | 44 | accuracy | 0.955556 |
| classification | nomad | Jango | single_day | Jango_20150809_001 | Jango_20150809_001 | 44 | accuracy | 0.777778 |
| classification | nomad | Jango | single_day | Jango_20150809_001 | Jango_20150809_001 | 44 | accuracy | 0.955556 |
| classification | nomad | Jango | single_day | Jango_20150820_001 | Jango_20150820_001 | 44 | accuracy | 0.755556 |
| classification | nomad | Jango | single_day | Jango_20150820_001 | Jango_20150820_001 | 44 | accuracy | 0.777778 |
| classification | nomad | Jango | single_day | Jango_20150824_001 | Jango_20150824_001 | 44 | accuracy | 0.727273 |
| classification | nomad | Jango | single_day | Jango_20150824_001 | Jango_20150824_001 | 44 | accuracy | 0.840909 |
| classification | nomad | Jango | single_day | Jango_20150825_001 | Jango_20150825_001 | 44 | accuracy | 0.804348 |
| classification | nomad | Jango | single_day | Jango_20150825_001 | Jango_20150825_001 | 44 | accuracy | 0.891304 |
| classification | nomad | Jango | single_day | Jango_20150826_001 | Jango_20150826_001 | 44 | accuracy | 0.822222 |
| classification | nomad | Jango | single_day | Jango_20150826_001 | Jango_20150826_001 | 44 | accuracy | 0.977778 |
| classification | nomad | Jango | single_day | Jango_20150827_001 | Jango_20150827_001 | 44 | accuracy | 0.913043 |
| classification | nomad | Jango | single_day | Jango_20150827_001 | Jango_20150827_001 | 44 | accuracy | 0.891304 |
| classification | nomad | Jango | single_day | Jango_20150828_001 | Jango_20150828_001 | 44 | accuracy | 0.674419 |
| classification | nomad | Jango | single_day | Jango_20150828_001 | Jango_20150828_001 | 44 | accuracy | 0.837209 |
| classification | nomad | Jango | single_day | Jango_20150831_001 | Jango_20150831_001 | 44 | accuracy | 0.837838 |
| classification | nomad | Jango | single_day | Jango_20150831_001 | Jango_20150831_001 | 44 | accuracy | 0.945946 |
| classification | nomad | Jango | single_day | Jango_20150905_001 | Jango_20150905_001 | 44 | accuracy | 0.804348 |
| classification | nomad | Jango | single_day | Jango_20150905_001 | Jango_20150905_001 | 44 | accuracy | 0.826087 |
| classification | nomad | Jango | single_day | Jango_20150906_001 | Jango_20150906_001 | 44 | accuracy | 0.902439 |
| classification | nomad | Jango | single_day | Jango_20150906_001 | Jango_20150906_001 | 44 | accuracy | 0.878049 |
| classification | nomad | Jango | single_day | Jango_20150908_001 | Jango_20150908_001 | 44 | accuracy | 0.465116 |
| classification | nomad | Jango | single_day | Jango_20150908_001 | Jango_20150908_001 | 44 | accuracy | 0.930233 |
| classification | nomad | Jango | single_day | Jango_20151029_001 | Jango_20151029_001 | 44 | accuracy | 0.772727 |
| classification | nomad | Jango | single_day | Jango_20151029_001 | Jango_20151029_001 | 44 | accuracy | 0.863636 |
| classification | nomad | Jango | single_day | Jango_20151102_001 | Jango_20151102_001 | 44 | accuracy | 0.704545 |
| classification | nomad | Jango | single_day | Jango_20151102_001 | Jango_20151102_001 | 44 | accuracy | 0.863636 |
| classification | mlp | Jango | single_day | Jango_20150730_001 | Jango_20150730_001 | 44 | accuracy | 0.939394 |
| classification | mlp | Jango | single_day | Jango_20150730_001 | Jango_20150730_001 | 46 | accuracy | 0.787879 |
| classification | mlp | Jango | single_day | Jango_20150730_001 | Jango_20150730_001 | 54 | accuracy | 0.878788 |
| classification | mlp | Jango | single_day | Jango_20150731_001 | Jango_20150731_001 | 44 | accuracy | 0.975 |
| classification | mlp | Jango | single_day | Jango_20150731_001 | Jango_20150731_001 | 46 | accuracy | 0.975 |
| classification | mlp | Jango | single_day | Jango_20150731_001 | Jango_20150731_001 | 54 | accuracy | 0.95 |
| classification | mlp | Jango | single_day | Jango_20150801_001 | Jango_20150801_001 | 44 | accuracy | 1 |
| classification | mlp | Jango | single_day | Jango_20150801_001 | Jango_20150801_001 | 46 | accuracy | 1 |
| classification | mlp | Jango | single_day | Jango_20150801_001 | Jango_20150801_001 | 54 | accuracy | 0.931818 |

| | | | | | | | | |
|---|---|---|---|---|---|---|---|---|
| classification | mlp | Jango | single_day | Jango_20150805_001 | Jango_20150805_001 | 44 | accuracy | 0.955556 |
| classification | mlp | Jango | single_day | Jango_20150805_001 | Jango_20150805_001 | 46 | accuracy | 0.955556 |
| classification | mlp | Jango | single_day | Jango_20150805_001 | Jango_20150805_001 | 54 | accuracy | 1 |
| classification | mlp | Jango | single_day | Jango_20150806_001 | Jango_20150806_001 | 44 | accuracy | 0.933333 |
| classification | mlp | Jango | single_day | Jango_20150806_001 | Jango_20150806_001 | 46 | accuracy | 1 |
| classification | mlp | Jango | single_day | Jango_20150806_001 | Jango_20150806_001 | 54 | accuracy | 0.933333 |
| classification | mlp | Jango | single_day | Jango_20150807_001 | Jango_20150807_001 | 44 | accuracy | 0.977778 |
| classification | mlp | Jango | single_day | Jango_20150807_001 | Jango_20150807_001 | 46 | accuracy | 0.911111 |
| classification | mlp | Jango | single_day | Jango_20150807_001 | Jango_20150807_001 | 54 | accuracy | 0.977778 |
| classification | mlp | Jango | single_day | Jango_20150808_001 | Jango_20150808_001 | 44 | accuracy | 0.933333 |
| classification | mlp | Jango | single_day | Jango_20150808_001 | Jango_20150808_001 | 46 | accuracy | 0.911111 |
| classification | mlp | Jango | single_day | Jango_20150808_001 | Jango_20150808_001 | 54 | accuracy | 0.844444 |
| classification | mlp | Jango | single_day | Jango_20150809_001 | Jango_20150809_001 | 44 | accuracy | 0.911111 |
| classification | mlp | Jango | single_day | Jango_20150809_001 | Jango_20150809_001 | 46 | accuracy | 0.977778 |
| classification | mlp | Jango | single_day | Jango_20150809_001 | Jango_20150809_001 | 54 | accuracy | 1 |
| classification | mlp | Jango | single_day | Jango_20150820_001 | Jango_20150820_001 | 44 | accuracy | 0.933333 |
| classification | mlp | Jango | single_day | Jango_20150820_001 | Jango_20150820_001 | 46 | accuracy | 1 |
| classification | mlp | Jango | single_day | Jango_20150820_001 | Jango_20150820_001 | 54 | accuracy | 0.955556 |
| classification | mlp | Jango | single_day | Jango_20150824_001 | Jango_20150824_001 | 44 | accuracy | 0.931818 |
| classification | mlp | Jango | single_day | Jango_20150824_001 | Jango_20150824_001 | 46 | accuracy | 0.931818 |
| classification | mlp | Jango | single_day | Jango_20150824_001 | Jango_20150824_001 | 54 | accuracy | 0.931818 |
| classification | mlp | Jango | single_day | Jango_20150825_001 | Jango_20150825_001 | 44 | accuracy | 0.978261 |
| classification | mlp | Jango | single_day | Jango_20150825_001 | Jango_20150825_001 | 46 | accuracy | 0.956522 |
| classification | mlp | Jango | single_day | Jango_20150825_001 | Jango_20150825_001 | 54 | accuracy | 0.978261 |
| classification | mlp | Jango | single_day | Jango_20150826_001 | Jango_20150826_001 | 44 | accuracy | 0.933333 |
| classification | mlp | Jango | single_day | Jango_20150826_001 | Jango_20150826_001 | 46 | accuracy | 0.977778 |
| classification | mlp | Jango | single_day | Jango_20150826_001 | Jango_20150826_001 | 54 | accuracy | 1 |
| classification | mlp | Jango | single_day | Jango_20150827_001 | Jango_20150827_001 | 44 | accuracy | 0.978261 |
| classification | mlp | Jango | single_day | Jango_20150827_001 | Jango_20150827_001 | 46 | accuracy | 0.978261 |
| classification | mlp | Jango | single_day | Jango_20150827_001 | Jango_20150827_001 | 54 | accuracy | 0.978261 |
| classification | mlp | Jango | single_day | Jango_20150828_001 | Jango_20150828_001 | 44 | accuracy | 1 |
| classification | mlp | Jango | single_day | Jango_20150828_001 | Jango_20150828_001 | 46 | accuracy | 0.860465 |
| classification | mlp | Jango | single_day | Jango_20150828_001 | Jango_20150828_001 | 54 | accuracy | 0.953488 |
| classification | mlp | Jango | single_day | Jango_20150831_001 | Jango_20150831_001 | 44 | accuracy | 0.972973 |
| classification | mlp | Jango | single_day | Jango_20150831_001 | Jango_20150831_001 | 46 | accuracy | 0.945946 |
| classification | mlp | Jango | single_day | Jango_20150831_001 | Jango_20150831_001 | 54 | accuracy | 0.945946 |
| classification | mlp | Jango | single_day | Jango_20150905_001 | Jango_20150905_001 | 44 | accuracy | 0.869565 |
| classification | mlp | Jango | single_day | Jango_20150905_001 | Jango_20150905_001 | 46 | accuracy | 0.869565 |
| classification | mlp | Jango | single_day | Jango_20150905_001 | Jango_20150905_001 | 54 | accuracy | 0.956522 |
| classification | mlp | Jango | single_day | Jango_20150906_001 | Jango_20150906_001 | 44 | accuracy | 0.95122 |
| classification | mlp | Jango | single_day | Jango_20150906_001 | Jango_20150906_001 | 46 | accuracy | 0.97561 |
| classification | mlp | Jango | single_day | Jango_20150906_001 | Jango_20150906_001 | 54 | accuracy | 0.97561 |
| classification | mlp | Jango | single_day | Jango_20150908_001 | Jango_20150908_001 | 44 | accuracy | 0.883721 |
| classification | mlp | Jango | single_day | Jango_20150908_001 | Jango_20150908_001 | 46 | accuracy | 0.976744 |
| classification | mlp | Jango | single_day | Jango_20150908_001 | Jango_20150908_001 | 54 | accuracy | 0.930233 |
| classification | mlp | Jango | single_day | Jango_20151029_001 | Jango_20151029_001 | 44 | accuracy | 0.931818 |
| classification | mlp | Jango | single_day | Jango_20151029_001 | Jango_20151029_001 | 46 | accuracy | 0.954545 |
| classification | mlp | Jango | single_day | Jango_20151029_001 | Jango_20151029_001 | 54 | accuracy | 0.977273 |
| classification | mlp | Jango | single_day | Jango_20151102_001 | Jango_20151102_001 | 44 | accuracy | 0.909091 |
| classification | mlp | Jango | single_day | Jango_20151102_001 | Jango_20151102_001 | 46 | accuracy | 0.931818 |
| classification | mlp | Jango | single_day | Jango_20151102_001 | Jango_20151102_001 | 54 | accuracy | 0.909091 |
| classification | convpi_vae | Jango | single_day | Jango_20150730_001 | Jango_20150730_001 | 31 | accuracy | 1 |
| classification | convpi_vae | Jango | single_day | Jango_20150730_001 | Jango_20150730_001 | 44 | accuracy | 1 |
| classification | convpi_vae | Jango | single_day | Jango_20150730_001 | Jango_20150730_001 | 46 | accuracy | 1 |
| classification | convpi_vae | Jango | single_day | Jango_20150730_001 | Jango_20150730_001 | 54 | accuracy | 1 |
| classification | convpi_vae | Jango | single_day | Jango_20150730_001 | Jango_20150730_001 | 69 | accuracy | 1 |
| classification | convpi_vae | Jango | single_day | Jango_20150731_001 | Jango_20150731_001 | 31 | accuracy | 1 |
| classification | convpi_vae | Jango | single_day | Jango_20150731_001 | Jango_20150731_001 | 44 | accuracy | 1 |
| classification | convpi_vae | Jango | single_day | Jango_20150731_001 | Jango_20150731_001 | 46 | accuracy | 1 |
| classification | convpi_vae | Jango | single_day | Jango_20150731_001 | Jango_20150731_001 | 54 | accuracy | 1 |
| classification | convpi_vae | Jango | single_day | Jango_20150731_001 | Jango_20150731_001 | 69 | accuracy | 1 |
| classification | convpi_vae | Jango | single_day | Jango_20150801_001 | Jango_20150801_001 | 31 | accuracy | 0.977273 |
| classification | convpi_vae | Jango | single_day | Jango_20150801_001 | Jango_20150801_001 | 44 | accuracy | 1 |

| | | | | | | | | |
|---|---|---|---|---|---|---|---|---|
| classification | convpi_vae | Jango | single_day | Jango_20150801_001 | Jango_20150801_001 | 46 | accuracy | 1 |
| classification | convpi_vae | Jango | single_day | Jango_20150801_001 | Jango_20150801_001 | 54 | accuracy | 1 |
| classification | convpi_vae | Jango | single_day | Jango_20150801_001 | Jango_20150801_001 | 69 | accuracy | 1 |
| classification | convpi_vae | Jango | single_day | Jango_20150805_001 | Jango_20150805_001 | 31 | accuracy | 1 |
| classification | convpi_vae | Jango | single_day | Jango_20150805_001 | Jango_20150805_001 | 44 | accuracy | 1 |
| classification | convpi_vae | Jango | single_day | Jango_20150805_001 | Jango_20150805_001 | 46 | accuracy | 1 |
| classification | convpi_vae | Jango | single_day | Jango_20150805_001 | Jango_20150805_001 | 54 | accuracy | 1 |
| classification | convpi_vae | Jango | single_day | Jango_20150805_001 | Jango_20150805_001 | 69 | accuracy | 1 |
| classification | convpi_vae | Jango | single_day | Jango_20150806_001 | Jango_20150806_001 | 31 | accuracy | 1 |
| classification | convpi_vae | Jango | single_day | Jango_20150806_001 | Jango_20150806_001 | 44 | accuracy | 1 |
| classification | convpi_vae | Jango | single_day | Jango_20150806_001 | Jango_20150806_001 | 46 | accuracy | 1 |
| classification | convpi_vae | Jango | single_day | Jango_20150806_001 | Jango_20150806_001 | 54 | accuracy | 1 |
| classification | convpi_vae | Jango | single_day | Jango_20150806_001 | Jango_20150806_001 | 69 | accuracy | 1 |
| classification | convpi_vae | Jango | single_day | Jango_20150807_001 | Jango_20150807_001 | 31 | accuracy | 1 |
| classification | convpi_vae | Jango | single_day | Jango_20150807_001 | Jango_20150807_001 | 44 | accuracy | 1 |
| classification | convpi_vae | Jango | single_day | Jango_20150807_001 | Jango_20150807_001 | 46 | accuracy | 1 |
| classification | convpi_vae | Jango | single_day | Jango_20150807_001 | Jango_20150807_001 | 54 | accuracy | 1 |
| classification | convpi_vae | Jango | single_day | Jango_20150807_001 | Jango_20150807_001 | 69 | accuracy | 1 |
| classification | convpi_vae | Jango | single_day | Jango_20150808_001 | Jango_20150808_001 | 31 | accuracy | 1 |
| classification | convpi_vae | Jango | single_day | Jango_20150808_001 | Jango_20150808_001 | 44 | accuracy | 1 |
| classification | convpi_vae | Jango | single_day | Jango_20150808_001 | Jango_20150808_001 | 46 | accuracy | 1 |
| classification | convpi_vae | Jango | single_day | Jango_20150808_001 | Jango_20150808_001 | 54 | accuracy | 1 |
| classification | convpi_vae | Jango | single_day | Jango_20150808_001 | Jango_20150808_001 | 69 | accuracy | 1 |
| classification | convpi_vae | Jango | single_day | Jango_20150809_001 | Jango_20150809_001 | 31 | accuracy | 1 |
| classification | convpi_vae | Jango | single_day | Jango_20150809_001 | Jango_20150809_001 | 44 | accuracy | 1 |
| classification | convpi_vae | Jango | single_day | Jango_20150809_001 | Jango_20150809_001 | 46 | accuracy | 1 |
| classification | convpi_vae | Jango | single_day | Jango_20150809_001 | Jango_20150809_001 | 54 | accuracy | 1 |
| classification | convpi_vae | Jango | single_day | Jango_20150809_001 | Jango_20150809_001 | 69 | accuracy | 1 |
| classification | convpi_vae | Jango | single_day | Jango_20150820_001 | Jango_20150820_001 | 31 | accuracy | 1 |
| classification | convpi_vae | Jango | single_day | Jango_20150820_001 | Jango_20150820_001 | 44 | accuracy | 1 |
| classification | convpi_vae | Jango | single_day | Jango_20150820_001 | Jango_20150820_001 | 46 | accuracy | 1 |
| classification | convpi_vae | Jango | single_day | Jango_20150820_001 | Jango_20150820_001 | 54 | accuracy | 1 |
| classification | convpi_vae | Jango | single_day | Jango_20150820_001 | Jango_20150820_001 | 69 | accuracy | 1 |
| classification | convpi_vae | Jango | single_day | Jango_20150824_001 | Jango_20150824_001 | 31 | accuracy | 1 |
| classification | convpi_vae | Jango | single_day | Jango_20150824_001 | Jango_20150824_001 | 44 | accuracy | 1 |
| classification | convpi_vae | Jango | single_day | Jango_20150824_001 | Jango_20150824_001 | 46 | accuracy | 1 |
| classification | convpi_vae | Jango | single_day | Jango_20150824_001 | Jango_20150824_001 | 54 | accuracy | 1 |
| classification | convpi_vae | Jango | single_day | Jango_20150824_001 | Jango_20150824_001 | 69 | accuracy | 1 |
| classification | convpi_vae | Jango | single_day | Jango_20150825_001 | Jango_20150825_001 | 31 | accuracy | 1 |
| classification | convpi_vae | Jango | single_day | Jango_20150825_001 | Jango_20150825_001 | 44 | accuracy | 1 |
| classification | convpi_vae | Jango | single_day | Jango_20150825_001 | Jango_20150825_001 | 46 | accuracy | 1 |
| classification | convpi_vae | Jango | single_day | Jango_20150825_001 | Jango_20150825_001 | 54 | accuracy | 1 |
| classification | convpi_vae | Jango | single_day | Jango_20150825_001 | Jango_20150825_001 | 69 | accuracy | 1 |
| classification | convpi_vae | Jango | single_day | Jango_20150826_001 | Jango_20150826_001 | 31 | accuracy | 1 |
| classification | convpi_vae | Jango | single_day | Jango_20150826_001 | Jango_20150826_001 | 44 | accuracy | 1 |
| classification | convpi_vae | Jango | single_day | Jango_20150826_001 | Jango_20150826_001 | 46 | accuracy | 1 |
| classification | convpi_vae | Jango | single_day | Jango_20150826_001 | Jango_20150826_001 | 54 | accuracy | 1 |
| classification | convpi_vae | Jango | single_day | Jango_20150826_001 | Jango_20150826_001 | 69 | accuracy | 1 |
| classification | convpi_vae | Jango | single_day | Jango_20150827_001 | Jango_20150827_001 | 31 | accuracy | 1 |
| classification | convpi_vae | Jango | single_day | Jango_20150827_001 | Jango_20150827_001 | 44 | accuracy | 1 |
| classification | convpi_vae | Jango | single_day | Jango_20150827_001 | Jango_20150827_001 | 46 | accuracy | 1 |
| classification | convpi_vae | Jango | single_day | Jango_20150827_001 | Jango_20150827_001 | 54 | accuracy | 1 |
| classification | convpi_vae | Jango | single_day | Jango_20150827_001 | Jango_20150827_001 | 69 | accuracy | 1 |
| classification | convpi_vae | Jango | single_day | Jango_20150828_001 | Jango_20150828_001 | 31 | accuracy | 1 |
| classification | convpi_vae | Jango | single_day | Jango_20150828_001 | Jango_20150828_001 | 44 | accuracy | 1 |
| classification | convpi_vae | Jango | single_day | Jango_20150828_001 | Jango_20150828_001 | 46 | accuracy | 1 |
| classification | convpi_vae | Jango | single_day | Jango_20150828_001 | Jango_20150828_001 | 54 | accuracy | 1 |
| classification | convpi_vae | Jango | single_day | Jango_20150828_001 | Jango_20150828_001 | 69 | accuracy | 1 |
| classification | convpi_vae | Jango | single_day | Jango_20150831_001 | Jango_20150831_001 | 31 | accuracy | 1 |
| classification | convpi_vae | Jango | single_day | Jango_20150831_001 | Jango_20150831_001 | 44 | accuracy | 1 |
| classification | convpi_vae | Jango | single_day | Jango_20150831_001 | Jango_20150831_001 | 46 | accuracy | 1 |
| classification | convpi_vae | Jango | single_day | Jango_20150831_001 | Jango_20150831_001 | 54 | accuracy | 1 |
| classification | convpi_vae | Jango | single_day | Jango_20150831_001 | Jango_20150831_001 | 69 | accuracy | 1 |

| | | | | | | | | |
|---|---|---|---|---|---|---|---|---|
| classification | convpi_vae | Jango | single_day | Jango_20150905_001 | Jango_20150905_001 | 31 | accuracy | 1 |
| classification | convpi_vae | Jango | single_day | Jango_20150905_001 | Jango_20150905_001 | 44 | accuracy | 1 |
| classification | convpi_vae | Jango | single_day | Jango_20150905_001 | Jango_20150905_001 | 46 | accuracy | 1 |
| classification | convpi_vae | Jango | single_day | Jango_20150905_001 | Jango_20150905_001 | 54 | accuracy | 1 |
| classification | convpi_vae | Jango | single_day | Jango_20150905_001 | Jango_20150905_001 | 69 | accuracy | 1 |
| classification | convpi_vae | Jango | single_day | Jango_20150906_001 | Jango_20150906_001 | 31 | accuracy | 1 |
| classification | convpi_vae | Jango | single_day | Jango_20150906_001 | Jango_20150906_001 | 44 | accuracy | 1 |
| classification | convpi_vae | Jango | single_day | Jango_20150906_001 | Jango_20150906_001 | 46 | accuracy | 1 |
| classification | convpi_vae | Jango | single_day | Jango_20150906_001 | Jango_20150906_001 | 54 | accuracy | 1 |
| classification | convpi_vae | Jango | single_day | Jango_20150906_001 | Jango_20150906_001 | 69 | accuracy | 1 |
| classification | convpi_vae | Jango | single_day | Jango_20150908_001 | Jango_20150908_001 | 31 | accuracy | 1 |
| classification | convpi_vae | Jango | single_day | Jango_20150908_001 | Jango_20150908_001 | 44 | accuracy | 1 |
| classification | convpi_vae | Jango | single_day | Jango_20150908_001 | Jango_20150908_001 | 46 | accuracy | 1 |
| classification | convpi_vae | Jango | single_day | Jango_20150908_001 | Jango_20150908_001 | 54 | accuracy | 1 |
| classification | convpi_vae | Jango | single_day | Jango_20150908_001 | Jango_20150908_001 | 69 | accuracy | 1 |
| classification | convpi_vae | Jango | single_day | Jango_20151029_001 | Jango_20151029_001 | 31 | accuracy | 1 |
| classification | convpi_vae | Jango | single_day | Jango_20151029_001 | Jango_20151029_001 | 44 | accuracy | 1 |
| classification | convpi_vae | Jango | single_day | Jango_20151029_001 | Jango_20151029_001 | 46 | accuracy | 1 |
| classification | convpi_vae | Jango | single_day | Jango_20151029_001 | Jango_20151029_001 | 54 | accuracy | 1 |
| classification | convpi_vae | Jango | single_day | Jango_20151029_001 | Jango_20151029_001 | 69 | accuracy | 1 |
| classification | convpi_vae | Jango | single_day | Jango_20151102_001 | Jango_20151102_001 | 31 | accuracy | 1 |
| classification | convpi_vae | Jango | single_day | Jango_20151102_001 | Jango_20151102_001 | 44 | accuracy | 1 |
| classification | convpi_vae | Jango | single_day | Jango_20151102_001 | Jango_20151102_001 | 46 | accuracy | 1 |
| classification | convpi_vae | Jango | single_day | Jango_20151102_001 | Jango_20151102_001 | 54 | accuracy | 1 |
| classification | convpi_vae | Jango | single_day | Jango_20151102_001 | Jango_20151102_001 | 69 | accuracy | 1 |
| classification | lstm | Jango | cross_day | Jango_20150730_001__to__Jango_20... | Jango_20150906_001__to__Jango_20... | 31 | accuracy | 0.367803 |
| classification | lstm | Jango | cross_day | Jango_20150730_001__to__Jango_20... | Jango_20150906_001__to__Jango_20... | 44 | accuracy | 0.341951 |
| classification | lstm | Jango | cross_day | Jango_20150730_001__to__Jango_20... | Jango_20150906_001__to__Jango_20... | 44 | accuracy | 0.342055 |
| classification | lstm | Jango | cross_day | Jango_20150730_001__to__Jango_20... | Jango_20150906_001__to__Jango_20... | 44 | accuracy | 0.465687 |
| classification | lstm | Jango | cross_day | Jango_20150730_001__to__Jango_20... | Jango_20150906_001__to__Jango_20... | 44 | accuracy | 0.341951 |
| classification | lstm | Jango | cross_day | Jango_20150730_001__to__Jango_20... | Jango_20150906_001__to__Jango_20... | 46 | accuracy | 0.403055 |
| classification | lstm | Jango | cross_day | Jango_20150730_001__to__Jango_20... | Jango_20150906_001__to__Jango_20... | 46 | accuracy | 0.404534 |
| classification | lstm | Jango | cross_day | Jango_20150730_001__to__Jango_20... | Jango_20150906_001__to__Jango_20... | 46 | accuracy | 0.443608 |
| classification | lstm | Jango | cross_day | Jango_20150730_001__to__Jango_20... | Jango_20150906_001__to__Jango_20... | 46 | accuracy | 0.403055 |
| classification | lstm | Jango | cross_day | Jango_20150730_001__to__Jango_20... | Jango_20150906_001__to__Jango_20... | 54 | accuracy | 0.374853 |
| classification | lstm | Jango | cross_day | Jango_20150730_001__to__Jango_20... | Jango_20150906_001__to__Jango_20... | 54 | accuracy | 0.376869 |
| classification | lstm | Jango | cross_day | Jango_20150730_001__to__Jango_20... | Jango_20150906_001__to__Jango_20... | 54 | accuracy | 0.441985 |
| classification | lstm | Jango | cross_day | Jango_20150730_001__to__Jango_20... | Jango_20150906_001__to__Jango_20... | 69 | accuracy | 0.387779 |
| classification | lstm | Jango | cross_day | Jango_20150730_001__to__Jango_20... | Jango_20150906_001__to__Jango_20... | 31 | accuracy | 0.475911 |
| classification | lstm | Jango | cross_day | Jango_20150730_001__to__Jango_20... | Jango_20150906_001__to__Jango_20... | 44 | accuracy | 0.540541 |
| classification | lstm | Jango | cross_day | Jango_20150730_001__to__Jango_20... | Jango_20150906_001__to__Jango_20... | 46 | accuracy | 0.498237 |
| classification | lstm | Jango | cross_day | Jango_20150730_001__to__Jango_20... | Jango_20150906_001__to__Jango_20... | 54 | accuracy | 0.391304 |
| classification | lstm | Jango | cross_day | Jango_20150730_001__to__Jango_20... | Jango_20150906_001__to__Jango_20... | 69 | accuracy | 0.475911 |
| classification | transformer | Jango | cross_day | Jango_20150730_001__to__Jango_20... | Jango_20150906_001__to__Jango_20... | 31 | accuracy | 0.525264 |
| classification | transformer | Jango | cross_day | Jango_20150730_001__to__Jango_20... | Jango_20150906_001__to__Jango_20... | 44 | accuracy | 0.473561 |
| classification | transformer | Jango | cross_day | Jango_20150730_001__to__Jango_20... | Jango_20150906_001__to__Jango_20... | 44 | accuracy | 0.473148 |
| classification | transformer | Jango | cross_day | Jango_20150730_001__to__Jango_20... | Jango_20150906_001__to__Jango_20... | 44 | accuracy | 0.628637 |
| classification | transformer | Jango | cross_day | Jango_20150730_001__to__Jango_20... | Jango_20150906_001__to__Jango_20... | 46 | accuracy | 0.502938 |
| classification | transformer | Jango | cross_day | Jango_20150730_001__to__Jango_20... | Jango_20150906_001__to__Jango_20... | 46 | accuracy | 0.502406 |
| classification | transformer | Jango | cross_day | Jango_20150730_001__to__Jango_20... | Jango_20150906_001__to__Jango_20... | 46 | accuracy | 0.612915 |
| classification | transformer | Jango | cross_day | Jango_20150730_001__to__Jango_20... | Jango_20150906_001__to__Jango_20... | 46 | accuracy | 0.502938 |
| classification | transformer | Jango | cross_day | Jango_20150730_001__to__Jango_20... | Jango_20150906_001__to__Jango_20... | 54 | accuracy | 0.508813 |
| classification | transformer | Jango | cross_day | Jango_20150730_001__to__Jango_20... | Jango_20150906_001__to__Jango_20... | 54 | accuracy | 0.507949 |
| classification | transformer | Jango | cross_day | Jango_20150730_001__to__Jango_20... | Jango_20150906_001__to__Jango_20... | 54 | accuracy | 0.48108 |
| classification | transformer | Jango | cross_day | Jango_20150730_001__to__Jango_20... | Jango_20150906_001__to__Jango_20... | 69 | accuracy | 0.545241 |
| classification | transformer | Jango | cross_day | Jango_20150730_001__to__Jango_20... | Jango_20150906_001__to__Jango_20... | 31 | accuracy | 0.582844 |
| classification | transformer | Jango | cross_day | Jango_20150730_001__to__Jango_20... | Jango_20150906_001__to__Jango_20... | 44 | accuracy | 0.581669 |
| classification | transformer | Jango | cross_day | Jango_20150730_001__to__Jango_20... | Jango_20150906_001__to__Jango_20... | 46 | accuracy | 0.537015 |
| classification | transformer | Jango | cross_day | Jango_20150730_001__to__Jango_20... | Jango_20150906_001__to__Jango_20... | 54 | accuracy | 0.509988 |
| classification | transformer | Jango | cross_day | Jango_20150730_001__to__Jango_20... | Jango_20150906_001__to__Jango_20... | 69 | accuracy | 0.506463 |
| classification | lfads | Jango | cross_day | Jango_20150730_001__to__Jango_20... | Jango_20150906_001__to__Jango_20... | 44 | accuracy | 0.504113 |
| classification | lfads | Jango | cross_day | Jango_20150730_001__to__Jango_20... | Jango_20150906_001__to__Jango_20... | 44 | accuracy | 0.539562 |
| classification | lfads | Jango | cross_day | Jango_20150730_001__to__Jango_20... | Jango_20150906_001__to__Jango_20... | 46 | accuracy | 0.512338 |

| | | | | | | | | |
|---|---|---|---|---|---|---|---|---|
| classification | lfads | Jango | cross_day | Jango_20150730_001__to__Jango_20... | Jango_20150906_001__to__Jango_20... | 46 | accuracy | 0.52788 |
| classification | lfads | Jango | cross_day | Jango_20150730_001__to__Jango_20... | Jango_20150906_001__to__Jango_20... | 54 | accuracy | 0.494712 |
| classification | lfads | Jango | cross_day | Jango_20150730_001__to__Jango_20... | Jango_20150906_001__to__Jango_20... | 54 | accuracy | 0.498297 |
| classification | cycle_gan | Jango | cross_day | Jango_20150730_001__to__Jango_20... | Jango_20150906_001__to__Jango_20... | 31 | accuracy | 0.453584 |
| classification | cycle_gan | Jango | cross_day | Jango_20150730_001__to__Jango_20... | Jango_20150906_001__to__Jango_20... | 44 | accuracy | 0.621194 |
| classification | cycle_gan | Jango | cross_day | Jango_20150730_001__to__Jango_20... | Jango_20150906_001__to__Jango_20... | 44 | accuracy | 0.200719 |
| classification | cycle_gan | Jango | cross_day | Jango_20150730_001__to__Jango_20... | Jango_20150906_001__to__Jango_20... | 44 | accuracy | 0.47585 |
| classification | cycle_gan | Jango | cross_day | Jango_20150730_001__to__Jango_20... | Jango_20150906_001__to__Jango_20... | 44 | accuracy | 0.294947 |
| classification | cycle_gan | Jango | cross_day | Jango_20150730_001__to__Jango_20... | Jango_20150906_001__to__Jango_20... | 46 | accuracy | 0.148599 |
| classification | cycle_gan | Jango | cross_day | Jango_20150730_001__to__Jango_20... | Jango_20150906_001__to__Jango_20... | 46 | accuracy | 0.217542 |
| classification | cycle_gan | Jango | cross_day | Jango_20150730_001__to__Jango_20... | Jango_20150906_001__to__Jango_20... | 46 | accuracy | 0.436098 |
| classification | cycle_gan | Jango | cross_day | Jango_20150730_001__to__Jango_20... | Jango_20150906_001__to__Jango_20... | 46 | accuracy | 0.431257 |
| classification | cycle_gan | Jango | cross_day | Jango_20150730_001__to__Jango_20... | Jango_20150906_001__to__Jango_20... | 54 | accuracy | 0.186358 |
| classification | cycle_gan | Jango | cross_day | Jango_20150730_001__to__Jango_20... | Jango_20150906_001__to__Jango_20... | 54 | accuracy | 0.206864 |
| classification | cycle_gan | Jango | cross_day | Jango_20150730_001__to__Jango_20... | Jango_20150906_001__to__Jango_20... | 54 | accuracy | 0.408062 |
| classification | cycle_gan | Jango | cross_day | Jango_20150730_001__to__Jango_20... | Jango_20150906_001__to__Jango_20... | 54 | accuracy | 0.354877 |
| classification | cycle_gan | Jango | cross_day | Jango_20150730_001__to__Jango_20... | Jango_20150906_001__to__Jango_20... | 69 | accuracy | 0.294947 |
| classification | cycle_gan | Jango | cross_day | Jango_20150730_001__to__Jango_20... | Jango_20150906_001__to__Jango_20... | 31 | accuracy | 0.682726 |
| classification | cycle_gan | Jango | cross_day | Jango_20150730_001__to__Jango_20... | Jango_20150906_001__to__Jango_20... | 44 | accuracy | 0.567568 |
| classification | cycle_gan | Jango | cross_day | Jango_20150730_001__to__Jango_20... | Jango_20150906_001__to__Jango_20... | 46 | accuracy | 0.556992 |
| classification | cycle_gan | Jango | cross_day | Jango_20150730_001__to__Jango_20... | Jango_20150906_001__to__Jango_20... | 54 | accuracy | 0.709753 |
| classification | cycle_gan | Jango | cross_day | Jango_20150730_001__to__Jango_20... | Jango_20150906_001__to__Jango_20... | 69 | accuracy | 0.710928 |
| classification | stabilization | Jango | cross_day | Jango_20150730_001__to__Jango_20... | Jango_20150906_001__to__Jango_20... | 44 | accuracy | 0.622412 |
| classification | stabilization | Jango | cross_day | Jango_20150730_001__to__Jango_20... | Jango_20150906_001__to__Jango_20... | 44 | accuracy | 0.750487 |
| classification | stabilization | Jango | cross_day | Jango_20150730_001__to__Jango_20... | Jango_20150906_001__to__Jango_20... | 44 | accuracy | 0.779761 |
| classification | stabilization | Jango | cross_day | Jango_20150730_001__to__Jango_20... | Jango_20150906_001__to__Jango_20... | 46 | accuracy | 0.674787 |
| classification | stabilization | Jango | cross_day | Jango_20150730_001__to__Jango_20... | Jango_20150906_001__to__Jango_20... | 46 | accuracy | 0.74436 |
| classification | stabilization | Jango | cross_day | Jango_20150730_001__to__Jango_20... | Jango_20150906_001__to__Jango_20... | 46 | accuracy | 0.777162 |
| classification | stabilization | Jango | cross_day | Jango_20150730_001__to__Jango_20... | Jango_20150906_001__to__Jango_20... | 54 | accuracy | 0.680877 |
| classification | stabilization | Jango | cross_day | Jango_20150730_001__to__Jango_20... | Jango_20150906_001__to__Jango_20... | 54 | accuracy | 0.717785 |
| classification | stabilization | Jango | cross_day | Jango_20150730_001__to__Jango_20... | Jango_20150906_001__to__Jango_20... | 54 | accuracy | 0.794937 |
| classification | mfsnn | Jango | cross_day | Jango_20150730_001__to__Jango_20... | Jango_20150906_001__to__Jango_20... | 44 | accuracy | 0.503087 |
| classification | mfsnn | Jango | cross_day | Jango_20150730_001__to__Jango_20... | Jango_20150906_001__to__Jango_20... | 44 | accuracy | 0.753069 |
| classification | mscformer | Jango | cross_day | Jango_20150730_001__to__Jango_20... | Jango_20150906_001 | 44 | accuracy | 0.946078 |
| classification | mscformer | Jango | cross_day | Jango_20150730_001__to__Jango_20... | Jango_20150908_001 | 44 | accuracy | 0.938967 |
| classification | mscformer | Jango | cross_day | Jango_20150730_001__to__Jango_20... | Jango_20151029_001 | 44 | accuracy | 0.87037 |
| classification | mscformer | Jango | cross_day | Jango_20150730_001__to__Jango_20... | Jango_20151102_001 | 44 | accuracy | 0.83945 |
| classification | nomad | Jango | cross_day | Jango_20150730_001__to__Jango_20... | Jango_20150906_001__to__Jango_20... | 44 | accuracy | 0.893934 |
| classification | nomad | Jango | cross_day | Jango_20150730_001__to__Jango_20... | Jango_20150906_001__to__Jango_20... | 44 | accuracy | 0.901811 |
| classification | mlp | Jango | cross_day | Jango_20150730_001__to__Jango_20... | Jango_20150906_001 | 44 | accuracy | 0.583333 |
| classification | mlp | Jango | cross_day | Jango_20150730_001__to__Jango_20... | Jango_20150906_001 | 46 | accuracy | 0.651961 |
| classification | mlp | Jango | cross_day | Jango_20150730_001__to__Jango_20... | Jango_20150906_001 | 54 | accuracy | 0.612745 |
| classification | mlp | Jango | cross_day | Jango_20150730_001__to__Jango_20... | Jango_20150908_001 | 44 | accuracy | 0.126761 |
| classification | mlp | Jango | cross_day | Jango_20150730_001__to__Jango_20... | Jango_20150908_001 | 46 | accuracy | 0.126761 |
| classification | mlp | Jango | cross_day | Jango_20150730_001__to__Jango_20... | Jango_20150908_001 | 54 | accuracy | 0.126761 |
| classification | mlp | Jango | cross_day | Jango_20150730_001__to__Jango_20... | Jango_20151029_001 | 44 | accuracy | 0.740741 |
| classification | mlp | Jango | cross_day | Jango_20150730_001__to__Jango_20... | Jango_20151029_001 | 46 | accuracy | 0.638889 |
| classification | mlp | Jango | cross_day | Jango_20150730_001__to__Jango_20... | Jango_20151029_001 | 54 | accuracy | 0.773148 |
| classification | mlp | Jango | cross_day | Jango_20150730_001__to__Jango_20... | Jango_20151102_001 | 44 | accuracy | 0.486239 |
| classification | mlp | Jango | cross_day | Jango_20150730_001__to__Jango_20... | Jango_20151102_001 | 46 | accuracy | 0.380734 |
| classification | mlp | Jango | cross_day | Jango_20150730_001__to__Jango_20... | Jango_20151102_001 | 54 | accuracy | 0.5 |
| classification | convpi_vae | Jango | cross_day | Jango_20150730_001__to__Jango_20... | Jango_20150906_001__to__Jango_20... | 31 | accuracy | 0.169213 |
| classification | convpi_vae | Jango | cross_day | Jango_20150730_001__to__Jango_20... | Jango_20150906_001__to__Jango_20... | 44 | accuracy | 0.139835 |
| classification | convpi_vae | Jango | cross_day | Jango_20150730_001__to__Jango_20... | Jango_20150906_001__to__Jango_20... | 46 | accuracy | 0.169213 |
| classification | convpi_vae | Jango | cross_day | Jango_20150730_001__to__Jango_20... | Jango_20150906_001__to__Jango_20... | 54 | accuracy | 0.176263 |
| classification | convpi_vae | Jango | cross_day | Jango_20150730_001__to__Jango_20... | Jango_20150906_001__to__Jango_20... | 69 | accuracy | 0.176263 |
| classification | convpi_vae | Jango | cross_day | Jango_20150730_001__to__Jango_20... | Jango_20150906_001__to__Jango_20... | 31 | accuracy | 1 |
| classification | convpi_vae | Jango | cross_day | Jango_20150730_001__to__Jango_20... | Jango_20150906_001__to__Jango_20... | 44 | accuracy | 1 |
| classification | convpi_vae | Jango | cross_day | Jango_20150730_001__to__Jango_20... | Jango_20150906_001__to__Jango_20... | 46 | accuracy | 0.875441 |
| classification | convpi_vae | Jango | cross_day | Jango_20150730_001__to__Jango_20... | Jango_20150906_001__to__Jango_20... | 54 | accuracy | 0.905993 |
| classification | convpi_vae | Jango | cross_day | Jango_20150730_001__to__Jango_20... | Jango_20150906_001__to__Jango_20... | 69 | accuracy | 0.990599 |
| classification | lstm | Visual Grating | single_day | ECoG-1-1015.mat | ECoG-1-1015.mat | 44 | accuracy | 0.615385 |
| classification | lstm | Visual Grating | single_day | ECoG-1-1015.mat | ECoG-1-1015.mat | 44 | accuracy | 0.615385 |

| | | | | | | | | |
|---|---|---|---|---|---|---|---|---|
| classification | lstm | Visual Grating | single_day | ECoG-1-1015.mat | ECoG-1-1015.mat | 46 | accuracy | 0.538462 |
| classification | lstm | Visual Grating | single_day | ECoG-1-1015.mat | ECoG-1-1015.mat | 46 | accuracy | 0.523077 |
| classification | lstm | Visual Grating | single_day | ECoG-1-1015.mat | ECoG-1-1015.mat | 54 | accuracy | 0.430769 |
| classification | lstm | Visual Grating | single_day | ECoG-1-1015.mat | ECoG-1-1015.mat | 54 | accuracy | 0.461538 |
| classification | lstm | Visual Grating | single_day | ECoG-1-1016.mat | ECoG-1-1016.mat | 44 | accuracy | 0.646154 |
| classification | lstm | Visual Grating | single_day | ECoG-1-1016.mat | ECoG-1-1016.mat | 44 | accuracy | 0.676923 |
| classification | lstm | Visual Grating | single_day | ECoG-1-1016.mat | ECoG-1-1016.mat | 46 | accuracy | 0.723077 |
| classification | lstm | Visual Grating | single_day | ECoG-1-1016.mat | ECoG-1-1016.mat | 46 | accuracy | 0.676923 |
| classification | lstm | Visual Grating | single_day | ECoG-1-1016.mat | ECoG-1-1016.mat | 54 | accuracy | 0.646154 |
| classification | lstm | Visual Grating | single_day | ECoG-1-1016.mat | ECoG-1-1016.mat | 54 | accuracy | 0.707692 |
| classification | lstm | Visual Grating | single_day | ECoG-1-1019.mat | ECoG-1-1019.mat | 44 | accuracy | 0.784615 |
| classification | lstm | Visual Grating | single_day | ECoG-1-1019.mat | ECoG-1-1019.mat | 44 | accuracy | 0.784615 |
| classification | lstm | Visual Grating | single_day | ECoG-1-1019.mat | ECoG-1-1019.mat | 46 | accuracy | 0.846154 |
| classification | lstm | Visual Grating | single_day | ECoG-1-1019.mat | ECoG-1-1019.mat | 46 | accuracy | 0.846154 |
| classification | lstm | Visual Grating | single_day | ECoG-1-1019.mat | ECoG-1-1019.mat | 54 | accuracy | 0.615385 |
| classification | lstm | Visual Grating | single_day | ECoG-1-1019.mat | ECoG-1-1019.mat | 54 | accuracy | 0.615385 |
| classification | lstm | Visual Grating | single_day | ECoG-2-0723.mat | ECoG-2-0723.mat | 44 | accuracy | 0.676923 |
| classification | lstm | Visual Grating | single_day | ECoG-2-0723.mat | ECoG-2-0723.mat | 44 | accuracy | 0.692308 |
| classification | lstm | Visual Grating | single_day | ECoG-2-0723.mat | ECoG-2-0723.mat | 46 | accuracy | 0.723077 |
| classification | lstm | Visual Grating | single_day | ECoG-2-0723.mat | ECoG-2-0723.mat | 46 | accuracy | 0.723077 |
| classification | lstm | Visual Grating | single_day | ECoG-2-0723.mat | ECoG-2-0723.mat | 54 | accuracy | 0.692308 |
| classification | lstm | Visual Grating | single_day | ECoG-2-0723.mat | ECoG-2-0723.mat | 54 | accuracy | 0.692308 |
| classification | lstm | Visual Grating | single_day | ECoG-2-0802.mat | ECoG-2-0802.mat | 44 | accuracy | 0.646154 |
| classification | lstm | Visual Grating | single_day | ECoG-2-0802.mat | ECoG-2-0802.mat | 44 | accuracy | 0.676923 |
| classification | lstm | Visual Grating | single_day | ECoG-2-0802.mat | ECoG-2-0802.mat | 46 | accuracy | 0.492308 |
| classification | lstm | Visual Grating | single_day | ECoG-2-0802.mat | ECoG-2-0802.mat | 46 | accuracy | 0.446154 |
| classification | lstm | Visual Grating | single_day | ECoG-2-0802.mat | ECoG-2-0802.mat | 54 | accuracy | 0.646154 |
| classification | lstm | Visual Grating | single_day | ECoG-2-0802.mat | ECoG-2-0802.mat | 54 | accuracy | 0.615385 |
| classification | lstm | Visual Grating | single_day | ECoG-2-0803.mat | ECoG-2-0803.mat | 44 | accuracy | 0.753846 |
| classification | lstm | Visual Grating | single_day | ECoG-2-0803.mat | ECoG-2-0803.mat | 44 | accuracy | 0.753846 |
| classification | lstm | Visual Grating | single_day | ECoG-2-0803.mat | ECoG-2-0803.mat | 46 | accuracy | 0.676923 |
| classification | lstm | Visual Grating | single_day | ECoG-2-0803.mat | ECoG-2-0803.mat | 46 | accuracy | 0.676923 |
| classification | lstm | Visual Grating | single_day | ECoG-2-0803.mat | ECoG-2-0803.mat | 54 | accuracy | 0.738462 |
| classification | lstm | Visual Grating | single_day | ECoG-2-0803.mat | ECoG-2-0803.mat | 54 | accuracy | 0.769231 |
| classification | lstm | Visual Grating | single_day | ECoG-2-0804.mat | ECoG-2-0804.mat | 44 | accuracy | 0.723077 |
| classification | lstm | Visual Grating | single_day | ECoG-2-0804.mat | ECoG-2-0804.mat | 44 | accuracy | 0.723077 |
| classification | lstm | Visual Grating | single_day | ECoG-2-0804.mat | ECoG-2-0804.mat | 46 | accuracy | 0.676923 |
| classification | lstm | Visual Grating | single_day | ECoG-2-0804.mat | ECoG-2-0804.mat | 46 | accuracy | 0.753846 |
| classification | lstm | Visual Grating | single_day | ECoG-2-0804.mat | ECoG-2-0804.mat | 54 | accuracy | 0.815385 |
| classification | lstm | Visual Grating | single_day | ECoG-2-0804.mat | ECoG-2-0804.mat | 54 | accuracy | 0.738462 |
| classification | lstm | Visual Grating | single_day | ECoG-2-0806.mat | ECoG-2-0806.mat | 44 | accuracy | 0.553846 |
| classification | lstm | Visual Grating | single_day | ECoG-2-0806.mat | ECoG-2-0806.mat | 44 | accuracy | 0.569231 |
| classification | lstm | Visual Grating | single_day | ECoG-2-0806.mat | ECoG-2-0806.mat | 46 | accuracy | 0.6 |
| classification | lstm | Visual Grating | single_day | ECoG-2-0806.mat | ECoG-2-0806.mat | 46 | accuracy | 0.692308 |
| classification | lstm | Visual Grating | single_day | ECoG-2-0806.mat | ECoG-2-0806.mat | 54 | accuracy | 0.584615 |
| classification | lstm | Visual Grating | single_day | ECoG-2-0806.mat | ECoG-2-0806.mat | 54 | accuracy | 0.584615 |
| classification | lstm | Visual Grating | single_day | ECoG-3-0722.mat | ECoG-3-0722.mat | 44 | accuracy | 0.865979 |
| classification | lstm | Visual Grating | single_day | ECoG-3-0722.mat | ECoG-3-0722.mat | 44 | accuracy | 0.865979 |
| classification | lstm | Visual Grating | single_day | ECoG-3-0722.mat | ECoG-3-0722.mat | 46 | accuracy | 0.845361 |
| classification | lstm | Visual Grating | single_day | ECoG-3-0722.mat | ECoG-3-0722.mat | 46 | accuracy | 0.85567 |
| classification | lstm | Visual Grating | single_day | ECoG-3-0722.mat | ECoG-3-0722.mat | 54 | accuracy | 0.865979 |
| classification | lstm | Visual Grating | single_day | ECoG-3-0722.mat | ECoG-3-0722.mat | 54 | accuracy | 0.876289 |
| classification | lstm | Visual Grating | single_day | ECoG-3-0805.mat | ECoG-3-0805.mat | 44 | accuracy | 0.692308 |
| classification | lstm | Visual Grating | single_day | ECoG-3-0805.mat | ECoG-3-0805.mat | 44 | accuracy | 0.676923 |
| classification | lstm | Visual Grating | single_day | ECoG-3-0805.mat | ECoG-3-0805.mat | 46 | accuracy | 0.692308 |
| classification | lstm | Visual Grating | single_day | ECoG-3-0805.mat | ECoG-3-0805.mat | 46 | accuracy | 0.692308 |
| classification | lstm | Visual Grating | single_day | ECoG-3-0805.mat | ECoG-3-0805.mat | 54 | accuracy | 0.6 |
| classification | lstm | Visual Grating | single_day | ECoG-3-0805.mat | ECoG-3-0805.mat | 54 | accuracy | 0.630769 |
| classification | transformer | Visual Grating | single_day | ECoG-1-1015.mat | ECoG-1-1015.mat | 44 | accuracy | 0.538462 |
| classification | transformer | Visual Grating | single_day | ECoG-1-1015.mat | ECoG-1-1015.mat | 44 | accuracy | 0.538462 |
| classification | transformer | Visual Grating | single_day | ECoG-1-1015.mat | ECoG-1-1015.mat | 46 | accuracy | 0.446154 |
| classification | transformer | Visual Grating | single_day | ECoG-1-1015.mat | ECoG-1-1015.mat | 46 | accuracy | 0.446154 |
| classification | transformer | Visual Grating | single_day | ECoG-1-1015.mat | ECoG-1-1015.mat | 54 | accuracy | 0.507692 |

| | | | | | | | | |
|---|---|---|---|---|---|---|---|---|
| classification | transformer | Visual Grating | single_day | ECoG-1-1015.mat | ECoG-1-1015.mat | 54 | accuracy | 0.476923 |
| classification | transformer | Visual Grating | single_day | ECoG-1-1016.mat | ECoG-1-1016.mat | 44 | accuracy | 0.646154 |
| classification | transformer | Visual Grating | single_day | ECoG-1-1016.mat | ECoG-1-1016.mat | 44 | accuracy | 0.676923 |
| classification | transformer | Visual Grating | single_day | ECoG-1-1016.mat | ECoG-1-1016.mat | 46 | accuracy | 0.615385 |
| classification | transformer | Visual Grating | single_day | ECoG-1-1016.mat | ECoG-1-1016.mat | 46 | accuracy | 0.692308 |
| classification | transformer | Visual Grating | single_day | ECoG-1-1016.mat | ECoG-1-1016.mat | 54 | accuracy | 0.538462 |
| classification | transformer | Visual Grating | single_day | ECoG-1-1016.mat | ECoG-1-1016.mat | 54 | accuracy | 0.538462 |
| classification | transformer | Visual Grating | single_day | ECoG-1-1019.mat | ECoG-1-1019.mat | 44 | accuracy | 0.830769 |
| classification | transformer | Visual Grating | single_day | ECoG-1-1019.mat | ECoG-1-1019.mat | 44 | accuracy | 0.830769 |
| classification | transformer | Visual Grating | single_day | ECoG-1-1019.mat | ECoG-1-1019.mat | 46 | accuracy | 0.676923 |
| classification | transformer | Visual Grating | single_day | ECoG-1-1019.mat | ECoG-1-1019.mat | 46 | accuracy | 0.692308 |
| classification | transformer | Visual Grating | single_day | ECoG-1-1019.mat | ECoG-1-1019.mat | 54 | accuracy | 0.615385 |
| classification | transformer | Visual Grating | single_day | ECoG-1-1019.mat | ECoG-1-1019.mat | 54 | accuracy | 0.630769 |
| classification | transformer | Visual Grating | single_day | ECoG-2-0723.mat | ECoG-2-0723.mat | 44 | accuracy | 0.538462 |
| classification | transformer | Visual Grating | single_day | ECoG-2-0723.mat | ECoG-2-0723.mat | 44 | accuracy | 0.538462 |
| classification | transformer | Visual Grating | single_day | ECoG-2-0723.mat | ECoG-2-0723.mat | 46 | accuracy | 0.661538 |
| classification | transformer | Visual Grating | single_day | ECoG-2-0723.mat | ECoG-2-0723.mat | 46 | accuracy | 0.692308 |
| classification | transformer | Visual Grating | single_day | ECoG-2-0723.mat | ECoG-2-0723.mat | 54 | accuracy | 0.538462 |
| classification | transformer | Visual Grating | single_day | ECoG-2-0723.mat | ECoG-2-0723.mat | 54 | accuracy | 0.507692 |
| classification | transformer | Visual Grating | single_day | ECoG-2-0802.mat | ECoG-2-0802.mat | 44 | accuracy | 0.553846 |
| classification | transformer | Visual Grating | single_day | ECoG-2-0802.mat | ECoG-2-0802.mat | 44 | accuracy | 0.584615 |
| classification | transformer | Visual Grating | single_day | ECoG-2-0802.mat | ECoG-2-0802.mat | 46 | accuracy | 0.415385 |
| classification | transformer | Visual Grating | single_day | ECoG-2-0802.mat | ECoG-2-0802.mat | 46 | accuracy | 0.415385 |
| classification | transformer | Visual Grating | single_day | ECoG-2-0802.mat | ECoG-2-0802.mat | 54 | accuracy | 0.661538 |
| classification | transformer | Visual Grating | single_day | ECoG-2-0802.mat | ECoG-2-0802.mat | 54 | accuracy | 0.676923 |
| classification | transformer | Visual Grating | single_day | ECoG-2-0803.mat | ECoG-2-0803.mat | 44 | accuracy | 0.769231 |
| classification | transformer | Visual Grating | single_day | ECoG-2-0803.mat | ECoG-2-0803.mat | 44 | accuracy | 0.784615 |
| classification | transformer | Visual Grating | single_day | ECoG-2-0803.mat | ECoG-2-0803.mat | 46 | accuracy | 0.630769 |
| classification | transformer | Visual Grating | single_day | ECoG-2-0803.mat | ECoG-2-0803.mat | 46 | accuracy | 0.661538 |
| classification | transformer | Visual Grating | single_day | ECoG-2-0803.mat | ECoG-2-0803.mat | 54 | accuracy | 0.569231 |
| classification | transformer | Visual Grating | single_day | ECoG-2-0803.mat | ECoG-2-0803.mat | 54 | accuracy | 0.569231 |
| classification | transformer | Visual Grating | single_day | ECoG-2-0804.mat | ECoG-2-0804.mat | 44 | accuracy | 0.646154 |
| classification | transformer | Visual Grating | single_day | ECoG-2-0804.mat | ECoG-2-0804.mat | 44 | accuracy | 0.723077 |
| classification | transformer | Visual Grating | single_day | ECoG-2-0804.mat | ECoG-2-0804.mat | 46 | accuracy | 0.723077 |
| classification | transformer | Visual Grating | single_day | ECoG-2-0804.mat | ECoG-2-0804.mat | 46 | accuracy | 0.723077 |
| classification | transformer | Visual Grating | single_day | ECoG-2-0804.mat | ECoG-2-0804.mat | 54 | accuracy | 0.646154 |
| classification | transformer | Visual Grating | single_day | ECoG-2-0804.mat | ECoG-2-0804.mat | 54 | accuracy | 0.646154 |
| classification | transformer | Visual Grating | single_day | ECoG-2-0806.mat | ECoG-2-0806.mat | 44 | accuracy | 0.646154 |
| classification | transformer | Visual Grating | single_day | ECoG-2-0806.mat | ECoG-2-0806.mat | 44 | accuracy | 0.630769 |
| classification | transformer | Visual Grating | single_day | ECoG-2-0806.mat | ECoG-2-0806.mat | 46 | accuracy | 0.523077 |
| classification | transformer | Visual Grating | single_day | ECoG-2-0806.mat | ECoG-2-0806.mat | 46 | accuracy | 0.507692 |
| classification | transformer | Visual Grating | single_day | ECoG-2-0806.mat | ECoG-2-0806.mat | 54 | accuracy | 0.630769 |
| classification | transformer | Visual Grating | single_day | ECoG-2-0806.mat | ECoG-2-0806.mat | 54 | accuracy | 0.646154 |
| classification | transformer | Visual Grating | single_day | ECoG-3-0722.mat | ECoG-3-0722.mat | 44 | accuracy | 0.845361 |
| classification | transformer | Visual Grating | single_day | ECoG-3-0722.mat | ECoG-3-0722.mat | 44 | accuracy | 0.824742 |
| classification | transformer | Visual Grating | single_day | ECoG-3-0722.mat | ECoG-3-0722.mat | 46 | accuracy | 0.865979 |
| classification | transformer | Visual Grating | single_day | ECoG-3-0722.mat | ECoG-3-0722.mat | 46 | accuracy | 0.804124 |
| classification | transformer | Visual Grating | single_day | ECoG-3-0722.mat | ECoG-3-0722.mat | 54 | accuracy | 0.793814 |
| classification | transformer | Visual Grating | single_day | ECoG-3-0805.mat | ECoG-3-0805.mat | 44 | accuracy | 0.646154 |
| classification | transformer | Visual Grating | single_day | ECoG-3-0805.mat | ECoG-3-0805.mat | 44 | accuracy | 0.676923 |
| classification | transformer | Visual Grating | single_day | ECoG-3-0805.mat | ECoG-3-0805.mat | 46 | accuracy | 0.707692 |
| classification | transformer | Visual Grating | single_day | ECoG-3-0805.mat | ECoG-3-0805.mat | 46 | accuracy | 0.615385 |
| classification | transformer | Visual Grating | single_day | ECoG-3-0805.mat | ECoG-3-0805.mat | 54 | accuracy | 0.584615 |
| classification | transformer | Visual Grating | single_day | ECoG-3-0805.mat | ECoG-3-0805.mat | 54 | accuracy | 0.615385 |
| classification | lfads | Visual Grating | single_day | ECoG-1-1015.mat | ECoG-1-1015.mat | 1 | accuracy | 0.476923 |
| classification | lfads | Visual Grating | single_day | ECoG-1-1015.mat | ECoG-1-1015.mat | 2 | accuracy | 0.6 |
| classification | lfads | Visual Grating | single_day | ECoG-1-1015.mat | ECoG-1-1015.mat | 3 | accuracy | 0.415385 |
| classification | lfads | Visual Grating | single_day | ECoG-1-1015.mat | ECoG-1-1015.mat | 4 | accuracy | 0.6 |
| classification | lfads | Visual Grating | single_day | ECoG-1-1015.mat | ECoG-1-1015.mat | 44 | accuracy | 0.615385 |
| classification | lfads | Visual Grating | single_day | ECoG-1-1015.mat | ECoG-1-1015.mat | 44 | accuracy | 0.523077 |
| classification | lfads | Visual Grating | single_day | ECoG-1-1015.mat | ECoG-1-1015.mat | 46 | accuracy | 0.476923 |
| classification | lfads | Visual Grating | single_day | ECoG-1-1015.mat | ECoG-1-1015.mat | 46 | accuracy | 0.461538 |
| classification | lfads | Visual Grating | single_day | ECoG-1-1015.mat | ECoG-1-1015.mat | 54 | accuracy | 0.507692 |

| | | | | | | | | |
|---|---|---|---|---|---|---|---|---|
| classification | lfads | Visual Grating | single_day | ECoG-1-1015.mat | ECoG-1-1015.mat | 54 | accuracy | 0.476923 |
| classification | lfads | Visual Grating | single_day | ECoG-1-1016.mat | ECoG-1-1016.mat | 1 | accuracy | 0.507692 |
| classification | lfads | Visual Grating | single_day | ECoG-1-1016.mat | ECoG-1-1016.mat | 2 | accuracy | 0.615385 |
| classification | lfads | Visual Grating | single_day | ECoG-1-1016.mat | ECoG-1-1016.mat | 3 | accuracy | 0.492308 |
| classification | lfads | Visual Grating | single_day | ECoG-1-1016.mat | ECoG-1-1016.mat | 4 | accuracy | 0.430769 |
| classification | lfads | Visual Grating | single_day | ECoG-1-1016.mat | ECoG-1-1016.mat | 44 | accuracy | 0.584615 |
| classification | lfads | Visual Grating | single_day | ECoG-1-1016.mat | ECoG-1-1016.mat | 44 | accuracy | 0.630769 |
| classification | lfads | Visual Grating | single_day | ECoG-1-1016.mat | ECoG-1-1016.mat | 46 | accuracy | 0.646154 |
| classification | lfads | Visual Grating | single_day | ECoG-1-1016.mat | ECoG-1-1016.mat | 46 | accuracy | 0.615385 |
| classification | lfads | Visual Grating | single_day | ECoG-1-1016.mat | ECoG-1-1016.mat | 54 | accuracy | 0.523077 |
| classification | lfads | Visual Grating | single_day | ECoG-1-1016.mat | ECoG-1-1016.mat | 54 | accuracy | 0.553846 |
| classification | lfads | Visual Grating | single_day | ECoG-1-1019.mat | ECoG-1-1019.mat | 1 | accuracy | 0.507692 |
| classification | lfads | Visual Grating | single_day | ECoG-1-1019.mat | ECoG-1-1019.mat | 2 | accuracy | 0.569231 |
| classification | lfads | Visual Grating | single_day | ECoG-1-1019.mat | ECoG-1-1019.mat | 3 | accuracy | 0.584615 |
| classification | lfads | Visual Grating | single_day | ECoG-1-1019.mat | ECoG-1-1019.mat | 4 | accuracy | 0.646154 |
| classification | lfads | Visual Grating | single_day | ECoG-1-1019.mat | ECoG-1-1019.mat | 44 | accuracy | 0.553846 |
| classification | lfads | Visual Grating | single_day | ECoG-1-1019.mat | ECoG-1-1019.mat | 44 | accuracy | 0.538462 |
| classification | lfads | Visual Grating | single_day | ECoG-1-1019.mat | ECoG-1-1019.mat | 46 | accuracy | 0.6 |
| classification | lfads | Visual Grating | single_day | ECoG-1-1019.mat | ECoG-1-1019.mat | 54 | accuracy | 0.615385 |
| classification | lfads | Visual Grating | single_day | ECoG-1-1019.mat | ECoG-1-1019.mat | 54 | accuracy | 0.523077 |
| classification | lfads | Visual Grating | single_day | ECoG-2-0723.mat | ECoG-2-0723.mat | 1 | accuracy | 0.646154 |
| classification | lfads | Visual Grating | single_day | ECoG-2-0723.mat | ECoG-2-0723.mat | 2 | accuracy | 0.661538 |
| classification | lfads | Visual Grating | single_day | ECoG-2-0723.mat | ECoG-2-0723.mat | 3 | accuracy | 0.476923 |
| classification | lfads | Visual Grating | single_day | ECoG-2-0723.mat | ECoG-2-0723.mat | 4 | accuracy | 0.661538 |
| classification | lfads | Visual Grating | single_day | ECoG-2-0723.mat | ECoG-2-0723.mat | 44 | accuracy | 0.553846 |
| classification | lfads | Visual Grating | single_day | ECoG-2-0723.mat | ECoG-2-0723.mat | 44 | accuracy | 0.538462 |
| classification | lfads | Visual Grating | single_day | ECoG-2-0723.mat | ECoG-2-0723.mat | 46 | accuracy | 0.492308 |
| classification | lfads | Visual Grating | single_day | ECoG-2-0723.mat | ECoG-2-0723.mat | 46 | accuracy | 0.492308 |
| classification | lfads | Visual Grating | single_day | ECoG-2-0723.mat | ECoG-2-0723.mat | 54 | accuracy | 0.569231 |
| classification | lfads | Visual Grating | single_day | ECoG-2-0723.mat | ECoG-2-0723.mat | 54 | accuracy | 0.569231 |
| classification | lfads | Visual Grating | single_day | ECoG-2-0802.mat | ECoG-2-0802.mat | 1 | accuracy | 0.492308 |
| classification | lfads | Visual Grating | single_day | ECoG-2-0802.mat | ECoG-2-0802.mat | 2 | accuracy | 0.492308 |
| classification | lfads | Visual Grating | single_day | ECoG-2-0802.mat | ECoG-2-0802.mat | 3 | accuracy | 0.553846 |
| classification | lfads | Visual Grating | single_day | ECoG-2-0802.mat | ECoG-2-0802.mat | 4 | accuracy | 0.430769 |
| classification | lfads | Visual Grating | single_day | ECoG-2-0802.mat | ECoG-2-0802.mat | 44 | accuracy | 0.630769 |
| classification | lfads | Visual Grating | single_day | ECoG-2-0802.mat | ECoG-2-0802.mat | 44 | accuracy | 0.615385 |
| classification | lfads | Visual Grating | single_day | ECoG-2-0802.mat | ECoG-2-0802.mat | 46 | accuracy | 0.538462 |
| classification | lfads | Visual Grating | single_day | ECoG-2-0802.mat | ECoG-2-0802.mat | 46 | accuracy | 0.446154 |
| classification | lfads | Visual Grating | single_day | ECoG-2-0802.mat | ECoG-2-0802.mat | 54 | accuracy | 0.630769 |
| classification | lfads | Visual Grating | single_day | ECoG-2-0802.mat | ECoG-2-0802.mat | 54 | accuracy | 0.538462 |
| classification | lfads | Visual Grating | single_day | ECoG-2-0803.mat | ECoG-2-0803.mat | 1 | accuracy | 0.584615 |
| classification | lfads | Visual Grating | single_day | ECoG-2-0803.mat | ECoG-2-0803.mat | 2 | accuracy | 0.615385 |
| classification | lfads | Visual Grating | single_day | ECoG-2-0803.mat | ECoG-2-0803.mat | 3 | accuracy | 0.615385 |
| classification | lfads | Visual Grating | single_day | ECoG-2-0803.mat | ECoG-2-0803.mat | 4 | accuracy | 0.569231 |
| classification | lfads | Visual Grating | single_day | ECoG-2-0803.mat | ECoG-2-0803.mat | 44 | accuracy | 0.707692 |
| classification | lfads | Visual Grating | single_day | ECoG-2-0803.mat | ECoG-2-0803.mat | 44 | accuracy | 0.676923 |
| classification | lfads | Visual Grating | single_day | ECoG-2-0803.mat | ECoG-2-0803.mat | 46 | accuracy | 0.646154 |
| classification | lfads | Visual Grating | single_day | ECoG-2-0803.mat | ECoG-2-0803.mat | 46 | accuracy | 0.6 |
| classification | lfads | Visual Grating | single_day | ECoG-2-0803.mat | ECoG-2-0803.mat | 54 | accuracy | 0.661538 |
| classification | lfads | Visual Grating | single_day | ECoG-2-0803.mat | ECoG-2-0803.mat | 54 | accuracy | 0.6 |
| classification | lfads | Visual Grating | single_day | ECoG-2-0804.mat | ECoG-2-0804.mat | 1 | accuracy | 0.769231 |
| classification | lfads | Visual Grating | single_day | ECoG-2-0804.mat | ECoG-2-0804.mat | 2 | accuracy | 0.630769 |
| classification | lfads | Visual Grating | single_day | ECoG-2-0804.mat | ECoG-2-0804.mat | 3 | accuracy | 0.646154 |
| classification | lfads | Visual Grating | single_day | ECoG-2-0804.mat | ECoG-2-0804.mat | 4 | accuracy | 0.615385 |
| classification | lfads | Visual Grating | single_day | ECoG-2-0804.mat | ECoG-2-0804.mat | 44 | accuracy | 0.707692 |
| classification | lfads | Visual Grating | single_day | ECoG-2-0804.mat | ECoG-2-0804.mat | 44 | accuracy | 0.676923 |
| classification | lfads | Visual Grating | single_day | ECoG-2-0804.mat | ECoG-2-0804.mat | 46 | accuracy | 0.661538 |
| classification | lfads | Visual Grating | single_day | ECoG-2-0804.mat | ECoG-2-0804.mat | 46 | accuracy | 0.646154 |
| classification | lfads | Visual Grating | single_day | ECoG-2-0804.mat | ECoG-2-0804.mat | 54 | accuracy | 0.661538 |
| classification | lfads | Visual Grating | single_day | ECoG-2-0804.mat | ECoG-2-0804.mat | 54 | accuracy | 0.676923 |
| classification | lfads | Visual Grating | single_day | ECoG-2-0806.mat | ECoG-2-0806.mat | 1 | accuracy | 0.553846 |
| classification | lfads | Visual Grating | single_day | ECoG-2-0806.mat | ECoG-2-0806.mat | 2 | accuracy | 0.523077 |
| classification | lfads | Visual Grating | single_day | ECoG-2-0806.mat | ECoG-2-0806.mat | 3 | accuracy | 0.492308 |

| | | | | | | | | |
|---|---|---|---|---|---|---|---|---|
| classification | lfads | Visual Grating | single_day | ECoG-2-0806.mat | ECoG-2-0806.mat | 4 | accuracy | 0.646154 |
| classification | lfads | Visual Grating | single_day | ECoG-2-0806.mat | ECoG-2-0806.mat | 44 | accuracy | 0.446154 |
| classification | lfads | Visual Grating | single_day | ECoG-2-0806.mat | ECoG-2-0806.mat | 44 | accuracy | 0.461538 |
| classification | lfads | Visual Grating | single_day | ECoG-2-0806.mat | ECoG-2-0806.mat | 46 | accuracy | 0.630769 |
| classification | lfads | Visual Grating | single_day | ECoG-2-0806.mat | ECoG-2-0806.mat | 46 | accuracy | 0.523077 |
| classification | lfads | Visual Grating | single_day | ECoG-2-0806.mat | ECoG-2-0806.mat | 54 | accuracy | 0.492308 |
| classification | lfads | Visual Grating | single_day | ECoG-2-0806.mat | ECoG-2-0806.mat | 54 | accuracy | 0.553846 |
| classification | lfads | Visual Grating | single_day | ECoG-3-0722.mat | ECoG-3-0722.mat | 1 | accuracy | 0.597938 |
| classification | lfads | Visual Grating | single_day | ECoG-3-0722.mat | ECoG-3-0722.mat | 2 | accuracy | 0.649485 |
| classification | lfads | Visual Grating | single_day | ECoG-3-0722.mat | ECoG-3-0722.mat | 3 | accuracy | 0.670103 |
| classification | lfads | Visual Grating | single_day | ECoG-3-0722.mat | ECoG-3-0722.mat | 4 | accuracy | 0.752577 |
| classification | lfads | Visual Grating | single_day | ECoG-3-0722.mat | ECoG-3-0722.mat | 44 | accuracy | 0.639175 |
| classification | lfads | Visual Grating | single_day | ECoG-3-0722.mat | ECoG-3-0722.mat | 44 | accuracy | 0.680412 |
| classification | lfads | Visual Grating | single_day | ECoG-3-0722.mat | ECoG-3-0722.mat | 46 | accuracy | 0.57732 |
| classification | lfads | Visual Grating | single_day | ECoG-3-0722.mat | ECoG-3-0722.mat | 46 | accuracy | 0.659794 |
| classification | lfads | Visual Grating | single_day | ECoG-3-0722.mat | ECoG-3-0722.mat | 54 | accuracy | 0.721649 |
| classification | lfads | Visual Grating | single_day | ECoG-3-0722.mat | ECoG-3-0722.mat | 54 | accuracy | 0.639175 |
| classification | lfads | Visual Grating | single_day | ECoG-3-0805.mat | ECoG-3-0805.mat | 1 | accuracy | 0.692308 |
| classification | lfads | Visual Grating | single_day | ECoG-3-0805.mat | ECoG-3-0805.mat | 2 | accuracy | 0.615385 |
| classification | lfads | Visual Grating | single_day | ECoG-3-0805.mat | ECoG-3-0805.mat | 3 | accuracy | 0.523077 |
| classification | lfads | Visual Grating | single_day | ECoG-3-0805.mat | ECoG-3-0805.mat | 4 | accuracy | 0.723077 |
| classification | lfads | Visual Grating | single_day | ECoG-3-0805.mat | ECoG-3-0805.mat | 44 | accuracy | 0.646154 |
| classification | lfads | Visual Grating | single_day | ECoG-3-0805.mat | ECoG-3-0805.mat | 44 | accuracy | 0.6 |
| classification | lfads | Visual Grating | single_day | ECoG-3-0805.mat | ECoG-3-0805.mat | 46 | accuracy | 0.569231 |
| classification | lfads | Visual Grating | single_day | ECoG-3-0805.mat | ECoG-3-0805.mat | 46 | accuracy | 0.569231 |
| classification | lfads | Visual Grating | single_day | ECoG-3-0805.mat | ECoG-3-0805.mat | 54 | accuracy | 0.538462 |
| classification | lfads | Visual Grating | single_day | ECoG-3-0805.mat | ECoG-3-0805.mat | 54 | accuracy | 0.630769 |
| classification | cycle_gan | Visual Grating | single_day | ECoG-1-1015.mat | ECoG-1-1015.mat | 44 | accuracy | 0.784615 |
| classification | cycle_gan | Visual Grating | single_day | ECoG-1-1015.mat | ECoG-1-1015.mat | 44 | accuracy | 0.784615 |
| classification | cycle_gan | Visual Grating | single_day | ECoG-1-1015.mat | ECoG-1-1015.mat | 46 | accuracy | 0.615385 |
| classification | cycle_gan | Visual Grating | single_day | ECoG-1-1015.mat | ECoG-1-1015.mat | 46 | accuracy | 0.615385 |
| classification | cycle_gan | Visual Grating | single_day | ECoG-1-1015.mat | ECoG-1-1015.mat | 54 | accuracy | 0.815385 |
| classification | cycle_gan | Visual Grating | single_day | ECoG-1-1015.mat | ECoG-1-1015.mat | 54 | accuracy | 0.815385 |
| classification | cycle_gan | Visual Grating | single_day | ECoG-1-1016.mat | ECoG-1-1016.mat | 44 | accuracy | 0.861538 |
| classification | cycle_gan | Visual Grating | single_day | ECoG-1-1016.mat | ECoG-1-1016.mat | 44 | accuracy | 0.861538 |
| classification | cycle_gan | Visual Grating | single_day | ECoG-1-1016.mat | ECoG-1-1016.mat | 46 | accuracy | 0.892308 |
| classification | cycle_gan | Visual Grating | single_day | ECoG-1-1016.mat | ECoG-1-1016.mat | 46 | accuracy | 0.892308 |
| classification | cycle_gan | Visual Grating | single_day | ECoG-1-1016.mat | ECoG-1-1016.mat | 54 | accuracy | 0.861538 |
| classification | cycle_gan | Visual Grating | single_day | ECoG-1-1016.mat | ECoG-1-1016.mat | 54 | accuracy | 0.861538 |
| classification | cycle_gan | Visual Grating | single_day | ECoG-1-1019.mat | ECoG-1-1019.mat | 44 | accuracy | 0.738462 |
| classification | cycle_gan | Visual Grating | single_day | ECoG-1-1019.mat | ECoG-1-1019.mat | 44 | accuracy | 0.738462 |
| classification | cycle_gan | Visual Grating | single_day | ECoG-1-1019.mat | ECoG-1-1019.mat | 46 | accuracy | 0.815385 |
| classification | cycle_gan | Visual Grating | single_day | ECoG-1-1019.mat | ECoG-1-1019.mat | 46 | accuracy | 0.815385 |
| classification | cycle_gan | Visual Grating | single_day | ECoG-1-1019.mat | ECoG-1-1019.mat | 54 | accuracy | 0.753846 |
| classification | cycle_gan | Visual Grating | single_day | ECoG-1-1019.mat | ECoG-1-1019.mat | 54 | accuracy | 0.753846 |
| classification | cycle_gan | Visual Grating | single_day | ECoG-2-0723.mat | ECoG-2-0723.mat | 44 | accuracy | 0.676923 |
| classification | cycle_gan | Visual Grating | single_day | ECoG-2-0723.mat | ECoG-2-0723.mat | 44 | accuracy | 0.676923 |
| classification | cycle_gan | Visual Grating | single_day | ECoG-2-0723.mat | ECoG-2-0723.mat | 46 | accuracy | 0.738462 |
| classification | cycle_gan | Visual Grating | single_day | ECoG-2-0723.mat | ECoG-2-0723.mat | 46 | accuracy | 0.738462 |
| classification | cycle_gan | Visual Grating | single_day | ECoG-2-0723.mat | ECoG-2-0723.mat | 54 | accuracy | 0.753846 |
| classification | cycle_gan | Visual Grating | single_day | ECoG-2-0723.mat | ECoG-2-0723.mat | 54 | accuracy | 0.753846 |
| classification | cycle_gan | Visual Grating | single_day | ECoG-2-0802.mat | ECoG-2-0802.mat | 44 | accuracy | 0.661538 |
| classification | cycle_gan | Visual Grating | single_day | ECoG-2-0802.mat | ECoG-2-0802.mat | 44 | accuracy | 0.661538 |
| classification | cycle_gan | Visual Grating | single_day | ECoG-2-0802.mat | ECoG-2-0802.mat | 46 | accuracy | 0.538462 |
| classification | cycle_gan | Visual Grating | single_day | ECoG-2-0802.mat | ECoG-2-0802.mat | 46 | accuracy | 0.538462 |
| classification | cycle_gan | Visual Grating | single_day | ECoG-2-0802.mat | ECoG-2-0802.mat | 54 | accuracy | 0.646154 |
| classification | cycle_gan | Visual Grating | single_day | ECoG-2-0802.mat | ECoG-2-0802.mat | 54 | accuracy | 0.646154 |
| classification | cycle_gan | Visual Grating | single_day | ECoG-2-0803.mat | ECoG-2-0803.mat | 44 | accuracy | 0.861538 |
| classification | cycle_gan | Visual Grating | single_day | ECoG-2-0803.mat | ECoG-2-0803.mat | 44 | accuracy | 0.861538 |
| classification | cycle_gan | Visual Grating | single_day | ECoG-2-0803.mat | ECoG-2-0803.mat | 46 | accuracy | 0.692308 |
| classification | cycle_gan | Visual Grating | single_day | ECoG-2-0803.mat | ECoG-2-0803.mat | 46 | accuracy | 0.692308 |
| classification | cycle_gan | Visual Grating | single_day | ECoG-2-0803.mat | ECoG-2-0803.mat | 54 | accuracy | 0.830769 |
| classification | cycle_gan | Visual Grating | single_day | ECoG-2-0803.mat | ECoG-2-0803.mat | 54 | accuracy | 0.830769 |

| | | | | | | | | |
|---|---|---|---|---|---|---|---|---|
| classification | cycle_gan | Visual Grating | single_day | ECoG-2-0804.mat | ECoG-2-0804.mat | 44 | accuracy | 0.892308 |
| classification | cycle_gan | Visual Grating | single_day | ECoG-2-0804.mat | ECoG-2-0804.mat | 44 | accuracy | 0.892308 |
| classification | cycle_gan | Visual Grating | single_day | ECoG-2-0804.mat | ECoG-2-0804.mat | 46 | accuracy | 0.815385 |
| classification | cycle_gan | Visual Grating | single_day | ECoG-2-0804.mat | ECoG-2-0804.mat | 46 | accuracy | 0.815385 |
| classification | cycle_gan | Visual Grating | single_day | ECoG-2-0804.mat | ECoG-2-0804.mat | 54 | accuracy | 0.830769 |
| classification | cycle_gan | Visual Grating | single_day | ECoG-2-0804.mat | ECoG-2-0804.mat | 54 | accuracy | 0.830769 |
| classification | cycle_gan | Visual Grating | single_day | ECoG-2-0806.mat | ECoG-2-0806.mat | 44 | accuracy | 0.753846 |
| classification | cycle_gan | Visual Grating | single_day | ECoG-2-0806.mat | ECoG-2-0806.mat | 44 | accuracy | 0.769231 |
| classification | cycle_gan | Visual Grating | single_day | ECoG-2-0806.mat | ECoG-2-0806.mat | 46 | accuracy | 0.738462 |
| classification | cycle_gan | Visual Grating | single_day | ECoG-2-0806.mat | ECoG-2-0806.mat | 46 | accuracy | 0.738462 |
| classification | cycle_gan | Visual Grating | single_day | ECoG-2-0806.mat | ECoG-2-0806.mat | 54 | accuracy | 0.738462 |
| classification | cycle_gan | Visual Grating | single_day | ECoG-2-0806.mat | ECoG-2-0806.mat | 54 | accuracy | 0.723077 |
| classification | cycle_gan | Visual Grating | single_day | ECoG-3-0722.mat | ECoG-3-0722.mat | 44 | accuracy | 0.896907 |
| classification | cycle_gan | Visual Grating | single_day | ECoG-3-0722.mat | ECoG-3-0722.mat | 44 | accuracy | 0.896907 |
| classification | cycle_gan | Visual Grating | single_day | ECoG-3-0722.mat | ECoG-3-0722.mat | 46 | accuracy | 0.865979 |
| classification | cycle_gan | Visual Grating | single_day | ECoG-3-0722.mat | ECoG-3-0722.mat | 46 | accuracy | 0.865979 |
| classification | cycle_gan | Visual Grating | single_day | ECoG-3-0722.mat | ECoG-3-0722.mat | 54 | accuracy | 0.886598 |
| classification | cycle_gan | Visual Grating | single_day | ECoG-3-0722.mat | ECoG-3-0722.mat | 54 | accuracy | 0.886598 |
| classification | cycle_gan | Visual Grating | single_day | ECoG-3-0805.mat | ECoG-3-0805.mat | 44 | accuracy | 0.784615 |
| classification | cycle_gan | Visual Grating | single_day | ECoG-3-0805.mat | ECoG-3-0805.mat | 44 | accuracy | 0.784615 |
| classification | cycle_gan | Visual Grating | single_day | ECoG-3-0805.mat | ECoG-3-0805.mat | 46 | accuracy | 0.830769 |
| classification | cycle_gan | Visual Grating | single_day | ECoG-3-0805.mat | ECoG-3-0805.mat | 46 | accuracy | 0.830769 |
| classification | cycle_gan | Visual Grating | single_day | ECoG-3-0805.mat | ECoG-3-0805.mat | 54 | accuracy | 0.784615 |
| classification | cycle_gan | Visual Grating | single_day | ECoG-3-0805.mat | ECoG-3-0805.mat | 54 | accuracy | 0.784615 |
| classification | stabilization | Visual Grating | single_day | ECoG-1-1015.mat | ECoG-1-1015.mat | 44 | accuracy | 0.738462 |
| classification | stabilization | Visual Grating | single_day | ECoG-1-1015.mat | ECoG-1-1015.mat | 44 | accuracy | 0.738462 |
| classification | stabilization | Visual Grating | single_day | ECoG-1-1015.mat | ECoG-1-1015.mat | 46 | accuracy | 0.707692 |
| classification | stabilization | Visual Grating | single_day | ECoG-1-1015.mat | ECoG-1-1015.mat | 46 | accuracy | 0.707692 |
| classification | stabilization | Visual Grating | single_day | ECoG-1-1015.mat | ECoG-1-1015.mat | 54 | accuracy | 0.738462 |
| classification | stabilization | Visual Grating | single_day | ECoG-1-1015.mat | ECoG-1-1015.mat | 54 | accuracy | 0.738462 |
| classification | stabilization | Visual Grating | single_day | ECoG-1-1016.mat | ECoG-1-1016.mat | 44 | accuracy | 0.784615 |
| classification | stabilization | Visual Grating | single_day | ECoG-1-1016.mat | ECoG-1-1016.mat | 44 | accuracy | 0.784615 |
| classification | stabilization | Visual Grating | single_day | ECoG-1-1016.mat | ECoG-1-1016.mat | 46 | accuracy | 0.892308 |
| classification | stabilization | Visual Grating | single_day | ECoG-1-1016.mat | ECoG-1-1016.mat | 46 | accuracy | 0.892308 |
| classification | stabilization | Visual Grating | single_day | ECoG-1-1016.mat | ECoG-1-1016.mat | 54 | accuracy | 0.661538 |
| classification | stabilization | Visual Grating | single_day | ECoG-1-1016.mat | ECoG-1-1016.mat | 54 | accuracy | 0.661538 |
| classification | stabilization | Visual Grating | single_day | ECoG-1-1019.mat | ECoG-1-1019.mat | 44 | accuracy | 0.8 |
| classification | stabilization | Visual Grating | single_day | ECoG-1-1019.mat | ECoG-1-1019.mat | 46 | accuracy | 0.784615 |
| classification | stabilization | Visual Grating | single_day | ECoG-1-1019.mat | ECoG-1-1019.mat | 46 | accuracy | 0.784615 |
| classification | stabilization | Visual Grating | single_day | ECoG-1-1019.mat | ECoG-1-1019.mat | 54 | accuracy | 0.723077 |
| classification | stabilization | Visual Grating | single_day | ECoG-1-1019.mat | ECoG-1-1019.mat | 54 | accuracy | 0.723077 |
| classification | stabilization | Visual Grating | single_day | ECoG-2-0723.mat | ECoG-2-0723.mat | 44 | accuracy | 0.692308 |
| classification | stabilization | Visual Grating | single_day | ECoG-2-0723.mat | ECoG-2-0723.mat | 44 | accuracy | 0.692308 |
| classification | stabilization | Visual Grating | single_day | ECoG-2-0723.mat | ECoG-2-0723.mat | 46 | accuracy | 0.830769 |
| classification | stabilization | Visual Grating | single_day | ECoG-2-0723.mat | ECoG-2-0723.mat | 46 | accuracy | 0.830769 |
| classification | stabilization | Visual Grating | single_day | ECoG-2-0723.mat | ECoG-2-0723.mat | 54 | accuracy | 0.707692 |
| classification | stabilization | Visual Grating | single_day | ECoG-2-0723.mat | ECoG-2-0723.mat | 54 | accuracy | 0.707692 |
| classification | stabilization | Visual Grating | single_day | ECoG-2-0802.mat | ECoG-2-0802.mat | 44 | accuracy | 0.738462 |
| classification | stabilization | Visual Grating | single_day | ECoG-2-0802.mat | ECoG-2-0802.mat | 44 | accuracy | 0.738462 |
| classification | stabilization | Visual Grating | single_day | ECoG-2-0802.mat | ECoG-2-0802.mat | 46 | accuracy | 0.661538 |
| classification | stabilization | Visual Grating | single_day | ECoG-2-0802.mat | ECoG-2-0802.mat | 46 | accuracy | 0.661538 |
| classification | stabilization | Visual Grating | single_day | ECoG-2-0802.mat | ECoG-2-0802.mat | 54 | accuracy | 0.676923 |
| classification | stabilization | Visual Grating | single_day | ECoG-2-0802.mat | ECoG-2-0802.mat | 54 | accuracy | 0.676923 |
| classification | stabilization | Visual Grating | single_day | ECoG-2-0803.mat | ECoG-2-0803.mat | 44 | accuracy | 0.815385 |
| classification | stabilization | Visual Grating | single_day | ECoG-2-0803.mat | ECoG-2-0803.mat | 44 | accuracy | 0.815385 |
| classification | stabilization | Visual Grating | single_day | ECoG-2-0803.mat | ECoG-2-0803.mat | 46 | accuracy | 0.692308 |
| classification | stabilization | Visual Grating | single_day | ECoG-2-0803.mat | ECoG-2-0803.mat | 46 | accuracy | 0.692308 |
| classification | stabilization | Visual Grating | single_day | ECoG-2-0803.mat | ECoG-2-0803.mat | 54 | accuracy | 0.8 |
| classification | stabilization | Visual Grating | single_day | ECoG-2-0804.mat | ECoG-2-0804.mat | 44 | accuracy | 0.815385 |
| classification | stabilization | Visual Grating | single_day | ECoG-2-0804.mat | ECoG-2-0804.mat | 44 | accuracy | 0.815385 |
| classification | stabilization | Visual Grating | single_day | ECoG-2-0804.mat | ECoG-2-0804.mat | 46 | accuracy | 0.784615 |
| classification | stabilization | Visual Grating | single_day | ECoG-2-0804.mat | ECoG-2-0804.mat | 46 | accuracy | 0.784615 |
| classification | stabilization | Visual Grating | single_day | ECoG-2-0804.mat | ECoG-2-0804.mat | 54 | accuracy | 0.769231 |

| | | | | | | | | |
|---|---|---|---|---|---|---|---|---|
| classification | stabilization | Visual Grating | single_day | ECoG-2-0804.mat | ECoG-2-0804.mat | 54 | accuracy | 0.769231 |
| classification | stabilization | Visual Grating | single_day | ECoG-2-0806.mat | ECoG-2-0806.mat | 44 | accuracy | 0.661538 |
| classification | stabilization | Visual Grating | single_day | ECoG-2-0806.mat | ECoG-2-0806.mat | 44 | accuracy | 0.661538 |
| classification | stabilization | Visual Grating | single_day | ECoG-2-0806.mat | ECoG-2-0806.mat | 46 | accuracy | 0.738462 |
| classification | stabilization | Visual Grating | single_day | ECoG-2-0806.mat | ECoG-2-0806.mat | 46 | accuracy | 0.738462 |
| classification | stabilization | Visual Grating | single_day | ECoG-2-0806.mat | ECoG-2-0806.mat | 54 | accuracy | 0.753846 |
| classification | stabilization | Visual Grating | single_day | ECoG-2-0806.mat | ECoG-2-0806.mat | 54 | accuracy | 0.753846 |
| classification | stabilization | Visual Grating | single_day | ECoG-3-0722.mat | ECoG-3-0722.mat | 44 | accuracy | 0.845361 |
| classification | stabilization | Visual Grating | single_day | ECoG-3-0722.mat | ECoG-3-0722.mat | 44 | accuracy | 0.845361 |
| classification | stabilization | Visual Grating | single_day | ECoG-3-0722.mat | ECoG-3-0722.mat | 46 | accuracy | 0.835052 |
| classification | stabilization | Visual Grating | single_day | ECoG-3-0722.mat | ECoG-3-0722.mat | 46 | accuracy | 0.835052 |
| classification | stabilization | Visual Grating | single_day | ECoG-3-0722.mat | ECoG-3-0722.mat | 54 | accuracy | 0.886598 |
| classification | stabilization | Visual Grating | single_day | ECoG-3-0722.mat | ECoG-3-0722.mat | 54 | accuracy | 0.886598 |
| classification | stabilization | Visual Grating | single_day | ECoG-3-0805.mat | ECoG-3-0805.mat | 44 | accuracy | 0.753846 |
| classification | stabilization | Visual Grating | single_day | ECoG-3-0805.mat | ECoG-3-0805.mat | 44 | accuracy | 0.753846 |
| classification | stabilization | Visual Grating | single_day | ECoG-3-0805.mat | ECoG-3-0805.mat | 46 | accuracy | 0.769231 |
| classification | stabilization | Visual Grating | single_day | ECoG-3-0805.mat | ECoG-3-0805.mat | 46 | accuracy | 0.769231 |
| classification | stabilization | Visual Grating | single_day | ECoG-3-0805.mat | ECoG-3-0805.mat | 54 | accuracy | 0.723077 |
| classification | stabilization | Visual Grating | single_day | ECoG-3-0805.mat | ECoG-3-0805.mat | 54 | accuracy | 0.723077 |
| classification | mfsnn | Visual Grating | single_day | ECoG-1-1015.mat | ECoG-1-1015.mat | 44 | accuracy | 0.492308 |
| classification | mfsnn | Visual Grating | single_day | ECoG-1-1015.mat | ECoG-1-1015.mat | 46 | accuracy | 0.507692 |
| classification | mfsnn | Visual Grating | single_day | ECoG-1-1015.mat | ECoG-1-1015.mat | 54 | accuracy | 0.507692 |
| classification | mfsnn | Visual Grating | single_day | ECoG-1-1016.mat | ECoG-1-1016.mat | 44 | accuracy | 0.492308 |
| classification | mfsnn | Visual Grating | single_day | ECoG-1-1016.mat | ECoG-1-1016.mat | 46 | accuracy | 0.492308 |
| classification | mfsnn | Visual Grating | single_day | ECoG-1-1016.mat | ECoG-1-1016.mat | 54 | accuracy | 0.507692 |
| classification | mfsnn | Visual Grating | single_day | ECoG-1-1019.mat | ECoG-1-1019.mat | 44 | accuracy | 0.492308 |
| classification | mfsnn | Visual Grating | single_day | ECoG-1-1019.mat | ECoG-1-1019.mat | 46 | accuracy | 0.492308 |
| classification | mfsnn | Visual Grating | single_day | ECoG-1-1019.mat | ECoG-1-1019.mat | 54 | accuracy | 0.507692 |
| classification | mfsnn | Visual Grating | single_day | ECoG-2-0723.mat | ECoG-2-0723.mat | 44 | accuracy | 0.507692 |
| classification | mfsnn | Visual Grating | single_day | ECoG-2-0723.mat | ECoG-2-0723.mat | 46 | accuracy | 0.507692 |
| classification | mfsnn | Visual Grating | single_day | ECoG-2-0723.mat | ECoG-2-0723.mat | 54 | accuracy | 0.492308 |
| classification | mfsnn | Visual Grating | single_day | ECoG-2-0802.mat | ECoG-2-0802.mat | 44 | accuracy | 0.492308 |
| classification | mfsnn | Visual Grating | single_day | ECoG-2-0802.mat | ECoG-2-0802.mat | 46 | accuracy | 0.492308 |
| classification | mfsnn | Visual Grating | single_day | ECoG-2-0802.mat | ECoG-2-0802.mat | 54 | accuracy | 0.492308 |
| classification | mfsnn | Visual Grating | single_day | ECoG-2-0803.mat | ECoG-2-0803.mat | 44 | accuracy | 0.492308 |
| classification | mfsnn | Visual Grating | single_day | ECoG-2-0803.mat | ECoG-2-0803.mat | 46 | accuracy | 0.507692 |
| classification | mfsnn | Visual Grating | single_day | ECoG-2-0803.mat | ECoG-2-0803.mat | 54 | accuracy | 0.507692 |
| classification | mfsnn | Visual Grating | single_day | ECoG-2-0804.mat | ECoG-2-0804.mat | 44 | accuracy | 0.507692 |
| classification | mfsnn | Visual Grating | single_day | ECoG-2-0804.mat | ECoG-2-0804.mat | 46 | accuracy | 0.507692 |
| classification | mfsnn | Visual Grating | single_day | ECoG-2-0804.mat | ECoG-2-0804.mat | 54 | accuracy | 0.492308 |
| classification | mfsnn | Visual Grating | single_day | ECoG-2-0806.mat | ECoG-2-0806.mat | 44 | accuracy | 0.492308 |
| classification | mfsnn | Visual Grating | single_day | ECoG-2-0806.mat | ECoG-2-0806.mat | 46 | accuracy | 0.507692 |
| classification | mfsnn | Visual Grating | single_day | ECoG-2-0806.mat | ECoG-2-0806.mat | 54 | accuracy | 0.507692 |
| classification | mfsnn | Visual Grating | single_day | ECoG-3-0722.mat | ECoG-3-0722.mat | 44 | accuracy | 0.505155 |
| classification | mfsnn | Visual Grating | single_day | ECoG-3-0722.mat | ECoG-3-0722.mat | 46 | accuracy | 0.494845 |
| classification | mfsnn | Visual Grating | single_day | ECoG-3-0722.mat | ECoG-3-0722.mat | 54 | accuracy | 0.494845 |
| classification | mfsnn | Visual Grating | single_day | ECoG-3-0805.mat | ECoG-3-0805.mat | 44 | accuracy | 0.553846 |
| classification | mfsnn | Visual Grating | single_day | ECoG-3-0805.mat | ECoG-3-0805.mat | 46 | accuracy | 0.492308 |
| classification | mfsnn | Visual Grating | single_day | ECoG-3-0805.mat | ECoG-3-0805.mat | 54 | accuracy | 0.492308 |
| classification | mscformer | Visual Grating | single_day | ECoG-1-1015.mat | ECoG-1-1015.mat | 44 | accuracy | 0.692308 |
| classification | mscformer | Visual Grating | single_day | ECoG-1-1015.mat | ECoG-1-1015.mat | 46 | accuracy | 0.615385 |
| classification | mscformer | Visual Grating | single_day | ECoG-1-1015.mat | ECoG-1-1015.mat | 54 | accuracy | 0.6 |
| classification | mscformer | Visual Grating | single_day | ECoG-1-1016.mat | ECoG-1-1016.mat | 44 | accuracy | 0.707692 |
| classification | mscformer | Visual Grating | single_day | ECoG-1-1016.mat | ECoG-1-1016.mat | 46 | accuracy | 0.553846 |
| classification | mscformer | Visual Grating | single_day | ECoG-1-1016.mat | ECoG-1-1016.mat | 54 | accuracy | 0.492308 |
| classification | mscformer | Visual Grating | single_day | ECoG-1-1019.mat | ECoG-1-1019.mat | 44 | accuracy | 0.553846 |
| classification | mscformer | Visual Grating | single_day | ECoG-1-1019.mat | ECoG-1-1019.mat | 46 | accuracy | 0.723077 |
| classification | mscformer | Visual Grating | single_day | ECoG-1-1019.mat | ECoG-1-1019.mat | 54 | accuracy | 0.584615 |
| classification | mscformer | Visual Grating | single_day | ECoG-2-0723.mat | ECoG-2-0723.mat | 44 | accuracy | 0.569231 |
| classification | mscformer | Visual Grating | single_day | ECoG-2-0723.mat | ECoG-2-0723.mat | 46 | accuracy | 0.676923 |
| classification | mscformer | Visual Grating | single_day | ECoG-2-0723.mat | ECoG-2-0723.mat | 54 | accuracy | 0.569231 |
| classification | mscformer | Visual Grating | single_day | ECoG-2-0802.mat | ECoG-2-0802.mat | 44 | accuracy | 0.676923 |
| classification | mscformer | Visual Grating | single_day | ECoG-2-0802.mat | ECoG-2-0802.mat | 46 | accuracy | 0.6 |

| | | | | | | | | |
|---|---|---|---|---|---|---|---|---|
| classification | mscformer | Visual Grating | single_day | ECoG-2-0802.mat | ECoG-2-0802.mat | 54 | accuracy | 0.723077 |
| classification | mscformer | Visual Grating | single_day | ECoG-2-0803.mat | ECoG-2-0803.mat | 44 | accuracy | 0.723077 |
| classification | mscformer | Visual Grating | single_day | ECoG-2-0803.mat | ECoG-2-0803.mat | 46 | accuracy | 0.738462 |
| classification | mscformer | Visual Grating | single_day | ECoG-2-0803.mat | ECoG-2-0803.mat | 54 | accuracy | 0.830769 |
| classification | mscformer | Visual Grating | single_day | ECoG-2-0804.mat | ECoG-2-0804.mat | 44 | accuracy | 0.723077 |
| classification | mscformer | Visual Grating | single_day | ECoG-2-0804.mat | ECoG-2-0804.mat | 46 | accuracy | 0.6 |
| classification | mscformer | Visual Grating | single_day | ECoG-2-0804.mat | ECoG-2-0804.mat | 54 | accuracy | 0.738462 |
| classification | mscformer | Visual Grating | single_day | ECoG-2-0806.mat | ECoG-2-0806.mat | 44 | accuracy | 0.461538 |
| classification | mscformer | Visual Grating | single_day | ECoG-2-0806.mat | ECoG-2-0806.mat | 46 | accuracy | 0.615385 |
| classification | mscformer | Visual Grating | single_day | ECoG-2-0806.mat | ECoG-2-0806.mat | 54 | accuracy | 0.692308 |
| classification | mscformer | Visual Grating | single_day | ECoG-3-0722.mat | ECoG-3-0722.mat | 44 | accuracy | 0.876289 |
| classification | mscformer | Visual Grating | single_day | ECoG-3-0722.mat | ECoG-3-0722.mat | 46 | accuracy | 0.876289 |
| classification | mscformer | Visual Grating | single_day | ECoG-3-0722.mat | ECoG-3-0722.mat | 54 | accuracy | 0.896907 |
| classification | mscformer | Visual Grating | single_day | ECoG-3-0805.mat | ECoG-3-0805.mat | 44 | accuracy | 0.692308 |
| classification | mscformer | Visual Grating | single_day | ECoG-3-0805.mat | ECoG-3-0805.mat | 46 | accuracy | 0.707692 |
| classification | mscformer | Visual Grating | single_day | ECoG-3-0805.mat | ECoG-3-0805.mat | 54 | accuracy | 0.723077 |
| classification | nomad | Visual Grating | single_day | ECoG-1-1015.mat | ECoG-1-1015.mat | 44 | accuracy | 0.584615 |
| classification | nomad | Visual Grating | single_day | ECoG-1-1015.mat | ECoG-1-1015.mat | 46 | accuracy | 0.630769 |
| classification | nomad | Visual Grating | single_day | ECoG-1-1015.mat | ECoG-1-1015.mat | 54 | accuracy | 0.553846 |
| classification | nomad | Visual Grating | single_day | ECoG-1-1016.mat | ECoG-1-1016.mat | 44 | accuracy | 0.723077 |
| classification | nomad | Visual Grating | single_day | ECoG-1-1016.mat | ECoG-1-1016.mat | 46 | accuracy | 0.523077 |
| classification | nomad | Visual Grating | single_day | ECoG-1-1016.mat | ECoG-1-1016.mat | 54 | accuracy | 0.553846 |
| classification | nomad | Visual Grating | single_day | ECoG-1-1019.mat | ECoG-1-1019.mat | 44 | accuracy | 0.692308 |
| classification | nomad | Visual Grating | single_day | ECoG-1-1019.mat | ECoG-1-1019.mat | 46 | accuracy | 0.830769 |
| classification | nomad | Visual Grating | single_day | ECoG-1-1019.mat | ECoG-1-1019.mat | 54 | accuracy | 0.784615 |
| classification | nomad | Visual Grating | single_day | ECoG-2-0723.mat | ECoG-2-0723.mat | 44 | accuracy | 0.615385 |
| classification | nomad | Visual Grating | single_day | ECoG-2-0723.mat | ECoG-2-0723.mat | 46 | accuracy | 0.692308 |
| classification | nomad | Visual Grating | single_day | ECoG-2-0723.mat | ECoG-2-0723.mat | 54 | accuracy | 0.661538 |
| classification | nomad | Visual Grating | single_day | ECoG-2-0802.mat | ECoG-2-0802.mat | 44 | accuracy | 0.523077 |
| classification | nomad | Visual Grating | single_day | ECoG-2-0802.mat | ECoG-2-0802.mat | 46 | accuracy | 0.523077 |
| classification | nomad | Visual Grating | single_day | ECoG-2-0802.mat | ECoG-2-0802.mat | 54 | accuracy | 0.615385 |
| classification | nomad | Visual Grating | single_day | ECoG-2-0803.mat | ECoG-2-0803.mat | 44 | accuracy | 0.738462 |
| classification | nomad | Visual Grating | single_day | ECoG-2-0803.mat | ECoG-2-0803.mat | 46 | accuracy | 0.692308 |
| classification | nomad | Visual Grating | single_day | ECoG-2-0803.mat | ECoG-2-0803.mat | 54 | accuracy | 0.676923 |
| classification | nomad | Visual Grating | single_day | ECoG-2-0804.mat | ECoG-2-0804.mat | 44 | accuracy | 0.753846 |
| classification | nomad | Visual Grating | single_day | ECoG-2-0804.mat | ECoG-2-0804.mat | 46 | accuracy | 0.8 |
| classification | nomad | Visual Grating | single_day | ECoG-2-0804.mat | ECoG-2-0804.mat | 54 | accuracy | 0.615385 |
| classification | nomad | Visual Grating | single_day | ECoG-2-0806.mat | ECoG-2-0806.mat | 44 | accuracy | 0.661538 |
| classification | nomad | Visual Grating | single_day | ECoG-2-0806.mat | ECoG-2-0806.mat | 46 | accuracy | 0.507692 |
| classification | nomad | Visual Grating | single_day | ECoG-2-0806.mat | ECoG-2-0806.mat | 54 | accuracy | 0.615385 |
| classification | nomad | Visual Grating | single_day | ECoG-3-0722.mat | ECoG-3-0722.mat | 44 | accuracy | 0.876289 |
| classification | nomad | Visual Grating | single_day | ECoG-3-0722.mat | ECoG-3-0722.mat | 46 | accuracy | 0.886598 |
| classification | nomad | Visual Grating | single_day | ECoG-3-0722.mat | ECoG-3-0722.mat | 54 | accuracy | 0.886598 |
| classification | nomad | Visual Grating | single_day | ECoG-3-0805.mat | ECoG-3-0805.mat | 44 | accuracy | 0.753846 |
| classification | nomad | Visual Grating | single_day | ECoG-3-0805.mat | ECoG-3-0805.mat | 46 | accuracy | 0.692308 |
| classification | nomad | Visual Grating | single_day | ECoG-3-0805.mat | ECoG-3-0805.mat | 54 | accuracy | 0.6 |
| classification | mlp | Visual Grating | single_day | ECoG-1-1015.mat | ECoG-1-1015.mat | 44 | accuracy | 0.769231 |
| classification | mlp | Visual Grating | single_day | ECoG-1-1015.mat | ECoG-1-1015.mat | 46 | accuracy | 0.661538 |
| classification | mlp | Visual Grating | single_day | ECoG-1-1015.mat | ECoG-1-1015.mat | 54 | accuracy | 0.830769 |
| classification | mlp | Visual Grating | single_day | ECoG-1-1016.mat | ECoG-1-1016.mat | 44 | accuracy | 0.8 |
| classification | mlp | Visual Grating | single_day | ECoG-1-1016.mat | ECoG-1-1016.mat | 46 | accuracy | 0.938462 |
| classification | mlp | Visual Grating | single_day | ECoG-1-1016.mat | ECoG-1-1016.mat | 54 | accuracy | 0.738462 |
| classification | mlp | Visual Grating | single_day | ECoG-1-1019.mat | ECoG-1-1019.mat | 44 | accuracy | 0.753846 |
| classification | mlp | Visual Grating | single_day | ECoG-1-1019.mat | ECoG-1-1019.mat | 46 | accuracy | 0.830769 |
| classification | mlp | Visual Grating | single_day | ECoG-1-1019.mat | ECoG-1-1019.mat | 54 | accuracy | 0.707692 |
| classification | mlp | Visual Grating | single_day | ECoG-2-0723.mat | ECoG-2-0723.mat | 44 | accuracy | 0.738462 |
| classification | mlp | Visual Grating | single_day | ECoG-2-0723.mat | ECoG-2-0723.mat | 46 | accuracy | 0.815385 |
| classification | mlp | Visual Grating | single_day | ECoG-2-0723.mat | ECoG-2-0723.mat | 54 | accuracy | 0.692308 |
| classification | mlp | Visual Grating | single_day | ECoG-2-0802.mat | ECoG-2-0802.mat | 44 | accuracy | 0.707692 |
| classification | mlp | Visual Grating | single_day | ECoG-2-0802.mat | ECoG-2-0802.mat | 46 | accuracy | 0.676923 |
| classification | mlp | Visual Grating | single_day | ECoG-2-0802.mat | ECoG-2-0802.mat | 54 | accuracy | 0.676923 |
| classification | mlp | Visual Grating | single_day | ECoG-2-0803.mat | ECoG-2-0803.mat | 44 | accuracy | 0.876923 |
| classification | mlp | Visual Grating | single_day | ECoG-2-0803.mat | ECoG-2-0803.mat | 46 | accuracy | 0.753846 |

| | | | | | | | | |
|---|---|---|---|---|---|---|---|---|
| classification | mlp | Visual Grating | single_day | ECoG-2-0803.mat | ECoG-2-0803.mat | 54 | accuracy | 0.784615 |
| classification | mlp | Visual Grating | single_day | ECoG-2-0804.mat | ECoG-2-0804.mat | 44 | accuracy | 0.815385 |
| classification | mlp | Visual Grating | single_day | ECoG-2-0804.mat | ECoG-2-0804.mat | 46 | accuracy | 0.830769 |
| classification | mlp | Visual Grating | single_day | ECoG-2-0804.mat | ECoG-2-0804.mat | 54 | accuracy | 0.769231 |
| classification | mlp | Visual Grating | single_day | ECoG-2-0806.mat | ECoG-2-0806.mat | 44 | accuracy | 0.692308 |
| classification | mlp | Visual Grating | single_day | ECoG-2-0806.mat | ECoG-2-0806.mat | 46 | accuracy | 0.723077 |
| classification | mlp | Visual Grating | single_day | ECoG-2-0806.mat | ECoG-2-0806.mat | 54 | accuracy | 0.723077 |
| classification | mlp | Visual Grating | single_day | ECoG-3-0722.mat | ECoG-3-0722.mat | 44 | accuracy | 0.896907 |
| classification | mlp | Visual Grating | single_day | ECoG-3-0722.mat | ECoG-3-0722.mat | 46 | accuracy | 0.845361 |
| classification | mlp | Visual Grating | single_day | ECoG-3-0722.mat | ECoG-3-0722.mat | 54 | accuracy | 0.886598 |
| classification | mlp | Visual Grating | single_day | ECoG-3-0805.mat | ECoG-3-0805.mat | 44 | accuracy | 0.738462 |
| classification | mlp | Visual Grating | single_day | ECoG-3-0805.mat | ECoG-3-0805.mat | 46 | accuracy | 0.815385 |
| classification | mlp | Visual Grating | single_day | ECoG-3-0805.mat | ECoG-3-0805.mat | 54 | accuracy | 0.738462 |
| classification | lstm | Visual Grating | cross_day | ECoG-3-0722.mat__to__ECoG-2-0804.... | ECoG-2-0806.mat__to__ECoG-2-0806.... | 44 | accuracy | 0.641745 |
| classification | lstm | Visual Grating | cross_day | ECoG-3-0722.mat__to__ECoG-2-0804.... | ECoG-2-0806.mat__to__ECoG-2-0806.... | 46 | accuracy | 0.623053 |
| classification | lstm | Visual Grating | cross_day | ECoG-3-0722.mat__to__ECoG-2-0804.... | ECoG-2-0806.mat__to__ECoG-2-0806.... | 54 | accuracy | 0.657321 |
| classification | transformer | Visual Grating | cross_day | ECoG-3-0722.mat__to__ECoG-2-0804.... | ECoG-2-0806.mat__to__ECoG-2-0806.... | 44 | accuracy | 0.682243 |
| classification | transformer | Visual Grating | cross_day | ECoG-3-0722.mat__to__ECoG-2-0804.... | ECoG-2-0806.mat__to__ECoG-2-0806.... | 46 | accuracy | 0.672897 |
| classification | transformer | Visual Grating | cross_day | ECoG-3-0722.mat__to__ECoG-2-0804.... | ECoG-2-0806.mat__to__ECoG-2-0806.... | 54 | accuracy | 0.676012 |
| classification | lfads | Visual Grating | cross_day | ECoG-3-0722.mat__to__ECoG-2-0804.... | ECoG-2-0806.mat__to__ECoG-2-0806.... | 44 | accuracy | 0.588785 |
| classification | lfads | Visual Grating | cross_day | ECoG-3-0722.mat__to__ECoG-2-0804.... | ECoG-2-0806.mat__to__ECoG-2-0806.... | 54 | accuracy | 0.570093 |
| classification | cycle_gan | Visual Grating | cross_day | ECoG-3-0722.mat__to__ECoG-2-0804.... | ECoG-2-0806.mat__to__ECoG-2-0806.... | 44 | accuracy | 0.611684 |
| classification | cycle_gan | Visual Grating | cross_day | ECoG-3-0722.mat__to__ECoG-2-0804.... | ECoG-2-0806.mat__to__ECoG-2-0806.... | 46 | accuracy | 0.494845 |
| classification | cycle_gan | Visual Grating | cross_day | ECoG-3-0722.mat__to__ECoG-2-0804.... | ECoG-2-0806.mat__to__ECoG-2-0806.... | 54 | accuracy | 0.525773 |
| classification | stabilization | Visual Grating | cross_day | ECoG-3-0722.mat__to__ECoG-2-0804.... | ECoG-2-0806.mat__to__ECoG-2-0806.... | 44 | accuracy | 0.690722 |
| classification | stabilization | Visual Grating | cross_day | ECoG-3-0722.mat__to__ECoG-2-0804.... | ECoG-2-0806.mat__to__ECoG-2-0806.... | 46 | accuracy | 0.735395 |
| classification | stabilization | Visual Grating | cross_day | ECoG-3-0722.mat__to__ECoG-2-0804.... | ECoG-2-0806.mat__to__ECoG-2-0806.... | 54 | accuracy | 0.714777 |
| classification | mfsnn | Visual Grating | cross_day | ECoG-3-0722.mat__to__ECoG-2-0804.... | ECoG-2-0806.mat | 44 | accuracy | 0.498442 |
| classification | mfsnn | Visual Grating | cross_day | ECoG-3-0722.mat__to__ECoG-2-0804.... | ECoG-2-0806.mat | 46 | accuracy | 0.498442 |
| classification | mfsnn | Visual Grating | cross_day | ECoG-3-0722.mat__to__ECoG-2-0804.... | ECoG-2-0806.mat | 54 | accuracy | 0.501558 |
| classification | mscformer | Visual Grating | cross_day | ECoG-3-0722.mat__to__ECoG-2-0804.... | ECoG-2-0806.mat | 44 | accuracy | 0.753894 |
| classification | mscformer | Visual Grating | cross_day | ECoG-3-0722.mat__to__ECoG-2-0804.... | ECoG-2-0806.mat | 46 | accuracy | 0.750779 |
| classification | mscformer | Visual Grating | cross_day | ECoG-3-0722.mat__to__ECoG-2-0804.... | ECoG-2-0806.mat | 54 | accuracy | 0.722741 |
| classification | nomad | Visual Grating | cross_day | ECoG-3-0722.mat__to__ECoG-2-0804.... | ECoG-2-0806.mat | 44 | accuracy | 0.71028 |
| classification | nomad | Visual Grating | cross_day | ECoG-3-0722.mat__to__ECoG-2-0804.... | ECoG-2-0806.mat | 46 | accuracy | 0.694704 |
| classification | nomad | Visual Grating | cross_day | ECoG-3-0722.mat__to__ECoG-2-0804.... | ECoG-2-0806.mat | 54 | accuracy | 0.700935 |
| classification | mlp | Visual Grating | cross_day | ECoG-3-0722.mat__to__ECoG-2-0804.... | ECoG-2-0806.mat | 44 | accuracy | 0.76947 |
| classification | mlp | Visual Grating | cross_day | ECoG-3-0722.mat__to__ECoG-2-0804.... | ECoG-2-0806.mat | 46 | accuracy | 0.76947 |
| classification | mlp | Visual Grating | cross_day | ECoG-3-0722.mat__to__ECoG-2-0804.... | ECoG-2-0806.mat | 54 | accuracy | 0.76324 |
| classification | lstm | Visual Coding | single_day | session_715093703_dg8_VISp | session_715093703_dg8_VISp | 44 | accuracy | 0.683333 |
| classification | lstm | Visual Coding | single_day | session_715093703_dg8_VISp | session_715093703_dg8_VISp | 44 | accuracy | 0.7 |
| classification | lstm | Visual Coding | single_day | session_715093703_dg8_VISp | session_715093703_dg8_VISp | 46 | accuracy | 0.666667 |
| classification | lstm | Visual Coding | single_day | session_715093703_dg8_VISp | session_715093703_dg8_VISp | 46 | accuracy | 0.683333 |
| classification | lstm | Visual Coding | single_day | session_715093703_dg8_VISp | session_715093703_dg8_VISp | 54 | accuracy | 0.708333 |
| classification | lstm | Visual Coding | single_day | session_715093703_dg8_VISp | session_715093703_dg8_VISp | 54 | accuracy | 0.691667 |
| classification | lstm | Visual Coding | single_day | session_719161530_dg8_VISp | session_719161530_dg8_VISp | 44 | accuracy | 0.741667 |
| classification | lstm | Visual Coding | single_day | session_719161530_dg8_VISp | session_719161530_dg8_VISp | 44 | accuracy | 0.825 |
| classification | lstm | Visual Coding | single_day | session_719161530_dg8_VISp | session_719161530_dg8_VISp | 46 | accuracy | 0.816667 |
| classification | lstm | Visual Coding | single_day | session_719161530_dg8_VISp | session_719161530_dg8_VISp | 46 | accuracy | 0.808333 |
| classification | lstm | Visual Coding | single_day | session_719161530_dg8_VISp | session_719161530_dg8_VISp | 54 | accuracy | 0.791667 |
| classification | lstm | Visual Coding | single_day | session_719161530_dg8_VISp | session_719161530_dg8_VISp | 54 | accuracy | 0.766667 |
| classification | lstm | Visual Coding | single_day | session_721123822_dg8_VISp | session_721123822_dg8_VISp | 44 | accuracy | 0.741667 |
| classification | lstm | Visual Coding | single_day | session_721123822_dg8_VISp | session_721123822_dg8_VISp | 44 | accuracy | 0.725 |
| classification | lstm | Visual Coding | single_day | session_721123822_dg8_VISp | session_721123822_dg8_VISp | 46 | accuracy | 0.716667 |
| classification | lstm | Visual Coding | single_day | session_721123822_dg8_VISp | session_721123822_dg8_VISp | 46 | accuracy | 0.733333 |
| classification | lstm | Visual Coding | single_day | session_721123822_dg8_VISp | session_721123822_dg8_VISp | 54 | accuracy | 0.716667 |
| classification | lstm | Visual Coding | single_day | session_732592105_dg8_VISp | session_732592105_dg8_VISp | 44 | accuracy | 0.858333 |
| classification | lstm | Visual Coding | single_day | session_732592105_dg8_VISp | session_732592105_dg8_VISp | 44 | accuracy | 0.866667 |
| classification | lstm | Visual Coding | single_day | session_732592105_dg8_VISp | session_732592105_dg8_VISp | 46 | accuracy | 0.933333 |
| classification | lstm | Visual Coding | single_day | session_732592105_dg8_VISp | session_732592105_dg8_VISp | 46 | accuracy | 0.825 |
| classification | lstm | Visual Coding | single_day | session_732592105_dg8_VISp | session_732592105_dg8_VISp | 54 | accuracy | 0.9 |
| classification | lstm | Visual Coding | single_day | session_732592105_dg8_VISp | session_732592105_dg8_VISp | 54 | accuracy | 0.916667 |
| classification | lstm | Visual Coding | single_day | session_737581020_dg8_VISp | session_737581020_dg8_VISp | 44 | accuracy | 0.791667 |

| | | | | | | | | |
|---|---|---|---|---|---|---|---|---|
| classification | lstm | Visual Coding | single_day | session_737581020_dg8_VISp | session_737581020_dg8_VISp | 46 | accuracy | 0.741667 |
| classification | lstm | Visual Coding | single_day | session_737581020_dg8_VISp | session_737581020_dg8_VISp | 46 | accuracy | 0.816667 |
| classification | lstm | Visual Coding | single_day | session_737581020_dg8_VISp | session_737581020_dg8_VISp | 54 | accuracy | 0.816667 |
| classification | lstm | Visual Coding | single_day | session_737581020_dg8_VISp | session_737581020_dg8_VISp | 54 | accuracy | 0.783333 |
| classification | lstm | Visual Coding | single_day | session_739448407_dg8_VISp | session_739448407_dg8_VISp | 44 | accuracy | 0.605042 |
| classification | lstm | Visual Coding | single_day | session_739448407_dg8_VISp | session_739448407_dg8_VISp | 44 | accuracy | 0.621849 |
| classification | lstm | Visual Coding | single_day | session_739448407_dg8_VISp | session_739448407_dg8_VISp | 46 | accuracy | 0.621849 |
| classification | lstm | Visual Coding | single_day | session_739448407_dg8_VISp | session_739448407_dg8_VISp | 54 | accuracy | 0.529412 |
| classification | lstm | Visual Coding | single_day | session_739448407_dg8_VISp | session_739448407_dg8_VISp | 54 | accuracy | 0.521008 |
| classification | lstm | Visual Coding | single_day | session_742951821_dg8_VISp | session_742951821_dg8_VISp | 44 | accuracy | 0.65 |
| classification | lstm | Visual Coding | single_day | session_742951821_dg8_VISp | session_742951821_dg8_VISp | 46 | accuracy | 0.591667 |
| classification | lstm | Visual Coding | single_day | session_742951821_dg8_VISp | session_742951821_dg8_VISp | 54 | accuracy | 0.725 |
| classification | lstm | Visual Coding | single_day | session_742951821_dg8_VISp | session_742951821_dg8_VISp | 54 | accuracy | 0.691667 |
| classification | lstm | Visual Coding | single_day | session_743475441_dg8_VISp | session_743475441_dg8_VISp | 44 | accuracy | 0.391667 |
| classification | lstm | Visual Coding | single_day | session_743475441_dg8_VISp | session_743475441_dg8_VISp | 44 | accuracy | 0.4 |
| classification | lstm | Visual Coding | single_day | session_743475441_dg8_VISp | session_743475441_dg8_VISp | 46 | accuracy | 0.416667 |
| classification | lstm | Visual Coding | single_day | session_743475441_dg8_VISp | session_743475441_dg8_VISp | 54 | accuracy | 0.408333 |
| classification | lstm | Visual Coding | single_day | session_743475441_dg8_VISp | session_743475441_dg8_VISp | 54 | accuracy | 0.391667 |
| classification | lstm | Visual Coding | single_day | session_744228101_dg8_VISp | session_744228101_dg8_VISp | 44 | accuracy | 0.8 |
| classification | lstm | Visual Coding | single_day | session_744228101_dg8_VISp | session_744228101_dg8_VISp | 44 | accuracy | 0.766667 |
| classification | lstm | Visual Coding | single_day | session_744228101_dg8_VISp | session_744228101_dg8_VISp | 46 | accuracy | 0.825 |
| classification | lstm | Visual Coding | single_day | session_744228101_dg8_VISp | session_744228101_dg8_VISp | 46 | accuracy | 0.816667 |
| classification | lstm | Visual Coding | single_day | session_744228101_dg8_VISp | session_744228101_dg8_VISp | 54 | accuracy | 0.816667 |
| classification | lstm | Visual Coding | single_day | session_744228101_dg8_VISp | session_744228101_dg8_VISp | 54 | accuracy | 0.841667 |
| classification | lstm | Visual Coding | single_day | session_746083955_dg8_VISp | session_746083955_dg8_VISp | 44 | accuracy | 0.433333 |
| classification | lstm | Visual Coding | single_day | session_746083955_dg8_VISp | session_746083955_dg8_VISp | 44 | accuracy | 0.441667 |
| classification | lstm | Visual Coding | single_day | session_746083955_dg8_VISp | session_746083955_dg8_VISp | 46 | accuracy | 0.458333 |
| classification | lstm | Visual Coding | single_day | session_746083955_dg8_VISp | session_746083955_dg8_VISp | 46 | accuracy | 0.475 |
| classification | lstm | Visual Coding | single_day | session_746083955_dg8_VISp | session_746083955_dg8_VISp | 54 | accuracy | 0.458333 |
| classification | lstm | Visual Coding | single_day | session_746083955_dg8_VISp | session_746083955_dg8_VISp | 54 | accuracy | 0.441667 |
| classification | lstm | Visual Coding | single_day | session_750332458_dg8_VISp | session_750332458_dg8_VISp | 44 | accuracy | 0.816667 |
| classification | lstm | Visual Coding | single_day | session_750332458_dg8_VISp | session_750332458_dg8_VISp | 44 | accuracy | 0.866667 |
| classification | lstm | Visual Coding | single_day | session_750332458_dg8_VISp | session_750332458_dg8_VISp | 46 | accuracy | 0.833333 |
| classification | lstm | Visual Coding | single_day | session_750332458_dg8_VISp | session_750332458_dg8_VISp | 46 | accuracy | 0.841667 |
| classification | lstm | Visual Coding | single_day | session_750332458_dg8_VISp | session_750332458_dg8_VISp | 54 | accuracy | 0.866667 |
| classification | lstm | Visual Coding | single_day | session_750332458_dg8_VISp | session_750332458_dg8_VISp | 54 | accuracy | 0.883333 |
| classification | lstm | Visual Coding | single_day | session_750749662_dg8_VISp | session_750749662_dg8_VISp | 44 | accuracy | 0.866667 |
| classification | lstm | Visual Coding | single_day | session_750749662_dg8_VISp | session_750749662_dg8_VISp | 44 | accuracy | 0.816667 |
| classification | lstm | Visual Coding | single_day | session_750749662_dg8_VISp | session_750749662_dg8_VISp | 46 | accuracy | 0.783333 |
| classification | lstm | Visual Coding | single_day | session_750749662_dg8_VISp | session_750749662_dg8_VISp | 46 | accuracy | 0.791667 |
| classification | lstm | Visual Coding | single_day | session_750749662_dg8_VISp | session_750749662_dg8_VISp | 54 | accuracy | 0.808333 |
| classification | lstm | Visual Coding | single_day | session_750749662_dg8_VISp | session_750749662_dg8_VISp | 54 | accuracy | 0.825 |
| classification | lstm | Visual Coding | single_day | session_751348571_dg8_VISp | session_751348571_dg8_VISp | 44 | accuracy | 0.766667 |
| classification | lstm | Visual Coding | single_day | session_751348571_dg8_VISp | session_751348571_dg8_VISp | 44 | accuracy | 0.783333 |
| classification | lstm | Visual Coding | single_day | session_751348571_dg8_VISp | session_751348571_dg8_VISp | 46 | accuracy | 0.816667 |
| classification | lstm | Visual Coding | single_day | session_751348571_dg8_VISp | session_751348571_dg8_VISp | 46 | accuracy | 0.783333 |
| classification | lstm | Visual Coding | single_day | session_751348571_dg8_VISp | session_751348571_dg8_VISp | 54 | accuracy | 0.783333 |
| classification | lstm | Visual Coding | single_day | session_751348571_dg8_VISp | session_751348571_dg8_VISp | 54 | accuracy | 0.791667 |
| classification | lstm | Visual Coding | single_day | session_754312389_dg8_VISp | session_754312389_dg8_VISp | 44 | accuracy | 0.933333 |
| classification | lstm | Visual Coding | single_day | session_754312389_dg8_VISp | session_754312389_dg8_VISp | 44 | accuracy | 0.95 |
| classification | lstm | Visual Coding | single_day | session_754312389_dg8_VISp | session_754312389_dg8_VISp | 46 | accuracy | 0.941667 |
| classification | lstm | Visual Coding | single_day | session_754312389_dg8_VISp | session_754312389_dg8_VISp | 46 | accuracy | 0.933333 |
| classification | lstm | Visual Coding | single_day | session_754312389_dg8_VISp | session_754312389_dg8_VISp | 54 | accuracy | 0.958333 |
| classification | lstm | Visual Coding | single_day | session_754312389_dg8_VISp | session_754312389_dg8_VISp | 54 | accuracy | 0.941667 |
| classification | lstm | Visual Coding | single_day | session_754829445_dg8_VISp | session_754829445_dg8_VISp | 44 | accuracy | 0.808333 |
| classification | lstm | Visual Coding | single_day | session_754829445_dg8_VISp | session_754829445_dg8_VISp | 44 | accuracy | 0.791667 |
| classification | lstm | Visual Coding | single_day | session_754829445_dg8_VISp | session_754829445_dg8_VISp | 46 | accuracy | 0.783333 |
| classification | lstm | Visual Coding | single_day | session_754829445_dg8_VISp | session_754829445_dg8_VISp | 54 | accuracy | 0.833333 |
| classification | lstm | Visual Coding | single_day | session_755434585_dg8_VISp | session_755434585_dg8_VISp | 44 | accuracy | 0.808333 |
| classification | lstm | Visual Coding | single_day | session_755434585_dg8_VISp | session_755434585_dg8_VISp | 44 | accuracy | 0.825 |
| classification | lstm | Visual Coding | single_day | session_755434585_dg8_VISp | session_755434585_dg8_VISp | 46 | accuracy | 0.75 |
| classification | lstm | Visual Coding | single_day | session_755434585_dg8_VISp | session_755434585_dg8_VISp | 46 | accuracy | 0.741667 |
| classification | lstm | Visual Coding | single_day | session_755434585_dg8_VISp | session_755434585_dg8_VISp | 54 | accuracy | 0.816667 |

| | | | | | | | | |
|---|---|---|---|---|---|---|---|---|
| classification | lstm | Visual Coding | single_day | session_755434585_dg8_VISp | session_755434585_dg8_VISp | 54 | accuracy | 0.825 |
| classification | lstm | Visual Coding | single_day | session_756029989_dg8_VISp | session_756029989_dg8_VISp | 44 | accuracy | 0.875 |
| classification | lstm | Visual Coding | single_day | session_756029989_dg8_VISp | session_756029989_dg8_VISp | 46 | accuracy | 0.85 |
| classification | lstm | Visual Coding | single_day | session_756029989_dg8_VISp | session_756029989_dg8_VISp | 54 | accuracy | 0.841667 |
| classification | lstm | Visual Coding | single_day | session_756029989_dg8_VISp | session_756029989_dg8_VISp | 54 | accuracy | 0.858333 |
| classification | lstm | Visual Coding | single_day | session_757216464_dg8_VISp | session_757216464_dg8_VISp | 44 | accuracy | 0.808333 |
| classification | lstm | Visual Coding | single_day | session_757216464_dg8_VISp | session_757216464_dg8_VISp | 44 | accuracy | 0.858333 |
| classification | lstm | Visual Coding | single_day | session_757216464_dg8_VISp | session_757216464_dg8_VISp | 46 | accuracy | 0.841667 |
| classification | lstm | Visual Coding | single_day | session_757216464_dg8_VISp | session_757216464_dg8_VISp | 46 | accuracy | 0.85 |
| classification | lstm | Visual Coding | single_day | session_757216464_dg8_VISp | session_757216464_dg8_VISp | 54 | accuracy | 0.808333 |
| classification | lstm | Visual Coding | single_day | session_757216464_dg8_VISp | session_757216464_dg8_VISp | 54 | accuracy | 0.775 |
| classification | lstm | Visual Coding | single_day | session_757970808_dg8_VISp | session_757970808_dg8_VISp | 44 | accuracy | 0.808333 |
| classification | lstm | Visual Coding | single_day | session_757970808_dg8_VISp | session_757970808_dg8_VISp | 44 | accuracy | 0.816667 |
| classification | lstm | Visual Coding | single_day | session_757970808_dg8_VISp | session_757970808_dg8_VISp | 46 | accuracy | 0.925 |
| classification | lstm | Visual Coding | single_day | session_757970808_dg8_VISp | session_757970808_dg8_VISp | 54 | accuracy | 0.841667 |
| classification | lstm | Visual Coding | single_day | session_757970808_dg8_VISp | session_757970808_dg8_VISp | 54 | accuracy | 0.833333 |
| classification | lstm | Visual Coding | single_day | session_758798717_dg8_VISp | session_758798717_dg8_VISp | 44 | accuracy | 0.641667 |
| classification | lstm | Visual Coding | single_day | session_758798717_dg8_VISp | session_758798717_dg8_VISp | 44 | accuracy | 0.6 |
| classification | lstm | Visual Coding | single_day | session_758798717_dg8_VISp | session_758798717_dg8_VISp | 46 | accuracy | 0.6 |
| classification | lstm | Visual Coding | single_day | session_758798717_dg8_VISp | session_758798717_dg8_VISp | 46 | accuracy | 0.625 |
| classification | lstm | Visual Coding | single_day | session_758798717_dg8_VISp | session_758798717_dg8_VISp | 54 | accuracy | 0.583333 |
| classification | lstm | Visual Coding | single_day | session_758798717_dg8_VISp | session_758798717_dg8_VISp | 54 | accuracy | 0.6 |
| classification | lstm | Visual Coding | single_day | session_759883607_dg8_VISp | session_759883607_dg8_VISp | 44 | accuracy | 0.866667 |
| classification | lstm | Visual Coding | single_day | session_759883607_dg8_VISp | session_759883607_dg8_VISp | 44 | accuracy | 0.816667 |
| classification | lstm | Visual Coding | single_day | session_759883607_dg8_VISp | session_759883607_dg8_VISp | 46 | accuracy | 0.808333 |
| classification | lstm | Visual Coding | single_day | session_759883607_dg8_VISp | session_759883607_dg8_VISp | 46 | accuracy | 0.8 |
| classification | lstm | Visual Coding | single_day | session_759883607_dg8_VISp | session_759883607_dg8_VISp | 54 | accuracy | 0.758333 |
| classification | lstm | Visual Coding | single_day | session_759883607_dg8_VISp | session_759883607_dg8_VISp | 54 | accuracy | 0.75 |
| classification | lstm | Visual Coding | single_day | session_760345702_dg8_VISp | session_760345702_dg8_VISp | 44 | accuracy | 0.958333 |
| classification | lstm | Visual Coding | single_day | session_760345702_dg8_VISp | session_760345702_dg8_VISp | 44 | accuracy | 0.95 |
| classification | lstm | Visual Coding | single_day | session_760345702_dg8_VISp | session_760345702_dg8_VISp | 46 | accuracy | 0.95 |
| classification | lstm | Visual Coding | single_day | session_760345702_dg8_VISp | session_760345702_dg8_VISp | 54 | accuracy | 0.891667 |
| classification | lstm | Visual Coding | single_day | session_760345702_dg8_VISp | session_760345702_dg8_VISp | 54 | accuracy | 0.908333 |
| classification | lstm | Visual Coding | single_day | session_760693773_dg8_VISp | session_760693773_dg8_VISp | 44 | accuracy | 0.908333 |
| classification | lstm | Visual Coding | single_day | session_760693773_dg8_VISp | session_760693773_dg8_VISp | 46 | accuracy | 0.875 |
| classification | lstm | Visual Coding | single_day | session_760693773_dg8_VISp | session_760693773_dg8_VISp | 46 | accuracy | 0.891667 |
| classification | lstm | Visual Coding | single_day | session_760693773_dg8_VISp | session_760693773_dg8_VISp | 54 | accuracy | 0.9 |
| classification | lstm | Visual Coding | single_day | session_760693773_dg8_VISp | session_760693773_dg8_VISp | 54 | accuracy | 0.891667 |
| classification | lstm | Visual Coding | single_day | session_761418226_dg8_VISp | session_761418226_dg8_VISp | 44 | accuracy | 0.791667 |
| classification | lstm | Visual Coding | single_day | session_761418226_dg8_VISp | session_761418226_dg8_VISp | 44 | accuracy | 0.775 |
| classification | lstm | Visual Coding | single_day | session_761418226_dg8_VISp | session_761418226_dg8_VISp | 46 | accuracy | 0.7 |
| classification | lstm | Visual Coding | single_day | session_761418226_dg8_VISp | session_761418226_dg8_VISp | 46 | accuracy | 0.716667 |
| classification | lstm | Visual Coding | single_day | session_761418226_dg8_VISp | session_761418226_dg8_VISp | 54 | accuracy | 0.691667 |
| classification | lstm | Visual Coding | single_day | session_761418226_dg8_VISp | session_761418226_dg8_VISp | 54 | accuracy | 0.7 |
| classification | lstm | Visual Coding | single_day | session_762120172_dg8_VISp | session_762120172_dg8_VISp | 44 | accuracy | 0.916667 |
| classification | lstm | Visual Coding | single_day | session_762120172_dg8_VISp | session_762120172_dg8_VISp | 46 | accuracy | 0.875 |
| classification | lstm | Visual Coding | single_day | session_762120172_dg8_VISp | session_762120172_dg8_VISp | 46 | accuracy | 0.85 |
| classification | lstm | Visual Coding | single_day | session_762120172_dg8_VISp | session_762120172_dg8_VISp | 54 | accuracy | 0.916667 |
| classification | lstm | Visual Coding | single_day | session_762120172_dg8_VISp | session_762120172_dg8_VISp | 54 | accuracy | 0.908333 |
| classification | lstm | Visual Coding | single_day | session_762602078_dg8_VISp | session_762602078_dg8_VISp | 44 | accuracy | 0.916667 |
| classification | lstm | Visual Coding | single_day | session_762602078_dg8_VISp | session_762602078_dg8_VISp | 46 | accuracy | 0.883333 |
| classification | lstm | Visual Coding | single_day | session_762602078_dg8_VISp | session_762602078_dg8_VISp | 46 | accuracy | 0.891667 |
| classification | lstm | Visual Coding | single_day | session_762602078_dg8_VISp | session_762602078_dg8_VISp | 54 | accuracy | 0.85 |
| classification | lstm | Visual Coding | single_day | session_763673393_dg8_VISp | session_763673393_dg8_VISp | 44 | accuracy | 0.691667 |
| classification | lstm | Visual Coding | single_day | session_763673393_dg8_VISp | session_763673393_dg8_VISp | 44 | accuracy | 0.7 |
| classification | lstm | Visual Coding | single_day | session_763673393_dg8_VISp | session_763673393_dg8_VISp | 46 | accuracy | 0.741667 |
| classification | lstm | Visual Coding | single_day | session_763673393_dg8_VISp | session_763673393_dg8_VISp | 46 | accuracy | 0.758333 |
| classification | lstm | Visual Coding | single_day | session_763673393_dg8_VISp | session_763673393_dg8_VISp | 54 | accuracy | 0.691667 |
| classification | lstm | Visual Coding | single_day | session_763673393_dg8_VISp | session_763673393_dg8_VISp | 54 | accuracy | 0.708333 |
| classification | lstm | Visual Coding | single_day | session_773418906_dg8_VISp | session_773418906_dg8_VISp | 44 | accuracy | 0.758333 |
| classification | lstm | Visual Coding | single_day | session_773418906_dg8_VISp | session_773418906_dg8_VISp | 44 | accuracy | 0.725 |
| classification | lstm | Visual Coding | single_day | session_773418906_dg8_VISp | session_773418906_dg8_VISp | 46 | accuracy | 0.683333 |
| classification | lstm | Visual Coding | single_day | session_773418906_dg8_VISp | session_773418906_dg8_VISp | 46 | accuracy | 0.7 |

| | | | | | | | | |
|---|---|---|---|---|---|---|---|---|
| classification | lstm | Visual Coding | single_day | session_773418906_dg8_VISp | session_773418906_dg8_VISp | 54 | accuracy | 0.708333 |
| classification | lstm | Visual Coding | single_day | session_773418906_dg8_VISp | session_773418906_dg8_VISp | 54 | accuracy | 0.691667 |
| classification | lstm | Visual Coding | single_day | session_791319847_dg8_VISp | session_791319847_dg8_VISp | 44 | accuracy | 0.966667 |
| classification | lstm | Visual Coding | single_day | session_791319847_dg8_VISp | session_791319847_dg8_VISp | 44 | accuracy | 0.958333 |
| classification | lstm | Visual Coding | single_day | session_791319847_dg8_VISp | session_791319847_dg8_VISp | 46 | accuracy | 0.933333 |
| classification | lstm | Visual Coding | single_day | session_791319847_dg8_VISp | session_791319847_dg8_VISp | 46 | accuracy | 0.925 |
| classification | lstm | Visual Coding | single_day | session_791319847_dg8_VISp | session_791319847_dg8_VISp | 54 | accuracy | 0.925 |
| classification | lstm | Visual Coding | single_day | session_797828357_dg8_VISp | session_797828357_dg8_VISp | 44 | accuracy | 0.891667 |
| classification | lstm | Visual Coding | single_day | session_797828357_dg8_VISp | session_797828357_dg8_VISp | 44 | accuracy | 0.866667 |
| classification | lstm | Visual Coding | single_day | session_797828357_dg8_VISp | session_797828357_dg8_VISp | 46 | accuracy | 0.833333 |
| classification | lstm | Visual Coding | single_day | session_797828357_dg8_VISp | session_797828357_dg8_VISp | 46 | accuracy | 0.791667 |
| classification | lstm | Visual Coding | single_day | session_797828357_dg8_VISp | session_797828357_dg8_VISp | 54 | accuracy | 0.85 |
| classification | lstm | Visual Coding | single_day | session_797828357_dg8_VISp | session_797828357_dg8_VISp | 54 | accuracy | 0.866667 |
| classification | lstm | Visual Coding | single_day | session_798911424_dg8_VISp | session_798911424_dg8_VISp | 44 | accuracy | 0.9 |
| classification | lstm | Visual Coding | single_day | session_798911424_dg8_VISp | session_798911424_dg8_VISp | 44 | accuracy | 0.941667 |
| classification | lstm | Visual Coding | single_day | session_798911424_dg8_VISp | session_798911424_dg8_VISp | 46 | accuracy | 0.891667 |
| classification | lstm | Visual Coding | single_day | session_798911424_dg8_VISp | session_798911424_dg8_VISp | 46 | accuracy | 0.883333 |
| classification | lstm | Visual Coding | single_day | session_798911424_dg8_VISp | session_798911424_dg8_VISp | 54 | accuracy | 0.866667 |
| classification | lstm | Visual Coding | single_day | session_798911424_dg8_VISp | session_798911424_dg8_VISp | 54 | accuracy | 0.841667 |
| classification | lstm | Visual Coding | single_day | session_799864342_dg8_VISp | session_799864342_dg8_VISp | 44 | accuracy | 0.925 |
| classification | lstm | Visual Coding | single_day | session_799864342_dg8_VISp | session_799864342_dg8_VISp | 46 | accuracy | 0.958333 |
| classification | lstm | Visual Coding | single_day | session_799864342_dg8_VISp | session_799864342_dg8_VISp | 46 | accuracy | 0.975 |
| classification | lstm | Visual Coding | single_day | session_799864342_dg8_VISp | session_799864342_dg8_VISp | 54 | accuracy | 0.866667 |
| classification | lstm | Visual Coding | single_day | session_799864342_dg8_VISp | session_799864342_dg8_VISp | 54 | accuracy | 0.883333 |
| classification | transformer | Visual Coding | single_day | session_715093703_dg8_VISp | session_715093703_dg8_VISp | 44 | accuracy | 0.758333 |
| classification | transformer | Visual Coding | single_day | session_715093703_dg8_VISp | session_715093703_dg8_VISp | 44 | accuracy | 0.8 |
| classification | transformer | Visual Coding | single_day | session_715093703_dg8_VISp | session_715093703_dg8_VISp | 46 | accuracy | 0.6 |
| classification | transformer | Visual Coding | single_day | session_715093703_dg8_VISp | session_715093703_dg8_VISp | 46 | accuracy | 0.608333 |
| classification | transformer | Visual Coding | single_day | session_715093703_dg8_VISp | session_715093703_dg8_VISp | 54 | accuracy | 0.65 |
| classification | transformer | Visual Coding | single_day | session_719161530_dg8_VISp | session_719161530_dg8_VISp | 44 | accuracy | 0.8 |
| classification | transformer | Visual Coding | single_day | session_719161530_dg8_VISp | session_719161530_dg8_VISp | 46 | accuracy | 0.783333 |
| classification | transformer | Visual Coding | single_day | session_719161530_dg8_VISp | session_719161530_dg8_VISp | 46 | accuracy | 0.775 |
| classification | transformer | Visual Coding | single_day | session_719161530_dg8_VISp | session_719161530_dg8_VISp | 54 | accuracy | 0.766667 |
| classification | transformer | Visual Coding | single_day | session_719161530_dg8_VISp | session_719161530_dg8_VISp | 54 | accuracy | 0.783333 |
| classification | transformer | Visual Coding | single_day | session_721123822_dg8_VISp | session_721123822_dg8_VISp | 44 | accuracy | 0.733333 |
| classification | transformer | Visual Coding | single_day | session_721123822_dg8_VISp | session_721123822_dg8_VISp | 44 | accuracy | 0.741667 |
| classification | transformer | Visual Coding | single_day | session_721123822_dg8_VISp | session_721123822_dg8_VISp | 46 | accuracy | 0.758333 |
| classification | transformer | Visual Coding | single_day | session_721123822_dg8_VISp | session_721123822_dg8_VISp | 46 | accuracy | 0.75 |
| classification | transformer | Visual Coding | single_day | session_721123822_dg8_VISp | session_721123822_dg8_VISp | 54 | accuracy | 0.725 |
| classification | transformer | Visual Coding | single_day | session_721123822_dg8_VISp | session_721123822_dg8_VISp | 54 | accuracy | 0.716667 |
| classification | transformer | Visual Coding | single_day | session_732592105_dg8_VISp | session_732592105_dg8_VISp | 44 | accuracy | 0.833333 |
| classification | transformer | Visual Coding | single_day | session_732592105_dg8_VISp | session_732592105_dg8_VISp | 46 | accuracy | 0.816667 |
| classification | transformer | Visual Coding | single_day | session_732592105_dg8_VISp | session_732592105_dg8_VISp | 46 | accuracy | 0.875 |
| classification | transformer | Visual Coding | single_day | session_732592105_dg8_VISp | session_732592105_dg8_VISp | 54 | accuracy | 0.858333 |
| classification | transformer | Visual Coding | single_day | session_732592105_dg8_VISp | session_732592105_dg8_VISp | 54 | accuracy | 0.866667 |
| classification | transformer | Visual Coding | single_day | session_737581020_dg8_VISp | session_737581020_dg8_VISp | 44 | accuracy | 0.841667 |
| classification | transformer | Visual Coding | single_day | session_737581020_dg8_VISp | session_737581020_dg8_VISp | 44 | accuracy | 0.85 |
| classification | transformer | Visual Coding | single_day | session_737581020_dg8_VISp | session_737581020_dg8_VISp | 46 | accuracy | 0.733333 |
| classification | transformer | Visual Coding | single_day | session_737581020_dg8_VISp | session_737581020_dg8_VISp | 46 | accuracy | 0.741667 |
| classification | transformer | Visual Coding | single_day | session_737581020_dg8_VISp | session_737581020_dg8_VISp | 54 | accuracy | 0.783333 |
| classification | transformer | Visual Coding | single_day | session_737581020_dg8_VISp | session_737581020_dg8_VISp | 54 | accuracy | 0.775 |
| classification | transformer | Visual Coding | single_day | session_739448407_dg8_VISp | session_739448407_dg8_VISp | 44 | accuracy | 0.596639 |
| classification | transformer | Visual Coding | single_day | session_739448407_dg8_VISp | session_739448407_dg8_VISp | 46 | accuracy | 0.689076 |
| classification | transformer | Visual Coding | single_day | session_739448407_dg8_VISp | session_739448407_dg8_VISp | 54 | accuracy | 0.638655 |
| classification | transformer | Visual Coding | single_day | session_739448407_dg8_VISp | session_739448407_dg8_VISp | 54 | accuracy | 0.621849 |
| classification | transformer | Visual Coding | single_day | session_742951821_dg8_VISp | session_742951821_dg8_VISp | 44 | accuracy | 0.591667 |
| classification | transformer | Visual Coding | single_day | session_742951821_dg8_VISp | session_742951821_dg8_VISp | 46 | accuracy | 0.6 |
| classification | transformer | Visual Coding | single_day | session_742951821_dg8_VISp | session_742951821_dg8_VISp | 46 | accuracy | 0.625 |
| classification | transformer | Visual Coding | single_day | session_742951821_dg8_VISp | session_742951821_dg8_VISp | 54 | accuracy | 0.691667 |
| classification | transformer | Visual Coding | single_day | session_742951821_dg8_VISp | session_742951821_dg8_VISp | 54 | accuracy | 0.625 |
| classification | transformer | Visual Coding | single_day | session_743475441_dg8_VISp | session_743475441_dg8_VISp | 44 | accuracy | 0.425 |
| classification | transformer | Visual Coding | single_day | session_743475441_dg8_VISp | session_743475441_dg8_VISp | 44 | accuracy | 0.416667 |
| classification | transformer | Visual Coding | single_day | session_743475441_dg8_VISp | session_743475441_dg8_VISp | 46 | accuracy | 0.441667 |

| | | | | | | | | |
|---|---|---|---|---|---|---|---|---|
| classification | transformer | Visual Coding | single_day | session_743475441_dg8_VISp | session_743475441_dg8_VISp | 54 | accuracy | 0.425 |
| classification | transformer | Visual Coding | single_day | session_744228101_dg8_VISp | session_744228101_dg8_VISp | 44 | accuracy | 0.816667 |
| classification | transformer | Visual Coding | single_day | session_744228101_dg8_VISp | session_744228101_dg8_VISp | 44 | accuracy | 0.8 |
| classification | transformer | Visual Coding | single_day | session_744228101_dg8_VISp | session_744228101_dg8_VISp | 46 | accuracy | 0.783333 |
| classification | transformer | Visual Coding | single_day | session_744228101_dg8_VISp | session_744228101_dg8_VISp | 46 | accuracy | 0.808333 |
| classification | transformer | Visual Coding | single_day | session_744228101_dg8_VISp | session_744228101_dg8_VISp | 54 | accuracy | 0.783333 |
| classification | transformer | Visual Coding | single_day | session_744228101_dg8_VISp | session_744228101_dg8_VISp | 54 | accuracy | 0.816667 |
| classification | transformer | Visual Coding | single_day | session_746083955_dg8_VISp | session_746083955_dg8_VISp | 44 | accuracy | 0.416667 |
| classification | transformer | Visual Coding | single_day | session_746083955_dg8_VISp | session_746083955_dg8_VISp | 44 | accuracy | 0.433333 |
| classification | transformer | Visual Coding | single_day | session_746083955_dg8_VISp | session_746083955_dg8_VISp | 46 | accuracy | 0.433333 |
| classification | transformer | Visual Coding | single_day | session_746083955_dg8_VISp | session_746083955_dg8_VISp | 46 | accuracy | 0.483333 |
| classification | transformer | Visual Coding | single_day | session_746083955_dg8_VISp | session_746083955_dg8_VISp | 54 | accuracy | 0.483333 |
| classification | transformer | Visual Coding | single_day | session_746083955_dg8_VISp | session_746083955_dg8_VISp | 54 | accuracy | 0.458333 |
| classification | transformer | Visual Coding | single_day | session_750332458_dg8_VISp | session_750332458_dg8_VISp | 44 | accuracy | 0.908333 |
| classification | transformer | Visual Coding | single_day | session_750332458_dg8_VISp | session_750332458_dg8_VISp | 44 | accuracy | 0.85 |
| classification | transformer | Visual Coding | single_day | session_750332458_dg8_VISp | session_750332458_dg8_VISp | 46 | accuracy | 0.925 |
| classification | transformer | Visual Coding | single_day | session_750332458_dg8_VISp | session_750332458_dg8_VISp | 46 | accuracy | 0.941667 |
| classification | transformer | Visual Coding | single_day | session_750332458_dg8_VISp | session_750332458_dg8_VISp | 54 | accuracy | 0.891667 |
| classification | transformer | Visual Coding | single_day | session_750332458_dg8_VISp | session_750332458_dg8_VISp | 54 | accuracy | 0.925 |
| classification | transformer | Visual Coding | single_day | session_750749662_dg8_VISp | session_750749662_dg8_VISp | 44 | accuracy | 0.825 |
| classification | transformer | Visual Coding | single_day | session_750749662_dg8_VISp | session_750749662_dg8_VISp | 46 | accuracy | 0.833333 |
| classification | transformer | Visual Coding | single_day | session_750749662_dg8_VISp | session_750749662_dg8_VISp | 46 | accuracy | 0.825 |
| classification | transformer | Visual Coding | single_day | session_750749662_dg8_VISp | session_750749662_dg8_VISp | 54 | accuracy | 0.866667 |
| classification | transformer | Visual Coding | single_day | session_750749662_dg8_VISp | session_750749662_dg8_VISp | 54 | accuracy | 0.858333 |
| classification | transformer | Visual Coding | single_day | session_751348571_dg8_VISp | session_751348571_dg8_VISp | 44 | accuracy | 0.875 |
| classification | transformer | Visual Coding | single_day | session_751348571_dg8_VISp | session_751348571_dg8_VISp | 44 | accuracy | 0.858333 |
| classification | transformer | Visual Coding | single_day | session_751348571_dg8_VISp | session_751348571_dg8_VISp | 46 | accuracy | 0.883333 |
| classification | transformer | Visual Coding | single_day | session_751348571_dg8_VISp | session_751348571_dg8_VISp | 46 | accuracy | 0.875 |
| classification | transformer | Visual Coding | single_day | session_751348571_dg8_VISp | session_751348571_dg8_VISp | 54 | accuracy | 0.85 |
| classification | transformer | Visual Coding | single_day | session_751348571_dg8_VISp | session_751348571_dg8_VISp | 54 | accuracy | 0.866667 |
| classification | transformer | Visual Coding | single_day | session_754312389_dg8_VISp | session_754312389_dg8_VISp | 44 | accuracy | 0.958333 |
| classification | transformer | Visual Coding | single_day | session_754312389_dg8_VISp | session_754312389_dg8_VISp | 46 | accuracy | 0.908333 |
| classification | transformer | Visual Coding | single_day | session_754312389_dg8_VISp | session_754312389_dg8_VISp | 46 | accuracy | 0.916667 |
| classification | transformer | Visual Coding | single_day | session_754312389_dg8_VISp | session_754312389_dg8_VISp | 54 | accuracy | 0.95 |
| classification | transformer | Visual Coding | single_day | session_754312389_dg8_VISp | session_754312389_dg8_VISp | 54 | accuracy | 0.925 |
| classification | transformer | Visual Coding | single_day | session_754829445_dg8_VISp | session_754829445_dg8_VISp | 44 | accuracy | 0.791667 |
| classification | transformer | Visual Coding | single_day | session_754829445_dg8_VISp | session_754829445_dg8_VISp | 46 | accuracy | 0.708333 |
| classification | transformer | Visual Coding | single_day | session_754829445_dg8_VISp | session_754829445_dg8_VISp | 46 | accuracy | 0.758333 |
| classification | transformer | Visual Coding | single_day | session_754829445_dg8_VISp | session_754829445_dg8_VISp | 54 | accuracy | 0.841667 |
| classification | transformer | Visual Coding | single_day | session_754829445_dg8_VISp | session_754829445_dg8_VISp | 54 | accuracy | 0.833333 |
| classification | transformer | Visual Coding | single_day | session_755434585_dg8_VISp | session_755434585_dg8_VISp | 44 | accuracy | 0.8 |
| classification | transformer | Visual Coding | single_day | session_755434585_dg8_VISp | session_755434585_dg8_VISp | 44 | accuracy | 0.841667 |
| classification | transformer | Visual Coding | single_day | session_755434585_dg8_VISp | session_755434585_dg8_VISp | 46 | accuracy | 0.775 |
| classification | transformer | Visual Coding | single_day | session_755434585_dg8_VISp | session_755434585_dg8_VISp | 46 | accuracy | 0.758333 |
| classification | transformer | Visual Coding | single_day | session_755434585_dg8_VISp | session_755434585_dg8_VISp | 54 | accuracy | 0.766667 |
| classification | transformer | Visual Coding | single_day | session_755434585_dg8_VISp | session_755434585_dg8_VISp | 54 | accuracy | 0.808333 |
| classification | transformer | Visual Coding | single_day | session_756029989_dg8_VISp | session_756029989_dg8_VISp | 44 | accuracy | 0.9 |
| classification | transformer | Visual Coding | single_day | session_756029989_dg8_VISp | session_756029989_dg8_VISp | 44 | accuracy | 0.908333 |
| classification | transformer | Visual Coding | single_day | session_756029989_dg8_VISp | session_756029989_dg8_VISp | 46 | accuracy | 0.891667 |
| classification | transformer | Visual Coding | single_day | session_756029989_dg8_VISp | session_756029989_dg8_VISp | 46 | accuracy | 0.9 |
| classification | transformer | Visual Coding | single_day | session_756029989_dg8_VISp | session_756029989_dg8_VISp | 54 | accuracy | 0.883333 |
| classification | transformer | Visual Coding | single_day | session_756029989_dg8_VISp | session_756029989_dg8_VISp | 54 | accuracy | 0.841667 |
| classification | transformer | Visual Coding | single_day | session_757216464_dg8_VISp | session_757216464_dg8_VISp | 44 | accuracy | 0.8 |
| classification | transformer | Visual Coding | single_day | session_757216464_dg8_VISp | session_757216464_dg8_VISp | 44 | accuracy | 0.808333 |
| classification | transformer | Visual Coding | single_day | session_757216464_dg8_VISp | session_757216464_dg8_VISp | 46 | accuracy | 0.858333 |
| classification | transformer | Visual Coding | single_day | session_757216464_dg8_VISp | session_757216464_dg8_VISp | 46 | accuracy | 0.808333 |
| classification | transformer | Visual Coding | single_day | session_757216464_dg8_VISp | session_757216464_dg8_VISp | 54 | accuracy | 0.808333 |
| classification | transformer | Visual Coding | single_day | session_757216464_dg8_VISp | session_757216464_dg8_VISp | 54 | accuracy | 0.825 |
| classification | transformer | Visual Coding | single_day | session_757970808_dg8_VISp | session_757970808_dg8_VISp | 44 | accuracy | 0.85 |
| classification | transformer | Visual Coding | single_day | session_757970808_dg8_VISp | session_757970808_dg8_VISp | 44 | accuracy | 0.866667 |
| classification | transformer | Visual Coding | single_day | session_757970808_dg8_VISp | session_757970808_dg8_VISp | 46 | accuracy | 0.9 |
| classification | transformer | Visual Coding | single_day | session_757970808_dg8_VISp | session_757970808_dg8_VISp | 46 | accuracy | 0.875 |
| classification | transformer | Visual Coding | single_day | session_757970808_dg8_VISp | session_757970808_dg8_VISp | 54 | accuracy | 0.858333 |

| | | | | | | | | |
|---|---|---|---|---|---|---|---|---|
| classification | transformer | Visual Coding | single_day | session_757970808_dg8_VISp | session_757970808_dg8_VISp | 54 | accuracy | 0.816667 |
| classification | transformer | Visual Coding | single_day | session_758798717_dg8_VISp | session_758798717_dg8_VISp | 44 | accuracy | 0.591667 |
| classification | transformer | Visual Coding | single_day | session_758798717_dg8_VISp | session_758798717_dg8_VISp | 46 | accuracy | 0.625 |
| classification | transformer | Visual Coding | single_day | session_758798717_dg8_VISp | session_758798717_dg8_VISp | 54 | accuracy | 0.566667 |
| classification | transformer | Visual Coding | single_day | session_758798717_dg8_VISp | session_758798717_dg8_VISp | 54 | accuracy | 0.575 |
| classification | transformer | Visual Coding | single_day | session_759883607_dg8_VISp | session_759883607_dg8_VISp | 44 | accuracy | 0.85 |
| classification | transformer | Visual Coding | single_day | session_759883607_dg8_VISp | session_759883607_dg8_VISp | 44 | accuracy | 0.866667 |
| classification | transformer | Visual Coding | single_day | session_759883607_dg8_VISp | session_759883607_dg8_VISp | 46 | accuracy | 0.808333 |
| classification | transformer | Visual Coding | single_day | session_759883607_dg8_VISp | session_759883607_dg8_VISp | 46 | accuracy | 0.825 |
| classification | transformer | Visual Coding | single_day | session_759883607_dg8_VISp | session_759883607_dg8_VISp | 54 | accuracy | 0.875 |
| classification | transformer | Visual Coding | single_day | session_759883607_dg8_VISp | session_759883607_dg8_VISp | 54 | accuracy | 0.825 |
| classification | transformer | Visual Coding | single_day | session_760345702_dg8_VISp | session_760345702_dg8_VISp | 44 | accuracy | 0.958333 |
| classification | transformer | Visual Coding | single_day | session_760345702_dg8_VISp | session_760345702_dg8_VISp | 44 | accuracy | 0.908333 |
| classification | transformer | Visual Coding | single_day | session_760345702_dg8_VISp | session_760345702_dg8_VISp | 46 | accuracy | 0.9 |
| classification | transformer | Visual Coding | single_day | session_760345702_dg8_VISp | session_760345702_dg8_VISp | 54 | accuracy | 0.925 |
| classification | transformer | Visual Coding | single_day | session_760345702_dg8_VISp | session_760345702_dg8_VISp | 54 | accuracy | 0.916667 |
| classification | transformer | Visual Coding | single_day | session_760693773_dg8_VISp | session_760693773_dg8_VISp | 44 | accuracy | 0.85 |
| classification | transformer | Visual Coding | single_day | session_760693773_dg8_VISp | session_760693773_dg8_VISp | 44 | accuracy | 0.883333 |
| classification | transformer | Visual Coding | single_day | session_760693773_dg8_VISp | session_760693773_dg8_VISp | 46 | accuracy | 0.85 |
| classification | transformer | Visual Coding | single_day | session_760693773_dg8_VISp | session_760693773_dg8_VISp | 46 | accuracy | 0.858333 |
| classification | transformer | Visual Coding | single_day | session_760693773_dg8_VISp | session_760693773_dg8_VISp | 54 | accuracy | 0.841667 |
| classification | transformer | Visual Coding | single_day | session_760693773_dg8_VISp | session_760693773_dg8_VISp | 54 | accuracy | 0.85 |
| classification | transformer | Visual Coding | single_day | session_761418226_dg8_VISp | session_761418226_dg8_VISp | 44 | accuracy | 0.758333 |
| classification | transformer | Visual Coding | single_day | session_761418226_dg8_VISp | session_761418226_dg8_VISp | 44 | accuracy | 0.766667 |
| classification | transformer | Visual Coding | single_day | session_761418226_dg8_VISp | session_761418226_dg8_VISp | 46 | accuracy | 0.766667 |
| classification | transformer | Visual Coding | single_day | session_761418226_dg8_VISp | session_761418226_dg8_VISp | 54 | accuracy | 0.775 |
| classification | transformer | Visual Coding | single_day | session_761418226_dg8_VISp | session_761418226_dg8_VISp | 54 | accuracy | 0.791667 |
| classification | transformer | Visual Coding | single_day | session_762120172_dg8_VISp | session_762120172_dg8_VISp | 44 | accuracy | 0.916667 |
| classification | transformer | Visual Coding | single_day | session_762120172_dg8_VISp | session_762120172_dg8_VISp | 44 | accuracy | 0.908333 |
| classification | transformer | Visual Coding | single_day | session_762120172_dg8_VISp | session_762120172_dg8_VISp | 46 | accuracy | 0.916667 |
| classification | transformer | Visual Coding | single_day | session_762120172_dg8_VISp | session_762120172_dg8_VISp | 46 | accuracy | 0.933333 |
| classification | transformer | Visual Coding | single_day | session_762120172_dg8_VISp | session_762120172_dg8_VISp | 54 | accuracy | 0.925 |
| classification | transformer | Visual Coding | single_day | session_762120172_dg8_VISp | session_762120172_dg8_VISp | 54 | accuracy | 0.916667 |
| classification | transformer | Visual Coding | single_day | session_762602078_dg8_VISp | session_762602078_dg8_VISp | 44 | accuracy | 0.933333 |
| classification | transformer | Visual Coding | single_day | session_762602078_dg8_VISp | session_762602078_dg8_VISp | 46 | accuracy | 0.916667 |
| classification | transformer | Visual Coding | single_day | session_762602078_dg8_VISp | session_762602078_dg8_VISp | 54 | accuracy | 0.891667 |
| classification | transformer | Visual Coding | single_day | session_763673393_dg8_VISp | session_763673393_dg8_VISp | 44 | accuracy | 0.775 |
| classification | transformer | Visual Coding | single_day | session_763673393_dg8_VISp | session_763673393_dg8_VISp | 44 | accuracy | 0.8 |
| classification | transformer | Visual Coding | single_day | session_763673393_dg8_VISp | session_763673393_dg8_VISp | 46 | accuracy | 0.808333 |
| classification | transformer | Visual Coding | single_day | session_763673393_dg8_VISp | session_763673393_dg8_VISp | 54 | accuracy | 0.675 |
| classification | transformer | Visual Coding | single_day | session_763673393_dg8_VISp | session_763673393_dg8_VISp | 54 | accuracy | 0.708333 |
| classification | transformer | Visual Coding | single_day | session_773418906_dg8_VISp | session_773418906_dg8_VISp | 44 | accuracy | 0.691667 |
| classification | transformer | Visual Coding | single_day | session_773418906_dg8_VISp | session_773418906_dg8_VISp | 44 | accuracy | 0.766667 |
| classification | transformer | Visual Coding | single_day | session_773418906_dg8_VISp | session_773418906_dg8_VISp | 46 | accuracy | 0.691667 |
| classification | transformer | Visual Coding | single_day | session_773418906_dg8_VISp | session_773418906_dg8_VISp | 46 | accuracy | 0.683333 |
| classification | transformer | Visual Coding | single_day | session_773418906_dg8_VISp | session_773418906_dg8_VISp | 54 | accuracy | 0.691667 |
| classification | transformer | Visual Coding | single_day | session_791319847_dg8_VISp | session_791319847_dg8_VISp | 44 | accuracy | 0.933333 |
| classification | transformer | Visual Coding | single_day | session_791319847_dg8_VISp | session_791319847_dg8_VISp | 46 | accuracy | 0.958333 |
| classification | transformer | Visual Coding | single_day | session_791319847_dg8_VISp | session_791319847_dg8_VISp | 46 | accuracy | 0.941667 |
| classification | transformer | Visual Coding | single_day | session_791319847_dg8_VISp | session_791319847_dg8_VISp | 54 | accuracy | 0.85 |
| classification | transformer | Visual Coding | single_day | session_791319847_dg8_VISp | session_791319847_dg8_VISp | 54 | accuracy | 0.858333 |
| classification | transformer | Visual Coding | single_day | session_797828357_dg8_VISp | session_797828357_dg8_VISp | 44 | accuracy | 0.766667 |
| classification | transformer | Visual Coding | single_day | session_797828357_dg8_VISp | session_797828357_dg8_VISp | 44 | accuracy | 0.775 |
| classification | transformer | Visual Coding | single_day | session_797828357_dg8_VISp | session_797828357_dg8_VISp | 46 | accuracy | 0.85 |
| classification | transformer | Visual Coding | single_day | session_797828357_dg8_VISp | session_797828357_dg8_VISp | 46 | accuracy | 0.841667 |
| classification | transformer | Visual Coding | single_day | session_797828357_dg8_VISp | session_797828357_dg8_VISp | 54 | accuracy | 0.841667 |
| classification | transformer | Visual Coding | single_day | session_797828357_dg8_VISp | session_797828357_dg8_VISp | 54 | accuracy | 0.833333 |
| classification | transformer | Visual Coding | single_day | session_798911424_dg8_VISp | session_798911424_dg8_VISp | 44 | accuracy | 0.916667 |
| classification | transformer | Visual Coding | single_day | session_798911424_dg8_VISp | session_798911424_dg8_VISp | 44 | accuracy | 0.933333 |
| classification | transformer | Visual Coding | single_day | session_798911424_dg8_VISp | session_798911424_dg8_VISp | 46 | accuracy | 0.875 |
| classification | transformer | Visual Coding | single_day | session_798911424_dg8_VISp | session_798911424_dg8_VISp | 46 | accuracy | 0.908333 |
| classification | transformer | Visual Coding | single_day | session_798911424_dg8_VISp | session_798911424_dg8_VISp | 54 | accuracy | 0.916667 |
| classification | transformer | Visual Coding | single_day | session_798911424_dg8_VISp | session_798911424_dg8_VISp | 54 | accuracy | 0.908333 |

| | | | | | | | | |
|---|---|---|---|---|---|---|---|---|
| classification | transformer | Visual Coding | single_day | session_799864342_dg8_VISp | session_799864342_dg8_VISp | 44 | accuracy | 0.941667 |
| classification | transformer | Visual Coding | single_day | session_799864342_dg8_VISp | session_799864342_dg8_VISp | 44 | accuracy | 0.908333 |
| classification | transformer | Visual Coding | single_day | session_799864342_dg8_VISp | session_799864342_dg8_VISp | 46 | accuracy | 0.966667 |
| classification | transformer | Visual Coding | single_day | session_799864342_dg8_VISp | session_799864342_dg8_VISp | 54 | accuracy | 0.958333 |
| classification | transformer | Visual Coding | single_day | session_799864342_dg8_VISp | session_799864342_dg8_VISp | 54 | accuracy | 0.95 |
| classification | lfads | Visual Coding | single_day | session_715093703_dg8_VISp | session_715093703_dg8_VISp | 44 | accuracy | 0.6 |
| classification | lfads | Visual Coding | single_day | session_715093703_dg8_VISp | session_715093703_dg8_VISp | 46 | accuracy | 0.516667 |
| classification | lfads | Visual Coding | single_day | session_715093703_dg8_VISp | session_715093703_dg8_VISp | 46 | accuracy | 0.55 |
| classification | lfads | Visual Coding | single_day | session_715093703_dg8_VISp | session_715093703_dg8_VISp | 54 | accuracy | 0.575 |
| classification | lfads | Visual Coding | single_day | session_715093703_dg8_VISp | session_715093703_dg8_VISp | 54 | accuracy | 0.591667 |
| classification | lfads | Visual Coding | single_day | session_719161530_dg8_VISp | session_719161530_dg8_VISp | 44 | accuracy | 0.675 |
| classification | lfads | Visual Coding | single_day | session_719161530_dg8_VISp | session_719161530_dg8_VISp | 44 | accuracy | 0.708333 |
| classification | lfads | Visual Coding | single_day | session_719161530_dg8_VISp | session_719161530_dg8_VISp | 46 | accuracy | 0.725 |
| classification | lfads | Visual Coding | single_day | session_719161530_dg8_VISp | session_719161530_dg8_VISp | 54 | accuracy | 0.7 |
| classification | lfads | Visual Coding | single_day | session_719161530_dg8_VISp | session_719161530_dg8_VISp | 54 | accuracy | 0.691667 |
| classification | lfads | Visual Coding | single_day | session_721123822_dg8_VISp | session_721123822_dg8_VISp | 44 | accuracy | 0.616667 |
| classification | lfads | Visual Coding | single_day | session_721123822_dg8_VISp | session_721123822_dg8_VISp | 46 | accuracy | 0.716667 |
| classification | lfads | Visual Coding | single_day | session_721123822_dg8_VISp | session_721123822_dg8_VISp | 46 | accuracy | 0.708333 |
| classification | lfads | Visual Coding | single_day | session_721123822_dg8_VISp | session_721123822_dg8_VISp | 54 | accuracy | 0.6 |
| classification | lfads | Visual Coding | single_day | session_721123822_dg8_VISp | session_721123822_dg8_VISp | 54 | accuracy | 0.608333 |
| classification | lfads | Visual Coding | single_day | session_732592105_dg8_VISp | session_732592105_dg8_VISp | 44 | accuracy | 0.633333 |
| classification | lfads | Visual Coding | single_day | session_732592105_dg8_VISp | session_732592105_dg8_VISp | 44 | accuracy | 0.658333 |
| classification | lfads | Visual Coding | single_day | session_732592105_dg8_VISp | session_732592105_dg8_VISp | 46 | accuracy | 0.658333 |
| classification | lfads | Visual Coding | single_day | session_732592105_dg8_VISp | session_732592105_dg8_VISp | 46 | accuracy | 0.625 |
| classification | lfads | Visual Coding | single_day | session_732592105_dg8_VISp | session_732592105_dg8_VISp | 54 | accuracy | 0.608333 |
| classification | lfads | Visual Coding | single_day | session_732592105_dg8_VISp | session_732592105_dg8_VISp | 54 | accuracy | 0.583333 |
| classification | lfads | Visual Coding | single_day | session_737581020_dg8_VISp | session_737581020_dg8_VISp | 44 | accuracy | 0.641667 |
| classification | lfads | Visual Coding | single_day | session_737581020_dg8_VISp | session_737581020_dg8_VISp | 46 | accuracy | 0.683333 |
| classification | lfads | Visual Coding | single_day | session_737581020_dg8_VISp | session_737581020_dg8_VISp | 54 | accuracy | 0.716667 |
| classification | lfads | Visual Coding | single_day | session_737581020_dg8_VISp | session_737581020_dg8_VISp | 54 | accuracy | 0.7 |
| classification | lfads | Visual Coding | single_day | session_739448407_dg8_VISp | session_739448407_dg8_VISp | 44 | accuracy | 0.453782 |
| classification | lfads | Visual Coding | single_day | session_739448407_dg8_VISp | session_739448407_dg8_VISp | 44 | accuracy | 0.420168 |
| classification | lfads | Visual Coding | single_day | session_739448407_dg8_VISp | session_739448407_dg8_VISp | 46 | accuracy | 0.529412 |
| classification | lfads | Visual Coding | single_day | session_739448407_dg8_VISp | session_739448407_dg8_VISp | 46 | accuracy | 0.504202 |
| classification | lfads | Visual Coding | single_day | session_739448407_dg8_VISp | session_739448407_dg8_VISp | 54 | accuracy | 0.495798 |
| classification | lfads | Visual Coding | single_day | session_742951821_dg8_VISp | session_742951821_dg8_VISp | 44 | accuracy | 0.491667 |
| classification | lfads | Visual Coding | single_day | session_742951821_dg8_VISp | session_742951821_dg8_VISp | 44 | accuracy | 0.516667 |
| classification | lfads | Visual Coding | single_day | session_742951821_dg8_VISp | session_742951821_dg8_VISp | 46 | accuracy | 0.483333 |
| classification | lfads | Visual Coding | single_day | session_742951821_dg8_VISp | session_742951821_dg8_VISp | 46 | accuracy | 0.491667 |
| classification | lfads | Visual Coding | single_day | session_742951821_dg8_VISp | session_742951821_dg8_VISp | 54 | accuracy | 0.558333 |
| classification | lfads | Visual Coding | single_day | session_742951821_dg8_VISp | session_742951821_dg8_VISp | 54 | accuracy | 0.566667 |
| classification | lfads | Visual Coding | single_day | session_743475441_dg8_VISp | session_743475441_dg8_VISp | 44 | accuracy | 0.266667 |
| classification | lfads | Visual Coding | single_day | session_743475441_dg8_VISp | session_743475441_dg8_VISp | 44 | accuracy | 0.233333 |
| classification | lfads | Visual Coding | single_day | session_743475441_dg8_VISp | session_743475441_dg8_VISp | 46 | accuracy | 0.291667 |
| classification | lfads | Visual Coding | single_day | session_743475441_dg8_VISp | session_743475441_dg8_VISp | 54 | accuracy | 0.341667 |
| classification | lfads | Visual Coding | single_day | session_743475441_dg8_VISp | session_743475441_dg8_VISp | 54 | accuracy | 0.3 |
| classification | lfads | Visual Coding | single_day | session_744228101_dg8_VISp | session_744228101_dg8_VISp | 44 | accuracy | 0.733333 |
| classification | lfads | Visual Coding | single_day | session_744228101_dg8_VISp | session_744228101_dg8_VISp | 46 | accuracy | 0.733333 |
| classification | lfads | Visual Coding | single_day | session_744228101_dg8_VISp | session_744228101_dg8_VISp | 46 | accuracy | 0.741667 |
| classification | lfads | Visual Coding | single_day | session_744228101_dg8_VISp | session_744228101_dg8_VISp | 54 | accuracy | 0.683333 |
| classification | lfads | Visual Coding | single_day | session_746083955_dg8_VISp | session_746083955_dg8_VISp | 44 | accuracy | 0.266667 |
| classification | lfads | Visual Coding | single_day | session_746083955_dg8_VISp | session_746083955_dg8_VISp | 44 | accuracy | 0.291667 |
| classification | lfads | Visual Coding | single_day | session_746083955_dg8_VISp | session_746083955_dg8_VISp | 46 | accuracy | 0.475 |
| classification | lfads | Visual Coding | single_day | session_746083955_dg8_VISp | session_746083955_dg8_VISp | 54 | accuracy | 0.3 |
| classification | lfads | Visual Coding | single_day | session_746083955_dg8_VISp | session_746083955_dg8_VISp | 54 | accuracy | 0.308333 |
| classification | lfads | Visual Coding | single_day | session_750332458_dg8_VISp | session_750332458_dg8_VISp | 44 | accuracy | 0.741667 |
| classification | lfads | Visual Coding | single_day | session_750332458_dg8_VISp | session_750332458_dg8_VISp | 44 | accuracy | 0.716667 |
| classification | lfads | Visual Coding | single_day | session_750332458_dg8_VISp | session_750332458_dg8_VISp | 46 | accuracy | 0.741667 |
| classification | lfads | Visual Coding | single_day | session_750332458_dg8_VISp | session_750332458_dg8_VISp | 46 | accuracy | 0.725 |
| classification | lfads | Visual Coding | single_day | session_750332458_dg8_VISp | session_750332458_dg8_VISp | 54 | accuracy | 0.766667 |
| classification | lfads | Visual Coding | single_day | session_750332458_dg8_VISp | session_750332458_dg8_VISp | 54 | accuracy | 0.8 |
| classification | lfads | Visual Coding | single_day | session_750749662_dg8_VISp | session_750749662_dg8_VISp | 44 | accuracy | 0.683333 |
| classification | lfads | Visual Coding | single_day | session_750749662_dg8_VISp | session_750749662_dg8_VISp | 44 | accuracy | 0.675 |

| | | | | | | | | |
|---|---|---|---|---|---|---|---|---|
| classification | lfads | Visual Coding | single_day | session_750749662_dg8_VISp | session_750749662_dg8_VISp | 46 | accuracy | 0.691667 |
| classification | lfads | Visual Coding | single_day | session_750749662_dg8_VISp | session_750749662_dg8_VISp | 46 | accuracy | 0.683333 |
| classification | lfads | Visual Coding | single_day | session_750749662_dg8_VISp | session_750749662_dg8_VISp | 54 | accuracy | 0.7 |
| classification | lfads | Visual Coding | single_day | session_750749662_dg8_VISp | session_750749662_dg8_VISp | 54 | accuracy | 0.691667 |
| classification | lfads | Visual Coding | single_day | session_751348571_dg8_VISp | session_751348571_dg8_VISp | 44 | accuracy | 0.783333 |
| classification | lfads | Visual Coding | single_day | session_751348571_dg8_VISp | session_751348571_dg8_VISp | 46 | accuracy | 0.783333 |
| classification | lfads | Visual Coding | single_day | session_751348571_dg8_VISp | session_751348571_dg8_VISp | 46 | accuracy | 0.791667 |
| classification | lfads | Visual Coding | single_day | session_751348571_dg8_VISp | session_751348571_dg8_VISp | 54 | accuracy | 0.841667 |
| classification | lfads | Visual Coding | single_day | session_751348571_dg8_VISp | session_751348571_dg8_VISp | 54 | accuracy | 0.833333 |
| classification | lfads | Visual Coding | single_day | session_754312389_dg8_VISp | session_754312389_dg8_VISp | 44 | accuracy | 0.95 |
| classification | lfads | Visual Coding | single_day | session_754312389_dg8_VISp | session_754312389_dg8_VISp | 44 | accuracy | 0.941667 |
| classification | lfads | Visual Coding | single_day | session_754312389_dg8_VISp | session_754312389_dg8_VISp | 46 | accuracy | 0.825 |
| classification | lfads | Visual Coding | single_day | session_754312389_dg8_VISp | session_754312389_dg8_VISp | 46 | accuracy | 0.883333 |
| classification | lfads | Visual Coding | single_day | session_754312389_dg8_VISp | session_754312389_dg8_VISp | 54 | accuracy | 0.866667 |
| classification | lfads | Visual Coding | single_day | session_754312389_dg8_VISp | session_754312389_dg8_VISp | 54 | accuracy | 0.858333 |
| classification | lfads | Visual Coding | single_day | session_754829445_dg8_VISp | session_754829445_dg8_VISp | 44 | accuracy | 0.708333 |
| classification | lfads | Visual Coding | single_day | session_754829445_dg8_VISp | session_754829445_dg8_VISp | 44 | accuracy | 0.691667 |
| classification | lfads | Visual Coding | single_day | session_754829445_dg8_VISp | session_754829445_dg8_VISp | 46 | accuracy | 0.55 |
| classification | lfads | Visual Coding | single_day | session_754829445_dg8_VISp | session_754829445_dg8_VISp | 46 | accuracy | 0.558333 |
| classification | lfads | Visual Coding | single_day | session_754829445_dg8_VISp | session_754829445_dg8_VISp | 54 | accuracy | 0.733333 |
| classification | lfads | Visual Coding | single_day | session_754829445_dg8_VISp | session_754829445_dg8_VISp | 54 | accuracy | 0.683333 |
| classification | lfads | Visual Coding | single_day | session_755434585_dg8_VISp | session_755434585_dg8_VISp | 44 | accuracy | 0.666667 |
| classification | lfads | Visual Coding | single_day | session_755434585_dg8_VISp | session_755434585_dg8_VISp | 44 | accuracy | 0.708333 |
| classification | lfads | Visual Coding | single_day | session_755434585_dg8_VISp | session_755434585_dg8_VISp | 46 | accuracy | 0.716667 |
| classification | lfads | Visual Coding | single_day | session_755434585_dg8_VISp | session_755434585_dg8_VISp | 46 | accuracy | 0.725 |
| classification | lfads | Visual Coding | single_day | session_755434585_dg8_VISp | session_755434585_dg8_VISp | 54 | accuracy | 0.708333 |
| classification | lfads | Visual Coding | single_day | session_755434585_dg8_VISp | session_755434585_dg8_VISp | 54 | accuracy | 0.716667 |
| classification | lfads | Visual Coding | single_day | session_756029989_dg8_VISp | session_756029989_dg8_VISp | 44 | accuracy | 0.725 |
| classification | lfads | Visual Coding | single_day | session_756029989_dg8_VISp | session_756029989_dg8_VISp | 44 | accuracy | 0.716667 |
| classification | lfads | Visual Coding | single_day | session_756029989_dg8_VISp | session_756029989_dg8_VISp | 46 | accuracy | 0.75 |
| classification | lfads | Visual Coding | single_day | session_756029989_dg8_VISp | session_756029989_dg8_VISp | 54 | accuracy | 0.75 |
| classification | lfads | Visual Coding | single_day | session_757216464_dg8_VISp | session_757216464_dg8_VISp | 44 | accuracy | 0.566667 |
| classification | lfads | Visual Coding | single_day | session_757216464_dg8_VISp | session_757216464_dg8_VISp | 44 | accuracy | 0.616667 |
| classification | lfads | Visual Coding | single_day | session_757216464_dg8_VISp | session_757216464_dg8_VISp | 46 | accuracy | 0.65 |
| classification | lfads | Visual Coding | single_day | session_757216464_dg8_VISp | session_757216464_dg8_VISp | 54 | accuracy | 0.583333 |
| classification | lfads | Visual Coding | single_day | session_757970808_dg8_VISp | session_757970808_dg8_VISp | 44 | accuracy | 0.625 |
| classification | lfads | Visual Coding | single_day | session_757970808_dg8_VISp | session_757970808_dg8_VISp | 44 | accuracy | 0.608333 |
| classification | lfads | Visual Coding | single_day | session_757970808_dg8_VISp | session_757970808_dg8_VISp | 46 | accuracy | 0.633333 |
| classification | lfads | Visual Coding | single_day | session_757970808_dg8_VISp | session_757970808_dg8_VISp | 46 | accuracy | 0.658333 |
| classification | lfads | Visual Coding | single_day | session_757970808_dg8_VISp | session_757970808_dg8_VISp | 54 | accuracy | 0.716667 |
| classification | lfads | Visual Coding | single_day | session_757970808_dg8_VISp | session_757970808_dg8_VISp | 54 | accuracy | 0.675 |
| classification | lfads | Visual Coding | single_day | session_758798717_dg8_VISp | session_758798717_dg8_VISp | 44 | accuracy | 0.608333 |
| classification | lfads | Visual Coding | single_day | session_758798717_dg8_VISp | session_758798717_dg8_VISp | 46 | accuracy | 0.625 |
| classification | lfads | Visual Coding | single_day | session_758798717_dg8_VISp | session_758798717_dg8_VISp | 46 | accuracy | 0.608333 |
| classification | lfads | Visual Coding | single_day | session_758798717_dg8_VISp | session_758798717_dg8_VISp | 54 | accuracy | 0.625 |
| classification | lfads | Visual Coding | single_day | session_759883607_dg8_VISp | session_759883607_dg8_VISp | 44 | accuracy | 0.658333 |
| classification | lfads | Visual Coding | single_day | session_759883607_dg8_VISp | session_759883607_dg8_VISp | 44 | accuracy | 0.683333 |
| classification | lfads | Visual Coding | single_day | session_759883607_dg8_VISp | session_759883607_dg8_VISp | 46 | accuracy | 0.591667 |
| classification | lfads | Visual Coding | single_day | session_759883607_dg8_VISp | session_759883607_dg8_VISp | 46 | accuracy | 0.6 |
| classification | lfads | Visual Coding | single_day | session_759883607_dg8_VISp | session_759883607_dg8_VISp | 54 | accuracy | 0.6 |
| classification | lfads | Visual Coding | single_day | session_760345702_dg8_VISp | session_760345702_dg8_VISp | 44 | accuracy | 0.783333 |
| classification | lfads | Visual Coding | single_day | session_760345702_dg8_VISp | session_760345702_dg8_VISp | 44 | accuracy | 0.841667 |
| classification | lfads | Visual Coding | single_day | session_760345702_dg8_VISp | session_760345702_dg8_VISp | 46 | accuracy | 0.775 |
| classification | lfads | Visual Coding | single_day | session_760345702_dg8_VISp | session_760345702_dg8_VISp | 46 | accuracy | 0.783333 |
| classification | lfads | Visual Coding | single_day | session_760345702_dg8_VISp | session_760345702_dg8_VISp | 54 | accuracy | 0.791667 |
| classification | lfads | Visual Coding | single_day | session_760345702_dg8_VISp | session_760345702_dg8_VISp | 54 | accuracy | 0.775 |
| classification | lfads | Visual Coding | single_day | session_760693773_dg8_VISp | session_760693773_dg8_VISp | 44 | accuracy | 0.8 |
| classification | lfads | Visual Coding | single_day | session_760693773_dg8_VISp | session_760693773_dg8_VISp | 44 | accuracy | 0.783333 |
| classification | lfads | Visual Coding | single_day | session_760693773_dg8_VISp | session_760693773_dg8_VISp | 46 | accuracy | 0.8 |
| classification | lfads | Visual Coding | single_day | session_760693773_dg8_VISp | session_760693773_dg8_VISp | 46 | accuracy | 0.808333 |
| classification | lfads | Visual Coding | single_day | session_760693773_dg8_VISp | session_760693773_dg8_VISp | 54 | accuracy | 0.791667 |
| classification | lfads | Visual Coding | single_day | session_760693773_dg8_VISp | session_760693773_dg8_VISp | 54 | accuracy | 0.741667 |
| classification | lfads | Visual Coding | single_day | session_761418226_dg8_VISp | session_761418226_dg8_VISp | 44 | accuracy | 0.708333 |

| | | | | | | | | |
|---|---|---|---|---|---|---|---|---|
| classification | lfads | Visual Coding | single_day | session_761418226_dg8_VISp | session_761418226_dg8_VISp | 44 | accuracy | 0.716667 |
| classification | lfads | Visual Coding | single_day | session_761418226_dg8_VISp | session_761418226_dg8_VISp | 46 | accuracy | 0.775 |
| classification | lfads | Visual Coding | single_day | session_761418226_dg8_VISp | session_761418226_dg8_VISp | 54 | accuracy | 0.758333 |
| classification | lfads | Visual Coding | single_day | session_762120172_dg8_VISp | session_762120172_dg8_VISp | 44 | accuracy | 0.741667 |
| classification | lfads | Visual Coding | single_day | session_762120172_dg8_VISp | session_762120172_dg8_VISp | 44 | accuracy | 0.716667 |
| classification | lfads | Visual Coding | single_day | session_762120172_dg8_VISp | session_762120172_dg8_VISp | 46 | accuracy | 0.658333 |
| classification | lfads | Visual Coding | single_day | session_762120172_dg8_VISp | session_762120172_dg8_VISp | 46 | accuracy | 0.666667 |
| classification | lfads | Visual Coding | single_day | session_762120172_dg8_VISp | session_762120172_dg8_VISp | 54 | accuracy | 0.708333 |
| classification | lfads | Visual Coding | single_day | session_762120172_dg8_VISp | session_762120172_dg8_VISp | 54 | accuracy | 0.691667 |
| classification | lfads | Visual Coding | single_day | session_762602078_dg8_VISp | session_762602078_dg8_VISp | 44 | accuracy | 0.9 |
| classification | lfads | Visual Coding | single_day | session_762602078_dg8_VISp | session_762602078_dg8_VISp | 44 | accuracy | 0.883333 |
| classification | lfads | Visual Coding | single_day | session_762602078_dg8_VISp | session_762602078_dg8_VISp | 46 | accuracy | 0.916667 |
| classification | lfads | Visual Coding | single_day | session_762602078_dg8_VISp | session_762602078_dg8_VISp | 46 | accuracy | 0.9 |
| classification | lfads | Visual Coding | single_day | session_762602078_dg8_VISp | session_762602078_dg8_VISp | 54 | accuracy | 0.725 |
| classification | lfads | Visual Coding | single_day | session_762602078_dg8_VISp | session_762602078_dg8_VISp | 54 | accuracy | 0.758333 |
| classification | lfads | Visual Coding | single_day | session_763673393_dg8_VISp | session_763673393_dg8_VISp | 44 | accuracy | 0.675 |
| classification | lfads | Visual Coding | single_day | session_763673393_dg8_VISp | session_763673393_dg8_VISp | 44 | accuracy | 0.608333 |
| classification | lfads | Visual Coding | single_day | session_763673393_dg8_VISp | session_763673393_dg8_VISp | 46 | accuracy | 0.666667 |
| classification | lfads | Visual Coding | single_day | session_763673393_dg8_VISp | session_763673393_dg8_VISp | 46 | accuracy | 0.683333 |
| classification | lfads | Visual Coding | single_day | session_763673393_dg8_VISp | session_763673393_dg8_VISp | 54 | accuracy | 0.7 |
| classification | lfads | Visual Coding | single_day | session_763673393_dg8_VISp | session_763673393_dg8_VISp | 54 | accuracy | 0.683333 |
| classification | lfads | Visual Coding | single_day | session_773418906_dg8_VISp | session_773418906_dg8_VISp | 44 | accuracy | 0.566667 |
| classification | lfads | Visual Coding | single_day | session_773418906_dg8_VISp | session_773418906_dg8_VISp | 44 | accuracy | 0.558333 |
| classification | lfads | Visual Coding | single_day | session_773418906_dg8_VISp | session_773418906_dg8_VISp | 46 | accuracy | 0.466667 |
| classification | lfads | Visual Coding | single_day | session_773418906_dg8_VISp | session_773418906_dg8_VISp | 54 | accuracy | 0.583333 |
| classification | lfads | Visual Coding | single_day | session_773418906_dg8_VISp | session_773418906_dg8_VISp | 54 | accuracy | 0.525 |
| classification | lfads | Visual Coding | single_day | session_791319847_dg8_VISp | session_791319847_dg8_VISp | 44 | accuracy | 0.808333 |
| classification | lfads | Visual Coding | single_day | session_791319847_dg8_VISp | session_791319847_dg8_VISp | 44 | accuracy | 0.85 |
| classification | lfads | Visual Coding | single_day | session_791319847_dg8_VISp | session_791319847_dg8_VISp | 46 | accuracy | 0.75 |
| classification | lfads | Visual Coding | single_day | session_791319847_dg8_VISp | session_791319847_dg8_VISp | 46 | accuracy | 0.775 |
| classification | lfads | Visual Coding | single_day | session_791319847_dg8_VISp | session_791319847_dg8_VISp | 54 | accuracy | 0.783333 |
| classification | lfads | Visual Coding | single_day | session_791319847_dg8_VISp | session_791319847_dg8_VISp | 54 | accuracy | 0.791667 |
| classification | lfads | Visual Coding | single_day | session_797828357_dg8_VISp | session_797828357_dg8_VISp | 44 | accuracy | 0.616667 |
| classification | lfads | Visual Coding | single_day | session_797828357_dg8_VISp | session_797828357_dg8_VISp | 44 | accuracy | 0.55 |
| classification | lfads | Visual Coding | single_day | session_797828357_dg8_VISp | session_797828357_dg8_VISp | 46 | accuracy | 0.491667 |
| classification | lfads | Visual Coding | single_day | session_797828357_dg8_VISp | session_797828357_dg8_VISp | 46 | accuracy | 0.508333 |
| classification | lfads | Visual Coding | single_day | session_797828357_dg8_VISp | session_797828357_dg8_VISp | 54 | accuracy | 0.558333 |
| classification | lfads | Visual Coding | single_day | session_798911424_dg8_VISp | session_798911424_dg8_VISp | 44 | accuracy | 0.658333 |
| classification | lfads | Visual Coding | single_day | session_798911424_dg8_VISp | session_798911424_dg8_VISp | 44 | accuracy | 0.691667 |
| classification | lfads | Visual Coding | single_day | session_798911424_dg8_VISp | session_798911424_dg8_VISp | 46 | accuracy | 0.525 |
| classification | lfads | Visual Coding | single_day | session_798911424_dg8_VISp | session_798911424_dg8_VISp | 46 | accuracy | 0.541667 |
| classification | lfads | Visual Coding | single_day | session_798911424_dg8_VISp | session_798911424_dg8_VISp | 54 | accuracy | 0.683333 |
| classification | lfads | Visual Coding | single_day | session_798911424_dg8_VISp | session_798911424_dg8_VISp | 54 | accuracy | 0.583333 |
| classification | lfads | Visual Coding | single_day | session_799864342_dg8_VISp | session_799864342_dg8_VISp | 44 | accuracy | 0.841667 |
| classification | lfads | Visual Coding | single_day | session_799864342_dg8_VISp | session_799864342_dg8_VISp | 44 | accuracy | 0.825 |
| classification | lfads | Visual Coding | single_day | session_799864342_dg8_VISp | session_799864342_dg8_VISp | 46 | accuracy | 0.858333 |
| classification | lfads | Visual Coding | single_day | session_799864342_dg8_VISp | session_799864342_dg8_VISp | 46 | accuracy | 0.85 |
| classification | lfads | Visual Coding | single_day | session_799864342_dg8_VISp | session_799864342_dg8_VISp | 54 | accuracy | 0.791667 |
| classification | lfads | Visual Coding | single_day | session_799864342_dg8_VISp | session_799864342_dg8_VISp | 54 | accuracy | 0.758333 |
| classification | cycle_gan | Visual Coding | single_day | session_715093703_dg8_VISp | session_715093703_dg8_VISp | 44 | accuracy | 0.342593 |
| classification | cycle_gan | Visual Coding | single_day | session_715093703_dg8_VISp | session_715093703_dg8_VISp | 44 | accuracy | 0.725 |
| classification | cycle_gan | Visual Coding | single_day | session_715093703_dg8_VISp | session_715093703_dg8_VISp | 46 | accuracy | 0.12963 |
| classification | cycle_gan | Visual Coding | single_day | session_715093703_dg8_VISp | session_715093703_dg8_VISp | 46 | accuracy | 0.675 |
| classification | cycle_gan | Visual Coding | single_day | session_715093703_dg8_VISp | session_715093703_dg8_VISp | 54 | accuracy | 0.0740741 |
| classification | cycle_gan | Visual Coding | single_day | session_715093703_dg8_VISp | session_715093703_dg8_VISp | 54 | accuracy | 0.758333 |
| classification | cycle_gan | Visual Coding | single_day | session_719161530_dg8_VISp | session_719161530_dg8_VISp | 44 | accuracy | 0.0555556 |
| classification | cycle_gan | Visual Coding | single_day | session_719161530_dg8_VISp | session_719161530_dg8_VISp | 44 | accuracy | 0.725 |
| classification | cycle_gan | Visual Coding | single_day | session_719161530_dg8_VISp | session_719161530_dg8_VISp | 46 | accuracy | 0.138889 |
| classification | cycle_gan | Visual Coding | single_day | session_719161530_dg8_VISp | session_719161530_dg8_VISp | 46 | accuracy | 0.741667 |
| classification | cycle_gan | Visual Coding | single_day | session_719161530_dg8_VISp | session_719161530_dg8_VISp | 54 | accuracy | 0.12963 |
| classification | cycle_gan | Visual Coding | single_day | session_719161530_dg8_VISp | session_719161530_dg8_VISp | 54 | accuracy | 0.758333 |
| classification | cycle_gan | Visual Coding | single_day | session_721123822_dg8_VISp | session_721123822_dg8_VISp | 44 | accuracy | 0.111111 |
| classification | cycle_gan | Visual Coding | single_day | session_721123822_dg8_VISp | session_721123822_dg8_VISp | 44 | accuracy | 0.616667 |

| | | | | | | | | |
|---|---|---|---|---|---|---|---|---|
| classification | cycle_gan | Visual Coding | single_day | session_721123822_dg8_VISp | session_721123822_dg8_VISp | 46 | accuracy | 0.12037 |
| classification | cycle_gan | Visual Coding | single_day | session_721123822_dg8_VISp | session_721123822_dg8_VISp | 46 | accuracy | 0.675 |
| classification | cycle_gan | Visual Coding | single_day | session_721123822_dg8_VISp | session_721123822_dg8_VISp | 54 | accuracy | 0.12037 |
| classification | cycle_gan | Visual Coding | single_day | session_721123822_dg8_VISp | session_721123822_dg8_VISp | 54 | accuracy | 0.583333 |
| classification | cycle_gan | Visual Coding | single_day | session_732592105_dg8_VISp | session_732592105_dg8_VISp | 44 | accuracy | 0.12963 |
| classification | cycle_gan | Visual Coding | single_day | session_732592105_dg8_VISp | session_732592105_dg8_VISp | 44 | accuracy | 0.683333 |
| classification | cycle_gan | Visual Coding | single_day | session_732592105_dg8_VISp | session_732592105_dg8_VISp | 46 | accuracy | 0.12037 |
| classification | cycle_gan | Visual Coding | single_day | session_732592105_dg8_VISp | session_732592105_dg8_VISp | 46 | accuracy | 0.758333 |
| classification | cycle_gan | Visual Coding | single_day | session_732592105_dg8_VISp | session_732592105_dg8_VISp | 54 | accuracy | 0.12963 |
| classification | cycle_gan | Visual Coding | single_day | session_732592105_dg8_VISp | session_732592105_dg8_VISp | 54 | accuracy | 0.666667 |
| classification | cycle_gan | Visual Coding | single_day | session_737581020_dg8_VISp | session_737581020_dg8_VISp | 44 | accuracy | 0.25 |
| classification | cycle_gan | Visual Coding | single_day | session_737581020_dg8_VISp | session_737581020_dg8_VISp | 44 | accuracy | 0.733333 |
| classification | cycle_gan | Visual Coding | single_day | session_737581020_dg8_VISp | session_737581020_dg8_VISp | 46 | accuracy | 0.259259 |
| classification | cycle_gan | Visual Coding | single_day | session_737581020_dg8_VISp | session_737581020_dg8_VISp | 46 | accuracy | 0.758333 |
| classification | cycle_gan | Visual Coding | single_day | session_737581020_dg8_VISp | session_737581020_dg8_VISp | 54 | accuracy | 0.148148 |
| classification | cycle_gan | Visual Coding | single_day | session_737581020_dg8_VISp | session_737581020_dg8_VISp | 54 | accuracy | 0.825 |
| classification | cycle_gan | Visual Coding | single_day | session_739448407_dg8_VISp | session_739448407_dg8_VISp | 44 | accuracy | 0.12963 |
| classification | cycle_gan | Visual Coding | single_day | session_739448407_dg8_VISp | session_739448407_dg8_VISp | 44 | accuracy | 0.529412 |
| classification | cycle_gan | Visual Coding | single_day | session_739448407_dg8_VISp | session_739448407_dg8_VISp | 46 | accuracy | 0.222222 |
| classification | cycle_gan | Visual Coding | single_day | session_739448407_dg8_VISp | session_739448407_dg8_VISp | 46 | accuracy | 0.521008 |
| classification | cycle_gan | Visual Coding | single_day | session_739448407_dg8_VISp | session_739448407_dg8_VISp | 54 | accuracy | 0.138889 |
| classification | cycle_gan | Visual Coding | single_day | session_739448407_dg8_VISp | session_739448407_dg8_VISp | 54 | accuracy | 0.546218 |
| classification | cycle_gan | Visual Coding | single_day | session_742951821_dg8_VISp | session_742951821_dg8_VISp | 44 | accuracy | 0.12963 |
| classification | cycle_gan | Visual Coding | single_day | session_742951821_dg8_VISp | session_742951821_dg8_VISp | 44 | accuracy | 0.408333 |
| classification | cycle_gan | Visual Coding | single_day | session_742951821_dg8_VISp | session_742951821_dg8_VISp | 46 | accuracy | 0.194444 |
| classification | cycle_gan | Visual Coding | single_day | session_742951821_dg8_VISp | session_742951821_dg8_VISp | 46 | accuracy | 0.558333 |
| classification | cycle_gan | Visual Coding | single_day | session_742951821_dg8_VISp | session_742951821_dg8_VISp | 54 | accuracy | 0.157407 |
| classification | cycle_gan | Visual Coding | single_day | session_742951821_dg8_VISp | session_742951821_dg8_VISp | 54 | accuracy | 0.5 |
| classification | cycle_gan | Visual Coding | single_day | session_743475441_dg8_VISp | session_743475441_dg8_VISp | 44 | accuracy | 0.111111 |
| classification | cycle_gan | Visual Coding | single_day | session_743475441_dg8_VISp | session_743475441_dg8_VISp | 44 | accuracy | 0.341667 |
| classification | cycle_gan | Visual Coding | single_day | session_743475441_dg8_VISp | session_743475441_dg8_VISp | 46 | accuracy | 0.12037 |
| classification | cycle_gan | Visual Coding | single_day | session_743475441_dg8_VISp | session_743475441_dg8_VISp | 46 | accuracy | 0.283333 |
| classification | cycle_gan | Visual Coding | single_day | session_743475441_dg8_VISp | session_743475441_dg8_VISp | 54 | accuracy | 0.12037 |
| classification | cycle_gan | Visual Coding | single_day | session_743475441_dg8_VISp | session_743475441_dg8_VISp | 54 | accuracy | 0.358333 |
| classification | cycle_gan | Visual Coding | single_day | session_744228101_dg8_VISp | session_744228101_dg8_VISp | 44 | accuracy | 0.12963 |
| classification | cycle_gan | Visual Coding | single_day | session_744228101_dg8_VISp | session_744228101_dg8_VISp | 44 | accuracy | 0.775 |
| classification | cycle_gan | Visual Coding | single_day | session_744228101_dg8_VISp | session_744228101_dg8_VISp | 46 | accuracy | 0.101852 |
| classification | cycle_gan | Visual Coding | single_day | session_744228101_dg8_VISp | session_744228101_dg8_VISp | 46 | accuracy | 0.741667 |
| classification | cycle_gan | Visual Coding | single_day | session_744228101_dg8_VISp | session_744228101_dg8_VISp | 54 | accuracy | 0.101852 |
| classification | cycle_gan | Visual Coding | single_day | session_744228101_dg8_VISp | session_744228101_dg8_VISp | 54 | accuracy | 0.75 |
| classification | cycle_gan | Visual Coding | single_day | session_746083955_dg8_VISp | session_746083955_dg8_VISp | 44 | accuracy | 0.222222 |
| classification | cycle_gan | Visual Coding | single_day | session_746083955_dg8_VISp | session_746083955_dg8_VISp | 44 | accuracy | 0.425 |
| classification | cycle_gan | Visual Coding | single_day | session_746083955_dg8_VISp | session_746083955_dg8_VISp | 46 | accuracy | 0.333333 |
| classification | cycle_gan | Visual Coding | single_day | session_746083955_dg8_VISp | session_746083955_dg8_VISp | 46 | accuracy | 0.425 |
| classification | cycle_gan | Visual Coding | single_day | session_746083955_dg8_VISp | session_746083955_dg8_VISp | 54 | accuracy | 0.138889 |
| classification | cycle_gan | Visual Coding | single_day | session_746083955_dg8_VISp | session_746083955_dg8_VISp | 54 | accuracy | 0.425 |
| classification | cycle_gan | Visual Coding | single_day | session_750332458_dg8_VISp | session_750332458_dg8_VISp | 44 | accuracy | 0.111111 |
| classification | cycle_gan | Visual Coding | single_day | session_750332458_dg8_VISp | session_750332458_dg8_VISp | 44 | accuracy | 0.816667 |
| classification | cycle_gan | Visual Coding | single_day | session_750332458_dg8_VISp | session_750332458_dg8_VISp | 46 | accuracy | 0.138889 |
| classification | cycle_gan | Visual Coding | single_day | session_750332458_dg8_VISp | session_750332458_dg8_VISp | 46 | accuracy | 0.875 |
| classification | cycle_gan | Visual Coding | single_day | session_750332458_dg8_VISp | session_750332458_dg8_VISp | 54 | accuracy | 0.0925926 |
| classification | cycle_gan | Visual Coding | single_day | session_750332458_dg8_VISp | session_750332458_dg8_VISp | 54 | accuracy | 0.808333 |
| classification | cycle_gan | Visual Coding | single_day | session_750749662_dg8_VISp | session_750749662_dg8_VISp | 44 | accuracy | 0.101852 |
| classification | cycle_gan | Visual Coding | single_day | session_750749662_dg8_VISp | session_750749662_dg8_VISp | 44 | accuracy | 0.691667 |
| classification | cycle_gan | Visual Coding | single_day | session_750749662_dg8_VISp | session_750749662_dg8_VISp | 46 | accuracy | 0.166667 |
| classification | cycle_gan | Visual Coding | single_day | session_750749662_dg8_VISp | session_750749662_dg8_VISp | 46 | accuracy | 0.708333 |
| classification | cycle_gan | Visual Coding | single_day | session_750749662_dg8_VISp | session_750749662_dg8_VISp | 54 | accuracy | 0.138889 |
| classification | cycle_gan | Visual Coding | single_day | session_750749662_dg8_VISp | session_750749662_dg8_VISp | 54 | accuracy | 0.75 |
| classification | cycle_gan | Visual Coding | single_day | session_751348571_dg8_VISp | session_751348571_dg8_VISp | 44 | accuracy | 0.185185 |
| classification | cycle_gan | Visual Coding | single_day | session_751348571_dg8_VISp | session_751348571_dg8_VISp | 44 | accuracy | 0.933333 |
| classification | cycle_gan | Visual Coding | single_day | session_751348571_dg8_VISp | session_751348571_dg8_VISp | 46 | accuracy | 0.175926 |
| classification | cycle_gan | Visual Coding | single_day | session_751348571_dg8_VISp | session_751348571_dg8_VISp | 46 | accuracy | 0.908333 |
| classification | cycle_gan | Visual Coding | single_day | session_751348571_dg8_VISp | session_751348571_dg8_VISp | 54 | accuracy | 0.101852 |

| | | | | | | | | |
|---|---|---|---|---|---|---|---|---|
| classification | cycle_gan | Visual Coding | single_day | session_751348571_dg8_VISp | session_751348571_dg8_VISp | 54 | accuracy | 0.825 |
| classification | cycle_gan | Visual Coding | single_day | session_754312389_dg8_VISp | session_754312389_dg8_VISp | 44 | accuracy | 0.12963 |
| classification | cycle_gan | Visual Coding | single_day | session_754312389_dg8_VISp | session_754312389_dg8_VISp | 44 | accuracy | 0.9 |
| classification | cycle_gan | Visual Coding | single_day | session_754312389_dg8_VISp | session_754312389_dg8_VISp | 46 | accuracy | 0.0833333 |
| classification | cycle_gan | Visual Coding | single_day | session_754312389_dg8_VISp | session_754312389_dg8_VISp | 46 | accuracy | 0.808333 |
| classification | cycle_gan | Visual Coding | single_day | session_754312389_dg8_VISp | session_754312389_dg8_VISp | 54 | accuracy | 0.138889 |
| classification | cycle_gan | Visual Coding | single_day | session_754312389_dg8_VISp | session_754312389_dg8_VISp | 54 | accuracy | 0.841667 |
| classification | cycle_gan | Visual Coding | single_day | session_754829445_dg8_VISp | session_754829445_dg8_VISp | 44 | accuracy | 0.166667 |
| classification | cycle_gan | Visual Coding | single_day | session_754829445_dg8_VISp | session_754829445_dg8_VISp | 44 | accuracy | 0.716667 |
| classification | cycle_gan | Visual Coding | single_day | session_754829445_dg8_VISp | session_754829445_dg8_VISp | 46 | accuracy | 0.148148 |
| classification | cycle_gan | Visual Coding | single_day | session_754829445_dg8_VISp | session_754829445_dg8_VISp | 46 | accuracy | 0.733333 |
| classification | cycle_gan | Visual Coding | single_day | session_754829445_dg8_VISp | session_754829445_dg8_VISp | 54 | accuracy | 0.101852 |
| classification | cycle_gan | Visual Coding | single_day | session_754829445_dg8_VISp | session_754829445_dg8_VISp | 54 | accuracy | 0.783333 |
| classification | cycle_gan | Visual Coding | single_day | session_755434585_dg8_VISp | session_755434585_dg8_VISp | 44 | accuracy | 0.166667 |
| classification | cycle_gan | Visual Coding | single_day | session_755434585_dg8_VISp | session_755434585_dg8_VISp | 44 | accuracy | 0.716667 |
| classification | cycle_gan | Visual Coding | single_day | session_755434585_dg8_VISp | session_755434585_dg8_VISp | 46 | accuracy | 0.157407 |
| classification | cycle_gan | Visual Coding | single_day | session_755434585_dg8_VISp | session_755434585_dg8_VISp | 46 | accuracy | 0.691667 |
| classification | cycle_gan | Visual Coding | single_day | session_755434585_dg8_VISp | session_755434585_dg8_VISp | 54 | accuracy | 0.12037 |
| classification | cycle_gan | Visual Coding | single_day | session_755434585_dg8_VISp | session_755434585_dg8_VISp | 54 | accuracy | 0.616667 |
| classification | cycle_gan | Visual Coding | single_day | session_756029989_dg8_VISp | session_756029989_dg8_VISp | 44 | accuracy | 0.268519 |
| classification | cycle_gan | Visual Coding | single_day | session_756029989_dg8_VISp | session_756029989_dg8_VISp | 44 | accuracy | 0.783333 |
| classification | cycle_gan | Visual Coding | single_day | session_756029989_dg8_VISp | session_756029989_dg8_VISp | 46 | accuracy | 0.12963 |
| classification | cycle_gan | Visual Coding | single_day | session_756029989_dg8_VISp | session_756029989_dg8_VISp | 46 | accuracy | 0.775 |
| classification | cycle_gan | Visual Coding | single_day | session_756029989_dg8_VISp | session_756029989_dg8_VISp | 54 | accuracy | 0.157407 |
| classification | cycle_gan | Visual Coding | single_day | session_756029989_dg8_VISp | session_756029989_dg8_VISp | 54 | accuracy | 0.758333 |
| classification | cycle_gan | Visual Coding | single_day | session_757216464_dg8_VISp | session_757216464_dg8_VISp | 44 | accuracy | 0.111111 |
| classification | cycle_gan | Visual Coding | single_day | session_757216464_dg8_VISp | session_757216464_dg8_VISp | 44 | accuracy | 0.566667 |
| classification | cycle_gan | Visual Coding | single_day | session_757216464_dg8_VISp | session_757216464_dg8_VISp | 46 | accuracy | 0.175926 |
| classification | cycle_gan | Visual Coding | single_day | session_757216464_dg8_VISp | session_757216464_dg8_VISp | 46 | accuracy | 0.708333 |
| classification | cycle_gan | Visual Coding | single_day | session_757216464_dg8_VISp | session_757216464_dg8_VISp | 54 | accuracy | 0.12963 |
| classification | cycle_gan | Visual Coding | single_day | session_757216464_dg8_VISp | session_757216464_dg8_VISp | 54 | accuracy | 0.566667 |
| classification | cycle_gan | Visual Coding | single_day | session_757970808_dg8_VISp | session_757970808_dg8_VISp | 44 | accuracy | 0.194444 |
| classification | cycle_gan | Visual Coding | single_day | session_757970808_dg8_VISp | session_757970808_dg8_VISp | 44 | accuracy | 0.75 |
| classification | cycle_gan | Visual Coding | single_day | session_757970808_dg8_VISp | session_757970808_dg8_VISp | 46 | accuracy | 0.138889 |
| classification | cycle_gan | Visual Coding | single_day | session_757970808_dg8_VISp | session_757970808_dg8_VISp | 46 | accuracy | 0.816667 |
| classification | cycle_gan | Visual Coding | single_day | session_757970808_dg8_VISp | session_757970808_dg8_VISp | 54 | accuracy | 0.166667 |
| classification | cycle_gan | Visual Coding | single_day | session_757970808_dg8_VISp | session_757970808_dg8_VISp | 54 | accuracy | 0.7 |
| classification | cycle_gan | Visual Coding | single_day | session_758798717_dg8_VISp | session_758798717_dg8_VISp | 44 | accuracy | 0.12037 |
| classification | cycle_gan | Visual Coding | single_day | session_758798717_dg8_VISp | session_758798717_dg8_VISp | 44 | accuracy | 0.625 |
| classification | cycle_gan | Visual Coding | single_day | session_758798717_dg8_VISp | session_758798717_dg8_VISp | 46 | accuracy | 0.12963 |
| classification | cycle_gan | Visual Coding | single_day | session_758798717_dg8_VISp | session_758798717_dg8_VISp | 46 | accuracy | 0.65 |
| classification | cycle_gan | Visual Coding | single_day | session_758798717_dg8_VISp | session_758798717_dg8_VISp | 54 | accuracy | 0.425926 |
| classification | cycle_gan | Visual Coding | single_day | session_758798717_dg8_VISp | session_758798717_dg8_VISp | 54 | accuracy | 0.691667 |
| classification | cycle_gan | Visual Coding | single_day | session_759883607_dg8_VISp | session_759883607_dg8_VISp | 44 | accuracy | 0.138889 |
| classification | cycle_gan | Visual Coding | single_day | session_759883607_dg8_VISp | session_759883607_dg8_VISp | 44 | accuracy | 0.808333 |
| classification | cycle_gan | Visual Coding | single_day | session_759883607_dg8_VISp | session_759883607_dg8_VISp | 46 | accuracy | 0.12963 |
| classification | cycle_gan | Visual Coding | single_day | session_759883607_dg8_VISp | session_759883607_dg8_VISp | 46 | accuracy | 0.716667 |
| classification | cycle_gan | Visual Coding | single_day | session_759883607_dg8_VISp | session_759883607_dg8_VISp | 54 | accuracy | 0.166667 |
| classification | cycle_gan | Visual Coding | single_day | session_759883607_dg8_VISp | session_759883607_dg8_VISp | 54 | accuracy | 0.716667 |
| classification | cycle_gan | Visual Coding | single_day | session_760345702_dg8_VISp | session_760345702_dg8_VISp | 44 | accuracy | 0.157407 |
| classification | cycle_gan | Visual Coding | single_day | session_760345702_dg8_VISp | session_760345702_dg8_VISp | 44 | accuracy | 0.866667 |
| classification | cycle_gan | Visual Coding | single_day | session_760345702_dg8_VISp | session_760345702_dg8_VISp | 46 | accuracy | 0.185185 |
| classification | cycle_gan | Visual Coding | single_day | session_760345702_dg8_VISp | session_760345702_dg8_VISp | 46 | accuracy | 0.825 |
| classification | cycle_gan | Visual Coding | single_day | session_760345702_dg8_VISp | session_760345702_dg8_VISp | 54 | accuracy | 0.12037 |
| classification | cycle_gan | Visual Coding | single_day | session_760345702_dg8_VISp | session_760345702_dg8_VISp | 54 | accuracy | 0.825 |
| classification | cycle_gan | Visual Coding | single_day | session_760693773_dg8_VISp | session_760693773_dg8_VISp | 44 | accuracy | 0.138889 |
| classification | cycle_gan | Visual Coding | single_day | session_760693773_dg8_VISp | session_760693773_dg8_VISp | 44 | accuracy | 0.916667 |
| classification | cycle_gan | Visual Coding | single_day | session_760693773_dg8_VISp | session_760693773_dg8_VISp | 46 | accuracy | 0.12963 |
| classification | cycle_gan | Visual Coding | single_day | session_760693773_dg8_VISp | session_760693773_dg8_VISp | 46 | accuracy | 0.875 |
| classification | cycle_gan | Visual Coding | single_day | session_760693773_dg8_VISp | session_760693773_dg8_VISp | 54 | accuracy | 0.25 |
| classification | cycle_gan | Visual Coding | single_day | session_760693773_dg8_VISp | session_760693773_dg8_VISp | 54 | accuracy | 0.883333 |
| classification | cycle_gan | Visual Coding | single_day | session_761418226_dg8_VISp | session_761418226_dg8_VISp | 44 | accuracy | 0.203704 |
| classification | cycle_gan | Visual Coding | single_day | session_761418226_dg8_VISp | session_761418226_dg8_VISp | 44 | accuracy | 0.733333 |

| | | | | | | | | |
|---|---|---|---|---|---|---|---|---|
| classification | cycle_gan | Visual Coding | single_day | session_761418226_dg8_VISp | session_761418226_dg8_VISp | 46 | accuracy | 0.111111 |
| classification | cycle_gan | Visual Coding | single_day | session_761418226_dg8_VISp | session_761418226_dg8_VISp | 46 | accuracy | 0.716667 |
| classification | cycle_gan | Visual Coding | single_day | session_761418226_dg8_VISp | session_761418226_dg8_VISp | 54 | accuracy | 0.185185 |
| classification | cycle_gan | Visual Coding | single_day | session_761418226_dg8_VISp | session_761418226_dg8_VISp | 54 | accuracy | 0.708333 |
| classification | cycle_gan | Visual Coding | single_day | session_762120172_dg8_VISp | session_762120172_dg8_VISp | 44 | accuracy | 0.12037 |
| classification | cycle_gan | Visual Coding | single_day | session_762120172_dg8_VISp | session_762120172_dg8_VISp | 44 | accuracy | 0.791667 |
| classification | cycle_gan | Visual Coding | single_day | session_762120172_dg8_VISp | session_762120172_dg8_VISp | 46 | accuracy | 0.148148 |
| classification | cycle_gan | Visual Coding | single_day | session_762120172_dg8_VISp | session_762120172_dg8_VISp | 46 | accuracy | 0.833333 |
| classification | cycle_gan | Visual Coding | single_day | session_762120172_dg8_VISp | session_762120172_dg8_VISp | 54 | accuracy | 0.12037 |
| classification | cycle_gan | Visual Coding | single_day | session_762120172_dg8_VISp | session_762120172_dg8_VISp | 54 | accuracy | 0.8 |
| classification | cycle_gan | Visual Coding | single_day | session_762602078_dg8_VISp | session_762602078_dg8_VISp | 44 | accuracy | 0.166667 |
| classification | cycle_gan | Visual Coding | single_day | session_762602078_dg8_VISp | session_762602078_dg8_VISp | 44 | accuracy | 0.983333 |
| classification | cycle_gan | Visual Coding | single_day | session_762602078_dg8_VISp | session_762602078_dg8_VISp | 46 | accuracy | 0.425926 |
| classification | cycle_gan | Visual Coding | single_day | session_762602078_dg8_VISp | session_762602078_dg8_VISp | 46 | accuracy | 0.966667 |
| classification | cycle_gan | Visual Coding | single_day | session_762602078_dg8_VISp | session_762602078_dg8_VISp | 54 | accuracy | 0.537037 |
| classification | cycle_gan | Visual Coding | single_day | session_762602078_dg8_VISp | session_762602078_dg8_VISp | 54 | accuracy | 0.941667 |
| classification | cycle_gan | Visual Coding | single_day | session_763673393_dg8_VISp | session_763673393_dg8_VISp | 44 | accuracy | 0.138889 |
| classification | cycle_gan | Visual Coding | single_day | session_763673393_dg8_VISp | session_763673393_dg8_VISp | 44 | accuracy | 0.7 |
| classification | cycle_gan | Visual Coding | single_day | session_763673393_dg8_VISp | session_763673393_dg8_VISp | 46 | accuracy | 0.0740741 |
| classification | cycle_gan | Visual Coding | single_day | session_763673393_dg8_VISp | session_763673393_dg8_VISp | 46 | accuracy | 0.741667 |
| classification | cycle_gan | Visual Coding | single_day | session_763673393_dg8_VISp | session_763673393_dg8_VISp | 54 | accuracy | 0.148148 |
| classification | cycle_gan | Visual Coding | single_day | session_763673393_dg8_VISp | session_763673393_dg8_VISp | 54 | accuracy | 0.716667 |
| classification | cycle_gan | Visual Coding | single_day | session_773418906_dg8_VISp | session_773418906_dg8_VISp | 44 | accuracy | 0.111111 |
| classification | cycle_gan | Visual Coding | single_day | session_773418906_dg8_VISp | session_773418906_dg8_VISp | 44 | accuracy | 0.533333 |
| classification | cycle_gan | Visual Coding | single_day | session_773418906_dg8_VISp | session_773418906_dg8_VISp | 46 | accuracy | 0.111111 |
| classification | cycle_gan | Visual Coding | single_day | session_773418906_dg8_VISp | session_773418906_dg8_VISp | 46 | accuracy | 0.508333 |
| classification | cycle_gan | Visual Coding | single_day | session_773418906_dg8_VISp | session_773418906_dg8_VISp | 54 | accuracy | 0.111111 |
| classification | cycle_gan | Visual Coding | single_day | session_773418906_dg8_VISp | session_773418906_dg8_VISp | 54 | accuracy | 0.516667 |
| classification | cycle_gan | Visual Coding | single_day | session_791319847_dg8_VISp | session_791319847_dg8_VISp | 44 | accuracy | 0.157407 |
| classification | cycle_gan | Visual Coding | single_day | session_791319847_dg8_VISp | session_791319847_dg8_VISp | 44 | accuracy | 0.833333 |
| classification | cycle_gan | Visual Coding | single_day | session_791319847_dg8_VISp | session_791319847_dg8_VISp | 46 | accuracy | 0.203704 |
| classification | cycle_gan | Visual Coding | single_day | session_791319847_dg8_VISp | session_791319847_dg8_VISp | 46 | accuracy | 0.883333 |
| classification | cycle_gan | Visual Coding | single_day | session_791319847_dg8_VISp | session_791319847_dg8_VISp | 54 | accuracy | 0.175926 |
| classification | cycle_gan | Visual Coding | single_day | session_791319847_dg8_VISp | session_791319847_dg8_VISp | 54 | accuracy | 0.883333 |
| classification | cycle_gan | Visual Coding | single_day | session_797828357_dg8_VISp | session_797828357_dg8_VISp | 44 | accuracy | 0.240741 |
| classification | cycle_gan | Visual Coding | single_day | session_797828357_dg8_VISp | session_797828357_dg8_VISp | 44 | accuracy | 0.608333 |
| classification | cycle_gan | Visual Coding | single_day | session_797828357_dg8_VISp | session_797828357_dg8_VISp | 46 | accuracy | 0.231481 |
| classification | cycle_gan | Visual Coding | single_day | session_797828357_dg8_VISp | session_797828357_dg8_VISp | 46 | accuracy | 0.675 |
| classification | cycle_gan | Visual Coding | single_day | session_797828357_dg8_VISp | session_797828357_dg8_VISp | 54 | accuracy | 0.148148 |
| classification | cycle_gan | Visual Coding | single_day | session_797828357_dg8_VISp | session_797828357_dg8_VISp | 54 | accuracy | 0.616667 |
| classification | cycle_gan | Visual Coding | single_day | session_798911424_dg8_VISp | session_798911424_dg8_VISp | 44 | accuracy | 0.12037 |
| classification | cycle_gan | Visual Coding | single_day | session_798911424_dg8_VISp | session_798911424_dg8_VISp | 44 | accuracy | 0.775 |
| classification | cycle_gan | Visual Coding | single_day | session_798911424_dg8_VISp | session_798911424_dg8_VISp | 46 | accuracy | 0.203704 |
| classification | cycle_gan | Visual Coding | single_day | session_798911424_dg8_VISp | session_798911424_dg8_VISp | 46 | accuracy | 0.875 |
| classification | cycle_gan | Visual Coding | single_day | session_798911424_dg8_VISp | session_798911424_dg8_VISp | 54 | accuracy | 0.111111 |
| classification | cycle_gan | Visual Coding | single_day | session_798911424_dg8_VISp | session_798911424_dg8_VISp | 54 | accuracy | 0.766667 |
| classification | cycle_gan | Visual Coding | single_day | session_799864342_dg8_VISp | session_799864342_dg8_VISp | 44 | accuracy | 0.222222 |
| classification | cycle_gan | Visual Coding | single_day | session_799864342_dg8_VISp | session_799864342_dg8_VISp | 44 | accuracy | 0.908333 |
| classification | cycle_gan | Visual Coding | single_day | session_799864342_dg8_VISp | session_799864342_dg8_VISp | 46 | accuracy | 0.101852 |
| classification | cycle_gan | Visual Coding | single_day | session_799864342_dg8_VISp | session_799864342_dg8_VISp | 46 | accuracy | 0.891667 |
| classification | cycle_gan | Visual Coding | single_day | session_799864342_dg8_VISp | session_799864342_dg8_VISp | 54 | accuracy | 0.12037 |
| classification | cycle_gan | Visual Coding | single_day | session_799864342_dg8_VISp | session_799864342_dg8_VISp | 54 | accuracy | 0.9 |
| classification | stabilization | Visual Coding | single_day | session_715093703_dg8_VISp | session_715093703_dg8_VISp | 44 | accuracy | 0.37037 |
| classification | stabilization | Visual Coding | single_day | session_715093703_dg8_VISp | session_715093703_dg8_VISp | 44 | accuracy | 0.683333 |
| classification | stabilization | Visual Coding | single_day | session_715093703_dg8_VISp | session_715093703_dg8_VISp | 46 | accuracy | 0.212963 |
| classification | stabilization | Visual Coding | single_day | session_715093703_dg8_VISp | session_715093703_dg8_VISp | 46 | accuracy | 0.583333 |
| classification | stabilization | Visual Coding | single_day | session_715093703_dg8_VISp | session_715093703_dg8_VISp | 54 | accuracy | 0.259259 |
| classification | stabilization | Visual Coding | single_day | session_715093703_dg8_VISp | session_715093703_dg8_VISp | 54 | accuracy | 0.641667 |
| classification | stabilization | Visual Coding | single_day | session_719161530_dg8_VISp | session_719161530_dg8_VISp | 44 | accuracy | 0.25 |
| classification | stabilization | Visual Coding | single_day | session_719161530_dg8_VISp | session_719161530_dg8_VISp | 44 | accuracy | 0.791667 |
| classification | stabilization | Visual Coding | single_day | session_719161530_dg8_VISp | session_719161530_dg8_VISp | 46 | accuracy | 0.296296 |
| classification | stabilization | Visual Coding | single_day | session_719161530_dg8_VISp | session_719161530_dg8_VISp | 46 | accuracy | 0.716667 |
| classification | stabilization | Visual Coding | single_day | session_719161530_dg8_VISp | session_719161530_dg8_VISp | 54 | accuracy | 0.351852 |

| | | | | | | | | |
|---|---|---|---|---|---|---|---|---|
| classification | stabilization | Visual Coding | single_day | session_719161530_dg8_VISp | session_719161530_dg8_VISp | 54 | accuracy | 0.816667 |
| classification | stabilization | Visual Coding | single_day | session_721123822_dg8_VISp | session_721123822_dg8_VISp | 44 | accuracy | 0.175926 |
| classification | stabilization | Visual Coding | single_day | session_721123822_dg8_VISp | session_721123822_dg8_VISp | 44 | accuracy | 0.741667 |
| classification | stabilization | Visual Coding | single_day | session_721123822_dg8_VISp | session_721123822_dg8_VISp | 46 | accuracy | 0.194444 |
| classification | stabilization | Visual Coding | single_day | session_721123822_dg8_VISp | session_721123822_dg8_VISp | 46 | accuracy | 0.658333 |
| classification | stabilization | Visual Coding | single_day | session_721123822_dg8_VISp | session_721123822_dg8_VISp | 54 | accuracy | 0.222222 |
| classification | stabilization | Visual Coding | single_day | session_721123822_dg8_VISp | session_721123822_dg8_VISp | 54 | accuracy | 0.616667 |
| classification | stabilization | Visual Coding | single_day | session_732592105_dg8_VISp | session_732592105_dg8_VISp | 44 | accuracy | 0.268519 |
| classification | stabilization | Visual Coding | single_day | session_732592105_dg8_VISp | session_732592105_dg8_VISp | 44 | accuracy | 0.683333 |
| classification | stabilization | Visual Coding | single_day | session_732592105_dg8_VISp | session_732592105_dg8_VISp | 46 | accuracy | 0.296296 |
| classification | stabilization | Visual Coding | single_day | session_732592105_dg8_VISp | session_732592105_dg8_VISp | 46 | accuracy | 0.641667 |
| classification | stabilization | Visual Coding | single_day | session_732592105_dg8_VISp | session_732592105_dg8_VISp | 54 | accuracy | 0.212963 |
| classification | stabilization | Visual Coding | single_day | session_732592105_dg8_VISp | session_732592105_dg8_VISp | 54 | accuracy | 0.675 |
| classification | stabilization | Visual Coding | single_day | session_737581020_dg8_VISp | session_737581020_dg8_VISp | 44 | accuracy | 0.435185 |
| classification | stabilization | Visual Coding | single_day | session_737581020_dg8_VISp | session_737581020_dg8_VISp | 44 | accuracy | 0.725 |
| classification | stabilization | Visual Coding | single_day | session_737581020_dg8_VISp | session_737581020_dg8_VISp | 46 | accuracy | 0.435185 |
| classification | stabilization | Visual Coding | single_day | session_737581020_dg8_VISp | session_737581020_dg8_VISp | 46 | accuracy | 0.733333 |
| classification | stabilization | Visual Coding | single_day | session_737581020_dg8_VISp | session_737581020_dg8_VISp | 54 | accuracy | 0.361111 |
| classification | stabilization | Visual Coding | single_day | session_737581020_dg8_VISp | session_737581020_dg8_VISp | 54 | accuracy | 0.766667 |
| classification | stabilization | Visual Coding | single_day | session_739448407_dg8_VISp | session_739448407_dg8_VISp | 44 | accuracy | 0.101852 |
| classification | stabilization | Visual Coding | single_day | session_739448407_dg8_VISp | session_739448407_dg8_VISp | 44 | accuracy | 0.462185 |
| classification | stabilization | Visual Coding | single_day | session_739448407_dg8_VISp | session_739448407_dg8_VISp | 46 | accuracy | 0.231481 |
| classification | stabilization | Visual Coding | single_day | session_739448407_dg8_VISp | session_739448407_dg8_VISp | 46 | accuracy | 0.546218 |
| classification | stabilization | Visual Coding | single_day | session_739448407_dg8_VISp | session_739448407_dg8_VISp | 54 | accuracy | 0.175926 |
| classification | stabilization | Visual Coding | single_day | session_739448407_dg8_VISp | session_739448407_dg8_VISp | 54 | accuracy | 0.478992 |
| classification | stabilization | Visual Coding | single_day | session_742951821_dg8_VISp | session_742951821_dg8_VISp | 44 | accuracy | 0.175926 |
| classification | stabilization | Visual Coding | single_day | session_742951821_dg8_VISp | session_742951821_dg8_VISp | 44 | accuracy | 0.466667 |
| classification | stabilization | Visual Coding | single_day | session_742951821_dg8_VISp | session_742951821_dg8_VISp | 46 | accuracy | 0.185185 |
| classification | stabilization | Visual Coding | single_day | session_742951821_dg8_VISp | session_742951821_dg8_VISp | 46 | accuracy | 0.466667 |
| classification | stabilization | Visual Coding | single_day | session_742951821_dg8_VISp | session_742951821_dg8_VISp | 54 | accuracy | 0.175926 |
| classification | stabilization | Visual Coding | single_day | session_742951821_dg8_VISp | session_742951821_dg8_VISp | 54 | accuracy | 0.508333 |
| classification | stabilization | Visual Coding | single_day | session_743475441_dg8_VISp | session_743475441_dg8_VISp | 44 | accuracy | 0.12963 |
| classification | stabilization | Visual Coding | single_day | session_743475441_dg8_VISp | session_743475441_dg8_VISp | 44 | accuracy | 0.258333 |
| classification | stabilization | Visual Coding | single_day | session_743475441_dg8_VISp | session_743475441_dg8_VISp | 46 | accuracy | 0.148148 |
| classification | stabilization | Visual Coding | single_day | session_743475441_dg8_VISp | session_743475441_dg8_VISp | 46 | accuracy | 0.325 |
| classification | stabilization | Visual Coding | single_day | session_743475441_dg8_VISp | session_743475441_dg8_VISp | 54 | accuracy | 0.157407 |
| classification | stabilization | Visual Coding | single_day | session_743475441_dg8_VISp | session_743475441_dg8_VISp | 54 | accuracy | 0.283333 |
| classification | stabilization | Visual Coding | single_day | session_744228101_dg8_VISp | session_744228101_dg8_VISp | 44 | accuracy | 0.222222 |
| classification | stabilization | Visual Coding | single_day | session_744228101_dg8_VISp | session_744228101_dg8_VISp | 44 | accuracy | 0.683333 |
| classification | stabilization | Visual Coding | single_day | session_744228101_dg8_VISp | session_744228101_dg8_VISp | 46 | accuracy | 0.203704 |
| classification | stabilization | Visual Coding | single_day | session_744228101_dg8_VISp | session_744228101_dg8_VISp | 46 | accuracy | 0.716667 |
| classification | stabilization | Visual Coding | single_day | session_744228101_dg8_VISp | session_744228101_dg8_VISp | 54 | accuracy | 0.231481 |
| classification | stabilization | Visual Coding | single_day | session_744228101_dg8_VISp | session_744228101_dg8_VISp | 54 | accuracy | 0.741667 |
| classification | stabilization | Visual Coding | single_day | session_746083955_dg8_VISp | session_746083955_dg8_VISp | 44 | accuracy | 0.175926 |
| classification | stabilization | Visual Coding | single_day | session_746083955_dg8_VISp | session_746083955_dg8_VISp | 44 | accuracy | 0.4 |
| classification | stabilization | Visual Coding | single_day | session_746083955_dg8_VISp | session_746083955_dg8_VISp | 46 | accuracy | 0.166667 |
| classification | stabilization | Visual Coding | single_day | session_746083955_dg8_VISp | session_746083955_dg8_VISp | 46 | accuracy | 0.45 |
| classification | stabilization | Visual Coding | single_day | session_746083955_dg8_VISp | session_746083955_dg8_VISp | 54 | accuracy | 0.12963 |
| classification | stabilization | Visual Coding | single_day | session_746083955_dg8_VISp | session_746083955_dg8_VISp | 54 | accuracy | 0.366667 |
| classification | stabilization | Visual Coding | single_day | session_750332458_dg8_VISp | session_750332458_dg8_VISp | 44 | accuracy | 0.5 |
| classification | stabilization | Visual Coding | single_day | session_750332458_dg8_VISp | session_750332458_dg8_VISp | 44 | accuracy | 0.75 |
| classification | stabilization | Visual Coding | single_day | session_750332458_dg8_VISp | session_750332458_dg8_VISp | 46 | accuracy | 0.324074 |
| classification | stabilization | Visual Coding | single_day | session_750332458_dg8_VISp | session_750332458_dg8_VISp | 46 | accuracy | 0.758333 |
| classification | stabilization | Visual Coding | single_day | session_750332458_dg8_VISp | session_750332458_dg8_VISp | 54 | accuracy | 0.314815 |
| classification | stabilization | Visual Coding | single_day | session_750332458_dg8_VISp | session_750332458_dg8_VISp | 54 | accuracy | 0.825 |
| classification | stabilization | Visual Coding | single_day | session_750749662_dg8_VISp | session_750749662_dg8_VISp | 44 | accuracy | 0.231481 |
| classification | stabilization | Visual Coding | single_day | session_750749662_dg8_VISp | session_750749662_dg8_VISp | 44 | accuracy | 0.7 |
| classification | stabilization | Visual Coding | single_day | session_750749662_dg8_VISp | session_750749662_dg8_VISp | 46 | accuracy | 0.203704 |
| classification | stabilization | Visual Coding | single_day | session_750749662_dg8_VISp | session_750749662_dg8_VISp | 46 | accuracy | 0.65 |
| classification | stabilization | Visual Coding | single_day | session_750749662_dg8_VISp | session_750749662_dg8_VISp | 54 | accuracy | 0.175926 |
| classification | stabilization | Visual Coding | single_day | session_750749662_dg8_VISp | session_750749662_dg8_VISp | 54 | accuracy | 0.625 |
| classification | stabilization | Visual Coding | single_day | session_751348571_dg8_VISp | session_751348571_dg8_VISp | 44 | accuracy | 0.138889 |
| classification | stabilization | Visual Coding | single_day | session_751348571_dg8_VISp | session_751348571_dg8_VISp | 44 | accuracy | 0.816667 |

| | | | | | | | | |
|---|---|---|---|---|---|---|---|---|
| classification | stabilization | Visual Coding | single_day | session_751348571_dg8_VISp | session_751348571_dg8_VISp | 46 | accuracy | 0.175926 |
| classification | stabilization | Visual Coding | single_day | session_751348571_dg8_VISp | session_751348571_dg8_VISp | 46 | accuracy | 0.808333 |
| classification | stabilization | Visual Coding | single_day | session_751348571_dg8_VISp | session_751348571_dg8_VISp | 54 | accuracy | 0.138889 |
| classification | stabilization | Visual Coding | single_day | session_751348571_dg8_VISp | session_751348571_dg8_VISp | 54 | accuracy | 0.75 |
| classification | stabilization | Visual Coding | single_day | session_754312389_dg8_VISp | session_754312389_dg8_VISp | 44 | accuracy | 0.37963 |
| classification | stabilization | Visual Coding | single_day | session_754312389_dg8_VISp | session_754312389_dg8_VISp | 44 | accuracy | 0.841667 |
| classification | stabilization | Visual Coding | single_day | session_754312389_dg8_VISp | session_754312389_dg8_VISp | 46 | accuracy | 0.296296 |
| classification | stabilization | Visual Coding | single_day | session_754312389_dg8_VISp | session_754312389_dg8_VISp | 46 | accuracy | 0.783333 |
| classification | stabilization | Visual Coding | single_day | session_754312389_dg8_VISp | session_754312389_dg8_VISp | 54 | accuracy | 0.342593 |
| classification | stabilization | Visual Coding | single_day | session_754312389_dg8_VISp | session_754312389_dg8_VISp | 54 | accuracy | 0.808333 |
| classification | stabilization | Visual Coding | single_day | session_754829445_dg8_VISp | session_754829445_dg8_VISp | 44 | accuracy | 0.398148 |
| classification | stabilization | Visual Coding | single_day | session_754829445_dg8_VISp | session_754829445_dg8_VISp | 44 | accuracy | 0.716667 |
| classification | stabilization | Visual Coding | single_day | session_754829445_dg8_VISp | session_754829445_dg8_VISp | 46 | accuracy | 0.342593 |
| classification | stabilization | Visual Coding | single_day | session_754829445_dg8_VISp | session_754829445_dg8_VISp | 46 | accuracy | 0.691667 |
| classification | stabilization | Visual Coding | single_day | session_754829445_dg8_VISp | session_754829445_dg8_VISp | 54 | accuracy | 0.546296 |
| classification | stabilization | Visual Coding | single_day | session_754829445_dg8_VISp | session_754829445_dg8_VISp | 54 | accuracy | 0.75 |
| classification | stabilization | Visual Coding | single_day | session_755434585_dg8_VISp | session_755434585_dg8_VISp | 44 | accuracy | 0.222222 |
| classification | stabilization | Visual Coding | single_day | session_755434585_dg8_VISp | session_755434585_dg8_VISp | 44 | accuracy | 0.725 |
| classification | stabilization | Visual Coding | single_day | session_755434585_dg8_VISp | session_755434585_dg8_VISp | 46 | accuracy | 0.342593 |
| classification | stabilization | Visual Coding | single_day | session_755434585_dg8_VISp | session_755434585_dg8_VISp | 46 | accuracy | 0.616667 |
| classification | stabilization | Visual Coding | single_day | session_755434585_dg8_VISp | session_755434585_dg8_VISp | 54 | accuracy | 0.351852 |
| classification | stabilization | Visual Coding | single_day | session_755434585_dg8_VISp | session_755434585_dg8_VISp | 54 | accuracy | 0.658333 |
| classification | stabilization | Visual Coding | single_day | session_756029989_dg8_VISp | session_756029989_dg8_VISp | 44 | accuracy | 0.12037 |
| classification | stabilization | Visual Coding | single_day | session_756029989_dg8_VISp | session_756029989_dg8_VISp | 44 | accuracy | 0.775 |
| classification | stabilization | Visual Coding | single_day | session_756029989_dg8_VISp | session_756029989_dg8_VISp | 46 | accuracy | 0.194444 |
| classification | stabilization | Visual Coding | single_day | session_756029989_dg8_VISp | session_756029989_dg8_VISp | 46 | accuracy | 0.791667 |
| classification | stabilization | Visual Coding | single_day | session_756029989_dg8_VISp | session_756029989_dg8_VISp | 54 | accuracy | 0.305556 |
| classification | stabilization | Visual Coding | single_day | session_756029989_dg8_VISp | session_756029989_dg8_VISp | 54 | accuracy | 0.783333 |
| classification | stabilization | Visual Coding | single_day | session_757216464_dg8_VISp | session_757216464_dg8_VISp | 44 | accuracy | 0.361111 |
| classification | stabilization | Visual Coding | single_day | session_757216464_dg8_VISp | session_757216464_dg8_VISp | 44 | accuracy | 0.7 |
| classification | stabilization | Visual Coding | single_day | session_757216464_dg8_VISp | session_757216464_dg8_VISp | 46 | accuracy | 0.5 |
| classification | stabilization | Visual Coding | single_day | session_757216464_dg8_VISp | session_757216464_dg8_VISp | 46 | accuracy | 0.658333 |
| classification | stabilization | Visual Coding | single_day | session_757216464_dg8_VISp | session_757216464_dg8_VISp | 54 | accuracy | 0.314815 |
| classification | stabilization | Visual Coding | single_day | session_757216464_dg8_VISp | session_757216464_dg8_VISp | 54 | accuracy | 0.633333 |
| classification | stabilization | Visual Coding | single_day | session_757970808_dg8_VISp | session_757970808_dg8_VISp | 44 | accuracy | 0.166667 |
| classification | stabilization | Visual Coding | single_day | session_757970808_dg8_VISp | session_757970808_dg8_VISp | 44 | accuracy | 0.733333 |
| classification | stabilization | Visual Coding | single_day | session_757970808_dg8_VISp | session_757970808_dg8_VISp | 46 | accuracy | 0.194444 |
| classification | stabilization | Visual Coding | single_day | session_757970808_dg8_VISp | session_757970808_dg8_VISp | 46 | accuracy | 0.791667 |
| classification | stabilization | Visual Coding | single_day | session_757970808_dg8_VISp | session_757970808_dg8_VISp | 54 | accuracy | 0.240741 |
| classification | stabilization | Visual Coding | single_day | session_757970808_dg8_VISp | session_757970808_dg8_VISp | 54 | accuracy | 0.75 |
| classification | stabilization | Visual Coding | single_day | session_758798717_dg8_VISp | session_758798717_dg8_VISp | 44 | accuracy | 0.342593 |
| classification | stabilization | Visual Coding | single_day | session_758798717_dg8_VISp | session_758798717_dg8_VISp | 44 | accuracy | 0.65 |
| classification | stabilization | Visual Coding | single_day | session_758798717_dg8_VISp | session_758798717_dg8_VISp | 46 | accuracy | 0.361111 |
| classification | stabilization | Visual Coding | single_day | session_758798717_dg8_VISp | session_758798717_dg8_VISp | 46 | accuracy | 0.633333 |
| classification | stabilization | Visual Coding | single_day | session_758798717_dg8_VISp | session_758798717_dg8_VISp | 54 | accuracy | 0.296296 |
| classification | stabilization | Visual Coding | single_day | session_758798717_dg8_VISp | session_758798717_dg8_VISp | 54 | accuracy | 0.666667 |
| classification | stabilization | Visual Coding | single_day | session_759883607_dg8_VISp | session_759883607_dg8_VISp | 44 | accuracy | 0.277778 |
| classification | stabilization | Visual Coding | single_day | session_759883607_dg8_VISp | session_759883607_dg8_VISp | 44 | accuracy | 0.675 |
| classification | stabilization | Visual Coding | single_day | session_759883607_dg8_VISp | session_759883607_dg8_VISp | 46 | accuracy | 0.166667 |
| classification | stabilization | Visual Coding | single_day | session_759883607_dg8_VISp | session_759883607_dg8_VISp | 46 | accuracy | 0.691667 |
| classification | stabilization | Visual Coding | single_day | session_759883607_dg8_VISp | session_759883607_dg8_VISp | 54 | accuracy | 0.194444 |
| classification | stabilization | Visual Coding | single_day | session_759883607_dg8_VISp | session_759883607_dg8_VISp | 54 | accuracy | 0.683333 |
| classification | stabilization | Visual Coding | single_day | session_760345702_dg8_VISp | session_760345702_dg8_VISp | 44 | accuracy | 0.361111 |
| classification | stabilization | Visual Coding | single_day | session_760345702_dg8_VISp | session_760345702_dg8_VISp | 44 | accuracy | 0.825 |
| classification | stabilization | Visual Coding | single_day | session_760345702_dg8_VISp | session_760345702_dg8_VISp | 46 | accuracy | 0.351852 |
| classification | stabilization | Visual Coding | single_day | session_760345702_dg8_VISp | session_760345702_dg8_VISp | 46 | accuracy | 0.816667 |
| classification | stabilization | Visual Coding | single_day | session_760345702_dg8_VISp | session_760345702_dg8_VISp | 54 | accuracy | 0.25 |
| classification | stabilization | Visual Coding | single_day | session_760345702_dg8_VISp | session_760345702_dg8_VISp | 54 | accuracy | 0.733333 |
| classification | stabilization | Visual Coding | single_day | session_760693773_dg8_VISp | session_760693773_dg8_VISp | 44 | accuracy | 0.361111 |
| classification | stabilization | Visual Coding | single_day | session_760693773_dg8_VISp | session_760693773_dg8_VISp | 44 | accuracy | 0.866667 |
| classification | stabilization | Visual Coding | single_day | session_760693773_dg8_VISp | session_760693773_dg8_VISp | 46 | accuracy | 0.333333 |
| classification | stabilization | Visual Coding | single_day | session_760693773_dg8_VISp | session_760693773_dg8_VISp | 46 | accuracy | 0.841667 |
| classification | stabilization | Visual Coding | single_day | session_760693773_dg8_VISp | session_760693773_dg8_VISp | 54 | accuracy | 0.564815 |

| | | | | | | | | |
|---|---|---|---|---|---|---|---|---|
| classification | stabilization | Visual Coding | single_day | session_760693773_dg8_VISp | session_760693773_dg8_VISp | 54 | accuracy | 0.816667 |
| classification | stabilization | Visual Coding | single_day | session_761418226_dg8_VISp | session_761418226_dg8_VISp | 44 | accuracy | 0.175926 |
| classification | stabilization | Visual Coding | single_day | session_761418226_dg8_VISp | session_761418226_dg8_VISp | 44 | accuracy | 0.741667 |
| classification | stabilization | Visual Coding | single_day | session_761418226_dg8_VISp | session_761418226_dg8_VISp | 46 | accuracy | 0.138889 |
| classification | stabilization | Visual Coding | single_day | session_761418226_dg8_VISp | session_761418226_dg8_VISp | 46 | accuracy | 0.708333 |
| classification | stabilization | Visual Coding | single_day | session_761418226_dg8_VISp | session_761418226_dg8_VISp | 54 | accuracy | 0.194444 |
| classification | stabilization | Visual Coding | single_day | session_761418226_dg8_VISp | session_761418226_dg8_VISp | 54 | accuracy | 0.716667 |
| classification | stabilization | Visual Coding | single_day | session_762120172_dg8_VISp | session_762120172_dg8_VISp | 44 | accuracy | 0.212963 |
| classification | stabilization | Visual Coding | single_day | session_762120172_dg8_VISp | session_762120172_dg8_VISp | 44 | accuracy | 0.766667 |
| classification | stabilization | Visual Coding | single_day | session_762120172_dg8_VISp | session_762120172_dg8_VISp | 46 | accuracy | 0.240741 |
| classification | stabilization | Visual Coding | single_day | session_762120172_dg8_VISp | session_762120172_dg8_VISp | 46 | accuracy | 0.733333 |
| classification | stabilization | Visual Coding | single_day | session_762120172_dg8_VISp | session_762120172_dg8_VISp | 54 | accuracy | 0.277778 |
| classification | stabilization | Visual Coding | single_day | session_762120172_dg8_VISp | session_762120172_dg8_VISp | 54 | accuracy | 0.708333 |
| classification | stabilization | Visual Coding | single_day | session_762602078_dg8_VISp | session_762602078_dg8_VISp | 44 | accuracy | 0.731481 |
| classification | stabilization | Visual Coding | single_day | session_762602078_dg8_VISp | session_762602078_dg8_VISp | 44 | accuracy | 0.95 |
| classification | stabilization | Visual Coding | single_day | session_762602078_dg8_VISp | session_762602078_dg8_VISp | 46 | accuracy | 0.407407 |
| classification | stabilization | Visual Coding | single_day | session_762602078_dg8_VISp | session_762602078_dg8_VISp | 46 | accuracy | 0.883333 |
| classification | stabilization | Visual Coding | single_day | session_762602078_dg8_VISp | session_762602078_dg8_VISp | 54 | accuracy | 0.675926 |
| classification | stabilization | Visual Coding | single_day | session_762602078_dg8_VISp | session_762602078_dg8_VISp | 54 | accuracy | 0.883333 |
| classification | stabilization | Visual Coding | single_day | session_763673393_dg8_VISp | session_763673393_dg8_VISp | 44 | accuracy | 0.509259 |
| classification | stabilization | Visual Coding | single_day | session_763673393_dg8_VISp | session_763673393_dg8_VISp | 44 | accuracy | 0.65 |
| classification | stabilization | Visual Coding | single_day | session_763673393_dg8_VISp | session_763673393_dg8_VISp | 46 | accuracy | 0.518519 |
| classification | stabilization | Visual Coding | single_day | session_763673393_dg8_VISp | session_763673393_dg8_VISp | 46 | accuracy | 0.733333 |
| classification | stabilization | Visual Coding | single_day | session_763673393_dg8_VISp | session_763673393_dg8_VISp | 54 | accuracy | 0.407407 |
| classification | stabilization | Visual Coding | single_day | session_763673393_dg8_VISp | session_763673393_dg8_VISp | 54 | accuracy | 0.625 |
| classification | stabilization | Visual Coding | single_day | session_773418906_dg8_VISp | session_773418906_dg8_VISp | 44 | accuracy | 0.194444 |
| classification | stabilization | Visual Coding | single_day | session_773418906_dg8_VISp | session_773418906_dg8_VISp | 44 | accuracy | 0.608333 |
| classification | stabilization | Visual Coding | single_day | session_773418906_dg8_VISp | session_773418906_dg8_VISp | 46 | accuracy | 0.185185 |
| classification | stabilization | Visual Coding | single_day | session_773418906_dg8_VISp | session_773418906_dg8_VISp | 46 | accuracy | 0.591667 |
| classification | stabilization | Visual Coding | single_day | session_773418906_dg8_VISp | session_773418906_dg8_VISp | 54 | accuracy | 0.185185 |
| classification | stabilization | Visual Coding | single_day | session_773418906_dg8_VISp | session_773418906_dg8_VISp | 54 | accuracy | 0.566667 |
| classification | stabilization | Visual Coding | single_day | session_791319847_dg8_VISp | session_791319847_dg8_VISp | 44 | accuracy | 0.351852 |
| classification | stabilization | Visual Coding | single_day | session_791319847_dg8_VISp | session_791319847_dg8_VISp | 44 | accuracy | 0.841667 |
| classification | stabilization | Visual Coding | single_day | session_791319847_dg8_VISp | session_791319847_dg8_VISp | 46 | accuracy | 0.37037 |
| classification | stabilization | Visual Coding | single_day | session_791319847_dg8_VISp | session_791319847_dg8_VISp | 46 | accuracy | 0.866667 |
| classification | stabilization | Visual Coding | single_day | session_791319847_dg8_VISp | session_791319847_dg8_VISp | 54 | accuracy | 0.203704 |
| classification | stabilization | Visual Coding | single_day | session_791319847_dg8_VISp | session_791319847_dg8_VISp | 54 | accuracy | 0.816667 |
| classification | stabilization | Visual Coding | single_day | session_797828357_dg8_VISp | session_797828357_dg8_VISp | 44 | accuracy | 0.472222 |
| classification | stabilization | Visual Coding | single_day | session_797828357_dg8_VISp | session_797828357_dg8_VISp | 44 | accuracy | 0.608333 |
| classification | stabilization | Visual Coding | single_day | session_797828357_dg8_VISp | session_797828357_dg8_VISp | 46 | accuracy | 0.435185 |
| classification | stabilization | Visual Coding | single_day | session_797828357_dg8_VISp | session_797828357_dg8_VISp | 46 | accuracy | 0.633333 |
| classification | stabilization | Visual Coding | single_day | session_797828357_dg8_VISp | session_797828357_dg8_VISp | 54 | accuracy | 0.324074 |
| classification | stabilization | Visual Coding | single_day | session_797828357_dg8_VISp | session_797828357_dg8_VISp | 54 | accuracy | 0.683333 |
| classification | stabilization | Visual Coding | single_day | session_798911424_dg8_VISp | session_798911424_dg8_VISp | 44 | accuracy | 0.277778 |
| classification | stabilization | Visual Coding | single_day | session_798911424_dg8_VISp | session_798911424_dg8_VISp | 44 | accuracy | 0.608333 |
| classification | stabilization | Visual Coding | single_day | session_798911424_dg8_VISp | session_798911424_dg8_VISp | 46 | accuracy | 0.37037 |
| classification | stabilization | Visual Coding | single_day | session_798911424_dg8_VISp | session_798911424_dg8_VISp | 46 | accuracy | 0.716667 |
| classification | stabilization | Visual Coding | single_day | session_798911424_dg8_VISp | session_798911424_dg8_VISp | 54 | accuracy | 0.314815 |
| classification | stabilization | Visual Coding | single_day | session_798911424_dg8_VISp | session_798911424_dg8_VISp | 54 | accuracy | 0.725 |
| classification | stabilization | Visual Coding | single_day | session_799864342_dg8_VISp | session_799864342_dg8_VISp | 44 | accuracy | 0.5 |
| classification | stabilization | Visual Coding | single_day | session_799864342_dg8_VISp | session_799864342_dg8_VISp | 44 | accuracy | 0.875 |
| classification | stabilization | Visual Coding | single_day | session_799864342_dg8_VISp | session_799864342_dg8_VISp | 46 | accuracy | 0.37037 |
| classification | stabilization | Visual Coding | single_day | session_799864342_dg8_VISp | session_799864342_dg8_VISp | 46 | accuracy | 0.933333 |
| classification | stabilization | Visual Coding | single_day | session_799864342_dg8_VISp | session_799864342_dg8_VISp | 54 | accuracy | 0.416667 |
| classification | stabilization | Visual Coding | single_day | session_799864342_dg8_VISp | session_799864342_dg8_VISp | 54 | accuracy | 0.858333 |
| classification | mfsnn | Visual Coding | single_day | session_715093703_dg8_VISp | session_715093703_dg8_VISp | 44 | accuracy | 0.516667 |
| classification | mfsnn | Visual Coding | single_day | session_715093703_dg8_VISp | session_715093703_dg8_VISp | 46 | accuracy | 0.4 |
| classification | mfsnn | Visual Coding | single_day | session_715093703_dg8_VISp | session_715093703_dg8_VISp | 54 | accuracy | 0.391667 |
| classification | mfsnn | Visual Coding | single_day | session_719161530_dg8_VISp | session_719161530_dg8_VISp | 44 | accuracy | 0.616667 |
| classification | mfsnn | Visual Coding | single_day | session_719161530_dg8_VISp | session_719161530_dg8_VISp | 46 | accuracy | 0.533333 |
| classification | mfsnn | Visual Coding | single_day | session_719161530_dg8_VISp | session_719161530_dg8_VISp | 54 | accuracy | 0.575 |
| classification | mfsnn | Visual Coding | single_day | session_721123822_dg8_VISp | session_721123822_dg8_VISp | 44 | accuracy | 0.525 |
| classification | mfsnn | Visual Coding | single_day | session_721123822_dg8_VISp | session_721123822_dg8_VISp | 46 | accuracy | 0.491667 |

| | | | | | | | | |
|---|---|---|---|---|---|---|---|---|
| classification | mfsnn | Visual Coding | single_day | session_721123822_dg8_VISp | session_721123822_dg8_VISp | 54 | accuracy | 0.558333 |
| classification | mfsnn | Visual Coding | single_day | session_732592105_dg8_VISp | session_732592105_dg8_VISp | 44 | accuracy | 0.441667 |
| classification | mfsnn | Visual Coding | single_day | session_732592105_dg8_VISp | session_732592105_dg8_VISp | 46 | accuracy | 0.366667 |
| classification | mfsnn | Visual Coding | single_day | session_732592105_dg8_VISp | session_732592105_dg8_VISp | 54 | accuracy | 0.441667 |
| classification | mfsnn | Visual Coding | single_day | session_737581020_dg8_VISp | session_737581020_dg8_VISp | 44 | accuracy | 0.558333 |
| classification | mfsnn | Visual Coding | single_day | session_737581020_dg8_VISp | session_737581020_dg8_VISp | 46 | accuracy | 0.466667 |
| classification | mfsnn | Visual Coding | single_day | session_737581020_dg8_VISp | session_737581020_dg8_VISp | 54 | accuracy | 0.65 |
| classification | mfsnn | Visual Coding | single_day | session_739448407_dg8_VISp | session_739448407_dg8_VISp | 44 | accuracy | 0.394958 |
| classification | mfsnn | Visual Coding | single_day | session_739448407_dg8_VISp | session_739448407_dg8_VISp | 46 | accuracy | 0.344538 |
| classification | mfsnn | Visual Coding | single_day | session_739448407_dg8_VISp | session_739448407_dg8_VISp | 54 | accuracy | 0.512605 |
| classification | mfsnn | Visual Coding | single_day | session_742951821_dg8_VISp | session_742951821_dg8_VISp | 44 | accuracy | 0.258333 |
| classification | mfsnn | Visual Coding | single_day | session_742951821_dg8_VISp | session_742951821_dg8_VISp | 46 | accuracy | 0.3 |
| classification | mfsnn | Visual Coding | single_day | session_742951821_dg8_VISp | session_742951821_dg8_VISp | 54 | accuracy | 0.375 |
| classification | mfsnn | Visual Coding | single_day | session_743475441_dg8_VISp | session_743475441_dg8_VISp | 44 | accuracy | 0.2 |
| classification | mfsnn | Visual Coding | single_day | session_743475441_dg8_VISp | session_743475441_dg8_VISp | 46 | accuracy | 0.2 |
| classification | mfsnn | Visual Coding | single_day | session_743475441_dg8_VISp | session_743475441_dg8_VISp | 54 | accuracy | 0.258333 |
| classification | mfsnn | Visual Coding | single_day | session_744228101_dg8_VISp | session_744228101_dg8_VISp | 44 | accuracy | 0.45 |
| classification | mfsnn | Visual Coding | single_day | session_744228101_dg8_VISp | session_744228101_dg8_VISp | 46 | accuracy | 0.55 |
| classification | mfsnn | Visual Coding | single_day | session_744228101_dg8_VISp | session_744228101_dg8_VISp | 54 | accuracy | 0.458333 |
| classification | mfsnn | Visual Coding | single_day | session_746083955_dg8_VISp | session_746083955_dg8_VISp | 44 | accuracy | 0.191667 |
| classification | mfsnn | Visual Coding | single_day | session_746083955_dg8_VISp | session_746083955_dg8_VISp | 46 | accuracy | 0.241667 |
| classification | mfsnn | Visual Coding | single_day | session_746083955_dg8_VISp | session_746083955_dg8_VISp | 54 | accuracy | 0.25 |
| classification | mfsnn | Visual Coding | single_day | session_750332458_dg8_VISp | session_750332458_dg8_VISp | 44 | accuracy | 0.575 |
| classification | mfsnn | Visual Coding | single_day | session_750332458_dg8_VISp | session_750332458_dg8_VISp | 46 | accuracy | 0.6 |
| classification | mfsnn | Visual Coding | single_day | session_750332458_dg8_VISp | session_750332458_dg8_VISp | 54 | accuracy | 0.591667 |
| classification | mfsnn | Visual Coding | single_day | session_750749662_dg8_VISp | session_750749662_dg8_VISp | 44 | accuracy | 0.158333 |
| classification | mfsnn | Visual Coding | single_day | session_750749662_dg8_VISp | session_750749662_dg8_VISp | 46 | accuracy | 0.625 |
| classification | mfsnn | Visual Coding | single_day | session_750749662_dg8_VISp | session_750749662_dg8_VISp | 54 | accuracy | 0.741667 |
| classification | mfsnn | Visual Coding | single_day | session_751348571_dg8_VISp | session_751348571_dg8_VISp | 44 | accuracy | 0.483333 |
| classification | mfsnn | Visual Coding | single_day | session_751348571_dg8_VISp | session_751348571_dg8_VISp | 46 | accuracy | 0.591667 |
| classification | mfsnn | Visual Coding | single_day | session_751348571_dg8_VISp | session_751348571_dg8_VISp | 54 | accuracy | 0.508333 |
| classification | mfsnn | Visual Coding | single_day | session_754312389_dg8_VISp | session_754312389_dg8_VISp | 44 | accuracy | 0.841667 |
| classification | mfsnn | Visual Coding | single_day | session_754312389_dg8_VISp | session_754312389_dg8_VISp | 46 | accuracy | 0.7 |
| classification | mfsnn | Visual Coding | single_day | session_754312389_dg8_VISp | session_754312389_dg8_VISp | 54 | accuracy | 0.733333 |
| classification | mfsnn | Visual Coding | single_day | session_754829445_dg8_VISp | session_754829445_dg8_VISp | 44 | accuracy | 0.683333 |
| classification | mfsnn | Visual Coding | single_day | session_754829445_dg8_VISp | session_754829445_dg8_VISp | 46 | accuracy | 0.616667 |
| classification | mfsnn | Visual Coding | single_day | session_754829445_dg8_VISp | session_754829445_dg8_VISp | 54 | accuracy | 0.391667 |
| classification | mfsnn | Visual Coding | single_day | session_755434585_dg8_VISp | session_755434585_dg8_VISp | 44 | accuracy | 0.433333 |
| classification | mfsnn | Visual Coding | single_day | session_755434585_dg8_VISp | session_755434585_dg8_VISp | 46 | accuracy | 0.575 |
| classification | mfsnn | Visual Coding | single_day | session_755434585_dg8_VISp | session_755434585_dg8_VISp | 54 | accuracy | 0.508333 |
| classification | mfsnn | Visual Coding | single_day | session_756029989_dg8_VISp | session_756029989_dg8_VISp | 44 | accuracy | 0.725 |
| classification | mfsnn | Visual Coding | single_day | session_756029989_dg8_VISp | session_756029989_dg8_VISp | 46 | accuracy | 0.758333 |
| classification | mfsnn | Visual Coding | single_day | session_756029989_dg8_VISp | session_756029989_dg8_VISp | 54 | accuracy | 0.608333 |
| classification | mfsnn | Visual Coding | single_day | session_757216464_dg8_VISp | session_757216464_dg8_VISp | 44 | accuracy | 0.508333 |
| classification | mfsnn | Visual Coding | single_day | session_757216464_dg8_VISp | session_757216464_dg8_VISp | 46 | accuracy | 0.3 |
| classification | mfsnn | Visual Coding | single_day | session_757216464_dg8_VISp | session_757216464_dg8_VISp | 54 | accuracy | 0.35 |
| classification | mfsnn | Visual Coding | single_day | session_757970808_dg8_VISp | session_757970808_dg8_VISp | 44 | accuracy | 0.483333 |
| classification | mfsnn | Visual Coding | single_day | session_757970808_dg8_VISp | session_757970808_dg8_VISp | 46 | accuracy | 0.583333 |
| classification | mfsnn | Visual Coding | single_day | session_757970808_dg8_VISp | session_757970808_dg8_VISp | 54 | accuracy | 0.608333 |
| classification | mfsnn | Visual Coding | single_day | session_758798717_dg8_VISp | session_758798717_dg8_VISp | 44 | accuracy | 0.533333 |
| classification | mfsnn | Visual Coding | single_day | session_758798717_dg8_VISp | session_758798717_dg8_VISp | 46 | accuracy | 0.416667 |
| classification | mfsnn | Visual Coding | single_day | session_758798717_dg8_VISp | session_758798717_dg8_VISp | 54 | accuracy | 0.483333 |
| classification | mfsnn | Visual Coding | single_day | session_759883607_dg8_VISp | session_759883607_dg8_VISp | 44 | accuracy | 0.483333 |
| classification | mfsnn | Visual Coding | single_day | session_759883607_dg8_VISp | session_759883607_dg8_VISp | 46 | accuracy | 0.516667 |
| classification | mfsnn | Visual Coding | single_day | session_759883607_dg8_VISp | session_759883607_dg8_VISp | 54 | accuracy | 0.391667 |
| classification | mfsnn | Visual Coding | single_day | session_760345702_dg8_VISp | session_760345702_dg8_VISp | 44 | accuracy | 0.65 |
| classification | mfsnn | Visual Coding | single_day | session_760345702_dg8_VISp | session_760345702_dg8_VISp | 46 | accuracy | 0.6 |
| classification | mfsnn | Visual Coding | single_day | session_760345702_dg8_VISp | session_760345702_dg8_VISp | 54 | accuracy | 0.758333 |
| classification | mfsnn | Visual Coding | single_day | session_760693773_dg8_VISp | session_760693773_dg8_VISp | 44 | accuracy | 0.683333 |
| classification | mfsnn | Visual Coding | single_day | session_760693773_dg8_VISp | session_760693773_dg8_VISp | 46 | accuracy | 0.475 |
| classification | mfsnn | Visual Coding | single_day | session_760693773_dg8_VISp | session_760693773_dg8_VISp | 54 | accuracy | 0.641667 |
| classification | mfsnn | Visual Coding | single_day | session_761418226_dg8_VISp | session_761418226_dg8_VISp | 44 | accuracy | 0.416667 |
| classification | mfsnn | Visual Coding | single_day | session_761418226_dg8_VISp | session_761418226_dg8_VISp | 46 | accuracy | 0.375 |

| | | | | | | | | |
|---|---|---|---|---|---|---|---|---|
| classification | mfsnn | Visual Coding | single_day | session_761418226_dg8_VISp | session_761418226_dg8_VISp | 54 | accuracy | 0.45 |
| classification | mfsnn | Visual Coding | single_day | session_762120172_dg8_VISp | session_762120172_dg8_VISp | 44 | accuracy | 0.308333 |
| classification | mfsnn | Visual Coding | single_day | session_762120172_dg8_VISp | session_762120172_dg8_VISp | 46 | accuracy | 0.358333 |
| classification | mfsnn | Visual Coding | single_day | session_762120172_dg8_VISp | session_762120172_dg8_VISp | 54 | accuracy | 0.541667 |
| classification | mfsnn | Visual Coding | single_day | session_762602078_dg8_VISp | session_762602078_dg8_VISp | 44 | accuracy | 0.883333 |
| classification | mfsnn | Visual Coding | single_day | session_762602078_dg8_VISp | session_762602078_dg8_VISp | 46 | accuracy | 0.766667 |
| classification | mfsnn | Visual Coding | single_day | session_762602078_dg8_VISp | session_762602078_dg8_VISp | 54 | accuracy | 0.833333 |
| classification | mfsnn | Visual Coding | single_day | session_763673393_dg8_VISp | session_763673393_dg8_VISp | 44 | accuracy | 0.558333 |
| classification | mfsnn | Visual Coding | single_day | session_763673393_dg8_VISp | session_763673393_dg8_VISp | 46 | accuracy | 0.566667 |
| classification | mfsnn | Visual Coding | single_day | session_763673393_dg8_VISp | session_763673393_dg8_VISp | 54 | accuracy | 0.55 |
| classification | mfsnn | Visual Coding | single_day | session_773418906_dg8_VISp | session_773418906_dg8_VISp | 44 | accuracy | 0.425 |
| classification | mfsnn | Visual Coding | single_day | session_773418906_dg8_VISp | session_773418906_dg8_VISp | 46 | accuracy | 0.4 |
| classification | mfsnn | Visual Coding | single_day | session_773418906_dg8_VISp | session_773418906_dg8_VISp | 54 | accuracy | 0.383333 |
| classification | mfsnn | Visual Coding | single_day | session_791319847_dg8_VISp | session_791319847_dg8_VISp | 44 | accuracy | 0.516667 |
| classification | mfsnn | Visual Coding | single_day | session_791319847_dg8_VISp | session_791319847_dg8_VISp | 46 | accuracy | 0.508333 |
| classification | mfsnn | Visual Coding | single_day | session_791319847_dg8_VISp | session_791319847_dg8_VISp | 54 | accuracy | 0.408333 |
| classification | mfsnn | Visual Coding | single_day | session_797828357_dg8_VISp | session_797828357_dg8_VISp | 44 | accuracy | 0.575 |
| classification | mfsnn | Visual Coding | single_day | session_797828357_dg8_VISp | session_797828357_dg8_VISp | 46 | accuracy | 0.591667 |
| classification | mfsnn | Visual Coding | single_day | session_797828357_dg8_VISp | session_797828357_dg8_VISp | 54 | accuracy | 0.491667 |
| classification | mfsnn | Visual Coding | single_day | session_798911424_dg8_VISp | session_798911424_dg8_VISp | 44 | accuracy | 0.533333 |
| classification | mfsnn | Visual Coding | single_day | session_798911424_dg8_VISp | session_798911424_dg8_VISp | 46 | accuracy | 0.491667 |
| classification | mfsnn | Visual Coding | single_day | session_798911424_dg8_VISp | session_798911424_dg8_VISp | 54 | accuracy | 0.558333 |
| classification | mfsnn | Visual Coding | single_day | session_799864342_dg8_VISp | session_799864342_dg8_VISp | 44 | accuracy | 0.741667 |
| classification | mfsnn | Visual Coding | single_day | session_799864342_dg8_VISp | session_799864342_dg8_VISp | 46 | accuracy | 0.708333 |
| classification | mfsnn | Visual Coding | single_day | session_799864342_dg8_VISp | session_799864342_dg8_VISp | 54 | accuracy | 0.658333 |
| classification | mscformer | Visual Coding | single_day | session_715093703_dg8_VISp | session_715093703_dg8_VISp | 44 | accuracy | 0.508333 |
| classification | mscformer | Visual Coding | single_day | session_715093703_dg8_VISp | session_715093703_dg8_VISp | 46 | accuracy | 0.366667 |
| classification | mscformer | Visual Coding | single_day | session_715093703_dg8_VISp | session_715093703_dg8_VISp | 54 | accuracy | 0.441667 |
| classification | mscformer | Visual Coding | single_day | session_719161530_dg8_VISp | session_719161530_dg8_VISp | 44 | accuracy | 0.516667 |
| classification | mscformer | Visual Coding | single_day | session_719161530_dg8_VISp | session_719161530_dg8_VISp | 46 | accuracy | 0.475 |
| classification | mscformer | Visual Coding | single_day | session_719161530_dg8_VISp | session_719161530_dg8_VISp | 54 | accuracy | 0.491667 |
| classification | mscformer | Visual Coding | single_day | session_721123822_dg8_VISp | session_721123822_dg8_VISp | 44 | accuracy | 0.175 |
| classification | mscformer | Visual Coding | single_day | session_721123822_dg8_VISp | session_721123822_dg8_VISp | 46 | accuracy | 0.35 |
| classification | mscformer | Visual Coding | single_day | session_721123822_dg8_VISp | session_721123822_dg8_VISp | 54 | accuracy | 0.383333 |
| classification | mscformer | Visual Coding | single_day | session_732592105_dg8_VISp | session_732592105_dg8_VISp | 44 | accuracy | 0.541667 |
| classification | mscformer | Visual Coding | single_day | session_732592105_dg8_VISp | session_732592105_dg8_VISp | 46 | accuracy | 0.491667 |
| classification | mscformer | Visual Coding | single_day | session_732592105_dg8_VISp | session_732592105_dg8_VISp | 54 | accuracy | 0.475 |
| classification | mscformer | Visual Coding | single_day | session_737581020_dg8_VISp | session_737581020_dg8_VISp | 44 | accuracy | 0.558333 |
| classification | mscformer | Visual Coding | single_day | session_737581020_dg8_VISp | session_737581020_dg8_VISp | 46 | accuracy | 0.458333 |
| classification | mscformer | Visual Coding | single_day | session_737581020_dg8_VISp | session_737581020_dg8_VISp | 54 | accuracy | 0.441667 |
| classification | mscformer | Visual Coding | single_day | session_739448407_dg8_VISp | session_739448407_dg8_VISp | 44 | accuracy | 0.310924 |
| classification | mscformer | Visual Coding | single_day | session_739448407_dg8_VISp | session_739448407_dg8_VISp | 46 | accuracy | 0.260504 |
| classification | mscformer | Visual Coding | single_day | session_739448407_dg8_VISp | session_739448407_dg8_VISp | 54 | accuracy | 0.294118 |
| classification | mscformer | Visual Coding | single_day | session_742951821_dg8_VISp | session_742951821_dg8_VISp | 44 | accuracy | 0.266667 |
| classification | mscformer | Visual Coding | single_day | session_742951821_dg8_VISp | session_742951821_dg8_VISp | 46 | accuracy | 0.258333 |
| classification | mscformer | Visual Coding | single_day | session_742951821_dg8_VISp | session_742951821_dg8_VISp | 54 | accuracy | 0.233333 |
| classification | mscformer | Visual Coding | single_day | session_743475441_dg8_VISp | session_743475441_dg8_VISp | 44 | accuracy | 0.166667 |
| classification | mscformer | Visual Coding | single_day | session_743475441_dg8_VISp | session_743475441_dg8_VISp | 46 | accuracy | 0.283333 |
| classification | mscformer | Visual Coding | single_day | session_743475441_dg8_VISp | session_743475441_dg8_VISp | 54 | accuracy | 0.108333 |
| classification | mscformer | Visual Coding | single_day | session_744228101_dg8_VISp | session_744228101_dg8_VISp | 44 | accuracy | 0.316667 |
| classification | mscformer | Visual Coding | single_day | session_744228101_dg8_VISp | session_744228101_dg8_VISp | 46 | accuracy | 0.408333 |
| classification | mscformer | Visual Coding | single_day | session_744228101_dg8_VISp | session_744228101_dg8_VISp | 54 | accuracy | 0.475 |
| classification | mscformer | Visual Coding | single_day | session_746083955_dg8_VISp | session_746083955_dg8_VISp | 44 | accuracy | 0.175 |
| classification | mscformer | Visual Coding | single_day | session_746083955_dg8_VISp | session_746083955_dg8_VISp | 46 | accuracy | 0.316667 |
| classification | mscformer | Visual Coding | single_day | session_746083955_dg8_VISp | session_746083955_dg8_VISp | 54 | accuracy | 0.191667 |
| classification | mscformer | Visual Coding | single_day | session_750332458_dg8_VISp | session_750332458_dg8_VISp | 44 | accuracy | 0.475 |
| classification | mscformer | Visual Coding | single_day | session_750332458_dg8_VISp | session_750332458_dg8_VISp | 46 | accuracy | 0.525 |
| classification | mscformer | Visual Coding | single_day | session_750332458_dg8_VISp | session_750332458_dg8_VISp | 54 | accuracy | 0.425 |
| classification | mscformer | Visual Coding | single_day | session_750749662_dg8_VISp | session_750749662_dg8_VISp | 44 | accuracy | 0.4 |
| classification | mscformer | Visual Coding | single_day | session_750749662_dg8_VISp | session_750749662_dg8_VISp | 46 | accuracy | 0.508333 |
| classification | mscformer | Visual Coding | single_day | session_750749662_dg8_VISp | session_750749662_dg8_VISp | 54 | accuracy | 0.416667 |
| classification | mscformer | Visual Coding | single_day | session_751348571_dg8_VISp | session_751348571_dg8_VISp | 44 | accuracy | 0.591667 |
| classification | mscformer | Visual Coding | single_day | session_751348571_dg8_VISp | session_751348571_dg8_VISp | 46 | accuracy | 0.616667 |

| | | | | | | | | |
|---|---|---|---|---|---|---|---|---|
| classification | mscformer | Visual Coding | single_day | session_751348571_dg8_VISp | session_751348571_dg8_VISp | 54 | accuracy | 0.541667 |
| classification | mscformer | Visual Coding | single_day | session_754312389_dg8_VISp | session_754312389_dg8_VISp | 44 | accuracy | 0.566667 |
| classification | mscformer | Visual Coding | single_day | session_754312389_dg8_VISp | session_754312389_dg8_VISp | 46 | accuracy | 0.633333 |
| classification | mscformer | Visual Coding | single_day | session_754312389_dg8_VISp | session_754312389_dg8_VISp | 54 | accuracy | 0.666667 |
| classification | mscformer | Visual Coding | single_day | session_754829445_dg8_VISp | session_754829445_dg8_VISp | 44 | accuracy | 0.416667 |
| classification | mscformer | Visual Coding | single_day | session_754829445_dg8_VISp | session_754829445_dg8_VISp | 46 | accuracy | 0.408333 |
| classification | mscformer | Visual Coding | single_day | session_754829445_dg8_VISp | session_754829445_dg8_VISp | 54 | accuracy | 0.425 |
| classification | mscformer | Visual Coding | single_day | session_755434585_dg8_VISp | session_755434585_dg8_VISp | 44 | accuracy | 0.375 |
| classification | mscformer | Visual Coding | single_day | session_755434585_dg8_VISp | session_755434585_dg8_VISp | 46 | accuracy | 0.408333 |
| classification | mscformer | Visual Coding | single_day | session_755434585_dg8_VISp | session_755434585_dg8_VISp | 54 | accuracy | 0.4 |
| classification | mscformer | Visual Coding | single_day | session_756029989_dg8_VISp | session_756029989_dg8_VISp | 44 | accuracy | 0.458333 |
| classification | mscformer | Visual Coding | single_day | session_756029989_dg8_VISp | session_756029989_dg8_VISp | 46 | accuracy | 0.508333 |
| classification | mscformer | Visual Coding | single_day | session_756029989_dg8_VISp | session_756029989_dg8_VISp | 54 | accuracy | 0.533333 |
| classification | mscformer | Visual Coding | single_day | session_757216464_dg8_VISp | session_757216464_dg8_VISp | 44 | accuracy | 0.45 |
| classification | mscformer | Visual Coding | single_day | session_757216464_dg8_VISp | session_757216464_dg8_VISp | 46 | accuracy | 0.308333 |
| classification | mscformer | Visual Coding | single_day | session_757216464_dg8_VISp | session_757216464_dg8_VISp | 54 | accuracy | 0.5 |
| classification | mscformer | Visual Coding | single_day | session_757970808_dg8_VISp | session_757970808_dg8_VISp | 44 | accuracy | 0.525 |
| classification | mscformer | Visual Coding | single_day | session_757970808_dg8_VISp | session_757970808_dg8_VISp | 46 | accuracy | 0.35 |
| classification | mscformer | Visual Coding | single_day | session_757970808_dg8_VISp | session_757970808_dg8_VISp | 54 | accuracy | 0.533333 |
| classification | mscformer | Visual Coding | single_day | session_758798717_dg8_VISp | session_758798717_dg8_VISp | 44 | accuracy | 0.325 |
| classification | mscformer | Visual Coding | single_day | session_758798717_dg8_VISp | session_758798717_dg8_VISp | 46 | accuracy | 0.383333 |
| classification | mscformer | Visual Coding | single_day | session_758798717_dg8_VISp | session_758798717_dg8_VISp | 54 | accuracy | 0.383333 |
| classification | mscformer | Visual Coding | single_day | session_759883607_dg8_VISp | session_759883607_dg8_VISp | 44 | accuracy | 0.466667 |
| classification | mscformer | Visual Coding | single_day | session_759883607_dg8_VISp | session_759883607_dg8_VISp | 46 | accuracy | 0.425 |
| classification | mscformer | Visual Coding | single_day | session_759883607_dg8_VISp | session_759883607_dg8_VISp | 54 | accuracy | 0.433333 |
| classification | mscformer | Visual Coding | single_day | session_760345702_dg8_VISp | session_760345702_dg8_VISp | 44 | accuracy | 0.575 |
| classification | mscformer | Visual Coding | single_day | session_760345702_dg8_VISp | session_760345702_dg8_VISp | 46 | accuracy | 0.541667 |
| classification | mscformer | Visual Coding | single_day | session_760345702_dg8_VISp | session_760345702_dg8_VISp | 54 | accuracy | 0.566667 |
| classification | mscformer | Visual Coding | single_day | session_760693773_dg8_VISp | session_760693773_dg8_VISp | 44 | accuracy | 0.45 |
| classification | mscformer | Visual Coding | single_day | session_760693773_dg8_VISp | session_760693773_dg8_VISp | 46 | accuracy | 0.675 |
| classification | mscformer | Visual Coding | single_day | session_760693773_dg8_VISp | session_760693773_dg8_VISp | 54 | accuracy | 0.708333 |
| classification | mscformer | Visual Coding | single_day | session_761418226_dg8_VISp | session_761418226_dg8_VISp | 44 | accuracy | 0.45 |
| classification | mscformer | Visual Coding | single_day | session_761418226_dg8_VISp | session_761418226_dg8_VISp | 46 | accuracy | 0.408333 |
| classification | mscformer | Visual Coding | single_day | session_761418226_dg8_VISp | session_761418226_dg8_VISp | 54 | accuracy | 0.433333 |
| classification | mscformer | Visual Coding | single_day | session_762120172_dg8_VISp | session_762120172_dg8_VISp | 44 | accuracy | 0.491667 |
| classification | mscformer | Visual Coding | single_day | session_762120172_dg8_VISp | session_762120172_dg8_VISp | 46 | accuracy | 0.491667 |
| classification | mscformer | Visual Coding | single_day | session_762120172_dg8_VISp | session_762120172_dg8_VISp | 54 | accuracy | 0.508333 |
| classification | mscformer | Visual Coding | single_day | session_762602078_dg8_VISp | session_762602078_dg8_VISp | 44 | accuracy | 0.541667 |
| classification | mscformer | Visual Coding | single_day | session_762602078_dg8_VISp | session_762602078_dg8_VISp | 46 | accuracy | 0.575 |
| classification | mscformer | Visual Coding | single_day | session_762602078_dg8_VISp | session_762602078_dg8_VISp | 54 | accuracy | 0.633333 |
| classification | mscformer | Visual Coding | single_day | session_763673393_dg8_VISp | session_763673393_dg8_VISp | 44 | accuracy | 0.375 |
| classification | mscformer | Visual Coding | single_day | session_763673393_dg8_VISp | session_763673393_dg8_VISp | 46 | accuracy | 0.441667 |
| classification | mscformer | Visual Coding | single_day | session_763673393_dg8_VISp | session_763673393_dg8_VISp | 54 | accuracy | 0.525 |
| classification | mscformer | Visual Coding | single_day | session_773418906_dg8_VISp | session_773418906_dg8_VISp | 44 | accuracy | 0.258333 |
| classification | mscformer | Visual Coding | single_day | session_773418906_dg8_VISp | session_773418906_dg8_VISp | 46 | accuracy | 0.291667 |
| classification | mscformer | Visual Coding | single_day | session_773418906_dg8_VISp | session_773418906_dg8_VISp | 54 | accuracy | 0.358333 |
| classification | mscformer | Visual Coding | single_day | session_791319847_dg8_VISp | session_791319847_dg8_VISp | 44 | accuracy | 0.4 |
| classification | mscformer | Visual Coding | single_day | session_791319847_dg8_VISp | session_791319847_dg8_VISp | 46 | accuracy | 0.466667 |
| classification | mscformer | Visual Coding | single_day | session_791319847_dg8_VISp | session_791319847_dg8_VISp | 54 | accuracy | 0.416667 |
| classification | mscformer | Visual Coding | single_day | session_797828357_dg8_VISp | session_797828357_dg8_VISp | 44 | accuracy | 0.416667 |
| classification | mscformer | Visual Coding | single_day | session_797828357_dg8_VISp | session_797828357_dg8_VISp | 46 | accuracy | 0.458333 |
| classification | mscformer | Visual Coding | single_day | session_797828357_dg8_VISp | session_797828357_dg8_VISp | 54 | accuracy | 0.458333 |
| classification | mscformer | Visual Coding | single_day | session_798911424_dg8_VISp | session_798911424_dg8_VISp | 44 | accuracy | 0.541667 |
| classification | mscformer | Visual Coding | single_day | session_798911424_dg8_VISp | session_798911424_dg8_VISp | 46 | accuracy | 0.508333 |
| classification | mscformer | Visual Coding | single_day | session_798911424_dg8_VISp | session_798911424_dg8_VISp | 54 | accuracy | 0.466667 |
| classification | mscformer | Visual Coding | single_day | session_799864342_dg8_VISp | session_799864342_dg8_VISp | 44 | accuracy | 0.625 |
| classification | mscformer | Visual Coding | single_day | session_799864342_dg8_VISp | session_799864342_dg8_VISp | 46 | accuracy | 0.683333 |
| classification | mscformer | Visual Coding | single_day | session_799864342_dg8_VISp | session_799864342_dg8_VISp | 54 | accuracy | 0.658333 |
| classification | nomad | Visual Coding | single_day | session_715093703_dg8_VISp | session_715093703_dg8_VISp | 44 | accuracy | 0.65 |
| classification | nomad | Visual Coding | single_day | session_715093703_dg8_VISp | session_715093703_dg8_VISp | 46 | accuracy | 0.658333 |
| classification | nomad | Visual Coding | single_day | session_715093703_dg8_VISp | session_715093703_dg8_VISp | 54 | accuracy | 0.608333 |
| classification | nomad | Visual Coding | single_day | session_719161530_dg8_VISp | session_719161530_dg8_VISp | 44 | accuracy | 0.775 |
| classification | nomad | Visual Coding | single_day | session_719161530_dg8_VISp | session_719161530_dg8_VISp | 46 | accuracy | 0.808333 |

| | | | | | | | | |
|---|---|---|---|---|---|---|---|---|
| classification | nomad | Visual Coding | single_day | session_719161530_dg8_VISp | session_719161530_dg8_VISp | 54 | accuracy | 0.8 |
| classification | nomad | Visual Coding | single_day | session_721123822_dg8_VISp | session_721123822_dg8_VISp | 44 | accuracy | 0.675 |
| classification | nomad | Visual Coding | single_day | session_721123822_dg8_VISp | session_721123822_dg8_VISp | 46 | accuracy | 0.583333 |
| classification | nomad | Visual Coding | single_day | session_721123822_dg8_VISp | session_721123822_dg8_VISp | 54 | accuracy | 0.65 |
| classification | nomad | Visual Coding | single_day | session_732592105_dg8_VISp | session_732592105_dg8_VISp | 44 | accuracy | 0.666667 |
| classification | nomad | Visual Coding | single_day | session_732592105_dg8_VISp | session_732592105_dg8_VISp | 46 | accuracy | 0.791667 |
| classification | nomad | Visual Coding | single_day | session_732592105_dg8_VISp | session_732592105_dg8_VISp | 54 | accuracy | 0.841667 |
| classification | nomad | Visual Coding | single_day | session_737581020_dg8_VISp | session_737581020_dg8_VISp | 44 | accuracy | 0.6 |
| classification | nomad | Visual Coding | single_day | session_737581020_dg8_VISp | session_737581020_dg8_VISp | 46 | accuracy | 0.575 |
| classification | nomad | Visual Coding | single_day | session_737581020_dg8_VISp | session_737581020_dg8_VISp | 54 | accuracy | 0.733333 |
| classification | nomad | Visual Coding | single_day | session_739448407_dg8_VISp | session_739448407_dg8_VISp | 44 | accuracy | 0.478992 |
| classification | nomad | Visual Coding | single_day | session_739448407_dg8_VISp | session_739448407_dg8_VISp | 46 | accuracy | 0.453782 |
| classification | nomad | Visual Coding | single_day | session_739448407_dg8_VISp | session_739448407_dg8_VISp | 54 | accuracy | 0.369748 |
| classification | nomad | Visual Coding | single_day | session_742951821_dg8_VISp | session_742951821_dg8_VISp | 44 | accuracy | 0.475 |
| classification | nomad | Visual Coding | single_day | session_742951821_dg8_VISp | session_742951821_dg8_VISp | 46 | accuracy | 0.441667 |
| classification | nomad | Visual Coding | single_day | session_742951821_dg8_VISp | session_742951821_dg8_VISp | 54 | accuracy | 0.591667 |
| classification | nomad | Visual Coding | single_day | session_743475441_dg8_VISp | session_743475441_dg8_VISp | 44 | accuracy | 0.3 |
| classification | nomad | Visual Coding | single_day | session_743475441_dg8_VISp | session_743475441_dg8_VISp | 46 | accuracy | 0.441667 |
| classification | nomad | Visual Coding | single_day | session_743475441_dg8_VISp | session_743475441_dg8_VISp | 54 | accuracy | 0.325 |
| classification | nomad | Visual Coding | single_day | session_744228101_dg8_VISp | session_744228101_dg8_VISp | 44 | accuracy | 0.75 |
| classification | nomad | Visual Coding | single_day | session_744228101_dg8_VISp | session_744228101_dg8_VISp | 46 | accuracy | 0.791667 |
| classification | nomad | Visual Coding | single_day | session_744228101_dg8_VISp | session_744228101_dg8_VISp | 54 | accuracy | 0.775 |
| classification | nomad | Visual Coding | single_day | session_746083955_dg8_VISp | session_746083955_dg8_VISp | 44 | accuracy | 0.441667 |
| classification | nomad | Visual Coding | single_day | session_746083955_dg8_VISp | session_746083955_dg8_VISp | 46 | accuracy | 0.491667 |
| classification | nomad | Visual Coding | single_day | session_746083955_dg8_VISp | session_746083955_dg8_VISp | 54 | accuracy | 0.4 |
| classification | nomad | Visual Coding | single_day | session_750332458_dg8_VISp | session_750332458_dg8_VISp | 44 | accuracy | 0.866667 |
| classification | nomad | Visual Coding | single_day | session_750332458_dg8_VISp | session_750332458_dg8_VISp | 46 | accuracy | 0.833333 |
| classification | nomad | Visual Coding | single_day | session_750332458_dg8_VISp | session_750332458_dg8_VISp | 54 | accuracy | 0.816667 |
| classification | nomad | Visual Coding | single_day | session_750749662_dg8_VISp | session_750749662_dg8_VISp | 44 | accuracy | 0.675 |
| classification | nomad | Visual Coding | single_day | session_750749662_dg8_VISp | session_750749662_dg8_VISp | 46 | accuracy | 0.766667 |
| classification | nomad | Visual Coding | single_day | session_750749662_dg8_VISp | session_750749662_dg8_VISp | 54 | accuracy | 0.8 |
| classification | nomad | Visual Coding | single_day | session_751348571_dg8_VISp | session_751348571_dg8_VISp | 44 | accuracy | 0.758333 |
| classification | nomad | Visual Coding | single_day | session_751348571_dg8_VISp | session_751348571_dg8_VISp | 46 | accuracy | 0.816667 |
| classification | nomad | Visual Coding | single_day | session_751348571_dg8_VISp | session_751348571_dg8_VISp | 54 | accuracy | 0.783333 |
| classification | nomad | Visual Coding | single_day | session_754312389_dg8_VISp | session_754312389_dg8_VISp | 44 | accuracy | 0.95 |
| classification | nomad | Visual Coding | single_day | session_754312389_dg8_VISp | session_754312389_dg8_VISp | 46 | accuracy | 0.9 |
| classification | nomad | Visual Coding | single_day | session_754312389_dg8_VISp | session_754312389_dg8_VISp | 54 | accuracy | 0.916667 |
| classification | nomad | Visual Coding | single_day | session_754829445_dg8_VISp | session_754829445_dg8_VISp | 44 | accuracy | 0.766667 |
| classification | nomad | Visual Coding | single_day | session_754829445_dg8_VISp | session_754829445_dg8_VISp | 46 | accuracy | 0.8 |
| classification | nomad | Visual Coding | single_day | session_754829445_dg8_VISp | session_754829445_dg8_VISp | 54 | accuracy | 0.758333 |
| classification | nomad | Visual Coding | single_day | session_755434585_dg8_VISp | session_755434585_dg8_VISp | 44 | accuracy | 0.725 |
| classification | nomad | Visual Coding | single_day | session_755434585_dg8_VISp | session_755434585_dg8_VISp | 46 | accuracy | 0.716667 |
| classification | nomad | Visual Coding | single_day | session_755434585_dg8_VISp | session_755434585_dg8_VISp | 54 | accuracy | 0.716667 |
| classification | nomad | Visual Coding | single_day | session_756029989_dg8_VISp | session_756029989_dg8_VISp | 44 | accuracy | 0.75 |
| classification | nomad | Visual Coding | single_day | session_756029989_dg8_VISp | session_756029989_dg8_VISp | 46 | accuracy | 0.841667 |
| classification | nomad | Visual Coding | single_day | session_756029989_dg8_VISp | session_756029989_dg8_VISp | 54 | accuracy | 0.85 |
| classification | nomad | Visual Coding | single_day | session_757216464_dg8_VISp | session_757216464_dg8_VISp | 44 | accuracy | 0.758333 |
| classification | nomad | Visual Coding | single_day | session_757216464_dg8_VISp | session_757216464_dg8_VISp | 46 | accuracy | 0.708333 |
| classification | nomad | Visual Coding | single_day | session_757216464_dg8_VISp | session_757216464_dg8_VISp | 54 | accuracy | 0.758333 |
| classification | nomad | Visual Coding | single_day | session_757970808_dg8_VISp | session_757970808_dg8_VISp | 44 | accuracy | 0.7 |
| classification | nomad | Visual Coding | single_day | session_757970808_dg8_VISp | session_757970808_dg8_VISp | 46 | accuracy | 0.841667 |
| classification | nomad | Visual Coding | single_day | session_757970808_dg8_VISp | session_757970808_dg8_VISp | 54 | accuracy | 0.825 |
| classification | nomad | Visual Coding | single_day | session_758798717_dg8_VISp | session_758798717_dg8_VISp | 44 | accuracy | 0.641667 |
| classification | nomad | Visual Coding | single_day | session_758798717_dg8_VISp | session_758798717_dg8_VISp | 46 | accuracy | 0.65 |
| classification | nomad | Visual Coding | single_day | session_758798717_dg8_VISp | session_758798717_dg8_VISp | 54 | accuracy | 0.591667 |
| classification | nomad | Visual Coding | single_day | session_759883607_dg8_VISp | session_759883607_dg8_VISp | 44 | accuracy | 0.758333 |
| classification | nomad | Visual Coding | single_day | session_759883607_dg8_VISp | session_759883607_dg8_VISp | 46 | accuracy | 0.75 |
| classification | nomad | Visual Coding | single_day | session_759883607_dg8_VISp | session_759883607_dg8_VISp | 54 | accuracy | 0.725 |
| classification | nomad | Visual Coding | single_day | session_760345702_dg8_VISp | session_760345702_dg8_VISp | 44 | accuracy | 0.891667 |
| classification | nomad | Visual Coding | single_day | session_760345702_dg8_VISp | session_760345702_dg8_VISp | 46 | accuracy | 0.891667 |
| classification | nomad | Visual Coding | single_day | session_760345702_dg8_VISp | session_760345702_dg8_VISp | 54 | accuracy | 0.875 |
| classification | nomad | Visual Coding | single_day | session_760693773_dg8_VISp | session_760693773_dg8_VISp | 44 | accuracy | 0.875 |
| classification | nomad | Visual Coding | single_day | session_760693773_dg8_VISp | session_760693773_dg8_VISp | 46 | accuracy | 0.833333 |

| | | | | | | | | |
|---|---|---|---|---|---|---|---|---|
| classification | nomad | Visual Coding | single_day | session_760693773_dg8_VISp | session_760693773_dg8_VISp | 54 | accuracy | 0.841667 |
| classification | nomad | Visual Coding | single_day | session_761418226_dg8_VISp | session_761418226_dg8_VISp | 44 | accuracy | 0.733333 |
| classification | nomad | Visual Coding | single_day | session_761418226_dg8_VISp | session_761418226_dg8_VISp | 46 | accuracy | 0.741667 |
| classification | nomad | Visual Coding | single_day | session_761418226_dg8_VISp | session_761418226_dg8_VISp | 54 | accuracy | 0.725 |
| classification | nomad | Visual Coding | single_day | session_762120172_dg8_VISp | session_762120172_dg8_VISp | 44 | accuracy | 0.766667 |
| classification | nomad | Visual Coding | single_day | session_762120172_dg8_VISp | session_762120172_dg8_VISp | 46 | accuracy | 0.891667 |
| classification | nomad | Visual Coding | single_day | session_762120172_dg8_VISp | session_762120172_dg8_VISp | 54 | accuracy | 0.8 |
| classification | nomad | Visual Coding | single_day | session_762602078_dg8_VISp | session_762602078_dg8_VISp | 44 | accuracy | 0.908333 |
| classification | nomad | Visual Coding | single_day | session_762602078_dg8_VISp | session_762602078_dg8_VISp | 46 | accuracy | 0.883333 |
| classification | nomad | Visual Coding | single_day | session_762602078_dg8_VISp | session_762602078_dg8_VISp | 54 | accuracy | 0.766667 |
| classification | nomad | Visual Coding | single_day | session_763673393_dg8_VISp | session_763673393_dg8_VISp | 44 | accuracy | 0.716667 |
| classification | nomad | Visual Coding | single_day | session_763673393_dg8_VISp | session_763673393_dg8_VISp | 46 | accuracy | 0.683333 |
| classification | nomad | Visual Coding | single_day | session_763673393_dg8_VISp | session_763673393_dg8_VISp | 54 | accuracy | 0.741667 |
| classification | nomad | Visual Coding | single_day | session_773418906_dg8_VISp | session_773418906_dg8_VISp | 44 | accuracy | 0.641667 |
| classification | nomad | Visual Coding | single_day | session_773418906_dg8_VISp | session_773418906_dg8_VISp | 46 | accuracy | 0.6 |
| classification | nomad | Visual Coding | single_day | session_773418906_dg8_VISp | session_773418906_dg8_VISp | 54 | accuracy | 0.608333 |
| classification | nomad | Visual Coding | single_day | session_791319847_dg8_VISp | session_791319847_dg8_VISp | 44 | accuracy | 0.916667 |
| classification | nomad | Visual Coding | single_day | session_791319847_dg8_VISp | session_791319847_dg8_VISp | 46 | accuracy | 0.925 |
| classification | nomad | Visual Coding | single_day | session_791319847_dg8_VISp | session_791319847_dg8_VISp | 54 | accuracy | 0.866667 |
| classification | nomad | Visual Coding | single_day | session_797828357_dg8_VISp | session_797828357_dg8_VISp | 44 | accuracy | 0.775 |
| classification | nomad | Visual Coding | single_day | session_797828357_dg8_VISp | session_797828357_dg8_VISp | 46 | accuracy | 0.75 |
| classification | nomad | Visual Coding | single_day | session_797828357_dg8_VISp | session_797828357_dg8_VISp | 54 | accuracy | 0.783333 |
| classification | nomad | Visual Coding | single_day | session_798911424_dg8_VISp | session_798911424_dg8_VISp | 44 | accuracy | 0.808333 |
| classification | nomad | Visual Coding | single_day | session_798911424_dg8_VISp | session_798911424_dg8_VISp | 46 | accuracy | 0.841667 |
| classification | nomad | Visual Coding | single_day | session_798911424_dg8_VISp | session_798911424_dg8_VISp | 54 | accuracy | 0.866667 |
| classification | nomad | Visual Coding | single_day | session_799864342_dg8_VISp | session_799864342_dg8_VISp | 44 | accuracy | 0.883333 |
| classification | nomad | Visual Coding | single_day | session_799864342_dg8_VISp | session_799864342_dg8_VISp | 46 | accuracy | 0.908333 |
| classification | nomad | Visual Coding | single_day | session_799864342_dg8_VISp | session_799864342_dg8_VISp | 54 | accuracy | 0.875 |
| classification | mlp | Visual Coding | single_day | session_715093703_dg8_VISp | session_715093703_dg8_VISp | 44 | accuracy | 0.7 |
| classification | mlp | Visual Coding | single_day | session_715093703_dg8_VISp | session_715093703_dg8_VISp | 46 | accuracy | 0.625 |
| classification | mlp | Visual Coding | single_day | session_715093703_dg8_VISp | session_715093703_dg8_VISp | 54 | accuracy | 0.683333 |
| classification | mlp | Visual Coding | single_day | session_719161530_dg8_VISp | session_719161530_dg8_VISp | 44 | accuracy | 0.766667 |
| classification | mlp | Visual Coding | single_day | session_719161530_dg8_VISp | session_719161530_dg8_VISp | 46 | accuracy | 0.725 |
| classification | mlp | Visual Coding | single_day | session_719161530_dg8_VISp | session_719161530_dg8_VISp | 54 | accuracy | 0.775 |
| classification | mlp | Visual Coding | single_day | session_721123822_dg8_VISp | session_721123822_dg8_VISp | 44 | accuracy | 0.725 |
| classification | mlp | Visual Coding | single_day | session_721123822_dg8_VISp | session_721123822_dg8_VISp | 46 | accuracy | 0.733333 |
| classification | mlp | Visual Coding | single_day | session_721123822_dg8_VISp | session_721123822_dg8_VISp | 54 | accuracy | 0.683333 |
| classification | mlp | Visual Coding | single_day | session_732592105_dg8_VISp | session_732592105_dg8_VISp | 44 | accuracy | 0.75 |
| classification | mlp | Visual Coding | single_day | session_732592105_dg8_VISp | session_732592105_dg8_VISp | 46 | accuracy | 0.758333 |
| classification | mlp | Visual Coding | single_day | session_732592105_dg8_VISp | session_732592105_dg8_VISp | 54 | accuracy | 0.658333 |
| classification | mlp | Visual Coding | single_day | session_737581020_dg8_VISp | session_737581020_dg8_VISp | 44 | accuracy | 0.758333 |
| classification | mlp | Visual Coding | single_day | session_737581020_dg8_VISp | session_737581020_dg8_VISp | 46 | accuracy | 0.791667 |
| classification | mlp | Visual Coding | single_day | session_737581020_dg8_VISp | session_737581020_dg8_VISp | 54 | accuracy | 0.808333 |
| classification | mlp | Visual Coding | single_day | session_739448407_dg8_VISp | session_739448407_dg8_VISp | 44 | accuracy | 0.546218 |
| classification | mlp | Visual Coding | single_day | session_739448407_dg8_VISp | session_739448407_dg8_VISp | 46 | accuracy | 0.571429 |
| classification | mlp | Visual Coding | single_day | session_739448407_dg8_VISp | session_739448407_dg8_VISp | 54 | accuracy | 0.436975 |
| classification | mlp | Visual Coding | single_day | session_742951821_dg8_VISp | session_742951821_dg8_VISp | 44 | accuracy | 0.441667 |
| classification | mlp | Visual Coding | single_day | session_742951821_dg8_VISp | session_742951821_dg8_VISp | 46 | accuracy | 0.5 |
| classification | mlp | Visual Coding | single_day | session_742951821_dg8_VISp | session_742951821_dg8_VISp | 54 | accuracy | 0.658333 |
| classification | mlp | Visual Coding | single_day | session_743475441_dg8_VISp | session_743475441_dg8_VISp | 44 | accuracy | 0.308333 |
| classification | mlp | Visual Coding | single_day | session_743475441_dg8_VISp | session_743475441_dg8_VISp | 46 | accuracy | 0.358333 |
| classification | mlp | Visual Coding | single_day | session_743475441_dg8_VISp | session_743475441_dg8_VISp | 54 | accuracy | 0.341667 |
| classification | mlp | Visual Coding | single_day | session_744228101_dg8_VISp | session_744228101_dg8_VISp | 44 | accuracy | 0.775 |
| classification | mlp | Visual Coding | single_day | session_744228101_dg8_VISp | session_744228101_dg8_VISp | 46 | accuracy | 0.775 |
| classification | mlp | Visual Coding | single_day | session_744228101_dg8_VISp | session_744228101_dg8_VISp | 54 | accuracy | 0.683333 |
| classification | mlp | Visual Coding | single_day | session_746083955_dg8_VISp | session_746083955_dg8_VISp | 44 | accuracy | 0.458333 |
| classification | mlp | Visual Coding | single_day | session_746083955_dg8_VISp | session_746083955_dg8_VISp | 46 | accuracy | 0.533333 |
| classification | mlp | Visual Coding | single_day | session_746083955_dg8_VISp | session_746083955_dg8_VISp | 54 | accuracy | 0.441667 |
| classification | mlp | Visual Coding | single_day | session_750332458_dg8_VISp | session_750332458_dg8_VISp | 44 | accuracy | 0.816667 |
| classification | mlp | Visual Coding | single_day | session_750332458_dg8_VISp | session_750332458_dg8_VISp | 46 | accuracy | 0.791667 |
| classification | mlp | Visual Coding | single_day | session_750332458_dg8_VISp | session_750332458_dg8_VISp | 54 | accuracy | 0.825 |
| classification | mlp | Visual Coding | single_day | session_750749662_dg8_VISp | session_750749662_dg8_VISp | 44 | accuracy | 0.766667 |
| classification | mlp | Visual Coding | single_day | session_750749662_dg8_VISp | session_750749662_dg8_VISp | 46 | accuracy | 0.7 |

| | | | | | | | | |
|---|---|---|---|---|---|---|---|---|
| classification | mlp | Visual Coding | single_day | session_750749662_dg8_VISp | session_750749662_dg8_VISp | 54 | accuracy | 0.716667 |
| classification | mlp | Visual Coding | single_day | session_751348571_dg8_VISp | session_751348571_dg8_VISp | 44 | accuracy | 0.908333 |
| classification | mlp | Visual Coding | single_day | session_751348571_dg8_VISp | session_751348571_dg8_VISp | 46 | accuracy | 0.833333 |
| classification | mlp | Visual Coding | single_day | session_751348571_dg8_VISp | session_751348571_dg8_VISp | 54 | accuracy | 0.825 |
| classification | mlp | Visual Coding | single_day | session_754312389_dg8_VISp | session_754312389_dg8_VISp | 44 | accuracy | 0.9 |
| classification | mlp | Visual Coding | single_day | session_754312389_dg8_VISp | session_754312389_dg8_VISp | 46 | accuracy | 0.866667 |
| classification | mlp | Visual Coding | single_day | session_754312389_dg8_VISp | session_754312389_dg8_VISp | 54 | accuracy | 0.833333 |
| classification | mlp | Visual Coding | single_day | session_754829445_dg8_VISp | session_754829445_dg8_VISp | 44 | accuracy | 0.766667 |
| classification | mlp | Visual Coding | single_day | session_754829445_dg8_VISp | session_754829445_dg8_VISp | 46 | accuracy | 0.708333 |
| classification | mlp | Visual Coding | single_day | session_754829445_dg8_VISp | session_754829445_dg8_VISp | 54 | accuracy | 0.8 |
| classification | mlp | Visual Coding | single_day | session_755434585_dg8_VISp | session_755434585_dg8_VISp | 44 | accuracy | 0.8 |
| classification | mlp | Visual Coding | single_day | session_755434585_dg8_VISp | session_755434585_dg8_VISp | 46 | accuracy | 0.65 |
| classification | mlp | Visual Coding | single_day | session_755434585_dg8_VISp | session_755434585_dg8_VISp | 54 | accuracy | 0.716667 |
| classification | mlp | Visual Coding | single_day | session_756029989_dg8_VISp | session_756029989_dg8_VISp | 44 | accuracy | 0.85 |
| classification | mlp | Visual Coding | single_day | session_756029989_dg8_VISp | session_756029989_dg8_VISp | 46 | accuracy | 0.8 |
| classification | mlp | Visual Coding | single_day | session_756029989_dg8_VISp | session_756029989_dg8_VISp | 54 | accuracy | 0.808333 |
| classification | mlp | Visual Coding | single_day | session_757216464_dg8_VISp | session_757216464_dg8_VISp | 44 | accuracy | 0.725 |
| classification | mlp | Visual Coding | single_day | session_757216464_dg8_VISp | session_757216464_dg8_VISp | 46 | accuracy | 0.741667 |
| classification | mlp | Visual Coding | single_day | session_757216464_dg8_VISp | session_757216464_dg8_VISp | 54 | accuracy | 0.691667 |
| classification | mlp | Visual Coding | single_day | session_757970808_dg8_VISp | session_757970808_dg8_VISp | 44 | accuracy | 0.733333 |
| classification | mlp | Visual Coding | single_day | session_757970808_dg8_VISp | session_757970808_dg8_VISp | 46 | accuracy | 0.808333 |
| classification | mlp | Visual Coding | single_day | session_757970808_dg8_VISp | session_757970808_dg8_VISp | 54 | accuracy | 0.775 |
| classification | mlp | Visual Coding | single_day | session_758798717_dg8_VISp | session_758798717_dg8_VISp | 44 | accuracy | 0.658333 |
| classification | mlp | Visual Coding | single_day | session_758798717_dg8_VISp | session_758798717_dg8_VISp | 46 | accuracy | 0.641667 |
| classification | mlp | Visual Coding | single_day | session_758798717_dg8_VISp | session_758798717_dg8_VISp | 54 | accuracy | 0.666667 |
| classification | mlp | Visual Coding | single_day | session_759883607_dg8_VISp | session_759883607_dg8_VISp | 44 | accuracy | 0.766667 |
| classification | mlp | Visual Coding | single_day | session_759883607_dg8_VISp | session_759883607_dg8_VISp | 46 | accuracy | 0.758333 |
| classification | mlp | Visual Coding | single_day | session_759883607_dg8_VISp | session_759883607_dg8_VISp | 54 | accuracy | 0.733333 |
| classification | mlp | Visual Coding | single_day | session_760345702_dg8_VISp | session_760345702_dg8_VISp | 44 | accuracy | 0.833333 |
| classification | mlp | Visual Coding | single_day | session_760345702_dg8_VISp | session_760345702_dg8_VISp | 46 | accuracy | 0.808333 |
| classification | mlp | Visual Coding | single_day | session_760345702_dg8_VISp | session_760345702_dg8_VISp | 54 | accuracy | 0.85 |
| classification | mlp | Visual Coding | single_day | session_760693773_dg8_VISp | session_760693773_dg8_VISp | 44 | accuracy | 0.9 |
| classification | mlp | Visual Coding | single_day | session_760693773_dg8_VISp | session_760693773_dg8_VISp | 46 | accuracy | 0.858333 |
| classification | mlp | Visual Coding | single_day | session_760693773_dg8_VISp | session_760693773_dg8_VISp | 54 | accuracy | 0.916667 |
| classification | mlp | Visual Coding | single_day | session_761418226_dg8_VISp | session_761418226_dg8_VISp | 44 | accuracy | 0.741667 |
| classification | mlp | Visual Coding | single_day | session_761418226_dg8_VISp | session_761418226_dg8_VISp | 46 | accuracy | 0.675 |
| classification | mlp | Visual Coding | single_day | session_761418226_dg8_VISp | session_761418226_dg8_VISp | 54 | accuracy | 0.725 |
| classification | mlp | Visual Coding | single_day | session_762120172_dg8_VISp | session_762120172_dg8_VISp | 44 | accuracy | 0.833333 |
| classification | mlp | Visual Coding | single_day | session_762120172_dg8_VISp | session_762120172_dg8_VISp | 46 | accuracy | 0.816667 |
| classification | mlp | Visual Coding | single_day | session_762120172_dg8_VISp | session_762120172_dg8_VISp | 54 | accuracy | 0.733333 |
| classification | mlp | Visual Coding | single_day | session_762602078_dg8_VISp | session_762602078_dg8_VISp | 44 | accuracy | 0.95 |
| classification | mlp | Visual Coding | single_day | session_762602078_dg8_VISp | session_762602078_dg8_VISp | 46 | accuracy | 0.908333 |
| classification | mlp | Visual Coding | single_day | session_762602078_dg8_VISp | session_762602078_dg8_VISp | 54 | accuracy | 0.875 |
| classification | mlp | Visual Coding | single_day | session_763673393_dg8_VISp | session_763673393_dg8_VISp | 44 | accuracy | 0.716667 |
| classification | mlp | Visual Coding | single_day | session_763673393_dg8_VISp | session_763673393_dg8_VISp | 46 | accuracy | 0.691667 |
| classification | mlp | Visual Coding | single_day | session_763673393_dg8_VISp | session_763673393_dg8_VISp | 54 | accuracy | 0.683333 |
| classification | mlp | Visual Coding | single_day | session_773418906_dg8_VISp | session_773418906_dg8_VISp | 44 | accuracy | 0.566667 |
| classification | mlp | Visual Coding | single_day | session_773418906_dg8_VISp | session_773418906_dg8_VISp | 46 | accuracy | 0.591667 |
| classification | mlp | Visual Coding | single_day | session_773418906_dg8_VISp | session_773418906_dg8_VISp | 54 | accuracy | 0.6 |
| classification | mlp | Visual Coding | single_day | session_791319847_dg8_VISp | session_791319847_dg8_VISp | 44 | accuracy | 0.925 |
| classification | mlp | Visual Coding | single_day | session_791319847_dg8_VISp | session_791319847_dg8_VISp | 46 | accuracy | 0.866667 |
| classification | mlp | Visual Coding | single_day | session_791319847_dg8_VISp | session_791319847_dg8_VISp | 54 | accuracy | 0.875 |
| classification | mlp | Visual Coding | single_day | session_797828357_dg8_VISp | session_797828357_dg8_VISp | 44 | accuracy | 0.758333 |
| classification | mlp | Visual Coding | single_day | session_797828357_dg8_VISp | session_797828357_dg8_VISp | 46 | accuracy | 0.683333 |
| classification | mlp | Visual Coding | single_day | session_797828357_dg8_VISp | session_797828357_dg8_VISp | 54 | accuracy | 0.775 |
| classification | mlp | Visual Coding | single_day | session_798911424_dg8_VISp | session_798911424_dg8_VISp | 44 | accuracy | 0.733333 |
| classification | mlp | Visual Coding | single_day | session_798911424_dg8_VISp | session_798911424_dg8_VISp | 46 | accuracy | 0.775 |
| classification | mlp | Visual Coding | single_day | session_798911424_dg8_VISp | session_798911424_dg8_VISp | 54 | accuracy | 0.775 |
| classification | mlp | Visual Coding | single_day | session_799864342_dg8_VISp | session_799864342_dg8_VISp | 44 | accuracy | 0.883333 |
| classification | mlp | Visual Coding | single_day | session_799864342_dg8_VISp | session_799864342_dg8_VISp | 46 | accuracy | 0.933333 |
| classification | mlp | Visual Coding | single_day | session_799864342_dg8_VISp | session_799864342_dg8_VISp | 54 | accuracy | 0.875 |
| classification | lstm | Visual Coding | cross_day | session_715093703_dg8_VISp__to__s... | session_762602078_dg8_VISp__to__s... | 44 | accuracy | 0.14881 |
| classification | lstm | Visual Coding | cross_day | session_715093703_dg8_VISp__to__s... | session_762602078_dg8_VISp__to__s... | 46 | accuracy | 0.130714 |

| | | | | | | | | |
|---|---|---|---|---|---|---|---|---|
| classification | lstm | Visual Coding | cross_day | session_715093703_dg8_VISp__to__s... | session_762602078_dg8_VISp__to__s... | 54 | accuracy | 0.12381 |
| classification | transformer | Visual Coding | cross_day | session_715093703_dg8_VISp__to__s... | session_762602078_dg8_VISp__to__s... | 44 | accuracy | 0.137857 |
| classification | transformer | Visual Coding | cross_day | session_715093703_dg8_VISp__to__s... | session_762602078_dg8_VISp__to__s... | 46 | accuracy | 0.11619 |
| classification | transformer | Visual Coding | cross_day | session_715093703_dg8_VISp__to__s... | session_762602078_dg8_VISp__to__s... | 54 | accuracy | 0.097381 |
| classification | lfads | Visual Coding | cross_day | session_715093703_dg8_VISp__to__s... | session_762602078_dg8_VISp__to__s... | 44 | accuracy | 0.127857 |
| classification | lfads | Visual Coding | cross_day | session_715093703_dg8_VISp__to__s... | session_762602078_dg8_VISp__to__s... | 46 | accuracy | 0.135952 |
| classification | lfads | Visual Coding | cross_day | session_715093703_dg8_VISp__to__s... | session_762602078_dg8_VISp__to__s... | 54 | accuracy | 0.147381 |
| classification | cycle_gan | Visual Coding | cross_day | session_715093703_dg8_VISp__to__s... | session_762602078_dg8_VISp__to__s... | 44 | accuracy | 0.149875 |
| classification | cycle_gan | Visual Coding | cross_day | session_715093703_dg8_VISp__to__s... | session_762602078_dg8_VISp__to__s... | 46 | accuracy | 0.1401 |
| classification | cycle_gan | Visual Coding | cross_day | session_715093703_dg8_VISp__to__s... | session_762602078_dg8_VISp__to__s... | 54 | accuracy | 0.154637 |
| classification | stabilization | Visual Coding | cross_day | session_715093703_dg8_VISp__to__s... | session_762602078_dg8_VISp__to__s... | 44 | accuracy | 0.122055 |
| classification | stabilization | Visual Coding | cross_day | session_715093703_dg8_VISp__to__s... | session_762602078_dg8_VISp__to__s... | 46 | accuracy | 0.106767 |
| classification | stabilization | Visual Coding | cross_day | session_715093703_dg8_VISp__to__s... | session_762602078_dg8_VISp__to__s... | 54 | accuracy | 0.107519 |
| classification | mfsnn | Visual Coding | cross_day | session_715093703_dg8_VISp__to__s... | session_762602078_dg8_VISp | 44 | accuracy | 0.11 |
| classification | mfsnn | Visual Coding | cross_day | session_715093703_dg8_VISp__to__s... | session_762602078_dg8_VISp | 46 | accuracy | 0.183333 |
| classification | mfsnn | Visual Coding | cross_day | session_715093703_dg8_VISp__to__s... | session_762602078_dg8_VISp | 54 | accuracy | 0.125 |
| classification | mfsnn | Visual Coding | cross_day | session_715093703_dg8_VISp__to__s... | session_763673393_dg8_VISp | 44 | accuracy | 0.126667 |
| classification | mfsnn | Visual Coding | cross_day | session_715093703_dg8_VISp__to__s... | session_763673393_dg8_VISp | 46 | accuracy | 0.133333 |
| classification | mfsnn | Visual Coding | cross_day | session_715093703_dg8_VISp__to__s... | session_763673393_dg8_VISp | 54 | accuracy | 0.125 |
| classification | mfsnn | Visual Coding | cross_day | session_715093703_dg8_VISp__to__s... | session_773418906_dg8_VISp | 44 | accuracy | 0.125 |
| classification | mfsnn | Visual Coding | cross_day | session_715093703_dg8_VISp__to__s... | session_773418906_dg8_VISp | 46 | accuracy | 0.111667 |
| classification | mfsnn | Visual Coding | cross_day | session_715093703_dg8_VISp__to__s... | session_773418906_dg8_VISp | 54 | accuracy | 0.125 |
| classification | mfsnn | Visual Coding | cross_day | session_715093703_dg8_VISp__to__s... | session_791319847_dg8_VISp | 44 | accuracy | 0.125 |
| classification | mfsnn | Visual Coding | cross_day | session_715093703_dg8_VISp__to__s... | session_791319847_dg8_VISp | 46 | accuracy | 0.141667 |
| classification | mfsnn | Visual Coding | cross_day | session_715093703_dg8_VISp__to__s... | session_791319847_dg8_VISp | 54 | accuracy | 0.125 |
| classification | mfsnn | Visual Coding | cross_day | session_715093703_dg8_VISp__to__s... | session_797828357_dg8_VISp | 44 | accuracy | 0.241667 |
| classification | mfsnn | Visual Coding | cross_day | session_715093703_dg8_VISp__to__s... | session_797828357_dg8_VISp | 46 | accuracy | 0.151667 |
| classification | mfsnn | Visual Coding | cross_day | session_715093703_dg8_VISp__to__s... | session_797828357_dg8_VISp | 54 | accuracy | 0.105 |
| classification | mfsnn | Visual Coding | cross_day | session_715093703_dg8_VISp__to__s... | session_798911424_dg8_VISp | 44 | accuracy | 0.125 |
| classification | mfsnn | Visual Coding | cross_day | session_715093703_dg8_VISp__to__s... | session_798911424_dg8_VISp | 46 | accuracy | 0.113333 |
| classification | mfsnn | Visual Coding | cross_day | session_715093703_dg8_VISp__to__s... | session_798911424_dg8_VISp | 54 | accuracy | 0.125 |
| classification | mfsnn | Visual Coding | cross_day | session_715093703_dg8_VISp__to__s... | session_799864342_dg8_VISp | 44 | accuracy | 0.0483333 |
| classification | mfsnn | Visual Coding | cross_day | session_715093703_dg8_VISp__to__s... | session_799864342_dg8_VISp | 46 | accuracy | 0.13 |
| classification | mfsnn | Visual Coding | cross_day | session_715093703_dg8_VISp__to__s... | session_799864342_dg8_VISp | 54 | accuracy | 0.125 |
| classification | mscformer | Visual Coding | cross_day | session_715093703_dg8_VISp__to__s... | session_762602078_dg8_VISp | 44 | accuracy | 0.11 |
| classification | mscformer | Visual Coding | cross_day | session_715093703_dg8_VISp__to__s... | session_762602078_dg8_VISp | 46 | accuracy | 0.14 |
| classification | mscformer | Visual Coding | cross_day | session_715093703_dg8_VISp__to__s... | session_762602078_dg8_VISp | 54 | accuracy | 0.128333 |
| classification | mscformer | Visual Coding | cross_day | session_715093703_dg8_VISp__to__s... | session_763673393_dg8_VISp | 44 | accuracy | 0.1 |
| classification | mscformer | Visual Coding | cross_day | session_715093703_dg8_VISp__to__s... | session_763673393_dg8_VISp | 46 | accuracy | 0.106667 |
| classification | mscformer | Visual Coding | cross_day | session_715093703_dg8_VISp__to__s... | session_763673393_dg8_VISp | 54 | accuracy | 0.12 |
| classification | mscformer | Visual Coding | cross_day | session_715093703_dg8_VISp__to__s... | session_773418906_dg8_VISp | 44 | accuracy | 0.16 |
| classification | mscformer | Visual Coding | cross_day | session_715093703_dg8_VISp__to__s... | session_773418906_dg8_VISp | 46 | accuracy | 0.158333 |
| classification | mscformer | Visual Coding | cross_day | session_715093703_dg8_VISp__to__s... | session_773418906_dg8_VISp | 54 | accuracy | 0.183333 |
| classification | mscformer | Visual Coding | cross_day | session_715093703_dg8_VISp__to__s... | session_791319847_dg8_VISp | 44 | accuracy | 0.0883333 |
| classification | mscformer | Visual Coding | cross_day | session_715093703_dg8_VISp__to__s... | session_791319847_dg8_VISp | 46 | accuracy | 0.0783333 |
| classification | mscformer | Visual Coding | cross_day | session_715093703_dg8_VISp__to__s... | session_791319847_dg8_VISp | 54 | accuracy | 0.075 |
| classification | mscformer | Visual Coding | cross_day | session_715093703_dg8_VISp__to__s... | session_797828357_dg8_VISp | 44 | accuracy | 0.0883333 |
| classification | mscformer | Visual Coding | cross_day | session_715093703_dg8_VISp__to__s... | session_797828357_dg8_VISp | 46 | accuracy | 0.085 |
| classification | mscformer | Visual Coding | cross_day | session_715093703_dg8_VISp__to__s... | session_797828357_dg8_VISp | 54 | accuracy | 0.115 |
| classification | mscformer | Visual Coding | cross_day | session_715093703_dg8_VISp__to__s... | session_798911424_dg8_VISp | 44 | accuracy | 0.131667 |
| classification | mscformer | Visual Coding | cross_day | session_715093703_dg8_VISp__to__s... | session_798911424_dg8_VISp | 46 | accuracy | 0.12 |
| classification | mscformer | Visual Coding | cross_day | session_715093703_dg8_VISp__to__s... | session_798911424_dg8_VISp | 54 | accuracy | 0.085 |
| classification | mscformer | Visual Coding | cross_day | session_715093703_dg8_VISp__to__s... | session_799864342_dg8_VISp | 44 | accuracy | 0.145 |
| classification | mscformer | Visual Coding | cross_day | session_715093703_dg8_VISp__to__s... | session_799864342_dg8_VISp | 46 | accuracy | 0.146667 |
| classification | mscformer | Visual Coding | cross_day | session_715093703_dg8_VISp__to__s... | session_799864342_dg8_VISp | 54 | accuracy | 0.165 |
| classification | nomad | Visual Coding | cross_day | session_715093703_dg8_VISp__to__s... | session_762602078_dg8_VISp | 44 | accuracy | 0.08 |
| classification | nomad | Visual Coding | cross_day | session_715093703_dg8_VISp__to__s... | session_762602078_dg8_VISp | 46 | accuracy | 0.125 |
| classification | nomad | Visual Coding | cross_day | session_715093703_dg8_VISp__to__s... | session_762602078_dg8_VISp | 54 | accuracy | 0.151667 |
| classification | nomad | Visual Coding | cross_day | session_715093703_dg8_VISp__to__s... | session_763673393_dg8_VISp | 44 | accuracy | 0.103333 |
| classification | nomad | Visual Coding | cross_day | session_715093703_dg8_VISp__to__s... | session_763673393_dg8_VISp | 46 | accuracy | 0.128333 |
| classification | nomad | Visual Coding | cross_day | session_715093703_dg8_VISp__to__s... | session_763673393_dg8_VISp | 54 | accuracy | 0.146667 |
| classification | nomad | Visual Coding | cross_day | session_715093703_dg8_VISp__to__s... | session_773418906_dg8_VISp | 44 | accuracy | 0.171667 |
| classification | nomad | Visual Coding | cross_day | session_715093703_dg8_VISp__to__s... | session_773418906_dg8_VISp | 46 | accuracy | 0.221667 |

| | | | | | | | | |
|---|---|---|---|---|---|---|---|---|
| classification | nomad | Visual Coding | cross_day | session_715093703_dg8_VISp__to__s... | session_773418906_dg8_VISp | 54 | accuracy | 0.203333 |
| classification | nomad | Visual Coding | cross_day | session_715093703_dg8_VISp__to__s... | session_791319847_dg8_VISp | 44 | accuracy | 0.16 |
| classification | nomad | Visual Coding | cross_day | session_715093703_dg8_VISp__to__s... | session_791319847_dg8_VISp | 46 | accuracy | 0.128333 |
| classification | nomad | Visual Coding | cross_day | session_715093703_dg8_VISp__to__s... | session_791319847_dg8_VISp | 54 | accuracy | 0.126667 |
| classification | nomad | Visual Coding | cross_day | session_715093703_dg8_VISp__to__s... | session_797828357_dg8_VISp | 44 | accuracy | 0.171667 |
| classification | nomad | Visual Coding | cross_day | session_715093703_dg8_VISp__to__s... | session_797828357_dg8_VISp | 46 | accuracy | 0.101667 |
| classification | nomad | Visual Coding | cross_day | session_715093703_dg8_VISp__to__s... | session_797828357_dg8_VISp | 54 | accuracy | 0.175 |
| classification | nomad | Visual Coding | cross_day | session_715093703_dg8_VISp__to__s... | session_798911424_dg8_VISp | 44 | accuracy | 0.0566667 |
| classification | nomad | Visual Coding | cross_day | session_715093703_dg8_VISp__to__s... | session_798911424_dg8_VISp | 46 | accuracy | 0.0866667 |
| classification | nomad | Visual Coding | cross_day | session_715093703_dg8_VISp__to__s... | session_798911424_dg8_VISp | 54 | accuracy | 0.143333 |
| classification | nomad | Visual Coding | cross_day | session_715093703_dg8_VISp__to__s... | session_799864342_dg8_VISp | 44 | accuracy | 0.0716667 |
| classification | nomad | Visual Coding | cross_day | session_715093703_dg8_VISp__to__s... | session_799864342_dg8_VISp | 46 | accuracy | 0.0433333 |
| classification | nomad | Visual Coding | cross_day | session_715093703_dg8_VISp__to__s... | session_799864342_dg8_VISp | 54 | accuracy | 0.0733333 |
| classification | mlp | Visual Coding | cross_day | session_715093703_dg8_VISp__to__s... | session_762602078_dg8_VISp | 44 | accuracy | 0.106667 |
| classification | mlp | Visual Coding | cross_day | session_715093703_dg8_VISp__to__s... | session_762602078_dg8_VISp | 46 | accuracy | 0.0966667 |
| classification | mlp | Visual Coding | cross_day | session_715093703_dg8_VISp__to__s... | session_762602078_dg8_VISp | 54 | accuracy | 0.208333 |
| classification | mlp | Visual Coding | cross_day | session_715093703_dg8_VISp__to__s... | session_763673393_dg8_VISp | 44 | accuracy | 0.19 |
| classification | mlp | Visual Coding | cross_day | session_715093703_dg8_VISp__to__s... | session_763673393_dg8_VISp | 46 | accuracy | 0.166667 |
| classification | mlp | Visual Coding | cross_day | session_715093703_dg8_VISp__to__s... | session_763673393_dg8_VISp | 54 | accuracy | 0.13 |
| classification | mlp | Visual Coding | cross_day | session_715093703_dg8_VISp__to__s... | session_773418906_dg8_VISp | 44 | accuracy | 0.171667 |
| classification | mlp | Visual Coding | cross_day | session_715093703_dg8_VISp__to__s... | session_773418906_dg8_VISp | 46 | accuracy | 0.155 |
| classification | mlp | Visual Coding | cross_day | session_715093703_dg8_VISp__to__s... | session_773418906_dg8_VISp | 54 | accuracy | 0.211667 |
| classification | mlp | Visual Coding | cross_day | session_715093703_dg8_VISp__to__s... | session_791319847_dg8_VISp | 44 | accuracy | 0.0466667 |
| classification | mlp | Visual Coding | cross_day | session_715093703_dg8_VISp__to__s... | session_791319847_dg8_VISp | 46 | accuracy | 0.055 |
| classification | mlp | Visual Coding | cross_day | session_715093703_dg8_VISp__to__s... | session_791319847_dg8_VISp | 54 | accuracy | 0.075 |
| classification | mlp | Visual Coding | cross_day | session_715093703_dg8_VISp__to__s... | session_797828357_dg8_VISp | 44 | accuracy | 0.136667 |
| classification | mlp | Visual Coding | cross_day | session_715093703_dg8_VISp__to__s... | session_797828357_dg8_VISp | 46 | accuracy | 0.128333 |
| classification | mlp | Visual Coding | cross_day | session_715093703_dg8_VISp__to__s... | session_797828357_dg8_VISp | 54 | accuracy | 0.08 |
| classification | mlp | Visual Coding | cross_day | session_715093703_dg8_VISp__to__s... | session_798911424_dg8_VISp | 44 | accuracy | 0.0933333 |
| classification | mlp | Visual Coding | cross_day | session_715093703_dg8_VISp__to__s... | session_798911424_dg8_VISp | 46 | accuracy | 0.0616667 |
| classification | mlp | Visual Coding | cross_day | session_715093703_dg8_VISp__to__s... | session_798911424_dg8_VISp | 54 | accuracy | 0.148333 |
| classification | mlp | Visual Coding | cross_day | session_715093703_dg8_VISp__to__s... | session_799864342_dg8_VISp | 44 | accuracy | 0.0583333 |
| classification | mlp | Visual Coding | cross_day | session_715093703_dg8_VISp__to__s... | session_799864342_dg8_VISp | 46 | accuracy | 0.0983333 |
| classification | mlp | Visual Coding | cross_day | session_715093703_dg8_VISp__to__s... | session_799864342_dg8_VISp | 54 | accuracy | 0.085 |
| classification | lstm | Music | single_day | All_RISE_IEEG_preprocessed_final | All_RISE_IEEG_preprocessed_final | 44 | accuracy | 0.829787 |
| classification | lstm | Music | single_day | All_RISE_IEEG_preprocessed_final | All_RISE_IEEG_preprocessed_final | 46 | accuracy | 0.837524 |
| classification | lstm | Music | single_day | All_RISE_IEEG_preprocessed_final | All_RISE_IEEG_preprocessed_final | 54 | accuracy | 0.823985 |
| classification | lstm | Music | single_day | Patient00 | Patient00 | 44 | accuracy | 0.944444 |
| classification | lstm | Music | single_day | Patient00 | Patient00 | 46 | accuracy | 0.944444 |
| classification | lstm | Music | single_day | Patient00 | Patient00 | 54 | accuracy | 0.944444 |
| classification | lstm | Music | single_day | Patient01 | Patient01 | 44 | accuracy | 0.944444 |
| classification | lstm | Music | single_day | Patient01 | Patient01 | 46 | accuracy | 0.944444 |
| classification | lstm | Music | single_day | Patient01 | Patient01 | 54 | accuracy | 0.888889 |
| classification | lstm | Music | single_day | Patient02 | Patient02 | 44 | accuracy | 0.944444 |
| classification | lstm | Music | single_day | Patient02 | Patient02 | 46 | accuracy | 0.944444 |
| classification | lstm | Music | single_day | Patient02 | Patient02 | 54 | accuracy | 1 |
| classification | lstm | Music | single_day | Patient03 | Patient03 | 44 | accuracy | 0.833333 |
| classification | lstm | Music | single_day | Patient03 | Patient03 | 46 | accuracy | 0.833333 |
| classification | lstm | Music | single_day | Patient03 | Patient03 | 54 | accuracy | 0.833333 |
| classification | lstm | Music | single_day | Patient04 | Patient04 | 44 | accuracy | 1 |
| classification | lstm | Music | single_day | Patient04 | Patient04 | 46 | accuracy | 0.833333 |
| classification | lstm | Music | single_day | Patient04 | Patient04 | 54 | accuracy | 0.833333 |
| classification | lstm | Music | single_day | Patient05 | Patient05 | 44 | accuracy | 0.888889 |
| classification | lstm | Music | single_day | Patient05 | Patient05 | 46 | accuracy | 1 |
| classification | lstm | Music | single_day | Patient05 | Patient05 | 54 | accuracy | 0.888889 |
| classification | lstm | Music | single_day | Patient06 | Patient06 | 44 | accuracy | 0.888889 |
| classification | lstm | Music | single_day | Patient06 | Patient06 | 46 | accuracy | 0.888889 |
| classification | lstm | Music | single_day | Patient06 | Patient06 | 54 | accuracy | 0.888889 |
| classification | lstm | Music | single_day | Patient07 | Patient07 | 44 | accuracy | 0.833333 |
| classification | lstm | Music | single_day | Patient07 | Patient07 | 46 | accuracy | 0.888889 |
| classification | lstm | Music | single_day | Patient07 | Patient07 | 54 | accuracy | 0.888889 |
| classification | lstm | Music | single_day | Patient08 | Patient08 | 44 | accuracy | 0.833333 |
| classification | lstm | Music | single_day | Patient08 | Patient08 | 46 | accuracy | 0.888889 |

| | | | | | | | | |
|---|---|---|---|---|---|---|---|---|
| classification | lstm | Music | single_day | Patient08 | Patient08 | 54 | accuracy | 0.944444 |
| classification | lstm | Music | single_day | Patient09 | Patient09 | 44 | accuracy | 0.833333 |
| classification | lstm | Music | single_day | Patient09 | Patient09 | 46 | accuracy | 0.833333 |
| classification | lstm | Music | single_day | Patient09 | Patient09 | 54 | accuracy | 0.833333 |
| classification | lstm | Music | single_day | Patient10 | Patient10 | 44 | accuracy | 0.833333 |
| classification | lstm | Music | single_day | Patient10 | Patient10 | 46 | accuracy | 0.944444 |
| classification | lstm | Music | single_day | Patient10 | Patient10 | 54 | accuracy | 0.833333 |
| classification | lstm | Music | single_day | Patient11 | Patient11 | 44 | accuracy | 0.944444 |
| classification | lstm | Music | single_day | Patient11 | Patient11 | 46 | accuracy | 0.888889 |
| classification | lstm | Music | single_day | Patient11 | Patient11 | 54 | accuracy | 0.777778 |
| classification | lstm | Music | single_day | Patient12 | Patient12 | 44 | accuracy | 0.833333 |
| classification | lstm | Music | single_day | Patient12 | Patient12 | 46 | accuracy | 0.833333 |
| classification | lstm | Music | single_day | Patient12 | Patient12 | 54 | accuracy | 0.777778 |
| classification | lstm | Music | single_day | Patient13 | Patient13 | 44 | accuracy | 0.833333 |
| classification | lstm | Music | single_day | Patient13 | Patient13 | 46 | accuracy | 0.888889 |
| classification | lstm | Music | single_day | Patient13 | Patient13 | 54 | accuracy | 0.888889 |
| classification | lstm | Music | single_day | Patient14 | Patient14 | 44 | accuracy | 0.888889 |
| classification | lstm | Music | single_day | Patient14 | Patient14 | 46 | accuracy | 1 |
| classification | lstm | Music | single_day | Patient14 | Patient14 | 54 | accuracy | 1 |
| classification | lstm | Music | single_day | Patient15 | Patient15 | 44 | accuracy | 0.944444 |
| classification | lstm | Music | single_day | Patient15 | Patient15 | 46 | accuracy | 1 |
| classification | lstm | Music | single_day | Patient15 | Patient15 | 54 | accuracy | 0.944444 |
| classification | lstm | Music | single_day | Patient16 | Patient16 | 44 | accuracy | 0.888889 |
| classification | lstm | Music | single_day | Patient16 | Patient16 | 46 | accuracy | 0.777778 |
| classification | lstm | Music | single_day | Patient16 | Patient16 | 54 | accuracy | 0.833333 |
| classification | lstm | Music | single_day | Patient17 | Patient17 | 44 | accuracy | 0.833333 |
| classification | lstm | Music | single_day | Patient17 | Patient17 | 46 | accuracy | 0.833333 |
| classification | lstm | Music | single_day | Patient17 | Patient17 | 54 | accuracy | 0.888889 |
| classification | lstm | Music | single_day | Patient18 | Patient18 | 44 | accuracy | 0.944444 |
| classification | lstm | Music | single_day | Patient18 | Patient18 | 46 | accuracy | 0.888889 |
| classification | lstm | Music | single_day | Patient18 | Patient18 | 54 | accuracy | 0.944444 |
| classification | lstm | Music | single_day | Patient19 | Patient19 | 44 | accuracy | 0.888889 |
| classification | lstm | Music | single_day | Patient19 | Patient19 | 46 | accuracy | 0.888889 |
| classification | lstm | Music | single_day | Patient19 | Patient19 | 54 | accuracy | 0.888889 |
| classification | lstm | Music | single_day | Patient20 | Patient20 | 44 | accuracy | 0.833333 |
| classification | lstm | Music | single_day | Patient20 | Patient20 | 46 | accuracy | 0.833333 |
| classification | lstm | Music | single_day | Patient20 | Patient20 | 54 | accuracy | 0.666667 |
| classification | lstm | Music | single_day | Patient21 | Patient21 | 44 | accuracy | 0.944444 |
| classification | lstm | Music | single_day | Patient21 | Patient21 | 46 | accuracy | 0.833333 |
| classification | lstm | Music | single_day | Patient21 | Patient21 | 54 | accuracy | 0.888889 |
| classification | lstm | Music | single_day | Patient22 | Patient22 | 44 | accuracy | 0.833333 |
| classification | lstm | Music | single_day | Patient22 | Patient22 | 46 | accuracy | 0.888889 |
| classification | lstm | Music | single_day | Patient22 | Patient22 | 54 | accuracy | 0.833333 |
| classification | lstm | Music | single_day | Patient23 | Patient23 | 44 | accuracy | 0.833333 |
| classification | lstm | Music | single_day | Patient23 | Patient23 | 46 | accuracy | 0.833333 |
| classification | lstm | Music | single_day | Patient23 | Patient23 | 54 | accuracy | 0.833333 |
| classification | lstm | Music | single_day | Patient24 | Patient24 | 44 | accuracy | 0.944444 |
| classification | lstm | Music | single_day | Patient24 | Patient24 | 46 | accuracy | 0.944444 |
| classification | lstm | Music | single_day | Patient24 | Patient24 | 54 | accuracy | 0.888889 |
| classification | lstm | Music | single_day | Patient25 | Patient25 | 44 | accuracy | 1 |
| classification | lstm | Music | single_day | Patient25 | Patient25 | 46 | accuracy | 0.888889 |
| classification | lstm | Music | single_day | Patient25 | Patient25 | 54 | accuracy | 0.944444 |
| classification | lstm | Music | single_day | Patient26 | Patient26 | 44 | accuracy | 0.833333 |
| classification | lstm | Music | single_day | Patient26 | Patient26 | 46 | accuracy | 0.833333 |
| classification | lstm | Music | single_day | Patient26 | Patient26 | 54 | accuracy | 0.833333 |
| classification | lstm | Music | single_day | Patient27 | Patient27 | 44 | accuracy | 0.833333 |
| classification | lstm | Music | single_day | Patient27 | Patient27 | 46 | accuracy | 0.833333 |
| classification | lstm | Music | single_day | Patient27 | Patient27 | 54 | accuracy | 0.833333 |
| classification | lstm | Music | single_day | Patient28 | Patient28 | 44 | accuracy | 0.944444 |
| classification | lstm | Music | single_day | Patient28 | Patient28 | 46 | accuracy | 1 |
| classification | lstm | Music | single_day | Patient28 | Patient28 | 54 | accuracy | 0.944444 |
| classification | transformer | Music | single_day | All_RISE_IEEG_preprocessed_final | All_RISE_IEEG_preprocessed_final | 44 | accuracy | 0.823985 |
| classification | transformer | Music | single_day | All_RISE_IEEG_preprocessed_final | All_RISE_IEEG_preprocessed_final | 46 | accuracy | 0.814313 |

| | | | | | | | | |
|---|---|---|---|---|---|---|---|---|
| classification | transformer | Music | single_day | All_RISE_IEEG_preprocessed_final | All_RISE_IEEG_preprocessed_final | 54 | accuracy | 0.820116 |
| classification | transformer | Music | single_day | Patient00 | Patient00 | 44 | accuracy | 0.888889 |
| classification | transformer | Music | single_day | Patient00 | Patient00 | 46 | accuracy | 0.944444 |
| classification | transformer | Music | single_day | Patient00 | Patient00 | 54 | accuracy | 0.888889 |
| classification | transformer | Music | single_day | Patient01 | Patient01 | 44 | accuracy | 0.833333 |
| classification | transformer | Music | single_day | Patient01 | Patient01 | 46 | accuracy | 0.833333 |
| classification | transformer | Music | single_day | Patient01 | Patient01 | 54 | accuracy | 0.833333 |
| classification | transformer | Music | single_day | Patient02 | Patient02 | 44 | accuracy | 0.944444 |
| classification | transformer | Music | single_day | Patient02 | Patient02 | 46 | accuracy | 1 |
| classification | transformer | Music | single_day | Patient02 | Patient02 | 54 | accuracy | 0.777778 |
| classification | transformer | Music | single_day | Patient03 | Patient03 | 44 | accuracy | 0.833333 |
| classification | transformer | Music | single_day | Patient03 | Patient03 | 46 | accuracy | 0.833333 |
| classification | transformer | Music | single_day | Patient03 | Patient03 | 54 | accuracy | 0.833333 |
| classification | transformer | Music | single_day | Patient04 | Patient04 | 44 | accuracy | 0.833333 |
| classification | transformer | Music | single_day | Patient04 | Patient04 | 46 | accuracy | 0.833333 |
| classification | transformer | Music | single_day | Patient04 | Patient04 | 54 | accuracy | 0.833333 |
| classification | transformer | Music | single_day | Patient05 | Patient05 | 44 | accuracy | 0.888889 |
| classification | transformer | Music | single_day | Patient05 | Patient05 | 46 | accuracy | 1 |
| classification | transformer | Music | single_day | Patient05 | Patient05 | 54 | accuracy | 0.888889 |
| classification | transformer | Music | single_day | Patient06 | Patient06 | 44 | accuracy | 0.888889 |
| classification | transformer | Music | single_day | Patient06 | Patient06 | 46 | accuracy | 0.833333 |
| classification | transformer | Music | single_day | Patient06 | Patient06 | 54 | accuracy | 0.888889 |
| classification | transformer | Music | single_day | Patient07 | Patient07 | 44 | accuracy | 0.833333 |
| classification | transformer | Music | single_day | Patient07 | Patient07 | 46 | accuracy | 0.833333 |
| classification | transformer | Music | single_day | Patient07 | Patient07 | 54 | accuracy | 0.833333 |
| classification | transformer | Music | single_day | Patient08 | Patient08 | 44 | accuracy | 0.944444 |
| classification | transformer | Music | single_day | Patient08 | Patient08 | 46 | accuracy | 0.833333 |
| classification | transformer | Music | single_day | Patient08 | Patient08 | 54 | accuracy | 0.833333 |
| classification | transformer | Music | single_day | Patient09 | Patient09 | 44 | accuracy | 0.833333 |
| classification | transformer | Music | single_day | Patient09 | Patient09 | 46 | accuracy | 0.833333 |
| classification | transformer | Music | single_day | Patient09 | Patient09 | 54 | accuracy | 0.833333 |
| classification | transformer | Music | single_day | Patient10 | Patient10 | 44 | accuracy | 0.777778 |
| classification | transformer | Music | single_day | Patient10 | Patient10 | 46 | accuracy | 0.944444 |
| classification | transformer | Music | single_day | Patient10 | Patient10 | 54 | accuracy | 0.888889 |
| classification | transformer | Music | single_day | Patient11 | Patient11 | 44 | accuracy | 0.833333 |
| classification | transformer | Music | single_day | Patient11 | Patient11 | 46 | accuracy | 0.833333 |
| classification | transformer | Music | single_day | Patient11 | Patient11 | 54 | accuracy | 0.833333 |
| classification | transformer | Music | single_day | Patient12 | Patient12 | 44 | accuracy | 0.833333 |
| classification | transformer | Music | single_day | Patient12 | Patient12 | 46 | accuracy | 0.833333 |
| classification | transformer | Music | single_day | Patient12 | Patient12 | 54 | accuracy | 0.833333 |
| classification | transformer | Music | single_day | Patient13 | Patient13 | 44 | accuracy | 0.888889 |
| classification | transformer | Music | single_day | Patient13 | Patient13 | 46 | accuracy | 0.944444 |
| classification | transformer | Music | single_day | Patient13 | Patient13 | 54 | accuracy | 0.888889 |
| classification | transformer | Music | single_day | Patient14 | Patient14 | 44 | accuracy | 0.888889 |
| classification | transformer | Music | single_day | Patient14 | Patient14 | 46 | accuracy | 1 |
| classification | transformer | Music | single_day | Patient14 | Patient14 | 54 | accuracy | 0.833333 |
| classification | transformer | Music | single_day | Patient15 | Patient15 | 44 | accuracy | 0.833333 |
| classification | transformer | Music | single_day | Patient15 | Patient15 | 46 | accuracy | 0.888889 |
| classification | transformer | Music | single_day | Patient15 | Patient15 | 54 | accuracy | 0.944444 |
| classification | transformer | Music | single_day | Patient16 | Patient16 | 44 | accuracy | 0.888889 |
| classification | transformer | Music | single_day | Patient16 | Patient16 | 46 | accuracy | 0.777778 |
| classification | transformer | Music | single_day | Patient16 | Patient16 | 54 | accuracy | 0.888889 |
| classification | transformer | Music | single_day | Patient17 | Patient17 | 44 | accuracy | 0.777778 |
| classification | transformer | Music | single_day | Patient17 | Patient17 | 46 | accuracy | 0.888889 |
| classification | transformer | Music | single_day | Patient17 | Patient17 | 54 | accuracy | 0.777778 |
| classification | transformer | Music | single_day | Patient18 | Patient18 | 44 | accuracy | 1 |
| classification | transformer | Music | single_day | Patient18 | Patient18 | 46 | accuracy | 0.888889 |
| classification | transformer | Music | single_day | Patient18 | Patient18 | 54 | accuracy | 0.888889 |
| classification | transformer | Music | single_day | Patient19 | Patient19 | 44 | accuracy | 0.833333 |
| classification | transformer | Music | single_day | Patient19 | Patient19 | 46 | accuracy | 0.833333 |
| classification | transformer | Music | single_day | Patient19 | Patient19 | 54 | accuracy | 0.833333 |
| classification | transformer | Music | single_day | Patient20 | Patient20 | 44 | accuracy | 0.833333 |
| classification | transformer | Music | single_day | Patient20 | Patient20 | 46 | accuracy | 0.833333 |

| | | | | | | | | |
|---|---|---|---|---|---|---|---|---|
| classification | transformer | Music | single_day | Patient20 | Patient20 | 54 | accuracy | 0.833333 |
| classification | transformer | Music | single_day | Patient21 | Patient21 | 44 | accuracy | 0.944444 |
| classification | transformer | Music | single_day | Patient21 | Patient21 | 46 | accuracy | 0.888889 |
| classification | transformer | Music | single_day | Patient21 | Patient21 | 54 | accuracy | 0.833333 |
| classification | transformer | Music | single_day | Patient22 | Patient22 | 44 | accuracy | 0.888889 |
| classification | transformer | Music | single_day | Patient22 | Patient22 | 46 | accuracy | 0.833333 |
| classification | transformer | Music | single_day | Patient22 | Patient22 | 54 | accuracy | 0.833333 |
| classification | transformer | Music | single_day | Patient23 | Patient23 | 44 | accuracy | 0.666667 |
| classification | transformer | Music | single_day | Patient23 | Patient23 | 46 | accuracy | 0.833333 |
| classification | transformer | Music | single_day | Patient23 | Patient23 | 54 | accuracy | 0.833333 |
| classification | transformer | Music | single_day | Patient24 | Patient24 | 44 | accuracy | 0.888889 |
| classification | transformer | Music | single_day | Patient24 | Patient24 | 46 | accuracy | 1 |
| classification | transformer | Music | single_day | Patient24 | Patient24 | 54 | accuracy | 0.888889 |
| classification | transformer | Music | single_day | Patient25 | Patient25 | 44 | accuracy | 0.944444 |
| classification | transformer | Music | single_day | Patient25 | Patient25 | 46 | accuracy | 0.888889 |
| classification | transformer | Music | single_day | Patient25 | Patient25 | 54 | accuracy | 0.888889 |
| classification | transformer | Music | single_day | Patient26 | Patient26 | 44 | accuracy | 0.833333 |
| classification | transformer | Music | single_day | Patient26 | Patient26 | 46 | accuracy | 0.833333 |
| classification | transformer | Music | single_day | Patient26 | Patient26 | 54 | accuracy | 0.888889 |
| classification | transformer | Music | single_day | Patient27 | Patient27 | 44 | accuracy | 0.888889 |
| classification | transformer | Music | single_day | Patient27 | Patient27 | 46 | accuracy | 0.777778 |
| classification | transformer | Music | single_day | Patient27 | Patient27 | 54 | accuracy | 0.833333 |
| classification | transformer | Music | single_day | Patient28 | Patient28 | 44 | accuracy | 1 |
| classification | transformer | Music | single_day | Patient28 | Patient28 | 46 | accuracy | 1 |
| classification | transformer | Music | single_day | Patient28 | Patient28 | 54 | accuracy | 0.888889 |
| classification | lfads | Music | single_day | All_RISE_IEEG_preprocessed_final | All_RISE_IEEG_preprocessed_final | 44 | accuracy | 0.729207 |
| classification | lfads | Music | single_day | Patient00 | Patient00 | 44 | accuracy | 0.944444 |
| classification | lfads | Music | single_day | Patient00 | Patient00 | 46 | accuracy | 0.888889 |
| classification | lfads | Music | single_day | Patient00 | Patient00 | 54 | accuracy | 0.833333 |
| classification | lfads | Music | single_day | Patient01 | Patient01 | 44 | accuracy | 0.777778 |
| classification | lfads | Music | single_day | Patient01 | Patient01 | 46 | accuracy | 0.888889 |
| classification | lfads | Music | single_day | Patient01 | Patient01 | 54 | accuracy | 0.777778 |
| classification | lfads | Music | single_day | Patient02 | Patient02 | 44 | accuracy | 0.888889 |
| classification | lfads | Music | single_day | Patient02 | Patient02 | 46 | accuracy | 0.944444 |
| classification | lfads | Music | single_day | Patient02 | Patient02 | 54 | accuracy | 0.833333 |
| classification | lfads | Music | single_day | Patient03 | Patient03 | 44 | accuracy | 0.833333 |
| classification | lfads | Music | single_day | Patient03 | Patient03 | 46 | accuracy | 0.722222 |
| classification | lfads | Music | single_day | Patient03 | Patient03 | 54 | accuracy | 0.888889 |
| classification | lfads | Music | single_day | Patient04 | Patient04 | 44 | accuracy | 0.833333 |
| classification | lfads | Music | single_day | Patient04 | Patient04 | 46 | accuracy | 0.833333 |
| classification | lfads | Music | single_day | Patient04 | Patient04 | 54 | accuracy | 0.888889 |
| classification | lfads | Music | single_day | Patient05 | Patient05 | 44 | accuracy | 1 |
| classification | lfads | Music | single_day | Patient05 | Patient05 | 46 | accuracy | 1 |
| classification | lfads | Music | single_day | Patient05 | Patient05 | 54 | accuracy | 0.888889 |
| classification | lfads | Music | single_day | Patient06 | Patient06 | 44 | accuracy | 0.833333 |
| classification | lfads | Music | single_day | Patient06 | Patient06 | 46 | accuracy | 0.777778 |
| classification | lfads | Music | single_day | Patient06 | Patient06 | 54 | accuracy | 0.888889 |
| classification | lfads | Music | single_day | Patient07 | Patient07 | 44 | accuracy | 0.888889 |
| classification | lfads | Music | single_day | Patient07 | Patient07 | 46 | accuracy | 0.722222 |
| classification | lfads | Music | single_day | Patient07 | Patient07 | 54 | accuracy | 0.777778 |
| classification | lfads | Music | single_day | Patient08 | Patient08 | 44 | accuracy | 0.888889 |
| classification | lfads | Music | single_day | Patient08 | Patient08 | 46 | accuracy | 0.833333 |
| classification | lfads | Music | single_day | Patient08 | Patient08 | 54 | accuracy | 0.833333 |
| classification | lfads | Music | single_day | Patient09 | Patient09 | 44 | accuracy | 0.833333 |
| classification | lfads | Music | single_day | Patient09 | Patient09 | 46 | accuracy | 0.722222 |
| classification | lfads | Music | single_day | Patient09 | Patient09 | 54 | accuracy | 0.888889 |
| classification | lfads | Music | single_day | Patient10 | Patient10 | 44 | accuracy | 0.833333 |
| classification | lfads | Music | single_day | Patient10 | Patient10 | 46 | accuracy | 0.722222 |
| classification | lfads | Music | single_day | Patient10 | Patient10 | 54 | accuracy | 0.777778 |
| classification | lfads | Music | single_day | Patient11 | Patient11 | 44 | accuracy | 0.722222 |
| classification | lfads | Music | single_day | Patient11 | Patient11 | 46 | accuracy | 0.833333 |
| classification | lfads | Music | single_day | Patient11 | Patient11 | 54 | accuracy | 0.833333 |
| classification | lfads | Music | single_day | Patient12 | Patient12 | 44 | accuracy | 0.888889 |

| | | | | | | | | |
|---|---|---|---|---|---|---|---|---|
| classification | lfads | Music | single_day | Patient12 | Patient12 | 46 | accuracy | 0.722222 |
| classification | lfads | Music | single_day | Patient12 | Patient12 | 54 | accuracy | 0.833333 |
| classification | lfads | Music | single_day | Patient13 | Patient13 | 44 | accuracy | 0.888889 |
| classification | lfads | Music | single_day | Patient13 | Patient13 | 46 | accuracy | 0.777778 |
| classification | lfads | Music | single_day | Patient13 | Patient13 | 54 | accuracy | 0.888889 |
| classification | lfads | Music | single_day | Patient14 | Patient14 | 44 | accuracy | 0.888889 |
| classification | lfads | Music | single_day | Patient14 | Patient14 | 46 | accuracy | 0.888889 |
| classification | lfads | Music | single_day | Patient14 | Patient14 | 54 | accuracy | 0.777778 |
| classification | lfads | Music | single_day | Patient15 | Patient15 | 44 | accuracy | 0.833333 |
| classification | lfads | Music | single_day | Patient15 | Patient15 | 46 | accuracy | 0.833333 |
| classification | lfads | Music | single_day | Patient15 | Patient15 | 54 | accuracy | 0.833333 |
| classification | lfads | Music | single_day | Patient16 | Patient16 | 44 | accuracy | 0.722222 |
| classification | lfads | Music | single_day | Patient16 | Patient16 | 46 | accuracy | 0.666667 |
| classification | lfads | Music | single_day | Patient16 | Patient16 | 54 | accuracy | 0.833333 |
| classification | lfads | Music | single_day | Patient17 | Patient17 | 44 | accuracy | 0.944444 |
| classification | lfads | Music | single_day | Patient17 | Patient17 | 46 | accuracy | 0.888889 |
| classification | lfads | Music | single_day | Patient17 | Patient17 | 54 | accuracy | 0.888889 |
| classification | lfads | Music | single_day | Patient18 | Patient18 | 44 | accuracy | 0.833333 |
| classification | lfads | Music | single_day | Patient18 | Patient18 | 46 | accuracy | 0.833333 |
| classification | lfads | Music | single_day | Patient18 | Patient18 | 54 | accuracy | 0.833333 |
| classification | lfads | Music | single_day | Patient19 | Patient19 | 44 | accuracy | 0.722222 |
| classification | lfads | Music | single_day | Patient19 | Patient19 | 46 | accuracy | 0.833333 |
| classification | lfads | Music | single_day | Patient19 | Patient19 | 54 | accuracy | 0.722222 |
| classification | lfads | Music | single_day | Patient20 | Patient20 | 44 | accuracy | 0.833333 |
| classification | lfads | Music | single_day | Patient20 | Patient20 | 46 | accuracy | 0.833333 |
| classification | lfads | Music | single_day | Patient20 | Patient20 | 54 | accuracy | 0.833333 |
| classification | lfads | Music | single_day | Patient21 | Patient21 | 44 | accuracy | 0.833333 |
| classification | lfads | Music | single_day | Patient21 | Patient21 | 46 | accuracy | 0.833333 |
| classification | lfads | Music | single_day | Patient21 | Patient21 | 54 | accuracy | 0.833333 |
| classification | lfads | Music | single_day | Patient22 | Patient22 | 44 | accuracy | 0.833333 |
| classification | lfads | Music | single_day | Patient22 | Patient22 | 46 | accuracy | 0.833333 |
| classification | lfads | Music | single_day | Patient22 | Patient22 | 54 | accuracy | 0.833333 |
| classification | lfads | Music | single_day | Patient23 | Patient23 | 44 | accuracy | 0.833333 |
| classification | lfads | Music | single_day | Patient23 | Patient23 | 46 | accuracy | 0.833333 |
| classification | lfads | Music | single_day | Patient23 | Patient23 | 54 | accuracy | 0.888889 |
| classification | lfads | Music | single_day | Patient24 | Patient24 | 44 | accuracy | 0.833333 |
| classification | lfads | Music | single_day | Patient24 | Patient24 | 46 | accuracy | 0.833333 |
| classification | lfads | Music | single_day | Patient24 | Patient24 | 54 | accuracy | 0.777778 |
| classification | lfads | Music | single_day | Patient25 | Patient25 | 44 | accuracy | 0.888889 |
| classification | lfads | Music | single_day | Patient25 | Patient25 | 46 | accuracy | 0.777778 |
| classification | lfads | Music | single_day | Patient25 | Patient25 | 54 | accuracy | 0.944444 |
| classification | lfads | Music | single_day | Patient26 | Patient26 | 44 | accuracy | 0.777778 |
| classification | lfads | Music | single_day | Patient26 | Patient26 | 46 | accuracy | 0.833333 |
| classification | lfads | Music | single_day | Patient26 | Patient26 | 54 | accuracy | 0.833333 |
| classification | lfads | Music | single_day | Patient27 | Patient27 | 44 | accuracy | 0.833333 |
| classification | lfads | Music | single_day | Patient27 | Patient27 | 46 | accuracy | 0.777778 |
| classification | lfads | Music | single_day | Patient27 | Patient27 | 54 | accuracy | 0.777778 |
| classification | lfads | Music | single_day | Patient28 | Patient28 | 44 | accuracy | 0.888889 |
| classification | lfads | Music | single_day | Patient28 | Patient28 | 46 | accuracy | 0.944444 |
| classification | lfads | Music | single_day | Patient28 | Patient28 | 54 | accuracy | 0.888889 |
| classification | cycle_gan | Music | single_day | All_RISE_IEEG_preprocessed_final | All_RISE_IEEG_preprocessed_final | 44 | accuracy | 0.820116 |
| classification | cycle_gan | Music | single_day | All_RISE_IEEG_preprocessed_final | All_RISE_IEEG_preprocessed_final | 46 | accuracy | 0.820116 |
| classification | cycle_gan | Music | single_day | All_RISE_IEEG_preprocessed_final | All_RISE_IEEG_preprocessed_final | 54 | accuracy | 0.818182 |
| classification | cycle_gan | Music | single_day | Patient00 | Patient00 | 44 | accuracy | 0.833333 |
| classification | cycle_gan | Music | single_day | Patient00 | Patient00 | 46 | accuracy | 0.833333 |
| classification | cycle_gan | Music | single_day | Patient00 | Patient00 | 54 | accuracy | 0.777778 |
| classification | cycle_gan | Music | single_day | Patient01 | Patient01 | 44 | accuracy | 0.833333 |
| classification | cycle_gan | Music | single_day | Patient01 | Patient01 | 46 | accuracy | 0.777778 |
| classification | cycle_gan | Music | single_day | Patient01 | Patient01 | 54 | accuracy | 0.833333 |
| classification | cycle_gan | Music | single_day | Patient02 | Patient02 | 44 | accuracy | 0.944444 |
| classification | cycle_gan | Music | single_day | Patient02 | Patient02 | 46 | accuracy | 1 |
| classification | cycle_gan | Music | single_day | Patient02 | Patient02 | 54 | accuracy | 0.833333 |
| classification | cycle_gan | Music | single_day | Patient03 | Patient03 | 44 | accuracy | 0.833333 |

| | | | | | | | | |
|---|---|---|---|---|---|---|---|---|
| classification | cycle_gan | Music | single_day | Patient03 | Patient03 | 46 | accuracy | 0.833333 |
| classification | cycle_gan | Music | single_day | Patient03 | Patient03 | 54 | accuracy | 0.833333 |
| classification | cycle_gan | Music | single_day | Patient04 | Patient04 | 44 | accuracy | 0.888889 |
| classification | cycle_gan | Music | single_day | Patient04 | Patient04 | 46 | accuracy | 0.833333 |
| classification | cycle_gan | Music | single_day | Patient04 | Patient04 | 54 | accuracy | 0.833333 |
| classification | cycle_gan | Music | single_day | Patient05 | Patient05 | 44 | accuracy | 0.888889 |
| classification | cycle_gan | Music | single_day | Patient05 | Patient05 | 46 | accuracy | 1 |
| classification | cycle_gan | Music | single_day | Patient05 | Patient05 | 54 | accuracy | 0.833333 |
| classification | cycle_gan | Music | single_day | Patient06 | Patient06 | 44 | accuracy | 0.833333 |
| classification | cycle_gan | Music | single_day | Patient06 | Patient06 | 46 | accuracy | 0.833333 |
| classification | cycle_gan | Music | single_day | Patient06 | Patient06 | 54 | accuracy | 0.833333 |
| classification | cycle_gan | Music | single_day | Patient07 | Patient07 | 44 | accuracy | 0.722222 |
| classification | cycle_gan | Music | single_day | Patient07 | Patient07 | 46 | accuracy | 0.833333 |
| classification | cycle_gan | Music | single_day | Patient07 | Patient07 | 54 | accuracy | 0.722222 |
| classification | cycle_gan | Music | single_day | Patient08 | Patient08 | 44 | accuracy | 0.888889 |
| classification | cycle_gan | Music | single_day | Patient08 | Patient08 | 46 | accuracy | 0.888889 |
| classification | cycle_gan | Music | single_day | Patient08 | Patient08 | 54 | accuracy | 0.777778 |
| classification | cycle_gan | Music | single_day | Patient09 | Patient09 | 44 | accuracy | 0.777778 |
| classification | cycle_gan | Music | single_day | Patient09 | Patient09 | 46 | accuracy | 0.777778 |
| classification | cycle_gan | Music | single_day | Patient09 | Patient09 | 54 | accuracy | 0.777778 |
| classification | cycle_gan | Music | single_day | Patient10 | Patient10 | 44 | accuracy | 0.777778 |
| classification | cycle_gan | Music | single_day | Patient10 | Patient10 | 46 | accuracy | 0.777778 |
| classification | cycle_gan | Music | single_day | Patient10 | Patient10 | 54 | accuracy | 0.833333 |
| classification | cycle_gan | Music | single_day | Patient11 | Patient11 | 44 | accuracy | 0.722222 |
| classification | cycle_gan | Music | single_day | Patient11 | Patient11 | 46 | accuracy | 0.833333 |
| classification | cycle_gan | Music | single_day | Patient11 | Patient11 | 54 | accuracy | 0.833333 |
| classification | cycle_gan | Music | single_day | Patient12 | Patient12 | 44 | accuracy | 0.777778 |
| classification | cycle_gan | Music | single_day | Patient12 | Patient12 | 46 | accuracy | 0.833333 |
| classification | cycle_gan | Music | single_day | Patient12 | Patient12 | 54 | accuracy | 0.833333 |
| classification | cycle_gan | Music | single_day | Patient13 | Patient13 | 44 | accuracy | 0.944444 |
| classification | cycle_gan | Music | single_day | Patient13 | Patient13 | 46 | accuracy | 0.888889 |
| classification | cycle_gan | Music | single_day | Patient13 | Patient13 | 54 | accuracy | 0.777778 |
| classification | cycle_gan | Music | single_day | Patient14 | Patient14 | 44 | accuracy | 0.888889 |
| classification | cycle_gan | Music | single_day | Patient14 | Patient14 | 46 | accuracy | 0.944444 |
| classification | cycle_gan | Music | single_day | Patient14 | Patient14 | 54 | accuracy | 0.833333 |
| classification | cycle_gan | Music | single_day | Patient15 | Patient15 | 44 | accuracy | 0.833333 |
| classification | cycle_gan | Music | single_day | Patient15 | Patient15 | 46 | accuracy | 0.777778 |
| classification | cycle_gan | Music | single_day | Patient15 | Patient15 | 54 | accuracy | 0.833333 |
| classification | cycle_gan | Music | single_day | Patient16 | Patient16 | 44 | accuracy | 0.833333 |
| classification | cycle_gan | Music | single_day | Patient16 | Patient16 | 46 | accuracy | 0.833333 |
| classification | cycle_gan | Music | single_day | Patient16 | Patient16 | 54 | accuracy | 0.833333 |
| classification | cycle_gan | Music | single_day | Patient17 | Patient17 | 44 | accuracy | 0.833333 |
| classification | cycle_gan | Music | single_day | Patient17 | Patient17 | 46 | accuracy | 0.833333 |
| classification | cycle_gan | Music | single_day | Patient17 | Patient17 | 54 | accuracy | 0.777778 |
| classification | cycle_gan | Music | single_day | Patient18 | Patient18 | 44 | accuracy | 0.722222 |
| classification | cycle_gan | Music | single_day | Patient18 | Patient18 | 46 | accuracy | 1 |
| classification | cycle_gan | Music | single_day | Patient18 | Patient18 | 54 | accuracy | 0.833333 |
| classification | cycle_gan | Music | single_day | Patient19 | Patient19 | 44 | accuracy | 0.833333 |
| classification | cycle_gan | Music | single_day | Patient19 | Patient19 | 46 | accuracy | 0.833333 |
| classification | cycle_gan | Music | single_day | Patient19 | Patient19 | 54 | accuracy | 0.888889 |
| classification | cycle_gan | Music | single_day | Patient20 | Patient20 | 44 | accuracy | 0.833333 |
| classification | cycle_gan | Music | single_day | Patient20 | Patient20 | 46 | accuracy | 0.777778 |
| classification | cycle_gan | Music | single_day | Patient20 | Patient20 | 54 | accuracy | 0.666667 |
| classification | cycle_gan | Music | single_day | Patient21 | Patient21 | 44 | accuracy | 0.944444 |
| classification | cycle_gan | Music | single_day | Patient21 | Patient21 | 46 | accuracy | 0.777778 |
| classification | cycle_gan | Music | single_day | Patient21 | Patient21 | 54 | accuracy | 0.833333 |
| classification | cycle_gan | Music | single_day | Patient22 | Patient22 | 44 | accuracy | 0.833333 |
| classification | cycle_gan | Music | single_day | Patient22 | Patient22 | 46 | accuracy | 0.833333 |
| classification | cycle_gan | Music | single_day | Patient22 | Patient22 | 54 | accuracy | 0.833333 |
| classification | cycle_gan | Music | single_day | Patient23 | Patient23 | 44 | accuracy | 0.777778 |
| classification | cycle_gan | Music | single_day | Patient23 | Patient23 | 46 | accuracy | 0.833333 |
| classification | cycle_gan | Music | single_day | Patient23 | Patient23 | 54 | accuracy | 0.777778 |
| classification | cycle_gan | Music | single_day | Patient24 | Patient24 | 44 | accuracy | 0.833333 |

| | | | | | | | | |
|---|---|---|---|---|---|---|---|---|
| classification | cycle_gan | Music | single_day | Patient24 | Patient24 | 46 | accuracy | 1 |
| classification | cycle_gan | Music | single_day | Patient24 | Patient24 | 54 | accuracy | 0.833333 |
| classification | cycle_gan | Music | single_day | Patient25 | Patient25 | 44 | accuracy | 0.888889 |
| classification | cycle_gan | Music | single_day | Patient25 | Patient25 | 46 | accuracy | 0.888889 |
| classification | cycle_gan | Music | single_day | Patient25 | Patient25 | 54 | accuracy | 0.833333 |
| classification | cycle_gan | Music | single_day | Patient26 | Patient26 | 44 | accuracy | 0.777778 |
| classification | cycle_gan | Music | single_day | Patient26 | Patient26 | 46 | accuracy | 0.777778 |
| classification | cycle_gan | Music | single_day | Patient26 | Patient26 | 54 | accuracy | 0.722222 |
| classification | cycle_gan | Music | single_day | Patient27 | Patient27 | 44 | accuracy | 0.833333 |
| classification | cycle_gan | Music | single_day | Patient27 | Patient27 | 46 | accuracy | 0.777778 |
| classification | cycle_gan | Music | single_day | Patient27 | Patient27 | 54 | accuracy | 0.833333 |
| classification | cycle_gan | Music | single_day | Patient28 | Patient28 | 44 | accuracy | 0.944444 |
| classification | cycle_gan | Music | single_day | Patient28 | Patient28 | 46 | accuracy | 0.888889 |
| classification | cycle_gan | Music | single_day | Patient28 | Patient28 | 54 | accuracy | 0.888889 |
| classification | stabilization | Music | single_day | All_RISE_IEEG_preprocessed_final | All_RISE_IEEG_preprocessed_final | 44 | accuracy | 0.810445 |
| classification | stabilization | Music | single_day | All_RISE_IEEG_preprocessed_final | All_RISE_IEEG_preprocessed_final | 46 | accuracy | 0.793037 |
| classification | stabilization | Music | single_day | All_RISE_IEEG_preprocessed_final | All_RISE_IEEG_preprocessed_final | 54 | accuracy | 0.758221 |
| classification | stabilization | Music | single_day | Patient00 | Patient00 | 44 | accuracy | 0.944444 |
| classification | stabilization | Music | single_day | Patient00 | Patient00 | 46 | accuracy | 0.944444 |
| classification | stabilization | Music | single_day | Patient00 | Patient00 | 54 | accuracy | 0.777778 |
| classification | stabilization | Music | single_day | Patient01 | Patient01 | 44 | accuracy | 0.722222 |
| classification | stabilization | Music | single_day | Patient01 | Patient01 | 46 | accuracy | 0.722222 |
| classification | stabilization | Music | single_day | Patient01 | Patient01 | 54 | accuracy | 0.888889 |
| classification | stabilization | Music | single_day | Patient02 | Patient02 | 44 | accuracy | 0.888889 |
| classification | stabilization | Music | single_day | Patient02 | Patient02 | 46 | accuracy | 0.833333 |
| classification | stabilization | Music | single_day | Patient02 | Patient02 | 54 | accuracy | 0.777778 |
| classification | stabilization | Music | single_day | Patient03 | Patient03 | 44 | accuracy | 0.888889 |
| classification | stabilization | Music | single_day | Patient03 | Patient03 | 46 | accuracy | 0.722222 |
| classification | stabilization | Music | single_day | Patient03 | Patient03 | 54 | accuracy | 0.833333 |
| classification | stabilization | Music | single_day | Patient04 | Patient04 | 44 | accuracy | 0.722222 |
| classification | stabilization | Music | single_day | Patient04 | Patient04 | 46 | accuracy | 0.722222 |
| classification | stabilization | Music | single_day | Patient04 | Patient04 | 54 | accuracy | 0.944444 |
| classification | stabilization | Music | single_day | Patient05 | Patient05 | 44 | accuracy | 0.722222 |
| classification | stabilization | Music | single_day | Patient05 | Patient05 | 46 | accuracy | 0.833333 |
| classification | stabilization | Music | single_day | Patient05 | Patient05 | 54 | accuracy | 0.833333 |
| classification | stabilization | Music | single_day | Patient06 | Patient06 | 44 | accuracy | 0.833333 |
| classification | stabilization | Music | single_day | Patient06 | Patient06 | 46 | accuracy | 0.833333 |
| classification | stabilization | Music | single_day | Patient06 | Patient06 | 54 | accuracy | 0.722222 |
| classification | stabilization | Music | single_day | Patient07 | Patient07 | 44 | accuracy | 0.555556 |
| classification | stabilization | Music | single_day | Patient07 | Patient07 | 46 | accuracy | 0.666667 |
| classification | stabilization | Music | single_day | Patient07 | Patient07 | 54 | accuracy | 0.611111 |
| classification | stabilization | Music | single_day | Patient08 | Patient08 | 44 | accuracy | 0.944444 |
| classification | stabilization | Music | single_day | Patient08 | Patient08 | 46 | accuracy | 0.722222 |
| classification | stabilization | Music | single_day | Patient08 | Patient08 | 54 | accuracy | 0.722222 |
| classification | stabilization | Music | single_day | Patient09 | Patient09 | 44 | accuracy | 0.611111 |
| classification | stabilization | Music | single_day | Patient09 | Patient09 | 46 | accuracy | 0.611111 |
| classification | stabilization | Music | single_day | Patient09 | Patient09 | 54 | accuracy | 0.777778 |
| classification | stabilization | Music | single_day | Patient10 | Patient10 | 44 | accuracy | 0.833333 |
| classification | stabilization | Music | single_day | Patient10 | Patient10 | 46 | accuracy | 0.777778 |
| classification | stabilization | Music | single_day | Patient10 | Patient10 | 54 | accuracy | 0.833333 |
| classification | stabilization | Music | single_day | Patient11 | Patient11 | 44 | accuracy | 0.722222 |
| classification | stabilization | Music | single_day | Patient11 | Patient11 | 46 | accuracy | 0.722222 |
| classification | stabilization | Music | single_day | Patient11 | Patient11 | 54 | accuracy | 0.722222 |
| classification | stabilization | Music | single_day | Patient12 | Patient12 | 44 | accuracy | 0.722222 |
| classification | stabilization | Music | single_day | Patient12 | Patient12 | 46 | accuracy | 0.833333 |
| classification | stabilization | Music | single_day | Patient12 | Patient12 | 54 | accuracy | 0.833333 |
| classification | stabilization | Music | single_day | Patient13 | Patient13 | 44 | accuracy | 0.888889 |
| classification | stabilization | Music | single_day | Patient13 | Patient13 | 46 | accuracy | 0.777778 |
| classification | stabilization | Music | single_day | Patient13 | Patient13 | 54 | accuracy | 0.833333 |
| classification | stabilization | Music | single_day | Patient14 | Patient14 | 44 | accuracy | 0.944444 |
| classification | stabilization | Music | single_day | Patient14 | Patient14 | 46 | accuracy | 0.777778 |
| classification | stabilization | Music | single_day | Patient14 | Patient14 | 54 | accuracy | 0.944444 |
| classification | stabilization | Music | single_day | Patient15 | Patient15 | 44 | accuracy | 0.944444 |

| | | | | | | | | |
|---|---|---|---|---|---|---|---|---|
| classification | stabilization | Music | single_day | Patient15 | Patient15 | 46 | accuracy | 1 |
| classification | stabilization | Music | single_day | Patient15 | Patient15 | 54 | accuracy | 0.944444 |
| classification | stabilization | Music | single_day | Patient16 | Patient16 | 44 | accuracy | 0.833333 |
| classification | stabilization | Music | single_day | Patient16 | Patient16 | 46 | accuracy | 0.722222 |
| classification | stabilization | Music | single_day | Patient16 | Patient16 | 54 | accuracy | 0.833333 |
| classification | stabilization | Music | single_day | Patient17 | Patient17 | 44 | accuracy | 0.833333 |
| classification | stabilization | Music | single_day | Patient17 | Patient17 | 46 | accuracy | 0.666667 |
| classification | stabilization | Music | single_day | Patient17 | Patient17 | 54 | accuracy | 0.722222 |
| classification | stabilization | Music | single_day | Patient18 | Patient18 | 44 | accuracy | 1 |
| classification | stabilization | Music | single_day | Patient18 | Patient18 | 46 | accuracy | 0.944444 |
| classification | stabilization | Music | single_day | Patient18 | Patient18 | 54 | accuracy | 0.777778 |
| classification | stabilization | Music | single_day | Patient19 | Patient19 | 44 | accuracy | 0.666667 |
| classification | stabilization | Music | single_day | Patient19 | Patient19 | 46 | accuracy | 0.666667 |
| classification | stabilization | Music | single_day | Patient19 | Patient19 | 54 | accuracy | 0.777778 |
| classification | stabilization | Music | single_day | Patient20 | Patient20 | 44 | accuracy | 0.722222 |
| classification | stabilization | Music | single_day | Patient20 | Patient20 | 46 | accuracy | 0.611111 |
| classification | stabilization | Music | single_day | Patient20 | Patient20 | 54 | accuracy | 0.666667 |
| classification | stabilization | Music | single_day | Patient21 | Patient21 | 44 | accuracy | 0.944444 |
| classification | stabilization | Music | single_day | Patient21 | Patient21 | 46 | accuracy | 0.833333 |
| classification | stabilization | Music | single_day | Patient21 | Patient21 | 54 | accuracy | 0.833333 |
| classification | stabilization | Music | single_day | Patient22 | Patient22 | 44 | accuracy | 0.833333 |
| classification | stabilization | Music | single_day | Patient22 | Patient22 | 46 | accuracy | 0.833333 |
| classification | stabilization | Music | single_day | Patient22 | Patient22 | 54 | accuracy | 0.777778 |
| classification | stabilization | Music | single_day | Patient23 | Patient23 | 44 | accuracy | 0.722222 |
| classification | stabilization | Music | single_day | Patient23 | Patient23 | 46 | accuracy | 0.722222 |
| classification | stabilization | Music | single_day | Patient23 | Patient23 | 54 | accuracy | 0.833333 |
| classification | stabilization | Music | single_day | Patient24 | Patient24 | 44 | accuracy | 0.888889 |
| classification | stabilization | Music | single_day | Patient24 | Patient24 | 46 | accuracy | 0.777778 |
| classification | stabilization | Music | single_day | Patient24 | Patient24 | 54 | accuracy | 0.777778 |
| classification | stabilization | Music | single_day | Patient25 | Patient25 | 44 | accuracy | 0.888889 |
| classification | stabilization | Music | single_day | Patient25 | Patient25 | 46 | accuracy | 0.722222 |
| classification | stabilization | Music | single_day | Patient25 | Patient25 | 54 | accuracy | 0.888889 |
| classification | stabilization | Music | single_day | Patient26 | Patient26 | 44 | accuracy | 0.833333 |
| classification | stabilization | Music | single_day | Patient26 | Patient26 | 46 | accuracy | 0.888889 |
| classification | stabilization | Music | single_day | Patient26 | Patient26 | 54 | accuracy | 0.777778 |
| classification | stabilization | Music | single_day | Patient27 | Patient27 | 44 | accuracy | 0.833333 |
| classification | stabilization | Music | single_day | Patient27 | Patient27 | 46 | accuracy | 0.833333 |
| classification | stabilization | Music | single_day | Patient27 | Patient27 | 54 | accuracy | 0.777778 |
| classification | stabilization | Music | single_day | Patient28 | Patient28 | 44 | accuracy | 0.944444 |
| classification | stabilization | Music | single_day | Patient28 | Patient28 | 46 | accuracy | 0.777778 |
| classification | stabilization | Music | single_day | Patient28 | Patient28 | 54 | accuracy | 0.888889 |
| classification | mfsnn | Music | single_day | Patient00 | Patient00 | 44 | accuracy | 0.833333 |
| classification | mfsnn | Music | single_day | Patient00 | Patient00 | 46 | accuracy | 0.833333 |
| classification | mfsnn | Music | single_day | Patient00 | Patient00 | 54 | accuracy | 0.833333 |
| classification | mfsnn | Music | single_day | Patient01 | Patient01 | 44 | accuracy | 0.833333 |
| classification | mfsnn | Music | single_day | Patient01 | Patient01 | 46 | accuracy | 0.833333 |
| classification | mfsnn | Music | single_day | Patient01 | Patient01 | 54 | accuracy | 0.833333 |
| classification | mfsnn | Music | single_day | Patient02 | Patient02 | 44 | accuracy | 0.833333 |
| classification | mfsnn | Music | single_day | Patient02 | Patient02 | 46 | accuracy | 0.833333 |
| classification | mfsnn | Music | single_day | Patient02 | Patient02 | 54 | accuracy | 0.833333 |
| classification | mfsnn | Music | single_day | Patient03 | Patient03 | 44 | accuracy | 0.833333 |
| classification | mfsnn | Music | single_day | Patient03 | Patient03 | 46 | accuracy | 0.833333 |
| classification | mfsnn | Music | single_day | Patient03 | Patient03 | 54 | accuracy | 0.833333 |
| classification | mfsnn | Music | single_day | Patient04 | Patient04 | 44 | accuracy | 0.833333 |
| classification | mfsnn | Music | single_day | Patient04 | Patient04 | 46 | accuracy | 0.777778 |
| classification | mfsnn | Music | single_day | Patient04 | Patient04 | 54 | accuracy | 0.833333 |
| classification | mfsnn | Music | single_day | Patient05 | Patient05 | 44 | accuracy | 0.944444 |
| classification | mfsnn | Music | single_day | Patient05 | Patient05 | 46 | accuracy | 0.888889 |
| classification | mfsnn | Music | single_day | Patient05 | Patient05 | 54 | accuracy | 0.833333 |
| classification | mfsnn | Music | single_day | Patient06 | Patient06 | 44 | accuracy | 0.944444 |
| classification | mfsnn | Music | single_day | Patient06 | Patient06 | 46 | accuracy | 0.888889 |
| classification | mfsnn | Music | single_day | Patient06 | Patient06 | 54 | accuracy | 0.944444 |
| classification | mfsnn | Music | single_day | Patient07 | Patient07 | 44 | accuracy | 0.833333 |

| | | | | | | | | |
|---|---|---|---|---|---|---|---|---|
| classification | mfsnn | Music | single_day | Patient07 | Patient07 | 46 | accuracy | 0.833333 |
| classification | mfsnn | Music | single_day | Patient07 | Patient07 | 54 | accuracy | 0.833333 |
| classification | mfsnn | Music | single_day | Patient08 | Patient08 | 44 | accuracy | 0.833333 |
| classification | mfsnn | Music | single_day | Patient08 | Patient08 | 46 | accuracy | 0.888889 |
| classification | mfsnn | Music | single_day | Patient08 | Patient08 | 54 | accuracy | 0.888889 |
| classification | mfsnn | Music | single_day | Patient09 | Patient09 | 44 | accuracy | 0.833333 |
| classification | mfsnn | Music | single_day | Patient09 | Patient09 | 46 | accuracy | 0.833333 |
| classification | mfsnn | Music | single_day | Patient09 | Patient09 | 54 | accuracy | 0.833333 |
| classification | mfsnn | Music | single_day | Patient10 | Patient10 | 44 | accuracy | 0.833333 |
| classification | mfsnn | Music | single_day | Patient10 | Patient10 | 46 | accuracy | 0.833333 |
| classification | mfsnn | Music | single_day | Patient10 | Patient10 | 54 | accuracy | 0.833333 |
| classification | mfsnn | Music | single_day | Patient11 | Patient11 | 44 | accuracy | 0.833333 |
| classification | mfsnn | Music | single_day | Patient11 | Patient11 | 46 | accuracy | 0.833333 |
| classification | mfsnn | Music | single_day | Patient11 | Patient11 | 54 | accuracy | 0.833333 |
| classification | mfsnn | Music | single_day | Patient12 | Patient12 | 44 | accuracy | 0.833333 |
| classification | mfsnn | Music | single_day | Patient12 | Patient12 | 46 | accuracy | 0.833333 |
| classification | mfsnn | Music | single_day | Patient12 | Patient12 | 54 | accuracy | 0.833333 |
| classification | mfsnn | Music | single_day | Patient13 | Patient13 | 44 | accuracy | 0.833333 |
| classification | mfsnn | Music | single_day | Patient13 | Patient13 | 46 | accuracy | 0.833333 |
| classification | mfsnn | Music | single_day | Patient13 | Patient13 | 54 | accuracy | 0.833333 |
| classification | mfsnn | Music | single_day | Patient14 | Patient14 | 44 | accuracy | 0.944444 |
| classification | mfsnn | Music | single_day | Patient14 | Patient14 | 46 | accuracy | 0.833333 |
| classification | mfsnn | Music | single_day | Patient14 | Patient14 | 54 | accuracy | 0.833333 |
| classification | mfsnn | Music | single_day | Patient15 | Patient15 | 44 | accuracy | 0.833333 |
| classification | mfsnn | Music | single_day | Patient15 | Patient15 | 46 | accuracy | 0.833333 |
| classification | mfsnn | Music | single_day | Patient15 | Patient15 | 54 | accuracy | 0.888889 |
| classification | mfsnn | Music | single_day | Patient16 | Patient16 | 44 | accuracy | 0.833333 |
| classification | mfsnn | Music | single_day | Patient16 | Patient16 | 46 | accuracy | 0.833333 |
| classification | mfsnn | Music | single_day | Patient16 | Patient16 | 54 | accuracy | 0.888889 |
| classification | mfsnn | Music | single_day | Patient17 | Patient17 | 44 | accuracy | 0.833333 |
| classification | mfsnn | Music | single_day | Patient17 | Patient17 | 46 | accuracy | 0.833333 |
| classification | mfsnn | Music | single_day | Patient17 | Patient17 | 54 | accuracy | 0.833333 |
| classification | mfsnn | Music | single_day | Patient18 | Patient18 | 44 | accuracy | 0.833333 |
| classification | mfsnn | Music | single_day | Patient18 | Patient18 | 46 | accuracy | 0.833333 |
| classification | mfsnn | Music | single_day | Patient18 | Patient18 | 54 | accuracy | 0.833333 |
| classification | mfsnn | Music | single_day | Patient19 | Patient19 | 44 | accuracy | 1 |
| classification | mfsnn | Music | single_day | Patient19 | Patient19 | 46 | accuracy | 0.833333 |
| classification | mfsnn | Music | single_day | Patient19 | Patient19 | 54 | accuracy | 0.833333 |
| classification | mfsnn | Music | single_day | Patient20 | Patient20 | 44 | accuracy | 0.833333 |
| classification | mfsnn | Music | single_day | Patient20 | Patient20 | 46 | accuracy | 0.777778 |
| classification | mfsnn | Music | single_day | Patient20 | Patient20 | 54 | accuracy | 0.833333 |
| classification | mfsnn | Music | single_day | Patient21 | Patient21 | 44 | accuracy | 0.833333 |
| classification | mfsnn | Music | single_day | Patient21 | Patient21 | 46 | accuracy | 0.944444 |
| classification | mfsnn | Music | single_day | Patient21 | Patient21 | 54 | accuracy | 0.888889 |
| classification | mfsnn | Music | single_day | Patient22 | Patient22 | 44 | accuracy | 0.833333 |
| classification | mfsnn | Music | single_day | Patient22 | Patient22 | 46 | accuracy | 0.833333 |
| classification | mfsnn | Music | single_day | Patient22 | Patient22 | 54 | accuracy | 0.833333 |
| classification | mfsnn | Music | single_day | Patient23 | Patient23 | 44 | accuracy | 0.833333 |
| classification | mfsnn | Music | single_day | Patient23 | Patient23 | 46 | accuracy | 0.833333 |
| classification | mfsnn | Music | single_day | Patient23 | Patient23 | 54 | accuracy | 0.833333 |
| classification | mfsnn | Music | single_day | Patient24 | Patient24 | 44 | accuracy | 0.833333 |
| classification | mfsnn | Music | single_day | Patient24 | Patient24 | 46 | accuracy | 0.833333 |
| classification | mfsnn | Music | single_day | Patient24 | Patient24 | 54 | accuracy | 0.888889 |
| classification | mfsnn | Music | single_day | Patient25 | Patient25 | 44 | accuracy | 0.833333 |
| classification | mfsnn | Music | single_day | Patient25 | Patient25 | 46 | accuracy | 0.833333 |
| classification | mfsnn | Music | single_day | Patient25 | Patient25 | 54 | accuracy | 0.777778 |
| classification | mfsnn | Music | single_day | Patient26 | Patient26 | 44 | accuracy | 0.833333 |
| classification | mfsnn | Music | single_day | Patient26 | Patient26 | 46 | accuracy | 0.777778 |
| classification | mfsnn | Music | single_day | Patient26 | Patient26 | 54 | accuracy | 0.833333 |
| classification | mfsnn | Music | single_day | Patient27 | Patient27 | 44 | accuracy | 0.833333 |
| classification | mfsnn | Music | single_day | Patient27 | Patient27 | 46 | accuracy | 0.777778 |
| classification | mfsnn | Music | single_day | Patient27 | Patient27 | 54 | accuracy | 0.888889 |
| classification | mfsnn | Music | single_day | Patient28 | Patient28 | 44 | accuracy | 0.833333 |

| | | | | | | | | |
|---|---|---|---|---|---|---|---|---|
| classification | mfsnn | Music | single_day | Patient28 | Patient28 | 46 | accuracy | 0.944444 |
| classification | mfsnn | Music | single_day | Patient28 | Patient28 | 54 | accuracy | 0.833333 |
| classification | mscformer | Music | single_day | Patient00 | Patient00 | 44 | accuracy | 0.833333 |
| classification | mscformer | Music | single_day | Patient00 | Patient00 | 46 | accuracy | 0.833333 |
| classification | mscformer | Music | single_day | Patient00 | Patient00 | 54 | accuracy | 0.833333 |
| classification | mscformer | Music | single_day | Patient01 | Patient01 | 44 | accuracy | 0.833333 |
| classification | mscformer | Music | single_day | Patient01 | Patient01 | 46 | accuracy | 0.833333 |
| classification | mscformer | Music | single_day | Patient01 | Patient01 | 54 | accuracy | 0.833333 |
| classification | mscformer | Music | single_day | Patient02 | Patient02 | 44 | accuracy | 0.833333 |
| classification | mscformer | Music | single_day | Patient02 | Patient02 | 46 | accuracy | 0.833333 |
| classification | mscformer | Music | single_day | Patient02 | Patient02 | 54 | accuracy | 0.833333 |
| classification | mscformer | Music | single_day | Patient03 | Patient03 | 44 | accuracy | 0.833333 |
| classification | mscformer | Music | single_day | Patient03 | Patient03 | 46 | accuracy | 0.833333 |
| classification | mscformer | Music | single_day | Patient03 | Patient03 | 54 | accuracy | 0.833333 |
| classification | mscformer | Music | single_day | Patient04 | Patient04 | 44 | accuracy | 0.833333 |
| classification | mscformer | Music | single_day | Patient04 | Patient04 | 46 | accuracy | 0.833333 |
| classification | mscformer | Music | single_day | Patient04 | Patient04 | 54 | accuracy | 0.833333 |
| classification | mscformer | Music | single_day | Patient05 | Patient05 | 44 | accuracy | 0.833333 |
| classification | mscformer | Music | single_day | Patient05 | Patient05 | 46 | accuracy | 0.833333 |
| classification | mscformer | Music | single_day | Patient05 | Patient05 | 54 | accuracy | 0.833333 |
| classification | mscformer | Music | single_day | Patient06 | Patient06 | 44 | accuracy | 0.833333 |
| classification | mscformer | Music | single_day | Patient06 | Patient06 | 46 | accuracy | 0.833333 |
| classification | mscformer | Music | single_day | Patient06 | Patient06 | 54 | accuracy | 0.833333 |
| classification | mscformer | Music | single_day | Patient07 | Patient07 | 44 | accuracy | 0.833333 |
| classification | mscformer | Music | single_day | Patient07 | Patient07 | 46 | accuracy | 0.833333 |
| classification | mscformer | Music | single_day | Patient07 | Patient07 | 54 | accuracy | 0.833333 |
| classification | mscformer | Music | single_day | Patient08 | Patient08 | 44 | accuracy | 0.833333 |
| classification | mscformer | Music | single_day | Patient08 | Patient08 | 46 | accuracy | 0.833333 |
| classification | mscformer | Music | single_day | Patient08 | Patient08 | 54 | accuracy | 0.833333 |
| classification | mscformer | Music | single_day | Patient09 | Patient09 | 44 | accuracy | 0.833333 |
| classification | mscformer | Music | single_day | Patient09 | Patient09 | 46 | accuracy | 0.833333 |
| classification | mscformer | Music | single_day | Patient09 | Patient09 | 54 | accuracy | 0.833333 |
| classification | mscformer | Music | single_day | Patient10 | Patient10 | 44 | accuracy | 0.833333 |
| classification | mscformer | Music | single_day | Patient10 | Patient10 | 46 | accuracy | 0.833333 |
| classification | mscformer | Music | single_day | Patient10 | Patient10 | 54 | accuracy | 0.833333 |
| classification | mscformer | Music | single_day | Patient11 | Patient11 | 44 | accuracy | 0.833333 |
| classification | mscformer | Music | single_day | Patient11 | Patient11 | 46 | accuracy | 0.833333 |
| classification | mscformer | Music | single_day | Patient11 | Patient11 | 54 | accuracy | 0.833333 |
| classification | mscformer | Music | single_day | Patient12 | Patient12 | 44 | accuracy | 0.833333 |
| classification | mscformer | Music | single_day | Patient12 | Patient12 | 46 | accuracy | 0.833333 |
| classification | mscformer | Music | single_day | Patient12 | Patient12 | 54 | accuracy | 0.833333 |
| classification | mscformer | Music | single_day | Patient13 | Patient13 | 44 | accuracy | 0.833333 |
| classification | mscformer | Music | single_day | Patient13 | Patient13 | 46 | accuracy | 0.833333 |
| classification | mscformer | Music | single_day | Patient13 | Patient13 | 54 | accuracy | 0.833333 |
| classification | mscformer | Music | single_day | Patient14 | Patient14 | 44 | accuracy | 0.833333 |
| classification | mscformer | Music | single_day | Patient14 | Patient14 | 46 | accuracy | 0.833333 |
| classification | mscformer | Music | single_day | Patient14 | Patient14 | 54 | accuracy | 0.833333 |
| classification | mscformer | Music | single_day | Patient15 | Patient15 | 44 | accuracy | 0.833333 |
| classification | mscformer | Music | single_day | Patient15 | Patient15 | 46 | accuracy | 0.833333 |
| classification | mscformer | Music | single_day | Patient15 | Patient15 | 54 | accuracy | 0.833333 |
| classification | mscformer | Music | single_day | Patient16 | Patient16 | 44 | accuracy | 0.833333 |
| classification | mscformer | Music | single_day | Patient16 | Patient16 | 46 | accuracy | 0.833333 |
| classification | mscformer | Music | single_day | Patient16 | Patient16 | 54 | accuracy | 0.833333 |
| classification | mscformer | Music | single_day | Patient17 | Patient17 | 44 | accuracy | 0.833333 |
| classification | mscformer | Music | single_day | Patient17 | Patient17 | 46 | accuracy | 0.833333 |
| classification | mscformer | Music | single_day | Patient17 | Patient17 | 54 | accuracy | 0.833333 |
| classification | mscformer | Music | single_day | Patient18 | Patient18 | 44 | accuracy | 0.833333 |
| classification | mscformer | Music | single_day | Patient18 | Patient18 | 46 | accuracy | 0.833333 |
| classification | mscformer | Music | single_day | Patient18 | Patient18 | 54 | accuracy | 0.833333 |
| classification | mscformer | Music | single_day | Patient19 | Patient19 | 44 | accuracy | 0.833333 |
| classification | mscformer | Music | single_day | Patient19 | Patient19 | 46 | accuracy | 0.833333 |
| classification | mscformer | Music | single_day | Patient19 | Patient19 | 54 | accuracy | 0.833333 |
| classification | mscformer | Music | single_day | Patient20 | Patient20 | 44 | accuracy | 0.833333 |

| | | | | | | | | |
|---|---|---|---|---|---|---|---|---|
| classification | mscformer | Music | single_day | Patient20 | Patient20 | 46 | accuracy | 0.833333 |
| classification | mscformer | Music | single_day | Patient20 | Patient20 | 54 | accuracy | 0.833333 |
| classification | mscformer | Music | single_day | Patient21 | Patient21 | 44 | accuracy | 0.833333 |
| classification | mscformer | Music | single_day | Patient21 | Patient21 | 46 | accuracy | 0.833333 |
| classification | mscformer | Music | single_day | Patient21 | Patient21 | 54 | accuracy | 0.833333 |
| classification | mscformer | Music | single_day | Patient22 | Patient22 | 44 | accuracy | 0.833333 |
| classification | mscformer | Music | single_day | Patient22 | Patient22 | 46 | accuracy | 0.833333 |
| classification | mscformer | Music | single_day | Patient22 | Patient22 | 54 | accuracy | 0.833333 |
| classification | mscformer | Music | single_day | Patient23 | Patient23 | 44 | accuracy | 0.833333 |
| classification | mscformer | Music | single_day | Patient23 | Patient23 | 46 | accuracy | 0.833333 |
| classification | mscformer | Music | single_day | Patient23 | Patient23 | 54 | accuracy | 0.833333 |
| classification | mscformer | Music | single_day | Patient24 | Patient24 | 44 | accuracy | 0.833333 |
| classification | mscformer | Music | single_day | Patient24 | Patient24 | 46 | accuracy | 0.833333 |
| classification | mscformer | Music | single_day | Patient24 | Patient24 | 54 | accuracy | 0.833333 |
| classification | mscformer | Music | single_day | Patient25 | Patient25 | 44 | accuracy | 0.833333 |
| classification | mscformer | Music | single_day | Patient25 | Patient25 | 46 | accuracy | 0.833333 |
| classification | mscformer | Music | single_day | Patient25 | Patient25 | 54 | accuracy | 0.833333 |
| classification | mscformer | Music | single_day | Patient26 | Patient26 | 44 | accuracy | 0.833333 |
| classification | mscformer | Music | single_day | Patient26 | Patient26 | 46 | accuracy | 0.833333 |
| classification | mscformer | Music | single_day | Patient26 | Patient26 | 54 | accuracy | 0.833333 |
| classification | mscformer | Music | single_day | Patient27 | Patient27 | 44 | accuracy | 0.833333 |
| classification | mscformer | Music | single_day | Patient27 | Patient27 | 46 | accuracy | 0.833333 |
| classification | mscformer | Music | single_day | Patient27 | Patient27 | 54 | accuracy | 0.833333 |
| classification | mscformer | Music | single_day | Patient28 | Patient28 | 44 | accuracy | 0.833333 |
| classification | mscformer | Music | single_day | Patient28 | Patient28 | 46 | accuracy | 0.833333 |
| classification | mscformer | Music | single_day | Patient28 | Patient28 | 54 | accuracy | 0.833333 |
| classification | nomad | Music | single_day | Patient00 | Patient00 | 44 | accuracy | 0.833333 |
| classification | nomad | Music | single_day | Patient00 | Patient00 | 46 | accuracy | 0.833333 |
| classification | nomad | Music | single_day | Patient00 | Patient00 | 54 | accuracy | 0.888889 |
| classification | nomad | Music | single_day | Patient01 | Patient01 | 44 | accuracy | 0.833333 |
| classification | nomad | Music | single_day | Patient01 | Patient01 | 46 | accuracy | 0.777778 |
| classification | nomad | Music | single_day | Patient01 | Patient01 | 54 | accuracy | 0.833333 |
| classification | nomad | Music | single_day | Patient02 | Patient02 | 44 | accuracy | 0.888889 |
| classification | nomad | Music | single_day | Patient02 | Patient02 | 46 | accuracy | 0.944444 |
| classification | nomad | Music | single_day | Patient02 | Patient02 | 54 | accuracy | 0.944444 |
| classification | nomad | Music | single_day | Patient03 | Patient03 | 44 | accuracy | 0.944444 |
| classification | nomad | Music | single_day | Patient03 | Patient03 | 46 | accuracy | 0.833333 |
| classification | nomad | Music | single_day | Patient03 | Patient03 | 54 | accuracy | 0.777778 |
| classification | nomad | Music | single_day | Patient04 | Patient04 | 44 | accuracy | 0.777778 |
| classification | nomad | Music | single_day | Patient04 | Patient04 | 46 | accuracy | 0.722222 |
| classification | nomad | Music | single_day | Patient04 | Patient04 | 54 | accuracy | 0.888889 |
| classification | nomad | Music | single_day | Patient05 | Patient05 | 44 | accuracy | 0.888889 |
| classification | nomad | Music | single_day | Patient05 | Patient05 | 46 | accuracy | 0.944444 |
| classification | nomad | Music | single_day | Patient05 | Patient05 | 54 | accuracy | 0.777778 |
| classification | nomad | Music | single_day | Patient06 | Patient06 | 44 | accuracy | 0.833333 |
| classification | nomad | Music | single_day | Patient06 | Patient06 | 46 | accuracy | 0.944444 |
| classification | nomad | Music | single_day | Patient06 | Patient06 | 54 | accuracy | 0.833333 |
| classification | nomad | Music | single_day | Patient07 | Patient07 | 44 | accuracy | 0.833333 |
| classification | nomad | Music | single_day | Patient07 | Patient07 | 46 | accuracy | 0.833333 |
| classification | nomad | Music | single_day | Patient07 | Patient07 | 54 | accuracy | 0.833333 |
| classification | nomad | Music | single_day | Patient08 | Patient08 | 44 | accuracy | 0.833333 |
| classification | nomad | Music | single_day | Patient08 | Patient08 | 46 | accuracy | 0.666667 |
| classification | nomad | Music | single_day | Patient08 | Patient08 | 54 | accuracy | 0.833333 |
| classification | nomad | Music | single_day | Patient09 | Patient09 | 44 | accuracy | 0.777778 |
| classification | nomad | Music | single_day | Patient09 | Patient09 | 46 | accuracy | 0.833333 |
| classification | nomad | Music | single_day | Patient09 | Patient09 | 54 | accuracy | 0.777778 |
| classification | nomad | Music | single_day | Patient10 | Patient10 | 44 | accuracy | 0.833333 |
| classification | nomad | Music | single_day | Patient10 | Patient10 | 46 | accuracy | 0.833333 |
| classification | nomad | Music | single_day | Patient10 | Patient10 | 54 | accuracy | 0.944444 |
| classification | nomad | Music | single_day | Patient11 | Patient11 | 44 | accuracy | 0.833333 |
| classification | nomad | Music | single_day | Patient11 | Patient11 | 46 | accuracy | 0.666667 |
| classification | nomad | Music | single_day | Patient11 | Patient11 | 54 | accuracy | 0.833333 |
| classification | nomad | Music | single_day | Patient12 | Patient12 | 44 | accuracy | 0.833333 |

| | | | | | | | | |
|---|---|---|---|---|---|---|---|---|
| classification | nomad | Music | single_day | Patient12 | Patient12 | 46 | accuracy | 0.777778 |
| classification | nomad | Music | single_day | Patient12 | Patient12 | 54 | accuracy | 0.722222 |
| classification | nomad | Music | single_day | Patient13 | Patient13 | 44 | accuracy | 0.833333 |
| classification | nomad | Music | single_day | Patient13 | Patient13 | 46 | accuracy | 0.833333 |
| classification | nomad | Music | single_day | Patient13 | Patient13 | 54 | accuracy | 0.5 |
| classification | nomad | Music | single_day | Patient14 | Patient14 | 44 | accuracy | 0.833333 |
| classification | nomad | Music | single_day | Patient14 | Patient14 | 46 | accuracy | 0.833333 |
| classification | nomad | Music | single_day | Patient14 | Patient14 | 54 | accuracy | 0.888889 |
| classification | nomad | Music | single_day | Patient15 | Patient15 | 44 | accuracy | 0.833333 |
| classification | nomad | Music | single_day | Patient15 | Patient15 | 46 | accuracy | 0.833333 |
| classification | nomad | Music | single_day | Patient15 | Patient15 | 54 | accuracy | 0.555556 |
| classification | nomad | Music | single_day | Patient16 | Patient16 | 44 | accuracy | 0.777778 |
| classification | nomad | Music | single_day | Patient16 | Patient16 | 46 | accuracy | 0.666667 |
| classification | nomad | Music | single_day | Patient16 | Patient16 | 54 | accuracy | 0.888889 |
| classification | nomad | Music | single_day | Patient17 | Patient17 | 44 | accuracy | 0.888889 |
| classification | nomad | Music | single_day | Patient17 | Patient17 | 46 | accuracy | 0.888889 |
| classification | nomad | Music | single_day | Patient17 | Patient17 | 54 | accuracy | 0.888889 |
| classification | nomad | Music | single_day | Patient18 | Patient18 | 44 | accuracy | 0.833333 |
| classification | nomad | Music | single_day | Patient18 | Patient18 | 46 | accuracy | 0.833333 |
| classification | nomad | Music | single_day | Patient18 | Patient18 | 54 | accuracy | 0.833333 |
| classification | nomad | Music | single_day | Patient19 | Patient19 | 44 | accuracy | 0.833333 |
| classification | nomad | Music | single_day | Patient19 | Patient19 | 46 | accuracy | 0.611111 |
| classification | nomad | Music | single_day | Patient19 | Patient19 | 54 | accuracy | 0.833333 |
| classification | nomad | Music | single_day | Patient20 | Patient20 | 44 | accuracy | 0.833333 |
| classification | nomad | Music | single_day | Patient20 | Patient20 | 46 | accuracy | 0.722222 |
| classification | nomad | Music | single_day | Patient20 | Patient20 | 54 | accuracy | 0.833333 |
| classification | nomad | Music | single_day | Patient21 | Patient21 | 44 | accuracy | 0.944444 |
| classification | nomad | Music | single_day | Patient21 | Patient21 | 46 | accuracy | 0.833333 |
| classification | nomad | Music | single_day | Patient21 | Patient21 | 54 | accuracy | 0.833333 |
| classification | nomad | Music | single_day | Patient22 | Patient22 | 44 | accuracy | 0.888889 |
| classification | nomad | Music | single_day | Patient22 | Patient22 | 46 | accuracy | 0.833333 |
| classification | nomad | Music | single_day | Patient22 | Patient22 | 54 | accuracy | 0.833333 |
| classification | nomad | Music | single_day | Patient23 | Patient23 | 44 | accuracy | 0.777778 |
| classification | nomad | Music | single_day | Patient23 | Patient23 | 46 | accuracy | 0.833333 |
| classification | nomad | Music | single_day | Patient23 | Patient23 | 54 | accuracy | 0.833333 |
| classification | nomad | Music | single_day | Patient24 | Patient24 | 44 | accuracy | 0.833333 |
| classification | nomad | Music | single_day | Patient24 | Patient24 | 46 | accuracy | 1 |
| classification | nomad | Music | single_day | Patient24 | Patient24 | 54 | accuracy | 0.833333 |
| classification | nomad | Music | single_day | Patient25 | Patient25 | 44 | accuracy | 0.888889 |
| classification | nomad | Music | single_day | Patient25 | Patient25 | 46 | accuracy | 0.888889 |
| classification | nomad | Music | single_day | Patient25 | Patient25 | 54 | accuracy | 0.833333 |
| classification | nomad | Music | single_day | Patient26 | Patient26 | 44 | accuracy | 0.833333 |
| classification | nomad | Music | single_day | Patient26 | Patient26 | 46 | accuracy | 0.777778 |
| classification | nomad | Music | single_day | Patient26 | Patient26 | 54 | accuracy | 0.833333 |
| classification | nomad | Music | single_day | Patient27 | Patient27 | 44 | accuracy | 0.888889 |
| classification | nomad | Music | single_day | Patient27 | Patient27 | 46 | accuracy | 0.833333 |
| classification | nomad | Music | single_day | Patient27 | Patient27 | 54 | accuracy | 0.833333 |
| classification | nomad | Music | single_day | Patient28 | Patient28 | 44 | accuracy | 0.888889 |
| classification | nomad | Music | single_day | Patient28 | Patient28 | 46 | accuracy | 0.944444 |
| classification | nomad | Music | single_day | Patient28 | Patient28 | 54 | accuracy | 0.888889 |
| classification | mlp | Music | single_day | Patient00 | Patient00 | 44 | accuracy | 0.888889 |
| classification | mlp | Music | single_day | Patient00 | Patient00 | 46 | accuracy | 1 |
| classification | mlp | Music | single_day | Patient00 | Patient00 | 54 | accuracy | 0.944444 |
| classification | mlp | Music | single_day | Patient01 | Patient01 | 44 | accuracy | 0.777778 |
| classification | mlp | Music | single_day | Patient01 | Patient01 | 46 | accuracy | 0.944444 |
| classification | mlp | Music | single_day | Patient01 | Patient01 | 54 | accuracy | 0.833333 |
| classification | mlp | Music | single_day | Patient02 | Patient02 | 44 | accuracy | 0.944444 |
| classification | mlp | Music | single_day | Patient02 | Patient02 | 46 | accuracy | 0.833333 |
| classification | mlp | Music | single_day | Patient02 | Patient02 | 54 | accuracy | 0.944444 |
| classification | mlp | Music | single_day | Patient03 | Patient03 | 44 | accuracy | 0.833333 |
| classification | mlp | Music | single_day | Patient03 | Patient03 | 46 | accuracy | 0.833333 |
| classification | mlp | Music | single_day | Patient03 | Patient03 | 54 | accuracy | 0.833333 |
| classification | mlp | Music | single_day | Patient04 | Patient04 | 44 | accuracy | 1 |

| | | | | | | | | |
|---|---|---|---|---|---|---|---|---|
| classification | mlp | Music | single_day | Patient04 | Patient04 | 46 | accuracy | 1 |
| classification | mlp | Music | single_day | Patient04 | Patient04 | 54 | accuracy | 1 |
| classification | mlp | Music | single_day | Patient05 | Patient05 | 44 | accuracy | 0.833333 |
| classification | mlp | Music | single_day | Patient05 | Patient05 | 46 | accuracy | 0.888889 |
| classification | mlp | Music | single_day | Patient05 | Patient05 | 54 | accuracy | 0.888889 |
| classification | mlp | Music | single_day | Patient06 | Patient06 | 44 | accuracy | 0.888889 |
| classification | mlp | Music | single_day | Patient06 | Patient06 | 46 | accuracy | 0.833333 |
| classification | mlp | Music | single_day | Patient06 | Patient06 | 54 | accuracy | 0.888889 |
| classification | mlp | Music | single_day | Patient07 | Patient07 | 44 | accuracy | 0.833333 |
| classification | mlp | Music | single_day | Patient07 | Patient07 | 46 | accuracy | 0.833333 |
| classification | mlp | Music | single_day | Patient07 | Patient07 | 54 | accuracy | 0.833333 |
| classification | mlp | Music | single_day | Patient08 | Patient08 | 44 | accuracy | 0.888889 |
| classification | mlp | Music | single_day | Patient08 | Patient08 | 46 | accuracy | 0.833333 |
| classification | mlp | Music | single_day | Patient08 | Patient08 | 54 | accuracy | 0.833333 |
| classification | mlp | Music | single_day | Patient09 | Patient09 | 44 | accuracy | 0.666667 |
| classification | mlp | Music | single_day | Patient09 | Patient09 | 46 | accuracy | 0.777778 |
| classification | mlp | Music | single_day | Patient09 | Patient09 | 54 | accuracy | 0.833333 |
| classification | mlp | Music | single_day | Patient10 | Patient10 | 44 | accuracy | 0.777778 |
| classification | mlp | Music | single_day | Patient10 | Patient10 | 46 | accuracy | 0.833333 |
| classification | mlp | Music | single_day | Patient10 | Patient10 | 54 | accuracy | 0.888889 |
| classification | mlp | Music | single_day | Patient11 | Patient11 | 44 | accuracy | 0.833333 |
| classification | mlp | Music | single_day | Patient11 | Patient11 | 46 | accuracy | 0.833333 |
| classification | mlp | Music | single_day | Patient11 | Patient11 | 54 | accuracy | 0.833333 |
| classification | mlp | Music | single_day | Patient12 | Patient12 | 44 | accuracy | 0.833333 |
| classification | mlp | Music | single_day | Patient12 | Patient12 | 46 | accuracy | 0.833333 |
| classification | mlp | Music | single_day | Patient12 | Patient12 | 54 | accuracy | 0.833333 |
| classification | mlp | Music | single_day | Patient13 | Patient13 | 44 | accuracy | 0.833333 |
| classification | mlp | Music | single_day | Patient13 | Patient13 | 46 | accuracy | 0.944444 |
| classification | mlp | Music | single_day | Patient13 | Patient13 | 54 | accuracy | 0.833333 |
| classification | mlp | Music | single_day | Patient14 | Patient14 | 44 | accuracy | 0.888889 |
| classification | mlp | Music | single_day | Patient14 | Patient14 | 46 | accuracy | 1 |
| classification | mlp | Music | single_day | Patient14 | Patient14 | 54 | accuracy | 0.833333 |
| classification | mlp | Music | single_day | Patient15 | Patient15 | 44 | accuracy | 0.833333 |
| classification | mlp | Music | single_day | Patient15 | Patient15 | 46 | accuracy | 0.833333 |
| classification | mlp | Music | single_day | Patient15 | Patient15 | 54 | accuracy | 0.888889 |
| classification | mlp | Music | single_day | Patient16 | Patient16 | 44 | accuracy | 0.833333 |
| classification | mlp | Music | single_day | Patient16 | Patient16 | 46 | accuracy | 0.777778 |
| classification | mlp | Music | single_day | Patient16 | Patient16 | 54 | accuracy | 0.777778 |
| classification | mlp | Music | single_day | Patient17 | Patient17 | 44 | accuracy | 0.833333 |
| classification | mlp | Music | single_day | Patient17 | Patient17 | 46 | accuracy | 0.833333 |
| classification | mlp | Music | single_day | Patient17 | Patient17 | 54 | accuracy | 0.944444 |
| classification | mlp | Music | single_day | Patient18 | Patient18 | 44 | accuracy | 0.944444 |
| classification | mlp | Music | single_day | Patient18 | Patient18 | 46 | accuracy | 1 |
| classification | mlp | Music | single_day | Patient18 | Patient18 | 54 | accuracy | 0.888889 |
| classification | mlp | Music | single_day | Patient19 | Patient19 | 44 | accuracy | 0.833333 |
| classification | mlp | Music | single_day | Patient19 | Patient19 | 46 | accuracy | 0.722222 |
| classification | mlp | Music | single_day | Patient19 | Patient19 | 54 | accuracy | 0.833333 |
| classification | mlp | Music | single_day | Patient20 | Patient20 | 44 | accuracy | 0.833333 |
| classification | mlp | Music | single_day | Patient20 | Patient20 | 46 | accuracy | 0.833333 |
| classification | mlp | Music | single_day | Patient20 | Patient20 | 54 | accuracy | 0.833333 |
| classification | mlp | Music | single_day | Patient21 | Patient21 | 44 | accuracy | 0.888889 |
| classification | mlp | Music | single_day | Patient21 | Patient21 | 46 | accuracy | 0.888889 |
| classification | mlp | Music | single_day | Patient21 | Patient21 | 54 | accuracy | 0.777778 |
| classification | mlp | Music | single_day | Patient22 | Patient22 | 44 | accuracy | 0.833333 |
| classification | mlp | Music | single_day | Patient22 | Patient22 | 46 | accuracy | 0.888889 |
| classification | mlp | Music | single_day | Patient22 | Patient22 | 54 | accuracy | 0.833333 |
| classification | mlp | Music | single_day | Patient23 | Patient23 | 44 | accuracy | 0.666667 |
| classification | mlp | Music | single_day | Patient23 | Patient23 | 46 | accuracy | 0.666667 |
| classification | mlp | Music | single_day | Patient23 | Patient23 | 54 | accuracy | 0.833333 |
| classification | mlp | Music | single_day | Patient24 | Patient24 | 44 | accuracy | 0.944444 |
| classification | mlp | Music | single_day | Patient24 | Patient24 | 46 | accuracy | 1 |
| classification | mlp | Music | single_day | Patient24 | Patient24 | 54 | accuracy | 0.944444 |
| classification | mlp | Music | single_day | Patient25 | Patient25 | 44 | accuracy | 0.888889 |

| | | | | | | | | |
|---|---|---|---|---|---|---|---|---|
| classification | mlp | Music | single_day | Patient25 | Patient25 | 46 | accuracy | 0.888889 |
| classification | mlp | Music | single_day | Patient25 | Patient25 | 54 | accuracy | 0.888889 |
| classification | mlp | Music | single_day | Patient26 | Patient26 | 44 | accuracy | 0.777778 |
| classification | mlp | Music | single_day | Patient26 | Patient26 | 46 | accuracy | 0.833333 |
| classification | mlp | Music | single_day | Patient26 | Patient26 | 54 | accuracy | 0.722222 |
| classification | mlp | Music | single_day | Patient27 | Patient27 | 44 | accuracy | 0.833333 |
| classification | mlp | Music | single_day | Patient27 | Patient27 | 46 | accuracy | 0.722222 |
| classification | mlp | Music | single_day | Patient27 | Patient27 | 54 | accuracy | 0.777778 |
| classification | mlp | Music | single_day | Patient28 | Patient28 | 44 | accuracy | 0.944444 |
| classification | mlp | Music | single_day | Patient28 | Patient28 | 46 | accuracy | 0.944444 |
| classification | mlp | Music | single_day | Patient28 | Patient28 | 54 | accuracy | 0.888889 |
| classification | lstm | Music | cross_day | Patient00__to__Patient20 | Patient23 | 44 | accuracy | 0.831461 |
| classification | lstm | Music | cross_day | Patient00__to__Patient20 | Patient23 | 46 | accuracy | 0.808989 |
| classification | lstm | Music | cross_day | Patient00__to__Patient20 | Patient23 | 54 | accuracy | 0.820225 |
| classification | lstm | Music | cross_day | Patient00__to__Patient20 | Patient24 | 44 | accuracy | 0.820225 |
| classification | lstm | Music | cross_day | Patient00__to__Patient20 | Patient24 | 46 | accuracy | 0.820225 |
| classification | lstm | Music | cross_day | Patient00__to__Patient20 | Patient24 | 54 | accuracy | 0.820225 |
| classification | lstm | Music | cross_day | Patient00__to__Patient20 | Patient25 | 44 | accuracy | 0.898876 |
| classification | lstm | Music | cross_day | Patient00__to__Patient20 | Patient25 | 46 | accuracy | 0.876404 |
| classification | lstm | Music | cross_day | Patient00__to__Patient20 | Patient25 | 54 | accuracy | 0.876404 |
| classification | lstm | Music | cross_day | Patient00__to__Patient20 | Patient26 | 44 | accuracy | 0.775281 |
| classification | lstm | Music | cross_day | Patient00__to__Patient20 | Patient26 | 46 | accuracy | 0.741573 |
| classification | lstm | Music | cross_day | Patient00__to__Patient20 | Patient26 | 54 | accuracy | 0.775281 |
| classification | lstm | Music | cross_day | Patient00__to__Patient20 | Patient27 | 44 | accuracy | 0.842697 |
| classification | lstm | Music | cross_day | Patient00__to__Patient20 | Patient27 | 46 | accuracy | 0.797753 |
| classification | lstm | Music | cross_day | Patient00__to__Patient20 | Patient27 | 54 | accuracy | 0.831461 |
| classification | lstm | Music | cross_day | Patient00__to__Patient20 | Patient28 | 44 | accuracy | 0.786517 |
| classification | lstm | Music | cross_day | Patient00__to__Patient20 | Patient28 | 46 | accuracy | 0.775281 |
| classification | lstm | Music | cross_day | Patient00__to__Patient20 | Patient28 | 54 | accuracy | 0.797753 |
| classification | transformer | Music | cross_day | Patient00__to__Patient20 | Patient23 | 44 | accuracy | 0.797753 |
| classification | transformer | Music | cross_day | Patient00__to__Patient20 | Patient23 | 46 | accuracy | 0.797753 |
| classification | transformer | Music | cross_day | Patient00__to__Patient20 | Patient23 | 54 | accuracy | 0.808989 |
| classification | transformer | Music | cross_day | Patient00__to__Patient20 | Patient24 | 44 | accuracy | 0.808989 |
| classification | transformer | Music | cross_day | Patient00__to__Patient20 | Patient24 | 46 | accuracy | 0.820225 |
| classification | transformer | Music | cross_day | Patient00__to__Patient20 | Patient24 | 54 | accuracy | 0.820225 |
| classification | transformer | Music | cross_day | Patient00__to__Patient20 | Patient25 | 44 | accuracy | 0.88764 |
| classification | transformer | Music | cross_day | Patient00__to__Patient20 | Patient25 | 46 | accuracy | 0.88764 |
| classification | transformer | Music | cross_day | Patient00__to__Patient20 | Patient25 | 54 | accuracy | 0.876404 |
| classification | transformer | Music | cross_day | Patient00__to__Patient20 | Patient26 | 44 | accuracy | 0.752809 |
| classification | transformer | Music | cross_day | Patient00__to__Patient20 | Patient26 | 46 | accuracy | 0.764045 |
| classification | transformer | Music | cross_day | Patient00__to__Patient20 | Patient26 | 54 | accuracy | 0.752809 |
| classification | transformer | Music | cross_day | Patient00__to__Patient20 | Patient27 | 44 | accuracy | 0.775281 |
| classification | transformer | Music | cross_day | Patient00__to__Patient20 | Patient27 | 46 | accuracy | 0.808989 |
| classification | transformer | Music | cross_day | Patient00__to__Patient20 | Patient27 | 54 | accuracy | 0.820225 |
| classification | transformer | Music | cross_day | Patient00__to__Patient20 | Patient28 | 44 | accuracy | 0.764045 |
| classification | transformer | Music | cross_day | Patient00__to__Patient20 | Patient28 | 46 | accuracy | 0.775281 |
| classification | transformer | Music | cross_day | Patient00__to__Patient20 | Patient28 | 54 | accuracy | 0.797753 |
| classification | lfads | Music | cross_day | Patient00__to__Patient20 | Patient23 | 44 | accuracy | 0.820225 |
| classification | lfads | Music | cross_day | Patient00__to__Patient20 | Patient23 | 46 | accuracy | 0.820225 |
| classification | lfads | Music | cross_day | Patient00__to__Patient20 | Patient23 | 54 | accuracy | 0.820225 |
| classification | lfads | Music | cross_day | Patient00__to__Patient20 | Patient24 | 44 | accuracy | 0.820225 |
| classification | lfads | Music | cross_day | Patient00__to__Patient20 | Patient24 | 46 | accuracy | 0.820225 |
| classification | lfads | Music | cross_day | Patient00__to__Patient20 | Patient24 | 54 | accuracy | 0.820225 |
| classification | lfads | Music | cross_day | Patient00__to__Patient20 | Patient25 | 44 | accuracy | 0.820225 |
| classification | lfads | Music | cross_day | Patient00__to__Patient20 | Patient25 | 46 | accuracy | 0.820225 |
| classification | lfads | Music | cross_day | Patient00__to__Patient20 | Patient25 | 54 | accuracy | 0.820225 |
| classification | lfads | Music | cross_day | Patient00__to__Patient20 | Patient26 | 44 | accuracy | 0.820225 |
| classification | lfads | Music | cross_day | Patient00__to__Patient20 | Patient26 | 46 | accuracy | 0.820225 |
| classification | lfads | Music | cross_day | Patient00__to__Patient20 | Patient26 | 54 | accuracy | 0.820225 |
| classification | lfads | Music | cross_day | Patient00__to__Patient20 | Patient27 | 44 | accuracy | 0.820225 |
| classification | lfads | Music | cross_day | Patient00__to__Patient20 | Patient27 | 46 | accuracy | 0.820225 |
| classification | lfads | Music | cross_day | Patient00__to__Patient20 | Patient27 | 54 | accuracy | 0.820225 |
| classification | lfads | Music | cross_day | Patient00__to__Patient20 | Patient28 | 44 | accuracy | 0.820225 |

| | | | | | | | | |
|---|---|---|---|---|---|---|---|---|
| classification | lfads | Music | cross_day | Patient00__to__Patient20 | Patient28 | 46 | accuracy | 0.820225 |
| classification | lfads | Music | cross_day | Patient00__to__Patient20 | Patient28 | 54 | accuracy | 0.820225 |
| classification | cycle_gan | Music | cross_day | Patient00__to__Patient20 | Patient23 | 44 | accuracy | 0.790123 |
| classification | cycle_gan | Music | cross_day | Patient00__to__Patient20 | Patient23 | 46 | accuracy | 0.802469 |
| classification | cycle_gan | Music | cross_day | Patient00__to__Patient20 | Patient23 | 54 | accuracy | 0.802469 |
| classification | cycle_gan | Music | cross_day | Patient00__to__Patient20 | Patient24 | 44 | accuracy | 0.802469 |
| classification | cycle_gan | Music | cross_day | Patient00__to__Patient20 | Patient24 | 46 | accuracy | 0.802469 |
| classification | cycle_gan | Music | cross_day | Patient00__to__Patient20 | Patient24 | 54 | accuracy | 0.802469 |
| classification | cycle_gan | Music | cross_day | Patient00__to__Patient20 | Patient25 | 44 | accuracy | 0.802469 |
| classification | cycle_gan | Music | cross_day | Patient00__to__Patient20 | Patient25 | 46 | accuracy | 0.802469 |
| classification | cycle_gan | Music | cross_day | Patient00__to__Patient20 | Patient25 | 54 | accuracy | 0.802469 |
| classification | cycle_gan | Music | cross_day | Patient00__to__Patient20 | Patient26 | 44 | accuracy | 0.777778 |
| classification | cycle_gan | Music | cross_day | Patient00__to__Patient20 | Patient26 | 46 | accuracy | 0.777778 |
| classification | cycle_gan | Music | cross_day | Patient00__to__Patient20 | Patient26 | 54 | accuracy | 0.802469 |
| classification | cycle_gan | Music | cross_day | Patient00__to__Patient20 | Patient27 | 44 | accuracy | 0.790123 |
| classification | cycle_gan | Music | cross_day | Patient00__to__Patient20 | Patient27 | 46 | accuracy | 0.802469 |
| classification | cycle_gan | Music | cross_day | Patient00__to__Patient20 | Patient27 | 54 | accuracy | 0.802469 |
| classification | cycle_gan | Music | cross_day | Patient00__to__Patient20 | Patient28 | 44 | accuracy | 0.802469 |
| classification | cycle_gan | Music | cross_day | Patient00__to__Patient20 | Patient28 | 46 | accuracy | 0.802469 |
| classification | cycle_gan | Music | cross_day | Patient00__to__Patient20 | Patient28 | 54 | accuracy | 0.802469 |
| classification | stabilization | Music | cross_day | Patient00__to__Patient20 | Patient23 | 44 | accuracy | 0.62963 |
| classification | stabilization | Music | cross_day | Patient00__to__Patient20 | Patient23 | 46 | accuracy | 0.580247 |
| classification | stabilization | Music | cross_day | Patient00__to__Patient20 | Patient23 | 54 | accuracy | 0.679012 |
| classification | stabilization | Music | cross_day | Patient00__to__Patient20 | Patient24 | 44 | accuracy | 0.82716 |
| classification | stabilization | Music | cross_day | Patient00__to__Patient20 | Patient24 | 46 | accuracy | 0.888889 |
| classification | stabilization | Music | cross_day | Patient00__to__Patient20 | Patient24 | 54 | accuracy | 0.82716 |
| classification | stabilization | Music | cross_day | Patient00__to__Patient20 | Patient25 | 44 | accuracy | 0.753086 |
| classification | stabilization | Music | cross_day | Patient00__to__Patient20 | Patient25 | 46 | accuracy | 0.790123 |
| classification | stabilization | Music | cross_day | Patient00__to__Patient20 | Patient25 | 54 | accuracy | 0.802469 |
| classification | stabilization | Music | cross_day | Patient00__to__Patient20 | Patient26 | 44 | accuracy | 0.666667 |
| classification | stabilization | Music | cross_day | Patient00__to__Patient20 | Patient26 | 46 | accuracy | 0.679012 |
| classification | stabilization | Music | cross_day | Patient00__to__Patient20 | Patient26 | 54 | accuracy | 0.740741 |
| classification | stabilization | Music | cross_day | Patient00__to__Patient20 | Patient27 | 44 | accuracy | 0.82716 |
| classification | stabilization | Music | cross_day | Patient00__to__Patient20 | Patient27 | 46 | accuracy | 0.790123 |
| classification | stabilization | Music | cross_day | Patient00__to__Patient20 | Patient27 | 54 | accuracy | 0.839506 |
| classification | stabilization | Music | cross_day | Patient00__to__Patient20 | Patient28 | 44 | accuracy | 0.703704 |
| classification | stabilization | Music | cross_day | Patient00__to__Patient20 | Patient28 | 46 | accuracy | 0.679012 |
| classification | stabilization | Music | cross_day | Patient00__to__Patient20 | Patient28 | 54 | accuracy | 0.740741 |
| classification | mfsnn | Music | cross_day | Patient00__to__Patient20 | Patient23 | 44 | accuracy | 0.820225 |
| classification | mfsnn | Music | cross_day | Patient00__to__Patient20 | Patient23 | 46 | accuracy | 0.820225 |
| classification | mfsnn | Music | cross_day | Patient00__to__Patient20 | Patient23 | 54 | accuracy | 0.820225 |
| classification | mfsnn | Music | cross_day | Patient00__to__Patient20 | Patient24 | 44 | accuracy | 0.820225 |
| classification | mfsnn | Music | cross_day | Patient00__to__Patient20 | Patient24 | 46 | accuracy | 0.820225 |
| classification | mfsnn | Music | cross_day | Patient00__to__Patient20 | Patient24 | 54 | accuracy | 0.820225 |
| classification | mfsnn | Music | cross_day | Patient00__to__Patient20 | Patient25 | 44 | accuracy | 0.820225 |
| classification | mfsnn | Music | cross_day | Patient00__to__Patient20 | Patient25 | 46 | accuracy | 0.820225 |
| classification | mfsnn | Music | cross_day | Patient00__to__Patient20 | Patient25 | 54 | accuracy | 0.820225 |
| classification | mfsnn | Music | cross_day | Patient00__to__Patient20 | Patient26 | 44 | accuracy | 0.820225 |
| classification | mfsnn | Music | cross_day | Patient00__to__Patient20 | Patient26 | 46 | accuracy | 0.820225 |
| classification | mfsnn | Music | cross_day | Patient00__to__Patient20 | Patient26 | 54 | accuracy | 0.820225 |
| classification | mfsnn | Music | cross_day | Patient00__to__Patient20 | Patient27 | 44 | accuracy | 0.820225 |
| classification | mfsnn | Music | cross_day | Patient00__to__Patient20 | Patient27 | 46 | accuracy | 0.808989 |
| classification | mfsnn | Music | cross_day | Patient00__to__Patient20 | Patient27 | 54 | accuracy | 0.820225 |
| classification | mfsnn | Music | cross_day | Patient00__to__Patient20 | Patient28 | 44 | accuracy | 0.820225 |
| classification | mfsnn | Music | cross_day | Patient00__to__Patient20 | Patient28 | 46 | accuracy | 0.820225 |
| classification | mfsnn | Music | cross_day | Patient00__to__Patient20 | Patient28 | 54 | accuracy | 0.820225 |
| classification | mscformer | Music | cross_day | Patient00__to__Patient20 | Patient23 | 44 | accuracy | 0.820225 |
| classification | mscformer | Music | cross_day | Patient00__to__Patient20 | Patient23 | 46 | accuracy | 0.820225 |
| classification | mscformer | Music | cross_day | Patient00__to__Patient20 | Patient23 | 54 | accuracy | 0.820225 |
| classification | mscformer | Music | cross_day | Patient00__to__Patient20 | Patient24 | 44 | accuracy | 0.820225 |
| classification | mscformer | Music | cross_day | Patient00__to__Patient20 | Patient24 | 46 | accuracy | 0.820225 |
| classification | mscformer | Music | cross_day | Patient00__to__Patient20 | Patient24 | 54 | accuracy | 0.820225 |
| classification | mscformer | Music | cross_day | Patient00__to__Patient20 | Patient25 | 44 | accuracy | 0.820225 |

| | | | | | | | | |
|---|---|---|---|---|---|---|---|---|
| classification | mscformer | Music | cross_day | Patient00__to__Patient20 | Patient25 | 46 | accuracy | 0.820225 |
| classification | mscformer | Music | cross_day | Patient00__to__Patient20 | Patient25 | 54 | accuracy | 0.820225 |
| classification | mscformer | Music | cross_day | Patient00__to__Patient20 | Patient26 | 44 | accuracy | 0.820225 |
| classification | mscformer | Music | cross_day | Patient00__to__Patient20 | Patient26 | 46 | accuracy | 0.820225 |
| classification | mscformer | Music | cross_day | Patient00__to__Patient20 | Patient26 | 54 | accuracy | 0.820225 |
| classification | mscformer | Music | cross_day | Patient00__to__Patient20 | Patient27 | 44 | accuracy | 0.820225 |
| classification | mscformer | Music | cross_day | Patient00__to__Patient20 | Patient27 | 46 | accuracy | 0.820225 |
| classification | mscformer | Music | cross_day | Patient00__to__Patient20 | Patient27 | 54 | accuracy | 0.820225 |
| classification | mscformer | Music | cross_day | Patient00__to__Patient20 | Patient28 | 44 | accuracy | 0.820225 |
| classification | mscformer | Music | cross_day | Patient00__to__Patient20 | Patient28 | 46 | accuracy | 0.820225 |
| classification | mscformer | Music | cross_day | Patient00__to__Patient20 | Patient28 | 54 | accuracy | 0.820225 |
| classification | nomad | Music | cross_day | Patient00__to__Patient20 | Patient23 | 44 | accuracy | 0.797753 |
| classification | nomad | Music | cross_day | Patient00__to__Patient20 | Patient23 | 46 | accuracy | 0.808989 |
| classification | nomad | Music | cross_day | Patient00__to__Patient20 | Patient23 | 54 | accuracy | 0.842697 |
| classification | nomad | Music | cross_day | Patient00__to__Patient20 | Patient24 | 44 | accuracy | 0.853933 |
| classification | nomad | Music | cross_day | Patient00__to__Patient20 | Patient24 | 46 | accuracy | 0.820225 |
| classification | nomad | Music | cross_day | Patient00__to__Patient20 | Patient24 | 54 | accuracy | 0.842697 |
| classification | nomad | Music | cross_day | Patient00__to__Patient20 | Patient25 | 44 | accuracy | 0.865169 |
| classification | nomad | Music | cross_day | Patient00__to__Patient20 | Patient25 | 46 | accuracy | 0.842697 |
| classification | nomad | Music | cross_day | Patient00__to__Patient20 | Patient25 | 54 | accuracy | 0.865169 |
| classification | nomad | Music | cross_day | Patient00__to__Patient20 | Patient26 | 44 | accuracy | 0.775281 |
| classification | nomad | Music | cross_day | Patient00__to__Patient20 | Patient26 | 46 | accuracy | 0.752809 |
| classification | nomad | Music | cross_day | Patient00__to__Patient20 | Patient26 | 54 | accuracy | 0.730337 |
| classification | nomad | Music | cross_day | Patient00__to__Patient20 | Patient27 | 44 | accuracy | 0.797753 |
| classification | nomad | Music | cross_day | Patient00__to__Patient20 | Patient27 | 46 | accuracy | 0.831461 |
| classification | nomad | Music | cross_day | Patient00__to__Patient20 | Patient27 | 54 | accuracy | 0.797753 |
| classification | nomad | Music | cross_day | Patient00__to__Patient20 | Patient28 | 44 | accuracy | 0.775281 |
| classification | nomad | Music | cross_day | Patient00__to__Patient20 | Patient28 | 46 | accuracy | 0.808989 |
| classification | nomad | Music | cross_day | Patient00__to__Patient20 | Patient28 | 54 | accuracy | 0.741573 |
| classification | mlp | Music | cross_day | Patient00__to__Patient20 | Patient23 | 44 | accuracy | 0.820225 |
| classification | mlp | Music | cross_day | Patient00__to__Patient20 | Patient23 | 46 | accuracy | 0.820225 |
| classification | mlp | Music | cross_day | Patient00__to__Patient20 | Patient23 | 54 | accuracy | 0.820225 |
| classification | mlp | Music | cross_day | Patient00__to__Patient20 | Patient24 | 44 | accuracy | 0.820225 |
| classification | mlp | Music | cross_day | Patient00__to__Patient20 | Patient24 | 46 | accuracy | 0.820225 |
| classification | mlp | Music | cross_day | Patient00__to__Patient20 | Patient24 | 54 | accuracy | 0.820225 |
| classification | mlp | Music | cross_day | Patient00__to__Patient20 | Patient25 | 44 | accuracy | 0.820225 |
| classification | mlp | Music | cross_day | Patient00__to__Patient20 | Patient25 | 46 | accuracy | 0.820225 |
| classification | mlp | Music | cross_day | Patient00__to__Patient20 | Patient25 | 54 | accuracy | 0.820225 |
| classification | mlp | Music | cross_day | Patient00__to__Patient20 | Patient26 | 44 | accuracy | 0.820225 |
| classification | mlp | Music | cross_day | Patient00__to__Patient20 | Patient26 | 46 | accuracy | 0.820225 |
| classification | mlp | Music | cross_day | Patient00__to__Patient20 | Patient26 | 54 | accuracy | 0.820225 |
| classification | mlp | Music | cross_day | Patient00__to__Patient20 | Patient27 | 44 | accuracy | 0.820225 |
| classification | mlp | Music | cross_day | Patient00__to__Patient20 | Patient27 | 46 | accuracy | 0.820225 |
| classification | mlp | Music | cross_day | Patient00__to__Patient20 | Patient27 | 54 | accuracy | 0.820225 |
| classification | mlp | Music | cross_day | Patient00__to__Patient20 | Patient28 | 44 | accuracy | 0.820225 |
| classification | mlp | Music | cross_day | Patient00__to__Patient20 | Patient28 | 46 | accuracy | 0.820225 |
| classification | mlp | Music | cross_day | Patient00__to__Patient20 | Patient28 | 54 | accuracy | 0.820225 |
| classification | lstm | Emotion | single_day | data_20100708 | data_20100708 | 44 | accuracy | 0.416667 |
| classification | lstm | Emotion | single_day | data_20100708 | data_20100708 | 46 | accuracy | 0.4 |
| classification | lstm | Emotion | single_day | data_20100708 | data_20100708 | 54 | accuracy | 0.316667 |
| classification | lstm | Emotion | single_day | data_20100709 | data_20100709 | 44 | accuracy | 0.3 |
| classification | lstm | Emotion | single_day | data_20100709 | data_20100709 | 46 | accuracy | 0.266667 |
| classification | lstm | Emotion | single_day | data_20100709 | data_20100709 | 54 | accuracy | 0.333333 |
| classification | lstm | Emotion | single_day | data_20100807 | data_20100807 | 44 | accuracy | 0.333333 |
| classification | lstm | Emotion | single_day | data_20100807 | data_20100807 | 46 | accuracy | 0.35 |
| classification | lstm | Emotion | single_day | data_20100807 | data_20100807 | 54 | accuracy | 0.266667 |
| classification | lstm | Emotion | single_day | data_20110509 | data_20110509 | 44 | accuracy | 0.3 |
| classification | lstm | Emotion | single_day | data_20110509 | data_20110509 | 46 | accuracy | 0.283333 |
| classification | lstm | Emotion | single_day | data_20110509 | data_20110509 | 54 | accuracy | 0.433333 |
| classification | lstm | Emotion | single_day | data_20110510 | data_20110510 | 44 | accuracy | 0.283333 |
| classification | lstm | Emotion | single_day | data_20110510 | data_20110510 | 46 | accuracy | 0.333333 |
| classification | lstm | Emotion | single_day | data_20110510 | data_20110510 | 54 | accuracy | 0.333333 |
| classification | lstm | Emotion | single_day | data_20110517 | data_20110517 | 44 | accuracy | 0.4 |

| | | | | | | | | |
|---|---|---|---|---|---|---|---|---|
| classification | lstm | Emotion | single_day | data_20110517 | data_20110517 | 46 | accuracy | 0.35 |
| classification | lstm | Emotion | single_day | data_20110517 | data_20110517 | 54 | accuracy | 0.366667 |
| classification | lstm | Emotion | single_day | data_20110704 | data_20110704 | 44 | accuracy | 0.3 |
| classification | lstm | Emotion | single_day | data_20110704 | data_20110704 | 46 | accuracy | 0.3 |
| classification | lstm | Emotion | single_day | data_20110704 | data_20110704 | 54 | accuracy | 0.333333 |
| classification | lstm | Emotion | single_day | data_20110705 | data_20110705 | 44 | accuracy | 0.383333 |
| classification | lstm | Emotion | single_day | data_20110705 | data_20110705 | 46 | accuracy | 0.333333 |
| classification | lstm | Emotion | single_day | data_20110705 | data_20110705 | 54 | accuracy | 0.366667 |
| classification | lstm | Emotion | single_day | data_20110706 | data_20110706 | 44 | accuracy | 0.383333 |
| classification | lstm | Emotion | single_day | data_20110706 | data_20110706 | 46 | accuracy | 0.35 |
| classification | lstm | Emotion | single_day | data_20110706 | data_20110706 | 54 | accuracy | 0.383333 |
| classification | transformer | Emotion | single_day | data_20100708 | data_20100708 | 44 | accuracy | 0.316667 |
| classification | transformer | Emotion | single_day | data_20100708 | data_20100708 | 46 | accuracy | 0.333333 |
| classification | transformer | Emotion | single_day | data_20100708 | data_20100708 | 54 | accuracy | 0.333333 |
| classification | transformer | Emotion | single_day | data_20100709 | data_20100709 | 44 | accuracy | 0.366667 |
| classification | transformer | Emotion | single_day | data_20100709 | data_20100709 | 46 | accuracy | 0.216667 |
| classification | transformer | Emotion | single_day | data_20100709 | data_20100709 | 54 | accuracy | 0.35 |
| classification | transformer | Emotion | single_day | data_20100807 | data_20100807 | 44 | accuracy | 0.35 |
| classification | transformer | Emotion | single_day | data_20100807 | data_20100807 | 46 | accuracy | 0.3 |
| classification | transformer | Emotion | single_day | data_20100807 | data_20100807 | 54 | accuracy | 0.266667 |
| classification | transformer | Emotion | single_day | data_20110509 | data_20110509 | 44 | accuracy | 0.316667 |
| classification | transformer | Emotion | single_day | data_20110509 | data_20110509 | 46 | accuracy | 0.283333 |
| classification | transformer | Emotion | single_day | data_20110509 | data_20110509 | 54 | accuracy | 0.333333 |
| classification | transformer | Emotion | single_day | data_20110510 | data_20110510 | 44 | accuracy | 0.333333 |
| classification | transformer | Emotion | single_day | data_20110510 | data_20110510 | 46 | accuracy | 0.35 |
| classification | transformer | Emotion | single_day | data_20110510 | data_20110510 | 54 | accuracy | 0.366667 |
| classification | transformer | Emotion | single_day | data_20110517 | data_20110517 | 44 | accuracy | 0.416667 |
| classification | transformer | Emotion | single_day | data_20110517 | data_20110517 | 46 | accuracy | 0.45 |
| classification | transformer | Emotion | single_day | data_20110517 | data_20110517 | 54 | accuracy | 0.35 |
| classification | transformer | Emotion | single_day | data_20110704 | data_20110704 | 44 | accuracy | 0.366667 |
| classification | transformer | Emotion | single_day | data_20110704 | data_20110704 | 46 | accuracy | 0.4 |
| classification | transformer | Emotion | single_day | data_20110704 | data_20110704 | 54 | accuracy | 0.316667 |
| classification | transformer | Emotion | single_day | data_20110705 | data_20110705 | 44 | accuracy | 0.35 |
| classification | transformer | Emotion | single_day | data_20110705 | data_20110705 | 46 | accuracy | 0.35 |
| classification | transformer | Emotion | single_day | data_20110705 | data_20110705 | 54 | accuracy | 0.3 |
| classification | transformer | Emotion | single_day | data_20110706 | data_20110706 | 44 | accuracy | 0.4 |
| classification | transformer | Emotion | single_day | data_20110706 | data_20110706 | 46 | accuracy | 0.333333 |
| classification | transformer | Emotion | single_day | data_20110706 | data_20110706 | 54 | accuracy | 0.333333 |
| classification | lfads | Emotion | single_day | data_20100708 | data_20100708 | 44 | accuracy | 0.4 |
| classification | lfads | Emotion | single_day | data_20100708 | data_20100708 | 46 | accuracy | 0.25 |
| classification | lfads | Emotion | single_day | data_20100708 | data_20100708 | 54 | accuracy | 0.3 |
| classification | lfads | Emotion | single_day | data_20100709 | data_20100709 | 44 | accuracy | 0.283333 |
| classification | lfads | Emotion | single_day | data_20100709 | data_20100709 | 46 | accuracy | 0.266667 |
| classification | lfads | Emotion | single_day | data_20100709 | data_20100709 | 54 | accuracy | 0.316667 |
| classification | lfads | Emotion | single_day | data_20100807 | data_20100807 | 44 | accuracy | 0.3 |
| classification | lfads | Emotion | single_day | data_20100807 | data_20100807 | 46 | accuracy | 0.333333 |
| classification | lfads | Emotion | single_day | data_20100807 | data_20100807 | 54 | accuracy | 0.366667 |
| classification | lfads | Emotion | single_day | data_20110509 | data_20110509 | 44 | accuracy | 0.3 |
| classification | lfads | Emotion | single_day | data_20110509 | data_20110509 | 46 | accuracy | 0.35 |
| classification | lfads | Emotion | single_day | data_20110509 | data_20110509 | 54 | accuracy | 0.383333 |
| classification | lfads | Emotion | single_day | data_20110510 | data_20110510 | 44 | accuracy | 0.4 |
| classification | lfads | Emotion | single_day | data_20110510 | data_20110510 | 46 | accuracy | 0.333333 |
| classification | lfads | Emotion | single_day | data_20110510 | data_20110510 | 54 | accuracy | 0.416667 |
| classification | lfads | Emotion | single_day | data_20110517 | data_20110517 | 44 | accuracy | 0.383333 |
| classification | lfads | Emotion | single_day | data_20110517 | data_20110517 | 46 | accuracy | 0.35 |
| classification | lfads | Emotion | single_day | data_20110517 | data_20110517 | 54 | accuracy | 0.3 |
| classification | lfads | Emotion | single_day | data_20110704 | data_20110704 | 44 | accuracy | 0.5 |
| classification | lfads | Emotion | single_day | data_20110704 | data_20110704 | 46 | accuracy | 0.466667 |
| classification | lfads | Emotion | single_day | data_20110704 | data_20110704 | 54 | accuracy | 0.466667 |
| classification | lfads | Emotion | single_day | data_20110705 | data_20110705 | 44 | accuracy | 0.35 |
| classification | lfads | Emotion | single_day | data_20110705 | data_20110705 | 46 | accuracy | 0.433333 |
| classification | lfads | Emotion | single_day | data_20110705 | data_20110705 | 54 | accuracy | 0.3 |
| classification | lfads | Emotion | single_day | data_20110706 | data_20110706 | 44 | accuracy | 0.383333 |

| | | | | | | | | |
|---|---|---|---|---|---|---|---|---|
| classification | lfads | Emotion | single_day | data_20110706 | data_20110706 | 46 | accuracy | 0.35 |
| classification | lfads | Emotion | single_day | data_20110706 | data_20110706 | 54 | accuracy | 0.416667 |
| classification | cycle_gan | Emotion | single_day | data_20100708 | data_20100708 | 44 | accuracy | 0.25 |
| classification | cycle_gan | Emotion | single_day | data_20100708 | data_20100708 | 46 | accuracy | 0.316667 |
| classification | cycle_gan | Emotion | single_day | data_20100708 | data_20100708 | 54 | accuracy | 0.366667 |
| classification | cycle_gan | Emotion | single_day | data_20100709 | data_20100709 | 44 | accuracy | 0.35 |
| classification | cycle_gan | Emotion | single_day | data_20100709 | data_20100709 | 46 | accuracy | 0.316667 |
| classification | cycle_gan | Emotion | single_day | data_20100709 | data_20100709 | 54 | accuracy | 0.366667 |
| classification | cycle_gan | Emotion | single_day | data_20100807 | data_20100807 | 44 | accuracy | 0.383333 |
| classification | cycle_gan | Emotion | single_day | data_20100807 | data_20100807 | 46 | accuracy | 0.35 |
| classification | cycle_gan | Emotion | single_day | data_20100807 | data_20100807 | 54 | accuracy | 0.383333 |
| classification | cycle_gan | Emotion | single_day | data_20110509 | data_20110509 | 44 | accuracy | 0.383333 |
| classification | cycle_gan | Emotion | single_day | data_20110509 | data_20110509 | 46 | accuracy | 0.416667 |
| classification | cycle_gan | Emotion | single_day | data_20110509 | data_20110509 | 54 | accuracy | 0.4 |
| classification | cycle_gan | Emotion | single_day | data_20110510 | data_20110510 | 44 | accuracy | 0.35 |
| classification | cycle_gan | Emotion | single_day | data_20110510 | data_20110510 | 46 | accuracy | 0.416667 |
| classification | cycle_gan | Emotion | single_day | data_20110510 | data_20110510 | 54 | accuracy | 0.35 |
| classification | cycle_gan | Emotion | single_day | data_20110517 | data_20110517 | 44 | accuracy | 0.316667 |
| classification | cycle_gan | Emotion | single_day | data_20110517 | data_20110517 | 46 | accuracy | 0.366667 |
| classification | cycle_gan | Emotion | single_day | data_20110517 | data_20110517 | 54 | accuracy | 0.333333 |
| classification | cycle_gan | Emotion | single_day | data_20110704 | data_20110704 | 44 | accuracy | 0.4 |
| classification | cycle_gan | Emotion | single_day | data_20110704 | data_20110704 | 46 | accuracy | 0.433333 |
| classification | cycle_gan | Emotion | single_day | data_20110704 | data_20110704 | 54 | accuracy | 0.3 |
| classification | cycle_gan | Emotion | single_day | data_20110705 | data_20110705 | 44 | accuracy | 0.4 |
| classification | cycle_gan | Emotion | single_day | data_20110705 | data_20110705 | 46 | accuracy | 0.416667 |
| classification | cycle_gan | Emotion | single_day | data_20110705 | data_20110705 | 54 | accuracy | 0.383333 |
| classification | cycle_gan | Emotion | single_day | data_20110706 | data_20110706 | 44 | accuracy | 0.35 |
| classification | cycle_gan | Emotion | single_day | data_20110706 | data_20110706 | 46 | accuracy | 0.333333 |
| classification | cycle_gan | Emotion | single_day | data_20110706 | data_20110706 | 54 | accuracy | 0.45 |
| classification | stabilization | Emotion | single_day | data_20100708 | data_20100708 | 44 | accuracy | 0.266667 |
| classification | stabilization | Emotion | single_day | data_20100708 | data_20100708 | 46 | accuracy | 0.366667 |
| classification | stabilization | Emotion | single_day | data_20100708 | data_20100708 | 54 | accuracy | 0.4 |
| classification | stabilization | Emotion | single_day | data_20100709 | data_20100709 | 44 | accuracy | 0.366667 |
| classification | stabilization | Emotion | single_day | data_20100709 | data_20100709 | 46 | accuracy | 0.283333 |
| classification | stabilization | Emotion | single_day | data_20100709 | data_20100709 | 54 | accuracy | 0.4 |
| classification | stabilization | Emotion | single_day | data_20100807 | data_20100807 | 44 | accuracy | 0.4 |
| classification | stabilization | Emotion | single_day | data_20100807 | data_20100807 | 46 | accuracy | 0.35 |
| classification | stabilization | Emotion | single_day | data_20100807 | data_20100807 | 54 | accuracy | 0.35 |
| classification | stabilization | Emotion | single_day | data_20110509 | data_20110509 | 44 | accuracy | 0.35 |
| classification | stabilization | Emotion | single_day | data_20110509 | data_20110509 | 46 | accuracy | 0.416667 |
| classification | stabilization | Emotion | single_day | data_20110509 | data_20110509 | 54 | accuracy | 0.366667 |
| classification | stabilization | Emotion | single_day | data_20110510 | data_20110510 | 44 | accuracy | 0.483333 |
| classification | stabilization | Emotion | single_day | data_20110510 | data_20110510 | 46 | accuracy | 0.433333 |
| classification | stabilization | Emotion | single_day | data_20110510 | data_20110510 | 54 | accuracy | 0.366667 |
| classification | stabilization | Emotion | single_day | data_20110517 | data_20110517 | 44 | accuracy | 0.366667 |
| classification | stabilization | Emotion | single_day | data_20110517 | data_20110517 | 46 | accuracy | 0.366667 |
| classification | stabilization | Emotion | single_day | data_20110517 | data_20110517 | 54 | accuracy | 0.383333 |
| classification | stabilization | Emotion | single_day | data_20110704 | data_20110704 | 44 | accuracy | 0.3 |
| classification | stabilization | Emotion | single_day | data_20110704 | data_20110704 | 46 | accuracy | 0.4 |
| classification | stabilization | Emotion | single_day | data_20110704 | data_20110704 | 54 | accuracy | 0.316667 |
| classification | stabilization | Emotion | single_day | data_20110705 | data_20110705 | 44 | accuracy | 0.333333 |
| classification | stabilization | Emotion | single_day | data_20110705 | data_20110705 | 46 | accuracy | 0.416667 |
| classification | stabilization | Emotion | single_day | data_20110705 | data_20110705 | 54 | accuracy | 0.4 |
| classification | stabilization | Emotion | single_day | data_20110706 | data_20110706 | 44 | accuracy | 0.383333 |
| classification | stabilization | Emotion | single_day | data_20110706 | data_20110706 | 46 | accuracy | 0.466667 |
| classification | stabilization | Emotion | single_day | data_20110706 | data_20110706 | 54 | accuracy | 0.416667 |
| classification | mfsnn | Emotion | single_day | data_20100708 | data_20100708 | 44 | accuracy | 0.333333 |
| classification | mfsnn | Emotion | single_day | data_20100708 | data_20100708 | 46 | accuracy | 0.333333 |
| classification | mfsnn | Emotion | single_day | data_20100708 | data_20100708 | 54 | accuracy | 0.333333 |
| classification | mfsnn | Emotion | single_day | data_20100709 | data_20100709 | 44 | accuracy | 0.333333 |
| classification | mfsnn | Emotion | single_day | data_20100709 | data_20100709 | 46 | accuracy | 0.4 |
| classification | mfsnn | Emotion | single_day | data_20100709 | data_20100709 | 54 | accuracy | 0.233333 |
| classification | mfsnn | Emotion | single_day | data_20100807 | data_20100807 | 44 | accuracy | 0.416667 |

| | | | | | | | | |
|---|---|---|---|---|---|---|---|---|
| classification | mfsnn | Emotion | single_day | data_20100807 | data_20100807 | 46 | accuracy | 0.366667 |
| classification | mfsnn | Emotion | single_day | data_20100807 | data_20100807 | 54 | accuracy | 0.283333 |
| classification | mfsnn | Emotion | single_day | data_20110509 | data_20110509 | 44 | accuracy | 0.283333 |
| classification | mfsnn | Emotion | single_day | data_20110509 | data_20110509 | 46 | accuracy | 0.383333 |
| classification | mfsnn | Emotion | single_day | data_20110509 | data_20110509 | 54 | accuracy | 0.333333 |
| classification | mfsnn | Emotion | single_day | data_20110510 | data_20110510 | 44 | accuracy | 0.333333 |
| classification | mfsnn | Emotion | single_day | data_20110510 | data_20110510 | 46 | accuracy | 0.333333 |
| classification | mfsnn | Emotion | single_day | data_20110510 | data_20110510 | 54 | accuracy | 0.316667 |
| classification | mfsnn | Emotion | single_day | data_20110517 | data_20110517 | 44 | accuracy | 0.266667 |
| classification | mfsnn | Emotion | single_day | data_20110517 | data_20110517 | 46 | accuracy | 0.333333 |
| classification | mfsnn | Emotion | single_day | data_20110517 | data_20110517 | 54 | accuracy | 0.333333 |
| classification | mfsnn | Emotion | single_day | data_20110704 | data_20110704 | 44 | accuracy | 0.35 |
| classification | mfsnn | Emotion | single_day | data_20110704 | data_20110704 | 46 | accuracy | 0.333333 |
| classification | mfsnn | Emotion | single_day | data_20110704 | data_20110704 | 54 | accuracy | 0.333333 |
| classification | mfsnn | Emotion | single_day | data_20110705 | data_20110705 | 44 | accuracy | 0.283333 |
| classification | mfsnn | Emotion | single_day | data_20110705 | data_20110705 | 46 | accuracy | 0.333333 |
| classification | mfsnn | Emotion | single_day | data_20110705 | data_20110705 | 54 | accuracy | 0.333333 |
| classification | mfsnn | Emotion | single_day | data_20110706 | data_20110706 | 44 | accuracy | 0.333333 |
| classification | mfsnn | Emotion | single_day | data_20110706 | data_20110706 | 46 | accuracy | 0.333333 |
| classification | mfsnn | Emotion | single_day | data_20110706 | data_20110706 | 54 | accuracy | 0.283333 |
| classification | mscformer | Emotion | single_day | data_20100708 | data_20100708 | 44 | accuracy | 0.366667 |
| classification | mscformer | Emotion | single_day | data_20100708 | data_20100708 | 46 | accuracy | 0.4 |
| classification | mscformer | Emotion | single_day | data_20100708 | data_20100708 | 54 | accuracy | 0.383333 |
| classification | mscformer | Emotion | single_day | data_20100709 | data_20100709 | 44 | accuracy | 0.3 |
| classification | mscformer | Emotion | single_day | data_20100709 | data_20100709 | 46 | accuracy | 0.25 |
| classification | mscformer | Emotion | single_day | data_20100709 | data_20100709 | 54 | accuracy | 0.3 |
| classification | mscformer | Emotion | single_day | data_20100807 | data_20100807 | 44 | accuracy | 0.383333 |
| classification | mscformer | Emotion | single_day | data_20100807 | data_20100807 | 46 | accuracy | 0.35 |
| classification | mscformer | Emotion | single_day | data_20100807 | data_20100807 | 54 | accuracy | 0.216667 |
| classification | mscformer | Emotion | single_day | data_20110509 | data_20110509 | 44 | accuracy | 0.35 |
| classification | mscformer | Emotion | single_day | data_20110509 | data_20110509 | 46 | accuracy | 0.4 |
| classification | mscformer | Emotion | single_day | data_20110509 | data_20110509 | 54 | accuracy | 0.4 |
| classification | mscformer | Emotion | single_day | data_20110510 | data_20110510 | 44 | accuracy | 0.3 |
| classification | mscformer | Emotion | single_day | data_20110510 | data_20110510 | 46 | accuracy | 0.366667 |
| classification | mscformer | Emotion | single_day | data_20110510 | data_20110510 | 54 | accuracy | 0.3 |
| classification | mscformer | Emotion | single_day | data_20110517 | data_20110517 | 44 | accuracy | 0.333333 |
| classification | mscformer | Emotion | single_day | data_20110517 | data_20110517 | 46 | accuracy | 0.5 |
| classification | mscformer | Emotion | single_day | data_20110517 | data_20110517 | 54 | accuracy | 0.4 |
| classification | mscformer | Emotion | single_day | data_20110704 | data_20110704 | 44 | accuracy | 0.366667 |
| classification | mscformer | Emotion | single_day | data_20110704 | data_20110704 | 46 | accuracy | 0.316667 |
| classification | mscformer | Emotion | single_day | data_20110704 | data_20110704 | 54 | accuracy | 0.416667 |
| classification | mscformer | Emotion | single_day | data_20110705 | data_20110705 | 44 | accuracy | 0.433333 |
| classification | mscformer | Emotion | single_day | data_20110705 | data_20110705 | 46 | accuracy | 0.416667 |
| classification | mscformer | Emotion | single_day | data_20110705 | data_20110705 | 54 | accuracy | 0.416667 |
| classification | mscformer | Emotion | single_day | data_20110706 | data_20110706 | 44 | accuracy | 0.383333 |
| classification | mscformer | Emotion | single_day | data_20110706 | data_20110706 | 46 | accuracy | 0.466667 |
| classification | mscformer | Emotion | single_day | data_20110706 | data_20110706 | 54 | accuracy | 0.366667 |
| classification | nomad | Emotion | single_day | data_20100708 | data_20100708 | 44 | accuracy | 0.333333 |
| classification | nomad | Emotion | single_day | data_20100708 | data_20100708 | 46 | accuracy | 0.316667 |
| classification | nomad | Emotion | single_day | data_20100708 | data_20100708 | 54 | accuracy | 0.333333 |
| classification | nomad | Emotion | single_day | data_20100709 | data_20100709 | 44 | accuracy | 0.45 |
| classification | nomad | Emotion | single_day | data_20100709 | data_20100709 | 46 | accuracy | 0.35 |
| classification | nomad | Emotion | single_day | data_20100709 | data_20100709 | 54 | accuracy | 0.35 |
| classification | nomad | Emotion | single_day | data_20100807 | data_20100807 | 44 | accuracy | 0.3 |
| classification | nomad | Emotion | single_day | data_20100807 | data_20100807 | 46 | accuracy | 0.4 |
| classification | nomad | Emotion | single_day | data_20100807 | data_20100807 | 54 | accuracy | 0.4 |
| classification | nomad | Emotion | single_day | data_20110509 | data_20110509 | 44 | accuracy | 0.433333 |
| classification | nomad | Emotion | single_day | data_20110509 | data_20110509 | 46 | accuracy | 0.366667 |
| classification | nomad | Emotion | single_day | data_20110509 | data_20110509 | 54 | accuracy | 0.333333 |
| classification | nomad | Emotion | single_day | data_20110510 | data_20110510 | 44 | accuracy | 0.35 |
| classification | nomad | Emotion | single_day | data_20110510 | data_20110510 | 46 | accuracy | 0.366667 |
| classification | nomad | Emotion | single_day | data_20110510 | data_20110510 | 54 | accuracy | 0.35 |
| classification | nomad | Emotion | single_day | data_20110517 | data_20110517 | 44 | accuracy | 0.35 |

| | | | | | | | | |
|---|---|---|---|---|---|---|---|---|
| classification | nomad | Emotion | single_day | data_20110517 | data_20110517 | 46 | accuracy | 0.333333 |
| classification | nomad | Emotion | single_day | data_20110517 | data_20110517 | 54 | accuracy | 0.316667 |
| classification | nomad | Emotion | single_day | data_20110704 | data_20110704 | 44 | accuracy | 0.266667 |
| classification | nomad | Emotion | single_day | data_20110704 | data_20110704 | 46 | accuracy | 0.366667 |
| classification | nomad | Emotion | single_day | data_20110704 | data_20110704 | 54 | accuracy | 0.3 |
| classification | nomad | Emotion | single_day | data_20110705 | data_20110705 | 44 | accuracy | 0.433333 |
| classification | nomad | Emotion | single_day | data_20110705 | data_20110705 | 46 | accuracy | 0.433333 |
| classification | nomad | Emotion | single_day | data_20110705 | data_20110705 | 54 | accuracy | 0.316667 |
| classification | nomad | Emotion | single_day | data_20110706 | data_20110706 | 44 | accuracy | 0.4 |
| classification | nomad | Emotion | single_day | data_20110706 | data_20110706 | 46 | accuracy | 0.383333 |
| classification | nomad | Emotion | single_day | data_20110706 | data_20110706 | 54 | accuracy | 0.566667 |
| classification | mlp | Emotion | single_day | data_20100708 | data_20100708 | 44 | accuracy | 0.283333 |
| classification | mlp | Emotion | single_day | data_20100708 | data_20100708 | 46 | accuracy | 0.416667 |
| classification | mlp | Emotion | single_day | data_20100708 | data_20100708 | 54 | accuracy | 0.35 |
| classification | mlp | Emotion | single_day | data_20100709 | data_20100709 | 44 | accuracy | 0.316667 |
| classification | mlp | Emotion | single_day | data_20100709 | data_20100709 | 46 | accuracy | 0.316667 |
| classification | mlp | Emotion | single_day | data_20100709 | data_20100709 | 54 | accuracy | 0.316667 |
| classification | mlp | Emotion | single_day | data_20100807 | data_20100807 | 44 | accuracy | 0.4 |
| classification | mlp | Emotion | single_day | data_20100807 | data_20100807 | 46 | accuracy | 0.366667 |
| classification | mlp | Emotion | single_day | data_20100807 | data_20100807 | 54 | accuracy | 0.316667 |
| classification | mlp | Emotion | single_day | data_20110509 | data_20110509 | 44 | accuracy | 0.3 |
| classification | mlp | Emotion | single_day | data_20110509 | data_20110509 | 46 | accuracy | 0.466667 |
| classification | mlp | Emotion | single_day | data_20110509 | data_20110509 | 54 | accuracy | 0.416667 |
| classification | mlp | Emotion | single_day | data_20110510 | data_20110510 | 44 | accuracy | 0.433333 |
| classification | mlp | Emotion | single_day | data_20110510 | data_20110510 | 46 | accuracy | 0.433333 |
| classification | mlp | Emotion | single_day | data_20110510 | data_20110510 | 54 | accuracy | 0.4 |
| classification | mlp | Emotion | single_day | data_20110517 | data_20110517 | 44 | accuracy | 0.416667 |
| classification | mlp | Emotion | single_day | data_20110517 | data_20110517 | 46 | accuracy | 0.45 |
| classification | mlp | Emotion | single_day | data_20110517 | data_20110517 | 54 | accuracy | 0.366667 |
| classification | mlp | Emotion | single_day | data_20110704 | data_20110704 | 44 | accuracy | 0.3 |
| classification | mlp | Emotion | single_day | data_20110704 | data_20110704 | 46 | accuracy | 0.35 |
| classification | mlp | Emotion | single_day | data_20110704 | data_20110704 | 54 | accuracy | 0.333333 |
| classification | mlp | Emotion | single_day | data_20110705 | data_20110705 | 44 | accuracy | 0.416667 |
| classification | mlp | Emotion | single_day | data_20110705 | data_20110705 | 46 | accuracy | 0.433333 |
| classification | mlp | Emotion | single_day | data_20110705 | data_20110705 | 54 | accuracy | 0.433333 |
| classification | mlp | Emotion | single_day | data_20110706 | data_20110706 | 44 | accuracy | 0.5 |
| classification | mlp | Emotion | single_day | data_20110706 | data_20110706 | 46 | accuracy | 0.383333 |
| classification | mlp | Emotion | single_day | data_20110706 | data_20110706 | 54 | accuracy | 0.383333 |
| classification | lstm | Emotion | cross_day | data_20100708__to__data_20110517 | data_20110705__to__data_20110706 | 44 | accuracy | 0.335 |
| classification | lstm | Emotion | cross_day | data_20100708__to__data_20110517 | data_20110705__to__data_20110706 | 46 | accuracy | 0.345 |
| classification | lstm | Emotion | cross_day | data_20100708__to__data_20110517 | data_20110705__to__data_20110706 | 54 | accuracy | 0.371667 |
| classification | transformer | Emotion | cross_day | data_20100708__to__data_20110517 | data_20110705__to__data_20110706 | 44 | accuracy | 0.323333 |
| classification | transformer | Emotion | cross_day | data_20100708__to__data_20110517 | data_20110705__to__data_20110706 | 46 | accuracy | 0.326667 |
| classification | transformer | Emotion | cross_day | data_20100708__to__data_20110517 | data_20110705__to__data_20110706 | 54 | accuracy | 0.351667 |
| classification | lfads | Emotion | cross_day | data_20100708__to__data_20110517 | data_20110705__to__data_20110706 | 44 | accuracy | 0.318333 |
| classification | lfads | Emotion | cross_day | data_20100708__to__data_20110517 | data_20110705__to__data_20110706 | 46 | accuracy | 0.391667 |
| classification | lfads | Emotion | cross_day | data_20100708__to__data_20110517 | data_20110705__to__data_20110706 | 54 | accuracy | 0.338333 |
| classification | cycle_gan | Emotion | cross_day | data_20100708__to__data_20110517 | data_20110705__to__data_20110706 | 44 | accuracy | 0.32963 |
| classification | cycle_gan | Emotion | cross_day | data_20100708__to__data_20110517 | data_20110705__to__data_20110706 | 46 | accuracy | 0.348148 |
| classification | cycle_gan | Emotion | cross_day | data_20100708__to__data_20110517 | data_20110705__to__data_20110706 | 54 | accuracy | 0.364815 |
| classification | stabilization | Emotion | cross_day | data_20100708__to__data_20110517 | data_20110705__to__data_20110706 | 44 | accuracy | 0.37963 |
| classification | stabilization | Emotion | cross_day | data_20100708__to__data_20110517 | data_20110705__to__data_20110706 | 46 | accuracy | 0.368519 |
| classification | stabilization | Emotion | cross_day | data_20100708__to__data_20110517 | data_20110705__to__data_20110706 | 54 | accuracy | 0.385185 |
| classification | mfsnn | Emotion | cross_day | data_20100708__to__data_20110517 | data_20110705 | 44 | accuracy | 0.333333 |
| classification | mfsnn | Emotion | cross_day | data_20100708__to__data_20110517 | data_20110705 | 46 | accuracy | 0.333333 |
| classification | mfsnn | Emotion | cross_day | data_20100708__to__data_20110517 | data_20110705 | 54 | accuracy | 0.333333 |
| classification | mfsnn | Emotion | cross_day | data_20100708__to__data_20110517 | data_20110706 | 44 | accuracy | 0.333333 |
| classification | mfsnn | Emotion | cross_day | data_20100708__to__data_20110517 | data_20110706 | 46 | accuracy | 0.333333 |
| classification | mfsnn | Emotion | cross_day | data_20100708__to__data_20110517 | data_20110706 | 54 | accuracy | 0.333333 |
| classification | mscformer | Emotion | cross_day | data_20100708__to__data_20110517 | data_20110705 | 44 | accuracy | 0.323333 |
| classification | mscformer | Emotion | cross_day | data_20100708__to__data_20110517 | data_20110705 | 46 | accuracy | 0.336667 |
| classification | mscformer | Emotion | cross_day | data_20100708__to__data_20110517 | data_20110705 | 54 | accuracy | 0.35 |
| classification | mscformer | Emotion | cross_day | data_20100708__to__data_20110517 | data_20110706 | 44 | accuracy | 0.38 |

| | | | | | | | | |
|---|---|---|---|---|---|---|---|---|
| classification | mscformer | Emotion | cross_day | data_20100708__to__data_20110517 | data_20110706 | 46 | accuracy | 0.36 |
| classification | mscformer | Emotion | cross_day | data_20100708__to__data_20110517 | data_20110706 | 54 | accuracy | 0.363333 |
| classification | nomad | Emotion | cross_day | data_20100708__to__data_20110517 | data_20110705 | 44 | accuracy | 0.353333 |
| classification | nomad | Emotion | cross_day | data_20100708__to__data_20110517 | data_20110705 | 46 | accuracy | 0.346667 |
| classification | nomad | Emotion | cross_day | data_20100708__to__data_20110517 | data_20110705 | 54 | accuracy | 0.326667 |
| classification | nomad | Emotion | cross_day | data_20100708__to__data_20110517 | data_20110706 | 44 | accuracy | 0.32 |
| classification | nomad | Emotion | cross_day | data_20100708__to__data_20110517 | data_20110706 | 46 | accuracy | 0.36 |
| classification | nomad | Emotion | cross_day | data_20100708__to__data_20110517 | data_20110706 | 54 | accuracy | 0.35 |
| classification | mlp | Emotion | cross_day | data_20100708__to__data_20110517 | data_20110705 | 44 | accuracy | 0.37 |
| classification | mlp | Emotion | cross_day | data_20100708__to__data_20110517 | data_20110705 | 46 | accuracy | 0.33 |
| classification | mlp | Emotion | cross_day | data_20100708__to__data_20110517 | data_20110705 | 54 | accuracy | 0.35 |
| classification | mlp | Emotion | cross_day | data_20100708__to__data_20110517 | data_20110706 | 44 | accuracy | 0.373333 |
| classification | mlp | Emotion | cross_day | data_20100708__to__data_20110517 | data_20110706 | 46 | accuracy | 0.36 |
| classification | mlp | Emotion | cross_day | data_20100708__to__data_20110517 | data_20110706 | 54 | accuracy | 0.376667 |
| classification | lstm | Speech | single_day | day0 | day0 | 44 | cer | 0.841163 |
| classification | lstm | Speech | single_day | day0 | day0 | 44 | cer | 0.508949 |
| classification | lstm | Speech | single_day | day0 | day0 | 46 | cer | 0.686801 |
| classification | lstm | Speech | single_day | day0 | day0 | 46 | cer | 0.493289 |
| classification | lstm | Speech | single_day | day0 | day0 | 54 | cer | 0.8434 |
| classification | lstm | Speech | single_day | day0 | day0 | 54 | cer | 0.525727 |
| classification | lstm | Speech | single_day | day1 | day1 | 44 | cer | 0.863218 |
| classification | lstm | Speech | single_day | day1 | day1 | 44 | cer | 0.442529 |
| classification | lstm | Speech | single_day | day1 | day1 | 46 | cer | 0.682759 |
| classification | lstm | Speech | single_day | day1 | day1 | 46 | cer | 0.44023 |
| classification | lstm | Speech | single_day | day1 | day1 | 54 | cer | 0.768966 |
| classification | lstm | Speech | single_day | day1 | day1 | 54 | cer | 0.436782 |
| classification | lstm | Speech | single_day | day10 | day10 | 44 | cer | 0.527559 |
| classification | lstm | Speech | single_day | day10 | day10 | 44 | cer | 0.430446 |
| classification | lstm | Speech | single_day | day10 | day10 | 46 | cer | 0.736658 |
| classification | lstm | Speech | single_day | day10 | day10 | 46 | cer | 0.447069 |
| classification | lstm | Speech | single_day | day10 | day10 | 54 | cer | 0.673666 |
| classification | lstm | Speech | single_day | day10 | day10 | 54 | cer | 0.463692 |
| classification | lstm | Speech | single_day | day11 | day11 | 44 | cer | 0.621324 |
| classification | lstm | Speech | single_day | day11 | day11 | 44 | cer | 0.393382 |
| classification | lstm | Speech | single_day | day11 | day11 | 46 | cer | 0.732537 |
| classification | lstm | Speech | single_day | day11 | day11 | 46 | cer | 0.384191 |
| classification | lstm | Speech | single_day | day11 | day11 | 54 | cer | 0.442096 |
| classification | lstm | Speech | single_day | day11 | day11 | 54 | cer | 0.392463 |
| classification | lstm | Speech | single_day | day12 | day12 | 44 | cer | 0.465613 |
| classification | lstm | Speech | single_day | day12 | day12 | 44 | cer | 0.420074 |
| classification | lstm | Speech | single_day | day12 | day12 | 46 | cer | 0.517658 |
| classification | lstm | Speech | single_day | day12 | day12 | 46 | cer | 0.427509 |
| classification | lstm | Speech | single_day | day12 | day12 | 54 | cer | 0.578067 |
| classification | lstm | Speech | single_day | day12 | day12 | 54 | cer | 0.407063 |
| classification | lstm | Speech | single_day | day13 | day13 | 44 | cer | 0.533409 |
| classification | lstm | Speech | single_day | day13 | day13 | 44 | cer | 0.381653 |
| classification | lstm | Speech | single_day | day13 | day13 | 46 | cer | 0.482446 |
| classification | lstm | Speech | single_day | day13 | day13 | 46 | cer | 0.382786 |
| classification | lstm | Speech | single_day | day13 | day13 | 54 | cer | 0.592299 |
| classification | lstm | Speech | single_day | day13 | day13 | 54 | cer | 0.364666 |
| classification | lstm | Speech | single_day | day14 | day14 | 44 | cer | 0.620874 |
| classification | lstm | Speech | single_day | day14 | day14 | 44 | cer | 0.397859 |
| classification | lstm | Speech | single_day | day14 | day14 | 46 | cer | 0.438894 |
| classification | lstm | Speech | single_day | day14 | day14 | 46 | cer | 0.388046 |
| classification | lstm | Speech | single_day | day14 | day14 | 54 | cer | 0.863515 |
| classification | lstm | Speech | single_day | day14 | day14 | 54 | cer | 0.380018 |
| classification | lstm | Speech | single_day | day15 | day15 | 44 | cer | 0.440433 |
| classification | lstm | Speech | single_day | day15 | day15 | 44 | cer | 0.40704 |
| classification | lstm | Speech | single_day | day15 | day15 | 46 | cer | 0.694043 |
| classification | lstm | Speech | single_day | day15 | day15 | 46 | cer | 0.354693 |
| classification | lstm | Speech | single_day | day15 | day15 | 54 | cer | 0.638087 |
| classification | lstm | Speech | single_day | day15 | day15 | 54 | cer | 0.368231 |
| classification | lstm | Speech | single_day | day16 | day16 | 44 | cer | 0.730159 |

| | | | | | | | | |
|---|---|---|---|---|---|---|---|---|
| classification | lstm | Speech | single_day | day16 | day16 | 44 | cer | 0.389418 |
| classification | lstm | Speech | single_day | day16 | day16 | 46 | cer | 0.569312 |
| classification | lstm | Speech | single_day | day16 | day16 | 46 | cer | 0.362963 |
| classification | lstm | Speech | single_day | day16 | day16 | 54 | cer | 0.674074 |
| classification | lstm | Speech | single_day | day16 | day16 | 54 | cer | 0.404233 |
| classification | lstm | Speech | single_day | day17 | day17 | 44 | cer | 0.88154 |
| classification | lstm | Speech | single_day | day17 | day17 | 44 | cer | 0.503455 |
| classification | lstm | Speech | single_day | day17 | day17 | 46 | cer | 0.711747 |
| classification | lstm | Speech | single_day | day17 | day17 | 46 | cer | 0.498519 |
| classification | lstm | Speech | single_day | day17 | day17 | 54 | cer | 0.869694 |
| classification | lstm | Speech | single_day | day17 | day17 | 54 | cer | 0.463968 |
| classification | lstm | Speech | single_day | day18 | day18 | 44 | cer | 0.67184 |
| classification | lstm | Speech | single_day | day18 | day18 | 44 | cer | 0.390244 |
| classification | lstm | Speech | single_day | day18 | day18 | 46 | cer | 0.437916 |
| classification | lstm | Speech | single_day | day18 | day18 | 46 | cer | 0.40133 |
| classification | lstm | Speech | single_day | day18 | day18 | 54 | cer | 0.590909 |
| classification | lstm | Speech | single_day | day18 | day18 | 54 | cer | 0.378049 |
| classification | lstm | Speech | single_day | day19 | day19 | 44 | cer | 0.701876 |
| classification | lstm | Speech | single_day | day19 | day19 | 44 | cer | 0.415597 |
| classification | lstm | Speech | single_day | day19 | day19 | 46 | cer | 0.684107 |
| classification | lstm | Speech | single_day | day19 | day19 | 46 | cer | 0.415597 |
| classification | lstm | Speech | single_day | day19 | day19 | 54 | cer | 0.501481 |
| classification | lstm | Speech | single_day | day19 | day19 | 54 | cer | 0.396841 |
| classification | lstm | Speech | single_day | day2 | day2 | 44 | cer | 0.504792 |
| classification | lstm | Speech | single_day | day2 | day2 | 44 | cer | 0.412141 |
| classification | lstm | Speech | single_day | day2 | day2 | 46 | cer | 0.666134 |
| classification | lstm | Speech | single_day | day2 | day2 | 46 | cer | 0.388179 |
| classification | lstm | Speech | single_day | day2 | day2 | 54 | cer | 0.683706 |
| classification | lstm | Speech | single_day | day2 | day2 | 54 | cer | 0.429712 |
| classification | lstm | Speech | single_day | day20 | day20 | 44 | cer | 0.754774 |
| classification | lstm | Speech | single_day | day20 | day20 | 44 | cer | 0.379899 |
| classification | lstm | Speech | single_day | day20 | day20 | 46 | cer | 0.742714 |
| classification | lstm | Speech | single_day | day20 | day20 | 46 | cer | 0.39397 |
| classification | lstm | Speech | single_day | day20 | day20 | 54 | cer | 0.716583 |
| classification | lstm | Speech | single_day | day20 | day20 | 54 | cer | 0.364824 |
| classification | lstm | Speech | single_day | day21 | day21 | 44 | cer | 0.432052 |
| classification | lstm | Speech | single_day | day21 | day21 | 44 | cer | 0.393627 |
| classification | lstm | Speech | single_day | day21 | day21 | 46 | cer | 0.518276 |
| classification | lstm | Speech | single_day | day21 | day21 | 46 | cer | 0.385192 |
| classification | lstm | Speech | single_day | day21 | day21 | 54 | cer | 0.656982 |
| classification | lstm | Speech | single_day | day21 | day21 | 54 | cer | 0.39269 |
| classification | lstm | Speech | single_day | day22 | day22 | 44 | cer | 0.642261 |
| classification | lstm | Speech | single_day | day22 | day22 | 44 | cer | 0.354032 |
| classification | lstm | Speech | single_day | day22 | day22 | 46 | cer | 0.543095 |
| classification | lstm | Speech | single_day | day22 | day22 | 46 | cer | 0.351251 |
| classification | lstm | Speech | single_day | day22 | day22 | 54 | cer | 0.382762 |
| classification | lstm | Speech | single_day | day22 | day22 | 54 | cer | 0.34569 |
| classification | lstm | Speech | single_day | day23 | day23 | 44 | cer | 0.43125 |
| classification | lstm | Speech | single_day | day23 | day23 | 44 | cer | 0.39375 |
| classification | lstm | Speech | single_day | day23 | day23 | 46 | cer | 0.522917 |
| classification | lstm | Speech | single_day | day23 | day23 | 46 | cer | 0.39375 |
| classification | lstm | Speech | single_day | day23 | day23 | 54 | cer | 0.427083 |
| classification | lstm | Speech | single_day | day23 | day23 | 54 | cer | 0.402083 |
| classification | lstm | Speech | single_day | day3 | day3 | 44 | cer | 0.769231 |
| classification | lstm | Speech | single_day | day3 | day3 | 44 | cer | 0.499215 |
| classification | lstm | Speech | single_day | day3 | day3 | 46 | cer | 0.759812 |
| classification | lstm | Speech | single_day | day3 | day3 | 46 | cer | 0.536892 |
| classification | lstm | Speech | single_day | day3 | day3 | 54 | cer | 0.858713 |
| classification | lstm | Speech | single_day | day3 | day3 | 54 | cer | 0.547881 |
| classification | lstm | Speech | single_day | day4 | day4 | 44 | cer | 0.636771 |
| classification | lstm | Speech | single_day | day4 | day4 | 44 | cer | 0.340807 |
| classification | lstm | Speech | single_day | day4 | day4 | 46 | cer | 0.492377 |
| classification | lstm | Speech | single_day | day4 | day4 | 46 | cer | 0.321973 |

| | | | | | | | | |
|---|---|---|---|---|---|---|---|---|
| classification | lstm | Speech | single_day | day4 | day4 | 54 | cer | 0.358744 |
| classification | lstm | Speech | single_day | day5 | day5 | 44 | cer | 0.46516 |
| classification | lstm | Speech | single_day | day5 | day5 | 44 | cer | 0.380414 |
| classification | lstm | Speech | single_day | day5 | day5 | 46 | cer | 0.473635 |
| classification | lstm | Speech | single_day | day5 | day5 | 46 | cer | 0.357815 |
| classification | lstm | Speech | single_day | day5 | day5 | 54 | cer | 0.647834 |
| classification | lstm | Speech | single_day | day5 | day5 | 54 | cer | 0.352166 |
| classification | lstm | Speech | single_day | day6 | day6 | 44 | cer | 0.448439 |
| classification | lstm | Speech | single_day | day6 | day6 | 44 | cer | 0.347209 |
| classification | lstm | Speech | single_day | day6 | day6 | 46 | cer | 0.707663 |
| classification | lstm | Speech | single_day | day6 | day6 | 46 | cer | 0.350047 |
| classification | lstm | Speech | single_day | day6 | day6 | 54 | cer | 0.636708 |
| classification | lstm | Speech | single_day | day6 | day6 | 54 | cer | 0.355724 |
| classification | lstm | Speech | single_day | day7 | day7 | 44 | cer | 0.534759 |
| classification | lstm | Speech | single_day | day7 | day7 | 44 | cer | 0.381462 |
| classification | lstm | Speech | single_day | day7 | day7 | 46 | cer | 0.663102 |
| classification | lstm | Speech | single_day | day7 | day7 | 46 | cer | 0.395722 |
| classification | lstm | Speech | single_day | day7 | day7 | 54 | cer | 0.646168 |
| classification | lstm | Speech | single_day | day7 | day7 | 54 | cer | 0.410873 |
| classification | lstm | Speech | single_day | day8 | day8 | 44 | cer | 0.393625 |
| classification | lstm | Speech | single_day | day8 | day8 | 44 | cer | 0.392032 |
| classification | lstm | Speech | single_day | day8 | day8 | 46 | cer | 0.660558 |
| classification | lstm | Speech | single_day | day8 | day8 | 46 | cer | 0.39761 |
| classification | lstm | Speech | single_day | day8 | day8 | 54 | cer | 0.798406 |
| classification | lstm | Speech | single_day | day8 | day8 | 54 | cer | 0.408765 |
| classification | lstm | Speech | single_day | day9 | day9 | 44 | cer | 0.675833 |
| classification | lstm | Speech | single_day | day9 | day9 | 44 | cer | 0.453333 |
| classification | lstm | Speech | single_day | day9 | day9 | 46 | cer | 0.5325 |
| classification | lstm | Speech | single_day | day9 | day9 | 46 | cer | 0.461667 |
| classification | lstm | Speech | single_day | day9 | day9 | 54 | cer | 0.648333 |
| classification | lstm | Speech | single_day | day9 | day9 | 54 | cer | 0.455833 |
| classification | transformer | Speech | single_day | day0 | day0 | 44 | cer | 1 |
| classification | transformer | Speech | single_day | day0 | day0 | 44 | cer | 0.592841 |
| classification | transformer | Speech | single_day | day0 | day0 | 46 | cer | 1 |
| classification | transformer | Speech | single_day | day0 | day0 | 46 | cer | 0.590604 |
| classification | transformer | Speech | single_day | day0 | day0 | 54 | cer | 0.804251 |
| classification | transformer | Speech | single_day | day0 | day0 | 54 | cer | 0.573826 |
| classification | transformer | Speech | single_day | day1 | day1 | 44 | cer | 0.968966 |
| classification | transformer | Speech | single_day | day1 | day1 | 44 | cer | 0.536782 |
| classification | transformer | Speech | single_day | day1 | day1 | 46 | cer | 1 |
| classification | transformer | Speech | single_day | day1 | day1 | 46 | cer | 0.547126 |
| classification | transformer | Speech | single_day | day1 | day1 | 54 | cer | 0.793103 |
| classification | transformer | Speech | single_day | day1 | day1 | 54 | cer | 0.698851 |
| classification | transformer | Speech | single_day | day10 | day10 | 44 | cer | 0.895013 |
| classification | transformer | Speech | single_day | day10 | day10 | 44 | cer | 0.565179 |
| classification | transformer | Speech | single_day | day10 | day10 | 46 | cer | 1 |
| classification | transformer | Speech | single_day | day10 | day10 | 46 | cer | 0.553806 |
| classification | transformer | Speech | single_day | day10 | day10 | 54 | cer | 0.83902 |
| classification | transformer | Speech | single_day | day10 | day10 | 54 | cer | 0.548556 |
| classification | transformer | Speech | single_day | day11 | day11 | 44 | cer | 0.835478 |
| classification | transformer | Speech | single_day | day11 | day11 | 44 | cer | 0.516544 |
| classification | transformer | Speech | single_day | day11 | day11 | 46 | cer | 0.795956 |
| classification | transformer | Speech | single_day | day11 | day11 | 46 | cer | 0.5 |
| classification | transformer | Speech | single_day | day11 | day11 | 54 | cer | 0.831801 |
| classification | transformer | Speech | single_day | day11 | day11 | 54 | cer | 0.513787 |
| classification | transformer | Speech | single_day | day12 | day12 | 44 | cer | 1 |
| classification | transformer | Speech | single_day | day12 | day12 | 44 | cer | 0.54368 |
| classification | transformer | Speech | single_day | day12 | day12 | 46 | cer | 1 |
| classification | transformer | Speech | single_day | day12 | day12 | 46 | cer | 0.532528 |
| classification | transformer | Speech | single_day | day12 | day12 | 54 | cer | 1 |
| classification | transformer | Speech | single_day | day12 | day12 | 54 | cer | 0.541822 |
| classification | transformer | Speech | single_day | day13 | day13 | 44 | cer | 0.851642 |
| classification | transformer | Speech | single_day | day13 | day13 | 44 | cer | 0.513024 |

| | | | | | | | | |
|---|---|---|---|---|---|---|---|---|
| classification | transformer | Speech | single_day | day13 | day13 | 46 | cer | 1 |
| classification | transformer | Speech | single_day | day13 | day13 | 46 | cer | 0.621744 |
| classification | transformer | Speech | single_day | day13 | day13 | 54 | cer | 1 |
| classification | transformer | Speech | single_day | day13 | day13 | 54 | cer | 0.687429 |
| classification | transformer | Speech | single_day | day14 | day14 | 44 | cer | 0.813559 |
| classification | transformer | Speech | single_day | day14 | day14 | 44 | cer | 0.491525 |
| classification | transformer | Speech | single_day | day14 | day14 | 46 | cer | 0.821588 |
| classification | transformer | Speech | single_day | day14 | day14 | 46 | cer | 0.505798 |
| classification | transformer | Speech | single_day | day14 | day14 | 54 | cer | 0.752899 |
| classification | transformer | Speech | single_day | day14 | day14 | 54 | cer | 0.504014 |
| classification | transformer | Speech | single_day | day15 | day15 | 44 | cer | 0.851083 |
| classification | transformer | Speech | single_day | day15 | day15 | 44 | cer | 0.480144 |
| classification | transformer | Speech | single_day | day15 | day15 | 46 | cer | 0.774368 |
| classification | transformer | Speech | single_day | day15 | day15 | 46 | cer | 0.544224 |
| classification | transformer | Speech | single_day | day15 | day15 | 54 | cer | 0.755415 |
| classification | transformer | Speech | single_day | day15 | day15 | 54 | cer | 0.481949 |
| classification | transformer | Speech | single_day | day16 | day16 | 44 | cer | 0.731217 |
| classification | transformer | Speech | single_day | day16 | day16 | 44 | cer | 0.49418 |
| classification | transformer | Speech | single_day | day16 | day16 | 46 | cer | 0.797884 |
| classification | transformer | Speech | single_day | day16 | day16 | 46 | cer | 0.496296 |
| classification | transformer | Speech | single_day | day16 | day16 | 54 | cer | 0.756614 |
| classification | transformer | Speech | single_day | day16 | day16 | 54 | cer | 0.488889 |
| classification | transformer | Speech | single_day | day17 | day17 | 44 | cer | 1 |
| classification | transformer | Speech | single_day | day17 | day17 | 44 | cer | 0.562685 |
| classification | transformer | Speech | single_day | day17 | day17 | 46 | cer | 1 |
| classification | transformer | Speech | single_day | day17 | day17 | 46 | cer | 0.631787 |
| classification | transformer | Speech | single_day | day17 | day17 | 54 | cer | 0.766041 |
| classification | transformer | Speech | single_day | day17 | day17 | 54 | cer | 0.553801 |
| classification | transformer | Speech | single_day | day18 | day18 | 44 | cer | 0.797118 |
| classification | transformer | Speech | single_day | day18 | day18 | 44 | cer | 0.51663 |
| classification | transformer | Speech | single_day | day18 | day18 | 46 | cer | 1 |
| classification | transformer | Speech | single_day | day18 | day18 | 46 | cer | 0.529933 |
| classification | transformer | Speech | single_day | day18 | day18 | 54 | cer | 1 |
| classification | transformer | Speech | single_day | day18 | day18 | 54 | cer | 0.513304 |
| classification | transformer | Speech | single_day | day19 | day19 | 44 | cer | 1 |
| classification | transformer | Speech | single_day | day19 | day19 | 44 | cer | 0.52616 |
| classification | transformer | Speech | single_day | day19 | day19 | 46 | cer | 1 |
| classification | transformer | Speech | single_day | day19 | day19 | 46 | cer | 0.527147 |
| classification | transformer | Speech | single_day | day19 | day19 | 54 | cer | 0.82231 |
| classification | transformer | Speech | single_day | day19 | day19 | 54 | cer | 0.532083 |
| classification | transformer | Speech | single_day | day2 | day2 | 44 | cer | 0.763578 |
| classification | transformer | Speech | single_day | day2 | day2 | 44 | cer | 0.538339 |
| classification | transformer | Speech | single_day | day2 | day2 | 46 | cer | 0.771565 |
| classification | transformer | Speech | single_day | day2 | day2 | 46 | cer | 0.535144 |
| classification | transformer | Speech | single_day | day2 | day2 | 54 | cer | 0.720447 |
| classification | transformer | Speech | single_day | day2 | day2 | 54 | cer | 0.533546 |
| classification | transformer | Speech | single_day | day20 | day20 | 44 | cer | 0.773869 |
| classification | transformer | Speech | single_day | day20 | day20 | 44 | cer | 0.478392 |
| classification | transformer | Speech | single_day | day20 | day20 | 46 | cer | 1 |
| classification | transformer | Speech | single_day | day20 | day20 | 46 | cer | 0.497487 |
| classification | transformer | Speech | single_day | day20 | day20 | 54 | cer | 0.747739 |
| classification | transformer | Speech | single_day | day20 | day20 | 54 | cer | 0.526633 |
| classification | transformer | Speech | single_day | day21 | day21 | 44 | cer | 0.923149 |
| classification | transformer | Speech | single_day | day21 | day21 | 44 | cer | 0.488285 |
| classification | transformer | Speech | single_day | day21 | day21 | 46 | cer | 0.790066 |
| classification | transformer | Speech | single_day | day21 | day21 | 46 | cer | 0.49672 |
| classification | transformer | Speech | single_day | day21 | day21 | 54 | cer | 0.869728 |
| classification | transformer | Speech | single_day | day21 | day21 | 54 | cer | 0.522962 |
| classification | transformer | Speech | single_day | day22 | day22 | 44 | cer | 0.756256 |
| classification | transformer | Speech | single_day | day22 | day22 | 44 | cer | 0.506024 |
| classification | transformer | Speech | single_day | day22 | day22 | 46 | cer | 0.759963 |
| classification | transformer | Speech | single_day | day22 | day22 | 46 | cer | 0.516219 |
| classification | transformer | Speech | single_day | day22 | day22 | 54 | cer | 0.759963 |

| | | | | | | | | |
|---|---|---|---|---|---|---|---|---|
| classification | transformer | Speech | single_day | day22 | day22 | 54 | cer | 0.504171 |
| classification | transformer | Speech | single_day | day23 | day23 | 44 | cer | 0.919792 |
| classification | transformer | Speech | single_day | day23 | day23 | 44 | cer | 0.516667 |
| classification | transformer | Speech | single_day | day23 | day23 | 46 | cer | 0.771875 |
| classification | transformer | Speech | single_day | day23 | day23 | 46 | cer | 0.55625 |
| classification | transformer | Speech | single_day | day23 | day23 | 54 | cer | 0.880208 |
| classification | transformer | Speech | single_day | day23 | day23 | 54 | cer | 0.529167 |
| classification | transformer | Speech | single_day | day3 | day3 | 44 | cer | 1 |
| classification | transformer | Speech | single_day | day3 | day3 | 44 | cer | 0.598116 |
| classification | transformer | Speech | single_day | day3 | day3 | 46 | cer | 1 |
| classification | transformer | Speech | single_day | day3 | day3 | 46 | cer | 0.601256 |
| classification | transformer | Speech | single_day | day3 | day3 | 54 | cer | 0.847724 |
| classification | transformer | Speech | single_day | day3 | day3 | 54 | cer | 0.621664 |
| classification | transformer | Speech | single_day | day4 | day4 | 44 | cer | 1 |
| classification | transformer | Speech | single_day | day4 | day4 | 44 | cer | 0.483408 |
| classification | transformer | Speech | single_day | day4 | day4 | 46 | cer | 1 |
| classification | transformer | Speech | single_day | day4 | day4 | 46 | cer | 0.527354 |
| classification | transformer | Speech | single_day | day4 | day4 | 54 | cer | 0.756951 |
| classification | transformer | Speech | single_day | day4 | day4 | 54 | cer | 0.492377 |
| classification | transformer | Speech | single_day | day5 | day5 | 44 | cer | 1 |
| classification | transformer | Speech | single_day | day5 | day5 | 44 | cer | 0.512241 |
| classification | transformer | Speech | single_day | day5 | day5 | 46 | cer | 1 |
| classification | transformer | Speech | single_day | day5 | day5 | 46 | cer | 0.519774 |
| classification | transformer | Speech | single_day | day5 | day5 | 54 | cer | 0.815443 |
| classification | transformer | Speech | single_day | day5 | day5 | 54 | cer | 0.481168 |
| classification | transformer | Speech | single_day | day6 | day6 | 44 | cer | 1 |
| classification | transformer | Speech | single_day | day6 | day6 | 44 | cer | 0.48439 |
| classification | transformer | Speech | single_day | day6 | day6 | 46 | cer | 0.823084 |
| classification | transformer | Speech | single_day | day6 | day6 | 46 | cer | 0.491012 |
| classification | transformer | Speech | single_day | day6 | day6 | 54 | cer | 0.937559 |
| classification | transformer | Speech | single_day | day6 | day6 | 54 | cer | 0.495743 |
| classification | transformer | Speech | single_day | day7 | day7 | 44 | cer | 1 |
| classification | transformer | Speech | single_day | day7 | day7 | 44 | cer | 0.496435 |
| classification | transformer | Speech | single_day | day7 | day7 | 46 | cer | 1 |
| classification | transformer | Speech | single_day | day7 | day7 | 46 | cer | 0.500891 |
| classification | transformer | Speech | single_day | day7 | day7 | 54 | cer | 0.911765 |
| classification | transformer | Speech | single_day | day7 | day7 | 54 | cer | 0.555258 |
| classification | transformer | Speech | single_day | day8 | day8 | 44 | cer | 0.772908 |
| classification | transformer | Speech | single_day | day8 | day8 | 44 | cer | 0.515538 |
| classification | transformer | Speech | single_day | day8 | day8 | 46 | cer | 1 |
| classification | transformer | Speech | single_day | day8 | day8 | 46 | cer | 0.522709 |
| classification | transformer | Speech | single_day | day8 | day8 | 54 | cer | 0.803984 |
| classification | transformer | Speech | single_day | day8 | day8 | 54 | cer | 0.494024 |
| classification | transformer | Speech | single_day | day9 | day9 | 44 | cer | 1 |
| classification | transformer | Speech | single_day | day9 | day9 | 44 | cer | 0.583333 |
| classification | transformer | Speech | single_day | day9 | day9 | 46 | cer | 0.8525 |
| classification | transformer | Speech | single_day | day9 | day9 | 46 | cer | 0.546667 |
| classification | transformer | Speech | single_day | day9 | day9 | 54 | cer | 0.863333 |
| classification | transformer | Speech | single_day | day9 | day9 | 54 | cer | 0.555 |
| classification | lstm | Speech | cross_day | day0__to__day18 | day19 | 44 | cer | 0.292201 |
| classification | lstm | Speech | cross_day | day0__to__day18 | day19 | 46 | cer | 0.281343 |
| classification | lstm | Speech | cross_day | day0__to__day18 | day19 | 54 | cer | 0.28233 |
| classification | lstm | Speech | cross_day | day0__to__day18 | day20 | 44 | cer | 0.268342 |
| classification | lstm | Speech | cross_day | day0__to__day18 | day20 | 46 | cer | 0.268342 |
| classification | lstm | Speech | cross_day | day0__to__day18 | day20 | 54 | cer | 0.268342 |
| classification | lstm | Speech | cross_day | day0__to__day18 | day21 | 44 | cer | 0.367385 |
| classification | lstm | Speech | cross_day | day0__to__day18 | day21 | 46 | cer | 0.367385 |
| classification | lstm | Speech | cross_day | day0__to__day18 | day21 | 54 | cer | 0.367385 |
| classification | lstm | Speech | cross_day | day0__to__day18 | day22 | 44 | cer | 0.386469 |
| classification | lstm | Speech | cross_day | day0__to__day18 | day22 | 46 | cer | 0.386469 |
| classification | lstm | Speech | cross_day | day0__to__day18 | day22 | 54 | cer | 0.386469 |
| classification | lstm | Speech | cross_day | day0__to__day18 | day23 | 44 | cer | 0.451042 |
| classification | lstm | Speech | cross_day | day0__to__day18 | day23 | 46 | cer | 0.451042 |

| | | | | | | | | |
|---|---|---|---|---|---|---|---|---|
| classification | lstm | Speech | cross_day | day0__to__day18 | day23 | 54 | cer | 0.451042 |
| classification | transformer | Speech | cross_day | day0__to__day18 | day19 | 44 | cer | 0.350444 |
| classification | transformer | Speech | cross_day | day0__to__day18 | day19 | 46 | cer | 0.35538 |
| classification | transformer | Speech | cross_day | day0__to__day18 | day19 | 54 | cer | 0.333662 |
| classification | transformer | Speech | cross_day | day0__to__day18 | day20 | 44 | cer | 0.308543 |
| classification | transformer | Speech | cross_day | day0__to__day18 | day20 | 46 | cer | 0.308543 |
| classification | transformer | Speech | cross_day | day0__to__day18 | day20 | 54 | cer | 0.308543 |
| classification | transformer | Speech | cross_day | day0__to__day18 | day21 | 44 | cer | 0.406748 |
| classification | transformer | Speech | cross_day | day0__to__day18 | day21 | 46 | cer | 0.406748 |
| classification | transformer | Speech | cross_day | day0__to__day18 | day21 | 54 | cer | 0.406748 |
| classification | transformer | Speech | cross_day | day0__to__day18 | day22 | 44 | cer | 0.445783 |
| classification | transformer | Speech | cross_day | day0__to__day18 | day22 | 46 | cer | 0.445783 |
| classification | transformer | Speech | cross_day | day0__to__day18 | day22 | 54 | cer | 0.445783 |
| classification | transformer | Speech | cross_day | day0__to__day18 | day23 | 44 | cer | 0.470833 |
| classification | transformer | Speech | cross_day | day0__to__day18 | day23 | 46 | cer | 0.470833 |
| classification | transformer | Speech | cross_day | day0__to__day18 | day23 | 54 | cer | 0.470833 |
