## Supplementary File 4 for "BCIJelly: An integrated ecosystem for brain–computer interface research"

| Task type | Algorithm | Dataset | Protocol | Training day | Test day | Seed | Metric | Score |
|---|---|---|---|---|---|---|---|---|
| regression | LSTM | FALCON M2 | single_day | 20201019 | 20201019 | 44 | r2 | 0.758371 |
| regression | LSTM | FALCON M2 | single_day | 20201019 | 20201019 | 44 | r2 | 0.750483 |
| regression | LSTM | FALCON M2 | single_day | 20201020 | 20201020 | 44 | r2 | 0.730236 |
| regression | LSTM | FALCON M2 | single_day | 20201020 | 20201020 | 44 | r2 | 0.758144 |
| regression | LSTM | FALCON M2 | single_day | 20201027 | 20201027 | 44 | r2 | 0.664683 |
| regression | LSTM | FALCON M2 | single_day | 20201030 | 20201030 | 44 | r2 | 0.710555 |
| regression | LSTM | FALCON M2 | single_day | 20201030 | 20201030 | 44 | r2 | 0.760956 |
| regression | LSTM | FALCON M2 | single_day | 20201118 | 20201118 | 44 | r2 | 0.655249 |
| regression | LSTM | FALCON M2 | single_day | 20201119 | 20201119 | 44 | r2 | 0.783061 |
| regression | LSTM | FALCON M2 | single_day | 20201124 | 20201124 | 44 | r2 | 0.707259 |
| regression | Transformer | FALCON M2 | single_day | 20201019 | 20201019 | 44 | r2 | 0.535024 |
| regression | Transformer | FALCON M2 | single_day | 20201019 | 20201019 | 44 | r2 | 0.532577 |
| regression | Transformer | FALCON M2 | single_day | 20201020 | 20201020 | 44 | r2 | 0.50946 |
| regression | Transformer | FALCON M2 | single_day | 20201020 | 20201020 | 44 | r2 | 0.538972 |
| regression | Transformer | FALCON M2 | single_day | 20201027 | 20201027 | 44 | r2 | 0.229077 |
| regression | Transformer | FALCON M2 | single_day | 20201027 | 20201027 | 44 | r2 | 0.501931 |
| regression | LFADS | FALCON M2 | single_day | 20201019 | 20201019 | 44 | r2 | 0.42879 |
| regression | LFADS | FALCON M2 | single_day | 20201019 | 20201019 | 44 | r2 | 0.447598 |
| regression | LFADS | FALCON M2 | single_day | 20201020 | 20201020 | 44 | r2 | 0.424644 |
| regression | LFADS | FALCON M2 | single_day | 20201020 | 20201020 | 44 | r2 | 0.408737 |
| regression | LFADS | FALCON M2 | single_day | 20201027 | 20201027 | 44 | r2 | 0.273813 |
| regression | LFADS | FALCON M2 | single_day | 20201027 | 20201027 | 44 | r2 | 0.365454 |
| regression | Cycle-GAN | FALCON M2 | single_day | 20201019 | 20201019 | 44 | r2 | 0.290035 |
| regression | Cycle-GAN | FALCON M2 | single_day | 20201019 | 20201019 | 44 | r2 | 0.259282 |
| regression | Cycle-GAN | FALCON M2 | single_day | 20201020 | 20201020 | 44 | r2 | 0.203882 |
| regression | Cycle-GAN | FALCON M2 | single_day | 20201020 | 20201020 | 44 | r2 | 0.201545 |
| regression | Cycle-GAN | FALCON M2 | single_day | 20201027 | 20201027 | 44 | r2 | -0.122948 |
| regression | Cycle-GAN | FALCON M2 | single_day | 20201027 | 20201027 | 44 | r2 | -0.296844 |
| regression | Stabilization | FALCON M2 | single_day | 20201019 | 20201019 | 44 | r2 | 0.210147 |
| regression | Stabilization | FALCON M2 | single_day | 20201019 | 20201019 | 44 | r2 | 0.264311 |
| regression | Stabilization | FALCON M2 | single_day | 20201020 | 20201020 | 44 | r2 | 0.20785 |
| regression | Stabilization | FALCON M2 | single_day | 20201020 | 20201020 | 44 | r2 | 0.258683 |
| regression | Stabilization | FALCON M2 | single_day | 20201027 | 20201027 | 44 | r2 | 0.109725 |
| regression | Stabilization | FALCON M2 | single_day | 20201027 | 20201027 | 44 | r2 | 0.202096 |
| regression | FENet | FALCON M2 | single_day | 20201019 | 20201019 | 44 | r2 | 0.395487 |
| regression | FENet | FALCON M2 | single_day | 20201019 | 20201019 | 44 | r2 | 0.346515 |
| regression | FENet | FALCON M2 | single_day | 20201020 | 20201020 | 44 | r2 | 0.388516 |
| regression | FENet | FALCON M2 | single_day | 20201020 | 20201020 | 44 | r2 | 0.377513 |
| regression | FENet | FALCON M2 | single_day | 20201027 | 20201027 | 44 | r2 | 0.405597 |
| regression | FENet | FALCON M2 | single_day | 20201027 | 20201027 | 44 | r2 | 0.441776 |
| regression | DFINE | FALCON M2 | single_day | 20201019 | 20201019 | 44 | r2 | 0.5165 |
| regression | DFINE | FALCON M2 | single_day | 20201019 | 20201019 | 44 | r2 | 0.499371 |
| regression | DFINE | FALCON M2 | single_day | 20201020 | 20201020 | 44 | r2 | 0.460262 |
| regression | DFINE | FALCON M2 | single_day | 20201020 | 20201020 | 44 | r2 | 0.469553 |
| regression | DFINE | FALCON M2 | single_day | 20201027 | 20201027 | 44 | r2 | 0.407212 |
| regression | DFINE | FALCON M2 | single_day | 20201030 | 20201030 | 44 | r2 | 0.434586 |
| regression | DFINE | FALCON M2 | single_day | 20201030 | 20201030 | 44 | r2 | 0.674921 |
| regression | DFINE | FALCON M2 | single_day | 20201118 | 20201118 | 44 | r2 | 0.483162 |
| regression | DFINE | FALCON M2 | single_day | 20201119 | 20201119 | 44 | r2 | 0.614019 |
| regression | DFINE | FALCON M2 | single_day | 20201124 | 20201124 | 44 | r2 | 0.529197 |
| regression | LSTM | Jango | single_day | 20150730 | 20150730 | 44 | r2 | 0.831824 |
| regression | LSTM | Jango | single_day | 20150801 | 20150801 | 44 | r2 | 0.938998 |
| regression | LSTM | Jango | single_day | 20150806 | 20150806 | 44 | r2 | 0.947855 |
| regression | LSTM | Jango | single_day | 20150808 | 20150808 | 44 | r2 | 0.915552 |
| regression | LSTM | Jango | single_day | 20150820 | 20150820 | 44 | r2 | 0.946348 |
| regression | LSTM | Jango | single_day | 20150825 | 20150825 | 44 | r2 | 0.928595 |
| regression | LSTM | Jango | single_day | 20150827 | 20150827 | 44 | r2 | 0.943488 |
| regression | LSTM | Jango | single_day | 20150831 | 20150831 | 44 | r2 | 0.940727 |
| regression | LSTM | Jango | single_day | 20150906 | 20150906 | 44 | r2 | 0.939265 |
| regression | LSTM | Jango | single_day | 20151102 | 20151102 | 44 | r2 | 0.920463 |

| | | | | | | | | |
|---|---|---|---|---|---|---|---|---|
| regression | Transformer | Jango | single_day | 20150730 | 20150730 | 44 | r2 | 0.777132 |
| regression | Transformer | Jango | single_day | 20150801 | 20150801 | 44 | r2 | 0.83916 |
| regression | Transformer | Jango | single_day | 20150806 | 20150806 | 44 | r2 | 0.895105 |
| regression | Transformer | Jango | single_day | 20150808 | 20150808 | 44 | r2 | 0.861165 |
| regression | Transformer | Jango | single_day | 20150820 | 20150820 | 44 | r2 | 0.851774 |
| regression | Transformer | Jango | single_day | 20150825 | 20150825 | 44 | r2 | 0.875632 |
| regression | Transformer | Jango | single_day | 20150827 | 20150827 | 44 | r2 | 0.878876 |
| regression | Transformer | Jango | single_day | 20150831 | 20150831 | 44 | r2 | 0.87368 |
| regression | Transformer | Jango | single_day | 20150906 | 20150906 | 44 | r2 | 0.869825 |
| regression | Transformer | Jango | single_day | 20151102 | 20151102 | 44 | r2 | 0.874794 |
| regression | LFADS | Jango | single_day | 20150730 | 20150730 | 44 | r2 | 0.567337 |
| regression | LFADS | Jango | single_day | 20150801 | 20150801 | 44 | r2 | 0.935768 |
| regression | LFADS | Jango | single_day | 20150806 | 20150806 | 44 | r2 | 0.953339 |
| regression | LFADS | Jango | single_day | 20150808 | 20150808 | 44 | r2 | 0.9376 |
| regression | LFADS | Jango | single_day | 20150820 | 20150820 | 44 | r2 | 0.946243 |
| regression | LFADS | Jango | single_day | 20150825 | 20150825 | 44 | r2 | 0.944765 |
| regression | LFADS | Jango | single_day | 20150827 | 20150827 | 44 | r2 | 0.935352 |
| regression | LFADS | Jango | single_day | 20150831 | 20150831 | 44 | r2 | 0.706182 |
| regression | LFADS | Jango | single_day | 20150906 | 20150906 | 44 | r2 | 0.895063 |
| regression | LFADS | Jango | single_day | 20151102 | 20151102 | 44 | r2 | 0.876596 |
| regression | Cycle-GAN | Jango | single_day | 20150730 | 20150730 | 44 | r2 | 0.727797 |
| regression | Cycle-GAN | Jango | single_day | 20150801 | 20150801 | 44 | r2 | 0.906996 |
| regression | Cycle-GAN | Jango | single_day | 20150806 | 20150806 | 44 | r2 | 0.902273 |
| regression | Cycle-GAN | Jango | single_day | 20150808 | 20150808 | 44 | r2 | 0.908129 |
| regression | Cycle-GAN | Jango | single_day | 20150820 | 20150820 | 44 | r2 | 0.896057 |
| regression | Cycle-GAN | Jango | single_day | 20150825 | 20150825 | 44 | r2 | 0.892239 |
| regression | Cycle-GAN | Jango | single_day | 20150827 | 20150827 | 44 | r2 | 0.909221 |
| regression | Cycle-GAN | Jango | single_day | 20150831 | 20150831 | 44 | r2 | 0.877366 |
| regression | Cycle-GAN | Jango | single_day | 20150906 | 20150906 | 44 | r2 | 0.920398 |
| regression | Cycle-GAN | Jango | single_day | 20151102 | 20151102 | 44 | r2 | 0.841204 |
| regression | Stabilization | Jango | single_day | 20150730 | 20150730 | 44 | r2 | 0.442793 |
| regression | Stabilization | Jango | single_day | 20150801 | 20150801 | 44 | r2 | 0.855684 |
| regression | Stabilization | Jango | single_day | 20150806 | 20150806 | 44 | r2 | 0.851486 |
| regression | Stabilization | Jango | single_day | 20150808 | 20150808 | 44 | r2 | 0.793243 |
| regression | Stabilization | Jango | single_day | 20150820 | 20150820 | 44 | r2 | 0.853224 |
| regression | Stabilization | Jango | single_day | 20150825 | 20150825 | 44 | r2 | 0.864082 |
| regression | Stabilization | Jango | single_day | 20150827 | 20150827 | 44 | r2 | 0.896821 |
| regression | Stabilization | Jango | single_day | 20150831 | 20150831 | 44 | r2 | 0.742719 |
| regression | Stabilization | Jango | single_day | 20150906 | 20150906 | 44 | r2 | 0.82634 |
| regression | Stabilization | Jango | single_day | 20151102 | 20151102 | 44 | r2 | 0.711973 |
| regression | FENet | Jango | single_day | 20150730 | 20150730 | 44 | r2 | 0.745768 |
| regression | FENet | Jango | single_day | 20150801 | 20150801 | 44 | r2 | 0.88536 |
| regression | FENet | Jango | single_day | 20150806 | 20150806 | 44 | r2 | 0.856309 |
| regression | FENet | Jango | single_day | 20150808 | 20150808 | 44 | r2 | 0.879245 |
| regression | FENet | Jango | single_day | 20150820 | 20150820 | 44 | r2 | 0.869249 |
| regression | FENet | Jango | single_day | 20150825 | 20150825 | 44 | r2 | 0.838904 |
| regression | FENet | Jango | single_day | 20150827 | 20150827 | 44 | r2 | 0.8283 |
| regression | FENet | Jango | single_day | 20150831 | 20150831 | 44 | r2 | 0.865207 |
| regression | FENet | Jango | single_day | 20150906 | 20150906 | 44 | r2 | 0.843232 |
| regression | FENet | Jango | single_day | 20151102 | 20151102 | 44 | r2 | 0.819976 |
| regression | DFINE | Jango | single_day | 20150730 | 20150730 | 44 | r2 | 0.836713 |
| regression | DFINE | Jango | single_day | 20150801 | 20150801 | 44 | r2 | 0.945177 |
| regression | DFINE | Jango | single_day | 20150806 | 20150806 | 44 | r2 | 0.944101 |
| regression | DFINE | Jango | single_day | 20150808 | 20150808 | 44 | r2 | 0.907899 |
| regression | DFINE | Jango | single_day | 20150820 | 20150820 | 44 | r2 | 0.920091 |
| regression | DFINE | Jango | single_day | 20150825 | 20150825 | 44 | r2 | 0.879689 |
| regression | DFINE | Jango | single_day | 20150827 | 20150827 | 44 | r2 | 0.924228 |
| regression | DFINE | Jango | single_day | 20150831 | 20150831 | 44 | r2 | 0.91978 |
| regression | DFINE | Jango | single_day | 20150906 | 20150906 | 44 | r2 | 0.911321 |
| regression | DFINE | Jango | single_day | 20151102 | 20151102 | 44 | r2 | 0.908094 |
| regression | KF | Jango | single_day | 20150730 | 20150730 | 44 | r2 | 0.562942 |
| regression | KF | Jango | single_day | 20150801 | 20150801 | 44 | r2 | 0.620772 |
| regression | KF | Jango | single_day | 20150806 | 20150806 | 44 | r2 | 0.783768 |

| | | | | | | | | |
|---|---|---|---|---|---|---|---|---|
| regression | KF | Jango | single_day | 20150808 | 20150808 | 44 | r2 | 0.704261 |
| regression | KF | Jango | single_day | 20150820 | 20150820 | 44 | r2 | 0.70576 |
| regression | KF | Jango | single_day | 20150825 | 20150825 | 44 | r2 | 0.637114 |
| regression | KF | Jango | single_day | 20150827 | 20150827 | 44 | r2 | 0.692507 |
| regression | KF | Jango | single_day | 20150831 | 20150831 | 44 | r2 | 0.753591 |
| regression | KF | Jango | single_day | 20150906 | 20150906 | 44 | r2 | 0.716347 |
| regression | KF | Jango | single_day | 20151102 | 20151102 | 44 | r2 | 0.637093 |
| regression | seqVAE | Jango | single_day | 20150730 | 20150730 | 44 | r2 | 0.789748 |
| regression | seqVAE | Jango | single_day | 20150801 | 20150801 | 44 | r2 | 0.922016 |
| regression | seqVAE | Jango | single_day | 20150806 | 20150806 | 44 | r2 | 0.916534 |
| regression | seqVAE | Jango | single_day | 20150808 | 20150808 | 44 | r2 | 0.89362 |
| regression | seqVAE | Jango | single_day | 20150820 | 20150820 | 44 | r2 | 0.906782 |
| regression | seqVAE | Jango | single_day | 20150825 | 20150825 | 44 | r2 | 0.903998 |
| regression | seqVAE | Jango | single_day | 20150827 | 20150827 | 44 | r2 | 0.856447 |
| regression | seqVAE | Jango | single_day | 20150831 | 20150831 | 44 | r2 | 0.816376 |
| regression | seqVAE | Jango | single_day | 20150906 | 20150906 | 44 | r2 | 0.896008 |
| regression | seqVAE | Jango | single_day | 20151102 | 20151102 | 44 | r2 | 0.839342 |
| regression | NoMAD | Jango | single_day | 20150730 | 20150730 | 44 | r2 | 0.866637 |
| regression | NoMAD | Jango | single_day | 20150801 | 20150801 | 44 | r2 | 0.945293 |
| regression | NoMAD | Jango | single_day | 20150806 | 20150806 | 44 | r2 | 0.938549 |
| regression | NoMAD | Jango | single_day | 20150808 | 20150808 | 44 | r2 | 0.926525 |
| regression | NoMAD | Jango | single_day | 20150820 | 20150820 | 44 | r2 | 0.938602 |
| regression | NoMAD | Jango | single_day | 20150825 | 20150825 | 44 | r2 | 0.913213 |
| regression | NoMAD | Jango | single_day | 20150827 | 20150827 | 44 | r2 | 0.943893 |
| regression | NoMAD | Jango | single_day | 20150831 | 20150831 | 44 | r2 | 0.922414 |
| regression | NoMAD | Jango | single_day | 20150906 | 20150906 | 44 | r2 | 0.885348 |
| regression | NoMAD | Jango | single_day | 20151102 | 20151102 | 44 | r2 | 0.871054 |
| regression | LSTM | LINK CO | single_day | 20211005 | 20211005 | 44 | r2 | 0.645925 |
| regression | LSTM | LINK CO | single_day | 20211007 | 20211007 | 44 | r2 | 0.740711 |
| regression | LSTM | LINK CO | single_day | 20211008 | 20211008 | 44 | r2 | 0.709078 |
| regression | LSTM | LINK CO | single_day | 20211015 | 20211015 | 44 | r2 | 0.752054 |
| regression | LSTM | LINK CO | single_day | 20211019 | 20211019 | 44 | r2 | 0.754929 |
| regression | LSTM | LINK CO | single_day | 20211025 | 20211025 | 44 | r2 | 0.629301 |
| regression | LSTM | LINK CO | single_day | 20211026 | 20211026 | 44 | r2 | 0.618575 |
| regression | LSTM | LINK CO | single_day | 20211027 | 20211027 | 44 | r2 | 0.685171 |
| regression | LSTM | LINK CO | single_day | 20211101 | 20211101 | 44 | r2 | 0.723175 |
| regression | LSTM | LINK CO | single_day | 20211102 | 20211102 | 44 | r2 | 0.640328 |
| regression | Transformer | LINK CO | single_day | 20211005 | 20211005 | 44 | r2 | 0.282224 |
| regression | Transformer | LINK CO | single_day | 20211007 | 20211007 | 44 | r2 | 0.412591 |
| regression | Transformer | LINK CO | single_day | 20211008 | 20211008 | 44 | r2 | 0.342206 |
| regression | Transformer | LINK CO | single_day | 20211015 | 20211015 | 44 | r2 | 0.376575 |
| regression | Transformer | LINK CO | single_day | 20211019 | 20211019 | 44 | r2 | 0.358324 |
| regression | Transformer | LINK CO | single_day | 20211025 | 20211025 | 44 | r2 | 0.319351 |
| regression | Transformer | LINK CO | single_day | 20211026 | 20211026 | 44 | r2 | 0.287403 |
| regression | Transformer | LINK CO | single_day | 20211027 | 20211027 | 44 | r2 | 0.262204 |
| regression | Transformer | LINK CO | single_day | 20211101 | 20211101 | 44 | r2 | 0.368475 |
| regression | Transformer | LINK CO | single_day | 20211102 | 20211102 | 44 | r2 | 0.31752 |
| regression | LFADS | LINK CO | single_day | 20211005 | 20211005 | 44 | r2 | 0.0663531 |
| regression | LFADS | LINK CO | single_day | 20211007 | 20211007 | 44 | r2 | 0.0514316 |
| regression | LFADS | LINK CO | single_day | 20211008 | 20211008 | 44 | r2 | 0.0368602 |
| regression | LFADS | LINK CO | single_day | 20211015 | 20211015 | 44 | r2 | 0.0479444 |
| regression | LFADS | LINK CO | single_day | 20211019 | 20211019 | 44 | r2 | 0.171545 |
| regression | LFADS | LINK CO | single_day | 20211025 | 20211025 | 44 | r2 | 0.0529906 |
| regression | LFADS | LINK CO | single_day | 20211026 | 20211026 | 44 | r2 | 0.075895 |
| regression | LFADS | LINK CO | single_day | 20211027 | 20211027 | 44 | r2 | 0.0621816 |
| regression | LFADS | LINK CO | single_day | 20211101 | 20211101 | 44 | r2 | 0.0942072 |
| regression | LFADS | LINK CO | single_day | 20211102 | 20211102 | 44 | r2 | 0.0647629 |
| regression | Cycle-GAN | LINK CO | single_day | 20211005 | 20211005 | 44 | r2 | 0.357787 |
| regression | Cycle-GAN | LINK CO | single_day | 20211007 | 20211007 | 44 | r2 | 0.327824 |
| regression | Cycle-GAN | LINK CO | single_day | 20211008 | 20211008 | 44 | r2 | 0.381063 |
| regression | Cycle-GAN | LINK CO | single_day | 20211015 | 20211015 | 44 | r2 | 0.429668 |
| regression | Cycle-GAN | LINK CO | single_day | 20211019 | 20211019 | 44 | r2 | 0.320971 |
| regression | Cycle-GAN | LINK CO | single_day | 20211025 | 20211025 | 44 | r2 | 0.2535 |

| | | | | | | | | |
|---|---|---|---|---|---|---|---|---|
| regression | Cycle-GAN | LINK CO | single_day | 20211026 | 20211026 | 44 | r2 | 0.262487 |
| regression | Cycle-GAN | LINK CO | single_day | 20211027 | 20211027 | 44 | r2 | 0.238338 |
| regression | Cycle-GAN | LINK CO | single_day | 20211101 | 20211101 | 44 | r2 | 0.273775 |
| regression | Cycle-GAN | LINK CO | single_day | 20211102 | 20211102 | 44 | r2 | 0.260092 |
| regression | Stabilization | LINK CO | single_day | 20211005 | 20211005 | 44 | r2 | -0.0674035 |
| regression | Stabilization | LINK CO | single_day | 20211007 | 20211007 | 44 | r2 | 0.0885418 |
| regression | Stabilization | LINK CO | single_day | 20211008 | 20211008 | 44 | r2 | 0.0749724 |
| regression | Stabilization | LINK CO | single_day | 20211015 | 20211015 | 44 | r2 | 0.279048 |
| regression | Stabilization | LINK CO | single_day | 20211019 | 20211019 | 44 | r2 | 0.0322547 |
| regression | Stabilization | LINK CO | single_day | 20211025 | 20211025 | 44 | r2 | 0.0916206 |
| regression | Stabilization | LINK CO | single_day | 20211026 | 20211026 | 44 | r2 | 0.0470613 |
| regression | Stabilization | LINK CO | single_day | 20211027 | 20211027 | 44 | r2 | -0.0454146 |
| regression | Stabilization | LINK CO | single_day | 20211101 | 20211101 | 44 | r2 | 0.0748011 |
| regression | Stabilization | LINK CO | single_day | 20211102 | 20211102 | 44 | r2 | 0.097728 |
| regression | FENet | LINK CO | single_day | 20211005 | 20211005 | 44 | r2 | 0.390299 |
| regression | FENet | LINK CO | single_day | 20211007 | 20211007 | 44 | r2 | 0.472688 |
| regression | FENet | LINK CO | single_day | 20211008 | 20211008 | 44 | r2 | 0.354626 |
| regression | FENet | LINK CO | single_day | 20211015 | 20211015 | 44 | r2 | 0.495588 |
| regression | FENet | LINK CO | single_day | 20211019 | 20211019 | 44 | r2 | 0.418069 |
| regression | FENet | LINK CO | single_day | 20211025 | 20211025 | 44 | r2 | 0.340527 |
| regression | FENet | LINK CO | single_day | 20211026 | 20211026 | 44 | r2 | 0.358637 |
| regression | FENet | LINK CO | single_day | 20211027 | 20211027 | 44 | r2 | 0.245845 |
| regression | FENet | LINK CO | single_day | 20211101 | 20211101 | 44 | r2 | 0.432046 |
| regression | FENet | LINK CO | single_day | 20211102 | 20211102 | 44 | r2 | 0.352533 |
| regression | DFINE | LINK CO | single_day | 20211005 | 20211005 | 44 | r2 | 0.336128 |
| regression | DFINE | LINK CO | single_day | 20211007 | 20211007 | 44 | r2 | 0.377177 |
| regression | DFINE | LINK CO | single_day | 20211008 | 20211008 | 44 | r2 | 0.42686 |
| regression | DFINE | LINK CO | single_day | 20211015 | 20211015 | 44 | r2 | 0.487478 |
| regression | DFINE | LINK CO | single_day | 20211019 | 20211019 | 44 | r2 | 0.39316 |
| regression | DFINE | LINK CO | single_day | 20211025 | 20211025 | 44 | r2 | 0.332963 |
| regression | DFINE | LINK CO | single_day | 20211026 | 20211026 | 44 | r2 | 0.332706 |
| regression | DFINE | LINK CO | single_day | 20211027 | 20211027 | 44 | r2 | 0.353613 |
| regression | DFINE | LINK CO | single_day | 20211101 | 20211101 | 44 | r2 | 0.399245 |
| regression | DFINE | LINK CO | single_day | 20211102 | 20211102 | 44 | r2 | 0.408971 |
| regression | LSTM | LINK RTT | single_day | 20230123 | 20230123 | 44 | r2 | 0.725137 |
| regression | LSTM | LINK RTT | single_day | 20230124 | 20230124 | 44 | r2 | 0.710819 |
| regression | LSTM | LINK RTT | single_day | 20230127 | 20230127 | 44 | r2 | 0.6926 |
| regression | LSTM | LINK RTT | single_day | 20230130 | 20230130 | 44 | r2 | 0.623518 |
| regression | LSTM | LINK RTT | single_day | 20230131 | 20230131 | 44 | r2 | 0.69421 |
| regression | LSTM | LINK RTT | single_day | 20230203 | 20230203 | 44 | r2 | 0.67427 |
| regression | LSTM | LINK RTT | single_day | 20230206 | 20230206 | 44 | r2 | 0.675036 |
| regression | LSTM | LINK RTT | single_day | 20230207 | 20230207 | 44 | r2 | 0.649979 |
| regression | LSTM | LINK RTT | single_day | 20230210 | 20230210 | 44 | r2 | 0.592075 |
| regression | LSTM | LINK RTT | single_day | 20230213 | 20230213 | 44 | r2 | 0.567346 |
| regression | Transformer | LINK RTT | single_day | 20220805 | 20220805 | 44 | r2 | 0.41084 |
| regression | Transformer | LINK RTT | single_day | 20220810 | 20220810 | 44 | r2 | 0.479132 |
| regression | Transformer | LINK RTT | single_day | 20220811 | 20220811 | 44 | r2 | 0.494307 |
| regression | Transformer | LINK RTT | single_day | 20220817 | 20220817 | 44 | r2 | 0.54412 |
| regression | Transformer | LINK RTT | single_day | 20220818 | 20220818 | 44 | r2 | 0.519283 |
| regression | Transformer | LINK RTT | single_day | 20220819 | 20220819 | 44 | r2 | 0.604868 |
| regression | Transformer | LINK RTT | single_day | 20220825 | 20220825 | 44 | r2 | 0.454865 |
| regression | LFADS | LINK RTT | single_day | 20220805 | 20220805 | 44 | r2 | 0.205533 |
| regression | LFADS | LINK RTT | single_day | 20220810 | 20220810 | 44 | r2 | 0.178302 |
| regression | LFADS | LINK RTT | single_day | 20220811 | 20220811 | 44 | r2 | 0.0456493 |
| regression | LFADS | LINK RTT | single_day | 20220817 | 20220817 | 44 | r2 | 0.0733404 |
| regression | LFADS | LINK RTT | single_day | 20220818 | 20220818 | 44 | r2 | 0.244431 |
| regression | LFADS | LINK RTT | single_day | 20220819 | 20220819 | 44 | r2 | 0.446436 |
| regression | LFADS | LINK RTT | single_day | 20220825 | 20220825 | 44 | r2 | 0.246151 |
| regression | Cycle-GAN | LINK RTT | single_day | 20220805 | 20220805 | 44 | r2 | 0.326197 |
| regression | Cycle-GAN | LINK RTT | single_day | 20220810 | 20220810 | 44 | r2 | 0.397281 |
| regression | Cycle-GAN | LINK RTT | single_day | 20220811 | 20220811 | 44 | r2 | 0.304211 |
| regression | Cycle-GAN | LINK RTT | single_day | 20220817 | 20220817 | 44 | r2 | 0.374449 |
| regression | Cycle-GAN | LINK RTT | single_day | 20220818 | 20220818 | 44 | r2 | 0.344537 |

| | | | | | | | | |
|---|---|---|---|---|---|---|---|---|
| regression | Cycle-GAN | LINK RTT | single_day | 20220819 | 20220819 | 44 | r2 | 0.354791 |
| regression | Cycle-GAN | LINK RTT | single_day | 20220825 | 20220825 | 44 | r2 | 0.29888 |
| regression | Stabilization | LINK RTT | single_day | 20220805 | 20220805 | 44 | r2 | 0.313264 |
| regression | Stabilization | LINK RTT | single_day | 20220810 | 20220810 | 44 | r2 | 0.336258 |
| regression | Stabilization | LINK RTT | single_day | 20220811 | 20220811 | 44 | r2 | 0.22225 |
| regression | Stabilization | LINK RTT | single_day | 20220817 | 20220817 | 44 | r2 | 0.310076 |
| regression | Stabilization | LINK RTT | single_day | 20220818 | 20220818 | 44 | r2 | 0.437879 |
| regression | Stabilization | LINK RTT | single_day | 20220819 | 20220819 | 44 | r2 | 0.508169 |
| regression | Stabilization | LINK RTT | single_day | 20220825 | 20220825 | 44 | r2 | 0.309501 |
| regression | FENet | LINK RTT | single_day | 20220805 | 20220805 | 44 | r2 | 0.393759 |
| regression | FENet | LINK RTT | single_day | 20220810 | 20220810 | 44 | r2 | 0.449276 |
| regression | FENet | LINK RTT | single_day | 20220811 | 20220811 | 44 | r2 | 0.418099 |
| regression | FENet | LINK RTT | single_day | 20220817 | 20220817 | 44 | r2 | 0.499671 |
| regression | FENet | LINK RTT | single_day | 20220818 | 20220818 | 44 | r2 | 0.531254 |
| regression | FENet | LINK RTT | single_day | 20220819 | 20220819 | 44 | r2 | 0.596104 |
| regression | FENet | LINK RTT | single_day | 20220825 | 20220825 | 44 | r2 | 0.389339 |
| regression | DFINE | LINK RTT | single_day | 20220805 | 20220805 | 44 | r2 | 0.41974 |
| regression | DFINE | LINK RTT | single_day | 20220810 | 20220810 | 44 | r2 | 0.486251 |
| regression | DFINE | LINK RTT | single_day | 20220811 | 20220811 | 44 | r2 | 0.473995 |
| regression | DFINE | LINK RTT | single_day | 20220817 | 20220817 | 44 | r2 | 0.607354 |
| regression | DFINE | LINK RTT | single_day | 20220818 | 20220818 | 44 | r2 | 0.513362 |
| regression | DFINE | LINK RTT | single_day | 20220819 | 20220819 | 44 | r2 | 0.617482 |
| regression | DFINE | LINK RTT | single_day | 20220825 | 20220825 | 44 | r2 | 0.487737 |
| regression | LSTM | Indy | single_day | 20161005 | 20161005 | 44 | r2 | 0.947049 |
| regression | LSTM | Indy | single_day | 20161006 | 20161006 | 44 | r2 | 0.959323 |
| regression | LSTM | Indy | single_day | 20161007 | 20161007 | 44 | r2 | 0.942652 |
| regression | LSTM | Indy | single_day | 20161011 | 20161011 | 44 | r2 | 0.948239 |
| regression | LSTM | Indy | single_day | 20161013 | 20161013 | 44 | r2 | 0.938754 |
| regression | LSTM | Indy | single_day | 20161014 | 20161014 | 44 | r2 | 0.946172 |
| regression | LSTM | Indy | single_day | 20161017 | 20161017 | 44 | r2 | 0.94955 |
| regression | LSTM | Indy | single_day | 20161024 | 20161024 | 44 | r2 | 0.937554 |
| regression | LSTM | Indy | single_day | 20161025 | 20161025 | 44 | r2 | 0.947666 |
| regression | LSTM | Indy | single_day | 20161026 | 20161026 | 44 | r2 | 0.94791 |
| regression | Transformer | Indy | single_day | 20161005 | 20161005 | 44 | r2 | 0.538408 |
| regression | Transformer | Indy | single_day | 20161006 | 20161006 | 44 | r2 | 0.590498 |
| regression | Transformer | Indy | single_day | 20161007 | 20161007 | 44 | r2 | 0.557462 |
| regression | Transformer | Indy | single_day | 20161011 | 20161011 | 44 | r2 | 0.634613 |
| regression | Transformer | Indy | single_day | 20161013 | 20161013 | 44 | r2 | 0.52579 |
| regression | Transformer | Indy | single_day | 20161014 | 20161014 | 44 | r2 | 0.596138 |
| regression | Transformer | Indy | single_day | 20161017 | 20161017 | 44 | r2 | 0.57165 |
| regression | Transformer | Indy | single_day | 20161024 | 20161024 | 44 | r2 | 0.555307 |
| regression | Transformer | Indy | single_day | 20161025 | 20161025 | 44 | r2 | 0.58369 |
| regression | Transformer | Indy | single_day | 20161026 | 20161026 | 44 | r2 | 0.54816 |
| regression | LFADS | Indy | single_day | 20161005 | 20161005 | 44 | r2 | 0.480425 |
| regression | LFADS | Indy | single_day | 20161006 | 20161006 | 44 | r2 | 0.50984 |
| regression | LFADS | Indy | single_day | 20161007 | 20161007 | 44 | r2 | 0.495146 |
| regression | LFADS | Indy | single_day | 20161011 | 20161011 | 44 | r2 | 0.495875 |
| regression | LFADS | Indy | single_day | 20161013 | 20161013 | 44 | r2 | 0.592668 |
| regression | LFADS | Indy | single_day | 20161014 | 20161014 | 44 | r2 | 0.51993 |
| regression | LFADS | Indy | single_day | 20161017 | 20161017 | 44 | r2 | 0.601023 |
| regression | LFADS | Indy | single_day | 20161024 | 20161024 | 44 | r2 | 0.408558 |
| regression | LFADS | Indy | single_day | 20161025 | 20161025 | 44 | r2 | 0.563965 |
| regression | LFADS | Indy | single_day | 20161026 | 20161026 | 44 | r2 | 0.530593 |
| regression | Cycle-GAN | Indy | single_day | 20161005 | 20161005 | 44 | r2 | 0.189724 |
| regression | Cycle-GAN | Indy | single_day | 20161006 | 20161006 | 44 | r2 | 0.26963 |
| regression | Cycle-GAN | Indy | single_day | 20161007 | 20161007 | 44 | r2 | 0.299029 |
| regression | Cycle-GAN | Indy | single_day | 20161011 | 20161011 | 44 | r2 | 0.411624 |
| regression | Cycle-GAN | Indy | single_day | 20161013 | 20161013 | 44 | r2 | 0.336467 |
| regression | Cycle-GAN | Indy | single_day | 20161014 | 20161014 | 44 | r2 | 0.321852 |
| regression | Cycle-GAN | Indy | single_day | 20161017 | 20161017 | 44 | r2 | 0.271495 |
| regression | Cycle-GAN | Indy | single_day | 20161024 | 20161024 | 44 | r2 | 0.26713 |
| regression | Cycle-GAN | Indy | single_day | 20161025 | 20161025 | 44 | r2 | 0.325581 |
| regression | Cycle-GAN | Indy | single_day | 20161026 | 20161026 | 44 | r2 | 0.313102 |

| | | | | | | | | |
|---|---|---|---|---|---|---|---|---|
| regression | Stabilization | Indy | single_day | 20161005 | 20161005 | 44 | r2 | 0.253827 |
| regression | Stabilization | Indy | single_day | 20161006 | 20161006 | 44 | r2 | 0.298724 |
| regression | Stabilization | Indy | single_day | 20161007 | 20161007 | 44 | r2 | 0.296334 |
| regression | Stabilization | Indy | single_day | 20161011 | 20161011 | 44 | r2 | 0.263636 |
| regression | Stabilization | Indy | single_day | 20161013 | 20161013 | 44 | r2 | 0.227158 |
| regression | Stabilization | Indy | single_day | 20161014 | 20161014 | 44 | r2 | 0.369309 |
| regression | Stabilization | Indy | single_day | 20161017 | 20161017 | 44 | r2 | 0.325784 |
| regression | Stabilization | Indy | single_day | 20161024 | 20161024 | 44 | r2 | 0.255508 |
| regression | Stabilization | Indy | single_day | 20161025 | 20161025 | 44 | r2 | 0.275941 |
| regression | Stabilization | Indy | single_day | 20161026 | 20161026 | 44 | r2 | 0.210883 |
| regression | FENet | Indy | single_day | 20161005 | 20161005 | 44 | r2 | 0.232211 |
| regression | FENet | Indy | single_day | 20161006 | 20161006 | 44 | r2 | 0.378854 |
| regression | FENet | Indy | single_day | 20161007 | 20161007 | 44 | r2 | 0.379868 |
| regression | FENet | Indy | single_day | 20161011 | 20161011 | 44 | r2 | 0.431243 |
| regression | FENet | Indy | single_day | 20161013 | 20161013 | 44 | r2 | 0.424382 |
| regression | FENet | Indy | single_day | 20161014 | 20161014 | 44 | r2 | 0.447581 |
| regression | FENet | Indy | single_day | 20161017 | 20161017 | 44 | r2 | 0.384486 |
| regression | FENet | Indy | single_day | 20161024 | 20161024 | 44 | r2 | 0.322164 |
| regression | FENet | Indy | single_day | 20161025 | 20161025 | 44 | r2 | 0.391775 |
| regression | FENet | Indy | single_day | 20161026 | 20161026 | 44 | r2 | 0.385602 |
| regression | DFINE | Indy | single_day | 20161005 | 20161005 | 44 | r2 | 0.82816 |
| regression | DFINE | Indy | single_day | 20161006 | 20161006 | 44 | r2 | 0.85947 |
| regression | DFINE | Indy | single_day | 20161007 | 20161007 | 44 | r2 | 0.773047 |
| regression | DFINE | Indy | single_day | 20161011 | 20161011 | 44 | r2 | 0.794907 |
| regression | DFINE | Indy | single_day | 20161013 | 20161013 | 44 | r2 | 0.826442 |
| regression | DFINE | Indy | single_day | 20161014 | 20161014 | 44 | r2 | 0.833754 |
| regression | DFINE | Indy | single_day | 20161017 | 20161017 | 44 | r2 | 0.820781 |
| regression | DFINE | Indy | single_day | 20161024 | 20161024 | 44 | r2 | 0.819746 |
| regression | DFINE | Indy | single_day | 20161025 | 20161025 | 44 | r2 | 0.816391 |
| regression | DFINE | Indy | single_day | 20161026 | 20161026 | 44 | r2 | 0.800679 |
| regression | LSTM | FALCON M2 | cross_day | 20201019__to__20201020 | 20201030 | 44 | r2 | 0.253233 |
| regression | LSTM | FALCON M2 | cross_day | 20201019__to__20201020 | 20201030 | 44 | r2 | 0.336091 |
| regression | LSTM | FALCON M2 | cross_day | 20201019__to__20201020 | 20201118 | 44 | r2 | 0.0497588 |
| regression | Transformer | FALCON M2 | cross_day | 20201019__to__20201020 | 20201030 | 44 | r2 | 0.20475 |
| regression | Transformer | FALCON M2 | cross_day | 20201019__to__20201020 | 20201030 | 44 | r2 | 0.317925 |
| regression | Transformer | FALCON M2 | cross_day | 20201019__to__20201020 | 20201118 | 44 | r2 | 0.0893122 |
| regression | LFADS | FALCON M2 | cross_day | 20201019__to__20201020 | 20201030 | 44 | r2 | 0.173432 |
| regression | LFADS | FALCON M2 | cross_day | 20201019__to__20201020 | 20201030 | 44 | r2 | 0.103201 |
| regression | LFADS | FALCON M2 | cross_day | 20201019__to__20201020 | 20201118 | 44 | r2 | -0.144605 |
| regression | LFADS | FALCON M2 | cross_day | 20201019__to__20201020 | 20201119 | 44 | r2 | -0.301255 |
| regression | LFADS | FALCON M2 | cross_day | 20201019__to__20201020 | 20201124 | 44 | r2 | -0.44669 |
| regression | LFADS | FALCON M2 | cross_day | 20201019__to__20201020 | 20201124 | 44 | r2 | 0.044935 |
| regression | Cycle-GAN | FALCON M2 | cross_day | 20201019__to__20201020 | 20201030 | 44 | r2 | -0.0621497 |
| regression | Cycle-GAN | FALCON M2 | cross_day | 20201019__to__20201020 | 20201030 | 44 | r2 | -0.0751908 |
| regression | Cycle-GAN | FALCON M2 | cross_day | 20201019__to__20201020 | 20201118 | 44 | r2 | -0.0308357 |
| regression | Cycle-GAN | FALCON M2 | cross_day | 20201019__to__20201020 | 20201119 | 44 | r2 | -0.0469251 |
| regression | Cycle-GAN | FALCON M2 | cross_day | 20201019__to__20201020 | 20201124 | 44 | r2 | -0.28927 |
| regression | Cycle-GAN | FALCON M2 | cross_day | 20201019__to__20201020 | 20201124 | 44 | r2 | -0.188342 |
| regression | Stabilization | FALCON M2 | cross_day | 20201019__to__20201020 | 20201030 | 44 | r2 | -0.00361882 |
| regression | Stabilization | FALCON M2 | cross_day | 20201019__to__20201020 | 20201030 | 44 | r2 | -0.00169497 |
| regression | Stabilization | FALCON M2 | cross_day | 20201019__to__20201020 | 20201118 | 44 | r2 | -0.140495 |
| regression | FENet | FALCON M2 | cross_day | 20201019__to__20201020 | 20201030 | 44 | r2 | 0.270789 |
| regression | FENet | FALCON M2 | cross_day | 20201019__to__20201020 | 20201030 | 44 | r2 | 0.2276 |
| regression | FENet | FALCON M2 | cross_day | 20201019__to__20201020 | 20201124 | 44 | r2 | 0.125294 |
| regression | DFINE | FALCON M2 | cross_day | 20201019__to__20201020 | 20201030 | 44 | r2 | 0.187857 |
| regression | DFINE | FALCON M2 | cross_day | 20201019__to__20201020 | 20201030 | 44 | r2 | 0.321775 |
| regression | DFINE | FALCON M2 | cross_day | 20201019__to__20201020 | 20201124 | 44 | r2 | 0.0917456 |
| regression | LSTM | Jango | cross_day | 20150730__to__20150828 | 20150906 | 44 | r2 | 0.838628 |
| regression | LSTM | Jango | cross_day | 20150730__to__20150828 | 20150908 | 44 | r2 | 0.00273141 |
| regression | LSTM | Jango | cross_day | 20150730__to__20150828 | 20151029 | 44 | r2 | 0.92922 |
| regression | LSTM | Jango | cross_day | 20150730__to__20150828 | 20151102 | 44 | r2 | 0.725579 |
| regression | Transformer | Jango | cross_day | 20150730__to__20150828 | 20150906 | 44 | r2 | 0.846504 |
| regression | Transformer | Jango | cross_day | 20150730__to__20150828 | 20150908 | 44 | r2 | -0.149524 |

| | | | | | | | | |
|---|---|---|---|---|---|---|---|---|
| regression | Transformer | Jango | cross_day | 20150730__to__20150828 | 20151029 | 44 | r2 | 0.830514 |
| regression | Transformer | Jango | cross_day | 20150730__to__20150828 | 20151102 | 44 | r2 | 0.514046 |
| regression | LFADS | Jango | cross_day | 20150730__to__20150828 | 20150906 | 44 | r2 | 0.749789 |
| regression | LFADS | Jango | cross_day | 20150730__to__20150828 | 20150908 | 44 | r2 | 0.264598 |
| regression | LFADS | Jango | cross_day | 20150730__to__20150828 | 20151029 | 44 | r2 | 0.802621 |
| regression | LFADS | Jango | cross_day | 20150730__to__20150828 | 20151102 | 44 | r2 | 0.128072 |
| regression | Cycle-GAN | Jango | cross_day | 20150730__to__20150828 | 20150906 | 44 | r2 | 0.302573 |
| regression | Cycle-GAN | Jango | cross_day | 20150730__to__20150828 | 20150908 | 44 | r2 | -0.0319751 |
| regression | Cycle-GAN | Jango | cross_day | 20150730__to__20150828 | 20151029 | 44 | r2 | 0.47228 |
| regression | Cycle-GAN | Jango | cross_day | 20150730__to__20150828 | 20151102 | 44 | r2 | 0.208836 |
| regression | Stabilization | Jango | cross_day | 20150730__to__20150828 | 20150906 | 44 | r2 | 0.836614 |
| regression | Stabilization | Jango | cross_day | 20150730__to__20150828 | 20150908 | 44 | r2 | 0.843365 |
| regression | Stabilization | Jango | cross_day | 20150730__to__20150828 | 20151029 | 44 | r2 | 0.869974 |
| regression | Stabilization | Jango | cross_day | 20150730__to__20150828 | 20151102 | 44 | r2 | 0.802046 |
| regression | FENet | Jango | cross_day | 20150730__to__20150828 | 20150906 | 44 | r2 | 0.794359 |
| regression | FENet | Jango | cross_day | 20150730__to__20150828 | 20150908 | 44 | r2 | -0.245218 |
| regression | FENet | Jango | cross_day | 20150730__to__20150828 | 20151029 | 44 | r2 | 0.722123 |
| regression | FENet | Jango | cross_day | 20150730__to__20150828 | 20151102 | 44 | r2 | 0.598277 |
| regression | DFINE | Jango | cross_day | 20150730__to__20150828 | 20150906 | 44 | r2 | 0.631758 |
| regression | DFINE | Jango | cross_day | 20150730__to__20150828 | 20150908 | 44 | r2 | -0.101577 |
| regression | DFINE | Jango | cross_day | 20150730__to__20150828 | 20151029 | 44 | r2 | 0.838573 |
| regression | DFINE | Jango | cross_day | 20150730__to__20150828 | 20151102 | 44 | r2 | -0.321916 |
| regression | KF | Jango | cross_day | 20150730__to__20150828 | 20150906 | 44 | r2 | 0.580046 |
| regression | KF | Jango | cross_day | 20150730__to__20150828 | 20150908 | 44 | r2 | 0.486745 |
| regression | KF | Jango | cross_day | 20150730__to__20150828 | 20151029 | 44 | r2 | 0.56702 |
| regression | KF | Jango | cross_day | 20150730__to__20150828 | 20151102 | 44 | r2 | 0.510479 |
| regression | seqVAE | Jango | cross_day | 20150730__to__20150828 | 20150906 | 44 | r2 | 0.946692 |
| regression | seqVAE | Jango | cross_day | 20150730__to__20150828 | 20150908 | 44 | r2 | 0.902291 |
| regression | seqVAE | Jango | cross_day | 20150730__to__20150828 | 20151029 | 44 | r2 | 0.912088 |
| regression | seqVAE | Jango | cross_day | 20150730__to__20150828 | 20151102 | 44 | r2 | 0.757964 |
| regression | NoMAD | Jango | cross_day | 20150730__to__20150828 | 20150906 | 44 | r2 | 0.952309 |
| regression | NoMAD | Jango | cross_day | 20150730__to__20150828 | 20150908 | 44 | r2 | 0.901141 |
| regression | NoMAD | Jango | cross_day | 20150730__to__20150828 | 20151029 | 44 | r2 | 0.926784 |
| regression | NoMAD | Jango | cross_day | 20150730__to__20150828 | 20151102 | 44 | r2 | 0.798989 |
| regression | LSTM | LINK CO | cross_day | 20210104__to__20210111 | 20210225 | 44 | r2 | 0.0161349 |
| regression | LSTM | LINK CO | cross_day | 20210104__to__20210111 | 20210226 | 44 | r2 | -0.0425307 |
| regression | LSTM | LINK CO | cross_day | 20210104__to__20210111 | 20210227 | 44 | r2 | 0.0167732 |
| regression | LSTM | LINK CO | cross_day | 20210104__to__20210111 | 20210309 | 44 | r2 | 0.127111 |
| regression | LSTM | LINK CO | cross_day | 20210104__to__20210111 | 20210313 | 44 | r2 | -0.13128 |
| regression | LSTM | LINK CO | cross_day | 20210104__to__20210111 | 20210329 | 44 | r2 | 0.188493 |
| regression | LSTM | LINK CO | cross_day | 20210104__to__20210111 | 20210330 | 44 | r2 | 0.0639888 |
| regression | LSTM | LINK CO | cross_day | 20210104__to__20210111 | 20210402 | 44 | r2 | -0.304444 |
| regression | LSTM | LINK CO | cross_day | 20210104__to__20210111 | 20210406 | 44 | r2 | 0.157634 |
| regression | LSTM | LINK CO | cross_day | 20210104__to__20210111 | 20210409 | 44 | r2 | -0.132362 |
| regression | LSTM | LINK CO | cross_day | 20210104__to__20210111 | 20210415 | 44 | r2 | -0.164775 |
| regression | LSTM | LINK CO | cross_day | 20210104__to__20210111 | 20210420 | 44 | r2 | 0.0180058 |
| regression | LSTM | LINK CO | cross_day | 20210104__to__20210111 | 20210503 | 44 | r2 | 0.00122645 |
| regression | LSTM | LINK CO | cross_day | 20210104__to__20210111 | 20210504 | 44 | r2 | -0.33297 |
| regression | LSTM | LINK CO | cross_day | 20210104__to__20210111 | 20210511 | 44 | r2 | 0.24193 |
| regression | LSTM | LINK CO | cross_day | 20210104__to__20210111 | 20210518 | 44 | r2 | -0.344467 |
| regression | LSTM | LINK CO | cross_day | 20210104__to__20210111 | 20210520 | 44 | r2 | 0.171184 |
| regression | LSTM | LINK CO | cross_day | 20210104__to__20210111 | 20210601 | 44 | r2 | 0.0579256 |
| regression | Transformer | LINK CO | cross_day | 20210104__to__20210111 | 20210225 | 44 | r2 | 0.299177 |
| regression | Transformer | LINK CO | cross_day | 20210104__to__20210111 | 20210226 | 44 | r2 | 0.131917 |
| regression | Transformer | LINK CO | cross_day | 20210104__to__20210111 | 20210227 | 44 | r2 | 0.146589 |
| regression | Transformer | LINK CO | cross_day | 20210104__to__20210111 | 20210309 | 44 | r2 | 0.177757 |
| regression | Transformer | LINK CO | cross_day | 20210104__to__20210111 | 20210313 | 44 | r2 | -0.0219252 |
| regression | Transformer | LINK CO | cross_day | 20210104__to__20210111 | 20210329 | 44 | r2 | 0.240438 |
| regression | Transformer | LINK CO | cross_day | 20210104__to__20210111 | 20210330 | 44 | r2 | 0.217023 |
| regression | Transformer | LINK CO | cross_day | 20210104__to__20210111 | 20210402 | 44 | r2 | -0.12853 |
| regression | Transformer | LINK CO | cross_day | 20210104__to__20210111 | 20210406 | 44 | r2 | 0.217562 |
| regression | Transformer | LINK CO | cross_day | 20210104__to__20210111 | 20210409 | 44 | r2 | 0.134938 |
| regression | Transformer | LINK CO | cross_day | 20210104__to__20210111 | 20210415 | 44 | r2 | 0.00174597 |

| | | | | | | | | |
|---|---|---|---|---|---|---|---|---|
| regression | Transformer | LINK CO | cross_day | 20210104__to__20210111 | 20210420 | 44 | r2 | -0.174715 |
| regression | Transformer | LINK CO | cross_day | 20210104__to__20210111 | 20210503 | 44 | r2 | -0.0651919 |
| regression | Transformer | LINK CO | cross_day | 20210104__to__20210111 | 20210504 | 44 | r2 | -0.240946 |
| regression | Transformer | LINK CO | cross_day | 20210104__to__20210111 | 20210511 | 44 | r2 | 0.430293 |
| regression | Transformer | LINK CO | cross_day | 20210104__to__20210111 | 20210518 | 44 | r2 | 0.324507 |
| regression | Transformer | LINK CO | cross_day | 20210104__to__20210111 | 20210520 | 44 | r2 | 0.26312 |
| regression | Transformer | LINK CO | cross_day | 20210104__to__20210111 | 20210601 | 44 | r2 | 0.232746 |
| regression | LFADS | LINK CO | cross_day | 20210104__to__20210111 | 20210225 | 44 | r2 | 0.11043 |
| regression | LFADS | LINK CO | cross_day | 20210104__to__20210111 | 20210226 | 44 | r2 | 0.096683 |
| regression | LFADS | LINK CO | cross_day | 20210104__to__20210111 | 20210227 | 44 | r2 | 0.0854396 |
| regression | LFADS | LINK CO | cross_day | 20210104__to__20210111 | 20210309 | 44 | r2 | 0.0927323 |
| regression | LFADS | LINK CO | cross_day | 20210104__to__20210111 | 20210313 | 44 | r2 | 0.105012 |
| regression | LFADS | LINK CO | cross_day | 20210104__to__20210111 | 20210329 | 44 | r2 | 0.116482 |
| regression | LFADS | LINK CO | cross_day | 20210104__to__20210111 | 20210330 | 44 | r2 | 0.0559374 |
| regression | LFADS | LINK CO | cross_day | 20210104__to__20210111 | 20210402 | 44 | r2 | 0.0859917 |
| regression | LFADS | LINK CO | cross_day | 20210104__to__20210111 | 20210406 | 44 | r2 | 0.0762648 |
| regression | LFADS | LINK CO | cross_day | 20210104__to__20210111 | 20210409 | 44 | r2 | 0.150402 |
| regression | LFADS | LINK CO | cross_day | 20210104__to__20210111 | 20210415 | 44 | r2 | 0.0848566 |
| regression | LFADS | LINK CO | cross_day | 20210104__to__20210111 | 20210420 | 44 | r2 | 0.0689788 |
| regression | LFADS | LINK CO | cross_day | 20210104__to__20210111 | 20210503 | 44 | r2 | -0.0144896 |
| regression | LFADS | LINK CO | cross_day | 20210104__to__20210111 | 20210504 | 44 | r2 | 0.00116304 |
| regression | LFADS | LINK CO | cross_day | 20210104__to__20210111 | 20210511 | 44 | r2 | 0.160578 |
| regression | LFADS | LINK CO | cross_day | 20210104__to__20210111 | 20210518 | 44 | r2 | 0.17886 |
| regression | LFADS | LINK CO | cross_day | 20210104__to__20210111 | 20210520 | 44 | r2 | 0.1803 |
| regression | LFADS | LINK CO | cross_day | 20210104__to__20210111 | 20210601 | 44 | r2 | 0.183782 |
| regression | Cycle-GAN | LINK CO | cross_day | 20210104__to__20210111 | 20210225 | 44 | r2 | 0.0167552 |
| regression | Cycle-GAN | LINK CO | cross_day | 20210104__to__20210111 | 20210226 | 44 | r2 | 0.0175945 |
| regression | Cycle-GAN | LINK CO | cross_day | 20210104__to__20210111 | 20210227 | 44 | r2 | 0.0217069 |
| regression | Cycle-GAN | LINK CO | cross_day | 20210104__to__20210111 | 20210309 | 44 | r2 | 0.0156749 |
| regression | Cycle-GAN | LINK CO | cross_day | 20210104__to__20210111 | 20210313 | 44 | r2 | -0.0117951 |
| regression | Cycle-GAN | LINK CO | cross_day | 20210104__to__20210111 | 20210329 | 44 | r2 | 0.00912379 |
| regression | Cycle-GAN | LINK CO | cross_day | 20210104__to__20210111 | 20210330 | 44 | r2 | -0.00293416 |
| regression | Cycle-GAN | LINK CO | cross_day | 20210104__to__20210111 | 20210402 | 44 | r2 | -0.0165726 |
| regression | Cycle-GAN | LINK CO | cross_day | 20210104__to__20210111 | 20210406 | 44 | r2 | 0.0222532 |
| regression | Cycle-GAN | LINK CO | cross_day | 20210104__to__20210111 | 20210409 | 44 | r2 | -0.0860877 |
| regression | Cycle-GAN | LINK CO | cross_day | 20210104__to__20210111 | 20210415 | 44 | r2 | -0.219692 |
| regression | Cycle-GAN | LINK CO | cross_day | 20210104__to__20210111 | 20210420 | 44 | r2 | -0.0420645 |
| regression | Cycle-GAN | LINK CO | cross_day | 20210104__to__20210111 | 20210503 | 44 | r2 | -0.0434931 |
| regression | Cycle-GAN | LINK CO | cross_day | 20210104__to__20210111 | 20210504 | 44 | r2 | -0.183362 |
| regression | Cycle-GAN | LINK CO | cross_day | 20210104__to__20210111 | 20210511 | 44 | r2 | 0.0524427 |
| regression | Cycle-GAN | LINK CO | cross_day | 20210104__to__20210111 | 20210518 | 44 | r2 | -0.0923774 |
| regression | Cycle-GAN | LINK CO | cross_day | 20210104__to__20210111 | 20210520 | 44 | r2 | -0.161142 |
| regression | Cycle-GAN | LINK CO | cross_day | 20210104__to__20210111 | 20210601 | 44 | r2 | -0.000165439 |
| regression | Stabilization | LINK CO | cross_day | 20210104__to__20210111 | 20210225 | 44 | r2 | -0.110436 |
| regression | Stabilization | LINK CO | cross_day | 20210104__to__20210111 | 20210226 | 44 | r2 | 0.0231499 |
| regression | Stabilization | LINK CO | cross_day | 20210104__to__20210111 | 20210227 | 44 | r2 | -0.0701098 |
| regression | Stabilization | LINK CO | cross_day | 20210104__to__20210111 | 20210309 | 44 | r2 | 0.118405 |
| regression | Stabilization | LINK CO | cross_day | 20210104__to__20210111 | 20210313 | 44 | r2 | -0.039555 |
| regression | Stabilization | LINK CO | cross_day | 20210104__to__20210111 | 20210329 | 44 | r2 | -0.285274 |
| regression | Stabilization | LINK CO | cross_day | 20210104__to__20210111 | 20210330 | 44 | r2 | -0.127619 |
| regression | Stabilization | LINK CO | cross_day | 20210104__to__20210111 | 20210402 | 44 | r2 | -0.501738 |
| regression | Stabilization | LINK CO | cross_day | 20210104__to__20210111 | 20210406 | 44 | r2 | -0.137454 |
| regression | Stabilization | LINK CO | cross_day | 20210104__to__20210111 | 20210409 | 44 | r2 | -0.115399 |
| regression | Stabilization | LINK CO | cross_day | 20210104__to__20210111 | 20210415 | 44 | r2 | 0.0607084 |
| regression | Stabilization | LINK CO | cross_day | 20210104__to__20210111 | 20210420 | 44 | r2 | -0.096563 |
| regression | Stabilization | LINK CO | cross_day | 20210104__to__20210111 | 20210503 | 44 | r2 | -0.103935 |
| regression | Stabilization | LINK CO | cross_day | 20210104__to__20210111 | 20210504 | 44 | r2 | -0.0153552 |
| regression | Stabilization | LINK CO | cross_day | 20210104__to__20210111 | 20210511 | 44 | r2 | -0.106903 |
| regression | Stabilization | LINK CO | cross_day | 20210104__to__20210111 | 20210518 | 44 | r2 | 0.190322 |
| regression | Stabilization | LINK CO | cross_day | 20210104__to__20210111 | 20210520 | 44 | r2 | 0.177713 |
| regression | Stabilization | LINK CO | cross_day | 20210104__to__20210111 | 20210601 | 44 | r2 | 0.12957 |
| regression | FENet | LINK CO | cross_day | 20210104__to__20210111 | 20210225 | 44 | r2 | 0.0103616 |
| regression | FENet | LINK CO | cross_day | 20210104__to__20210111 | 20210226 | 44 | r2 | 0.0118539 |

| | | | | | | | | |
|---|---|---|---|---|---|---|---|---|
| regression | FENet | LINK CO | cross_day | 20210104__to__20210111 | 20210227 | 44 | r2 | 0.0235517 |
| regression | FENet | LINK CO | cross_day | 20210104__to__20210111 | 20210309 | 44 | r2 | 0.0295054 |
| regression | FENet | LINK CO | cross_day | 20210104__to__20210111 | 20210313 | 44 | r2 | 0.0439342 |
| regression | FENet | LINK CO | cross_day | 20210104__to__20210111 | 20210329 | 44 | r2 | 0.018755 |
| regression | FENet | LINK CO | cross_day | 20210104__to__20210111 | 20210330 | 44 | r2 | -0.00338247 |
| regression | FENet | LINK CO | cross_day | 20210104__to__20210111 | 20210402 | 44 | r2 | -0.0895113 |
| regression | FENet | LINK CO | cross_day | 20210104__to__20210111 | 20210406 | 44 | r2 | 0.0245756 |
| regression | FENet | LINK CO | cross_day | 20210104__to__20210111 | 20210409 | 44 | r2 | -0.0568016 |
| regression | FENet | LINK CO | cross_day | 20210104__to__20210111 | 20210415 | 44 | r2 | 0.0184886 |
| regression | FENet | LINK CO | cross_day | 20210104__to__20210111 | 20210420 | 44 | r2 | -0.0144637 |
| regression | FENet | LINK CO | cross_day | 20210104__to__20210111 | 20210503 | 44 | r2 | -0.0110661 |
| regression | FENet | LINK CO | cross_day | 20210104__to__20210111 | 20210504 | 44 | r2 | -0.024497 |
| regression | FENet | LINK CO | cross_day | 20210104__to__20210111 | 20210511 | 44 | r2 | 0.0171013 |
| regression | FENet | LINK CO | cross_day | 20210104__to__20210111 | 20210518 | 44 | r2 | 0.0148087 |
| regression | FENet | LINK CO | cross_day | 20210104__to__20210111 | 20210520 | 44 | r2 | 0.024677 |
| regression | FENet | LINK CO | cross_day | 20210104__to__20210111 | 20210601 | 44 | r2 | 0.00319275 |
| regression | DFINE | LINK CO | cross_day | 20210104__to__20210111 | 20210225 | 44 | r2 | 0.0450032 |
| regression | DFINE | LINK CO | cross_day | 20210104__to__20210111 | 20210226 | 44 | r2 | 0.0691303 |
| regression | DFINE | LINK CO | cross_day | 20210104__to__20210111 | 20210227 | 44 | r2 | 0.0512044 |
| regression | DFINE | LINK CO | cross_day | 20210104__to__20210111 | 20210309 | 44 | r2 | 0.0464436 |
| regression | DFINE | LINK CO | cross_day | 20210104__to__20210111 | 20210313 | 44 | r2 | -0.0136222 |
| regression | DFINE | LINK CO | cross_day | 20210104__to__20210111 | 20210329 | 44 | r2 | 0.0835767 |
| regression | DFINE | LINK CO | cross_day | 20210104__to__20210111 | 20210330 | 44 | r2 | 0.061935 |
| regression | DFINE | LINK CO | cross_day | 20210104__to__20210111 | 20210402 | 44 | r2 | 0.0135789 |
| regression | DFINE | LINK CO | cross_day | 20210104__to__20210111 | 20210406 | 44 | r2 | 0.0117635 |
| regression | DFINE | LINK CO | cross_day | 20210104__to__20210111 | 20210409 | 44 | r2 | 0.0263099 |
| regression | DFINE | LINK CO | cross_day | 20210104__to__20210111 | 20210415 | 44 | r2 | 0.076992 |
| regression | DFINE | LINK CO | cross_day | 20210104__to__20210111 | 20210420 | 44 | r2 | 3.15309e-05 |
| regression | DFINE | LINK CO | cross_day | 20210104__to__20210111 | 20210503 | 44 | r2 | -0.0190808 |
| regression | DFINE | LINK CO | cross_day | 20210104__to__20210111 | 20210504 | 44 | r2 | 0.00275508 |
| regression | DFINE | LINK CO | cross_day | 20210104__to__20210111 | 20210511 | 44 | r2 | 0.159118 |
| regression | DFINE | LINK CO | cross_day | 20210104__to__20210111 | 20210518 | 44 | r2 | 0.127861 |
| regression | DFINE | LINK CO | cross_day | 20210104__to__20210111 | 20210520 | 44 | r2 | 0.147238 |
| regression | DFINE | LINK CO | cross_day | 20210104__to__20210111 | 20210601 | 44 | r2 | 0.101203 |
| regression | LSTM | LINK RTT | cross_day | 20210305__to__20210322 | 20210412 | 44 | r2 | 0.36361 |
| regression | LSTM | LINK RTT | cross_day | 20210305__to__20210322 | 20210414 | 44 | r2 | -0.365453 |
| regression | LSTM | LINK RTT | cross_day | 20210305__to__20210322 | 20210421 | 44 | r2 | -0.0207578 |
| regression | LSTM | LINK RTT | cross_day | 20210305__to__20210322 | 20210428 | 44 | r2 | 0.268231 |
| regression | LSTM | LINK RTT | cross_day | 20210305__to__20210322 | 20210505 | 44 | r2 | -0.084742 |
| regression | LSTM | LINK RTT | cross_day | 20210305__to__20210322 | 20210519 | 44 | r2 | 0.304149 |
| regression | LSTM | LINK RTT | cross_day | 20210305__to__20210322 | 20210525 | 44 | r2 | -0.745366 |
| regression | LSTM | LINK RTT | cross_day | 20210305__to__20210322 | 20210526 | 44 | r2 | -0.204042 |
| regression | LSTM | LINK RTT | cross_day | 20210305__to__20210322 | 20210604 | 44 | r2 | -0.108275 |
| regression | LSTM | LINK RTT | cross_day | 20210305__to__20210322 | 20210608 | 44 | r2 | 0.133575 |
| regression | LSTM | LINK RTT | cross_day | 20210305__to__20210322 | 20210616 | 44 | r2 | 0.0924142 |
| regression | LSTM | LINK RTT | cross_day | 20210305__to__20210322 | 20210617 | 44 | r2 | 0.187363 |
| regression | LSTM | LINK RTT | cross_day | 20210305__to__20210322 | 20210624 | 44 | r2 | 0.108491 |
| regression | LSTM | LINK RTT | cross_day | 20210305__to__20210322 | 20210626 | 44 | r2 | 0.228737 |
| regression | LSTM | LINK RTT | cross_day | 20210305__to__20210322 | 20210629 | 44 | r2 | 0.218377 |
| regression | LSTM | LINK RTT | cross_day | 20210305__to__20210322 | 20210630 | 44 | r2 | 0.150254 |
| regression | LSTM | LINK RTT | cross_day | 20210305__to__20210322 | 20210701 | 44 | r2 | 0.0808493 |
| regression | LSTM | LINK RTT | cross_day | 20210305__to__20210322 | 20210703 | 44 | r2 | -0.209451 |
| regression | Transformer | LINK RTT | cross_day | 20210305__to__20210322 | 20210412 | 44 | r2 | 0.168808 |
| regression | Transformer | LINK RTT | cross_day | 20210305__to__20210322 | 20210414 | 44 | r2 | 0.139142 |
| regression | Transformer | LINK RTT | cross_day | 20210305__to__20210322 | 20210421 | 44 | r2 | 0.0237371 |
| regression | Transformer | LINK RTT | cross_day | 20210305__to__20210322 | 20210428 | 44 | r2 | 0.027941 |
| regression | Transformer | LINK RTT | cross_day | 20210305__to__20210322 | 20210505 | 44 | r2 | 0.158689 |
| regression | Transformer | LINK RTT | cross_day | 20210305__to__20210322 | 20210519 | 44 | r2 | 0.210895 |
| regression | Transformer | LINK RTT | cross_day | 20210305__to__20210322 | 20210525 | 44 | r2 | 0.127801 |
| regression | Transformer | LINK RTT | cross_day | 20210305__to__20210322 | 20210526 | 44 | r2 | 0.198582 |
| regression | Transformer | LINK RTT | cross_day | 20210305__to__20210322 | 20210604 | 44 | r2 | 0.223465 |
| regression | Transformer | LINK RTT | cross_day | 20210305__to__20210322 | 20210608 | 44 | r2 | 0.0863834 |
| regression | Transformer | LINK RTT | cross_day | 20210305__to__20210322 | 20210616 | 44 | r2 | 0.163117 |

| | | | | | | | | |
|---|---|---|---|---|---|---|---|---|
| regression | Transformer | LINK RTT | cross_day | 20210305__to__20210322 | 20210617 | 44 | r2 | 0.127492 |
| regression | Transformer | LINK RTT | cross_day | 20210305__to__20210322 | 20210624 | 44 | r2 | 0.139301 |
| regression | Transformer | LINK RTT | cross_day | 20210305__to__20210322 | 20210626 | 44 | r2 | 0.1358 |
| regression | Transformer | LINK RTT | cross_day | 20210305__to__20210322 | 20210629 | 44 | r2 | 0.174265 |
| regression | Transformer | LINK RTT | cross_day | 20210305__to__20210322 | 20210630 | 44 | r2 | 0.183141 |
| regression | Transformer | LINK RTT | cross_day | 20210305__to__20210322 | 20210701 | 44 | r2 | 0.18318 |
| regression | Transformer | LINK RTT | cross_day | 20210305__to__20210322 | 20210703 | 44 | r2 | 0.14929 |
| regression | LFADS | LINK RTT | cross_day | 20210305__to__20210322 | 20210412 | 44 | r2 | 0.00400373 |
| regression | LFADS | LINK RTT | cross_day | 20210305__to__20210322 | 20210414 | 44 | r2 | -0.00336504 |
| regression | LFADS | LINK RTT | cross_day | 20210305__to__20210322 | 20210421 | 44 | r2 | -0.025605 |
| regression | LFADS | LINK RTT | cross_day | 20210305__to__20210322 | 20210428 | 44 | r2 | -0.0154891 |
| regression | LFADS | LINK RTT | cross_day | 20210305__to__20210322 | 20210505 | 44 | r2 | -0.00302991 |
| regression | LFADS | LINK RTT | cross_day | 20210305__to__20210322 | 20210519 | 44 | r2 | -0.00445929 |
| regression | LFADS | LINK RTT | cross_day | 20210305__to__20210322 | 20210525 | 44 | r2 | -0.00496066 |
| regression | LFADS | LINK RTT | cross_day | 20210305__to__20210322 | 20210526 | 44 | r2 | -0.000525475 |
| regression | LFADS | LINK RTT | cross_day | 20210305__to__20210322 | 20210604 | 44 | r2 | -0.00164649 |
| regression | LFADS | LINK RTT | cross_day | 20210305__to__20210322 | 20210608 | 44 | r2 | -0.0224053 |
| regression | LFADS | LINK RTT | cross_day | 20210305__to__20210322 | 20210616 | 44 | r2 | -0.00350609 |
| regression | LFADS | LINK RTT | cross_day | 20210305__to__20210322 | 20210617 | 44 | r2 | 0.00392056 |
| regression | LFADS | LINK RTT | cross_day | 20210305__to__20210322 | 20210624 | 44 | r2 | -0.0062772 |
| regression | LFADS | LINK RTT | cross_day | 20210305__to__20210322 | 20210626 | 44 | r2 | -0.000312418 |
| regression | LFADS | LINK RTT | cross_day | 20210305__to__20210322 | 20210629 | 44 | r2 | 0.00170395 |
| regression | LFADS | LINK RTT | cross_day | 20210305__to__20210322 | 20210630 | 44 | r2 | -0.00113064 |
| regression | LFADS | LINK RTT | cross_day | 20210305__to__20210322 | 20210701 | 44 | r2 | 0.00274864 |
| regression | LFADS | LINK RTT | cross_day | 20210305__to__20210322 | 20210703 | 44 | r2 | 0.00362578 |
| regression | Cycle-GAN | LINK RTT | cross_day | 20210305__to__20210322 | 20210412 | 44 | r2 | -0.0218894 |
| regression | Cycle-GAN | LINK RTT | cross_day | 20210305__to__20210322 | 20210414 | 44 | r2 | -0.0108796 |
| regression | Cycle-GAN | LINK RTT | cross_day | 20210305__to__20210322 | 20210421 | 44 | r2 | -0.0276461 |
| regression | Cycle-GAN | LINK RTT | cross_day | 20210305__to__20210322 | 20210428 | 44 | r2 | -0.0166493 |
| regression | Cycle-GAN | LINK RTT | cross_day | 20210305__to__20210322 | 20210505 | 44 | r2 | -0.0828091 |
| regression | Cycle-GAN | LINK RTT | cross_day | 20210305__to__20210322 | 20210519 | 44 | r2 | 0.0558698 |
| regression | Cycle-GAN | LINK RTT | cross_day | 20210305__to__20210322 | 20210525 | 44 | r2 | -0.0791968 |
| regression | Cycle-GAN | LINK RTT | cross_day | 20210305__to__20210322 | 20210526 | 44 | r2 | 0.0407389 |
| regression | Cycle-GAN | LINK RTT | cross_day | 20210305__to__20210322 | 20210604 | 44 | r2 | 0.0181526 |
| regression | Cycle-GAN | LINK RTT | cross_day | 20210305__to__20210322 | 20210608 | 44 | r2 | 0.0629203 |
| regression | Cycle-GAN | LINK RTT | cross_day | 20210305__to__20210322 | 20210616 | 44 | r2 | 0.030329 |
| regression | Cycle-GAN | LINK RTT | cross_day | 20210305__to__20210322 | 20210617 | 44 | r2 | 0.0789772 |
| regression | Cycle-GAN | LINK RTT | cross_day | 20210305__to__20210322 | 20210624 | 44 | r2 | 0.0557829 |
| regression | Cycle-GAN | LINK RTT | cross_day | 20210305__to__20210322 | 20210626 | 44 | r2 | -0.00766853 |
| regression | Cycle-GAN | LINK RTT | cross_day | 20210305__to__20210322 | 20210629 | 44 | r2 | 0.0390834 |
| regression | Cycle-GAN | LINK RTT | cross_day | 20210305__to__20210322 | 20210630 | 44 | r2 | 0.0609642 |
| regression | Cycle-GAN | LINK RTT | cross_day | 20210305__to__20210322 | 20210701 | 44 | r2 | 0.0489943 |
| regression | Cycle-GAN | LINK RTT | cross_day | 20210305__to__20210322 | 20210703 | 44 | r2 | 0.0325692 |
| regression | Stabilization | LINK RTT | cross_day | 20210305__to__20210322 | 20210412 | 44 | r2 | 0.0137867 |
| regression | Stabilization | LINK RTT | cross_day | 20210305__to__20210322 | 20210414 | 44 | r2 | 0.0165355 |
| regression | Stabilization | LINK RTT | cross_day | 20210305__to__20210322 | 20210421 | 44 | r2 | 0.00148211 |
| regression | Stabilization | LINK RTT | cross_day | 20210305__to__20210322 | 20210428 | 44 | r2 | -0.0239331 |
| regression | Stabilization | LINK RTT | cross_day | 20210305__to__20210322 | 20210505 | 44 | r2 | 0.028447 |
| regression | Stabilization | LINK RTT | cross_day | 20210305__to__20210322 | 20210519 | 44 | r2 | 0.0707525 |
| regression | Stabilization | LINK RTT | cross_day | 20210305__to__20210322 | 20210525 | 44 | r2 | 0.0107163 |
| regression | Stabilization | LINK RTT | cross_day | 20210305__to__20210322 | 20210526 | 44 | r2 | 0.0488347 |
| regression | Stabilization | LINK RTT | cross_day | 20210305__to__20210322 | 20210604 | 44 | r2 | 0.0753134 |
| regression | Stabilization | LINK RTT | cross_day | 20210305__to__20210322 | 20210608 | 44 | r2 | 0.0186165 |
| regression | Stabilization | LINK RTT | cross_day | 20210305__to__20210322 | 20210616 | 44 | r2 | 0.0582023 |
| regression | Stabilization | LINK RTT | cross_day | 20210305__to__20210322 | 20210617 | 44 | r2 | 0.0446551 |
| regression | Stabilization | LINK RTT | cross_day | 20210305__to__20210322 | 20210624 | 44 | r2 | 0.0315138 |
| regression | Stabilization | LINK RTT | cross_day | 20210305__to__20210322 | 20210626 | 44 | r2 | 0.0344144 |
| regression | Stabilization | LINK RTT | cross_day | 20210305__to__20210322 | 20210629 | 44 | r2 | 0.0466064 |
| regression | Stabilization | LINK RTT | cross_day | 20210305__to__20210322 | 20210630 | 44 | r2 | 0.0176325 |
| regression | Stabilization | LINK RTT | cross_day | 20210305__to__20210322 | 20210701 | 44 | r2 | 0.0366368 |
| regression | Stabilization | LINK RTT | cross_day | 20210305__to__20210322 | 20210703 | 44 | r2 | 0.0300473 |
| regression | FENet | LINK RTT | cross_day | 20210305__to__20210322 | 20210412 | 44 | r2 | 0.340752 |
| regression | FENet | LINK RTT | cross_day | 20210305__to__20210322 | 20210414 | 44 | r2 | 0.26175 |

| | | | | | | | | |
|---|---|---|---|---|---|---|---|---|
| regression | FENet | LINK RTT | cross_day | 20210305__to__20210322 | 20210421 | 44 | r2 | 0.0647864 |
| regression | FENet | LINK RTT | cross_day | 20210305__to__20210322 | 20210428 | 44 | r2 | 0.181949 |
| regression | FENet | LINK RTT | cross_day | 20210305__to__20210322 | 20210505 | 44 | r2 | 0.186428 |
| regression | FENet | LINK RTT | cross_day | 20210305__to__20210322 | 20210519 | 44 | r2 | 0.369551 |
| regression | FENet | LINK RTT | cross_day | 20210305__to__20210322 | 20210525 | 44 | r2 | 0.245984 |
| regression | FENet | LINK RTT | cross_day | 20210305__to__20210322 | 20210526 | 44 | r2 | 0.300345 |
| regression | FENet | LINK RTT | cross_day | 20210305__to__20210322 | 20210604 | 44 | r2 | 0.338878 |
| regression | FENet | LINK RTT | cross_day | 20210305__to__20210322 | 20210608 | 44 | r2 | 0.141789 |
| regression | FENet | LINK RTT | cross_day | 20210305__to__20210322 | 20210616 | 44 | r2 | 0.242499 |
| regression | FENet | LINK RTT | cross_day | 20210305__to__20210322 | 20210617 | 44 | r2 | 0.211676 |
| regression | FENet | LINK RTT | cross_day | 20210305__to__20210322 | 20210624 | 44 | r2 | 0.2347 |
| regression | FENet | LINK RTT | cross_day | 20210305__to__20210322 | 20210626 | 44 | r2 | 0.272014 |
| regression | FENet | LINK RTT | cross_day | 20210305__to__20210322 | 20210629 | 44 | r2 | 0.219859 |
| regression | FENet | LINK RTT | cross_day | 20210305__to__20210322 | 20210630 | 44 | r2 | 0.270075 |
| regression | FENet | LINK RTT | cross_day | 20210305__to__20210322 | 20210701 | 44 | r2 | 0.240729 |
| regression | FENet | LINK RTT | cross_day | 20210305__to__20210322 | 20210703 | 44 | r2 | 0.196697 |
| regression | DFINE | LINK RTT | cross_day | 20210305__to__20210322 | 20210412 | 44 | r2 | 0.228338 |
| regression | DFINE | LINK RTT | cross_day | 20210305__to__20210322 | 20210414 | 44 | r2 | 0.151519 |
| regression | DFINE | LINK RTT | cross_day | 20210305__to__20210322 | 20210421 | 44 | r2 | 0.0351931 |
| regression | DFINE | LINK RTT | cross_day | 20210305__to__20210322 | 20210428 | 44 | r2 | 0.087609 |
| regression | DFINE | LINK RTT | cross_day | 20210305__to__20210322 | 20210505 | 44 | r2 | 0.117561 |
| regression | DFINE | LINK RTT | cross_day | 20210305__to__20210322 | 20210519 | 44 | r2 | 0.245047 |
| regression | DFINE | LINK RTT | cross_day | 20210305__to__20210322 | 20210525 | 44 | r2 | 0.274508 |
| regression | DFINE | LINK RTT | cross_day | 20210305__to__20210322 | 20210526 | 44 | r2 | 0.272271 |
| regression | DFINE | LINK RTT | cross_day | 20210305__to__20210322 | 20210604 | 44 | r2 | 0.299386 |
| regression | DFINE | LINK RTT | cross_day | 20210305__to__20210322 | 20210608 | 44 | r2 | 0.0842716 |
| regression | DFINE | LINK RTT | cross_day | 20210305__to__20210322 | 20210616 | 44 | r2 | 0.262258 |
| regression | DFINE | LINK RTT | cross_day | 20210305__to__20210322 | 20210617 | 44 | r2 | 0.231775 |
| regression | DFINE | LINK RTT | cross_day | 20210305__to__20210322 | 20210624 | 44 | r2 | 0.204139 |
| regression | DFINE | LINK RTT | cross_day | 20210305__to__20210322 | 20210626 | 44 | r2 | 0.31216 |
| regression | DFINE | LINK RTT | cross_day | 20210305__to__20210322 | 20210629 | 44 | r2 | 0.248405 |
| regression | DFINE | LINK RTT | cross_day | 20210305__to__20210322 | 20210630 | 44 | r2 | 0.285454 |
| regression | DFINE | LINK RTT | cross_day | 20210305__to__20210322 | 20210701 | 44 | r2 | 0.201907 |
| regression | DFINE | LINK RTT | cross_day | 20210305__to__20210322 | 20210703 | 44 | r2 | 0.230937 |
| regression | LSTM | Indy | cross_day | 20160418__to__20160622 | 20160630 | 44 | r2 | -0.00201088 |
| regression | LSTM | Indy | cross_day | 20160418__to__20160622 | 20160915 | 44 | r2 | 0.248038 |
| regression | LSTM | Indy | cross_day | 20160418__to__20160622 | 20160916 | 44 | r2 | 0.455376 |
| regression | LSTM | Indy | cross_day | 20160418__to__20160622 | 20160921 | 44 | r2 | 0.474134 |
| regression | LSTM | Indy | cross_day | 20160418__to__20160622 | 20160927 | 44 | r2 | 0.397523 |
| regression | LSTM | Indy | cross_day | 20160418__to__20160622 | 20160927 | 44 | r2 | 0.141076 |
| regression | LSTM | Indy | cross_day | 20160418__to__20160622 | 20160930 | 44 | r2 | 0.321428 |
| regression | LSTM | Indy | cross_day | 20160418__to__20160622 | 20160930 | 44 | r2 | 0.0556978 |
| regression | LSTM | Indy | cross_day | 20160418__to__20160622 | 20161005 | 44 | r2 | 0.218908 |
| regression | LSTM | Indy | cross_day | 20160418__to__20160622 | 20161006 | 44 | r2 | 0.077132 |
| regression | LSTM | Indy | cross_day | 20160418__to__20160622 | 20161007 | 44 | r2 | 0.351406 |
| regression | LSTM | Indy | cross_day | 20160418__to__20160622 | 20161011 | 44 | r2 | 0.260486 |
| regression | LSTM | Indy | cross_day | 20160418__to__20160622 | 20161013 | 44 | r2 | 0.00814357 |
| regression | LSTM | Indy | cross_day | 20160418__to__20160622 | 20161014 | 44 | r2 | 0.36163 |
| regression | LSTM | Indy | cross_day | 20160418__to__20160622 | 20161017 | 44 | r2 | 0.254448 |
| regression | LSTM | Indy | cross_day | 20160418__to__20160622 | 20161024 | 44 | r2 | 0.199862 |
| regression | LSTM | Indy | cross_day | 20160418__to__20160622 | 20161025 | 44 | r2 | -1.31938 |
| regression | LSTM | Indy | cross_day | 20160418__to__20160622 | 20161026 | 44 | r2 | 0.201217 |
| regression | Transformer | Indy | cross_day | 20160418__to__20160622 | 20160630 | 44 | r2 | -0.323174 |
| regression | Transformer | Indy | cross_day | 20160418__to__20160622 | 20160915 | 44 | r2 | 0.116196 |
| regression | Transformer | Indy | cross_day | 20160418__to__20160622 | 20160916 | 44 | r2 | 0.351111 |
| regression | Transformer | Indy | cross_day | 20160418__to__20160622 | 20160921 | 44 | r2 | 0.330069 |
| regression | Transformer | Indy | cross_day | 20160418__to__20160622 | 20160927 | 44 | r2 | 0.325338 |
| regression | Transformer | Indy | cross_day | 20160418__to__20160622 | 20160927 | 44 | r2 | 0.187373 |
| regression | Transformer | Indy | cross_day | 20160418__to__20160622 | 20160930 | 44 | r2 | 0.146258 |
| regression | Transformer | Indy | cross_day | 20160418__to__20160622 | 20160930 | 44 | r2 | 0.0328398 |
| regression | Transformer | Indy | cross_day | 20160418__to__20160622 | 20161005 | 44 | r2 | 0.216984 |
| regression | Transformer | Indy | cross_day | 20160418__to__20160622 | 20161006 | 44 | r2 | 0.0737364 |
| regression | Transformer | Indy | cross_day | 20160418__to__20160622 | 20161007 | 44 | r2 | 0.221176 |

| | | | | | | | | |
|---|---|---|---|---|---|---|---|---|
| regression | Transformer | Indy | cross_day | 20160418__to__20160622 | 20161011 | 44 | r2 | 0.128632 |
| regression | Transformer | Indy | cross_day | 20160418__to__20160622 | 20161013 | 44 | r2 | -0.116647 |
| regression | Transformer | Indy | cross_day | 20160418__to__20160622 | 20161014 | 44 | r2 | 0.247003 |
| regression | Transformer | Indy | cross_day | 20160418__to__20160622 | 20161017 | 44 | r2 | 0.138783 |
| regression | Transformer | Indy | cross_day | 20160418__to__20160622 | 20161024 | 44 | r2 | 0.100832 |
| regression | Transformer | Indy | cross_day | 20160418__to__20160622 | 20161025 | 44 | r2 | -0.454201 |
| regression | Transformer | Indy | cross_day | 20160418__to__20160622 | 20161026 | 44 | r2 | 0.105795 |
| regression | LFADS | Indy | cross_day | 20160418__to__20160622 | 20160630 | 44 | r2 | -0.171879 |
| regression | LFADS | Indy | cross_day | 20160418__to__20160622 | 20160915 | 44 | r2 | 0.0935764 |
| regression | LFADS | Indy | cross_day | 20160418__to__20160622 | 20160916 | 44 | r2 | 0.144676 |
| regression | LFADS | Indy | cross_day | 20160418__to__20160622 | 20160921 | 44 | r2 | 0.0861428 |
| regression | LFADS | Indy | cross_day | 20160418__to__20160622 | 20160927 | 44 | r2 | 0.10035 |
| regression | LFADS | Indy | cross_day | 20160418__to__20160622 | 20160927 | 44 | r2 | 0.0909157 |
| regression | LFADS | Indy | cross_day | 20160418__to__20160622 | 20160930 | 44 | r2 | 0.0604661 |
| regression | LFADS | Indy | cross_day | 20160418__to__20160622 | 20160930 | 44 | r2 | 0.107493 |
| regression | LFADS | Indy | cross_day | 20160418__to__20160622 | 20161005 | 44 | r2 | 0.0177424 |
| regression | LFADS | Indy | cross_day | 20160418__to__20160622 | 20161006 | 44 | r2 | 0.0806054 |
| regression | LFADS | Indy | cross_day | 20160418__to__20160622 | 20161007 | 44 | r2 | 0.0616133 |
| regression | LFADS | Indy | cross_day | 20160418__to__20160622 | 20161011 | 44 | r2 | 0.0131089 |
| regression | LFADS | Indy | cross_day | 20160418__to__20160622 | 20161013 | 44 | r2 | -0.0899647 |
| regression | LFADS | Indy | cross_day | 20160418__to__20160622 | 20161014 | 44 | r2 | 0.0644026 |
| regression | LFADS | Indy | cross_day | 20160418__to__20160622 | 20161017 | 44 | r2 | 0.0297894 |
| regression | LFADS | Indy | cross_day | 20160418__to__20160622 | 20161024 | 44 | r2 | 0.025188 |
| regression | LFADS | Indy | cross_day | 20160418__to__20160622 | 20161025 | 44 | r2 | 0.0583931 |
| regression | LFADS | Indy | cross_day | 20160418__to__20160622 | 20161026 | 44 | r2 | 0.0636937 |
| regression | Cycle-GAN | Indy | cross_day | 20160418__to__20160622 | 20160630 | 44 | r2 | -0.0024083 |
| regression | Cycle-GAN | Indy | cross_day | 20160418__to__20160622 | 20160915 | 44 | r2 | 0.0744658 |
| regression | Cycle-GAN | Indy | cross_day | 20160418__to__20160622 | 20160916 | 44 | r2 | 0.1645 |
| regression | Cycle-GAN | Indy | cross_day | 20160418__to__20160622 | 20160921 | 44 | r2 | 0.162079 |
| regression | Cycle-GAN | Indy | cross_day | 20160418__to__20160622 | 20160927 | 44 | r2 | 0.125684 |
| regression | Cycle-GAN | Indy | cross_day | 20160418__to__20160622 | 20160927 | 44 | r2 | 0.0866755 |
| regression | Cycle-GAN | Indy | cross_day | 20160418__to__20160622 | 20160930 | 44 | r2 | 0.0474066 |
| regression | Cycle-GAN | Indy | cross_day | 20160418__to__20160622 | 20160930 | 44 | r2 | 0.0202571 |
| regression | Cycle-GAN | Indy | cross_day | 20160418__to__20160622 | 20161005 | 44 | r2 | 0.0136131 |
| regression | Cycle-GAN | Indy | cross_day | 20160418__to__20160622 | 20161006 | 44 | r2 | 0.0193144 |
| regression | Cycle-GAN | Indy | cross_day | 20160418__to__20160622 | 20161007 | 44 | r2 | 0.0405895 |
| regression | Cycle-GAN | Indy | cross_day | 20160418__to__20160622 | 20161011 | 44 | r2 | 0.0527747 |
| regression | Cycle-GAN | Indy | cross_day | 20160418__to__20160622 | 20161013 | 44 | r2 | 0.000140871 |
| regression | Cycle-GAN | Indy | cross_day | 20160418__to__20160622 | 20161014 | 44 | r2 | 0.0783987 |
| regression | Cycle-GAN | Indy | cross_day | 20160418__to__20160622 | 20161017 | 44 | r2 | 0.0272866 |
| regression | Cycle-GAN | Indy | cross_day | 20160418__to__20160622 | 20161024 | 44 | r2 | 0.0223172 |
| regression | Cycle-GAN | Indy | cross_day | 20160418__to__20160622 | 20161025 | 44 | r2 | 0.0239181 |
| regression | Cycle-GAN | Indy | cross_day | 20160418__to__20160622 | 20161026 | 44 | r2 | 0.0433266 |
| regression | Stabilization | Indy | cross_day | 20160418__to__20160622 | 20160630 | 44 | r2 | 0.407089 |
| regression | Stabilization | Indy | cross_day | 20160418__to__20160622 | 20160915 | 44 | r2 | 0.333046 |
| regression | Stabilization | Indy | cross_day | 20160418__to__20160622 | 20160916 | 44 | r2 | 0.37578 |
| regression | Stabilization | Indy | cross_day | 20160418__to__20160622 | 20160921 | 44 | r2 | 0.34945 |
| regression | Stabilization | Indy | cross_day | 20160418__to__20160622 | 20160927 | 44 | r2 | 0.346949 |
| regression | Stabilization | Indy | cross_day | 20160418__to__20160622 | 20160927 | 44 | r2 | 0.343404 |
| regression | Stabilization | Indy | cross_day | 20160418__to__20160622 | 20160930 | 44 | r2 | 0.292824 |
| regression | Stabilization | Indy | cross_day | 20160418__to__20160622 | 20160930 | 44 | r2 | 0.336062 |
| regression | Stabilization | Indy | cross_day | 20160418__to__20160622 | 20161005 | 44 | r2 | 0.284864 |
| regression | Stabilization | Indy | cross_day | 20160418__to__20160622 | 20161006 | 44 | r2 | 0.22173 |
| regression | Stabilization | Indy | cross_day | 20160418__to__20160622 | 20161007 | 44 | r2 | 0.230313 |
| regression | Stabilization | Indy | cross_day | 20160418__to__20160622 | 20161011 | 44 | r2 | 0.235923 |
| regression | Stabilization | Indy | cross_day | 20160418__to__20160622 | 20161013 | 44 | r2 | 0.251379 |
| regression | Stabilization | Indy | cross_day | 20160418__to__20160622 | 20161014 | 44 | r2 | 0.353857 |
| regression | Stabilization | Indy | cross_day | 20160418__to__20160622 | 20161017 | 44 | r2 | 0.27955 |
| regression | Stabilization | Indy | cross_day | 20160418__to__20160622 | 20161024 | 44 | r2 | 0.268395 |
| regression | Stabilization | Indy | cross_day | 20160418__to__20160622 | 20161025 | 44 | r2 | 0.307411 |
| regression | Stabilization | Indy | cross_day | 20160418__to__20160622 | 20161026 | 44 | r2 | 0.283654 |
| regression | FENet | Indy | cross_day | 20160418__to__20160622 | 20160630 | 44 | r2 | -0.0143867 |
| regression | FENet | Indy | cross_day | 20160418__to__20160622 | 20160915 | 44 | r2 | 0.27263 |

| | | | | | | | | |
|---|---|---|---|---|---|---|---|---|
| regression | FENet | Indy | cross_day | 20160418__to__20160622 | 20160916 | 44 | r2 | 0.312919 |
| regression | FENet | Indy | cross_day | 20160418__to__20160622 | 20160921 | 44 | r2 | 0.281539 |
| regression | FENet | Indy | cross_day | 20160418__to__20160622 | 20160927 | 44 | r2 | 0.232261 |
| regression | FENet | Indy | cross_day | 20160418__to__20160622 | 20160927 | 44 | r2 | 0.195227 |
| regression | FENet | Indy | cross_day | 20160418__to__20160622 | 20160930 | 44 | r2 | 0.228701 |
| regression | FENet | Indy | cross_day | 20160418__to__20160622 | 20160930 | 44 | r2 | 0.209185 |
| regression | FENet | Indy | cross_day | 20160418__to__20160622 | 20161005 | 44 | r2 | 0.126536 |
| regression | FENet | Indy | cross_day | 20160418__to__20160622 | 20161006 | 44 | r2 | 0.188601 |
| regression | FENet | Indy | cross_day | 20160418__to__20160622 | 20161007 | 44 | r2 | 0.227073 |
| regression | FENet | Indy | cross_day | 20160418__to__20160622 | 20161011 | 44 | r2 | 0.18476 |
| regression | FENet | Indy | cross_day | 20160418__to__20160622 | 20161013 | 44 | r2 | 0.0177647 |
| regression | FENet | Indy | cross_day | 20160418__to__20160622 | 20161014 | 44 | r2 | 0.291773 |
| regression | FENet | Indy | cross_day | 20160418__to__20160622 | 20161017 | 44 | r2 | 0.176034 |
| regression | FENet | Indy | cross_day | 20160418__to__20160622 | 20161024 | 44 | r2 | 0.167238 |
| regression | FENet | Indy | cross_day | 20160418__to__20160622 | 20161025 | 44 | r2 | -0.00930917 |
| regression | FENet | Indy | cross_day | 20160418__to__20160622 | 20161026 | 44 | r2 | 0.213588 |
| regression | DFINE | Indy | cross_day | 20160418__to__20160622 | 20160630 | 44 | r2 | -0.00221771 |
| regression | DFINE | Indy | cross_day | 20160418__to__20160622 | 20160915 | 44 | r2 | 0.0804488 |
| regression | DFINE | Indy | cross_day | 20160418__to__20160622 | 20160916 | 44 | r2 | 0.17643 |
| regression | DFINE | Indy | cross_day | 20160418__to__20160622 | 20160921 | 44 | r2 | 0.16574 |
| regression | DFINE | Indy | cross_day | 20160418__to__20160622 | 20160927 | 44 | r2 | -0.684535 |
| regression | DFINE | Indy | cross_day | 20160418__to__20160622 | 20160927 | 44 | r2 | -0.721079 |
| regression | DFINE | Indy | cross_day | 20160418__to__20160622 | 20160930 | 44 | r2 | 0.0385812 |
| regression | DFINE | Indy | cross_day | 20160418__to__20160622 | 20160930 | 44 | r2 | 0.0824206 |
| regression | DFINE | Indy | cross_day | 20160418__to__20160622 | 20161005 | 44 | r2 | -0.539513 |
| regression | DFINE | Indy | cross_day | 20160418__to__20160622 | 20161006 | 44 | r2 | 0.0837018 |
| regression | DFINE | Indy | cross_day | 20160418__to__20160622 | 20161007 | 44 | r2 | 0.0877905 |
| regression | DFINE | Indy | cross_day | 20160418__to__20160622 | 20161011 | 44 | r2 | 0.0712675 |
| regression | DFINE | Indy | cross_day | 20160418__to__20160622 | 20161013 | 44 | r2 | -0.0028289 |
| regression | DFINE | Indy | cross_day | 20160418__to__20160622 | 20161014 | 44 | r2 | 0.0511863 |
| regression | DFINE | Indy | cross_day | 20160418__to__20160622 | 20161017 | 44 | r2 | 0.126681 |
| regression | DFINE | Indy | cross_day | 20160418__to__20160622 | 20161024 | 44 | r2 | -0.331931 |
| regression | DFINE | Indy | cross_day | 20160418__to__20160622 | 20161025 | 44 | r2 | 0.00257114 |
| regression | DFINE | Indy | cross_day | 20160418__to__20160622 | 20161026 | 44 | r2 | 0.0241018 |
| regression | NoMAD | FALCON M2 | cross_day | 20201019__to__20201020 | 20201030 | 44 | r2 | 0.301495 |
| regression | seqVAE | FALCON M2 | cross_day | 20201019__to__20201020 | 20201030 | 44 | r2 | 0.285369 |
| regression | NoMAD | FALCON M2 | cross_day | 20201019__to__20201020 | 20201030 | 44 | r2 | 0.352724 |
| regression | seqVAE | FALCON M2 | cross_day | 20201019__to__20201020 | 20201030 | 44 | r2 | 0.356611 |
| regression | NoMAD | FALCON M2 | cross_day | 20201019__to__20201020 | 20201118 | 44 | r2 | 0.170311 |
| regression | seqVAE | FALCON M2 | cross_day | 20201019__to__20201020 | 20201118 | 44 | r2 | 0.17382 |
| regression | NoMAD | FALCON M2 | cross_day | 20201019__to__20201020 | 20201119 | 44 | r2 | 0.0404992 |
| regression | seqVAE | FALCON M2 | cross_day | 20201019__to__20201020 | 20201119 | 44 | r2 | 0.0176189 |
| regression | NoMAD | FALCON M2 | cross_day | 20201019__to__20201020 | 20201124 | 44 | r2 | 0.0689135 |
| regression | seqVAE | FALCON M2 | cross_day | 20201019__to__20201020 | 20201124 | 44 | r2 | 0.0554186 |
| regression | NoMAD | FALCON M2 | cross_day | 20201019__to__20201020 | 20201124 | 44 | r2 | 0.0921189 |
| regression | seqVAE | FALCON M2 | cross_day | 20201019__to__20201020 | 20201124 | 44 | r2 | 0.0926863 |
| regression | NoMAD | FALCON M2 | single_day | 20201019 | 20201019 | 44 | r2 | 0.608766 |
| regression | seqVAE | FALCON M2 | single_day | 20201019 | 20201019 | 44 | r2 | 0.475843 |
| regression | NoMAD | FALCON M2 | single_day | 20201019 | 20201019 | 44 | r2 | 0.519187 |
| regression | seqVAE | FALCON M2 | single_day | 20201019 | 20201019 | 44 | r2 | 0.424663 |
| regression | NoMAD | FALCON M2 | single_day | 20201020 | 20201020 | 44 | r2 | 0.544238 |
| regression | seqVAE | FALCON M2 | single_day | 20201020 | 20201020 | 44 | r2 | 0.421571 |
| regression | NoMAD | FALCON M2 | single_day | 20201020 | 20201020 | 44 | r2 | 0.570287 |
| regression | seqVAE | FALCON M2 | single_day | 20201020 | 20201020 | 44 | r2 | 0.44381 |
| regression | NoMAD | FALCON M2 | single_day | 20201027 | 20201027 | 44 | r2 | 0.374841 |
| regression | seqVAE | FALCON M2 | single_day | 20201027 | 20201027 | 44 | r2 | 0.194215 |
| regression | NoMAD | FALCON M2 | single_day | 20201027 | 20201027 | 44 | r2 | 0.506761 |
| regression | seqVAE | FALCON M2 | single_day | 20201027 | 20201027 | 44 | r2 | 0.466839 |
| regression | NoMAD | LINK CO | cross_day | 20210104__to__20210111 | 20210225 | 44 | r2 | 0.434184 |
| regression | seqVAE | LINK CO | cross_day | 20210104__to__20210111 | 20210225 | 44 | r2 | 0.399929 |
| regression | NoMAD | LINK CO | cross_day | 20210104__to__20210111 | 20210226 | 44 | r2 | 0.388783 |
| regression | seqVAE | LINK CO | cross_day | 20210104__to__20210111 | 20210226 | 44 | r2 | 0.407255 |
| regression | NoMAD | LINK CO | cross_day | 20210104__to__20210111 | 20210227 | 44 | r2 | 0.305188 |

| | | | | | | | | |
|---|---|---|---|---|---|---|---|---|
| regression | seqVAE | LINK CO | cross_day | 20210104__to__20210111 | 20210227 | 44 | r2 | 0.32635 |
| regression | NoMAD | LINK CO | cross_day | 20210104__to__20210111 | 20210309 | 44 | r2 | 0.371004 |
| regression | seqVAE | LINK CO | cross_day | 20210104__to__20210111 | 20210309 | 44 | r2 | 0.338548 |
| regression | NoMAD | LINK CO | cross_day | 20210104__to__20210111 | 20210313 | 44 | r2 | 0.321825 |
| regression | seqVAE | LINK CO | cross_day | 20210104__to__20210111 | 20210313 | 44 | r2 | 0.284853 |
| regression | NoMAD | LINK CO | cross_day | 20210104__to__20210111 | 20210329 | 44 | r2 | 0.399922 |
| regression | seqVAE | LINK CO | cross_day | 20210104__to__20210111 | 20210329 | 44 | r2 | 0.352002 |
| regression | NoMAD | LINK CO | cross_day | 20210104__to__20210111 | 20210330 | 44 | r2 | 0.257329 |
| regression | seqVAE | LINK CO | cross_day | 20210104__to__20210111 | 20210330 | 44 | r2 | 0.276489 |
| regression | NoMAD | LINK CO | cross_day | 20210104__to__20210111 | 20210402 | 44 | r2 | 0.323946 |
| regression | seqVAE | LINK CO | cross_day | 20210104__to__20210111 | 20210402 | 44 | r2 | 0.278707 |
| regression | NoMAD | LINK CO | cross_day | 20210104__to__20210111 | 20210406 | 44 | r2 | 0.392404 |
| regression | seqVAE | LINK CO | cross_day | 20210104__to__20210111 | 20210406 | 44 | r2 | 0.344086 |
| regression | NoMAD | LINK CO | cross_day | 20210104__to__20210111 | 20210409 | 44 | r2 | 0.448553 |
| regression | seqVAE | LINK CO | cross_day | 20210104__to__20210111 | 20210409 | 44 | r2 | 0.404811 |
| regression | NoMAD | LINK CO | cross_day | 20210104__to__20210111 | 20210415 | 44 | r2 | 0.332801 |
| regression | seqVAE | LINK CO | cross_day | 20210104__to__20210111 | 20210415 | 44 | r2 | 0.390905 |
| regression | NoMAD | LINK CO | cross_day | 20210104__to__20210111 | 20210420 | 44 | r2 | 0.310842 |
| regression | seqVAE | LINK CO | cross_day | 20210104__to__20210111 | 20210420 | 44 | r2 | 0.309696 |
| regression | NoMAD | LINK CO | cross_day | 20210104__to__20210111 | 20210503 | 44 | r2 | 0.153431 |
| regression | seqVAE | LINK CO | cross_day | 20210104__to__20210111 | 20210503 | 44 | r2 | 0.199001 |
| regression | NoMAD | LINK CO | cross_day | 20210104__to__20210111 | 20210504 | 44 | r2 | 0.0535179 |
| regression | seqVAE | LINK CO | cross_day | 20210104__to__20210111 | 20210504 | 44 | r2 | 0.093131 |
| regression | NoMAD | LINK CO | cross_day | 20210104__to__20210111 | 20210511 | 44 | r2 | 0.32028 |
| regression | seqVAE | LINK CO | cross_day | 20210104__to__20210111 | 20210511 | 44 | r2 | 0.386474 |
| regression | NoMAD | LINK CO | cross_day | 20210104__to__20210111 | 20210518 | 44 | r2 | 0.397811 |
| regression | seqVAE | LINK CO | cross_day | 20210104__to__20210111 | 20210518 | 44 | r2 | 0.400356 |
| regression | NoMAD | LINK CO | cross_day | 20210104__to__20210111 | 20210520 | 44 | r2 | 0.489954 |
| regression | seqVAE | LINK CO | cross_day | 20210104__to__20210111 | 20210520 | 44 | r2 | 0.448068 |
| regression | NoMAD | LINK CO | cross_day | 20210104__to__20210111 | 20210601 | 44 | r2 | 0.491878 |
| regression | seqVAE | LINK CO | cross_day | 20210104__to__20210111 | 20210601 | 44 | r2 | 0.459575 |
| regression | NoMAD | LINK CO | single_day | 20210104 | 20210104 | 44 | r2 | 0.673084 |
| regression | seqVAE | LINK CO | single_day | 20210104 | 20210104 | 44 | r2 | 0.61359 |
| regression | NoMAD | LINK CO | single_day | 20210105 | 20210105 | 44 | r2 | 0.630644 |
| regression | seqVAE | LINK CO | single_day | 20210105 | 20210105 | 44 | r2 | 0.575493 |
| regression | NoMAD | LINK CO | single_day | 20210106 | 20210106 | 44 | r2 | 0.620698 |
| regression | seqVAE | LINK CO | single_day | 20210106 | 20210106 | 44 | r2 | 0.568633 |
| regression | NoMAD | LINK CO | single_day | 20210108 | 20210108 | 44 | r2 | 0.742411 |
| regression | seqVAE | LINK CO | single_day | 20210108 | 20210108 | 44 | r2 | 0.652003 |
| regression | NoMAD | LINK CO | single_day | 20210111 | 20210111 | 44 | r2 | 0.559896 |
| regression | seqVAE | LINK CO | single_day | 20210111 | 20210111 | 44 | r2 | 0.508475 |
| regression | NoMAD | LINK CO | single_day | 20210123 | 20210123 | 44 | r2 | 0.59891 |
| regression | seqVAE | LINK CO | single_day | 20210123 | 20210123 | 44 | r2 | 0.554397 |
| regression | NoMAD | LINK CO | single_day | 20210223 | 20210223 | 44 | r2 | 0.572937 |
| regression | seqVAE | LINK CO | single_day | 20210223 | 20210223 | 44 | r2 | 0.459776 |
| regression | NoMAD | LINK CO | single_day | 20210225 | 20210225 | 44 | r2 | 0.597433 |
| regression | seqVAE | LINK CO | single_day | 20210225 | 20210225 | 44 | r2 | 0.494629 |
| regression | NoMAD | LINK CO | single_day | 20210226 | 20210226 | 44 | r2 | 0.577779 |
| regression | seqVAE | LINK CO | single_day | 20210226 | 20210226 | 44 | r2 | 0.44171 |
| regression | NoMAD | LINK CO | single_day | 20210227 | 20210227 | 44 | r2 | 0.563145 |
| regression | seqVAE | LINK CO | single_day | 20210227 | 20210227 | 44 | r2 | 0.425318 |
| regression | NoMAD | LINK CO | single_day | 20210309 | 20210309 | 44 | r2 | 0.559885 |
| regression | seqVAE | LINK CO | single_day | 20210309 | 20210309 | 44 | r2 | 0.418179 |
| regression | NoMAD | LINK CO | single_day | 20210313 | 20210313 | 44 | r2 | 0.539463 |
| regression | seqVAE | LINK CO | single_day | 20210313 | 20210313 | 44 | r2 | 0.424341 |
| regression | NoMAD | LINK CO | single_day | 20210329 | 20210329 | 44 | r2 | 0.589238 |
| regression | seqVAE | LINK CO | single_day | 20210329 | 20210329 | 44 | r2 | 0.439344 |
| regression | NoMAD | LINK CO | single_day | 20210330 | 20210330 | 44 | r2 | 0.539166 |
| regression | seqVAE | LINK CO | single_day | 20210330 | 20210330 | 44 | r2 | 0.483515 |
| regression | NoMAD | LINK CO | single_day | 20210402 | 20210402 | 44 | r2 | 0.477593 |
| regression | seqVAE | LINK CO | single_day | 20210402 | 20210402 | 44 | r2 | 0.367181 |
| regression | NoMAD | LINK CO | single_day | 20210406 | 20210406 | 44 | r2 | 0.586416 |
| regression | seqVAE | LINK CO | single_day | 20210406 | 20210406 | 44 | r2 | 0.478439 |

| | | | | | | | | |
|---|---|---|---|---|---|---|---|---|
| regression | NoMAD | LINK CO | single_day | 20210409 | 20210409 | 44 | r2 | 0.614081 |
| regression | seqVAE | LINK CO | single_day | 20210409 | 20210409 | 44 | r2 | 0.445256 |
| regression | NoMAD | LINK CO | single_day | 20210415 | 20210415 | 44 | r2 | 0.599084 |
| regression | seqVAE | LINK CO | single_day | 20210415 | 20210415 | 44 | r2 | 0.462964 |
| regression | NoMAD | LINK CO | single_day | 20210420 | 20210420 | 44 | r2 | 0.556291 |
| regression | seqVAE | LINK CO | single_day | 20210420 | 20210420 | 44 | r2 | 0.450761 |
| regression | NoMAD | LINK CO | single_day | 20210503 | 20210503 | 44 | r2 | 0.594677 |
| regression | seqVAE | LINK CO | single_day | 20210503 | 20210503 | 44 | r2 | 0.396986 |
| regression | NoMAD | LINK CO | single_day | 20210504 | 20210504 | 44 | r2 | 0.584506 |
| regression | seqVAE | LINK CO | single_day | 20210504 | 20210504 | 44 | r2 | 0.472606 |
| regression | NoMAD | LINK CO | single_day | 20210511 | 20210511 | 44 | r2 | 0.631486 |
| regression | seqVAE | LINK CO | single_day | 20210511 | 20210511 | 44 | r2 | 0.572313 |
| regression | NoMAD | LINK CO | single_day | 20210518 | 20210518 | 44 | r2 | 0.692602 |
| regression | seqVAE | LINK CO | single_day | 20210518 | 20210518 | 44 | r2 | 0.659197 |
| regression | NoMAD | LINK CO | single_day | 20210520 | 20210520 | 44 | r2 | 0.689123 |
| regression | seqVAE | LINK CO | single_day | 20210520 | 20210520 | 44 | r2 | 0.565408 |
| regression | NoMAD | LINK CO | single_day | 20210601 | 20210601 | 44 | r2 | 0.666391 |
| regression | seqVAE | LINK CO | single_day | 20210601 | 20210601 | 44 | r2 | 0.560831 |
| regression | NoMAD | LINK RTT | cross_day | 20210305__to__20210322 | 20210412 | 44 | r2 | 0.474825 |
| regression | seqVAE | LINK RTT | cross_day | 20210305__to__20210322 | 20210412 | 44 | r2 | 0.494238 |
| regression | NoMAD | LINK RTT | cross_day | 20210305__to__20210322 | 20210414 | 44 | r2 | 0.482624 |
| regression | seqVAE | LINK RTT | cross_day | 20210305__to__20210322 | 20210414 | 44 | r2 | 0.526152 |
| regression | NoMAD | LINK RTT | cross_day | 20210305__to__20210322 | 20210421 | 44 | r2 | 0.233136 |
| regression | seqVAE | LINK RTT | cross_day | 20210305__to__20210322 | 20210421 | 44 | r2 | 0.263927 |
| regression | NoMAD | LINK RTT | cross_day | 20210305__to__20210322 | 20210428 | 44 | r2 | 0.470305 |
| regression | seqVAE | LINK RTT | cross_day | 20210305__to__20210322 | 20210428 | 44 | r2 | 0.459791 |
| regression | NoMAD | LINK RTT | cross_day | 20210305__to__20210322 | 20210505 | 44 | r2 | 0.387672 |
| regression | seqVAE | LINK RTT | cross_day | 20210305__to__20210322 | 20210505 | 44 | r2 | 0.397343 |
| regression | NoMAD | LINK RTT | cross_day | 20210305__to__20210322 | 20210519 | 44 | r2 | 0.36009 |
| regression | seqVAE | LINK RTT | cross_day | 20210305__to__20210322 | 20210519 | 44 | r2 | 0.410032 |
| regression | NoMAD | LINK RTT | cross_day | 20210305__to__20210322 | 20210525 | 44 | r2 | 0.422131 |
| regression | seqVAE | LINK RTT | cross_day | 20210305__to__20210322 | 20210525 | 44 | r2 | 0.45767 |
| regression | NoMAD | LINK RTT | cross_day | 20210305__to__20210322 | 20210526 | 44 | r2 | 0.518705 |
| regression | seqVAE | LINK RTT | cross_day | 20210305__to__20210322 | 20210526 | 44 | r2 | 0.507213 |
| regression | NoMAD | LINK RTT | cross_day | 20210305__to__20210322 | 20210604 | 44 | r2 | 0.52678 |
| regression | seqVAE | LINK RTT | cross_day | 20210305__to__20210322 | 20210604 | 44 | r2 | 0.52003 |
| regression | NoMAD | LINK RTT | cross_day | 20210305__to__20210322 | 20210608 | 44 | r2 | 0.325096 |
| regression | seqVAE | LINK RTT | cross_day | 20210305__to__20210322 | 20210608 | 44 | r2 | 0.291022 |
| regression | NoMAD | LINK RTT | cross_day | 20210305__to__20210322 | 20210616 | 44 | r2 | 0.327394 |
| regression | seqVAE | LINK RTT | cross_day | 20210305__to__20210322 | 20210616 | 44 | r2 | 0.328802 |
| regression | NoMAD | LINK RTT | cross_day | 20210305__to__20210322 | 20210617 | 44 | r2 | 0.393204 |
| regression | seqVAE | LINK RTT | cross_day | 20210305__to__20210322 | 20210617 | 44 | r2 | 0.391923 |
| regression | NoMAD | LINK RTT | cross_day | 20210305__to__20210322 | 20210624 | 44 | r2 | 0.415159 |
| regression | seqVAE | LINK RTT | cross_day | 20210305__to__20210322 | 20210624 | 44 | r2 | 0.417427 |
| regression | NoMAD | LINK RTT | cross_day | 20210305__to__20210322 | 20210626 | 44 | r2 | 0.394194 |
| regression | seqVAE | LINK RTT | cross_day | 20210305__to__20210322 | 20210626 | 44 | r2 | 0.363338 |
| regression | NoMAD | LINK RTT | cross_day | 20210305__to__20210322 | 20210629 | 44 | r2 | 0.378102 |
| regression | seqVAE | LINK RTT | cross_day | 20210305__to__20210322 | 20210629 | 44 | r2 | 0.335828 |
| regression | NoMAD | LINK RTT | cross_day | 20210305__to__20210322 | 20210630 | 44 | r2 | 0.522762 |
| regression | seqVAE | LINK RTT | cross_day | 20210305__to__20210322 | 20210630 | 44 | r2 | 0.470143 |
| regression | NoMAD | LINK RTT | cross_day | 20210305__to__20210322 | 20210701 | 44 | r2 | 0.485547 |
| regression | seqVAE | LINK RTT | cross_day | 20210305__to__20210322 | 20210701 | 44 | r2 | 0.471514 |
| regression | NoMAD | LINK RTT | cross_day | 20210305__to__20210322 | 20210703 | 44 | r2 | 0.405797 |
| regression | seqVAE | LINK RTT | cross_day | 20210305__to__20210322 | 20210703 | 44 | r2 | 0.342053 |
| regression | NoMAD | LINK RTT | single_day | 20210305 | 20210305 | 44 | r2 | 0.617064 |
| regression | seqVAE | LINK RTT | single_day | 20210305 | 20210305 | 44 | r2 | 0.612998 |
| regression | NoMAD | LINK RTT | single_day | 20210312 | 20210312 | 44 | r2 | 0.557834 |
| regression | seqVAE | LINK RTT | single_day | 20210312 | 20210312 | 44 | r2 | 0.490607 |
| regression | NoMAD | LINK RTT | single_day | 20210316 | 20210316 | 44 | r2 | 0.54628 |
| regression | seqVAE | LINK RTT | single_day | 20210316 | 20210316 | 44 | r2 | 0.509578 |
| regression | NoMAD | LINK RTT | single_day | 20210319 | 20210319 | 44 | r2 | 0.536267 |
| regression | seqVAE | LINK RTT | single_day | 20210319 | 20210319 | 44 | r2 | 0.467337 |
| regression | NoMAD | LINK RTT | single_day | 20210322 | 20210322 | 44 | r2 | 0.527818 |

| | | | | | | | | |
|---|---|---|---|---|---|---|---|---|
| regression | seqVAE | LINK RTT | single_day | 20210322 | 20210322 | 44 | r2 | 0.500392 |
| regression | NoMAD | LINK RTT | single_day | 20210405 | 20210405 | 44 | r2 | 0.557904 |
| regression | seqVAE | LINK RTT | single_day | 20210405 | 20210405 | 44 | r2 | 0.521715 |
| regression | NoMAD | LINK RTT | single_day | 20210407 | 20210407 | 44 | r2 | 0.589546 |
| regression | seqVAE | LINK RTT | single_day | 20210407 | 20210407 | 44 | r2 | 0.564096 |
| regression | NoMAD | LINK RTT | single_day | 20210412 | 20210412 | 44 | r2 | 0.55239 |
| regression | seqVAE | LINK RTT | single_day | 20210412 | 20210412 | 44 | r2 | 0.514104 |
| regression | NoMAD | LINK RTT | single_day | 20210414 | 20210414 | 44 | r2 | 0.615954 |
| regression | seqVAE | LINK RTT | single_day | 20210414 | 20210414 | 44 | r2 | 0.56171 |
| regression | NoMAD | LINK RTT | single_day | 20210421 | 20210421 | 44 | r2 | 0.481196 |
| regression | seqVAE | LINK RTT | single_day | 20210421 | 20210421 | 44 | r2 | 0.348661 |
| regression | NoMAD | LINK RTT | single_day | 20210428 | 20210428 | 44 | r2 | 0.5687 |
| regression | seqVAE | LINK RTT | single_day | 20210428 | 20210428 | 44 | r2 | 0.419345 |
| regression | NoMAD | LINK RTT | single_day | 20210505 | 20210505 | 44 | r2 | 0.57148 |
| regression | seqVAE | LINK RTT | single_day | 20210505 | 20210505 | 44 | r2 | 0.527525 |
| regression | NoMAD | LINK RTT | single_day | 20210519 | 20210519 | 44 | r2 | 0.685725 |
| regression | seqVAE | LINK RTT | single_day | 20210519 | 20210519 | 44 | r2 | 0.656374 |
| regression | NoMAD | LINK RTT | single_day | 20210525 | 20210525 | 44 | r2 | 0.670597 |
| regression | seqVAE | LINK RTT | single_day | 20210525 | 20210525 | 44 | r2 | 0.636756 |
| regression | NoMAD | LINK RTT | single_day | 20210526 | 20210526 | 44 | r2 | 0.638977 |
| regression | seqVAE | LINK RTT | single_day | 20210526 | 20210526 | 44 | r2 | 0.535854 |
| regression | NoMAD | LINK RTT | single_day | 20210604 | 20210604 | 44 | r2 | 0.609144 |
| regression | seqVAE | LINK RTT | single_day | 20210604 | 20210604 | 44 | r2 | 0.582662 |
| regression | NoMAD | LINK RTT | single_day | 20210608 | 20210608 | 44 | r2 | 0.659429 |
| regression | seqVAE | LINK RTT | single_day | 20210608 | 20210608 | 44 | r2 | 0.602656 |
| regression | NoMAD | LINK RTT | single_day | 20210616 | 20210616 | 44 | r2 | 0.693289 |
| regression | seqVAE | LINK RTT | single_day | 20210616 | 20210616 | 44 | r2 | 0.581949 |
| regression | NoMAD | LINK RTT | single_day | 20210617 | 20210617 | 44 | r2 | 0.652886 |
| regression | seqVAE | LINK RTT | single_day | 20210617 | 20210617 | 44 | r2 | 0.609014 |
| regression | NoMAD | LINK RTT | single_day | 20210624 | 20210624 | 44 | r2 | 0.617803 |
| regression | seqVAE | LINK RTT | single_day | 20210624 | 20210624 | 44 | r2 | 0.541713 |
| regression | NoMAD | LINK RTT | single_day | 20210626 | 20210626 | 44 | r2 | 0.621901 |
| regression | seqVAE | LINK RTT | single_day | 20210626 | 20210626 | 44 | r2 | 0.5686 |
| regression | NoMAD | LINK RTT | single_day | 20210629 | 20210629 | 44 | r2 | 0.630689 |
| regression | seqVAE | LINK RTT | single_day | 20210629 | 20210629 | 44 | r2 | 0.568308 |
| regression | NoMAD | LINK RTT | single_day | 20210630 | 20210630 | 44 | r2 | 0.629388 |
| regression | seqVAE | LINK RTT | single_day | 20210630 | 20210630 | 44 | r2 | 0.563454 |
| regression | NoMAD | LINK RTT | single_day | 20210701 | 20210701 | 44 | r2 | 0.636256 |
| regression | seqVAE | LINK RTT | single_day | 20210701 | 20210701 | 44 | r2 | 0.522151 |
| regression | NoMAD | LINK RTT | single_day | 20210703 | 20210703 | 44 | r2 | 0.627665 |
| regression | seqVAE | LINK RTT | single_day | 20210703 | 20210703 | 44 | r2 | 0.568932 |
| regression | NoMAD | Indy | cross_day | 20160418__to__20160622 | 20160630 | 44 | r2 | -0.111046 |
| regression | seqVAE | Indy | cross_day | 20160418__to__20160622 | 20160630 | 44 | r2 | -0.13356 |
| regression | NoMAD | Indy | cross_day | 20160418__to__20160622 | 20160915 | 44 | r2 | 0.539451 |
| regression | seqVAE | Indy | cross_day | 20160418__to__20160622 | 20160915 | 44 | r2 | 0.502752 |
| regression | NoMAD | Indy | cross_day | 20160418__to__20160622 | 20160916 | 44 | r2 | 0.611013 |
| regression | seqVAE | Indy | cross_day | 20160418__to__20160622 | 20160916 | 44 | r2 | 0.608598 |
| regression | NoMAD | Indy | cross_day | 20160418__to__20160622 | 20160921 | 44 | r2 | 0.567633 |
| regression | seqVAE | Indy | cross_day | 20160418__to__20160622 | 20160921 | 44 | r2 | 0.543245 |
| regression | NoMAD | Indy | cross_day | 20160418__to__20160622 | 20160927 | 44 | r2 | 0.521843 |
| regression | seqVAE | Indy | cross_day | 20160418__to__20160622 | 20160927 | 44 | r2 | 0.50026 |
| regression | NoMAD | Indy | cross_day | 20160418__to__20160622 | 20160927 | 44 | r2 | 0.500005 |
| regression | seqVAE | Indy | cross_day | 20160418__to__20160622 | 20160927 | 44 | r2 | 0.474929 |
| regression | NoMAD | Indy | cross_day | 20160418__to__20160622 | 20160930 | 44 | r2 | 0.497385 |
| regression | seqVAE | Indy | cross_day | 20160418__to__20160622 | 20160930 | 44 | r2 | 0.484329 |
| regression | NoMAD | Indy | cross_day | 20160418__to__20160622 | 20160930 | 44 | r2 | 0.465332 |
| regression | seqVAE | Indy | cross_day | 20160418__to__20160622 | 20160930 | 44 | r2 | 0.470033 |
| regression | NoMAD | Indy | cross_day | 20160418__to__20160622 | 20161005 | 44 | r2 | 0.400928 |
| regression | seqVAE | Indy | cross_day | 20160418__to__20160622 | 20161005 | 44 | r2 | 0.385929 |
| regression | NoMAD | Indy | cross_day | 20160418__to__20160622 | 20161006 | 44 | r2 | 0.473465 |
| regression | seqVAE | Indy | cross_day | 20160418__to__20160622 | 20161006 | 44 | r2 | 0.469656 |
| regression | NoMAD | Indy | cross_day | 20160418__to__20160622 | 20161007 | 44 | r2 | 0.421523 |
| regression | seqVAE | Indy | cross_day | 20160418__to__20160622 | 20161007 | 44 | r2 | 0.410988 |

| | | | | | | | | |
|---|---|---|---|---|---|---|---|---|
| regression | NoMAD | Indy | cross_day | 20160418__to__20160622 | 20161011 | 44 | r2 | 0.519567 |
| regression | seqVAE | Indy | cross_day | 20160418__to__20160622 | 20161011 | 44 | r2 | 0.501978 |
| regression | NoMAD | Indy | cross_day | 20160418__to__20160622 | 20161013 | 44 | r2 | -0.0402122 |
| regression | seqVAE | Indy | cross_day | 20160418__to__20160622 | 20161013 | 44 | r2 | -0.0603734 |
| regression | NoMAD | Indy | cross_day | 20160418__to__20160622 | 20161014 | 44 | r2 | 0.560339 |
| regression | seqVAE | Indy | cross_day | 20160418__to__20160622 | 20161014 | 44 | r2 | 0.555242 |
| regression | NoMAD | Indy | cross_day | 20160418__to__20160622 | 20161017 | 44 | r2 | 0.387825 |
| regression | seqVAE | Indy | cross_day | 20160418__to__20160622 | 20161017 | 44 | r2 | 0.369611 |
| regression | NoMAD | Indy | cross_day | 20160418__to__20160622 | 20161024 | 44 | r2 | 0.313252 |
| regression | seqVAE | Indy | cross_day | 20160418__to__20160622 | 20161024 | 44 | r2 | 0.285183 |
| regression | NoMAD | Indy | cross_day | 20160418__to__20160622 | 20161025 | 44 | r2 | 0.491527 |
| regression | seqVAE | Indy | cross_day | 20160418__to__20160622 | 20161025 | 44 | r2 | 0.466525 |
| regression | NoMAD | Indy | cross_day | 20160418__to__20160622 | 20161026 | 44 | r2 | 0.448015 |
| regression | seqVAE | Indy | cross_day | 20160418__to__20160622 | 20161026 | 44 | r2 | 0.41864 |
| regression | NoMAD | Indy | single_day | 20160418 | 20160418 | 44 | r2 | 0.880362 |
| regression | seqVAE | Indy | single_day | 20160418 | 20160418 | 44 | r2 | 0.814591 |
| regression | NoMAD | Indy | single_day | 20160419 | 20160419 | 44 | r2 | 0.847792 |
| regression | seqVAE | Indy | single_day | 20160419 | 20160419 | 44 | r2 | 0.760211 |
| regression | NoMAD | Indy | single_day | 20160420 | 20160420 | 44 | r2 | 0.885856 |
| regression | seqVAE | Indy | single_day | 20160420 | 20160420 | 44 | r2 | 0.837038 |
| regression | NoMAD | Indy | single_day | 20160426 | 20160426 | 44 | r2 | 0.883829 |
| regression | seqVAE | Indy | single_day | 20160426 | 20160426 | 44 | r2 | 0.835791 |
| regression | NoMAD | Indy | single_day | 20160622 | 20160622 | 44 | r2 | 0.882078 |
| regression | seqVAE | Indy | single_day | 20160622 | 20160622 | 44 | r2 | 0.830103 |
| regression | NoMAD | Indy | single_day | 20160624 | 20160624 | 44 | r2 | 0.826053 |
| regression | seqVAE | Indy | single_day | 20160624 | 20160624 | 44 | r2 | 0.736873 |
| regression | NoMAD | Indy | single_day | 20160627 | 20160627 | 44 | r2 | 0.892024 |
| regression | seqVAE | Indy | single_day | 20160627 | 20160627 | 44 | r2 | 0.851682 |
| regression | NoMAD | Indy | single_day | 20160630 | 20160630 | 44 | r2 | 0.853863 |
| regression | seqVAE | Indy | single_day | 20160630 | 20160630 | 44 | r2 | 0.800714 |
| regression | NoMAD | Indy | single_day | 20160915 | 20160915 | 44 | r2 | 0.793817 |
| regression | seqVAE | Indy | single_day | 20160915 | 20160915 | 44 | r2 | 0.708469 |
| regression | NoMAD | Indy | single_day | 20160916 | 20160916 | 44 | r2 | 0.842718 |
| regression | seqVAE | Indy | single_day | 20160916 | 20160916 | 44 | r2 | 0.73652 |
| regression | NoMAD | Indy | single_day | 20160921 | 20160921 | 44 | r2 | 0.82869 |
| regression | seqVAE | Indy | single_day | 20160921 | 20160921 | 44 | r2 | 0.74407 |
| regression | NoMAD | Indy | single_day | 20160927 | 20160927 | 44 | r2 | 0.820578 |
| regression | seqVAE | Indy | single_day | 20160927 | 20160927 | 44 | r2 | 0.745211 |
| regression | NoMAD | Indy | single_day | 20160927 | 20160927 | 44 | r2 | 0.847234 |
| regression | seqVAE | Indy | single_day | 20160927 | 20160927 | 44 | r2 | 0.751819 |
| regression | NoMAD | Indy | single_day | 20160930 | 20160930 | 44 | r2 | 0.818916 |
| regression | seqVAE | Indy | single_day | 20160930 | 20160930 | 44 | r2 | 0.734384 |
| regression | NoMAD | Indy | single_day | 20160930 | 20160930 | 44 | r2 | 0.768404 |
| regression | seqVAE | Indy | single_day | 20160930 | 20160930 | 44 | r2 | 0.678974 |
| regression | NoMAD | Indy | single_day | 20161005 | 20161005 | 44 | r2 | 0.845455 |
| regression | seqVAE | Indy | single_day | 20161005 | 20161005 | 44 | r2 | 0.721106 |
| regression | NoMAD | Indy | single_day | 20161006 | 20161006 | 44 | r2 | 0.833169 |
| regression | seqVAE | Indy | single_day | 20161006 | 20161006 | 44 | r2 | 0.76361 |
| regression | NoMAD | Indy | single_day | 20161007 | 20161007 | 44 | r2 | 0.804189 |
| regression | seqVAE | Indy | single_day | 20161007 | 20161007 | 44 | r2 | 0.746481 |
| regression | NoMAD | Indy | single_day | 20161011 | 20161011 | 44 | r2 | 0.839636 |
| regression | seqVAE | Indy | single_day | 20161011 | 20161011 | 44 | r2 | 0.764661 |
| regression | NoMAD | Indy | single_day | 20161013 | 20161013 | 44 | r2 | 0.837181 |
| regression | seqVAE | Indy | single_day | 20161013 | 20161013 | 44 | r2 | 0.756196 |
| regression | NoMAD | Indy | single_day | 20161014 | 20161014 | 44 | r2 | 0.838952 |
| regression | seqVAE | Indy | single_day | 20161014 | 20161014 | 44 | r2 | 0.759054 |
| regression | NoMAD | Indy | single_day | 20161017 | 20161017 | 44 | r2 | 0.852892 |
| regression | seqVAE | Indy | single_day | 20161017 | 20161017 | 44 | r2 | 0.748719 |
| regression | NoMAD | Indy | single_day | 20161024 | 20161024 | 44 | r2 | 0.831775 |
| regression | seqVAE | Indy | single_day | 20161024 | 20161024 | 44 | r2 | 0.736019 |
| regression | NoMAD | Indy | single_day | 20161025 | 20161025 | 44 | r2 | 0.835974 |
| regression | seqVAE | Indy | single_day | 20161025 | 20161025 | 44 | r2 | 0.746651 |
| regression | NoMAD | Indy | single_day | 20161026 | 20161026 | 44 | r2 | 0.839993 |

| | | | | | | | | |
|---|---|---|---|---|---|---|---|---|
| regression | seqVAE | Indy | single_day | 20161026 | 20161026 | 44 | r2 | 0.749343 |
| regression | KF | Indy | single_day | 20160418 | 20160418 | 44 | r2 | 0.0822149 |
| regression | KF | Indy | single_day | 20160419 | 20160419 | 44 | r2 | 0.0678259 |
| regression | KF | Indy | single_day | 20160420 | 20160420 | 44 | r2 | 0.0681195 |
| regression | KF | Indy | single_day | 20160426 | 20160426 | 44 | r2 | 0.0685835 |
| regression | KF | Indy | single_day | 20160622 | 20160622 | 44 | r2 | 0.0614104 |
| regression | KF | Indy | single_day | 20160624 | 20160624 | 44 | r2 | 0.0788981 |
| regression | KF | Indy | single_day | 20160627 | 20160627 | 44 | r2 | 0.0684208 |
| regression | KF | Indy | single_day | 20160630 | 20160630 | 44 | r2 | 0.039156 |
| regression | KF | Indy | single_day | 20160915 | 20160915 | 44 | r2 | 0.0479508 |
| regression | KF | Indy | single_day | 20160916 | 20160916 | 44 | r2 | 0.0563467 |
| regression | KF | Indy | single_day | 20160921 | 20160921 | 44 | r2 | 0.0433015 |
| regression | KF | Indy | single_day | 20160927 | 20160927 | 44 | r2 | 0.046103 |
| regression | KF | Indy | single_day | 20160927 | 20160927 | 44 | r2 | 0.0411553 |
| regression | KF | Indy | single_day | 20160930 | 20160930 | 44 | r2 | 0.0390882 |
| regression | KF | Indy | single_day | 20160930 | 20160930 | 44 | r2 | 0.0503583 |
| regression | KF | Indy | single_day | 20161005 | 20161005 | 44 | r2 | 0.0383194 |
| regression | KF | Indy | single_day | 20161006 | 20161006 | 44 | r2 | 0.0533272 |
| regression | KF | Indy | single_day | 20161007 | 20161007 | 44 | r2 | 0.0402298 |
| regression | KF | Indy | single_day | 20161011 | 20161011 | 44 | r2 | 0.059258 |
| regression | KF | Indy | single_day | 20161013 | 20161013 | 44 | r2 | 0.0429611 |
| regression | KF | Indy | single_day | 20161014 | 20161014 | 44 | r2 | 0.071239 |
| regression | KF | Indy | single_day | 20161017 | 20161017 | 44 | r2 | 0.0318737 |
| regression | KF | Indy | single_day | 20161024 | 20161024 | 44 | r2 | 0.0290894 |
| regression | KF | Indy | single_day | 20161025 | 20161025 | 44 | r2 | 0.0372359 |
| regression | KF | Indy | single_day | 20161026 | 20161026 | 44 | r2 | 0.0483609 |
| regression | KF | LINK CO | single_day | 20210104 | 20210104 | 44 | r2 | 0.390276 |
| regression | KF | LINK CO | single_day | 20210105 | 20210105 | 44 | r2 | 0.372782 |
| regression | KF | LINK CO | single_day | 20210106 | 20210106 | 44 | r2 | 0.297696 |
| regression | KF | LINK CO | single_day | 20210108 | 20210108 | 44 | r2 | 0.46593 |
| regression | KF | LINK CO | single_day | 20210111 | 20210111 | 44 | r2 | 0.357844 |
| regression | KF | LINK CO | single_day | 20210123 | 20210123 | 44 | r2 | 0.417039 |
| regression | KF | LINK CO | single_day | 20210223 | 20210223 | 44 | r2 | 0.270765 |
| regression | KF | LINK CO | single_day | 20210225 | 20210225 | 44 | r2 | 0.332804 |
| regression | KF | LINK CO | single_day | 20210226 | 20210226 | 44 | r2 | 0.318936 |
| regression | KF | LINK CO | single_day | 20210227 | 20210227 | 44 | r2 | 0.241844 |
| regression | KF | LINK CO | single_day | 20210309 | 20210309 | 44 | r2 | 0.267256 |
| regression | KF | LINK CO | single_day | 20210313 | 20210313 | 44 | r2 | 0.309335 |
| regression | KF | LINK CO | single_day | 20210329 | 20210329 | 44 | r2 | 0.203443 |
| regression | KF | LINK CO | single_day | 20210330 | 20210330 | 44 | r2 | 0.263682 |
| regression | KF | LINK CO | single_day | 20210402 | 20210402 | 44 | r2 | 0.13859 |
| regression | KF | LINK CO | single_day | 20210406 | 20210406 | 44 | r2 | 0.317572 |
| regression | KF | LINK CO | single_day | 20210409 | 20210409 | 44 | r2 | 0.267571 |
| regression | KF | LINK CO | single_day | 20210415 | 20210415 | 44 | r2 | 0.278713 |
| regression | KF | LINK CO | single_day | 20210420 | 20210420 | 44 | r2 | 0.253574 |
| regression | KF | LINK CO | single_day | 20210503 | 20210503 | 44 | r2 | 0.267261 |
| regression | KF | LINK CO | single_day | 20210504 | 20210504 | 44 | r2 | 0.31574 |
| regression | KF | LINK CO | single_day | 20210511 | 20210511 | 44 | r2 | 0.348329 |
| regression | KF | LINK CO | single_day | 20210518 | 20210518 | 44 | r2 | 0.453662 |
| regression | KF | LINK CO | single_day | 20210520 | 20210520 | 44 | r2 | 0.425847 |
| regression | KF | LINK CO | single_day | 20210601 | 20210601 | 44 | r2 | 0.396442 |
| regression | KF | LINK RTT | single_day | 20210305 | 20210305 | 44 | r2 | 0.354521 |
| regression | KF | LINK RTT | single_day | 20210312 | 20210312 | 44 | r2 | 0.327289 |
| regression | KF | LINK RTT | single_day | 20210316 | 20210316 | 44 | r2 | 0.333491 |
| regression | KF | LINK RTT | single_day | 20210319 | 20210319 | 44 | r2 | 0.327306 |
| regression | KF | LINK RTT | single_day | 20210322 | 20210322 | 44 | r2 | 0.313053 |
| regression | KF | LINK RTT | single_day | 20210405 | 20210405 | 44 | r2 | 0.319567 |
| regression | KF | LINK RTT | single_day | 20210407 | 20210407 | 44 | r2 | 0.325296 |
| regression | KF | LINK RTT | single_day | 20210412 | 20210412 | 44 | r2 | 0.345339 |
| regression | KF | LINK RTT | single_day | 20210414 | 20210414 | 44 | r2 | 0.331093 |
| regression | KF | LINK RTT | single_day | 20210421 | 20210421 | 44 | r2 | 0.273394 |
| regression | KF | LINK RTT | single_day | 20210428 | 20210428 | 44 | r2 | 0.231761 |
| regression | KF | LINK RTT | single_day | 20210505 | 20210505 | 44 | r2 | 0.29031 |

| | | | | | | | | |
|---|---|---|---|---|---|---|---|---|
| regression | KF | LINK RTT | single_day | 20210519 | 20210519 | 44 | r2 | 0.47651 |
| regression | KF | LINK RTT | single_day | 20210525 | 20210525 | 44 | r2 | 0.409695 |
| regression | KF | LINK RTT | single_day | 20210526 | 20210526 | 44 | r2 | 0.36176 |
| regression | KF | LINK RTT | single_day | 20210604 | 20210604 | 44 | r2 | 0.364126 |
| regression | KF | LINK RTT | single_day | 20210608 | 20210608 | 44 | r2 | 0.415884 |
| regression | KF | LINK RTT | single_day | 20210616 | 20210616 | 44 | r2 | 0.437525 |
| regression | KF | LINK RTT | single_day | 20210617 | 20210617 | 44 | r2 | 0.480843 |
| regression | KF | LINK RTT | single_day | 20210624 | 20210624 | 44 | r2 | 0.35149 |
| regression | KF | LINK RTT | single_day | 20210626 | 20210626 | 44 | r2 | 0.431258 |
| regression | KF | LINK RTT | single_day | 20210629 | 20210629 | 44 | r2 | 0.32149 |
| regression | KF | LINK RTT | single_day | 20210630 | 20210630 | 44 | r2 | 0.486494 |
| regression | KF | LINK RTT | single_day | 20210701 | 20210701 | 44 | r2 | 0.384536 |
| regression | KF | LINK RTT | single_day | 20210703 | 20210703 | 44 | r2 | 0.421623 |
| regression | KF | FALCON M2 | single_day | 20201019 | 20201019 | 44 | r2 | 0.295143 |
| regression | KF | FALCON M2 | single_day | 20201019 | 20201019 | 44 | r2 | 0.27465 |
| regression | KF | FALCON M2 | single_day | 20201020 | 20201020 | 44 | r2 | 0.334889 |
| regression | KF | FALCON M2 | single_day | 20201020 | 20201020 | 44 | r2 | 0.33405 |
| regression | KF | FALCON M2 | single_day | 20201027 | 20201027 | 44 | r2 | 0.244504 |
| regression | KF | FALCON M2 | single_day | 20201027 | 20201027 | 44 | r2 | 0.28429 |
| regression | KF | FALCON M2 | cross_day | 20201019__to__20201020 | 20201030 | 44 | r2 | 0.194073 |
| regression | KF | FALCON M2 | cross_day | 20201019__to__20201020 | 20201030 | 44 | r2 | 0.211126 |
| regression | KF | FALCON M2 | cross_day | 20201019__to__20201020 | 20201118 | 44 | r2 | 0.100299 |
| regression | KF | FALCON M2 | cross_day | 20201019__to__20201020 | 20201119 | 44 | r2 | 0.0333668 |
| regression | KF | FALCON M2 | cross_day | 20201019__to__20201020 | 20201124 | 44 | r2 | -0.0570163 |
| regression | KF | FALCON M2 | cross_day | 20201019__to__20201020 | 20201124 | 44 | r2 | 0.0202587 |
| regression | KF | LINK CO | cross_day | 20210104__to__20210111 | 20210225 | 44 | r2 | 0.136307 |
| regression | KF | LINK CO | cross_day | 20210104__to__20210111 | 20210226 | 44 | r2 | 0.144682 |
| regression | KF | LINK CO | cross_day | 20210104__to__20210111 | 20210227 | 44 | r2 | 0.138381 |
| regression | KF | LINK CO | cross_day | 20210104__to__20210111 | 20210309 | 44 | r2 | 0.155194 |
| regression | KF | LINK CO | cross_day | 20210104__to__20210111 | 20210313 | 44 | r2 | 0.149119 |
| regression | KF | LINK CO | cross_day | 20210104__to__20210111 | 20210329 | 44 | r2 | 0.0907232 |
| regression | KF | LINK CO | cross_day | 20210104__to__20210111 | 20210330 | 44 | r2 | 0.0502138 |
| regression | KF | LINK CO | cross_day | 20210104__to__20210111 | 20210402 | 44 | r2 | 0.0973771 |
| regression | KF | LINK CO | cross_day | 20210104__to__20210111 | 20210406 | 44 | r2 | 0.130058 |
| regression | KF | LINK CO | cross_day | 20210104__to__20210111 | 20210409 | 44 | r2 | 0.166423 |
| regression | KF | LINK CO | cross_day | 20210104__to__20210111 | 20210415 | 44 | r2 | 0.0953895 |
| regression | KF | LINK CO | cross_day | 20210104__to__20210111 | 20210420 | 44 | r2 | 0.144627 |
| regression | KF | LINK CO | cross_day | 20210104__to__20210111 | 20210503 | 44 | r2 | 0.0822616 |
| regression | KF | LINK CO | cross_day | 20210104__to__20210111 | 20210504 | 44 | r2 | 0.0300179 |
| regression | KF | LINK CO | cross_day | 20210104__to__20210111 | 20210511 | 44 | r2 | 0.231559 |
| regression | KF | LINK CO | cross_day | 20210104__to__20210111 | 20210518 | 44 | r2 | 0.215053 |
| regression | KF | LINK CO | cross_day | 20210104__to__20210111 | 20210520 | 44 | r2 | 0.259852 |
| regression | KF | LINK CO | cross_day | 20210104__to__20210111 | 20210601 | 44 | r2 | 0.208716 |
| regression | KF | LINK RTT | cross_day | 20210305__to__20210322 | 20210412 | 44 | r2 | 0.217476 |
| regression | KF | LINK RTT | cross_day | 20210305__to__20210322 | 20210414 | 44 | r2 | 0.212839 |
| regression | KF | LINK RTT | cross_day | 20210305__to__20210322 | 20210421 | 44 | r2 | 0.10158 |
| regression | KF | LINK RTT | cross_day | 20210305__to__20210322 | 20210428 | 44 | r2 | 0.179511 |
| regression | KF | LINK RTT | cross_day | 20210305__to__20210322 | 20210505 | 44 | r2 | 0.240664 |
| regression | KF | LINK RTT | cross_day | 20210305__to__20210322 | 20210519 | 44 | r2 | 0.260018 |
| regression | KF | LINK RTT | cross_day | 20210305__to__20210322 | 20210525 | 44 | r2 | 0.295445 |
| regression | KF | LINK RTT | cross_day | 20210305__to__20210322 | 20210526 | 44 | r2 | 0.248651 |
| regression | KF | LINK RTT | cross_day | 20210305__to__20210322 | 20210604 | 44 | r2 | 0.309073 |
| regression | KF | LINK RTT | cross_day | 20210305__to__20210322 | 20210608 | 44 | r2 | 0.227123 |
| regression | KF | LINK RTT | cross_day | 20210305__to__20210322 | 20210616 | 44 | r2 | 0.230246 |
| regression | KF | LINK RTT | cross_day | 20210305__to__20210322 | 20210617 | 44 | r2 | 0.234816 |
| regression | KF | LINK RTT | cross_day | 20210305__to__20210322 | 20210624 | 44 | r2 | 0.22806 |
| regression | KF | LINK RTT | cross_day | 20210305__to__20210322 | 20210626 | 44 | r2 | 0.283531 |
| regression | KF | LINK RTT | cross_day | 20210305__to__20210322 | 20210629 | 44 | r2 | 0.231466 |
| regression | KF | LINK RTT | cross_day | 20210305__to__20210322 | 20210630 | 44 | r2 | 0.271551 |
| regression | KF | LINK RTT | cross_day | 20210305__to__20210322 | 20210701 | 44 | r2 | 0.244824 |
| regression | KF | LINK RTT | cross_day | 20210305__to__20210322 | 20210703 | 44 | r2 | 0.247063 |
| regression | KF | Indy | cross_day | 20160418__to__20160622 | 20160630 | 44 | r2 | -0.271936 |
| regression | KF | Indy | cross_day | 20160418__to__20160622 | 20160915 | 44 | r2 | 0.334653 |

| | | | | | | | | |
|---|---|---|---|---|---|---|---|---|
| regression | KF | Indy | cross_day | 20160418__to__20160622 | 20160916 | 44 | r2 | 0.318763 |
| regression | KF | Indy | cross_day | 20160418__to__20160622 | 20160921 | 44 | r2 | 0.296787 |
| regression | KF | Indy | cross_day | 20160418__to__20160622 | 20160927 | 44 | r2 | 0.196879 |
| regression | KF | Indy | cross_day | 20160418__to__20160622 | 20160927 | 44 | r2 | 0.195968 |
| regression | KF | Indy | cross_day | 20160418__to__20160622 | 20160930 | 44 | r2 | 0.203192 |
| regression | KF | Indy | cross_day | 20160418__to__20160622 | 20160930 | 44 | r2 | 0.240289 |
| regression | KF | Indy | cross_day | 20160418__to__20160622 | 20161005 | 44 | r2 | 0.162992 |
| regression | KF | Indy | cross_day | 20160418__to__20160622 | 20161006 | 44 | r2 | 0.323878 |
| regression | KF | Indy | cross_day | 20160418__to__20160622 | 20161007 | 44 | r2 | 0.258674 |
| regression | KF | Indy | cross_day | 20160418__to__20160622 | 20161011 | 44 | r2 | 0.182978 |
| regression | KF | Indy | cross_day | 20160418__to__20160622 | 20161013 | 44 | r2 | 0.0301049 |
| regression | KF | Indy | cross_day | 20160418__to__20160622 | 20161014 | 44 | r2 | 0.285039 |
| regression | KF | Indy | cross_day | 20160418__to__20160622 | 20161017 | 44 | r2 | 0.221235 |
| regression | KF | Indy | cross_day | 20160418__to__20160622 | 20161024 | 44 | r2 | 0.0878173 |
| regression | KF | Indy | cross_day | 20160418__to__20160622 | 20161025 | 44 | r2 | 0.288327 |
| regression | KF | Indy | cross_day | 20160418__to__20160622 | 20161026 | 44 | r2 | 0.198614 |
| regression | MLP | FALCON M2 | cross_day | 20201019__to__20201020 | 20201030 | 44 | r2 | 0.240396 |
| regression | MLP | FALCON M2 | cross_day | 20201019__to__20201020 | 20201030 | 44 | r2 | 0.310066 |
| regression | MLP | FALCON M2 | cross_day | 20201019__to__20201020 | 20201118 | 44 | r2 | 0.0478222 |
| regression | MLP | FALCON M2 | cross_day | 20201019__to__20201020 | 20201119 | 44 | r2 | 0.061923 |
| regression | MLP | FALCON M2 | cross_day | 20201019__to__20201020 | 20201124 | 44 | r2 | 0.0484977 |
| regression | MLP | FALCON M2 | cross_day | 20201019__to__20201020 | 20201124 | 44 | r2 | 0.0172608 |
| regression | MLP | FALCON M2 | single_day | 20201019 | 20201019 | 44 | r2 | 0.314356 |
| regression | MLP | FALCON M2 | single_day | 20201019 | 20201019 | 44 | r2 | 0.329889 |
| regression | MLP | FALCON M2 | single_day | 20201020 | 20201020 | 44 | r2 | 0.290463 |
| regression | MLP | FALCON M2 | single_day | 20201020 | 20201020 | 44 | r2 | 0.265533 |
| regression | MLP | FALCON M2 | single_day | 20201027 | 20201027 | 44 | r2 | 0.158746 |
| regression | MLP | FALCON M2 | single_day | 20201027 | 20201027 | 44 | r2 | 0.226665 |
| regression | MLP | Jango | cross_day | 20150730__to__20150828 | 20150906 | 44 | r2 | 0.90532 |
| regression | MLP | Jango | cross_day | 20150730__to__20150828 | 20150908 | 44 | r2 | 0.850776 |
| regression | MLP | Jango | cross_day | 20150730__to__20150828 | 20151029 | 44 | r2 | 0.892563 |
| regression | MLP | Jango | cross_day | 20150730__to__20150828 | 20151102 | 44 | r2 | 0.739406 |
| regression | MLP | Jango | single_day | 20150730 | 20150730 | 44 | r2 | 0.83515 |
| regression | MLP | Jango | single_day | 20150801 | 20150801 | 44 | r2 | 0.906526 |
| regression | MLP | Jango | single_day | 20150806 | 20150806 | 44 | r2 | 0.869034 |
| regression | MLP | Jango | single_day | 20150808 | 20150808 | 44 | r2 | 0.879722 |
| regression | MLP | Jango | single_day | 20150820 | 20150820 | 44 | r2 | 0.870891 |
| regression | MLP | Jango | single_day | 20150825 | 20150825 | 44 | r2 | 0.868624 |
| regression | MLP | Jango | single_day | 20150827 | 20150827 | 44 | r2 | 0.859966 |
| regression | MLP | Jango | single_day | 20150831 | 20150831 | 44 | r2 | 0.880751 |
| regression | MLP | Jango | single_day | 20150906 | 20150906 | 44 | r2 | 0.902983 |
| regression | MLP | Jango | single_day | 20151102 | 20151102 | 44 | r2 | 0.828123 |
| regression | MLP | LINK CO | cross_day | 20210104__to__20210111 | 20210225 | 44 | r2 | 0.336536 |
| regression | MLP | LINK CO | cross_day | 20210104__to__20210111 | 20210226 | 44 | r2 | 0.352739 |
| regression | MLP | LINK CO | cross_day | 20210104__to__20210111 | 20210227 | 44 | r2 | 0.302554 |
| regression | MLP | LINK CO | cross_day | 20210104__to__20210111 | 20210309 | 44 | r2 | 0.280495 |
| regression | MLP | LINK CO | cross_day | 20210104__to__20210111 | 20210313 | 44 | r2 | 0.262205 |
| regression | MLP | LINK CO | cross_day | 20210104__to__20210111 | 20210329 | 44 | r2 | 0.346998 |
| regression | MLP | LINK CO | cross_day | 20210104__to__20210111 | 20210330 | 44 | r2 | 0.199717 |
| regression | MLP | LINK CO | cross_day | 20210104__to__20210111 | 20210402 | 44 | r2 | 0.22113 |
| regression | MLP | LINK CO | cross_day | 20210104__to__20210111 | 20210406 | 44 | r2 | 0.307817 |
| regression | MLP | LINK CO | cross_day | 20210104__to__20210111 | 20210409 | 44 | r2 | 0.35081 |
| regression | MLP | LINK CO | cross_day | 20210104__to__20210111 | 20210415 | 44 | r2 | 0.295824 |
| regression | MLP | LINK CO | cross_day | 20210104__to__20210111 | 20210420 | 44 | r2 | 0.282387 |
| regression | MLP | LINK CO | cross_day | 20210104__to__20210111 | 20210503 | 44 | r2 | 0.160396 |
| regression | MLP | LINK CO | cross_day | 20210104__to__20210111 | 20210504 | 44 | r2 | 0.0792794 |
| regression | MLP | LINK CO | cross_day | 20210104__to__20210111 | 20210511 | 44 | r2 | 0.368877 |
| regression | MLP | LINK CO | cross_day | 20210104__to__20210111 | 20210518 | 44 | r2 | 0.430257 |
| regression | MLP | LINK CO | cross_day | 20210104__to__20210111 | 20210520 | 44 | r2 | 0.423928 |
| regression | MLP | LINK CO | cross_day | 20210104__to__20210111 | 20210601 | 44 | r2 | 0.353582 |
| regression | MLP | LINK CO | single_day | 20210104 | 20210104 | 44 | r2 | 0.439649 |
| regression | MLP | LINK CO | single_day | 20210105 | 20210105 | 44 | r2 | 0.379505 |
| regression | MLP | LINK CO | single_day | 20210106 | 20210106 | 44 | r2 | 0.420014 |

| | | | | | | | | |
|---|---|---|---|---|---|---|---|---|
| regression | MLP | LINK CO | single_day | 20210108 | 20210108 | 44 | r2 | 0.40032 |
| regression | MLP | LINK CO | single_day | 20210111 | 20210111 | 44 | r2 | 0.327894 |
| regression | MLP | LINK CO | single_day | 20210123 | 20210123 | 44 | r2 | 0.3934 |
| regression | MLP | LINK CO | single_day | 20210223 | 20210223 | 44 | r2 | 0.286433 |
| regression | MLP | LINK CO | single_day | 20210225 | 20210225 | 44 | r2 | 0.234621 |
| regression | MLP | LINK CO | single_day | 20210226 | 20210226 | 44 | r2 | 0.380251 |
| regression | MLP | LINK CO | single_day | 20210227 | 20210227 | 44 | r2 | 0.317116 |
| regression | MLP | LINK CO | single_day | 20210309 | 20210309 | 44 | r2 | 0.294457 |
| regression | MLP | LINK CO | single_day | 20210313 | 20210313 | 44 | r2 | 0.293557 |
| regression | MLP | LINK CO | single_day | 20210329 | 20210329 | 44 | r2 | 0.321894 |
| regression | MLP | LINK CO | single_day | 20210330 | 20210330 | 44 | r2 | 0.271378 |
| regression | MLP | LINK CO | single_day | 20210402 | 20210402 | 44 | r2 | 0.26236 |
| regression | MLP | LINK CO | single_day | 20210406 | 20210406 | 44 | r2 | 0.346216 |
| regression | MLP | LINK CO | single_day | 20210409 | 20210409 | 44 | r2 | 0.338832 |
| regression | MLP | LINK CO | single_day | 20210415 | 20210415 | 44 | r2 | 0.277444 |
| regression | MLP | LINK CO | single_day | 20210420 | 20210420 | 44 | r2 | 0.310713 |
| regression | MLP | LINK CO | single_day | 20210503 | 20210503 | 44 | r2 | 0.312868 |
| regression | MLP | LINK CO | single_day | 20210504 | 20210504 | 44 | r2 | 0.320186 |
| regression | MLP | LINK CO | single_day | 20210511 | 20210511 | 44 | r2 | 0.386355 |
| regression | MLP | LINK CO | single_day | 20210518 | 20210518 | 44 | r2 | 0.398917 |
| regression | MLP | LINK CO | single_day | 20210520 | 20210520 | 44 | r2 | 0.431977 |
| regression | MLP | LINK CO | single_day | 20210601 | 20210601 | 44 | r2 | 0.393943 |
| regression | MLP | LINK RTT | cross_day | 20210305__to__20210322 | 20210412 | 44 | r2 | 0.43636 |
| regression | MLP | LINK RTT | cross_day | 20210305__to__20210322 | 20210414 | 44 | r2 | 0.438605 |
| regression | MLP | LINK RTT | cross_day | 20210305__to__20210322 | 20210421 | 44 | r2 | 0.236917 |
| regression | MLP | LINK RTT | cross_day | 20210305__to__20210322 | 20210428 | 44 | r2 | 0.380798 |
| regression | MLP | LINK RTT | cross_day | 20210305__to__20210322 | 20210505 | 44 | r2 | 0.354752 |
| regression | MLP | LINK RTT | cross_day | 20210305__to__20210322 | 20210519 | 44 | r2 | 0.423822 |
| regression | MLP | LINK RTT | cross_day | 20210305__to__20210322 | 20210525 | 44 | r2 | 0.416882 |
| regression | MLP | LINK RTT | cross_day | 20210305__to__20210322 | 20210526 | 44 | r2 | 0.441022 |
| regression | MLP | LINK RTT | cross_day | 20210305__to__20210322 | 20210604 | 44 | r2 | 0.453482 |
| regression | MLP | LINK RTT | cross_day | 20210305__to__20210322 | 20210608 | 44 | r2 | 0.310056 |
| regression | MLP | LINK RTT | cross_day | 20210305__to__20210322 | 20210616 | 44 | r2 | 0.37359 |
| regression | MLP | LINK RTT | cross_day | 20210305__to__20210322 | 20210617 | 44 | r2 | 0.375087 |
| regression | MLP | LINK RTT | cross_day | 20210305__to__20210322 | 20210624 | 44 | r2 | 0.343802 |
| regression | MLP | LINK RTT | cross_day | 20210305__to__20210322 | 20210626 | 44 | r2 | 0.362006 |
| regression | MLP | LINK RTT | cross_day | 20210305__to__20210322 | 20210629 | 44 | r2 | 0.339909 |
| regression | MLP | LINK RTT | cross_day | 20210305__to__20210322 | 20210630 | 44 | r2 | 0.443357 |
| regression | MLP | LINK RTT | cross_day | 20210305__to__20210322 | 20210701 | 44 | r2 | 0.417015 |
| regression | MLP | LINK RTT | cross_day | 20210305__to__20210322 | 20210703 | 44 | r2 | 0.353951 |
| regression | MLP | LINK RTT | single_day | 20210305 | 20210305 | 44 | r2 | 0.406671 |
| regression | MLP | LINK RTT | single_day | 20210312 | 20210312 | 44 | r2 | 0.402602 |
| regression | MLP | LINK RTT | single_day | 20210316 | 20210316 | 44 | r2 | 0.373202 |
| regression | MLP | LINK RTT | single_day | 20210319 | 20210319 | 44 | r2 | 0.397527 |
| regression | MLP | LINK RTT | single_day | 20210322 | 20210322 | 44 | r2 | 0.377103 |
| regression | MLP | LINK RTT | single_day | 20210405 | 20210405 | 44 | r2 | 0.367457 |
| regression | MLP | LINK RTT | single_day | 20210407 | 20210407 | 44 | r2 | 0.380262 |
| regression | MLP | LINK RTT | single_day | 20210412 | 20210412 | 44 | r2 | 0.382827 |
| regression | MLP | LINK RTT | single_day | 20210414 | 20210414 | 44 | r2 | 0.424121 |
| regression | MLP | LINK RTT | single_day | 20210421 | 20210421 | 44 | r2 | 0.329877 |
| regression | MLP | LINK RTT | single_day | 20210428 | 20210428 | 44 | r2 | 0.34832 |
| regression | MLP | LINK RTT | single_day | 20210505 | 20210505 | 44 | r2 | 0.385225 |
| regression | MLP | LINK RTT | single_day | 20210519 | 20210519 | 44 | r2 | 0.48606 |
| regression | MLP | LINK RTT | single_day | 20210525 | 20210525 | 44 | r2 | 0.462042 |
| regression | MLP | LINK RTT | single_day | 20210526 | 20210526 | 44 | r2 | 0.414265 |
| regression | MLP | LINK RTT | single_day | 20210604 | 20210604 | 44 | r2 | 0.45344 |
| regression | MLP | LINK RTT | single_day | 20210608 | 20210608 | 44 | r2 | 0.457063 |
| regression | MLP | LINK RTT | single_day | 20210616 | 20210616 | 44 | r2 | 0.492652 |
| regression | MLP | LINK RTT | single_day | 20210617 | 20210617 | 44 | r2 | 0.471387 |
| regression | MLP | LINK RTT | single_day | 20210624 | 20210624 | 44 | r2 | 0.414498 |
| regression | MLP | LINK RTT | single_day | 20210626 | 20210626 | 44 | r2 | 0.375067 |
| regression | MLP | LINK RTT | single_day | 20210629 | 20210629 | 44 | r2 | 0.394678 |
| regression | MLP | LINK RTT | single_day | 20210630 | 20210630 | 44 | r2 | 0.468809 |

| | | | | | | | | |
|---|---|---|---|---|---|---|---|---|
| regression | MLP | LINK RTT | single_day | 20210701 | 20210701 | 44 | r2 | 0.430904 |
| regression | MLP | LINK RTT | single_day | 20210703 | 20210703 | 44 | r2 | 0.394449 |
| regression | MLP | Indy | cross_day | 20160418__to__20160622 | 20160630 | 44 | r2 | -0.0447903 |
| regression | MLP | Indy | cross_day | 20160418__to__20160622 | 20160915 | 44 | r2 | 0.399162 |
| regression | MLP | Indy | cross_day | 20160418__to__20160622 | 20160916 | 44 | r2 | 0.469665 |
| regression | MLP | Indy | cross_day | 20160418__to__20160622 | 20160921 | 44 | r2 | 0.415022 |
| regression | MLP | Indy | cross_day | 20160418__to__20160622 | 20160927 | 44 | r2 | 0.382163 |
| regression | MLP | Indy | cross_day | 20160418__to__20160622 | 20160927 | 44 | r2 | 0.373223 |
| regression | MLP | Indy | cross_day | 20160418__to__20160622 | 20160930 | 44 | r2 | 0.380247 |
| regression | MLP | Indy | cross_day | 20160418__to__20160622 | 20160930 | 44 | r2 | 0.375341 |
| regression | MLP | Indy | cross_day | 20160418__to__20160622 | 20161005 | 44 | r2 | 0.284231 |
| regression | MLP | Indy | cross_day | 20160418__to__20160622 | 20161006 | 44 | r2 | 0.371553 |
| regression | MLP | Indy | cross_day | 20160418__to__20160622 | 20161007 | 44 | r2 | 0.304098 |
| regression | MLP | Indy | cross_day | 20160418__to__20160622 | 20161011 | 44 | r2 | 0.402978 |
| regression | MLP | Indy | cross_day | 20160418__to__20160622 | 20161013 | 44 | r2 | -0.00358295 |
| regression | MLP | Indy | cross_day | 20160418__to__20160622 | 20161014 | 44 | r2 | 0.449377 |
| regression | MLP | Indy | cross_day | 20160418__to__20160622 | 20161017 | 44 | r2 | 0.304398 |
| regression | MLP | Indy | cross_day | 20160418__to__20160622 | 20161024 | 44 | r2 | 0.219999 |
| regression | MLP | Indy | cross_day | 20160418__to__20160622 | 20161025 | 44 | r2 | 0.359286 |
| regression | MLP | Indy | cross_day | 20160418__to__20160622 | 20161026 | 44 | r2 | 0.352169 |
| regression | MLP | Indy | single_day | 20160418 | 20160418 | 44 | r2 | 0.674252 |
| regression | MLP | Indy | single_day | 20160419 | 20160419 | 44 | r2 | 0.537803 |
| regression | MLP | Indy | single_day | 20160420 | 20160420 | 44 | r2 | 0.718064 |
| regression | MLP | Indy | single_day | 20160426 | 20160426 | 44 | r2 | 0.714723 |
| regression | MLP | Indy | single_day | 20160622 | 20160622 | 44 | r2 | 0.717868 |
| regression | MLP | Indy | single_day | 20160624 | 20160624 | 44 | r2 | 0.568837 |
| regression | MLP | Indy | single_day | 20160627 | 20160627 | 44 | r2 | 0.759948 |
| regression | MLP | Indy | single_day | 20160630 | 20160630 | 44 | r2 | 0.629267 |
| regression | MLP | Indy | single_day | 20160915 | 20160915 | 44 | r2 | 0.47692 |
| regression | MLP | Indy | single_day | 20160916 | 20160916 | 44 | r2 | 0.544322 |
| regression | MLP | Indy | single_day | 20160921 | 20160921 | 44 | r2 | 0.487058 |
| regression | MLP | Indy | single_day | 20160927 | 20160927 | 44 | r2 | 0.54373 |
| regression | MLP | Indy | single_day | 20160927 | 20160927 | 44 | r2 | 0.546978 |
| regression | MLP | Indy | single_day | 20160930 | 20160930 | 44 | r2 | 0.52197 |
| regression | MLP | Indy | single_day | 20160930 | 20160930 | 44 | r2 | 0.504504 |
| regression | MLP | Indy | single_day | 20161005 | 20161005 | 44 | r2 | 0.480713 |
| regression | MLP | Indy | single_day | 20161006 | 20161006 | 44 | r2 | 0.553293 |
| regression | MLP | Indy | single_day | 20161007 | 20161007 | 44 | r2 | 0.533066 |
| regression | MLP | Indy | single_day | 20161011 | 20161011 | 44 | r2 | 0.56585 |
| regression | MLP | Indy | single_day | 20161013 | 20161013 | 44 | r2 | 0.575752 |
| regression | MLP | Indy | single_day | 20161014 | 20161014 | 44 | r2 | 0.587929 |
| regression | MLP | Indy | single_day | 20161017 | 20161017 | 44 | r2 | 0.556252 |
| regression | MLP | Indy | single_day | 20161024 | 20161024 | 44 | r2 | 0.562038 |
| regression | MLP | Indy | single_day | 20161025 | 20161025 | 44 | r2 | 0.534759 |
| regression | MLP | Indy | single_day | 20161026 | 20161026 | 44 | r2 | 0.572944 |
